# Shared Neural Codes for Emotion Recognition in Emoji and Human Faces

**DOI:** 10.1101/2025.03.21.640322

**Authors:** Madeline Molly Ely, Chloe Ann Kelsey, Géza Gergely Ambrus

## Abstract

Facial expressions are critical social signals, essential for human communication. This study used EEG to investigate the neural dynamics of the processing of emotional expressions in real and emoji faces, using a data-driven approach. Across two experiments with identical paradigms, two separate sets of participants viewed facial expressions (happy, angry, sad, neutral) in real faces (4 female and 4 male identities, *n* = 24) or emojis (6 platforms, *n* = 25) while performing a two-alternative forced-choice emotion recognition task. Time-resolved multivariate classification and spatio-temporal searchlight analyses revealed robust decoding of emotional expressions within and across experiments. Consistent effects emerged early and peaked between 145–160 ms over posterior-occipital and parietal regions. Notably, robust cross-classification between real and emoji faces demonstrated that face-like emoji stimuli evoke neural responses comparable to those elicited by real faces, with more sustained effects over right posterior sites. These findings suggest that the brain uses overlapping spatio-temporal codes for naturalistic and symbolic facial expressions, providing new insights into the neural coding of social signals and the representational overlap between natural and artificial emotional expressions.

## Introduction

Facial expression recognition is a fundamental aspect of human social interaction, enabling one to infer emotions and intentions from others’ facial cues (Frith, 2009; Jack & Schyns, 2015, 2017; Krumhuber et al., 2023; Schmidt & Cohn, 2001). In parallel, the advent of digital communication introduced emojis as established tools for conveying emotions in online interactions (Kaye et al., 2017). These simplified, iconic representations serve as surrogates for facial expressions, raising questions about how the brain processes such abstracted emotional cues compared to real human faces (Kaye et al., 2021). This data-driven study aims to address the adaptability and generalization of the neural mechanisms involved in emotion recognition by investigating whether real and emoji faces are processed similarly at the neural level.

Previous studies have found that the onset of human facial expression decoding occurs approximately 100 milliseconds after stimulus presentation. Smith and Smith (2019) reported that both facial expression and identity effects peaked within 90–170 ms over posterior electrodes, with task context influencing identity decoding at early stages but not expression decoding. Similarly, Muukkonen et al. (2020) observed that emotion information became available around 100 ms, spreading from occipital to temporal areas within the first 100–250 ms, with signals for happy faces peaking earlier than those for angry or fearful faces. Li et al. (2022) found that emotion processing began at approximately 120 ms, preceding identity extraction, which occurred around 235 ms. Additionally, Zhang et al. (2023) demonstrated that regions such as the lateral occipital cortex and inferior parietal cortex differentiated between neutral and expressive faces within the 100–150 ms window, representing categorical information as early as 100 ms.

Recent fMRI studies investigating the neural processing of emojis have largely approached the topic from linguistic or semantic perspectives (Chatzichristos et al., 2020; Dalski et al., 2024). One early fMRI study by Yuasa, Saito, and Mukawa (2011) explored brain activity related to the emotional valence of graphic emoticons (happy or sad). Their findings showed activation in regions such as the right inferior frontal gyrus, cingulate gyrus, and fusiform gyrus, suggesting that abstract emoticons are perceived as dynamic agents, processed similarly to human faces.

Most EEG studies that compare real and emoji faces have concentrated on identifying differences in neural processing between these two stimulus types. Notably, participants tend to recognize emotions expressed by emojis faster and more accurately than those depicted by real faces, which hints at potential differences in how the brain decodes these visual stimuli. For instance, Dalle, Nogare, and Proverbio (2023) conducted a source-localized EEG study showing that neural responses to emojis differed significantly from those elicited by human faces. Specifically, while real faces activated the fusiform face area (FFA), emojis elicited N170 components associated with occipital and object-processing areas. This suggests that the schematic nature of emojis may influence the brain’s processing strategies, with the N170 peak occurring earlier for emojis compared to human faces. In addition, the study revealed that both emojis and real faces engaged the limbic system and orbitofrontal cortex, areas related to emotional processing and anthropomorphization.

Further studies have examined different components of the ERP, such as the P100 and LPP, in response to real and emoji faces. Gantiva et al. (2020) found that human faces elicited larger P100 and LPP amplitudes compared to emoji faces, while emojis produced a larger N170 amplitude. Similarly, Yu et al. (2022) investigated emotional violations in facial expressions, emojis, and emotion words, observing classic N400 effects indicative of semantic processing across all stimuli. Notably, the N400 amplitude was smaller for both faces and emojis than for emotion words, suggesting shared neural mechanisms for processing emotional content across these stimulus types.

Complementing these findings, behavioral and physiological studies demonstrate that emojis can elicit affective and autonomic responses comparable to real faces. For example, Gantiva and collaborators (2021) have shown that happy, angry, and neutral emojis evoke zygomatic muscle activity and skin conductance responses akin to human faces, inducing similar patterns of pleasant and unpleasant affective states. Behavioral results are mixed: Recent studies by Kaye and colleagues explored the emotional and associative properties of emojis through behavioral paradigms. In an Emoji Spatial Stroop Task, Kaye et al. (2022) observed congruency effects for positive emojis presented in upper vertical spaces and negative emojis in lower spaces, suggesting that emojis may carry explicit emotional valence tied to spatial positions. However, follow-up work (Kaye, MacKenzie, et al., 2023) revealed no implicit effects of emoji valence on response accuracy or latency. Further investigations (Kaye, Rocabado, et al., 2023) demonstrated no facilitative effects of emojis on word processing or memory recall, with emojis failing to enhance associative links to emotion concepts. These findings collectively suggest that while emojis may evoke explicit emotional judgments, their role in implicit cognitive or associative processes remains limited.

Despite these insights, gaps remain in our understanding of how the brain processes emotional information conveyed by real faces versus emojis, particularly regarding their temporal dynamics and neural representations. Additionally, while previous studies have identified some overlap in the processing of real faces and emojis, fewer have directly tested whether these two stimulus types share neural codes during emotional expression processing.

In our previous study (Ely & Ambrus, 2025), we found significant and robust effects associated with facial expression processing, emerging as early as 120 ms and peaking around 160 ms. These findings, consistent across time-resolved decoding and representational similarity analyses, persisted even when controlling for low- and high-level image properties, suggesting that the observed emotional expression effects are independent of visual image characteristics. This raises the intriguing possibility that face-like stimuli, such as emojis, may elicit comparable neural patterns in response to emotional expressions.

To test this hypothesis, our study employs MVPA on EEG data to investigate the neural dynamics of facial expression processing across the two distinct stimulus types: real human faces and emoji faces, to determine whether the neural patterns associated with facial expressions are consistent and transferable between these different representations. We hypothesized that if the neural encoding of facial expressions is abstract and robust, it should be possible to decode displays of emotion not only within each experiment but also across the two experiments. Such findings would suggest that the brain utilizes a generalized neural code for emotion recognition, irrespective of the visual form of the facial stimulus.

## Methods

### Datasets

Datasets from two experiments were used, one for real human faces, one for emoji faces. The real faces dataset was taken from Ely and Ambrus (2025), the emoji dataset is novel and was acquired for the purposes of this study. The experiments were conducted in accordance with the guidelines of the Declaration of Helsinki. The study was approved by the ethics committee of Bournemouth University [Ethics ID: #52261]. The experiments were designed and implemented using PsychoPy (Peirce, 2007; Peirce et al., 2019). Participants in both experiments provided written informed consent before the experiment and took part in the study for partial course credits or volunteered their time. Participants disclosed no history of neurological conditions, had normal or corrected-to-normal vision, have not reported taking CNS-acting medication, and were right-handed.

### Experimental Design

#### Real faces experiment

The stimuli comprised frontal color photographs of eight individuals selected from the KDEF database (Lundqvist et al., 1998). It included four male faces (AM10, AM17, AM24, AM31) and four female faces (AF07, AF15, AF26, AF28), each displaying one of four posed facial expressions: happy, angry, sad, and neutral. This resulted in a total of 32 unique images, which were presented in a randomized sequence.

In a two-alternative forced choice (2AFC) paradigm, each trial began with the display of a fixation cross for 200 ms, followed by the presentation of a face image for 1000 ms. Afterward, a choice screen appeared, presenting one correct and one incorrect emotion label. The interstimulus interval varied between 500 and 1000 ms (**Figure 1**). Participants (*n* = 24, 8 male, age: 21.21 ± 4.17 years) were tasked with identifying the emotion conveyed by the face by pressing either the left or right key, with no time constraints imposed on their decision. Each face image was shown 12 times. For each presentation, the accurate facial expression was paired with an incorrect option four times, with response key assignments counterbalanced. To maintain an equal number of correct trials, incorrect responses triggered the reinsertion of the trial into the sequence for later presentation. Consequently, the dataset for each participant included 12 repetitions per image, 96 presentations per facial expression, and 48 presentations per identity, summing to a total of 384 trials.

**Figure 1.**
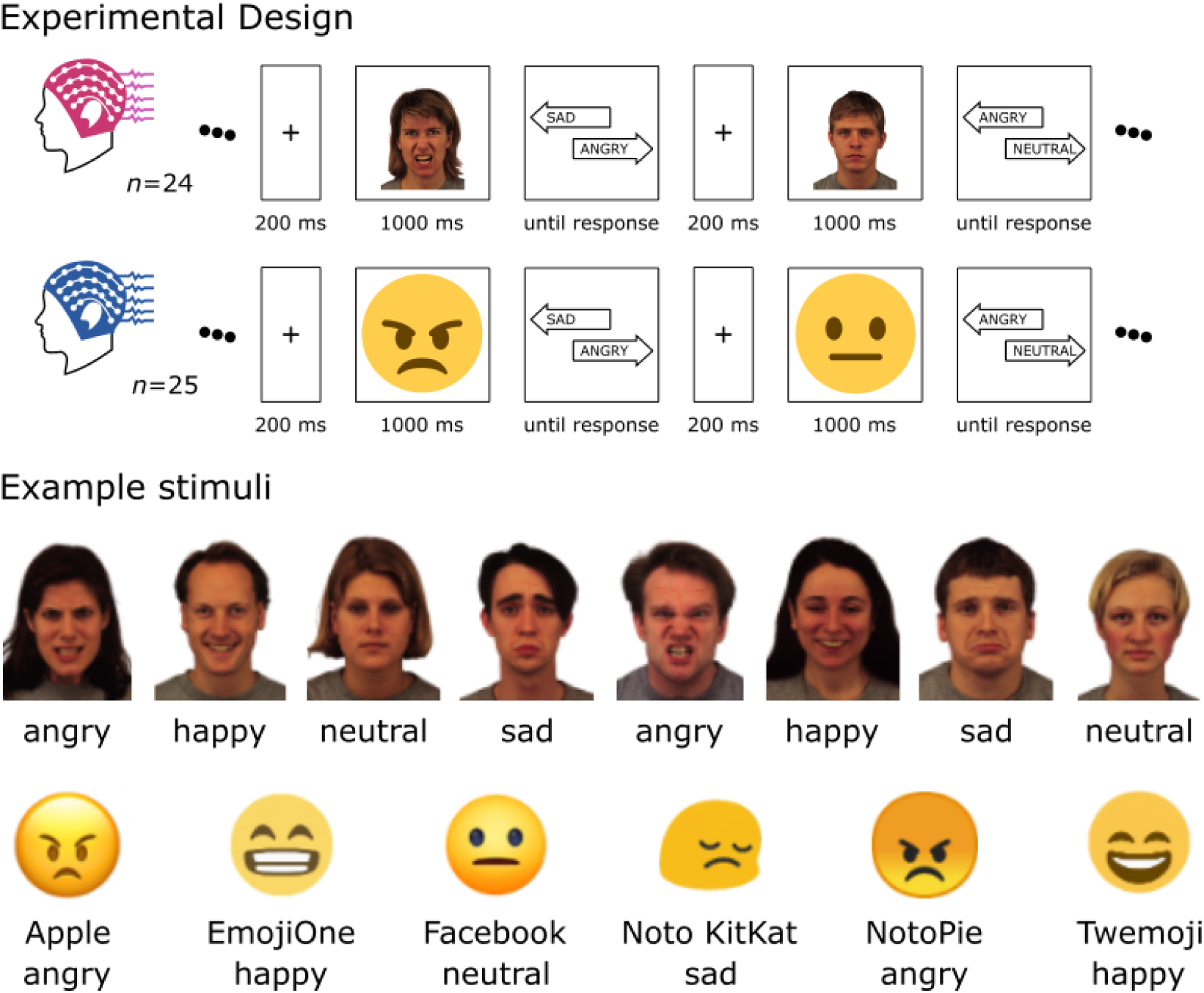
Experimental design and example stimuli. Two experiments were conducted to investigate emotion recognition using a two-alternative forced choice (2AFC) design. The first study (*n* = 24, reported in Ely & Ambrus, 2025) used frontal color photographs of eight individuals (four male, four female) from the KDEF database, each displaying one of four facial expressions—happy, angry, sad, and neutral—resulting in 32 unique images. The second study (*n* = 25) used 24 emoji images sourced from six platforms (Apple, Emoji One, Facebook, Noto KitKat, Noto Pie, and Twemoji) depicting the same four expressions. In both experiments, each trial began with a 200 ms fixation cross, followed by a 1000 ms stimulus presentation, and a choice screen displaying one correct and one incorrect emotion label, with an interstimulus interval between 500 and 1000 ms. Participants selected the correct emotion using a keypress with no time limit. For permission to use sample images, see the footnote. ^1^ The image elements are not to scale.

#### Emoji faces experiment

The design of the emoji experiment followed that of the real faces study. The stimuli comprised color emoji images sourced from six platforms: Apple, Emoji One, Facebook, Noto KitKat, Noto Pie, and Twemoji. Each emoji depicted one of four facial expressions—happy, angry, sad, and neutral—resulting in a total of 24 unique images. These images were presented in a randomized sequence.

The experiment employed a two-alternative forced choice design. Each trial began with a fixation cross displayed for 200 ms, followed by the presentation of an emoji image for 1000 ms. Subsequently, a choice screen appeared, presenting one correct and one incorrect emotion label, with an interstimulus interval ranging from 500 to 1000 ms (**Figure 1**). Participants (*n* = 25, 5 male, age: 21.08 ± 4.68 years) were instructed to identify the correct emotion depicted by the emoji by pressing the left or right key, with no time constraints on their response.

Each image was repeated four times, with the veridical facial expression paired with an incorrect option for response key assignment across trials to ensure counterbalancing. Incorrect responses prompted the reinsertion of the trial into the sequence for later presentation, ensuring an equal number of correct trials. As a result, each participant’s dataset included 12 repetitions per image, 72 presentations per facial expression, and 48 presentations per emoji platform, totaling 288 trials.

### EEG Recording and Processing

EEG data were recorded in the EEG laboratory of the Department of Psychology at Bournemouth University using a 64-channel BioSemi Active-Two device, with electrode placement based on the international 10-20 system. The recorded data were processed using MNE-Python (Gramfort et al., 2013, 2014). Preprocessing included bandpass filtering between 0.1 and 40 Hz, segmentation from −200 to 1200 ms relative to stimulus onset, baseline correction using the 200 ms preceding stimulus presentation, and down-sampling to 200 Hz. To improve the signal-to-noise ratio and optimize computation (Grootswagers et al., 2017), evoked responses for trials with the same image were grouped into bins of three and averaged for each participant. No additional processing steps were applied (Carlson et al., 2020; Delorme, 2023; Grootswagers et al., 2017). Data manipulation was performed using the numpy and scipy packages (Harris et al., 2020; Virtanen et al., 2020).

### Analysis Pipelines

Within-and cross-dataset classification analyses were carried out. Within-experiment analyses were conducted to estimate the information content related to the classes of interest for the two stimulus types (emoji and real faces). Cross-dataset analyses were done to probe the shared neural signals related to the processing of facial expressions displayed by real and emoji faces. Only correct trials were included. Linear discriminant analysis (LDA) classifiers were systematically trained across participants and datasets to categorize the facial expression, as well as the combinations of facial expressions of the presented stimuli. For a similar approach, see Ambrus (2024).

The **within-dataset analyses** employed a leave-one-participant-out approach. For the four-class emotion classification in the real-face experiment, the training data included six identities (three male and three female) aggregated from all but one participant, with testing conducted on the left-out identity in the excluded participant. Similarly, in the emoji experiment, training utilized data from all but one emoji platform aggregated across all but one participant, while testing was performed on the excluded platform in the left-out participant. The classification of emotional expression pairs followed a similar approach, with the key difference being that only two emotional expressions were included in each classification procedure. In the real-face experiment, training was performed on six identities (three male and three female), and testing was conducted on one identity excluded during training. In the emoji experiment, training utilized data for all but one platform, while the omitted platform was iteratively used for testing. These processes were repeated iteratively, ensuring that each identity or platform was excluded once for testing in each participant, and every participant was also left out once for testing within the experimental dataset. For a similar approach, see Klink et al. (2023).

The **cross-experiment analyses** were conducted following the procedures outlined in Dalski et al. (2022a). Classifiers were trained on aggregated data from one dataset and subsequently tested on each participant individually from the other dataset. This analysis was performed in both directions: from emojis to real faces and from real faces to emojis.

Time-resolved classification was performed across all electrodes and pre-defined regions of interest (ROIs), covering six scalp locations distributed along the median (left and right) and coronal (anterior, center, and posterior) planes (Ambrus et al. 2019, 2021). A spatio-temporal searchlight procedure systematically evaluated each channel by training and testing on data from the selected channel and its neighboring electrodes, employing the same time-resolved analysis framework as described in previous studies (Ambrus, 2024; Dalski, Kovács, & Ambrus, 2022). Classification analyses were carried out using scikit-learn (Pedregosa et al., 2011).

### Statistical analyses

A 35 ms moving average (covering 7 consecutive time points) was applied to all participant-level classification accuracy data (Ambrus, 2024; Ambrus et al., 2019, 2021; Dalski, Kovács, & Ambrus, 2022; Klink et al., 2023). Statistical analysis of the results included cluster permutation tests as well as Bayesian statistical methods. Classification accuracies were assessed using two-sided, one-sample cluster permutation tests (10,000 iterations) against chance (0.25 in the four-class facial expression classification, 0.5 in the two-class facial expression pair analyses), implemented in MNE-Python. The Bayesian analyses (Teichmann et al., 2022) also employed a two-sided approach, utilizing a non-directional whole-Cauchy prior with medium width (*r* = 0.707) and excluding the interval δ = -0.5 to +0.5. Bayes factors were then thresholded, with values greater than 10 considered as strong evidence (Moerel et al., 2022; Wetzels et al., 2011). Bayesian analyses were conducted using the BayesFactor R package (Morey et al., 2015).

## Results

### Effect onsets

Effect onsets were estimated by averaging decoding accuracies in the searchlight analyses across all electrodes for each time point. For a similar approach, see Ely and Ambrus (2025). **Figure 2** shows these results for 4-class facial expression classification (**Figure 2A**) and classification of emotion pairs (**Figure 2B-E**). See **Supplementary Information Table 1** for further details.

**Figure 2.**
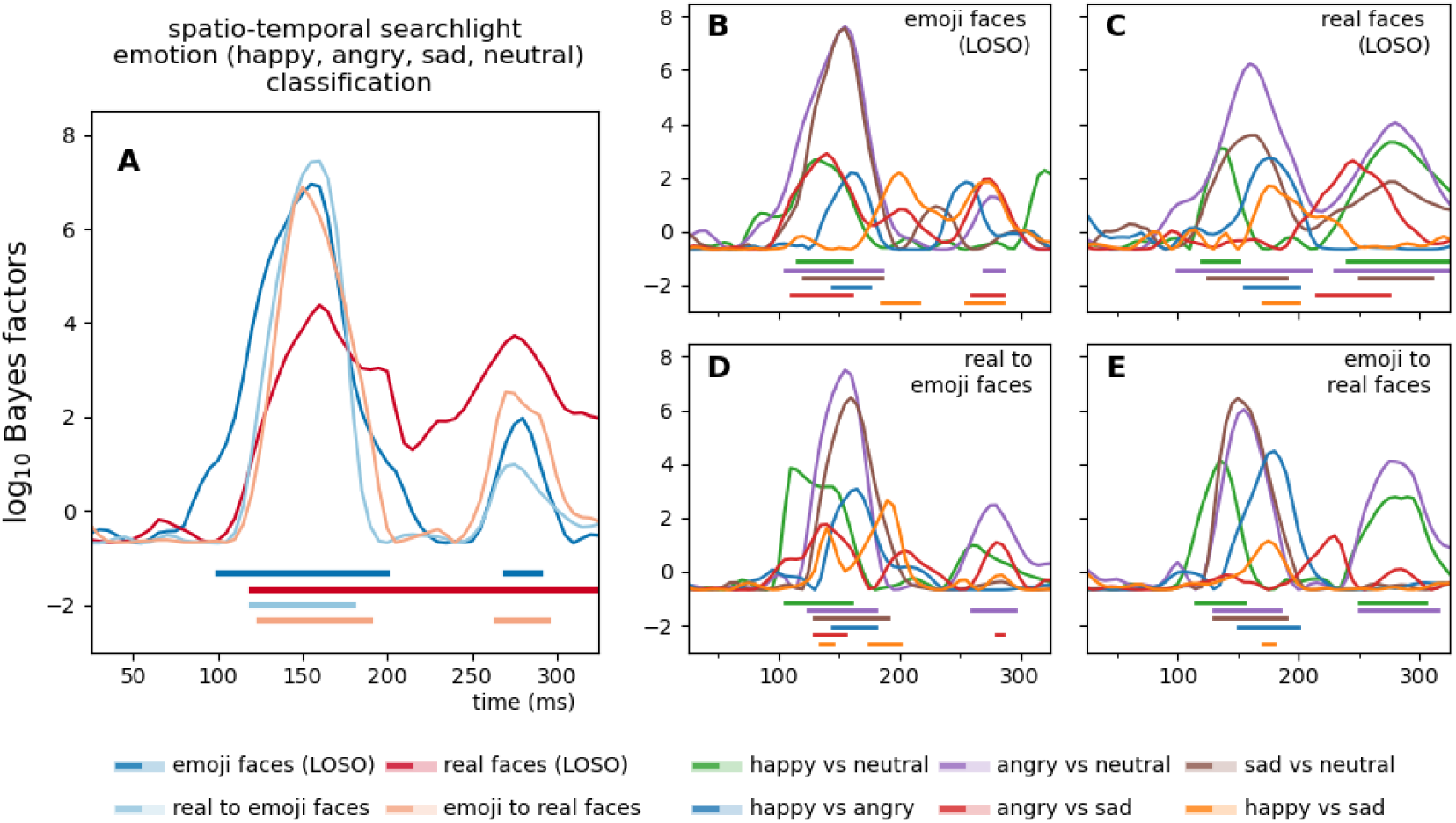
Effect onsets. Time-resolved spatio-temporal searchlight classification was performed on all channels and their neighboring electrodes. Classification performance was averaged across all sensors to probe the onset of the effects. Onsets were determined by two-tailed Bayesian *t*-tests against chance, with BF values exceeding 10 considered indicative of strong evidence (horizontal significance markers). The figure depicts time-resolved Bayes factors. For classification accuracies, see **Supplementary Information Figure 1**.

**Figure 3.**
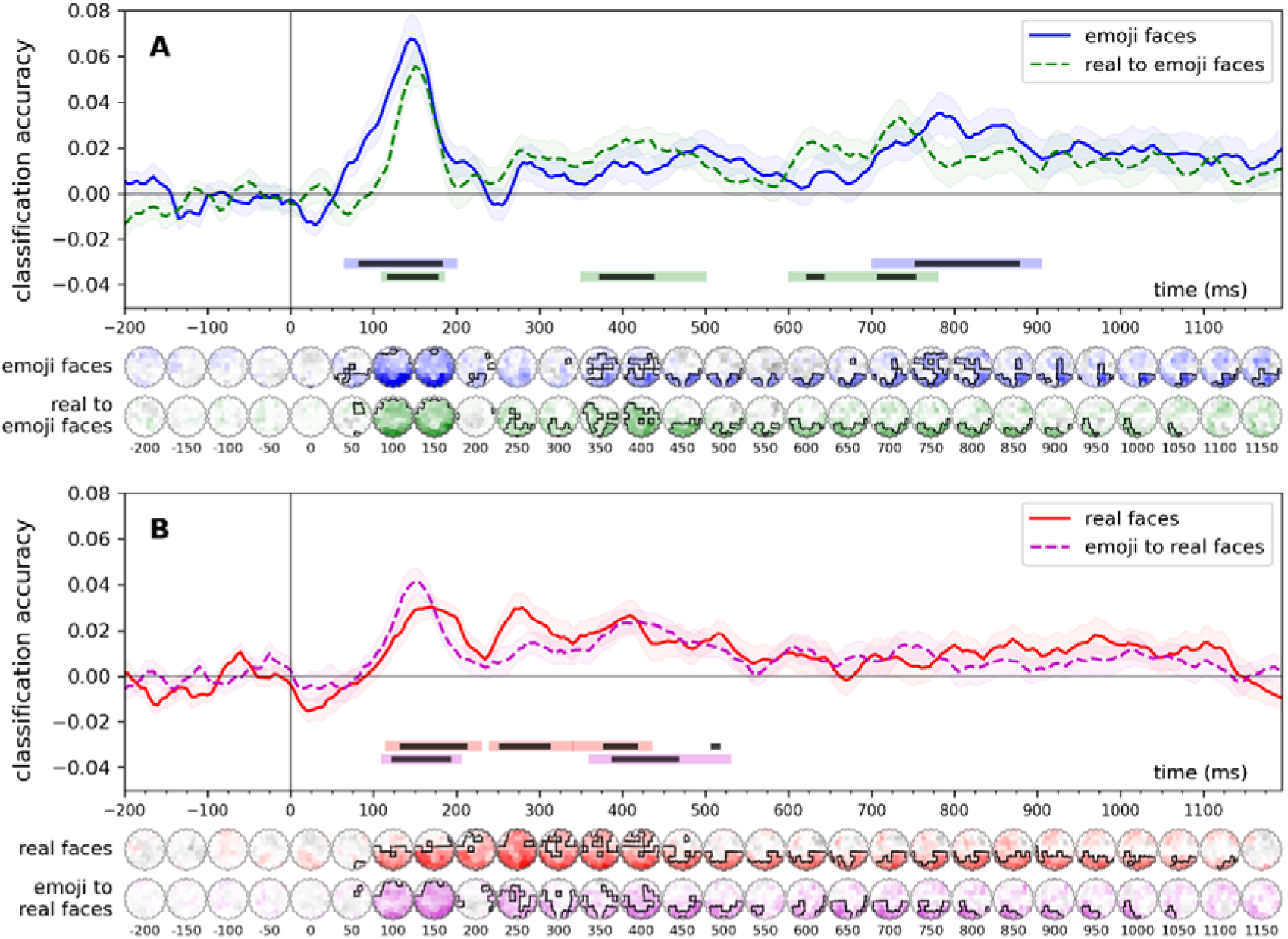
Time-resolved, within-experiment (leave-one-subject-out) and cross-experiment classification of facial expressions. In the within-experiment analyses (solid lines), classifiers were trained, in a leave-one-participant-out scheme to categorize the facial expressions of the stimuli, separately for the real and the emoji faces datasets. In the cross-experiment analyses (dashed lines), classifiers were trained on one dataset and were tested on the other. Error ranges represent ±SEM. Light lines denote significant clusters revealed by the two-sided cluster permutation tests, *p* < 0.05; dark lines denote results of the Bayesian statistical analyses, two-sided one-sample Bayesian *t*-tests, BF > 10, against chance (0.25). Results over all electrodes are presented here. See **Supplementary Information 2**. for the results of the analyses conducted on the pre-defined regions of interest. Spatio-temporal searchlight results are shown as scalp maps, with classification accuracy scores averaged in 50 ms steps. Sensors and time points that belong to significant positive clusters are marked. (Two-sided spatio-temporal cluster permutation tests, *p* < 0.05). For detailed statistics, see **Supplementary Table 1**.

Time-resolved spatio-temporal searchlight analyses revealed consistent decoding of emotional expressions within and across experiments. For emoji faces in the within-experiment leave-one-subject-out (LOSO) classification, emotion decoding onset at 100 ms peaked at 155 ms (BF > 10□), with pairwise contrasts showing early and robust effects. Happy vs. neutral decoding began at 115 ms, peaking at 130 ms (BF = 458.97), while angry vs. neutral started at 105 ms and peaked at 155 ms (BF > 10□). Sad vs. neutral had an onset at 120 ms, peaking similarly at 155 ms (BF > 10□). Other contrasts, such as happy vs. angry and happy vs. sad, showed later onsets and peaks, typically around 145–200 ms.

For real faces, within-experiment LOSO classification showed an emotion onset at 120 ms with a peak at 160 ms (BF > 10□). Pairwise contrasts largely mirrored those seen in emojis, though some contrasts, such as angry vs. sad (onset: 215 ms, peak: 245 ms; BF = 430.02), exhibited later effects.

In cross-experiment analyses, the four-class emotion decoding showed comparable temporal profiles across real-to-emoji and emoji-to-real transfers. Onsets ranged from 120 to 125 ms with peaks at 150–160 ms (BF > 10□), confirming shared neural coding of emotional expressions. Pairwise contrasts revealed earlier decoding for real-to-emoji classifications, such as happy vs. neutral (onset: 105 ms, peak: 110 ms; BF > 10³) and angry vs. neutral (onset: 125 ms, peak: 155 ms; BF > 10□). Emoji-to-real classification showed later effects, with decoding onset at 115–170 ms and peaks up to 180 ms, though angry vs. sad showed no significant effects.

### Multivariate classification

#### Facial expressions

Time-resolved classification of facial expressions (happy, angry, sad, neutral) revealed significant decoding across within- and cross-experiment conditions (**Figure 4**). For emoji stimuli, two clusters emerged: an early window (70–195 ms, p < 0.01, peak Cohen’s *d* = 1.21) and a late window (705–900 ms, *p* < 0.01, Cohen’s *d* = 0.80). In the real face experiment, three significant clusters were observed between 120–430 ms (cluster *p* < 0.05, peak Cohen’s *d*s = 0.78–1.20). Cross-experiment decoding (2AFC to emoji and emoji to 2AFC) yielded early (115–200 ms) and mid-late (355–525 ms) clusters with robust effect sizes (peak Cohen’s *d*s= 0.77–1.56).

**Figure 4.**
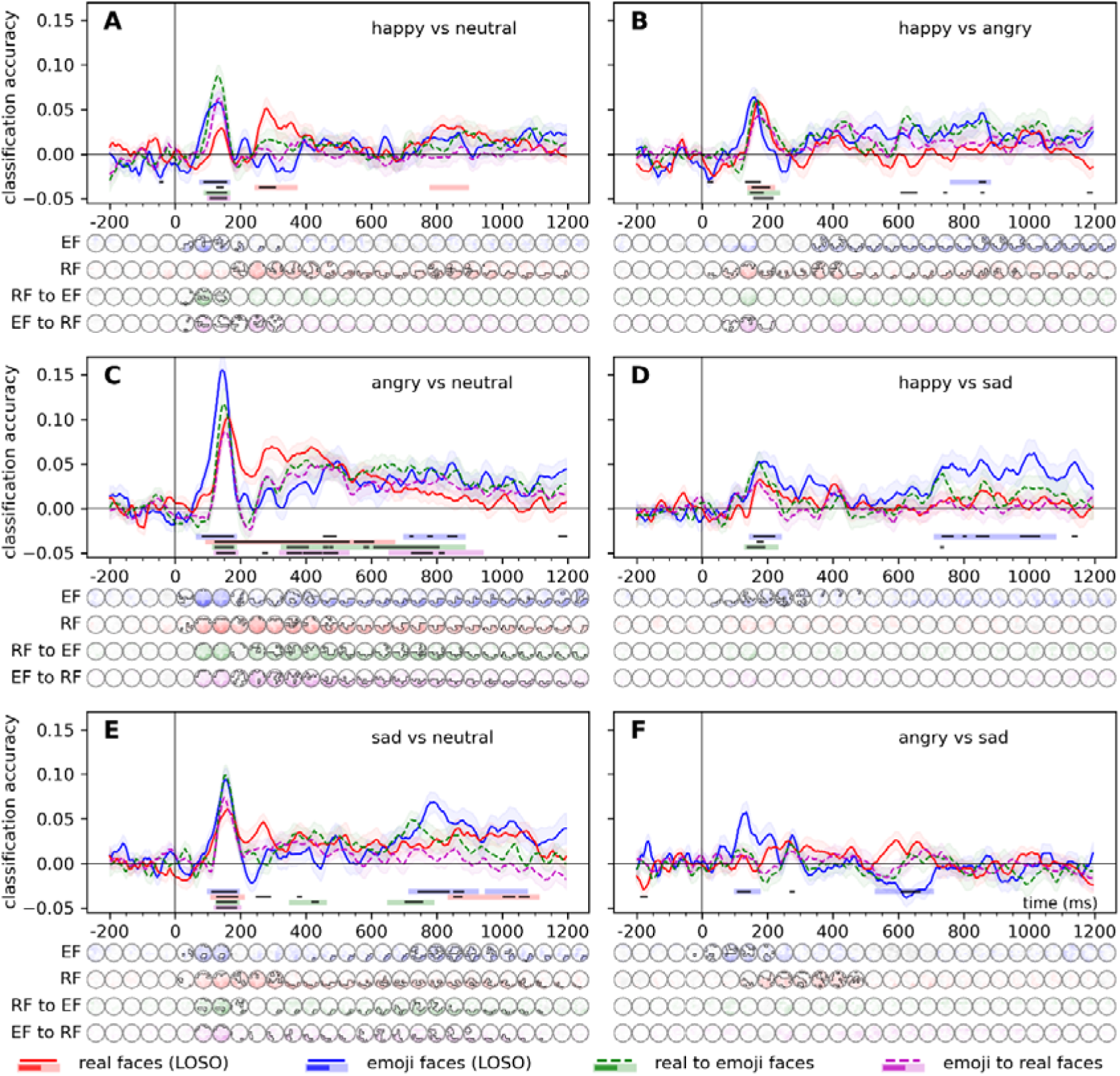
Time-resolved, within-experiment (leave-one-subject-out) and cross-experiment classification of facial expression pairs. For the within-experiment classification (solid lines), training was iteratively performed on six identities (3 male and 3 female) and tested on one left out in the real faces experiment, while in the emoji faces experiment, training was iteratively performed on five platforms and tested one platform left out. In the cross-experiment analyses (dashed lines), classifiers were trained on one dataset and were tested on the other. Error ranges represent ±SEM. Light lines denote significant clusters revealed by the two-sided cluster permutation tests, *p* < 0.05; dark lines denote results of the Bayesian statistical analyses, two-sided one-sample Bayesian *t*-tests, BF > 10, against chance (0.5). Results over all electrodes are presented here. See **Supplementary Information 2**. for the results of the analyses conducted on the pre-defined regions of interest. Spatio-temporal searchlight results are shown as scalp maps, with classification accuracy scores averaged in 50 ms steps. Sensors and time points that belong to significant positive clusters are marked. (Two-sided spatio-temporal cluster permutation tests, *p* < 0.05). For detailed statistics, see **Supplementary Table 1**.

Similarly, spatio-temporal searchlight classification of facial expressions revealed robust decoding in all analyses. For emoji faces, two significant clusters were identified (50–225 ms and 350–1195 ms; cluster *p*s < 0.01) with peak decoding at P6 (peak Cohen’s *d* = 1.91) and PO4 (*d* = 1.14). Real faces showed one broad cluster (80–1145 ms; *p* < 0.0001) peaking at PO7 (*d* = 1.65). Cross-experiment results indicated significant decoding for real-to-emoji (90–1065 ms; *p*s < 0.01) and emoji-to-real faces (75–1060 ms; *p*s < 0.05) with strong effects across occipital and parietal channels (Cohen’s *d*s ≤ 1.0).

#### Facial expression pairs

Time-resolved classification revealed significant decoding of facial expressions across within- and cross-experiment analyses (**Figure 4**). For real faces, decoding was most robust for angry vs. neutral pairs, showing a broad temporal window (100–665 ms, cluster *p* < 0.001, peak Cohen’s *d* = 2.11). Other pairs, such as happy vs. neutral and sad vs. neutral, exhibited narrower windows with positive decoding clusters peaking within 115–280 ms. For emoji faces, early significant decoding was observed for happy vs. neutral and angry vs. neutral (70–180 ms, Cohen’s *d*s > 1.5), with additional late-stage clusters. Cross-experiment results confirmed early decoding (95–160 ms, Cohen’s *ds* > 1.51) for real-to-emoji transfer, particularly for neutral and angry expressions.

Similarly, the spatio-temporal searchlight analysis revealed robust and significant neural differentiation for various facial expression pairs across the experimental conditions. For real faces, decoding of facial expression pairs such as happy vs. neutral, angry vs. neutral, and sad vs. neutral showed early and sustained differentiation, with clusters typically beginning within 75–215 ms and peaking around 150–290 ms at posterior channels (e.g., POz, PO8), with Cohen’s *d* values exceeding 1.6 in some cases. Similarly, for emoji faces, contrasts such as happy vs. neutral and angry vs. neutral demonstrated early neural differentiation, with clusters starting as early as 55 ms and showing multiple clusters of significant decoding throughout the trial, with Cohen’s *d* values exceeding 2.1. Cross-experiment analysis confirmed significant generalization, particularly for angry vs. neutral and sad vs. neutral pairs, with decoding onset within 85–120 ms.

More detailed results can be found in **Supplementary Table 1.**

## Discussion

The present study investigated the shared neural mechanisms underlying emotion recognition in real human faces and emojis using time-resolved EEG decoding. Our findings reveal robust spatio-temporal overlap in the neural processing of emotional expressions across these stimulus types, with implications for understanding how the brain generalizes social signal processing to symbolic representations.

Our results demonstrate that emotional expressions in both real faces and emojis can be decoded from EEG signals with high temporal precision, emerging as early as 100–120 ms and peaking around 145–160 ms over posterior-occipital and parietal regions. This aligns with prior studies showing early emotion discrimination in real faces (Y. Li et al., 2022; Muukkonen et al., 2020; Zhang et al., 2023) and extends these findings to emojis, suggesting that schematic facial representations engage rapid, feedforward mechanisms akin to naturalistic face processing (Motlagh et al., 2024). Notably, cross-classification analyses revealed significant generalization between real and emoji faces, particularly for neutral versus emotional contrasts (e.g., angry vs. neutral). This indicates that the brain employs overlapping neural codes for emotional content, irrespective of stimulus realism. The weaker decoding of fine-grained distinctions (e.g., angry vs. sad) mirrors challenges in real-face emotion discrimination (Smith & Smith, 2019), suggesting that categorical boundaries in both stimulus types may prioritize valence over exact emotional states. This hypothesis needs to be tested in future studies.

The early onset of emotion decoding (∼100 ms) for emojis challenges the notion that schematic faces require prolonged processing to extract affective meaning. Instead, our data suggest that emojis capitalize on pre-existing neural circuits optimized for rapid social signal extraction, in line with behavioral studies that find a speed advantage for both real and emoji faces when compared to word stimuli (Kaye et al., 2021).

The observed overlapping cross-classification results in the ca. 350-500 ms window suggest the involvement of higher-order cognitive and evaluative processes beyond early perceptual encoding. While the early decoding effects discussed above likely reflect rapid, feedforward mechanisms early visual categorization, the mid-late effects in this window indicate that real and emoji faces may share neural representations at a more abstract, integrative level of emotion processing. These later shared effects may capture ongoing emotional processing, which may reflect cognitive appraisal of the stimuli (Hajcak et al., 2010). As both our experiments used the same emotion recognition task, this similarity in neural codes could indicate the engagement of more controlled processes involved in decision-making. Future cross-classification studies using different procedures, e.g., tasks that do not involve the evaluation of facial expression, or using cognitive load paradigms, should investigate this possibility.

Our findings suggest that real and emoji faces share overlapping neural codes for emotion recognition, raising the question of whether these representations reflect domain-general emotion processing or are specific to faces and face-like stimuli. One possibility is that the observed cross-classification effects stem from a supra-modal emotion recognition system, in which the brain encodes emotional meaning independently of the stimulus format (Klasen et al., 2011; Peelen et al., 2010). This perspective aligns with theories proposing abstract, high-level representations of affect, where emotional signals are processed within a common neural network, potentially involving regions such as the superior temporal sulcus, anterior insula, and amygdala (Vytal & Hamann, 2010).

Alternatively, the shared neural responses between real and emoji faces may indicate a face-specific mechanism that extends to simplified, schematic representations. In our study, the time window that includes the N170 component was robustly implicated in both real face and emoji classification. This component is traditionally associated with face processing (Bentin et al., 1996; Hinojosa et al., 2015), suggesting that the symbolic emoji stimuli tap into the same perceptual and categorical processing pathways as naturalistic faces. Studies on other types of affective stimuli report later latencies in emotional content (Bo et al., 2022). This would support the notion that the brain treats emojis as a subset of facial stimuli, rather than engaging a more general emotional processing network at this stage. Future studies should explore these possibilities using cross-stimulus and cross-modal classification approaches (see Ambrus, 2024) that compare faces, emojis, and other emotionally expressive stimuli, such as words, body postures, scenes, and sounds.

### Future directions

Our study opens several promising avenues for further research into the neural mechanisms underlying affective processing. One important direction is to extend the range of emotional expressions examined, moving beyond basic categories such as happiness, sadness, anger, and neutrality, to include complex or mixed emotions. This is especially relevant in aiding the design of interactive affective systems and instruments aiming to reduce linguistic confounds (Kaye & Schweiger, 2023).

To disentangle the role of exposure-dependent learning, future work could investigate developmental cohorts: early sensitivity to emojis as faces in infants or young children with minimal cultural exposure would suggest innate biases, while delayed emergence would point to learned associations. Similarly, testing older adults who first encountered emojis after critical periods for face perception could reveal whether cross-classification diminishes with late exposure, implicating a more learning-dependent effect. Additionally, employing novel, culturally unfamiliar symbols - such as artificial emojis or platform-specific designs absent from participants’ prior experience - could clarify whether preserved cross-decoding relies on evolved mechanisms for schematic face-like patterns.

Complementing these investigations, cross-cultural studies could further explore whether the neural correspondence between emojis and real faces reflects universal mechanisms or culturally mediated processes. The comparison of neural responses to emotional expressions across populations with differing norms in emoji usage or interpretation could help us assess the universality of shared spatio-temporal codes. For example, cultures that attribute distinct meanings to specific emojis may exhibit reduced cross-classification for culturally ambiguous symbols. Conversely, preserved neural overlap for universally recognized emotions (e.g., happiness) across cultures would strengthen claims of evolved, domain-general mechanisms. Such work would not only clarify the global applicability of these findings but also identify cultural biases in how symbolic representations engage neural systems, bridging gaps between biological predispositions and sociocultural influences on emotion processing.

### Strengths of the current approach

The methodological framework of this study offers several key strengths. First, multivariate pattern analysis to EEG data provides unparalleled flexibility in probing shared neural representations across distinct stimulus types (Ambrus, 2024; Dalski, Kovács, & Ambrus, 2022a, 2022b; Dalski, Kovács, Wiese, et al., 2022; Klink et al., 2023; C. Li et al., 2022; Ozdemir & Ambrus, 2025). By training classifiers on one dataset (e.g., real faces) and testing on another (e.g., emojis) we isolated emotion-specific neural codes independent of low-level visual features. This approach inherently controlled for image properties (e.g., color, luminance, spatial frequency), as real faces and emojis differ markedly in their visual characteristics but converge in their emotional content. The use of identical experimental paradigms across the two experiments further minimized confounds related to the task.

Second, the study’s design is highly generalizable. The bidirectional cross-classification framework (Kaplan et al., 2015), here, testing both real-to-emoji and emoji-to-real decoding, provides a robust template for future research. For example, this approach could be extended to developmental cohorts to investigate how neural representations of symbolic stimuli emerge with age, or to cross-cultural studies comparing populations with divergent emoji interpretation norms. Similarly, clinical populations (e.g., individuals with autism spectrum disorder or prosopagnosia) could be tested to determine whether atypical face-processing mechanisms extend to schematic stimuli like emojis.

The data-driven nature of MVPA, free from *a priori* assumptions about regions or components of interest, enables the discovery of unanticipated patterns. This methodological agnosticism is particularly advantageous for studying novel stimuli like emojis, where established neural markers may not fully apply (Carlson et al., 2020; Grootswagers et al., 2017).

### Summary

This study investigated the neural dynamics underlying emotion recognition in emoji faces and contrasted it with that of real human faces, using EEG and multivariate pattern analysis. Across two experiments, participants performed emotion classification tasks while viewing happy, angry, sad, and neutral expressions in real faces or emojis. Robust decoding of emotional expressions emerged as early as 100–120 ms, peaking at 145–160 ms over posterior-occipital and parietal regions, with spatio-temporal patterns mirroring the N170 component - a hallmark of face-specific processing. Effects between the 350–500 ms post-stimulus time window were also observed. Critically, cross-classification analyses revealed bidirectional generalization of neural codes between real and emoji faces, particularly for emotion pairs involving neutral expressions (e.g., angry vs. neutral). This suggests that schematic emojis engage neural mechanisms tuned for facial expression processing, despite their artificiality. These findings advance our understanding of how symbolic and naturalistic social signals are processed, demonstrating the brain’s flexibility in the interpretation of abstract emotional cues. Our findings add to the growing body of work in semiotics that explores how humans assign emotional and semantic meaning to symbolic objects (Logi & Zappavigna, 2023), emphasizing the importance of emojis as more than mere decorative elements in communication. The results hold implications for digital communication, where emojis serve as proxies for facial expressions, and for designing emotionally responsive technologies that align with human social cognition. Future work should explore the extent of evolved and learned contributions to this overlap, including cross-cultural variability in emoji interpretation and developmental trajectories of symbolic emotion processing. This research further demonstrates the adaptability of human social perception in an increasingly online world.

## Supplementary material

**Supplementary Information.** Supplementary table and figures supplementing the results.

**Supplementary Table 1.** Detailed results of the statistical analysis in tabular form.

## Supplementary Information

## Supplementary Information 1

### Effect onsets

**Supplementary Information Table 1.** Onsets and peaks. Time-resolved spatio-temporal searchlight classification averaged across all sensors. Onsets were determined by two-tailed Bayesian *t*-tests against chance, with BF values exceeding 10 considered indicative of strong evidence. Supplements **Figure 2** in the main text.

|  |  | within-experiment |  | cross-experiment |  |
| --- | --- | --- | --- | --- | --- |
|  |  | emoji<br>faces | real<br>faces | real to<br>emoji faces | emoji to<br>real faces |
| emotion<br>4-class | onset | 100 ms | 120 ms | 120 ms | 125 ms |
|  | peak | 155 ms | 160 ms | 160 ms | 150 ms |
| | peak BF | $>10^6$ | $>10^4$ | $>10^7$ | $>10^6$ |
| happy vs neutral | onset | 115 ms | 120 ms | 105 ms | 115 ms |
|  | peak | 130 ms | 135 ms | 110 ms | 135 ms |
| | peak BF | 458.97 | $>10^3$ | $>10^3$ | $>10^4$ |
| angry vs neutral | onset | 105 ms | 100 ms | 125 ms | 130 ms |
|  | peak | 155 ms | 160 ms | 155 ms | 155 ms |
| | peak BF | $>10^7$ | $>10^6$ | $>10^7$ | $>10^6$ |
| sad vs neutral | onset | 120 ms | 125 ms | 130 ms | 130 ms |
|  | peak | 155 ms | 165 ms | 160 ms | 150 ms |
| | peak BF | $>10^7$ | $>10^3$ | $>10^6$ | $>10^6$ |
| happy vs angry | onset | 145 ms | 155 ms | 145 ms | 150 ms |
|  | peak | 160 ms | 175 ms | 165 ms | 180 ms |
| | peak BF | 156.02 | 543.72 | $>10^3$ | $>10^4$ |
| angry vs sad | onset | 110 ms | 215 ms | 130 ms | - |
|  | peak | 140 ms | 245 ms | 140 ms | - |
|  | peak BF | 782.21 | 430.02 | 57.1 | - |
| happy vs sad | onset | 185 ms | 170 ms | 135 ms | 170 ms |
|  | peak | 200 ms | 175 ms | 140 ms | 175 ms |
|  | peak BF | 155.56 | 49.46 | 41.68 | 13.9 |

**Supplementary Information Figure 1.**
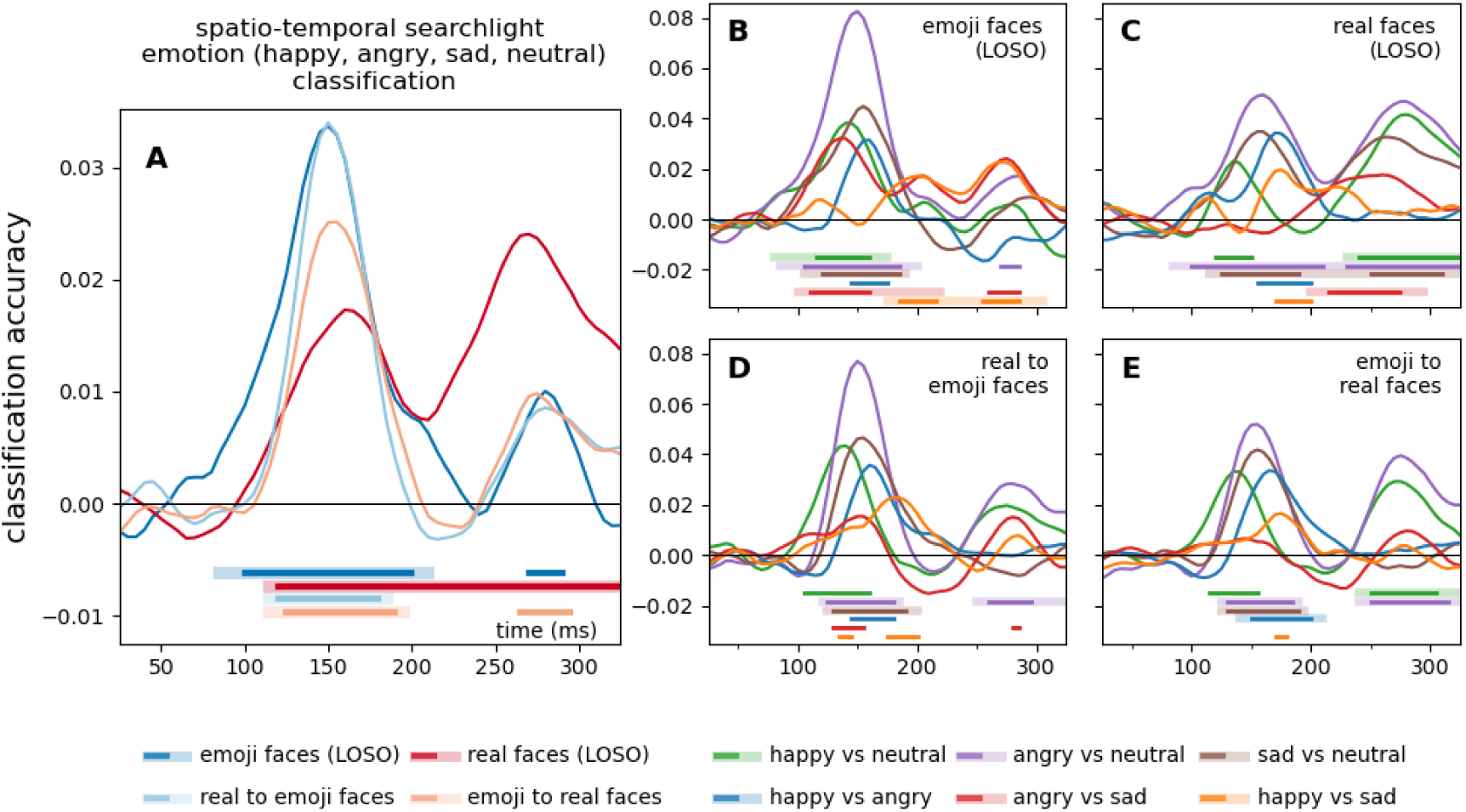
Spatiotemporal searchlight classification accuracies over all electrodes. Time-resolved spatio-temporal searchlight classification was performed on all channels and their neighboring electrodes. Classification performance was averaged across all sensors. Dark significance markers represent two-tailed Bayesian *t*-tests against chance, with BF values exceeding 10 considered indicative of strong evidence (horizontal significance markers). Light lines denote significant clusters revealed by the two-sided cluster permutation tests, *p* < 0.05. Supplements **Figure 2** in the main text.

## Supplementary Information 2

### Multivariate classification

**Supplementary Information Figure 2.**
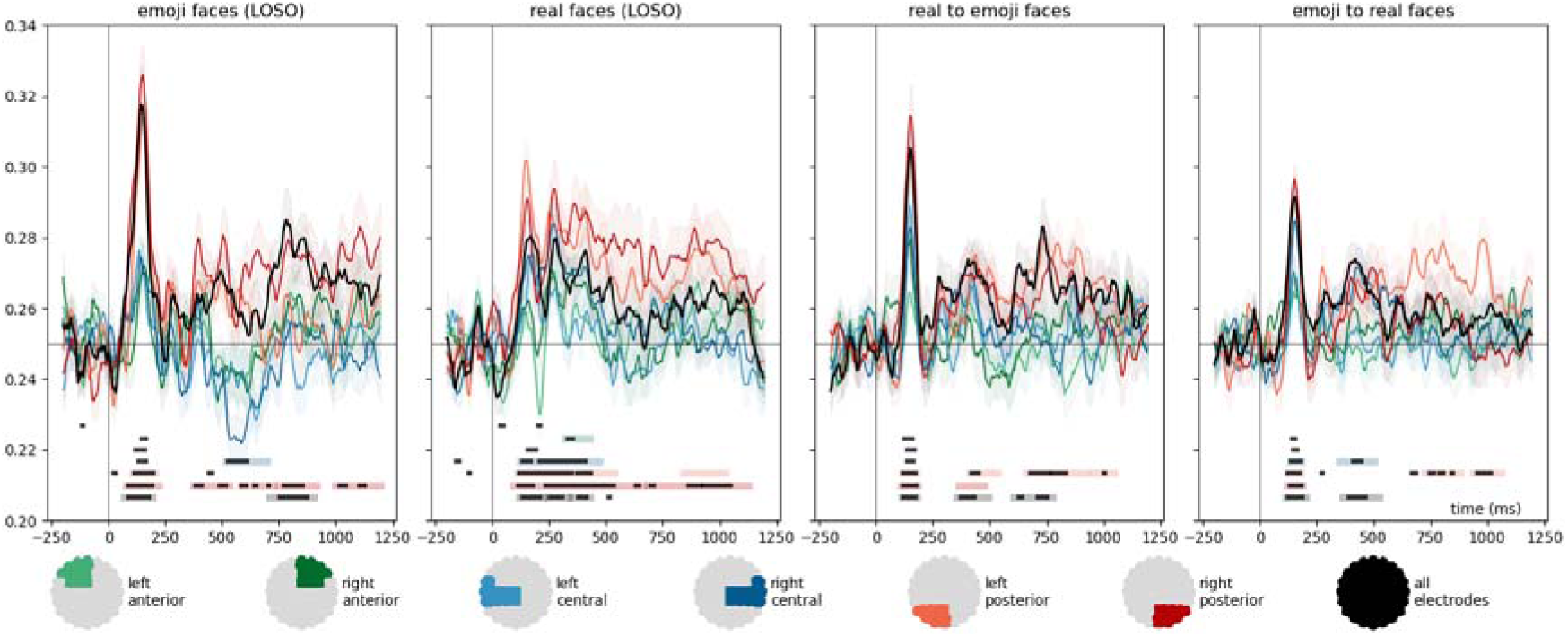
Time-resolved, within-experiment (leave-one-subject-out) and cross-experiment classification of facial expressions. In the within-experiment analyses (LOSO), classifiers were trained, in a leave-one-participant-out scheme to categorize the facial expressions of the stimuli, separately for the real and the emoji faces datasets. In the cross-experiment analyses, classifiers were trained on one dataset and were tested on the other. Error ranges represent ±SEM. Light lines denote significant clusters revealed by the two-sided cluster permutation tests, *p* < 0.05; dark lines denote results of the Bayesian statistical analyses, two-sided one-sample Bayesian *t*-tests, BF > 10, against chance (0.25). Results over all electrodes and pre-defined regions of interest are presented here. For detailed statistics, see **Supplementary Table 1**. Supplements **Figure 3** in the main text.

**Supplementary Information Figure 3.**
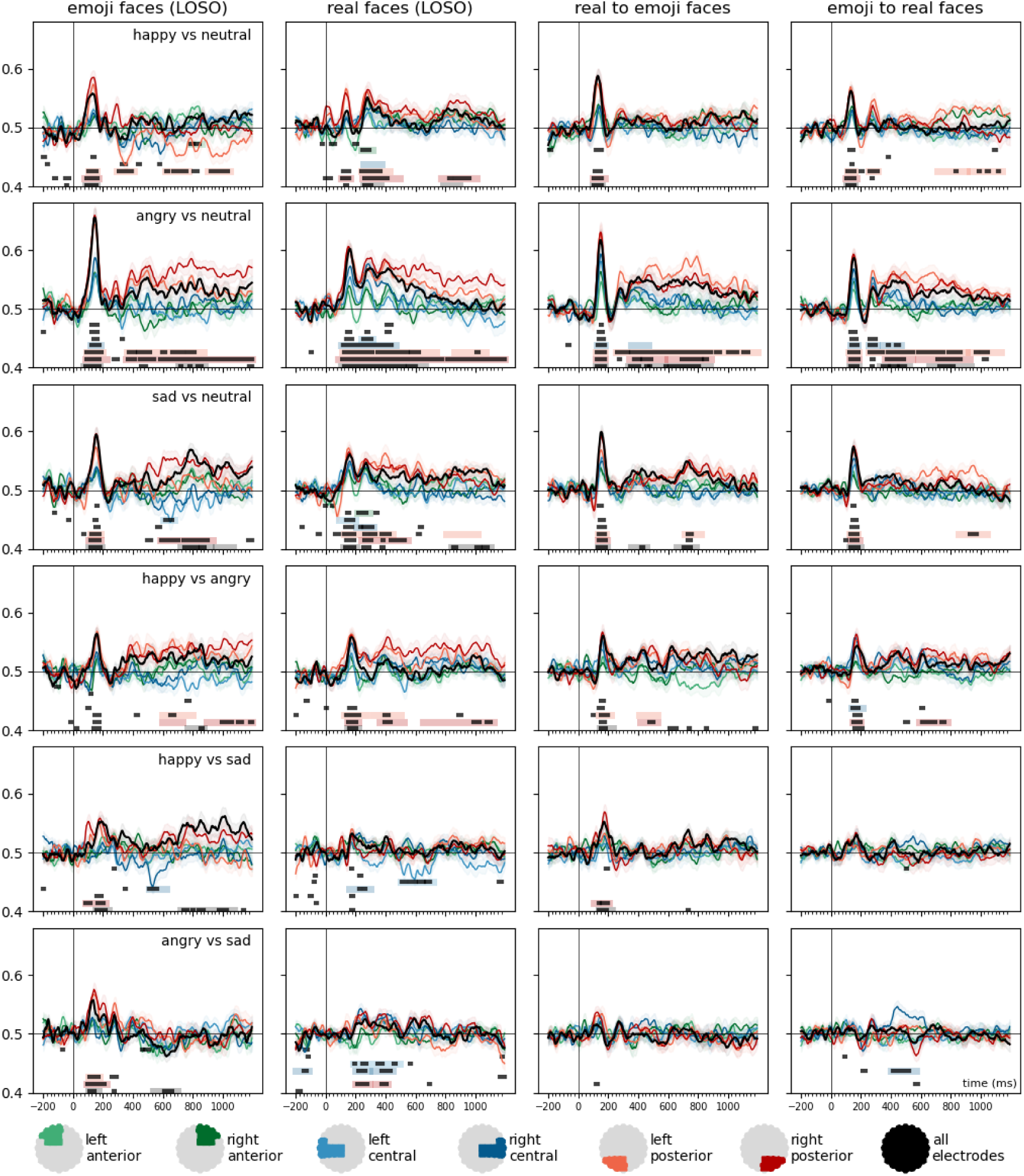
Time-resolved, within-experiment (leave-one-subject-out) and cross-experiment classification of facial expression pairs. For the within-experiment classification (LOSO), training was iteratively performed on six identities (3 male and 3 female) and tested on one left out in the real faces experiment, while in the emoji faces experiment, training was iteratively performed on five platforms and tested one platform left out. In the cross-experiment analyses, classifiers were trained on one dataset and were tested on the other. Error ranges represent ±SEM. Light lines denote significant clusters revealed by the two-sided cluster permutation tests, *p* < 0.05; dark lines denote results of the Bayesian statistical analyses, two-sided one-sample Bayesian *t*-tests, BF > 10, against chance (0.5). Results over all electrodes and pre-defined regions of interest are presented here. For detailed statistics, see **Supplementary Table 1**. Supplements **Figure 4** in the main text.

## Footnotes

1 Permission to display sample stimuli from the KDEF: “To the publisher: Researchers may always include sample images from KDEF or AKDEF in his/her manuscript when said manuscript is a doctoral thesis OR is a manuscript submitted to a scientific journal.” https://kdef.se/faq/using-and-publishing-kdef-and-akdef

## Notes

**Conflict of Interest**: The authors declare that they have no known competing financial interests or personal relationships that could have appeared to influence the work reported in this paper.

### Competing Interest Statement

The authors have declared no competing interest.

