## Supplementary Table 1 for "Shared Neural Codes for Emotion Recognition in Emoji and Human Faces": SupplementaryTable_1.html

  


### Supplementary Table 1.

  

Classification analyses

  


**A** emoji faces (LOSO) - emotion

**B** emoji faces (LOSO) - happy vs angry

**C** emoji faces (LOSO) - happy vs sad

**D** emoji faces (LOSO) - neutral vs happy

**E** emoji faces (LOSO) - angry vs sad

**F** emoji faces (LOSO) - neutral vs angry

**G** emoji faces (LOSO) - neutral vs sad

**H** real faces (LOSO) - emotion

**I** real faces (LOSO) - happy vs angry

**J** real faces (LOSO) - happy vs sad

**K** real faces (LOSO) - neutral vs happy

**L** real faces (LOSO) - angry vs sad

**M** real faces (LOSO) - neutral vs angry

**N** real faces (LOSO) - neutral vs sad

**O** real to emoji faces - emotion

**P** real to emoji faces - happy vs angry

**Q** real to emoji faces - happy vs sad

**R** real to emoji faces - neutral vs happy

**S** real to emoji faces - angry vs sad

**T** real to emoji faces - neutral vs angry

**U** real to emoji faces - neutral vs sad

**V** emoji to real faces - emotion

**W** emoji to real faces - happy vs angry

**X** emoji to real faces - happy vs sad

**Y** emoji to real faces - neutral vs happy

**Z** emoji to real faces - angry vs sad

**AA** emoji to real faces - neutral vs angry

**AB** emoji to real faces - neutral vs sad

A) emoji faces (LOSO) - emotion

  
|  | time window | peak latency | cluster *p* | peak Cohen's *d* |  | | | |  |
| **all electrodes** | 70 - 195 ms | 145 ms | 0.0086 | 1.2134 |  | | | |  |
 705 - 900 ms | 780 ms | 0.0054 | 0.7997 |  | | | ||  |  | | | | | | | | |

Time-resolved classification, cluster permutation tests

|  | **left hemisphere** | | | | **right hemisphere** | | | |  |
|  | time window | peak latency | cluster *p* | peak Cohen's *d* | time window | peak latency | cluster *p* | peak Cohen's *d* |  |
| **anterior** |  | | | |  | | | |  |
| **central** |  | | | | 515 - 695 ms | 685 ms | 0.001 | -0.4374 |  |
| **posterior** | 105 - 205 ms | 150 ms | 0.0104 | 1.7153 | 80 - 220 ms | 150 ms | 0.0056 | 1.7358 |  |
  | | | | 375 - 530 ms | 505 ms | 0.0171 | 0.7426 |  | | | | 580 - 725 ms | 655 ms | 0.0224 | 0.6267 |  | | | | 750 - 915 ms | 790 ms | 0.0144 | 0.8313 |  | | | | 1000 - 1195 ms | 1105 ms | 0.0121 | 0.6553 |

  

Time-resolved classification, Bayesian statistics

|  | -200 | -195 | -190 | -185 | -180 | -175 | -170 | -165 | -160 | -155 | -150 | -145 | -140 | -135 | -130 | -125 | -120 | -115 | -110 | -105 | -100 | -95 | -90 | -85 | -80 | -75 | -70 | -65 | -60 | -55 | -50 | -45 | -40 | -35 | -30 | -25 | -20 | -15 | -10 | -5 | 0 | 5 | 10 | 15 | 20 | 25 | 30 | 35 | 40 | 45 | 50 | 55 | 60 | 65 | 70 | 75 | 80 | 85 | 90 | 95 | 100 | 105 | 110 | 115 | 120 | 125 | 130 | 135 | 140 | 145 | 150 | 155 | 160 | 165 | 170 | 175 | 180 | 185 | 190 | 195 | 200 | 205 | 210 | 215 | 220 | 225 | 230 | 235 | 240 | 245 | 250 | 255 | 260 | 265 | 270 | 275 | 280 | 285 | 290 | 295 | 300 | 305 | 310 | 315 | 320 | 325 | 330 | 335 | 340 | 345 | 350 | 355 | 360 | 365 | 370 | 375 | 380 | 385 | 390 | 395 | 400 | 405 | 410 | 415 | 420 | 425 | 430 | 435 | 440 | 445 | 450 | 455 | 460 | 465 | 470 | 475 | 480 | 485 | 490 | 495 | 500 | 505 | 510 | 515 | 520 | 525 | 530 | 535 | 540 | 545 | 550 | 555 | 560 | 565 | 570 | 575 | 580 | 585 | 590 | 595 | 600 | 605 | 610 | 615 | 620 | 625 | 630 | 635 | 640 | 645 | 650 | 655 | 660 | 665 | 670 | 675 | 680 | 685 | 690 | 695 | 700 | 705 | 710 | 715 | 720 | 725 | 730 | 735 | 740 | 745 | 750 | 755 | 760 | 765 | 770 | 775 | 780 | 785 | 790 | 795 | 800 | 805 | 810 | 815 | 820 | 825 | 830 | 835 | 840 | 845 | 850 | 855 | 860 | 865 | 870 | 875 | 880 | 885 | 890 | 895 | 900 | 905 | 910 | 915 | 920 | 925 | 930 | 935 | 940 | 945 | 950 | 955 | 960 | 965 | 970 | 975 | 980 | 985 | 990 | 995 | 1000 | 1005 | 1010 | 1015 | 1020 | 1025 | 1030 | 1035 | 1040 | 1045 | 1050 | 1055 | 1060 | 1065 | 1070 | 1075 | 1080 | 1085 | 1090 | 1095 | 1100 | 1105 | 1110 | 1115 | 1120 | 1125 | 1130 | 1135 | 1140 | 1145 | 1150 | 1155 | 1160 | 1165 | 1170 | 1175 | 1180 | 1185 | 1190 | 1195 |
| --- | --- | --- | --- | --- | --- | --- | --- | --- | --- | --- | --- | --- | --- | --- | --- | --- | --- | --- | --- | --- | --- | --- | --- | --- | --- | --- | --- | --- | --- | --- | --- | --- | --- | --- | --- | --- | --- | --- | --- | --- | --- | --- | --- | --- | --- | --- | --- | --- | --- | --- | --- | --- | --- | --- | --- | --- | --- | --- | --- | --- | --- | --- | --- | --- | --- | --- | --- | --- | --- | --- | --- | --- | --- | --- | --- | --- | --- | --- | --- | --- | --- | --- | --- | --- | --- | --- | --- | --- | --- | --- | --- | --- | --- | --- | --- | --- | --- | --- | --- | --- | --- | --- | --- | --- | --- | --- | --- | --- | --- | --- | --- | --- | --- | --- | --- | --- | --- | --- | --- | --- | --- | --- | --- | --- | --- | --- | --- | --- | --- | --- | --- | --- | --- | --- | --- | --- | --- | --- | --- | --- | --- | --- | --- | --- | --- | --- | --- | --- | --- | --- | --- | --- | --- | --- | --- | --- | --- | --- | --- | --- | --- | --- | --- | --- | --- | --- | --- | --- | --- | --- | --- | --- | --- | --- | --- | --- | --- | --- | --- | --- | --- | --- | --- | --- | --- | --- | --- | --- | --- | --- | --- | --- | --- | --- | --- | --- | --- | --- | --- | --- | --- | --- | --- | --- | --- | --- | --- | --- | --- | --- | --- | --- | --- | --- | --- | --- | --- | --- | --- | --- | --- | --- | --- | --- | --- | --- | --- | --- | --- | --- | --- | --- | --- | --- | --- | --- | --- | --- | --- | --- | --- | --- | --- | --- | --- | --- | --- | --- | --- | --- | --- | --- | --- | --- | --- | --- | --- | --- | --- | --- | --- | --- | --- | --- | --- | --- | --- | --- | --- | --- | --- | --- | --- | --- | --- | --- | --- | --- | --- | --- |
| left anterior | 0.456571 | 0.566976 | 0.457029 | 0.437363 | 0.295597 | 0.386213 | 0.602464 | 0.502407 | 0.389310 | 0.423444 | 0.537324 | 0.577476 | 0.230589 | 0.215523 | 0.255863 | 0.624652 | 2.706883 | 18.056547 | 19.229512 | 2.439931 | 1.470845 | 0.522286 | 0.210974 | 0.245805 | 0.343738 | 0.489076 | 0.494639 | 0.563228 | 0.379179 | 0.294642 | 0.292111 | 0.262636 | 0.230055 | 0.232831 | 0.223918 | 0.246092 | 0.228174 | 0.215827 | 0.211899 | 0.213129 | 0.239012 | 0.327389 | 0.538116 | 0.821431 | 1.596168 | 2.149815 | 1.262988 | 0.446875 | 0.258331 | 0.223209 | 0.218003 | 0.211433 | 0.220196 | 0.224983 | 0.211703 | 0.228636 | 0.246201 | 0.215770 | 0.242485 | 0.317548 | 0.452787 | 0.568246 | 1.001346 | 1.526489 | 0.951639 | 0.669043 | 0.897594 | 1.219816 | 1.732347 | 2.044564e+00 | 2.114682e+00 | 2.980488e+00 | 3.329967e+00 | 2.575344 | 1.519997 | 1.384239 | 1.751822 | 1.547716 | 1.160723 | 0.890036 | 0.772230 | 0.724882 | 0.519770 | 0.290767 | 0.227831 | 0.224973 | 0.254972 | 0.256207 | 0.219127 | 0.211810 | 0.226494 | 0.280221 | 0.368591 | 0.382990 | 0.409067 | 0.517167 | 0.626331 | 0.651365 | 0.550945 | 0.393576 | 0.241359 | 0.210835 | 0.226639 | 0.241359 | 0.249059 | 0.253383 | 0.245319 | 0.213115 | 0.230079 | 0.321707 | 0.307859 | 0.307540 | 0.333543 | 0.312911 | 0.254313 | 0.233957 | 0.226937 | 0.254878 | 0.255563 | 0.242281 | 0.235438 | 0.239893 | 0.257908 | 0.272072 | 0.287422 | 0.277742 | 0.264028 | 0.231417 | 0.219037 | 0.212698 | 0.245049 | 0.319012 | 0.362643 | 0.403987 | 0.351673 | 0.264209 | 0.249629 | 0.250463 | 0.245757 | 0.253206 | 0.273205 | 0.322608 | 0.728711 | 0.990037 | 0.825664 | 0.845838 | 0.602446 | 0.447366 | 0.405190 | 0.355934 | 0.357383 | 0.443475 | 0.505689 | 0.715894 | 0.726305 | 0.570069 | 0.405513 | 0.402424 | 0.428698 | 0.676473 | 0.701874 | 0.587185 | 0.497386 | 0.530635 | 0.466480 | 0.302420 | 0.222889 | 0.211180 | 0.212048 | 0.211516 | 0.212098 | 0.211423 | 0.211796 | 0.211389 | 0.211384 | 0.211104 | 0.215739 | 0.223738 | 0.246564 | 0.234218 | 0.216649 | 0.210806 | 0.211048 | 0.215914 | 0.230198 | 0.248046 | 0.276835 | 0.261916 | 0.242801 | 0.251728 | 0.257008 | 0.262643 | 0.251709 | 0.255150 | 0.303537 | 0.356341 | 0.398934 | 0.401974 | 0.529839 | 0.713267 | 0.965331 | 0.903295 | 0.759729 | 0.681499 | 0.816560 | 0.702504 | 0.531052 | 0.401076 | 0.294222 | 0.244530 | 0.239338 | 0.216282 | 0.211211 | 0.212092 | 0.210833 | 0.219150 | 0.252241 | 0.234679 | 0.235260 | 0.241146 | 0.246790 | 0.243745 | 0.235748 | 0.240971 | 0.301503 | 0.283134 | 0.251011 | 0.217913 | 0.211293 | 0.211738 | 0.219547 | 0.276439 | 0.257820 | 0.324495 | 0.314529 | 0.297504 | 0.374919 | 0.348273 | 0.328623 | 0.293206 | 0.242236 | 0.237077 | 0.229729 | 0.233201 | 0.234719 | 0.242129 | 0.243948 | 0.228214 | 0.219687 | 0.217592 | 0.211112 | 0.211053 | 0.211071 | 0.213058 | 0.214324 | 0.211472 | 0.212817 | 0.212895 | 0.215970 | 0.231919 | 0.248996 | 0.247759 | 0.244562 | 0.229778 | 0.226669 | 0.226474 | 0.218843 | 0.217678 | 0.214120 | 0.217214 | 0.215623 | 0.213340 | 0.210813 | 0.210814 | 0.218653 | 0.218657 | 0.224260 | 0.236552 | 0.247997 | 0.296268 |
| right anterior | 3.897791 | 1.452660 | 0.979762 | 0.803648 | 0.409116 | 0.271539 | 0.213703 | 0.216106 | 0.220884 | 0.225907 | 0.258117 | 0.244271 | 0.232387 | 0.301510 | 0.353135 | 0.255100 | 0.214395 | 0.213614 | 0.215091 | 0.210950 | 0.245283 | 0.309343 | 0.362913 | 0.324102 | 0.329332 | 0.416841 | 1.898148 | 3.706817 | 1.638187 | 1.093919 | 1.488265 | 1.825840 | 0.855955 | 0.239108 | 0.218889 | 0.211642 | 0.217239 | 0.234979 | 0.275492 | 0.257697 | 0.230218 | 0.212176 | 0.234416 | 0.272736 | 0.364655 | 0.413954 | 0.493514 | 0.382849 | 0.384027 | 0.361163 | 0.344783 | 0.341475 | 0.304743 | 0.261868 | 0.349074 | 0.355342 | 0.286873 | 0.298086 | 0.317803 | 0.316420 | 0.285504 | 0.242608 | 0.218174 | 0.211206 | 0.233788 | 0.381796 | 0.765902 | 1.532888 | 4.713345 | 1.538519e+01 | 2.453081e+01 | 1.871921e+01 | 1.377089e+01 | 12.522686 | 5.653482 | 1.901019 | 0.599202 | 0.304404 | 0.235386 | 0.217755 | 0.211384 | 0.214540 | 0.260662 | 0.350588 | 0.373363 | 0.482290 | 0.791387 | 1.158112 | 1.203985 | 0.723119 | 0.432765 | 0.393734 | 0.275739 | 0.214145 | 0.211999 | 0.212491 | 0.211511 | 0.245970 | 0.306153 | 0.375822 | 0.544100 | 0.561232 | 0.404416 | 0.259960 | 0.211746 | 0.239996 | 0.282964 | 0.304844 | 0.387734 | 0.389272 | 0.315693 | 0.289554 | 0.324151 | 0.329997 | 0.395905 | 0.365610 | 0.347483 | 0.451465 | 0.413960 | 0.358004 | 0.359086 | 0.387418 | 0.491946 | 0.548166 | 0.793305 | 0.914840 | 0.985235 | 0.599237 | 0.402227 | 0.276657 | 0.213124 | 0.264378 | 0.347467 | 0.619950 | 0.774025 | 1.341104 | 1.642795 | 0.952885 | 0.834219 | 0.814002 | 0.743205 | 0.744801 | 0.532828 | 0.361734 | 0.390225 | 0.418790 | 0.417640 | 0.390200 | 0.311895 | 0.319246 | 0.328688 | 0.312707 | 0.344492 | 0.365963 | 0.332324 | 0.360059 | 0.365642 | 0.373904 | 0.507012 | 0.524983 | 0.657664 | 0.988414 | 1.231494 | 0.836712 | 0.684333 | 0.496484 | 0.348769 | 0.278220 | 0.235997 | 0.219168 | 0.241532 | 0.250571 | 0.246045 | 0.250724 | 0.289356 | 0.272388 | 0.261877 | 0.237395 | 0.224234 | 0.215206 | 0.219984 | 0.213816 | 0.219042 | 0.218094 | 0.221975 | 0.264198 | 0.328568 | 0.340345 | 0.277506 | 0.240020 | 0.248944 | 0.244749 | 0.222611 | 0.217524 | 0.259908 | 0.296891 | 0.357341 | 0.450489 | 0.636680 | 0.549129 | 0.464892 | 0.416797 | 0.398276 | 0.416424 | 0.430714 | 0.483597 | 0.572367 | 0.609134 | 0.612794 | 0.741727 | 0.683331 | 0.589617 | 0.406039 | 0.276028 | 0.270621 | 0.244633 | 0.224216 | 0.230675 | 0.224799 | 0.248449 | 0.409580 | 0.559958 | 0.782535 | 1.027206 | 1.513888 | 2.551695 | 1.685897 | 0.897183 | 0.719977 | 0.767740 | 0.802775 | 0.673662 | 0.688579 | 0.594387 | 0.463145 | 0.364873 | 0.295668 | 0.257293 | 0.247761 | 0.237680 | 0.263892 | 0.324225 | 0.338330 | 0.274079 | 0.249021 | 0.236369 | 0.230331 | 0.216319 | 0.210901 | 0.210830 | 0.219355 | 0.232004 | 0.245773 | 0.267084 | 0.295596 | 0.311303 | 0.368848 | 0.430292 | 0.497960 | 0.828828 | 1.402218 | 2.045340 | 2.774937 | 2.001256 | 1.737641 | 2.233791 | 0.903451 | 0.495302 | 0.289379 | 0.255435 | 0.247531 | 0.256621 | 0.256546 | 0.288286 | 0.354904 | 0.451612 | 0.404925 | 0.471185 | 0.446990 | 0.381362 |
| left central | 1.092882 | 1.621793 | 0.859299 | 0.396909 | 0.263528 | 0.211151 | 0.241009 | 0.266851 | 0.384982 | 0.482889 | 0.676932 | 0.400530 | 0.243750 | 0.243348 | 0.337023 | 0.417305 | 0.421167 | 0.312567 | 0.348528 | 0.458679 | 0.440368 | 0.310270 | 0.238968 | 0.235011 | 0.213252 | 0.218906 | 0.273507 | 0.328747 | 0.350002 | 0.616126 | 0.702957 | 0.463572 | 0.497043 | 0.447558 | 0.477117 | 0.556909 | 0.496269 | 0.398017 | 0.342201 | 0.255221 | 0.221666 | 0.212015 | 0.232387 | 0.282380 | 0.289066 | 0.253952 | 0.269211 | 0.270727 | 0.232842 | 0.210806 | 0.222596 | 0.266204 | 0.413118 | 0.610237 | 0.599597 | 0.444018 | 0.374869 | 0.351825 | 0.366240 | 0.340674 | 0.477196 | 1.153958 | 1.697527 | 3.703073 | 18.205754 | 125.971957 | 6016.535456 | 54475.324199 | 5236.467747 | 1.167239e+03 | 3.640855e+02 | 4.812322e+01 | 2.469547e+01 | 7.014606 | 0.823413 | 0.499980 | 0.314363 | 0.344612 | 0.403591 | 0.289005 | 0.256976 | 0.281905 | 0.246138 | 0.219981 | 0.210960 | 0.229721 | 0.254962 | 0.493146 | 0.881555 | 0.799166 | 0.653655 | 0.582239 | 0.421249 | 0.364507 | 0.242381 | 0.241532 | 0.235927 | 0.233442 | 0.286066 | 0.317775 | 0.352415 | 0.430583 | 0.379517 | 0.400969 | 0.425589 | 0.450580 | 0.604836 | 0.951441 | 1.289624 | 1.225753 | 1.224505 | 1.137459 | 0.723356 | 0.470849 | 0.316149 | 0.259224 | 0.221550 | 0.210829 | 0.212879 | 0.213079 | 0.212176 | 0.211320 | 0.211225 | 0.231772 | 0.278834 | 0.261455 | 0.233937 | 0.222784 | 0.219886 | 0.221029 | 0.227606 | 0.260408 | 0.344807 | 0.549449 | 0.876608 | 1.534055 | 1.676575 | 1.457377 | 0.763338 | 0.525615 | 0.441514 | 0.451061 | 0.369756 | 0.387634 | 0.382175 | 0.357810 | 0.327983 | 0.262736 | 0.241517 | 0.276069 | 0.297282 | 0.326122 | 0.396326 | 0.498853 | 0.535575 | 0.476547 | 0.390239 | 0.317959 | 0.296528 | 0.322982 | 0.369740 | 0.429834 | 0.468941 | 0.575805 | 0.747321 | 1.172174 | 1.508907 | 2.367819 | 3.630052 | 7.199177 | 8.576537 | 9.309328 | 7.185016 | 3.699618 | 1.520077 | 0.902431 | 0.518170 | 0.382009 | 0.339128 | 0.278613 | 0.273489 | 0.255722 | 0.234483 | 0.223394 | 0.215835 | 0.211877 | 0.211942 | 0.211478 | 0.215136 | 0.212805 | 0.210948 | 0.210892 | 0.211360 | 0.211483 | 0.225635 | 0.242679 | 0.255797 | 0.289131 | 0.276220 | 0.264776 | 0.222103 | 0.218812 | 0.254567 | 0.289204 | 0.376593 | 0.534693 | 0.872008 | 0.979676 | 0.710776 | 0.878682 | 0.941371 | 0.772964 | 0.597981 | 0.476677 | 0.364727 | 0.314845 | 0.279831 | 0.283719 | 0.305410 | 0.260224 | 0.233467 | 0.229470 | 0.230014 | 0.222788 | 0.216761 | 0.213132 | 0.219449 | 0.235168 | 0.246225 | 0.239191 | 0.221232 | 0.218079 | 0.251780 | 0.261326 | 0.307987 | 0.372253 | 0.463632 | 0.597826 | 0.685469 | 0.445337 | 0.381448 | 0.368685 | 0.335065 | 0.309688 | 0.347775 | 0.331625 | 0.327853 | 0.301787 | 0.242844 | 0.218340 | 0.211171 | 0.228353 | 0.244686 | 0.268322 | 0.282832 | 0.259729 | 0.256039 | 0.256598 | 0.264152 | 0.262310 | 0.250769 | 0.255154 | 0.271207 | 0.257056 | 0.230707 | 0.210806 | 0.224231 | 0.237416 | 0.244575 | 0.247863 | 0.245915 | 0.234056 | 0.219942 | 0.212079 | 0.211341 | 0.215577 | 0.223693 | 0.224788 | 0.222361 | 0.243266 |
| right central | 0.495846 | 0.280320 | 0.223001 | 0.211338 | 0.220898 | 0.221789 | 0.211105 | 0.224108 | 0.214810 | 0.211263 | 0.218110 | 0.211863 | 0.236605 | 0.408918 | 0.256412 | 0.279307 | 0.317409 | 0.389123 | 0.721481 | 0.724751 | 0.326348 | 0.296694 | 0.216233 | 0.214910 | 0.230306 | 0.255599 | 0.318606 | 0.249655 | 0.216514 | 0.256596 | 0.288357 | 0.248723 | 0.215732 | 0.232032 | 0.235138 | 0.219361 | 0.211762 | 0.213399 | 0.251007 | 0.221534 | 0.218296 | 0.277259 | 0.327147 | 0.288213 | 0.247792 | 0.210856 | 0.289442 | 0.364210 | 0.245275 | 0.210837 | 0.214698 | 0.217226 | 0.357649 | 0.687936 | 1.001309 | 0.622456 | 0.423903 | 0.372702 | 0.479265 | 0.418077 | 0.338256 | 0.318979 | 0.404351 | 0.907738 | 1.489091 | 1.689184 | 4.797725 | 20.156505 | 47.923068 | 9.531859e+01 | 7.267011e+01 | 4.098251e+01 | 5.800486e+01 | 15.770390 | 6.029469 | 2.063326 | 1.383175 | 1.266690 | 1.202604 | 0.904227 | 0.653070 | 0.356353 | 0.227556 | 0.235294 | 0.406340 | 0.607124 | 0.834283 | 0.740955 | 0.437591 | 0.265641 | 0.210817 | 0.216619 | 0.224673 | 0.251542 | 0.298642 | 0.305499 | 0.297404 | 0.256237 | 0.218215 | 0.220801 | 0.304761 | 0.404255 | 0.601986 | 1.030144 | 1.166217 | 1.833390 | 2.034085 | 2.720469 | 3.723716 | 3.757311 | 1.792959 | 1.221911 | 0.496574 | 0.237953 | 0.211003 | 0.256084 | 0.394084 | 0.638225 | 1.128916 | 1.515364 | 0.850248 | 0.423866 | 0.248950 | 0.210806 | 0.272639 | 0.499170 | 0.782951 | 0.976802 | 0.930248 | 0.708581 | 0.424473 | 0.322194 | 0.313051 | 0.403257 | 0.473439 | 0.425589 | 0.453295 | 0.517632 | 0.424527 | 0.382818 | 0.378745 | 0.502152 | 0.901373 | 2.024931 | 8.424144 | 47.102619 | 119.411295 | 212.961575 | 225.709792 | 167.550578 | 120.751882 | 148.200415 | 146.980571 | 245.841156 | 242.970192 | 177.168118 | 134.826938 | 185.408124 | 129.200983 | 117.528752 | 60.200329 | 28.160233 | 14.976163 | 7.160927 | 6.308012 | 5.603683 | 3.062573 | 3.987128 | 4.441592 | 4.842416 | 5.987371 | 6.805478 | 5.842386 | 5.497733 | 4.443668 | 3.372601 | 2.289705 | 1.569557 | 1.449434 | 1.599022 | 1.103431 | 0.607983 | 0.361060 | 0.324877 | 0.370378 | 0.462198 | 0.446596 | 0.590594 | 0.789514 | 1.294318 | 1.204449 | 0.824640 | 0.387437 | 0.244863 | 0.215556 | 0.210944 | 0.219248 | 0.215840 | 0.232518 | 0.257865 | 0.312624 | 0.319266 | 0.277092 | 0.263677 | 0.311623 | 0.260669 | 0.276850 | 0.285285 | 0.303314 | 0.308188 | 0.276595 | 0.266370 | 0.269502 | 0.249488 | 0.220470 | 0.219668 | 0.217147 | 0.219479 | 0.226011 | 0.223741 | 0.214615 | 0.217124 | 0.212934 | 0.219619 | 0.218114 | 0.214200 | 0.224501 | 0.248444 | 0.296553 | 0.289471 | 0.234805 | 0.219918 | 0.211543 | 0.213740 | 0.241799 | 0.295134 | 0.288675 | 0.230524 | 0.212886 | 0.249018 | 0.279391 | 0.316580 | 0.273026 | 0.231452 | 0.215065 | 0.219083 | 0.284138 | 0.367948 | 0.346906 | 0.285817 | 0.249639 | 0.237780 | 0.222305 | 0.230606 | 0.229931 | 0.248634 | 0.259923 | 0.246944 | 0.248688 | 0.283631 | 0.311843 | 0.382517 | 0.315509 | 0.274069 | 0.289500 | 0.297846 | 0.267583 | 0.221237 | 0.211681 | 0.212686 | 0.213202 | 0.212104 | 0.211339 | 0.210981 | 0.211998 | 0.242634 | 0.310652 | 0.383454 | 0.451224 | 0.523531 |
| left posterior | 0.217495 | 0.218464 | 0.277097 | 0.355526 | 0.358303 | 0.410384 | 0.716499 | 0.863608 | 0.675262 | 0.485874 | 0.351667 | 0.297107 | 0.239138 | 0.210879 | 0.211397 | 0.215804 | 0.234378 | 0.319329 | 0.396816 | 0.435020 | 0.418477 | 0.425233 | 0.547083 | 0.941218 | 1.050323 | 0.844951 | 0.787986 | 1.408684 | 1.438763 | 0.336034 | 0.213895 | 0.241468 | 0.411949 | 0.482398 | 0.550994 | 0.471543 | 0.248433 | 0.261786 | 0.220526 | 0.231001 | 0.248758 | 0.365364 | 0.976089 | 2.801467 | 12.931214 | 39.334229 | 19.217423 | 5.588591 | 0.548316 | 0.222773 | 0.210883 | 0.221452 | 0.226485 | 0.212518 | 0.312149 | 0.724761 | 0.697600 | 0.251265 | 0.231618 | 0.383905 | 0.768671 | 2.680799 | 7.845538 | 41.956582 | 201.116693 | 840.541690 | 2669.241203 | 14579.782372 | 83994.548440 | 5.907211e+05 | 1.419779e+06 | 5.652353e+06 | 3.290231e+06 | 136488.922348 | 9684.242182 | 954.907121 | 207.897690 | 43.670684 | 26.375951 | 12.864254 | 10.363332 | 3.969519 | 1.247889 | 0.521743 | 0.319120 | 0.238528 | 0.219957 | 0.222024 | 0.222773 | 0.228936 | 0.302729 | 0.430067 | 0.652054 | 1.457483 | 2.710942 | 2.890035 | 3.120257 | 2.182583 | 1.379714 | 0.881677 | 0.517476 | 0.356720 | 0.266396 | 0.220298 | 0.212228 | 0.211113 | 0.216746 | 0.235579 | 0.285339 | 0.319875 | 0.263624 | 0.264793 | 0.220491 | 0.219088 | 0.306254 | 0.579783 | 1.081709 | 1.242919 | 2.165583 | 2.543360 | 2.354818 | 1.832178 | 1.368952 | 1.248378 | 1.370320 | 1.910224 | 3.013881 | 5.448285 | 5.968921 | 10.606857 | 12.775039 | 11.492348 | 4.264952 | 2.059178 | 1.338590 | 1.195789 | 1.448170 | 1.743055 | 2.074983 | 2.016046 | 1.946154 | 1.420688 | 1.094498 | 0.790440 | 0.593240 | 0.430106 | 0.341084 | 0.298044 | 0.293291 | 0.317737 | 0.366397 | 0.421254 | 0.512011 | 0.725696 | 0.705838 | 0.693768 | 0.619166 | 0.571629 | 0.654829 | 0.552017 | 0.520707 | 0.686635 | 0.789619 | 0.913405 | 0.641685 | 0.382816 | 0.377574 | 0.356987 | 0.379732 | 0.377016 | 0.324250 | 0.283974 | 0.261778 | 0.242064 | 0.220674 | 0.214585 | 0.250475 | 0.298001 | 0.298188 | 0.243226 | 0.211960 | 0.214336 | 0.236083 | 0.248823 | 0.244779 | 0.231405 | 0.218739 | 0.217112 | 0.235986 | 0.239889 | 0.223218 | 0.210900 | 0.225376 | 0.295122 | 0.434480 | 0.732189 | 0.795959 | 0.736152 | 0.872804 | 0.659679 | 0.420376 | 0.351348 | 0.264010 | 0.240276 | 0.220313 | 0.210811 | 0.211560 | 0.217445 | 0.221520 | 0.215355 | 0.210829 | 0.224548 | 0.249515 | 0.276467 | 0.309484 | 0.344272 | 0.297032 | 0.278983 | 0.282468 | 0.285274 | 0.270617 | 0.258545 | 0.242468 | 0.261309 | 0.264329 | 0.253899 | 0.249375 | 0.249927 | 0.230911 | 0.233948 | 0.234415 | 0.217928 | 0.210812 | 0.214782 | 0.218815 | 0.221119 | 0.239252 | 0.258256 | 0.260203 | 0.262367 | 0.223371 | 0.211183 | 0.214063 | 0.222883 | 0.233791 | 0.231856 | 0.234239 | 0.225402 | 0.223911 | 0.231683 | 0.252227 | 0.267192 | 0.319868 | 0.397139 | 0.510197 | 0.684050 | 0.819460 | 0.887075 | 1.018801 | 0.889522 | 0.703682 | 0.498512 | 0.356285 | 0.291702 | 0.273084 | 0.236944 | 0.236060 | 0.240262 | 0.241742 | 0.243915 | 0.252442 | 0.263385 | 0.317812 | 0.338142 | 0.354689 | 0.405618 | 0.482675 | 0.540540 | 0.570593 | 0.512234 |
| right posterior | 0.365645 | 0.425612 | 0.306308 | 0.259534 | 0.250158 | 0.220577 | 0.211338 | 0.212645 | 0.216103 | 0.215460 | 0.219100 | 0.246002 | 0.257164 | 0.437524 | 1.452292 | 1.664451 | 0.875707 | 1.079176 | 0.552615 | 0.576575 | 0.347457 | 0.217356 | 0.219619 | 0.304136 | 0.571138 | 4.521889 | 9.073171 | 8.297730 | 3.021891 | 2.499437 | 2.732623 | 1.423639 | 0.493245 | 0.401939 | 0.312959 | 0.342439 | 0.386460 | 0.366443 | 0.406440 | 0.314354 | 0.243239 | 0.325011 | 0.393521 | 0.486102 | 0.474452 | 0.366886 | 0.296285 | 0.304515 | 0.223816 | 0.216604 | 0.226604 | 0.221605 | 0.219723 | 0.253926 | 0.398689 | 0.681968 | 1.710767 | 17.169592 | 143.902372 | 838.897324 | 2159.742115 | 5082.389365 | 13786.456486 | 37488.900045 | 50779.115458 | 50836.012012 | 49712.398937 | 169323.438498 | 826651.427131 | 1.152271e+06 | 1.741844e+06 | 9.348315e+05 | 3.902234e+05 | 287944.085525 | 41868.553075 | 5740.356407 | 715.390217 | 71.917817 | 23.020692 | 13.042894 | 5.215163 | 3.503381 | 2.626363 | 1.962233 | 1.297150 | 0.729870 | 0.412411 | 0.312369 | 0.271270 | 0.290992 | 0.419615 | 0.765442 | 1.507684 | 2.742862 | 2.103041 | 2.230014 | 2.781006 | 1.944322 | 1.965533 | 2.231036 | 1.644876 | 1.914232 | 0.895227 | 0.297794 | 0.211477 | 0.227358 | 0.257646 | 0.272039 | 0.261087 | 0.234047 | 0.225426 | 0.211171 | 0.230944 | 0.331495 | 0.939492 | 3.191088 | 8.736713 | 26.419215 | 40.936587 | 72.337647 | 75.573319 | 35.339293 | 16.877021 | 9.079911 | 6.606708 | 6.422778 | 5.847725 | 4.308878 | 3.965385 | 3.996721 | 5.111539 | 4.796903 | 5.129510 | 6.927394 | 5.815139 | 6.287091 | 8.273783 | 9.671020 | 10.131552 | 12.617183 | 15.549454 | 32.343518 | 29.503746 | 18.336262 | 8.468869 | 4.324522 | 2.036306 | 1.112255 | 0.662717 | 0.428658 | 0.308160 | 0.273053 | 0.262053 | 0.268427 | 0.352158 | 0.726357 | 3.304353 | 15.369328 | 20.990139 | 31.010457 | 29.839106 | 13.709351 | 7.561779 | 4.088460 | 3.509946 | 4.417698 | 5.127557 | 9.812741 | 14.068593 | 14.350944 | 14.736062 | 9.396339 | 8.994958 | 7.342871 | 6.206294 | 5.178001 | 4.866287 | 5.231950 | 5.783209 | 5.925519 | 14.773378 | 19.232007 | 8.414304 | 3.600919 | 1.841813 | 1.594525 | 1.207863 | 0.780181 | 0.718931 | 0.995127 | 1.422862 | 1.987385 | 2.700122 | 5.560618 | 15.630682 | 26.119890 | 34.126591 | 44.916615 | 86.724070 | 67.841054 | 42.176621 | 32.271827 | 37.949299 | 27.084596 | 33.553883 | 28.761936 | 32.133128 | 29.119224 | 22.004228 | 13.234412 | 10.896476 | 11.388640 | 9.751341 | 7.999422 | 5.927853 | 4.813772 | 4.738648 | 5.950692 | 5.082915 | 4.842563 | 4.315075 | 2.979944 | 2.023026 | 1.809773 | 1.182103 | 0.886621 | 0.752508 | 0.771948 | 0.896891 | 1.178230 | 1.169703 | 1.049333 | 1.075507 | 1.022761 | 0.872673 | 0.745345 | 0.607946 | 0.507361 | 0.672211 | 0.991759 | 1.851309 | 3.606471 | 7.873985 | 16.481558 | 19.530360 | 19.682703 | 23.038270 | 17.651134 | 14.869965 | 14.001070 | 9.481074 | 7.595382 | 5.651571 | 4.545419 | 4.142006 | 5.356246 | 5.088229 | 6.399832 | 6.703143 | 6.298199 | 7.462241 | 12.653172 | 15.965743 | 16.457307 | 11.977452 | 10.018578 | 8.109879 | 6.502315 | 3.075669 | 1.781727 | 1.534535 | 1.323559 | 1.363626 | 1.811006 | 2.585033 | 4.350417 | 5.117303 | 4.734200 | 6.550208 | 7.843637 |
| all electrodes | 0.314738 | 0.272329 | 0.324261 | 0.477840 | 0.308626 | 0.273153 | 0.327576 | 0.397450 | 0.293328 | 0.255565 | 0.236842 | 0.213257 | 0.344547 | 2.362624 | 2.382153 | 0.887531 | 0.633417 | 0.781636 | 0.746223 | 0.336450 | 0.212522 | 0.211593 | 0.211111 | 0.212234 | 0.212151 | 0.215422 | 0.239331 | 0.256244 | 0.261600 | 0.269253 | 0.280513 | 0.226293 | 0.214777 | 0.221779 | 0.213844 | 0.223815 | 0.212079 | 0.213647 | 0.308991 | 0.259756 | 0.236867 | 0.281158 | 0.419279 | 2.037154 | 3.969885 | 2.314767 | 5.164023 | 2.839992 | 0.895364 | 0.374974 | 0.244367 | 0.210863 | 0.321314 | 0.806272 | 1.681085 | 1.854005 | 2.796832 | 11.337206 | 38.226756 | 42.290666 | 30.562620 | 45.895016 | 102.047369 | 555.418931 | 1361.592850 | 2998.778450 | 3533.786838 | 8317.382546 | 6824.371167 | 6.728311e+03 | 8.828764e+03 | 4.938235e+03 | 3.943655e+03 | 3243.335078 | 565.771706 | 91.920671 | 19.195121 | 5.727550 | 2.605552 | 1.377591 | 1.051118 | 1.433957 | 1.568951 | 1.205714 | 0.946565 | 0.516472 | 0.316065 | 0.216920 | 0.228354 | 0.267340 | 0.252093 | 0.276420 | 0.231457 | 0.213397 | 0.299244 | 0.720965 | 0.972936 | 0.734465 | 0.782267 | 0.522142 | 0.478210 | 0.475618 | 0.363443 | 0.365558 | 0.390244 | 0.408441 | 0.471832 | 0.301501 | 0.273129 | 0.251962 | 0.255485 | 0.249900 | 0.286924 | 0.349177 | 0.496327 | 0.460368 | 0.782600 | 1.124196 | 1.351622 | 0.901716 | 0.608379 | 0.589613 | 0.772666 | 0.640671 | 0.610010 | 0.439570 | 0.413182 | 0.505155 | 0.695634 | 0.791789 | 1.093899 | 1.245151 | 2.680214 | 3.928748 | 6.502237 | 7.514165 | 7.982723 | 8.145453 | 8.426914 | 5.905389 | 3.919241 | 2.535482 | 1.955162 | 1.535815 | 1.582047 | 1.384136 | 1.261551 | 1.927230 | 2.623420 | 3.114537 | 1.327179 | 0.724781 | 0.507138 | 0.411872 | 0.335310 | 0.328998 | 0.322551 | 0.374427 | 0.365199 | 0.290409 | 0.265207 | 0.252503 | 0.233618 | 0.219058 | 0.221070 | 0.226413 | 0.293176 | 0.352374 | 0.382584 | 0.367024 | 0.350953 | 0.329218 | 0.300965 | 0.262252 | 0.240302 | 0.242084 | 0.258659 | 0.311811 | 0.427548 | 0.476532 | 0.682901 | 1.528568 | 2.342846 | 3.847004 | 4.466252 | 6.306111 | 9.761854 | 7.867305 | 7.174042 | 8.249775 | 9.520231 | 16.329295 | 14.432064 | 16.621693 | 38.294763 | 54.616356 | 60.865769 | 48.420159 | 35.270016 | 58.637020 | 70.575312 | 51.907951 | 20.809126 | 13.178154 | 11.398296 | 17.104815 | 18.394877 | 22.863623 | 22.171618 | 31.432713 | 50.413929 | 84.275684 | 59.402738 | 45.642301 | 25.910465 | 14.795887 | 8.843709 | 4.336966 | 2.650475 | 1.973902 | 1.387919 | 0.978062 | 0.860445 | 0.852840 | 1.210263 | 1.642958 | 2.329614 | 2.756926 | 3.629979 | 7.062841 | 10.420697 | 8.461966 | 8.782380 | 5.434853 | 3.616009 | 2.804944 | 1.722567 | 1.975254 | 1.713712 | 1.253427 | 1.123563 | 1.292200 | 1.369806 | 1.526379 | 1.357408 | 1.427804 | 1.898651 | 2.615947 | 2.417938 | 1.736003 | 1.276111 | 1.261397 | 1.204347 | 1.100547 | 1.074740 | 1.096748 | 1.146023 | 1.163462 | 1.277590 | 1.160349 | 0.905098 | 0.681016 | 0.576702 | 0.731170 | 1.014162 | 1.203320 | 1.436990 | 1.348963 | 1.525778 | 1.278260 | 0.859792 | 0.532183 | 0.461664 | 0.493100 | 0.601519 | 0.625452 | 0.785135 | 0.785636 | 1.041283 | 1.245018 |

Searchlight, spatiotemporal cluster permutation test

|  | start time | stop time | peak time | peak channel | cluster p | peak Cohen's d | direction |
| --- | --- | --- | --- | --- | --- | --- | --- |
| #1 | 50 | 225 | 145 | P6 | 0.0031 | 1.90512 | positive |
| #2 | 350 | 1195 | 1020 | PO4 | 0.0006 | 1.140252 | positive |

B) emoji faces (LOSO) - happy vs angry

  
|  | time window | peak latency | cluster *p* | peak Cohen's *d* |  | | | |
| **all electrodes** | 765 - 875 ms | 855 ms | 0.045 | 0.7939 |  | | | |
|  | | | | | | | | |

Time-resolved classification, cluster permutation tests

|  | **left hemisphere** | | | | **right hemisphere** | | | |
|  | time window | peak latency | cluster *p* | peak Cohen's *d* | time window | peak latency | cluster *p* | peak Cohen's *d* |
| **anterior** |  | | | |  | | | |
| **central** |  | | | |  | | | |
| **posterior** | 595 - 800 ms | 665 ms | 0.0258 | 0.7921 | 595 - 730 ms | 670 ms | 0.0389 | 0.5342 |
  | | | | 895 - 1195 ms | 1010 ms | 0.0074 | 0.7912 |

  

Time-resolved classification, Bayesian statistics

|  | -200 | -195 | -190 | -185 | -180 | -175 | -170 | -165 | -160 | -155 | -150 | -145 | -140 | -135 | -130 | -125 | -120 | -115 | -110 | -105 | -100 | -95 | -90 | -85 | -80 | -75 | -70 | -65 | -60 | -55 | -50 | -45 | -40 | -35 | -30 | -25 | -20 | -15 | -10 | -5 | 0 | 5 | 10 | 15 | 20 | 25 | 30 | 35 | 40 | 45 | 50 | 55 | 60 | 65 | 70 | 75 | 80 | 85 | 90 | 95 | 100 | 105 | 110 | 115 | 120 | 125 | 130 | 135 | 140 | 145 | 150 | 155 | 160 | 165 | 170 | 175 | 180 | 185 | 190 | 195 | 200 | 205 | 210 | 215 | 220 | 225 | 230 | 235 | 240 | 245 | 250 | 255 | 260 | 265 | 270 | 275 | 280 | 285 | 290 | 295 | 300 | 305 | 310 | 315 | 320 | 325 | 330 | 335 | 340 | 345 | 350 | 355 | 360 | 365 | 370 | 375 | 380 | 385 | 390 | 395 | 400 | 405 | 410 | 415 | 420 | 425 | 430 | 435 | 440 | 445 | 450 | 455 | 460 | 465 | 470 | 475 | 480 | 485 | 490 | 495 | 500 | 505 | 510 | 515 | 520 | 525 | 530 | 535 | 540 | 545 | 550 | 555 | 560 | 565 | 570 | 575 | 580 | 585 | 590 | 595 | 600 | 605 | 610 | 615 | 620 | 625 | 630 | 635 | 640 | 645 | 650 | 655 | 660 | 665 | 670 | 675 | 680 | 685 | 690 | 695 | 700 | 705 | 710 | 715 | 720 | 725 | 730 | 735 | 740 | 745 | 750 | 755 | 760 | 765 | 770 | 775 | 780 | 785 | 790 | 795 | 800 | 805 | 810 | 815 | 820 | 825 | 830 | 835 | 840 | 845 | 850 | 855 | 860 | 865 | 870 | 875 | 880 | 885 | 890 | 895 | 900 | 905 | 910 | 915 | 920 | 925 | 930 | 935 | 940 | 945 | 950 | 955 | 960 | 965 | 970 | 975 | 980 | 985 | 990 | 995 | 1000 | 1005 | 1010 | 1015 | 1020 | 1025 | 1030 | 1035 | 1040 | 1045 | 1050 | 1055 | 1060 | 1065 | 1070 | 1075 | 1080 | 1085 | 1090 | 1095 | 1100 | 1105 | 1110 | 1115 | 1120 | 1125 | 1130 | 1135 | 1140 | 1145 | 1150 | 1155 | 1160 | 1165 | 1170 | 1175 | 1180 | 1185 | 1190 | 1195 |
| --- | --- | --- | --- | --- | --- | --- | --- | --- | --- | --- | --- | --- | --- | --- | --- | --- | --- | --- | --- | --- | --- | --- | --- | --- | --- | --- | --- | --- | --- | --- | --- | --- | --- | --- | --- | --- | --- | --- | --- | --- | --- | --- | --- | --- | --- | --- | --- | --- | --- | --- | --- | --- | --- | --- | --- | --- | --- | --- | --- | --- | --- | --- | --- | --- | --- | --- | --- | --- | --- | --- | --- | --- | --- | --- | --- | --- | --- | --- | --- | --- | --- | --- | --- | --- | --- | --- | --- | --- | --- | --- | --- | --- | --- | --- | --- | --- | --- | --- | --- | --- | --- | --- | --- | --- | --- | --- | --- | --- | --- | --- | --- | --- | --- | --- | --- | --- | --- | --- | --- | --- | --- | --- | --- | --- | --- | --- | --- | --- | --- | --- | --- | --- | --- | --- | --- | --- | --- | --- | --- | --- | --- | --- | --- | --- | --- | --- | --- | --- | --- | --- | --- | --- | --- | --- | --- | --- | --- | --- | --- | --- | --- | --- | --- | --- | --- | --- | --- | --- | --- | --- | --- | --- | --- | --- | --- | --- | --- | --- | --- | --- | --- | --- | --- | --- | --- | --- | --- | --- | --- | --- | --- | --- | --- | --- | --- | --- | --- | --- | --- | --- | --- | --- | --- | --- | --- | --- | --- | --- | --- | --- | --- | --- | --- | --- | --- | --- | --- | --- | --- | --- | --- | --- | --- | --- | --- | --- | --- | --- | --- | --- | --- | --- | --- | --- | --- | --- | --- | --- | --- | --- | --- | --- | --- | --- | --- | --- | --- | --- | --- | --- | --- | --- | --- | --- | --- | --- | --- | --- | --- | --- | --- | --- | --- | --- | --- | --- | --- | --- | --- | --- | --- | --- | --- | --- | --- | --- | --- | --- | --- | --- |
| left anterior | 3.008320 | 2.459525 | 1.553811 | 0.817516 | 0.426543 | 0.251291 | 0.210939 | 0.253040 | 0.398240 | 0.663610 | 0.949606 | 0.691193 | 0.595878 | 1.169396 | 1.489183 | 1.924730 | 4.729736 | 14.794718 | 30.806204 | 24.397052 | 67.527555 | 104.153590 | 21.016473 | 3.520486 | 0.584781 | 0.241575 | 0.236165 | 0.486035 | 1.298686 | 1.168986 | 0.320200 | 0.215972 | 0.210822 | 0.214249 | 0.239644 | 0.225999 | 0.216704 | 0.213715 | 0.214485 | 0.212361 | 0.215435 | 0.227948 | 0.218911 | 0.222166 | 0.250571 | 0.288526 | 0.308913 | 0.287984 | 0.290890 | 0.359517 | 0.350228 | 0.259630 | 0.237180 | 0.232706 | 0.268546 | 0.255348 | 0.238177 | 0.253915 | 0.379834 | 0.386985 | 0.318668 | 0.240296 | 0.243617 | 0.217017 | 0.217310 | 0.280087 | 0.240129 | 0.211695 | 0.233042 | 0.253758 | 0.290314 | 0.342504 | 0.448810 | 0.398022 | 0.311705 | 0.260669 | 0.289801 | 0.310021 | 0.267260 | 0.223333 | 0.213103 | 0.212396 | 0.212649 | 0.219231 | 0.238786 | 0.212900 | 0.210882 | 0.217066 | 0.275483 | 0.644771 | 0.724984 | 0.558416 | 0.709645 | 0.433759 | 0.285753 | 0.226168 | 0.214699 | 0.228220 | 0.248992 | 0.236065 | 0.268031 | 0.258241 | 0.265960 | 0.249988 | 0.230829 | 0.226790 | 0.258468 | 0.250261 | 0.284236 | 0.296417 | 0.353377 | 0.498574 | 0.706108 | 0.797016 | 0.871118 | 0.752550 | 0.398177 | 0.277858 | 0.227137 | 0.213197 | 0.216694 | 0.220030 | 0.218358 | 0.251499 | 0.269400 | 0.327173 | 0.481059 | 0.613854 | 0.663860 | 0.890848 | 0.851856 | 0.929975 | 0.915746 | 0.576983 | 0.317101 | 0.245953 | 0.212803 | 0.211322 | 0.223240 | 0.225761 | 0.241764 | 0.351975 | 0.448092 | 0.636268 | 0.746038 | 0.344375 | 0.264190 | 0.226530 | 0.210874 | 0.212333 | 0.220270 | 0.241142 | 0.219634 | 0.212973 | 0.230919 | 0.225453 | 0.228442 | 0.241104 | 0.283935 | 0.266449 | 0.211084 | 0.253907 | 0.301658 | 0.329491 | 0.365797 | 0.727146 | 1.211241 | 1.992635 | 1.491308 | 1.611212 | 1.679215 | 1.270204 | 0.894218 | 0.708595 | 0.417292 | 0.360104 | 0.296323 | 0.260104 | 0.264745 | 0.281168 | 0.291357 | 0.298257 | 0.279590 | 0.290510 | 0.299507 | 0.320916 | 0.289675 | 0.282279 | 0.287535 | 0.360339 | 0.422316 | 0.331038 | 0.244374 | 0.228045 | 0.223268 | 0.220852 | 0.214333 | 0.212570 | 0.235717 | 0.312518 | 0.377826 | 0.501841 | 0.581527 | 0.542806 | 0.421061 | 0.327119 | 0.278100 | 0.272655 | 0.244796 | 0.234344 | 0.227501 | 0.221113 | 0.217494 | 0.217959 | 0.214298 | 0.210978 | 0.225421 | 0.274331 | 0.302648 | 0.320560 | 0.329640 | 0.399229 | 0.373660 | 0.307737 | 0.228679 | 0.216440 | 0.215116 | 0.210977 | 0.213942 | 0.218032 | 0.210806 | 0.246354 | 0.301028 | 0.316113 | 0.384375 | 0.472133 | 0.567638 | 0.476600 | 0.398462 | 0.373517 | 0.349955 | 0.319930 | 0.274537 | 0.259089 | 0.271243 | 0.271120 | 0.286412 | 0.265060 | 0.261340 | 0.279892 | 0.269550 | 0.243275 | 0.226806 | 0.215273 | 0.222356 | 0.226095 | 0.219682 | 0.213788 | 0.213973 | 0.215443 | 0.214931 | 0.213093 | 0.214125 | 0.229160 | 0.277979 | 0.352867 | 0.425685 | 0.559404 | 0.792974 | 0.935557 | 0.727287 | 0.651617 | 0.490408 | 0.420999 | 0.333134 | 0.263558 | 0.232476 | 0.221041 | 0.211041 | 0.215092 |
| right anterior | 0.467790 | 0.576810 | 0.543746 | 0.428156 | 0.338378 | 0.287276 | 0.287537 | 0.222936 | 0.214178 | 0.210847 | 0.211249 | 0.210821 | 0.235680 | 0.233951 | 0.249139 | 0.240831 | 0.273283 | 0.312072 | 0.225505 | 0.227336 | 0.222268 | 0.210960 | 0.216740 | 0.249501 | 0.275020 | 0.576873 | 2.227484 | 3.732300 | 1.917935 | 0.730202 | 0.539826 | 0.417204 | 0.310707 | 0.246624 | 0.219957 | 0.231206 | 0.292014 | 0.371948 | 0.403880 | 0.360914 | 0.377186 | 0.576219 | 0.532978 | 0.306241 | 0.216733 | 0.211759 | 0.214810 | 0.228593 | 0.224531 | 0.222701 | 0.252450 | 0.222874 | 0.210997 | 0.265516 | 0.354929 | 0.496360 | 0.609517 | 0.993289 | 0.979567 | 1.404650 | 2.153649 | 6.217881 | 7.657959 | 12.344924 | 12.241153 | 15.764109 | 5.804845 | 0.580525 | 0.211254 | 0.317538 | 0.765834 | 1.489548 | 4.888831 | 3.462590 | 1.144167 | 0.364949 | 0.212345 | 0.250753 | 0.212281 | 0.217481 | 0.246455 | 0.246836 | 0.220465 | 0.216208 | 0.223338 | 0.210843 | 0.314523 | 0.814330 | 2.513869 | 2.236974 | 3.173802 | 3.832591 | 8.119002 | 7.660083 | 3.245315 | 2.131155 | 1.236675 | 1.482403 | 2.034781 | 0.954940 | 0.313335 | 0.304178 | 0.268783 | 0.290335 | 0.244002 | 0.217809 | 0.241268 | 0.266714 | 0.220269 | 0.210943 | 0.216487 | 0.222009 | 0.260385 | 0.360405 | 0.430596 | 0.582710 | 1.005943 | 1.700925 | 2.536459 | 1.730415 | 1.222196 | 0.999386 | 0.523342 | 0.316550 | 0.274818 | 0.299102 | 0.303958 | 0.256689 | 0.216566 | 0.212075 | 0.222390 | 0.277067 | 0.447221 | 0.709790 | 1.106134 | 0.882264 | 0.497150 | 0.405595 | 0.291392 | 0.245102 | 0.237071 | 0.215091 | 0.212681 | 0.220311 | 0.231485 | 0.271038 | 0.319298 | 0.308514 | 0.254992 | 0.223346 | 0.212629 | 0.210855 | 0.213500 | 0.218508 | 0.237054 | 0.244067 | 0.241123 | 0.239188 | 0.233674 | 0.237646 | 0.241484 | 0.220863 | 0.213752 | 0.217627 | 0.227218 | 0.245536 | 0.269119 | 0.335652 | 0.416354 | 0.427210 | 0.354224 | 0.310377 | 0.270571 | 0.250702 | 0.224035 | 0.219302 | 0.220923 | 0.223317 | 0.243524 | 0.283551 | 0.307164 | 0.311263 | 0.311612 | 0.314358 | 0.298949 | 0.273970 | 0.260371 | 0.253331 | 0.268602 | 0.280356 | 0.318752 | 0.405921 | 0.481202 | 0.492806 | 0.478608 | 0.473058 | 0.443765 | 0.383456 | 0.333526 | 0.324816 | 0.297820 | 0.307389 | 0.307177 | 0.326137 | 0.345395 | 0.390572 | 0.379644 | 0.463019 | 0.468136 | 0.543558 | 0.638575 | 0.641241 | 0.497261 | 0.425652 | 0.333329 | 0.286340 | 0.246091 | 0.233212 | 0.227450 | 0.245276 | 0.274400 | 0.278385 | 0.273187 | 0.280662 | 0.277773 | 0.297592 | 0.304228 | 0.289856 | 0.295221 | 0.305716 | 0.285676 | 0.254462 | 0.231842 | 0.223415 | 0.227799 | 0.227182 | 0.237119 | 0.247989 | 0.261932 | 0.291865 | 0.313426 | 0.303242 | 0.318759 | 0.301255 | 0.297972 | 0.311006 | 0.298750 | 0.276552 | 0.267990 | 0.256767 | 0.269690 | 0.303938 | 0.350666 | 0.397994 | 0.357197 | 0.311019 | 0.318581 | 0.314486 | 0.273056 | 0.248331 | 0.243480 | 0.289263 | 0.365623 | 0.378610 | 0.316079 | 0.315086 | 0.280417 | 0.232570 | 0.212575 | 0.211649 | 0.222000 | 0.216017 | 0.214334 | 0.210876 | 0.224367 | 0.231977 | 0.223683 | 0.239635 | 0.246575 | 0.243389 |
| left central | 0.216654 | 0.210806 | 0.225522 | 0.247136 | 0.216617 | 0.210835 | 0.210895 | 0.232381 | 0.281919 | 0.236263 | 0.250230 | 0.231626 | 0.212676 | 0.217737 | 0.289853 | 0.342289 | 0.444599 | 1.316304 | 0.761109 | 0.348722 | 0.263278 | 0.360724 | 0.432606 | 0.256300 | 0.214799 | 0.211220 | 0.218797 | 0.230475 | 0.248252 | 0.260317 | 0.211446 | 0.217583 | 0.240981 | 0.295905 | 0.259718 | 0.215731 | 0.211253 | 0.238329 | 0.308335 | 0.467173 | 0.427790 | 0.256627 | 0.213513 | 0.218073 | 0.223367 | 0.227881 | 0.254921 | 0.233432 | 0.215225 | 0.210820 | 0.210867 | 0.212929 | 0.221444 | 0.238654 | 0.267054 | 0.256036 | 0.230941 | 0.248065 | 0.239275 | 0.226594 | 0.229773 | 0.224227 | 0.256919 | 0.318018 | 0.317486 | 0.352075 | 0.668363 | 1.060915 | 1.133116 | 0.906343 | 0.711806 | 0.676835 | 0.838912 | 0.572626 | 0.341690 | 0.241032 | 0.213419 | 0.218723 | 0.219506 | 0.293995 | 0.337676 | 0.264606 | 0.246107 | 0.242569 | 0.225752 | 0.226913 | 0.219815 | 0.273788 | 0.329084 | 0.343669 | 0.277261 | 0.300148 | 0.327574 | 0.308741 | 0.210829 | 0.217637 | 0.229945 | 0.238748 | 0.210824 | 0.224789 | 0.219154 | 0.237633 | 0.296883 | 0.503553 | 0.653713 | 0.489923 | 0.368666 | 0.403439 | 0.477934 | 0.557991 | 0.396014 | 0.399975 | 0.340735 | 0.390034 | 0.412099 | 0.312244 | 0.233948 | 0.221189 | 0.216028 | 0.216302 | 0.210841 | 0.215896 | 0.211698 | 0.211566 | 0.212965 | 0.210892 | 0.210842 | 0.210954 | 0.212092 | 0.210915 | 0.217311 | 0.248464 | 0.296664 | 0.334786 | 0.394833 | 0.611059 | 0.832356 | 0.575839 | 0.501669 | 0.483679 | 0.486253 | 0.544079 | 0.491495 | 0.424988 | 0.451593 | 0.407086 | 0.378616 | 0.379728 | 0.370531 | 0.462430 | 0.526470 | 0.544335 | 0.602296 | 0.681746 | 0.766916 | 0.752691 | 0.640211 | 0.584845 | 0.516029 | 0.589086 | 0.682801 | 0.776752 | 0.785459 | 0.808509 | 1.101161 | 1.548430 | 1.205265 | 1.003216 | 1.141935 | 2.420365 | 2.841090 | 2.836614 | 2.891691 | 3.471191 | 7.100251 | 5.028019 | 3.011069 | 2.749217 | 1.903788 | 1.173622 | 0.740031 | 0.433353 | 0.374010 | 0.323406 | 0.322202 | 0.297220 | 0.393066 | 0.545629 | 0.787956 | 1.431077 | 4.698674 | 7.174682 | 20.680590 | 32.854859 | 37.889878 | 18.084965 | 8.982847 | 3.495788 | 2.014757 | 1.301584 | 0.977072 | 1.096624 | 1.777361 | 1.878448 | 1.949422 | 2.175454 | 2.509912 | 2.116241 | 1.166602 | 0.591449 | 0.456521 | 0.399119 | 0.367479 | 0.318720 | 0.283939 | 0.334237 | 0.420197 | 0.505292 | 0.603747 | 0.803085 | 0.956796 | 1.191158 | 0.837586 | 0.514628 | 0.331342 | 0.270790 | 0.285992 | 0.424301 | 0.818284 | 0.994891 | 1.488487 | 4.800392 | 8.014840 | 5.456436 | 3.917734 | 1.826277 | 2.150807 | 2.909378 | 2.718412 | 3.743727 | 4.529289 | 3.246341 | 2.679674 | 1.758469 | 1.307871 | 0.966158 | 0.604134 | 0.409512 | 0.362832 | 0.360906 | 0.317531 | 0.251237 | 0.214992 | 0.211024 | 0.211768 | 0.210850 | 0.211086 | 0.211214 | 0.227384 | 0.298671 | 0.534705 | 0.848640 | 1.891854 | 3.700444 | 5.492373 | 6.367296 | 6.741213 | 4.126072 | 2.936186 | 2.617833 | 1.462142 | 1.234198 | 1.073814 | 0.838644 | 0.758731 | 0.961889 | 0.856397 | 0.869437 | 0.747911 | 0.611879 |
| right central | 11.321327 | 3.258336 | 2.601126 | 1.737263 | 0.952172 | 0.577930 | 0.317999 | 0.232817 | 0.211583 | 0.215208 | 0.210883 | 0.213238 | 0.269597 | 0.337658 | 0.333073 | 0.360986 | 0.340866 | 0.367070 | 0.311683 | 0.268966 | 0.237409 | 0.228776 | 0.224796 | 0.266902 | 0.292953 | 0.275120 | 0.290199 | 0.265794 | 0.211277 | 0.311344 | 0.648659 | 1.306477 | 1.732878 | 2.010038 | 1.792501 | 0.775814 | 0.440901 | 0.416859 | 0.561145 | 0.312961 | 0.219811 | 0.217395 | 0.253720 | 0.273463 | 0.253038 | 0.228383 | 0.223746 | 0.412509 | 0.488747 | 0.595239 | 1.163726 | 2.848164 | 1.883660 | 1.363307 | 0.552961 | 0.812439 | 4.370107 | 9.655771 | 7.182213 | 14.534091 | 25.779177 | 74.405636 | 46.301268 | 3.142745 | 0.771557 | 0.311692 | 0.210822 | 0.282414 | 0.629575 | 0.976629 | 1.493111 | 2.200963 | 3.255755 | 2.771629 | 1.295421 | 0.585909 | 0.327536 | 0.233052 | 0.218859 | 0.219372 | 0.217454 | 0.216882 | 0.210872 | 0.231151 | 0.226552 | 0.230402 | 0.230083 | 0.253666 | 0.303022 | 0.431677 | 0.495812 | 0.752991 | 1.212334 | 1.232937 | 1.024297 | 0.612047 | 0.414812 | 0.374538 | 0.423999 | 0.516156 | 0.843985 | 0.936554 | 2.545443 | 3.119018 | 2.259421 | 0.685685 | 0.256291 | 0.217587 | 0.220436 | 0.222406 | 0.236008 | 0.234488 | 0.249523 | 0.264994 | 0.232497 | 0.211234 | 0.237660 | 0.318866 | 0.481474 | 0.655342 | 0.585446 | 0.386611 | 0.280275 | 0.224795 | 0.210976 | 0.223496 | 0.255333 | 0.265443 | 0.271344 | 0.275932 | 0.261453 | 0.268148 | 0.281376 | 0.357828 | 0.414854 | 0.384635 | 0.407837 | 0.324766 | 0.258850 | 0.218806 | 0.211414 | 0.239823 | 0.319486 | 0.383987 | 0.504149 | 0.559130 | 0.865130 | 1.323855 | 1.060495 | 0.974622 | 0.544406 | 0.542812 | 0.830224 | 1.432623 | 1.884534 | 2.101121 | 1.772278 | 1.782716 | 2.098150 | 2.095036 | 1.374892 | 1.068938 | 0.953869 | 0.620566 | 0.698858 | 0.398749 | 0.306222 | 0.285758 | 0.273618 | 0.313096 | 0.448177 | 0.476565 | 0.547169 | 0.691124 | 0.763166 | 0.767742 | 0.560444 | 0.427061 | 0.385626 | 0.564902 | 0.375723 | 0.251914 | 0.211428 | 0.212066 | 0.213978 | 0.218445 | 0.226854 | 0.211476 | 0.224814 | 0.249229 | 0.236198 | 0.220477 | 0.213117 | 0.211602 | 0.253108 | 0.284507 | 0.308957 | 0.277235 | 0.239155 | 0.228307 | 0.238371 | 0.244316 | 0.251639 | 0.260482 | 0.285874 | 0.379736 | 0.637782 | 0.966481 | 1.340715 | 1.509366 | 1.602126 | 1.470213 | 1.247497 | 0.729339 | 0.491812 | 0.317015 | 0.262776 | 0.223583 | 0.217950 | 0.217552 | 0.232836 | 0.245063 | 0.274797 | 0.291144 | 0.350682 | 0.489841 | 0.625434 | 0.687441 | 0.673009 | 0.440114 | 0.309975 | 0.263090 | 0.237607 | 0.220126 | 0.214420 | 0.230846 | 0.211520 | 0.261494 | 0.358006 | 0.334757 | 0.312769 | 0.357338 | 0.367866 | 0.281058 | 0.240859 | 0.216153 | 0.219530 | 0.239205 | 0.289083 | 0.335461 | 0.324729 | 0.281037 | 0.263896 | 0.231458 | 0.232668 | 0.224360 | 0.224017 | 0.222354 | 0.220801 | 0.222110 | 0.219541 | 0.215021 | 0.263661 | 0.437940 | 0.518646 | 0.469742 | 0.449757 | 0.420039 | 0.353610 | 0.290195 | 0.264493 | 0.263994 | 0.252276 | 0.224237 | 0.216479 | 0.211296 | 0.210956 | 0.212283 | 0.223179 | 0.225747 |
| left posterior | 0.291304 | 0.399776 | 0.315685 | 0.375746 | 0.400700 | 0.755213 | 0.932680 | 0.709680 | 0.576264 | 0.733386 | 0.437385 | 0.353036 | 0.252089 | 0.211013 | 0.214926 | 0.235155 | 0.247019 | 0.239562 | 0.246063 | 0.227107 | 0.217194 | 0.218262 | 0.232414 | 0.256067 | 0.224787 | 0.210940 | 0.210822 | 0.230630 | 0.332743 | 0.901325 | 2.993894 | 2.625943 | 0.418172 | 0.214902 | 0.236817 | 0.357910 | 0.829409 | 1.468492 | 1.050183 | 0.750796 | 0.538394 | 0.496564 | 0.680632 | 0.969992 | 0.839283 | 0.934485 | 0.530686 | 0.287222 | 0.224875 | 0.211090 | 0.217404 | 0.237119 | 0.321516 | 0.382323 | 0.310803 | 0.254330 | 0.226186 | 0.327086 | 0.489138 | 0.516497 | 0.513352 | 0.449962 | 0.397030 | 0.320079 | 0.277051 | 0.262029 | 0.346634 | 0.653777 | 1.956570 | 16.067475 | 193.219023 | 1492.553746 | 5956.024787 | 4279.755780 | 993.336383 | 353.547560 | 50.476058 | 9.339977 | 2.796776 | 1.308702 | 0.680290 | 0.522960 | 0.387192 | 0.303150 | 0.227066 | 0.217241 | 0.402788 | 1.097512 | 5.606449 | 7.814167 | 5.193097 | 3.573534 | 2.857464 | 1.188751 | 0.533731 | 0.328742 | 0.278198 | 0.281771 | 0.280772 | 0.258238 | 0.228829 | 0.221154 | 0.219496 | 0.211651 | 0.220966 | 0.242628 | 0.236943 | 0.213193 | 0.211486 | 0.215634 | 0.211157 | 0.213985 | 0.235224 | 0.322425 | 0.578287 | 1.148250 | 1.960544 | 1.901761 | 2.097445 | 2.974754 | 3.673911 | 4.788607 | 6.175138 | 9.035757 | 12.809547 | 16.117391 | 17.409149 | 12.824901 | 9.660529 | 4.922481 | 2.326658 | 1.451013 | 1.087101 | 0.918674 | 0.999368 | 1.097440 | 1.367107 | 2.103911 | 2.726723 | 2.982553 | 3.301176 | 3.350067 | 2.709746 | 3.001162 | 3.345944 | 3.955856 | 4.758297 | 4.521425 | 3.501004 | 2.653741 | 1.359997 | 0.700097 | 0.502717 | 0.409805 | 0.386784 | 0.423230 | 0.515945 | 0.764303 | 1.104816 | 1.450013 | 1.652290 | 1.917123 | 2.070601 | 1.874381 | 1.603078 | 1.345088 | 1.371764 | 2.082342 | 4.000262 | 7.132445 | 10.828947 | 12.293658 | 26.137658 | 55.917373 | 52.618422 | 33.224760 | 21.402093 | 9.569199 | 7.555466 | 5.834046 | 4.835080 | 6.307814 | 6.428304 | 5.739144 | 6.913811 | 6.161643 | 5.064849 | 3.796419 | 2.646953 | 2.637327 | 3.017977 | 3.715165 | 4.459106 | 5.270828 | 7.228427 | 7.195521 | 6.042287 | 4.139581 | 2.836335 | 2.326100 | 1.541848 | 1.040377 | 0.802563 | 0.693257 | 0.619818 | 0.497176 | 0.448396 | 0.436233 | 0.421020 | 0.453149 | 0.575606 | 0.811357 | 1.374810 | 1.698325 | 1.608563 | 1.428854 | 1.160274 | 0.931955 | 0.746586 | 0.669706 | 0.630974 | 0.763002 | 1.249655 | 2.275145 | 2.584558 | 2.825847 | 2.030823 | 1.763347 | 1.764355 | 1.462333 | 1.465297 | 1.520894 | 1.335208 | 1.464786 | 1.670014 | 1.388026 | 1.100252 | 0.736484 | 0.586048 | 0.539003 | 0.498526 | 0.455675 | 0.474840 | 0.505650 | 0.533690 | 0.514739 | 0.505083 | 0.482329 | 0.516287 | 0.563678 | 0.632602 | 0.787547 | 1.219869 | 1.605279 | 1.786390 | 1.700352 | 1.375636 | 1.172836 | 1.034255 | 0.864207 | 0.689863 | 0.589690 | 0.466068 | 0.440613 | 0.428328 | 0.429412 | 0.421114 | 0.437005 | 0.545195 | 0.576642 | 0.643687 | 0.663120 | 0.733284 | 0.920106 | 1.244133 | 1.665231 | 3.437648 | 5.227896 | 8.231411 | 11.059038 |
| right posterior | 0.836428 | 0.601592 | 0.562439 | 0.515961 | 0.281182 | 0.247017 | 0.220728 | 0.212642 | 0.210888 | 0.212215 | 0.227328 | 0.234957 | 0.232207 | 0.229822 | 0.265642 | 0.267700 | 0.274748 | 0.226878 | 0.223946 | 0.294449 | 0.306812 | 0.245359 | 0.300020 | 0.314820 | 0.342801 | 0.240846 | 0.214828 | 0.210806 | 0.222373 | 0.270172 | 0.252637 | 0.215956 | 0.220177 | 0.301618 | 0.582836 | 2.168445 | 13.305198 | 23.661548 | 24.041376 | 8.663329 | 1.931432 | 0.876551 | 0.668181 | 0.574201 | 0.584858 | 0.425883 | 0.554222 | 0.868912 | 0.952162 | 1.130579 | 1.063447 | 0.831820 | 0.907509 | 0.601339 | 0.464571 | 0.392546 | 0.251901 | 0.211832 | 0.267305 | 0.378342 | 0.485508 | 0.657809 | 0.945880 | 0.753263 | 0.768832 | 1.237560 | 1.845635 | 3.461150 | 11.906136 | 31.189496 | 105.625424 | 287.837036 | 410.088184 | 554.303155 | 329.679015 | 41.486278 | 5.253198 | 1.047434 | 0.377711 | 0.250784 | 0.217526 | 0.211913 | 0.218177 | 0.230279 | 0.275504 | 0.352942 | 0.449785 | 0.813414 | 1.030013 | 0.864542 | 0.726310 | 0.408396 | 0.282514 | 0.253246 | 0.219997 | 0.213685 | 0.215146 | 0.216765 | 0.233699 | 0.247265 | 0.265062 | 0.294975 | 0.288050 | 0.276043 | 0.263675 | 0.244480 | 0.230024 | 0.216762 | 0.210917 | 0.215173 | 0.230641 | 0.280735 | 0.438271 | 0.846186 | 1.633642 | 2.813630 | 3.834060 | 3.723125 | 3.309479 | 2.410636 | 1.593832 | 1.059316 | 0.714968 | 0.584279 | 0.644991 | 0.620414 | 0.528901 | 0.512083 | 0.545620 | 0.672706 | 0.670393 | 0.625082 | 0.608292 | 0.673951 | 0.660481 | 0.719974 | 0.691581 | 0.709808 | 0.905176 | 1.259491 | 1.348976 | 1.819647 | 1.610382 | 1.336711 | 1.387268 | 1.065189 | 0.797374 | 0.621968 | 0.517936 | 0.567805 | 0.585805 | 0.422643 | 0.374394 | 0.336063 | 0.392371 | 0.458839 | 0.510332 | 0.593088 | 0.925443 | 1.725929 | 2.853626 | 4.442105 | 4.326573 | 4.772394 | 4.732558 | 4.604693 | 3.325292 | 3.410258 | 3.354947 | 3.882882 | 3.448925 | 3.765482 | 3.540442 | 3.613296 | 3.747438 | 3.269410 | 3.566550 | 3.704398 | 3.353992 | 3.041698 | 3.532336 | 3.438573 | 3.072654 | 2.092646 | 1.750044 | 1.698522 | 1.334961 | 0.943186 | 0.820363 | 0.801511 | 0.841392 | 0.871903 | 1.023834 | 1.975779 | 3.406422 | 4.278093 | 4.975373 | 5.599057 | 4.907860 | 4.965813 | 3.844665 | 3.332437 | 2.764896 | 2.535733 | 3.327810 | 4.163370 | 2.984068 | 2.659933 | 2.888388 | 4.982832 | 5.751874 | 3.142203 | 1.757436 | 1.689140 | 1.504257 | 1.248974 | 0.844853 | 0.676542 | 0.798603 | 1.328656 | 2.570358 | 5.361874 | 8.705771 | 9.898667 | 8.630738 | 6.711762 | 5.842298 | 3.501885 | 2.848580 | 2.468024 | 2.392377 | 2.526069 | 2.877496 | 2.846591 | 3.500844 | 4.188881 | 4.509653 | 4.804718 | 8.132774 | 12.462802 | 23.702204 | 43.438225 | 55.319774 | 71.181895 | 98.828030 | 120.273123 | 196.289036 | 144.720496 | 109.860052 | 62.530440 | 37.821299 | 23.253320 | 14.232788 | 6.538648 | 3.735576 | 3.290073 | 2.971468 | 2.552898 | 3.285583 | 5.020931 | 8.212405 | 16.817128 | 19.722499 | 24.858449 | 21.785385 | 16.588020 | 13.760971 | 9.422690 | 6.384286 | 5.592575 | 4.600160 | 4.296666 | 4.618402 | 4.232937 | 5.886569 | 8.789591 | 13.714793 | 20.434310 | 34.794793 | 36.180417 |
| all electrodes | 0.235859 | 0.242350 | 0.282359 | 0.302175 | 0.210822 | 0.219242 | 0.210919 | 0.211770 | 0.250119 | 0.394535 | 0.527907 | 0.748216 | 1.665373 | 6.493078 | 5.187678 | 2.168836 | 2.181825 | 2.549689 | 1.273575 | 0.776496 | 0.442301 | 0.621040 | 0.428025 | 0.237811 | 0.216692 | 0.216304 | 0.235100 | 0.282683 | 0.355424 | 0.282369 | 0.212363 | 0.213572 | 0.248035 | 0.325304 | 0.480412 | 0.381925 | 0.305313 | 0.282853 | 0.378026 | 0.451779 | 0.489641 | 0.461365 | 0.848525 | 3.738970 | 24.691984 | 36.428079 | 56.046911 | 9.212494 | 1.528970 | 0.543106 | 0.327019 | 0.231129 | 0.211350 | 0.228131 | 0.224543 | 0.210860 | 0.211866 | 0.217193 | 0.232390 | 0.246379 | 0.249213 | 0.274037 | 0.396534 | 0.655936 | 1.137224 | 1.904255 | 4.219314 | 15.536977 | 83.160898 | 317.209789 | 653.096934 | 1423.202016 | 2334.529673 | 1674.294937 | 242.087479 | 14.740881 | 2.279329 | 0.667502 | 0.412949 | 0.321946 | 0.255256 | 0.232470 | 0.227764 | 0.225760 | 0.228707 | 0.211585 | 0.240284 | 0.408216 | 0.679249 | 0.847057 | 0.683747 | 0.862024 | 0.949881 | 0.420757 | 0.226771 | 0.220004 | 0.251575 | 0.217671 | 0.211741 | 0.212674 | 0.213975 | 0.217508 | 0.229886 | 0.291644 | 0.601842 | 1.130982 | 1.356931 | 1.126709 | 0.668914 | 0.439581 | 0.398331 | 0.397644 | 0.453949 | 0.496655 | 0.688923 | 1.402844 | 2.657079 | 2.892179 | 1.812822 | 1.325999 | 1.124689 | 1.216603 | 1.294660 | 1.006003 | 0.685535 | 0.575526 | 0.505510 | 0.404533 | 0.375784 | 0.335755 | 0.312981 | 0.310212 | 0.319193 | 0.302503 | 0.337135 | 0.325891 | 0.408284 | 0.563258 | 0.630878 | 0.709131 | 1.137105 | 1.198053 | 1.599231 | 1.366823 | 1.185918 | 1.273221 | 1.271646 | 0.944673 | 1.025454 | 0.972652 | 0.839438 | 0.631073 | 0.561494 | 0.450282 | 0.461549 | 0.452732 | 0.426856 | 0.447261 | 0.438398 | 0.499917 | 0.575605 | 0.730440 | 0.785926 | 0.944571 | 0.821658 | 0.710468 | 0.558926 | 0.742112 | 1.202596 | 1.607200 | 1.626985 | 1.688289 | 1.549633 | 1.699777 | 1.285996 | 0.705335 | 0.473629 | 0.349670 | 0.292518 | 0.297175 | 0.331451 | 0.405208 | 0.485374 | 0.504858 | 0.673182 | 0.853808 | 0.874529 | 0.609272 | 0.483268 | 0.545125 | 0.685282 | 0.788053 | 1.101542 | 1.803341 | 2.435757 | 1.955648 | 1.432045 | 1.462536 | 1.286213 | 1.339706 | 1.345928 | 1.485633 | 2.260225 | 1.931663 | 1.695797 | 1.871488 | 2.529353 | 4.289579 | 7.154971 | 9.937098 | 41.733741 | 57.034004 | 48.080491 | 21.572939 | 7.327839 | 2.392180 | 0.842246 | 0.359310 | 0.278999 | 0.281810 | 0.300098 | 0.285361 | 0.319572 | 0.366863 | 0.464686 | 0.602612 | 0.768890 | 0.886159 | 0.881710 | 0.679343 | 0.685481 | 0.533368 | 0.499795 | 0.381911 | 0.315007 | 0.356538 | 0.529278 | 0.548303 | 0.685283 | 0.646668 | 0.766004 | 1.038636 | 1.183337 | 0.976940 | 0.822474 | 0.751192 | 0.798654 | 0.709292 | 0.494932 | 0.291602 | 0.242432 | 0.242702 | 0.247637 | 0.244869 | 0.281783 | 0.322416 | 0.434755 | 0.448154 | 0.475844 | 0.438543 | 0.471730 | 0.384577 | 0.428131 | 0.472984 | 0.572653 | 0.601915 | 0.581917 | 0.472666 | 0.405146 | 0.294068 | 0.266764 | 0.249813 | 0.255423 | 0.271963 | 0.292005 | 0.393582 | 0.584652 | 0.622345 | 0.861678 | 0.967622 |

Searchlight, spatiotemporal cluster permutation test

|  | start time | stop time | peak time | peak channel | cluster p | peak Cohen's d | direction |
| --- | --- | --- | --- | --- | --- | --- | --- |
| #1 | 365 | 1195 | 665 | P9 | 0.004 | 0.790684 | positive |

C) emoji faces (LOSO) - happy vs sad

  
|  | time window | peak latency | cluster *p* | peak Cohen's *d* |  | | | |  |
| **all electrodes** | 150 - 235 ms | 190 ms | 0.0401 | 0.8928 |  | | | |  |
 715 - 1075 ms | 1015 ms | 0.0043 | 0.8234 |  | | | ||  |  | | | | | | | | |

Time-resolved classification, cluster permutation tests

|  | **left hemisphere** | | | | **right hemisphere** | | | |  |
|  | time window | peak latency | cluster *p* | peak Cohen's *d* | time window | peak latency | cluster *p* | peak Cohen's *d* |  |
| **anterior** |  | | | |  | | | |  |
| **central** |  | | | | 505 - 620 ms | 620 ms | 0.0151 | -0.4652 |  |
| **posterior** |  | | | | 80 - 220 ms | 180 ms | 0.0183 | 0.9119 |  |

  

Time-resolved classification, Bayesian statistics

|  | -200 | -195 | -190 | -185 | -180 | -175 | -170 | -165 | -160 | -155 | -150 | -145 | -140 | -135 | -130 | -125 | -120 | -115 | -110 | -105 | -100 | -95 | -90 | -85 | -80 | -75 | -70 | -65 | -60 | -55 | -50 | -45 | -40 | -35 | -30 | -25 | -20 | -15 | -10 | -5 | 0 | 5 | 10 | 15 | 20 | 25 | 30 | 35 | 40 | 45 | 50 | 55 | 60 | 65 | 70 | 75 | 80 | 85 | 90 | 95 | 100 | 105 | 110 | 115 | 120 | 125 | 130 | 135 | 140 | 145 | 150 | 155 | 160 | 165 | 170 | 175 | 180 | 185 | 190 | 195 | 200 | 205 | 210 | 215 | 220 | 225 | 230 | 235 | 240 | 245 | 250 | 255 | 260 | 265 | 270 | 275 | 280 | 285 | 290 | 295 | 300 | 305 | 310 | 315 | 320 | 325 | 330 | 335 | 340 | 345 | 350 | 355 | 360 | 365 | 370 | 375 | 380 | 385 | 390 | 395 | 400 | 405 | 410 | 415 | 420 | 425 | 430 | 435 | 440 | 445 | 450 | 455 | 460 | 465 | 470 | 475 | 480 | 485 | 490 | 495 | 500 | 505 | 510 | 515 | 520 | 525 | 530 | 535 | 540 | 545 | 550 | 555 | 560 | 565 | 570 | 575 | 580 | 585 | 590 | 595 | 600 | 605 | 610 | 615 | 620 | 625 | 630 | 635 | 640 | 645 | 650 | 655 | 660 | 665 | 670 | 675 | 680 | 685 | 690 | 695 | 700 | 705 | 710 | 715 | 720 | 725 | 730 | 735 | 740 | 745 | 750 | 755 | 760 | 765 | 770 | 775 | 780 | 785 | 790 | 795 | 800 | 805 | 810 | 815 | 820 | 825 | 830 | 835 | 840 | 845 | 850 | 855 | 860 | 865 | 870 | 875 | 880 | 885 | 890 | 895 | 900 | 905 | 910 | 915 | 920 | 925 | 930 | 935 | 940 | 945 | 950 | 955 | 960 | 965 | 970 | 975 | 980 | 985 | 990 | 995 | 1000 | 1005 | 1010 | 1015 | 1020 | 1025 | 1030 | 1035 | 1040 | 1045 | 1050 | 1055 | 1060 | 1065 | 1070 | 1075 | 1080 | 1085 | 1090 | 1095 | 1100 | 1105 | 1110 | 1115 | 1120 | 1125 | 1130 | 1135 | 1140 | 1145 | 1150 | 1155 | 1160 | 1165 | 1170 | 1175 | 1180 | 1185 | 1190 | 1195 |
| --- | --- | --- | --- | --- | --- | --- | --- | --- | --- | --- | --- | --- | --- | --- | --- | --- | --- | --- | --- | --- | --- | --- | --- | --- | --- | --- | --- | --- | --- | --- | --- | --- | --- | --- | --- | --- | --- | --- | --- | --- | --- | --- | --- | --- | --- | --- | --- | --- | --- | --- | --- | --- | --- | --- | --- | --- | --- | --- | --- | --- | --- | --- | --- | --- | --- | --- | --- | --- | --- | --- | --- | --- | --- | --- | --- | --- | --- | --- | --- | --- | --- | --- | --- | --- | --- | --- | --- | --- | --- | --- | --- | --- | --- | --- | --- | --- | --- | --- | --- | --- | --- | --- | --- | --- | --- | --- | --- | --- | --- | --- | --- | --- | --- | --- | --- | --- | --- | --- | --- | --- | --- | --- | --- | --- | --- | --- | --- | --- | --- | --- | --- | --- | --- | --- | --- | --- | --- | --- | --- | --- | --- | --- | --- | --- | --- | --- | --- | --- | --- | --- | --- | --- | --- | --- | --- | --- | --- | --- | --- | --- | --- | --- | --- | --- | --- | --- | --- | --- | --- | --- | --- | --- | --- | --- | --- | --- | --- | --- | --- | --- | --- | --- | --- | --- | --- | --- | --- | --- | --- | --- | --- | --- | --- | --- | --- | --- | --- | --- | --- | --- | --- | --- | --- | --- | --- | --- | --- | --- | --- | --- | --- | --- | --- | --- | --- | --- | --- | --- | --- | --- | --- | --- | --- | --- | --- | --- | --- | --- | --- | --- | --- | --- | --- | --- | --- | --- | --- | --- | --- | --- | --- | --- | --- | --- | --- | --- | --- | --- | --- | --- | --- | --- | --- | --- | --- | --- | --- | --- | --- | --- | --- | --- | --- | --- | --- | --- | --- | --- | --- | --- | --- | --- | --- | --- | --- | --- | --- | --- | --- | --- |
| left anterior | 0.220328 | 0.214722 | 0.257945 | 0.235646 | 0.263602 | 0.242177 | 0.236910 | 0.222185 | 0.215001 | 0.215288 | 0.293983 | 0.881901 | 1.895042 | 1.646726 | 4.414797 | 6.464583 | 3.746408 | 3.716619 | 2.670565 | 1.483837 | 0.953271 | 0.349137 | 0.225115 | 0.237299 | 0.223763 | 0.223281 | 0.223136 | 0.215200 | 0.210956 | 0.219227 | 0.340234 | 0.560868 | 0.597812 | 0.510948 | 0.401985 | 0.259407 | 0.210873 | 0.236796 | 0.235002 | 0.239156 | 0.267339 | 0.271815 | 0.247678 | 0.237084 | 0.219488 | 0.217143 | 0.212996 | 0.293082 | 0.280691 | 0.263151 | 0.246322 | 0.211401 | 0.215644 | 0.232651 | 0.304112 | 0.409995 | 0.445353 | 0.382106 | 0.230559 | 0.220630 | 0.217161 | 0.210961 | 0.215096 | 0.256044 | 0.222938 | 0.211930 | 0.216511 | 0.210822 | 0.217726 | 0.219374 | 0.228841 | 0.216251 | 0.214908 | 0.217600 | 0.311900 | 0.411741 | 0.831455 | 1.385782 | 2.060803 | 2.866766 | 1.807379 | 1.563859 | 1.699035 | 1.249974 | 1.097448 | 0.760786 | 0.756925 | 0.870501 | 1.035098 | 1.078045 | 1.353610 | 1.997747 | 3.783175 | 8.910871 | 18.790109 | 21.588085 | 13.127902 | 4.927029 | 2.163290 | 1.312365 | 0.882695 | 0.571004 | 0.535850 | 0.433362 | 0.387658 | 0.383477 | 0.427183 | 0.395672 | 0.352569 | 0.268804 | 0.231999 | 0.218872 | 0.211022 | 0.214783 | 0.230755 | 0.287895 | 0.292233 | 0.258230 | 0.218293 | 0.211578 | 0.220239 | 0.225121 | 0.255322 | 0.264429 | 0.232076 | 0.212721 | 0.210861 | 0.214268 | 0.254319 | 0.259141 | 0.233818 | 0.261701 | 0.255440 | 0.222998 | 0.213548 | 0.269983 | 0.303850 | 0.327958 | 0.427239 | 0.728415 | 1.140944 | 0.831394 | 0.636103 | 0.683887 | 0.532366 | 0.424484 | 0.257615 | 0.211936 | 0.211236 | 0.211932 | 0.215536 | 0.223194 | 0.223553 | 0.228159 | 0.215991 | 0.210815 | 0.210958 | 0.211072 | 0.214017 | 0.220097 | 0.248642 | 0.234656 | 0.223319 | 0.226755 | 0.229632 | 0.228763 | 0.216662 | 0.213574 | 0.211765 | 0.213836 | 0.215457 | 0.232137 | 0.311542 | 0.501675 | 0.466974 | 0.476676 | 0.548768 | 0.678813 | 0.745121 | 0.645691 | 0.465861 | 0.451263 | 0.526946 | 0.536781 | 0.550908 | 0.463920 | 0.398210 | 0.351262 | 0.330529 | 0.320933 | 0.341585 | 0.376045 | 0.374968 | 0.361943 | 0.366946 | 0.398372 | 0.444867 | 0.419205 | 0.335152 | 0.288091 | 0.300797 | 0.312873 | 0.261606 | 0.233671 | 0.216990 | 0.233370 | 0.253035 | 0.233025 | 0.214897 | 0.213601 | 0.219830 | 0.244449 | 0.272105 | 0.260309 | 0.283369 | 0.308130 | 0.272414 | 0.239288 | 0.216371 | 0.211574 | 0.213221 | 0.211069 | 0.210814 | 0.217111 | 0.232750 | 0.294202 | 0.248516 | 0.216516 | 0.211164 | 0.219602 | 0.219971 | 0.225748 | 0.214095 | 0.217207 | 0.232094 | 0.229402 | 0.221056 | 0.219977 | 0.213956 | 0.215278 | 0.220126 | 0.224674 | 0.257519 | 0.254206 | 0.246338 | 0.227549 | 0.235785 | 0.232499 | 0.224569 | 0.213029 | 0.214504 | 0.210953 | 0.213312 | 0.221818 | 0.213412 | 0.211194 | 0.216711 | 0.219027 | 0.239603 | 0.278762 | 0.365877 | 0.356435 | 0.294720 | 0.298669 | 0.270027 | 0.238648 | 0.247923 | 0.236569 | 0.213673 | 0.212948 | 0.212779 | 0.221036 | 0.245778 | 0.260024 | 0.257257 | 0.297402 | 0.252974 | 0.250756 | 0.246744 | 0.236917 |
| right anterior | 0.212395 | 0.216893 | 0.216314 | 0.211787 | 0.215018 | 0.216241 | 0.211945 | 0.234536 | 0.303058 | 0.520813 | 1.353960 | 1.521637 | 1.112656 | 0.723215 | 0.931518 | 0.369271 | 0.212622 | 0.221939 | 0.211233 | 0.210836 | 0.215480 | 0.210833 | 0.210833 | 0.225566 | 0.245308 | 0.210831 | 0.221127 | 0.230053 | 0.284648 | 0.377073 | 0.589388 | 0.725330 | 0.336289 | 0.234504 | 0.223026 | 0.395177 | 0.552333 | 0.781693 | 0.898646 | 1.248528 | 0.796472 | 0.398199 | 0.290185 | 0.453178 | 0.680976 | 1.002747 | 0.444089 | 0.282078 | 0.211940 | 0.233250 | 0.296241 | 0.284302 | 0.289890 | 0.313829 | 0.288981 | 0.222791 | 0.214432 | 0.215618 | 0.250160 | 0.280317 | 0.388230 | 0.306447 | 0.240495 | 0.213309 | 0.213000 | 0.221501 | 0.213134 | 0.225139 | 0.221701 | 0.210881 | 0.247965 | 0.323619 | 0.284240 | 0.231258 | 0.217301 | 0.217644 | 0.216535 | 0.213427 | 0.212296 | 0.212981 | 0.218130 | 0.217497 | 0.225005 | 0.248205 | 0.275357 | 0.311913 | 0.326811 | 0.299347 | 0.325186 | 0.445871 | 0.652185 | 1.177486 | 1.465960 | 3.047174 | 4.432931 | 2.701263 | 1.015738 | 0.447693 | 0.289722 | 0.252770 | 0.256587 | 0.314161 | 0.316629 | 0.395680 | 0.501870 | 0.777064 | 0.963223 | 0.840881 | 0.992889 | 0.937365 | 0.550518 | 0.429510 | 0.328045 | 0.286835 | 0.253551 | 0.221787 | 0.223933 | 0.234679 | 0.246989 | 0.249229 | 0.270945 | 0.335142 | 0.462404 | 0.752870 | 1.333089 | 2.582289 | 4.377943 | 5.532856 | 4.251633 | 2.589923 | 1.203235 | 0.816803 | 0.553832 | 0.346793 | 0.254785 | 0.233336 | 0.235448 | 0.261824 | 0.247941 | 0.233059 | 0.226653 | 0.230327 | 0.242746 | 0.265960 | 0.262931 | 0.283547 | 0.288231 | 0.289526 | 0.327937 | 0.311706 | 0.266200 | 0.235474 | 0.230724 | 0.237247 | 0.257145 | 0.277263 | 0.320481 | 0.342618 | 0.318577 | 0.307348 | 0.269137 | 0.249681 | 0.217637 | 0.210849 | 0.210818 | 0.216456 | 0.217292 | 0.213965 | 0.211358 | 0.218166 | 0.268232 | 0.437468 | 0.884094 | 1.047226 | 1.108362 | 0.520180 | 0.299774 | 0.219473 | 0.210937 | 0.212239 | 0.217085 | 0.220991 | 0.248776 | 0.277028 | 0.254028 | 0.219195 | 0.210869 | 0.215078 | 0.214406 | 0.215927 | 0.213864 | 0.214078 | 0.212589 | 0.214281 | 0.218066 | 0.239204 | 0.253577 | 0.297975 | 0.339389 | 0.364660 | 0.332655 | 0.285928 | 0.271476 | 0.281848 | 0.315418 | 0.395156 | 0.410628 | 0.357125 | 0.290303 | 0.256427 | 0.221132 | 0.218524 | 0.340002 | 0.542629 | 0.417596 | 0.256398 | 0.261301 | 0.229554 | 0.211841 | 0.228326 | 0.271945 | 0.266105 | 0.217848 | 0.213837 | 0.211556 | 0.210818 | 0.222670 | 0.230995 | 0.228660 | 0.210937 | 0.215995 | 0.214424 | 0.213798 | 0.229463 | 0.240593 | 0.253370 | 0.256026 | 0.232493 | 0.239102 | 0.296031 | 0.315694 | 0.356403 | 0.331795 | 0.269392 | 0.290589 | 0.256327 | 0.212233 | 0.221925 | 0.320207 | 0.452092 | 0.380958 | 0.390824 | 0.359259 | 0.288457 | 0.280358 | 0.223864 | 0.211331 | 0.210983 | 0.210817 | 0.211203 | 0.210817 | 0.210907 | 0.219294 | 0.257763 | 0.293588 | 0.320987 | 0.355134 | 0.377000 | 0.370584 | 0.387254 | 0.314073 | 0.265822 | 0.253491 | 0.285588 | 0.378049 | 0.418378 | 0.388429 | 0.450594 | 0.652243 | 0.910015 |
| left central | 0.528961 | 0.390608 | 0.324370 | 0.298893 | 0.243655 | 0.231601 | 0.211848 | 0.215261 | 0.212000 | 0.213784 | 0.245401 | 0.224922 | 0.233217 | 0.329396 | 0.498317 | 0.709921 | 0.625860 | 0.408058 | 0.407279 | 0.485301 | 0.365463 | 0.224237 | 0.210947 | 0.215345 | 0.249191 | 0.325691 | 0.529137 | 0.580872 | 1.009118 | 1.006966 | 0.998842 | 0.588525 | 0.663296 | 0.498242 | 0.493749 | 0.403144 | 0.283727 | 0.218477 | 0.210895 | 0.215095 | 0.225761 | 0.244232 | 0.224521 | 0.220753 | 0.217217 | 0.212075 | 0.212115 | 0.211802 | 0.210806 | 0.212568 | 0.246364 | 0.271407 | 0.407014 | 0.562285 | 0.683683 | 0.607634 | 0.426293 | 0.383578 | 0.353246 | 0.290277 | 0.272489 | 0.259456 | 0.253886 | 0.299526 | 0.271434 | 0.250619 | 0.238206 | 0.232054 | 0.212656 | 0.212097 | 0.212622 | 0.210818 | 0.218170 | 0.266250 | 0.304607 | 0.361771 | 0.306691 | 0.294731 | 0.274080 | 0.244846 | 0.217866 | 0.212058 | 0.211295 | 0.210806 | 0.210850 | 0.212887 | 0.213456 | 0.212835 | 0.212121 | 0.211462 | 0.240111 | 0.290788 | 0.344878 | 0.371915 | 0.280344 | 0.217232 | 0.226599 | 0.336480 | 0.864085 | 1.103525 | 0.811572 | 0.503583 | 0.304109 | 0.236481 | 0.222607 | 0.214891 | 0.214240 | 0.228631 | 0.247993 | 0.284184 | 0.262584 | 0.255616 | 0.236124 | 0.221512 | 0.211295 | 0.211845 | 0.219753 | 0.212879 | 0.211564 | 0.215675 | 0.252979 | 0.323867 | 0.355027 | 0.335812 | 0.303990 | 0.264432 | 0.213036 | 0.212752 | 0.219937 | 0.211489 | 0.221538 | 0.260870 | 0.357471 | 0.499105 | 0.458779 | 0.426275 | 0.358253 | 0.393100 | 0.461978 | 0.469609 | 0.510326 | 0.520648 | 0.468219 | 0.399771 | 0.352621 | 0.315885 | 0.280334 | 0.245169 | 0.220359 | 0.218900 | 0.226662 | 0.221799 | 0.217886 | 0.220740 | 0.224822 | 0.259058 | 0.283864 | 0.294642 | 0.302223 | 0.284748 | 0.281617 | 0.278109 | 0.256068 | 0.229599 | 0.215117 | 0.211559 | 0.215491 | 0.215379 | 0.215009 | 0.219884 | 0.238532 | 0.245626 | 0.221204 | 0.211811 | 0.210806 | 0.213740 | 0.220234 | 0.243136 | 0.240069 | 0.225097 | 0.217780 | 0.218415 | 0.212742 | 0.211970 | 0.211154 | 0.211386 | 0.210882 | 0.212363 | 0.220021 | 0.235781 | 0.246929 | 0.239934 | 0.239496 | 0.244749 | 0.268394 | 0.302873 | 0.310193 | 0.293500 | 0.298572 | 0.306069 | 0.326692 | 0.290807 | 0.261859 | 0.245176 | 0.233229 | 0.217099 | 0.210806 | 0.214880 | 0.219830 | 0.242330 | 0.271007 | 0.273190 | 0.247977 | 0.230125 | 0.216060 | 0.215119 | 0.216445 | 0.214405 | 0.212361 | 0.210892 | 0.211164 | 0.211788 | 0.210862 | 0.216275 | 0.227939 | 0.232498 | 0.223134 | 0.219235 | 0.211206 | 0.211039 | 0.212410 | 0.215164 | 0.231179 | 0.226267 | 0.215329 | 0.211064 | 0.211055 | 0.211153 | 0.210921 | 0.218066 | 0.226815 | 0.235218 | 0.254016 | 0.259458 | 0.276402 | 0.289533 | 0.265671 | 0.241073 | 0.234043 | 0.228680 | 0.228496 | 0.233476 | 0.227539 | 0.217469 | 0.214380 | 0.212531 | 0.212681 | 0.216121 | 0.213508 | 0.234723 | 0.326303 | 0.507887 | 0.807458 | 1.049397 | 1.032703 | 1.267693 | 0.936064 | 0.681991 | 0.633663 | 0.671454 | 0.730306 | 0.726165 | 0.740634 | 0.655092 | 0.717225 | 0.569470 | 0.477465 | 0.381090 | 0.349297 | 0.324147 |
| right central | 10.633432 | 26.615661 | 9.084356 | 7.137194 | 9.078008 | 5.263889 | 2.570456 | 1.836719 | 0.876101 | 0.850713 | 1.616761 | 1.776254 | 2.254733 | 1.038271 | 0.677351 | 0.738057 | 0.370534 | 0.222836 | 0.211647 | 0.212629 | 0.216975 | 0.210806 | 0.250527 | 0.289910 | 0.260724 | 0.343383 | 0.382917 | 0.622237 | 0.450310 | 0.237770 | 0.243629 | 0.352050 | 0.529448 | 0.378798 | 0.292644 | 0.260332 | 0.257102 | 0.211971 | 0.224959 | 0.213226 | 0.224292 | 0.336854 | 0.464149 | 0.380581 | 0.378031 | 0.305163 | 0.227911 | 0.212027 | 0.211227 | 0.213426 | 0.211020 | 0.222123 | 0.219199 | 0.214405 | 0.216591 | 0.229337 | 0.305377 | 0.596840 | 0.679290 | 0.670588 | 1.254874 | 1.249423 | 0.614793 | 0.307120 | 0.256290 | 0.244851 | 0.269952 | 0.245466 | 0.250274 | 0.252424 | 0.262125 | 0.226143 | 0.215247 | 0.211528 | 0.212462 | 0.225323 | 0.241506 | 0.299043 | 0.400371 | 0.551340 | 0.580203 | 0.363164 | 0.305860 | 0.289363 | 0.237486 | 0.210847 | 0.211848 | 0.211318 | 0.237147 | 0.280117 | 0.313370 | 0.523546 | 1.235824 | 1.491089 | 1.688273 | 0.860611 | 0.380124 | 0.213148 | 0.324085 | 0.570761 | 0.729405 | 1.355257 | 1.311135 | 1.113133 | 1.024197 | 0.835665 | 1.097265 | 2.333810 | 4.885027 | 14.001028 | 23.460140 | 16.525185 | 6.328528 | 1.755979 | 0.861095 | 0.536841 | 0.312132 | 0.283820 | 0.235177 | 0.231485 | 0.233156 | 0.252632 | 0.312549 | 0.486235 | 0.529056 | 0.689708 | 0.962535 | 2.065383 | 2.464650 | 2.309622 | 1.350131 | 1.099008 | 1.379464 | 1.496505 | 0.882961 | 0.598826 | 0.427587 | 0.427246 | 0.461843 | 0.533229 | 0.805503 | 2.419439 | 14.175255 | 99.352296 | 388.452390 | 507.535537 | 809.402313 | 556.689839 | 239.950319 | 55.521032 | 27.091049 | 19.420166 | 13.365997 | 7.088855 | 5.131758 | 4.950584 | 3.627997 | 2.864818 | 2.704673 | 2.772153 | 2.822410 | 2.582942 | 1.959761 | 1.883629 | 1.992991 | 1.100733 | 0.694419 | 0.496239 | 0.443431 | 0.506012 | 0.524133 | 0.479869 | 0.433175 | 0.393238 | 0.345615 | 0.284186 | 0.222984 | 0.211836 | 0.210818 | 0.215454 | 0.242224 | 0.246382 | 0.231335 | 0.231122 | 0.253552 | 0.289234 | 0.270902 | 0.237091 | 0.235079 | 0.256306 | 0.278060 | 0.251999 | 0.227864 | 0.222484 | 0.229790 | 0.225140 | 0.217929 | 0.218208 | 0.219509 | 0.218180 | 0.229883 | 0.221628 | 0.215879 | 0.218820 | 0.214762 | 0.220261 | 0.216335 | 0.212219 | 0.216071 | 0.211308 | 0.211346 | 0.219650 | 0.226517 | 0.236849 | 0.299570 | 0.434778 | 0.827677 | 1.075766 | 0.780892 | 0.606469 | 0.554792 | 0.379973 | 0.287048 | 0.228582 | 0.223249 | 0.231730 | 0.238815 | 0.249576 | 0.286029 | 0.323456 | 0.344086 | 0.296689 | 0.273437 | 0.257166 | 0.245520 | 0.213276 | 0.216928 | 0.272647 | 0.364960 | 0.501208 | 0.650463 | 0.770963 | 0.537131 | 0.396119 | 0.300340 | 0.229500 | 0.212808 | 0.243395 | 0.242086 | 0.216659 | 0.211193 | 0.214709 | 0.215816 | 0.216459 | 0.226582 | 0.275540 | 0.353783 | 0.413731 | 0.339188 | 0.327858 | 0.356767 | 0.376243 | 0.353772 | 0.331317 | 0.275046 | 0.259605 | 0.231527 | 0.213535 | 0.212835 | 0.220200 | 0.227811 | 0.230732 | 0.216225 | 0.210815 | 0.212075 | 0.226901 | 0.271281 | 0.321892 | 0.556755 | 0.814213 |
| left posterior | 0.258173 | 0.267933 | 0.279883 | 0.260437 | 0.269296 | 0.414042 | 0.582191 | 0.817843 | 0.608223 | 0.362245 | 0.247650 | 0.213469 | 0.262389 | 0.389183 | 0.891049 | 0.975704 | 0.598843 | 0.332315 | 0.327642 | 0.363954 | 0.451651 | 0.528805 | 0.391082 | 0.296121 | 0.270872 | 0.223242 | 0.210856 | 0.253335 | 0.544580 | 0.566456 | 0.525376 | 0.340648 | 0.243425 | 0.228575 | 0.237838 | 0.239830 | 0.261516 | 0.236845 | 0.221626 | 0.221902 | 0.226275 | 0.235535 | 0.287856 | 0.465475 | 0.928902 | 1.495830 | 1.913514 | 1.778947 | 0.860070 | 0.663574 | 0.515631 | 0.409385 | 0.468036 | 0.437886 | 0.502945 | 0.911413 | 0.682621 | 0.332303 | 0.232885 | 0.210941 | 0.248284 | 0.398767 | 1.046994 | 2.737829 | 1.903005 | 1.211828 | 0.704393 | 0.599041 | 0.716603 | 0.910327 | 2.161289 | 12.077124 | 137.416094 | 400.129087 | 255.697222 | 124.614170 | 50.618001 | 20.571986 | 11.432216 | 6.466354 | 3.316707 | 1.985214 | 0.741852 | 0.479230 | 0.356861 | 0.299944 | 0.297193 | 0.313114 | 0.320816 | 0.381221 | 0.420011 | 0.566928 | 0.864980 | 1.208309 | 1.670488 | 1.942053 | 2.864552 | 3.059970 | 1.987364 | 1.228608 | 0.764797 | 0.487730 | 0.368349 | 0.269001 | 0.228740 | 0.213325 | 0.216347 | 0.290325 | 0.482209 | 0.472773 | 0.349063 | 0.267251 | 0.230558 | 0.213636 | 0.211630 | 0.215462 | 0.211001 | 0.210814 | 0.211927 | 0.219350 | 0.228970 | 0.255519 | 0.301503 | 0.329693 | 0.276708 | 0.259086 | 0.266478 | 0.253340 | 0.218859 | 0.222475 | 0.283383 | 0.310075 | 0.371166 | 0.424272 | 0.371103 | 0.295382 | 0.211921 | 0.232837 | 0.250835 | 0.259637 | 0.236928 | 0.225047 | 0.223837 | 0.215800 | 0.215646 | 0.217877 | 0.213132 | 0.215348 | 0.219943 | 0.228329 | 0.233856 | 0.228046 | 0.238940 | 0.285811 | 0.321007 | 0.308321 | 0.282947 | 0.274684 | 0.280760 | 0.230995 | 0.211451 | 0.211615 | 0.210943 | 0.210885 | 0.218818 | 0.229484 | 0.271958 | 0.375890 | 0.523014 | 0.520955 | 0.476971 | 0.356884 | 0.284822 | 0.266754 | 0.221545 | 0.213460 | 0.236566 | 0.261563 | 0.327607 | 0.331888 | 0.365520 | 0.319082 | 0.280506 | 0.241716 | 0.213453 | 0.235848 | 0.274757 | 0.308948 | 0.368585 | 0.434516 | 0.507765 | 0.564789 | 0.528755 | 0.501485 | 0.435777 | 0.333636 | 0.296034 | 0.285072 | 0.268216 | 0.274960 | 0.279576 | 0.273449 | 0.327844 | 0.376969 | 0.346678 | 0.362333 | 0.363385 | 0.363732 | 0.356235 | 0.304986 | 0.276371 | 0.241957 | 0.212735 | 0.240375 | 0.332186 | 0.421145 | 0.625115 | 0.785285 | 0.716676 | 0.484515 | 0.279560 | 0.222687 | 0.211758 | 0.214259 | 0.215248 | 0.215288 | 0.210995 | 0.218940 | 0.237597 | 0.295076 | 0.376942 | 0.468759 | 0.525400 | 0.482126 | 0.420606 | 0.396736 | 0.296478 | 0.259462 | 0.245076 | 0.248963 | 0.252280 | 0.267671 | 0.273312 | 0.313746 | 0.346773 | 0.379665 | 0.402617 | 0.428059 | 0.367041 | 0.384970 | 0.374106 | 0.419151 | 0.391295 | 0.303385 | 0.269558 | 0.284387 | 0.302068 | 0.318673 | 0.351966 | 0.420647 | 0.732874 | 1.048768 | 1.281444 | 1.130040 | 0.689550 | 0.355948 | 0.252318 | 0.215331 | 0.211092 | 0.217429 | 0.220340 | 0.210806 | 0.237102 | 0.355706 | 0.714721 | 1.306654 | 1.748198 | 2.427266 | 2.629355 | 2.450294 |
| right posterior | 0.219294 | 0.219407 | 0.216164 | 0.214441 | 0.221383 | 0.319783 | 0.312722 | 0.351973 | 0.303866 | 0.303658 | 0.308191 | 0.217724 | 0.227733 | 0.212263 | 0.226020 | 0.330444 | 0.313772 | 0.259537 | 0.278894 | 0.309183 | 0.216906 | 0.229430 | 0.243391 | 0.219016 | 0.214192 | 0.296215 | 0.386024 | 0.761881 | 0.522151 | 0.284763 | 0.222820 | 0.211145 | 0.236315 | 0.240011 | 0.233917 | 0.211323 | 0.219260 | 0.266908 | 0.401211 | 0.456884 | 0.493896 | 0.452733 | 0.383238 | 0.299426 | 0.261579 | 0.225002 | 0.212118 | 0.239823 | 0.357499 | 0.398903 | 0.266402 | 0.239656 | 0.249105 | 0.273664 | 0.313108 | 0.496523 | 1.849222 | 26.375544 | 104.197361 | 87.336674 | 102.792613 | 58.755571 | 25.606831 | 12.581239 | 7.109804 | 6.734101 | 4.318693 | 2.520594 | 1.756153 | 2.022150 | 2.202725 | 3.038946 | 9.812374 | 64.594871 | 251.913791 | 295.107229 | 216.326886 | 86.999135 | 31.362384 | 25.021037 | 14.051493 | 12.118493 | 8.144442 | 3.717862 | 1.949786 | 1.280594 | 0.631949 | 0.413188 | 0.346068 | 0.346581 | 0.486099 | 0.849122 | 1.435238 | 1.546926 | 1.223324 | 0.822674 | 0.532313 | 0.309018 | 0.242358 | 0.223854 | 0.239741 | 0.284929 | 0.304444 | 0.300627 | 0.289702 | 0.255021 | 0.226068 | 0.212273 | 0.210963 | 0.211192 | 0.211301 | 0.211554 | 0.213500 | 0.219541 | 0.245761 | 0.284530 | 0.294240 | 0.300790 | 0.310677 | 0.395993 | 0.435131 | 0.394947 | 0.386258 | 0.376476 | 0.378450 | 0.345907 | 0.309427 | 0.278390 | 0.252199 | 0.224367 | 0.212356 | 0.212283 | 0.232117 | 0.257969 | 0.251103 | 0.230845 | 0.222600 | 0.220822 | 0.216468 | 0.210806 | 0.213507 | 0.220149 | 0.215813 | 0.217785 | 0.245676 | 0.263117 | 0.242271 | 0.225354 | 0.215097 | 0.215470 | 0.214301 | 0.214586 | 0.221755 | 0.247017 | 0.285546 | 0.362030 | 0.500080 | 0.981149 | 1.616782 | 2.418792 | 3.032121 | 3.666862 | 5.018201 | 4.426731 | 2.879535 | 2.435351 | 2.605803 | 1.949000 | 1.585951 | 1.013689 | 1.097059 | 1.283359 | 1.643454 | 1.675875 | 2.315752 | 1.813802 | 1.579423 | 0.852295 | 0.510852 | 0.376175 | 0.272661 | 0.240649 | 0.244027 | 0.246914 | 0.287183 | 0.354162 | 0.415850 | 0.510221 | 0.617682 | 0.834221 | 1.231563 | 1.827516 | 2.645106 | 4.095218 | 4.825292 | 5.274702 | 6.718252 | 9.805042 | 11.475019 | 8.416647 | 4.613587 | 2.779588 | 2.145753 | 1.591338 | 1.351374 | 1.298115 | 1.179369 | 1.251353 | 1.597187 | 1.462270 | 1.292713 | 1.232360 | 1.079708 | 1.009914 | 1.128714 | 1.046833 | 0.972114 | 0.982779 | 0.797289 | 0.695806 | 0.557663 | 0.423978 | 0.368012 | 0.378479 | 0.424789 | 0.480546 | 0.474682 | 0.484115 | 0.426209 | 0.445866 | 0.412513 | 0.366611 | 0.349078 | 0.359302 | 0.382326 | 0.492028 | 0.712789 | 1.074631 | 1.651586 | 2.346038 | 2.795065 | 2.951354 | 2.508982 | 1.903977 | 1.525812 | 1.114198 | 1.134296 | 1.173861 | 1.135299 | 1.247302 | 1.256408 | 1.305217 | 1.209762 | 0.901235 | 0.720874 | 0.652530 | 0.548089 | 0.493656 | 0.396837 | 0.418170 | 0.452904 | 0.582460 | 0.734457 | 1.280958 | 2.241071 | 3.347391 | 3.796417 | 3.944396 | 3.465538 | 3.695607 | 2.013564 | 1.199861 | 1.335921 | 1.466959 | 1.550113 | 1.667839 | 1.756880 | 2.021863 | 2.589715 | 2.415314 |
| all electrodes | 0.214392 | 0.252472 | 0.296869 | 0.334449 | 0.426457 | 1.183846 | 1.072946 | 0.618994 | 0.413093 | 0.252278 | 0.214879 | 0.234959 | 0.224081 | 0.223663 | 0.231613 | 0.210911 | 0.234627 | 0.230238 | 0.223630 | 0.218623 | 0.232255 | 0.228290 | 0.303640 | 0.256138 | 0.356892 | 0.910995 | 1.129675 | 1.119092 | 0.557935 | 0.356111 | 0.251229 | 0.212296 | 0.263536 | 0.249259 | 0.246437 | 0.216081 | 0.279901 | 0.383249 | 0.575847 | 0.470252 | 0.332964 | 0.268592 | 0.223061 | 0.213746 | 0.213292 | 0.226978 | 0.244372 | 0.233917 | 0.216639 | 0.213422 | 0.224230 | 0.259811 | 0.246029 | 0.225774 | 0.219839 | 0.219291 | 0.210856 | 0.255095 | 0.383268 | 0.441534 | 0.492224 | 0.451830 | 0.419123 | 0.371859 | 0.276879 | 0.259117 | 0.314349 | 0.363629 | 0.555147 | 1.267762 | 2.155690 | 7.051207 | 17.169600 | 43.361089 | 85.719755 | 135.381573 | 141.807918 | 172.764438 | 174.005230 | 398.632205 | 271.403589 | 112.557671 | 91.694070 | 44.622659 | 21.201485 | 7.610745 | 2.627985 | 1.350615 | 1.153963 | 0.823429 | 0.675763 | 0.665711 | 0.779293 | 1.176506 | 1.570950 | 1.489434 | 0.968080 | 0.555657 | 0.322070 | 0.291838 | 0.389681 | 0.635082 | 0.859767 | 1.464929 | 2.465516 | 4.833973 | 6.739594 | 3.516072 | 2.144693 | 1.791656 | 1.447392 | 1.763516 | 1.892165 | 2.412320 | 2.666791 | 2.603276 | 2.369002 | 1.254583 | 0.778731 | 0.623947 | 0.724512 | 0.733982 | 0.628677 | 0.441411 | 0.449691 | 0.394661 | 0.321646 | 0.266590 | 0.251807 | 0.242164 | 0.217181 | 0.211072 | 0.210853 | 0.216885 | 0.234746 | 0.245586 | 0.288648 | 0.402419 | 0.418690 | 0.404642 | 0.346510 | 0.247867 | 0.224417 | 0.216761 | 0.217796 | 0.221478 | 0.221358 | 0.227055 | 0.259945 | 0.278971 | 0.265074 | 0.232584 | 0.223155 | 0.239191 | 0.243948 | 0.251144 | 0.257535 | 0.281794 | 0.342842 | 0.458794 | 0.464124 | 0.556238 | 0.500911 | 0.536210 | 0.560074 | 0.693041 | 0.796337 | 0.938238 | 0.912900 | 0.790020 | 0.696558 | 0.555897 | 0.465371 | 0.348056 | 0.304764 | 0.301849 | 0.331929 | 0.357540 | 0.335777 | 0.304264 | 0.389424 | 0.717258 | 1.274348 | 1.914940 | 3.084830 | 5.252880 | 8.235851 | 15.460122 | 12.131520 | 12.647377 | 13.084038 | 10.694017 | 15.861590 | 13.333901 | 7.962380 | 8.201423 | 5.776591 | 5.900915 | 6.732605 | 6.753876 | 11.433954 | 12.424932 | 6.053004 | 5.713702 | 6.114565 | 8.878622 | 10.157566 | 8.644397 | 10.061868 | 15.017558 | 18.321247 | 22.972361 | 18.118253 | 19.875328 | 22.432116 | 17.664792 | 9.527230 | 4.722771 | 2.705211 | 2.224718 | 1.994714 | 1.677209 | 1.732911 | 2.129808 | 3.370975 | 5.280830 | 6.090367 | 4.827272 | 4.124439 | 4.433844 | 7.453983 | 6.931310 | 5.600377 | 5.955243 | 7.187901 | 11.931075 | 13.513234 | 13.929022 | 18.762787 | 21.225448 | 27.467014 | 29.092529 | 43.580357 | 79.377640 | 68.916706 | 31.355082 | 18.358814 | 9.459219 | 7.647287 | 5.863706 | 3.508270 | 2.120457 | 1.658551 | 1.471970 | 1.420043 | 1.373237 | 1.119013 | 0.977457 | 1.198040 | 1.468756 | 1.335283 | 1.441045 | 1.757049 | 2.458483 | 4.583790 | 6.373973 | 8.940058 | 14.647360 | 18.779753 | 13.754817 | 7.858169 | 4.100038 | 3.208150 | 2.137586 | 1.428680 | 1.011081 | 0.751324 | 0.699866 | 0.640707 | 0.482129 |

Searchlight, spatiotemporal cluster permutation test

|  | start time | stop time | peak time | peak channel | cluster p | peak Cohen's d | direction |
| --- | --- | --- | --- | --- | --- | --- | --- |
| #1 | 75 | 470 | 175 | O1 | 0.0099 | 1.085016 | positive |

D) emoji faces (LOSO) - neutral vs happy

  
|  | time window | peak latency | cluster *p* | peak Cohen's *d* |  | | | |
| **all electrodes** | 80 - 160 ms | 130 ms | 0.0296 | 0.974 |  | | | |
|  | | | | | | | | |

Time-resolved classification, cluster permutation tests

|  | **left hemisphere** | | | | **right hemisphere** | | | |
|  | time window | peak latency | cluster *p* | peak Cohen's *d* | time window | peak latency | cluster *p* | peak Cohen's *d* |
| **anterior** | 775 - 860 ms | 835 ms | 0.0381 | 0.7821 |  | | | |
| **central** |  | | | |  | | | |
| **posterior** | 100 - 160 ms | 135 ms | 0.04 | 1.1164 | 75 - 170 ms | 140 ms | 0.0167 | 1.4459 |
 295 - 405 ms | 295 ms | 0.0079 | -0.4208 |  | | | | 615 - 770 ms | 615 ms | 0.0069 | -0.5316 |  | | | | 900 - 1050 ms | 900 ms | 0.0096 | -0.4653 |  | | | |

  

Time-resolved classification, Bayesian statistics

|  | -200 | -195 | -190 | -185 | -180 | -175 | -170 | -165 | -160 | -155 | -150 | -145 | -140 | -135 | -130 | -125 | -120 | -115 | -110 | -105 | -100 | -95 | -90 | -85 | -80 | -75 | -70 | -65 | -60 | -55 | -50 | -45 | -40 | -35 | -30 | -25 | -20 | -15 | -10 | -5 | 0 | 5 | 10 | 15 | 20 | 25 | 30 | 35 | 40 | 45 | 50 | 55 | 60 | 65 | 70 | 75 | 80 | 85 | 90 | 95 | 100 | 105 | 110 | 115 | 120 | 125 | 130 | 135 | 140 | 145 | 150 | 155 | 160 | 165 | 170 | 175 | 180 | 185 | 190 | 195 | 200 | 205 | 210 | 215 | 220 | 225 | 230 | 235 | 240 | 245 | 250 | 255 | 260 | 265 | 270 | 275 | 280 | 285 | 290 | 295 | 300 | 305 | 310 | 315 | 320 | 325 | 330 | 335 | 340 | 345 | 350 | 355 | 360 | 365 | 370 | 375 | 380 | 385 | 390 | 395 | 400 | 405 | 410 | 415 | 420 | 425 | 430 | 435 | 440 | 445 | 450 | 455 | 460 | 465 | 470 | 475 | 480 | 485 | 490 | 495 | 500 | 505 | 510 | 515 | 520 | 525 | 530 | 535 | 540 | 545 | 550 | 555 | 560 | 565 | 570 | 575 | 580 | 585 | 590 | 595 | 600 | 605 | 610 | 615 | 620 | 625 | 630 | 635 | 640 | 645 | 650 | 655 | 660 | 665 | 670 | 675 | 680 | 685 | 690 | 695 | 700 | 705 | 710 | 715 | 720 | 725 | 730 | 735 | 740 | 745 | 750 | 755 | 760 | 765 | 770 | 775 | 780 | 785 | 790 | 795 | 800 | 805 | 810 | 815 | 820 | 825 | 830 | 835 | 840 | 845 | 850 | 855 | 860 | 865 | 870 | 875 | 880 | 885 | 890 | 895 | 900 | 905 | 910 | 915 | 920 | 925 | 930 | 935 | 940 | 945 | 950 | 955 | 960 | 965 | 970 | 975 | 980 | 985 | 990 | 995 | 1000 | 1005 | 1010 | 1015 | 1020 | 1025 | 1030 | 1035 | 1040 | 1045 | 1050 | 1055 | 1060 | 1065 | 1070 | 1075 | 1080 | 1085 | 1090 | 1095 | 1100 | 1105 | 1110 | 1115 | 1120 | 1125 | 1130 | 1135 | 1140 | 1145 | 1150 | 1155 | 1160 | 1165 | 1170 | 1175 | 1180 | 1185 | 1190 | 1195 |
| --- | --- | --- | --- | --- | --- | --- | --- | --- | --- | --- | --- | --- | --- | --- | --- | --- | --- | --- | --- | --- | --- | --- | --- | --- | --- | --- | --- | --- | --- | --- | --- | --- | --- | --- | --- | --- | --- | --- | --- | --- | --- | --- | --- | --- | --- | --- | --- | --- | --- | --- | --- | --- | --- | --- | --- | --- | --- | --- | --- | --- | --- | --- | --- | --- | --- | --- | --- | --- | --- | --- | --- | --- | --- | --- | --- | --- | --- | --- | --- | --- | --- | --- | --- | --- | --- | --- | --- | --- | --- | --- | --- | --- | --- | --- | --- | --- | --- | --- | --- | --- | --- | --- | --- | --- | --- | --- | --- | --- | --- | --- | --- | --- | --- | --- | --- | --- | --- | --- | --- | --- | --- | --- | --- | --- | --- | --- | --- | --- | --- | --- | --- | --- | --- | --- | --- | --- | --- | --- | --- | --- | --- | --- | --- | --- | --- | --- | --- | --- | --- | --- | --- | --- | --- | --- | --- | --- | --- | --- | --- | --- | --- | --- | --- | --- | --- | --- | --- | --- | --- | --- | --- | --- | --- | --- | --- | --- | --- | --- | --- | --- | --- | --- | --- | --- | --- | --- | --- | --- | --- | --- | --- | --- | --- | --- | --- | --- | --- | --- | --- | --- | --- | --- | --- | --- | --- | --- | --- | --- | --- | --- | --- | --- | --- | --- | --- | --- | --- | --- | --- | --- | --- | --- | --- | --- | --- | --- | --- | --- | --- | --- | --- | --- | --- | --- | --- | --- | --- | --- | --- | --- | --- | --- | --- | --- | --- | --- | --- | --- | --- | --- | --- | --- | --- | --- | --- | --- | --- | --- | --- | --- | --- | --- | --- | --- | --- | --- | --- | --- | --- | --- | --- | --- | --- | --- | --- | --- | --- | --- | --- | --- |
| left anterior | 0.410517 | 0.677200 | 0.593426 | 0.571806 | 0.427052 | 0.322101 | 0.237299 | 0.217276 | 0.346925 | 0.366353 | 0.296061 | 0.259369 | 0.231267 | 0.239201 | 0.213051 | 0.226961 | 0.210911 | 0.232518 | 0.267398 | 0.285151 | 0.237102 | 0.231551 | 0.253891 | 0.221212 | 0.214325 | 0.213391 | 0.244424 | 0.271321 | 0.242875 | 0.220693 | 0.213701 | 0.220798 | 0.216901 | 0.231932 | 0.223728 | 0.214727 | 0.214957 | 0.217857 | 0.241989 | 0.363799 | 0.749343 | 0.576069 | 0.619996 | 0.425937 | 0.284548 | 0.214284 | 0.235793 | 0.336267 | 0.387110 | 0.356898 | 0.259352 | 0.227339 | 0.210952 | 0.304103 | 0.862642 | 1.246656 | 0.979349 | 0.403620 | 0.311308 | 0.214174 | 0.231827 | 0.258745 | 0.305499 | 0.388747 | 0.383785 | 0.453661 | 0.450301 | 0.442885 | 0.480907 | 0.420135 | 0.462783 | 0.440645 | 0.442390 | 0.396542 | 0.403423 | 0.540919 | 0.670535 | 0.506905 | 0.462653 | 0.393462 | 0.392241 | 0.394557 | 0.317589 | 0.236398 | 0.217728 | 0.212178 | 0.210907 | 0.215657 | 0.235818 | 0.241276 | 0.213222 | 0.214059 | 0.247619 | 0.266621 | 0.292711 | 0.311904 | 0.316065 | 0.305296 | 0.276456 | 0.216816 | 0.210881 | 0.219505 | 0.256255 | 0.294644 | 0.270870 | 0.267868 | 0.225409 | 0.218505 | 0.213512 | 0.219309 | 0.217562 | 0.210806 | 0.210822 | 0.225917 | 0.216969 | 0.211289 | 0.210827 | 0.265891 | 0.500220 | 0.646926 | 0.829576 | 0.831711 | 0.895318 | 1.007477 | 0.928917 | 0.818983 | 0.700235 | 0.748859 | 0.872773 | 0.801782 | 1.360655 | 1.837885 | 1.536411 | 1.318474 | 1.077926 | 1.571635 | 2.533534 | 1.795407 | 1.186216 | 1.061124 | 1.047269 | 1.032273 | 0.714461 | 0.513346 | 0.427617 | 0.555298 | 0.772320 | 1.593776 | 2.440704 | 2.164512 | 2.816087 | 2.916511 | 1.851760 | 1.229304 | 0.632275 | 0.472790 | 0.413651 | 0.320251 | 0.244657 | 0.213859 | 0.211086 | 0.212993 | 0.235259 | 0.245745 | 0.272073 | 0.402323 | 0.750572 | 1.680932 | 2.565925 | 2.100169 | 1.411053 | 1.098637 | 0.796944 | 0.534548 | 0.403958 | 0.428744 | 0.361455 | 0.469708 | 0.658757 | 0.872852 | 1.767770 | 2.219161 | 1.961772 | 1.997853 | 1.775727 | 1.934403 | 3.698937 | 4.623298 | 2.961166 | 1.685113 | 1.352830 | 1.377728 | 0.949141 | 0.870493 | 1.151177 | 3.282315 | 7.950302 | 16.712046 | 70.521598 | 158.652519 | 63.911508 | 20.119543 | 9.719444 | 14.316293 | 36.008523 | 26.641455 | 37.869973 | 50.030538 | 41.425352 | 17.111192 | 6.137790 | 3.160075 | 1.586134 | 0.614042 | 0.361880 | 0.290420 | 0.238605 | 0.236380 | 0.216911 | 0.212832 | 0.211540 | 0.213155 | 0.219963 | 0.244779 | 0.256311 | 0.265911 | 0.290688 | 0.298665 | 0.278182 | 0.295458 | 0.268144 | 0.250990 | 0.231545 | 0.219579 | 0.217415 | 0.211783 | 0.212285 | 0.212999 | 0.211216 | 0.210822 | 0.210822 | 0.211535 | 0.230361 | 0.251629 | 0.277900 | 0.267736 | 0.319830 | 0.347933 | 0.448948 | 0.474882 | 0.447359 | 0.308131 | 0.239642 | 0.242238 | 0.284187 | 0.290653 | 0.287954 | 0.282494 | 0.369307 | 0.608610 | 0.829441 | 0.680677 | 0.548765 | 0.478729 | 0.559944 | 0.615075 | 0.648825 | 0.538643 | 0.726419 | 0.879851 | 1.148779 | 1.125506 | 1.058938 | 0.973059 | 0.960166 | 0.579278 | 0.403762 | 0.299260 | 0.256946 | 0.240074 |
| right anterior | 1.836642 | 4.038557 | 4.751660 | 2.444004 | 1.421778 | 0.881278 | 0.467183 | 0.258159 | 0.226899 | 0.210890 | 0.211189 | 0.211058 | 0.210806 | 0.296056 | 0.923878 | 3.556369 | 2.257253 | 2.566469 | 1.875663 | 1.784067 | 0.481033 | 0.226669 | 0.211325 | 0.229585 | 0.282011 | 0.338437 | 0.381461 | 0.283410 | 0.214710 | 0.331579 | 0.511124 | 0.565917 | 0.805579 | 1.313303 | 1.054856 | 0.413201 | 0.275426 | 0.280444 | 0.300043 | 0.341961 | 0.313170 | 0.392983 | 0.577518 | 0.529295 | 0.580300 | 0.772004 | 0.661085 | 0.534521 | 0.375644 | 0.391411 | 0.769835 | 0.608922 | 0.399084 | 0.333442 | 0.269461 | 0.217971 | 0.211336 | 0.238358 | 0.246861 | 0.256518 | 0.278811 | 0.243771 | 0.220827 | 0.251167 | 0.332307 | 0.475323 | 0.469514 | 0.462145 | 0.515134 | 0.797871 | 0.772587 | 0.551757 | 0.438117 | 0.367528 | 0.305963 | 0.298868 | 0.218471 | 0.211599 | 0.211280 | 0.249148 | 0.275059 | 0.341439 | 0.281502 | 0.238628 | 0.210806 | 0.217615 | 0.252059 | 0.262886 | 0.393361 | 0.484127 | 0.662146 | 0.520682 | 0.420468 | 0.287307 | 0.275154 | 0.246748 | 0.261085 | 0.246010 | 0.301107 | 0.575407 | 2.063372 | 3.645378 | 5.929242 | 3.595333 | 1.894346 | 0.947385 | 0.469768 | 0.365796 | 0.351626 | 0.291642 | 0.350617 | 0.398829 | 0.400514 | 0.391128 | 0.409055 | 0.315160 | 0.270009 | 0.233834 | 0.232661 | 0.264055 | 0.306535 | 0.273083 | 0.274209 | 0.269359 | 0.228745 | 0.215294 | 0.220888 | 0.253249 | 0.274830 | 0.278813 | 0.236421 | 0.228660 | 0.400654 | 0.824606 | 2.008976 | 4.170096 | 4.866808 | 2.597637 | 1.537080 | 1.301798 | 1.735128 | 1.601605 | 1.503464 | 1.139891 | 0.693961 | 0.409405 | 0.228580 | 0.210869 | 0.221190 | 0.260705 | 0.274394 | 0.249626 | 0.231529 | 0.212031 | 0.221243 | 0.221592 | 0.211415 | 0.211659 | 0.224551 | 0.281610 | 0.335471 | 0.348989 | 0.405039 | 0.524099 | 0.448184 | 0.396279 | 0.326527 | 0.311942 | 0.298554 | 0.278744 | 0.277193 | 0.292329 | 0.280821 | 0.299872 | 0.271584 | 0.300510 | 0.372380 | 0.582039 | 0.850541 | 0.931787 | 0.591609 | 0.486691 | 0.399257 | 0.382135 | 0.279764 | 0.229832 | 0.215488 | 0.212453 | 0.210818 | 0.212498 | 0.235421 | 0.256389 | 0.277534 | 0.270832 | 0.274285 | 0.338538 | 0.436661 | 0.548677 | 0.488993 | 0.407873 | 0.428854 | 0.573879 | 0.769619 | 1.223887 | 1.244146 | 1.707818 | 1.686988 | 1.252651 | 0.968872 | 0.571034 | 0.307470 | 0.231630 | 0.213331 | 0.231894 | 0.238341 | 0.281317 | 0.300261 | 0.282440 | 0.246916 | 0.218197 | 0.215325 | 0.220305 | 0.225355 | 0.250447 | 0.289529 | 0.347222 | 0.368188 | 0.288095 | 0.237395 | 0.215248 | 0.212125 | 0.225660 | 0.235395 | 0.235429 | 0.212008 | 0.217522 | 0.235726 | 0.252907 | 0.291950 | 0.299864 | 0.283349 | 0.255578 | 0.239127 | 0.237319 | 0.250250 | 0.282485 | 0.337054 | 0.464482 | 0.522422 | 0.497255 | 0.430544 | 0.401594 | 0.388981 | 0.344780 | 0.288144 | 0.263023 | 0.255761 | 0.261951 | 0.229931 | 0.213001 | 0.211190 | 0.213644 | 0.216360 | 0.215573 | 0.225601 | 0.233375 | 0.283654 | 0.315043 | 0.287635 | 0.256755 | 0.264704 | 0.247628 | 0.232342 | 0.217710 | 0.214643 | 0.211081 | 0.215894 | 0.224484 | 0.227836 | 0.252356 |
| left central | 12.739329 | 76.664472 | 65.279686 | 11.334081 | 1.499251 | 0.453765 | 0.255956 | 0.237485 | 0.324382 | 0.265583 | 0.214907 | 0.227067 | 0.237973 | 0.211154 | 0.225379 | 0.323931 | 0.552571 | 0.725719 | 1.352842 | 3.034575 | 2.099116 | 0.682925 | 0.264959 | 0.225475 | 0.264337 | 0.222510 | 0.216580 | 0.250922 | 0.267332 | 0.232092 | 0.227227 | 0.391767 | 0.391337 | 0.406244 | 0.426728 | 0.377838 | 0.275798 | 0.262351 | 0.384749 | 0.338834 | 0.273101 | 0.238553 | 0.239173 | 0.237296 | 0.215420 | 0.211972 | 0.211750 | 0.216985 | 0.218777 | 0.221865 | 0.234918 | 0.279024 | 0.255805 | 0.232034 | 0.391952 | 0.294650 | 0.314491 | 0.262263 | 0.237779 | 0.248783 | 0.232728 | 0.247214 | 0.379619 | 0.585127 | 1.894642 | 5.671575 | 10.486074 | 14.841394 | 10.127556 | 9.218445 | 5.981412 | 2.368939 | 1.543464 | 0.729352 | 0.466950 | 0.392198 | 0.349916 | 0.323345 | 0.320565 | 0.360227 | 0.391217 | 0.330123 | 0.244148 | 0.211002 | 0.265149 | 0.371710 | 0.728937 | 2.401604 | 2.112084 | 1.172524 | 0.630146 | 0.318789 | 0.250897 | 0.248802 | 0.232046 | 0.225386 | 0.249417 | 0.306350 | 0.575253 | 1.570865 | 1.975388 | 2.208532 | 4.599051 | 5.316464 | 5.309108 | 2.910709 | 1.624341 | 1.370026 | 1.038090 | 0.849283 | 0.596268 | 0.494719 | 0.467737 | 0.478404 | 0.409457 | 0.353333 | 0.273531 | 0.226886 | 0.211744 | 0.212094 | 0.212423 | 0.210931 | 0.210820 | 0.218294 | 0.239851 | 0.225618 | 0.210868 | 0.242309 | 0.352220 | 0.457079 | 0.546748 | 0.377836 | 0.244499 | 0.222366 | 0.216957 | 0.211551 | 0.210887 | 0.216273 | 0.212383 | 0.211097 | 0.225476 | 0.280958 | 0.285970 | 0.277727 | 0.265304 | 0.250151 | 0.216370 | 0.218256 | 0.267063 | 0.304510 | 0.352012 | 0.424075 | 0.461305 | 0.350060 | 0.327909 | 0.400474 | 0.501329 | 0.502530 | 0.449650 | 0.364348 | 0.335616 | 0.263401 | 0.225972 | 0.213016 | 0.211906 | 0.228121 | 0.231749 | 0.229163 | 0.219781 | 0.219625 | 0.224777 | 0.222563 | 0.225089 | 0.215449 | 0.210836 | 0.211523 | 0.216164 | 0.234073 | 0.254060 | 0.303040 | 0.270353 | 0.237952 | 0.221015 | 0.211636 | 0.211860 | 0.211703 | 0.210948 | 0.219114 | 0.234157 | 0.256213 | 0.266251 | 0.251125 | 0.241533 | 0.235804 | 0.236474 | 0.246672 | 0.232650 | 0.228569 | 0.224554 | 0.215873 | 0.211194 | 0.214958 | 0.216974 | 0.220183 | 0.219728 | 0.212739 | 0.213474 | 0.210806 | 0.212206 | 0.215777 | 0.239395 | 0.261079 | 0.278262 | 0.335969 | 0.352107 | 0.297468 | 0.278551 | 0.248539 | 0.254519 | 0.254021 | 0.278198 | 0.304801 | 0.341845 | 0.340126 | 0.419808 | 0.399972 | 0.385259 | 0.328427 | 0.313825 | 0.323217 | 0.343342 | 0.310038 | 0.301734 | 0.327698 | 0.373630 | 0.369066 | 0.364874 | 0.361525 | 0.411500 | 0.497265 | 0.466058 | 0.420283 | 0.433647 | 0.442588 | 0.414920 | 0.416512 | 0.385565 | 0.383551 | 0.428925 | 0.405309 | 0.447496 | 0.477981 | 0.413549 | 0.391317 | 0.418965 | 0.399985 | 0.417479 | 0.381911 | 0.391826 | 0.406086 | 0.461245 | 0.485659 | 0.508093 | 0.534088 | 0.515884 | 0.524812 | 0.524205 | 0.462244 | 0.399633 | 0.381168 | 0.427936 | 0.493814 | 0.529816 | 0.706859 | 1.021375 | 1.578657 | 1.763565 | 1.601517 | 1.613207 | 1.968276 |
| right central | 0.269132 | 0.660745 | 1.720953 | 2.989407 | 9.800161 | 43.051258 | 89.879087 | 47.545959 | 6.541305 | 1.217340 | 0.590046 | 0.349966 | 0.288815 | 0.326595 | 0.368690 | 0.504341 | 0.549293 | 0.504662 | 0.447332 | 0.273524 | 0.213974 | 0.297847 | 0.677716 | 1.062235 | 1.664193 | 1.237137 | 0.311878 | 0.214566 | 0.402050 | 1.452497 | 2.462021 | 2.690329 | 0.678065 | 0.312718 | 0.243668 | 0.232605 | 0.270623 | 0.404827 | 1.024745 | 3.141548 | 1.407613 | 0.594991 | 0.336462 | 0.224459 | 0.211135 | 0.211634 | 0.242938 | 0.335160 | 0.551753 | 0.666885 | 0.624971 | 0.542757 | 0.222554 | 0.365856 | 0.869295 | 1.891585 | 2.308602 | 1.468927 | 0.756227 | 0.497785 | 0.353277 | 0.302503 | 0.298647 | 0.386145 | 0.636901 | 1.718561 | 2.864369 | 2.541669 | 1.790756 | 0.984351 | 0.559853 | 0.353805 | 0.307015 | 0.255763 | 0.251690 | 0.277241 | 0.302933 | 0.299801 | 0.399003 | 0.440261 | 0.465693 | 0.352956 | 0.276368 | 0.223776 | 0.211155 | 0.242972 | 0.300824 | 0.344923 | 0.371845 | 0.371915 | 0.335745 | 0.335103 | 0.313415 | 0.344073 | 0.261195 | 0.230399 | 0.234795 | 0.377279 | 0.747326 | 2.054814 | 2.743932 | 2.951631 | 2.904522 | 4.273750 | 5.515470 | 23.678214 | 37.454721 | 43.089722 | 187.842483 | 223.729466 | 109.570811 | 17.295410 | 2.773165 | 1.089559 | 1.182029 | 1.364989 | 2.067619 | 1.986676 | 0.786322 | 0.380197 | 0.338838 | 0.359420 | 0.462708 | 0.763002 | 1.781156 | 3.995245 | 9.120426 | 18.571766 | 33.387155 | 14.564670 | 4.187341 | 1.432831 | 0.966822 | 1.008331 | 0.780991 | 0.583265 | 0.597444 | 0.571863 | 0.526470 | 0.658540 | 0.785449 | 1.131728 | 1.134764 | 2.218884 | 4.896816 | 7.603031 | 5.868565 | 3.696649 | 1.758268 | 1.342072 | 0.984653 | 1.572735 | 1.930075 | 2.582010 | 3.863387 | 11.135704 | 17.911336 | 20.757251 | 9.997867 | 6.968874 | 5.122590 | 3.529628 | 2.038073 | 1.160884 | 0.807558 | 0.679765 | 0.585810 | 0.382694 | 0.343036 | 0.381308 | 0.408362 | 0.387202 | 0.374345 | 0.364507 | 0.406547 | 0.354966 | 0.274489 | 0.254990 | 0.252354 | 0.232599 | 0.216034 | 0.217108 | 0.232326 | 0.296696 | 0.335993 | 0.367692 | 0.400722 | 0.496415 | 0.450729 | 0.342750 | 0.266820 | 0.234349 | 0.215307 | 0.212973 | 0.210954 | 0.210956 | 0.212444 | 0.210845 | 0.211077 | 0.211457 | 0.210806 | 0.210990 | 0.212083 | 0.224693 | 0.258245 | 0.271337 | 0.301708 | 0.366699 | 0.443596 | 0.382729 | 0.325350 | 0.305877 | 0.363823 | 0.358868 | 0.281684 | 0.266881 | 0.268713 | 0.280105 | 0.307199 | 0.259261 | 0.236115 | 0.277071 | 0.345771 | 0.420849 | 0.457554 | 0.502893 | 0.523093 | 0.466659 | 0.344665 | 0.304910 | 0.286495 | 0.303364 | 0.277084 | 0.271233 | 0.275627 | 0.296070 | 0.326820 | 0.367185 | 0.389232 | 0.562562 | 0.629382 | 0.672347 | 0.612698 | 0.543912 | 0.418846 | 0.316108 | 0.256281 | 0.250587 | 0.244476 | 0.245749 | 0.227936 | 0.260076 | 0.308589 | 0.297160 | 0.259373 | 0.225496 | 0.214182 | 0.212113 | 0.213864 | 0.223585 | 0.220410 | 0.223603 | 0.212689 | 0.216034 | 0.250963 | 0.345098 | 0.439958 | 0.491811 | 0.585823 | 0.558990 | 0.510015 | 0.465758 | 0.371255 | 0.319201 | 0.324893 | 0.338705 | 0.352131 | 0.335201 | 0.283399 | 0.257582 |
| left posterior | 0.271902 | 0.286196 | 0.247153 | 0.223221 | 0.225285 | 0.226200 | 0.213961 | 0.210806 | 0.214206 | 0.223980 | 0.303798 | 0.294477 | 0.252350 | 0.298564 | 0.258644 | 0.290045 | 0.260112 | 0.276073 | 0.309776 | 0.572848 | 0.721641 | 1.192349 | 1.265767 | 1.400051 | 2.283235 | 6.211613 | 4.471433 | 4.435453 | 3.427401 | 3.505532 | 4.448579 | 2.601672 | 0.601357 | 0.296905 | 0.215094 | 0.215759 | 0.219697 | 0.211526 | 0.212260 | 0.242168 | 0.281403 | 0.430678 | 1.025766 | 1.068895 | 0.819795 | 0.456572 | 0.311626 | 0.263032 | 0.217213 | 0.219759 | 0.270253 | 0.300303 | 0.231734 | 0.211170 | 0.216323 | 0.217048 | 0.211626 | 0.260603 | 0.342487 | 0.873602 | 3.335696 | 39.798957 | 255.253822 | 1907.802874 | 6537.540897 | 6908.053779 | 5579.571853 | 2243.792580 | 648.445384 | 257.668869 | 80.594231 | 18.760314 | 4.591057 | 1.080364 | 0.474407 | 0.296826 | 0.298742 | 0.275269 | 0.280536 | 0.315704 | 0.314296 | 0.265747 | 0.240584 | 0.214347 | 0.211146 | 0.235116 | 0.260212 | 0.264469 | 0.234653 | 0.227805 | 0.234108 | 0.234857 | 0.217315 | 0.222068 | 0.214170 | 0.225994 | 0.233872 | 0.303620 | 0.573057 | 1.367247 | 4.353655 | 17.030815 | 55.354844 | 135.664478 | 338.368845 | 968.388188 | 6916.060479 | 41404.756389 | 342201.672576 | 298727.882018 | 108426.948897 | 115806.555052 | 68779.879544 | 11306.931144 | 3293.834135 | 1175.209876 | 352.155549 | 307.171589 | 94.573416 | 30.647487 | 6.334151 | 1.309729 | 0.916336 | 0.684705 | 0.401318 | 0.348927 | 0.319097 | 0.329286 | 0.341446 | 0.293122 | 0.250038 | 0.247155 | 0.235367 | 0.245246 | 0.242930 | 0.251946 | 0.232091 | 0.213657 | 0.212559 | 0.225635 | 0.238014 | 0.266526 | 0.291015 | 0.306804 | 0.366547 | 0.309529 | 0.248249 | 0.226680 | 0.214100 | 0.211314 | 0.212891 | 0.219124 | 0.230383 | 0.232893 | 0.247418 | 0.276308 | 0.273403 | 0.252425 | 0.229498 | 0.211542 | 0.229780 | 0.309106 | 0.819934 | 3.658570 | 9.120819 | 26.255118 | 32.279194 | 14.556533 | 7.321427 | 3.917411 | 2.782600 | 3.080051 | 4.232504 | 5.963897 | 11.557399 | 21.338319 | 29.478104 | 87.407669 | 218.809254 | 351.235690 | 398.009977 | 232.453755 | 196.429903 | 252.127138 | 273.278882 | 480.865944 | 1111.233343 | 3225.594486 | 2490.814286 | 667.622367 | 246.413098 | 73.330736 | 17.337981 | 4.252761 | 1.732574 | 1.151257 | 1.002290 | 0.876468 | 1.135129 | 1.874347 | 6.165194 | 11.606150 | 17.867304 | 23.668332 | 21.269954 | 20.454458 | 20.957416 | 15.209546 | 8.045539 | 3.671330 | 2.112597 | 2.201401 | 1.269458 | 0.916280 | 0.748621 | 0.899954 | 1.248380 | 1.303929 | 1.074811 | 1.279170 | 1.995804 | 2.848813 | 3.139615 | 6.165711 | 10.083505 | 24.120932 | 37.086725 | 59.560409 | 75.263554 | 82.330710 | 36.986088 | 27.511277 | 18.172182 | 18.114108 | 20.933923 | 33.222593 | 42.233677 | 59.034497 | 74.395806 | 109.550788 | 132.563791 | 74.589525 | 62.317071 | 42.899604 | 25.527953 | 22.618460 | 14.741041 | 10.390933 | 6.628093 | 3.643085 | 2.080490 | 1.271510 | 0.791292 | 0.802204 | 0.735790 | 0.820210 | 0.929596 | 1.343842 | 1.553688 | 1.501035 | 1.334562 | 1.118600 | 0.859917 | 0.700619 | 0.606583 | 0.703193 | 0.637565 | 0.507828 | 0.472174 | 0.629861 | 0.880882 | 1.105194 | 0.811175 | 0.680041 | 0.520347 | 0.439673 | 0.368696 | 0.304089 | 0.244347 | 0.232541 |
| right posterior | 1.135432 | 1.234571 | 1.390945 | 1.110230 | 1.083155 | 0.457094 | 0.267649 | 0.219415 | 0.213085 | 0.212081 | 0.220028 | 0.266121 | 0.430741 | 1.663544 | 12.849824 | 29.947694 | 25.694474 | 15.647016 | 8.117861 | 3.908007 | 0.592882 | 0.216147 | 0.211380 | 0.218149 | 0.211038 | 0.243334 | 0.349177 | 0.803807 | 1.833494 | 4.611165 | 17.722923 | 53.141263 | 33.101379 | 28.831213 | 39.052230 | 15.760836 | 9.026349 | 3.238988 | 1.017895 | 0.469156 | 0.311157 | 0.270922 | 0.273084 | 0.243178 | 0.216525 | 0.210821 | 0.214948 | 0.211228 | 0.212447 | 0.225314 | 0.254365 | 0.234649 | 0.240565 | 0.338113 | 0.532433 | 1.340642 | 2.942017 | 9.325905 | 92.174480 | 774.157268 | 2744.101623 | 6305.729588 | 10895.309727 | 32429.499548 | 55870.719761 | 51108.606464 | 54080.582880 | 55841.719846 | 87061.424975 | 72696.098336 | 27393.681332 | 4821.764362 | 477.526593 | 46.804190 | 3.896465 | 0.580972 | 0.247192 | 0.211794 | 0.217686 | 0.252180 | 0.298607 | 0.384879 | 0.482722 | 0.661312 | 0.732085 | 0.432863 | 0.292857 | 0.250574 | 0.234886 | 0.240821 | 0.237844 | 0.237066 | 0.270433 | 0.391709 | 0.550432 | 1.014894 | 1.811605 | 3.737941 | 6.744344 | 7.098609 | 3.907432 | 1.398548 | 0.521244 | 0.288075 | 0.227441 | 0.213130 | 0.211665 | 0.210956 | 0.214715 | 0.220445 | 0.231234 | 0.218666 | 0.216246 | 0.226921 | 0.240268 | 0.267845 | 0.334449 | 0.379017 | 0.559695 | 0.640022 | 0.492231 | 0.431794 | 0.384930 | 0.367909 | 0.406420 | 0.413883 | 0.386240 | 0.397132 | 0.461913 | 0.538660 | 0.529767 | 0.519250 | 0.507412 | 0.535903 | 0.411639 | 0.332464 | 0.266215 | 0.236340 | 0.225093 | 0.224839 | 0.230696 | 0.260363 | 0.283956 | 0.335090 | 0.332570 | 0.310529 | 0.266334 | 0.237944 | 0.223669 | 0.212373 | 0.211792 | 0.217798 | 0.236134 | 0.255884 | 0.261017 | 0.282603 | 0.257059 | 0.241458 | 0.245887 | 0.233683 | 0.234967 | 0.245571 | 0.269913 | 0.317581 | 0.342849 | 0.299660 | 0.293993 | 0.289597 | 0.272008 | 0.241363 | 0.238214 | 0.250602 | 0.284754 | 0.294543 | 0.286256 | 0.292603 | 0.266481 | 0.232752 | 0.223133 | 0.218359 | 0.214933 | 0.211286 | 0.210927 | 0.221134 | 0.250874 | 0.263674 | 0.252365 | 0.253989 | 0.277418 | 0.287472 | 0.246356 | 0.228457 | 0.239133 | 0.249313 | 0.256516 | 0.254765 | 0.256663 | 0.305204 | 0.327616 | 0.273008 | 0.262028 | 0.263299 | 0.247655 | 0.262767 | 0.249888 | 0.231433 | 0.220920 | 0.212620 | 0.212820 | 0.219125 | 0.242246 | 0.321124 | 0.372769 | 0.390000 | 0.369450 | 0.317028 | 0.301441 | 0.287702 | 0.247467 | 0.222539 | 0.212472 | 0.210920 | 0.210915 | 0.210833 | 0.211356 | 0.214284 | 0.211259 | 0.214520 | 0.219127 | 0.219377 | 0.218267 | 0.219108 | 0.219778 | 0.217920 | 0.211422 | 0.216671 | 0.221201 | 0.225487 | 0.216900 | 0.210915 | 0.211739 | 0.221841 | 0.221254 | 0.216550 | 0.211896 | 0.211243 | 0.217481 | 0.225175 | 0.230708 | 0.218150 | 0.212312 | 0.212055 | 0.215816 | 0.217268 | 0.213558 | 0.215641 | 0.213246 | 0.211990 | 0.212004 | 0.211031 | 0.210864 | 0.210813 | 0.212184 | 0.210814 | 0.210806 | 0.211164 | 0.211261 | 0.211243 | 0.215280 | 0.218740 | 0.222212 | 0.216049 | 0.214002 | 0.221486 | 0.224312 | 0.217831 | 0.218761 | 0.223576 | 0.237283 | 0.271204 |
| all electrodes | 0.211361 | 0.211035 | 0.219073 | 0.238008 | 0.418244 | 0.564155 | 0.564808 | 0.883204 | 1.931093 | 1.583882 | 0.691336 | 0.560240 | 0.815029 | 3.575292 | 3.943597 | 1.571622 | 0.980717 | 3.313093 | 6.444175 | 2.861960 | 0.547550 | 0.286643 | 0.257969 | 0.218239 | 0.228213 | 0.212178 | 0.286644 | 0.424699 | 0.821306 | 1.578619 | 4.175664 | 16.962872 | 13.247146 | 1.949306 | 0.943843 | 0.427280 | 0.324191 | 0.400670 | 0.636641 | 0.517807 | 0.513955 | 0.622001 | 0.793864 | 1.246058 | 0.984031 | 0.665545 | 0.761793 | 0.836326 | 0.883641 | 0.822908 | 0.436730 | 0.310443 | 0.233180 | 0.227151 | 0.366334 | 1.107340 | 2.148162 | 4.023234 | 11.264484 | 20.154184 | 46.653826 | 108.316643 | 256.025387 | 568.945403 | 610.691352 | 528.626671 | 440.336521 | 251.642564 | 222.208061 | 137.420149 | 45.890036 | 13.741535 | 1.875583 | 0.517580 | 0.235047 | 0.212530 | 0.258272 | 0.234339 | 0.216227 | 0.224740 | 0.324546 | 0.540790 | 0.655687 | 0.714121 | 0.631060 | 0.322314 | 0.217499 | 0.228103 | 0.293634 | 0.403232 | 0.504311 | 0.507264 | 0.350815 | 0.240518 | 0.223838 | 0.210914 | 0.211275 | 0.211415 | 0.239361 | 0.352453 | 0.530876 | 0.750470 | 1.283095 | 0.836105 | 0.568657 | 0.410241 | 0.408989 | 0.662187 | 0.924636 | 1.095724 | 1.618020 | 2.125127 | 1.466804 | 0.639322 | 0.299956 | 0.220825 | 0.212944 | 0.246411 | 0.441454 | 0.700393 | 0.461217 | 0.323385 | 0.292114 | 0.257795 | 0.234378 | 0.219099 | 0.225097 | 0.268873 | 0.306995 | 0.330220 | 0.414901 | 0.533379 | 0.496174 | 0.491624 | 0.405100 | 0.410803 | 0.352979 | 0.287985 | 0.287151 | 0.299080 | 0.244347 | 0.231687 | 0.234120 | 0.286168 | 0.470363 | 0.468362 | 0.491221 | 0.686800 | 0.947289 | 1.196883 | 1.025729 | 0.704669 | 0.560595 | 0.339007 | 0.287307 | 0.276286 | 0.258869 | 0.234041 | 0.211249 | 0.234577 | 0.222343 | 0.219553 | 0.214310 | 0.218498 | 0.213825 | 0.224170 | 0.316908 | 0.480724 | 0.592370 | 0.471873 | 0.396838 | 0.301670 | 0.243439 | 0.214047 | 0.224595 | 0.262124 | 0.267814 | 0.254234 | 0.240412 | 0.231859 | 0.215063 | 0.212495 | 0.211780 | 0.210929 | 0.210815 | 0.210843 | 0.216175 | 0.215806 | 0.217702 | 0.250340 | 0.320165 | 0.351652 | 0.421773 | 0.435583 | 0.479178 | 0.488616 | 0.480087 | 0.482523 | 0.620935 | 0.674388 | 0.669011 | 0.689330 | 0.778980 | 0.810455 | 0.655457 | 0.492225 | 0.444186 | 0.429701 | 0.524282 | 0.522653 | 0.512384 | 0.718182 | 0.913017 | 1.187319 | 1.247856 | 0.707940 | 0.466427 | 0.307918 | 0.256295 | 0.228295 | 0.211898 | 0.212381 | 0.217669 | 0.212415 | 0.221903 | 0.244576 | 0.260691 | 0.251835 | 0.255478 | 0.280458 | 0.291212 | 0.278379 | 0.287129 | 0.286314 | 0.327747 | 0.422524 | 0.531635 | 0.546336 | 0.498688 | 0.422494 | 0.431838 | 0.399067 | 0.340370 | 0.300435 | 0.274113 | 0.254003 | 0.257638 | 0.286989 | 0.356995 | 0.449288 | 0.579676 | 0.772305 | 1.030835 | 1.575542 | 1.601040 | 1.936720 | 2.261504 | 2.487847 | 2.329461 | 1.520068 | 0.755680 | 0.546158 | 0.442184 | 0.486246 | 0.645064 | 1.004170 | 1.103948 | 1.454105 | 1.670199 | 1.884389 | 2.147732 | 1.279702 | 1.028999 | 1.697888 | 2.095465 | 2.283410 | 1.621372 | 1.161174 | 1.360618 | 1.034900 |

Searchlight, spatiotemporal cluster permutation test

|  | start time | stop time | peak time | peak channel | cluster p | peak Cohen's d | direction |
| --- | --- | --- | --- | --- | --- | --- | --- |
| #1 | 55 | 320 | 140 | PO4 | 0.0013 | 1.488123 | positive |
| #2 | 295 | 1095 | 620 | FC4 | 0.0003 | -0.817317 | negative |
| #3 | 595 | 1170 | 700 | CP3 | 0.0013 | -0.86133 | negative |
| #4 | 840 | 1195 | 1050 | FC4 | 0.0447 | -1.141569 | negative |

E) emoji faces (LOSO) - angry vs sad

  
|  | time window | peak latency | cluster *p* | peak Cohen's *d* |  | | | |  |
| **all electrodes** | 105 - 170 ms | 135 ms | 0.0476 | 0.9389 |  | | | |  |
 535 - 700 ms | 545 ms | 0.0042 | -0.421 |  | | | ||  |  | | | | | | | | |

Time-resolved classification, cluster permutation tests

|  | **left hemisphere** | | | | **right hemisphere** | | | |  |
|  | time window | peak latency | cluster *p* | peak Cohen's *d* | time window | peak latency | cluster *p* | peak Cohen's *d* |  |
| **anterior** |  | | | |  | | | |  |
| **central** |  | | | |  | | | |  |
| **posterior** | 105 - 175 ms | 140 ms | 0.0218 | 1.2834 | 90 - 225 ms | 140 ms | 0.008 | 1.2281 |  |

  

Time-resolved classification, Bayesian statistics

|  | -200 | -195 | -190 | -185 | -180 | -175 | -170 | -165 | -160 | -155 | -150 | -145 | -140 | -135 | -130 | -125 | -120 | -115 | -110 | -105 | -100 | -95 | -90 | -85 | -80 | -75 | -70 | -65 | -60 | -55 | -50 | -45 | -40 | -35 | -30 | -25 | -20 | -15 | -10 | -5 | 0 | 5 | 10 | 15 | 20 | 25 | 30 | 35 | 40 | 45 | 50 | 55 | 60 | 65 | 70 | 75 | 80 | 85 | 90 | 95 | 100 | 105 | 110 | 115 | 120 | 125 | 130 | 135 | 140 | 145 | 150 | 155 | 160 | 165 | 170 | 175 | 180 | 185 | 190 | 195 | 200 | 205 | 210 | 215 | 220 | 225 | 230 | 235 | 240 | 245 | 250 | 255 | 260 | 265 | 270 | 275 | 280 | 285 | 290 | 295 | 300 | 305 | 310 | 315 | 320 | 325 | 330 | 335 | 340 | 345 | 350 | 355 | 360 | 365 | 370 | 375 | 380 | 385 | 390 | 395 | 400 | 405 | 410 | 415 | 420 | 425 | 430 | 435 | 440 | 445 | 450 | 455 | 460 | 465 | 470 | 475 | 480 | 485 | 490 | 495 | 500 | 505 | 510 | 515 | 520 | 525 | 530 | 535 | 540 | 545 | 550 | 555 | 560 | 565 | 570 | 575 | 580 | 585 | 590 | 595 | 600 | 605 | 610 | 615 | 620 | 625 | 630 | 635 | 640 | 645 | 650 | 655 | 660 | 665 | 670 | 675 | 680 | 685 | 690 | 695 | 700 | 705 | 710 | 715 | 720 | 725 | 730 | 735 | 740 | 745 | 750 | 755 | 760 | 765 | 770 | 775 | 780 | 785 | 790 | 795 | 800 | 805 | 810 | 815 | 820 | 825 | 830 | 835 | 840 | 845 | 850 | 855 | 860 | 865 | 870 | 875 | 880 | 885 | 890 | 895 | 900 | 905 | 910 | 915 | 920 | 925 | 930 | 935 | 940 | 945 | 950 | 955 | 960 | 965 | 970 | 975 | 980 | 985 | 990 | 995 | 1000 | 1005 | 1010 | 1015 | 1020 | 1025 | 1030 | 1035 | 1040 | 1045 | 1050 | 1055 | 1060 | 1065 | 1070 | 1075 | 1080 | 1085 | 1090 | 1095 | 1100 | 1105 | 1110 | 1115 | 1120 | 1125 | 1130 | 1135 | 1140 | 1145 | 1150 | 1155 | 1160 | 1165 | 1170 | 1175 | 1180 | 1185 | 1190 | 1195 |
| --- | --- | --- | --- | --- | --- | --- | --- | --- | --- | --- | --- | --- | --- | --- | --- | --- | --- | --- | --- | --- | --- | --- | --- | --- | --- | --- | --- | --- | --- | --- | --- | --- | --- | --- | --- | --- | --- | --- | --- | --- | --- | --- | --- | --- | --- | --- | --- | --- | --- | --- | --- | --- | --- | --- | --- | --- | --- | --- | --- | --- | --- | --- | --- | --- | --- | --- | --- | --- | --- | --- | --- | --- | --- | --- | --- | --- | --- | --- | --- | --- | --- | --- | --- | --- | --- | --- | --- | --- | --- | --- | --- | --- | --- | --- | --- | --- | --- | --- | --- | --- | --- | --- | --- | --- | --- | --- | --- | --- | --- | --- | --- | --- | --- | --- | --- | --- | --- | --- | --- | --- | --- | --- | --- | --- | --- | --- | --- | --- | --- | --- | --- | --- | --- | --- | --- | --- | --- | --- | --- | --- | --- | --- | --- | --- | --- | --- | --- | --- | --- | --- | --- | --- | --- | --- | --- | --- | --- | --- | --- | --- | --- | --- | --- | --- | --- | --- | --- | --- | --- | --- | --- | --- | --- | --- | --- | --- | --- | --- | --- | --- | --- | --- | --- | --- | --- | --- | --- | --- | --- | --- | --- | --- | --- | --- | --- | --- | --- | --- | --- | --- | --- | --- | --- | --- | --- | --- | --- | --- | --- | --- | --- | --- | --- | --- | --- | --- | --- | --- | --- | --- | --- | --- | --- | --- | --- | --- | --- | --- | --- | --- | --- | --- | --- | --- | --- | --- | --- | --- | --- | --- | --- | --- | --- | --- | --- | --- | --- | --- | --- | --- | --- | --- | --- | --- | --- | --- | --- | --- | --- | --- | --- | --- | --- | --- | --- | --- | --- | --- | --- | --- | --- | --- | --- | --- | --- | --- | --- | --- | --- | --- |
| left anterior | 0.527191 | 0.848714 | 1.260059 | 1.318361 | 0.866612 | 1.261019 | 2.257116 | 2.861908 | 0.982356 | 0.452212 | 0.259587 | 0.232874 | 0.227317 | 0.392528 | 0.674889 | 1.017425 | 0.701568 | 0.636857 | 1.993507 | 1.002539 | 0.289415 | 0.229505 | 0.448577 | 0.590959 | 0.981240 | 5.726719 | 11.249053 | 16.084801 | 4.719919 | 2.887423 | 0.803127 | 0.463936 | 0.359336 | 0.301189 | 0.282471 | 0.311885 | 0.371563 | 0.704226 | 1.127887 | 0.881789 | 0.667951 | 0.373438 | 0.239051 | 0.231626 | 0.531048 | 1.406710 | 0.823433 | 0.840753 | 0.882487 | 0.716285 | 0.411457 | 0.316133 | 0.273729 | 0.257041 | 0.223653 | 0.211095 | 0.212373 | 0.230768 | 0.334918 | 0.385661 | 0.362291 | 0.341988 | 0.498552 | 0.779868 | 0.655263 | 0.389779 | 0.450745 | 0.549906 | 0.687191 | 0.437892 | 0.389177 | 0.405623 | 0.393407 | 0.296185 | 0.235221 | 0.227039 | 0.239371 | 0.274728 | 0.329757 | 0.421067 | 0.467036 | 0.454548 | 0.421489 | 0.282187 | 0.216789 | 0.213532 | 0.222165 | 0.239409 | 0.228293 | 0.233880 | 0.215905 | 0.361971 | 0.836145 | 1.152668 | 1.556881 | 1.208782 | 0.848285 | 0.501689 | 0.244625 | 0.210862 | 0.217410 | 0.245017 | 0.274096 | 0.410998 | 0.688252 | 0.865446 | 1.084148 | 1.122268 | 0.687644 | 0.446551 | 0.245552 | 0.213332 | 0.234208 | 0.261837 | 0.248914 | 0.243721 | 0.259195 | 0.291864 | 0.383925 | 0.317327 | 0.273771 | 0.268229 | 0.262686 | 0.265834 | 0.245302 | 0.230359 | 0.219349 | 0.212787 | 0.261474 | 0.417227 | 1.127718 | 4.078500 | 14.904428 | 20.648477 | 10.137669 | 7.377237 | 12.049156 | 19.641803 | 16.290135 | 22.648063 | 30.682596 | 24.553681 | 10.892999 | 7.167858 | 3.731707 | 2.352059 | 1.341808 | 0.908086 | 1.217635 | 2.481933 | 3.699689 | 8.531619 | 12.669110 | 11.546222 | 9.610526 | 4.122728 | 1.603766 | 0.934183 | 0.574082 | 0.631252 | 1.046801 | 1.723283 | 1.943465 | 1.927104 | 1.764044 | 1.242375 | 1.115908 | 1.310133 | 1.585057 | 1.448404 | 1.464061 | 0.737376 | 0.663933 | 0.602281 | 0.432072 | 0.352203 | 0.343296 | 0.303939 | 0.455783 | 0.530316 | 0.459482 | 0.436332 | 0.367002 | 0.313398 | 0.309363 | 0.265991 | 0.256461 | 0.257264 | 0.265867 | 0.270546 | 0.285960 | 0.259385 | 0.238602 | 0.226913 | 0.219756 | 0.210874 | 0.212058 | 0.216986 | 0.229964 | 0.224142 | 0.220679 | 0.215885 | 0.210884 | 0.212405 | 0.223887 | 0.270295 | 0.292033 | 0.274538 | 0.253010 | 0.218851 | 0.211004 | 0.212446 | 0.219646 | 0.227995 | 0.230186 | 0.223185 | 0.219965 | 0.220041 | 0.227800 | 0.225708 | 0.239391 | 0.219220 | 0.212146 | 0.218352 | 0.258746 | 0.434946 | 0.738652 | 1.766427 | 1.587308 | 1.293388 | 2.075159 | 4.124648 | 6.015079 | 5.033319 | 2.962898 | 2.652584 | 3.243953 | 1.883534 | 0.900732 | 0.496498 | 0.419707 | 0.584819 | 0.868961 | 1.090146 | 1.196593 | 1.319585 | 1.522771 | 1.227740 | 0.824227 | 0.464713 | 0.288946 | 0.233323 | 0.220113 | 0.213251 | 0.212664 | 0.211754 | 0.219081 | 0.222790 | 0.222735 | 0.218845 | 0.210818 | 0.213182 | 0.230297 | 0.278761 | 0.332329 | 0.469669 | 0.726634 | 1.031915 | 1.121817 | 0.823095 | 1.222150 | 2.626294 | 4.367100 | 2.027917 | 0.705826 | 0.585801 | 0.790957 | 0.606012 | 0.355448 | 0.240575 |
| right anterior | 1.772208 | 1.331831 | 1.589214 | 2.160677 | 1.542211 | 0.593326 | 0.375322 | 0.283390 | 0.257585 | 0.211503 | 0.228606 | 0.329866 | 0.620634 | 0.563132 | 0.531231 | 0.464102 | 0.291247 | 0.279411 | 0.238842 | 0.219668 | 0.217699 | 0.219771 | 0.222392 | 0.213372 | 0.212096 | 0.245001 | 0.550140 | 0.602797 | 0.544858 | 0.515118 | 0.623903 | 0.923292 | 0.581021 | 0.223200 | 0.223760 | 0.270019 | 0.276719 | 0.222342 | 0.211208 | 0.210876 | 0.220356 | 0.210825 | 0.228498 | 0.225194 | 0.226060 | 0.257933 | 0.315506 | 0.349494 | 0.377366 | 0.308552 | 0.248760 | 0.217210 | 0.211488 | 0.225151 | 0.233224 | 0.272748 | 0.261606 | 0.216148 | 0.210820 | 0.211238 | 0.217696 | 0.241820 | 0.286041 | 0.444735 | 0.783777 | 1.335552 | 2.501538 | 2.134594 | 1.914564 | 1.554772 | 0.856408 | 0.691778 | 0.450487 | 0.379581 | 0.406383 | 0.374904 | 0.317068 | 0.242440 | 0.212226 | 0.234854 | 0.312870 | 0.486098 | 0.825974 | 0.781365 | 0.606076 | 0.473932 | 0.369455 | 0.317747 | 0.270757 | 0.217383 | 0.212412 | 0.277095 | 0.365260 | 0.456072 | 0.710544 | 1.016279 | 0.906531 | 0.524967 | 0.322958 | 0.281748 | 0.258521 | 0.223432 | 0.212990 | 0.211782 | 0.244168 | 0.255330 | 0.259673 | 0.239812 | 0.268793 | 0.279736 | 0.291988 | 0.271750 | 0.272032 | 0.288095 | 0.306545 | 0.303484 | 0.311921 | 0.322581 | 0.343423 | 0.397395 | 0.382346 | 0.460334 | 0.747460 | 1.172175 | 1.947956 | 2.685686 | 2.080906 | 1.537355 | 0.820852 | 0.406228 | 0.262632 | 0.215216 | 0.210925 | 0.219550 | 0.241400 | 0.272559 | 0.392299 | 0.488495 | 0.625275 | 0.988711 | 1.657528 | 2.552881 | 1.394976 | 0.930080 | 0.884580 | 1.690592 | 3.259701 | 3.555029 | 1.612610 | 1.277447 | 1.415537 | 1.601172 | 0.887912 | 0.637913 | 0.577741 | 1.155990 | 2.737678 | 3.065098 | 3.254046 | 3.589704 | 3.655028 | 5.482389 | 4.769179 | 2.779771 | 1.679436 | 1.073784 | 1.002504 | 1.047859 | 1.008409 | 0.649737 | 0.524797 | 0.547695 | 0.494478 | 0.415542 | 0.399997 | 0.468653 | 0.491606 | 0.445882 | 0.352072 | 0.297414 | 0.268664 | 0.215193 | 0.222876 | 0.227620 | 0.214193 | 0.213762 | 0.228306 | 0.268703 | 0.368431 | 0.656445 | 1.017734 | 1.666243 | 1.330642 | 0.862798 | 0.592221 | 0.441035 | 0.288787 | 0.236530 | 0.211469 | 0.210817 | 0.213706 | 0.217405 | 0.219090 | 0.240594 | 0.245184 | 0.264311 | 0.272418 | 0.299117 | 0.289776 | 0.278439 | 0.299263 | 0.364068 | 0.414311 | 0.513407 | 0.557508 | 0.573898 | 0.564865 | 0.424068 | 0.333541 | 0.331889 | 0.319166 | 0.283365 | 0.347055 | 0.429972 | 0.410973 | 0.439263 | 0.412977 | 0.379740 | 0.478468 | 0.469672 | 0.447572 | 0.504663 | 0.581362 | 0.656871 | 0.675428 | 0.401495 | 0.302647 | 0.248463 | 0.218870 | 0.212774 | 0.210806 | 0.210876 | 0.214090 | 0.229157 | 0.254254 | 0.271563 | 0.236551 | 0.215577 | 0.211945 | 0.242696 | 0.322620 | 0.508318 | 0.838812 | 1.271242 | 1.404456 | 1.727948 | 1.553346 | 2.012449 | 1.803505 | 1.074021 | 0.713888 | 0.495115 | 0.418500 | 0.455872 | 0.398616 | 0.312320 | 0.256257 | 0.221181 | 0.211330 | 0.211169 | 0.216948 | 0.224898 | 0.223667 | 0.220514 | 0.213376 | 0.211308 | 0.213988 | 0.216218 | 0.220383 | 0.226209 |
| left central | 1.086079 | 0.861781 | 0.470382 | 0.359705 | 0.284144 | 0.237000 | 0.232886 | 0.240871 | 0.239777 | 0.308963 | 0.282490 | 0.315688 | 0.331871 | 0.267866 | 0.217821 | 0.215978 | 0.211443 | 0.232475 | 0.264264 | 0.267896 | 0.313109 | 0.522227 | 0.693392 | 0.503942 | 0.297504 | 0.212471 | 0.212012 | 0.278960 | 0.374219 | 0.362780 | 0.337624 | 0.294631 | 0.230588 | 0.235233 | 0.214660 | 0.254080 | 0.256295 | 0.228768 | 0.211612 | 0.213141 | 0.219091 | 0.231869 | 0.389373 | 0.676084 | 0.747024 | 0.522537 | 0.400742 | 0.339526 | 0.242123 | 0.210980 | 0.231396 | 0.265417 | 0.327427 | 0.439905 | 0.591116 | 0.569825 | 0.449267 | 0.536518 | 0.679096 | 0.878255 | 0.677654 | 0.651943 | 0.884614 | 1.438542 | 1.541063 | 1.412290 | 1.179596 | 1.226918 | 0.875597 | 0.628809 | 0.500560 | 0.386495 | 0.260170 | 0.235544 | 0.218626 | 0.238382 | 0.294737 | 0.444548 | 1.005626 | 2.556211 | 3.882974 | 4.145963 | 2.660274 | 1.971279 | 1.512834 | 0.739033 | 0.446291 | 0.288744 | 0.284569 | 0.271578 | 0.237274 | 0.227053 | 0.236405 | 0.267166 | 0.369748 | 0.370322 | 0.489794 | 0.549766 | 0.330264 | 0.244861 | 0.221612 | 0.216345 | 0.216380 | 0.211549 | 0.211449 | 0.217798 | 0.257990 | 0.287484 | 0.301694 | 0.325385 | 0.316565 | 0.399108 | 0.575066 | 0.750720 | 0.952630 | 1.157078 | 1.640678 | 1.952319 | 1.331318 | 1.394542 | 0.819156 | 0.590519 | 0.409547 | 0.334004 | 0.306108 | 0.294677 | 0.280412 | 0.411711 | 0.566239 | 0.542811 | 0.503792 | 0.616084 | 1.287250 | 2.283737 | 2.506911 | 1.382378 | 1.095141 | 0.608196 | 0.500544 | 0.368284 | 0.354734 | 0.358458 | 0.356155 | 0.348237 | 0.352462 | 0.315161 | 0.271402 | 0.242414 | 0.241943 | 0.232572 | 0.213941 | 0.216862 | 0.273338 | 0.351704 | 0.426138 | 0.445890 | 0.428716 | 0.348406 | 0.251344 | 0.218627 | 0.213914 | 0.221577 | 0.254956 | 0.307358 | 0.450531 | 0.784479 | 1.516927 | 2.172788 | 2.718471 | 3.350715 | 2.990361 | 1.981870 | 1.120494 | 0.647016 | 0.405148 | 0.282196 | 0.219161 | 0.210923 | 0.243206 | 0.318558 | 0.419655 | 0.351075 | 0.345005 | 0.349960 | 0.327410 | 0.267408 | 0.253053 | 0.249667 | 0.255150 | 0.242915 | 0.234820 | 0.228167 | 0.221111 | 0.233326 | 0.232003 | 0.243809 | 0.258051 | 0.264371 | 0.285188 | 0.322436 | 0.297261 | 0.275895 | 0.240465 | 0.216941 | 0.213071 | 0.211132 | 0.211660 | 0.230791 | 0.235659 | 0.237419 | 0.216192 | 0.210900 | 0.217739 | 0.246154 | 0.345505 | 0.499157 | 0.738217 | 0.790774 | 0.748015 | 0.708909 | 0.513116 | 0.386200 | 0.324203 | 0.307969 | 0.346333 | 0.452531 | 0.422225 | 0.507459 | 0.575839 | 0.579730 | 0.527211 | 0.489319 | 0.392271 | 0.372878 | 0.288115 | 0.237209 | 0.218576 | 0.214636 | 0.212246 | 0.217717 | 0.222459 | 0.236953 | 0.274518 | 0.380486 | 0.640856 | 0.767857 | 0.837594 | 0.925705 | 1.041898 | 1.996294 | 3.250475 | 3.944378 | 5.591439 | 6.338808 | 8.867257 | 10.638841 | 8.630122 | 6.458848 | 4.644999 | 2.867081 | 1.772447 | 1.408379 | 1.110927 | 0.813977 | 0.543665 | 0.412829 | 0.372165 | 0.367044 | 0.320564 | 0.311689 | 0.279755 | 0.290521 | 0.308613 | 0.331496 | 0.354927 | 0.411322 | 0.464948 | 0.575458 | 0.613105 | 0.616663 |
| right central | 0.232681 | 0.211065 | 0.217380 | 0.224842 | 0.226156 | 0.226226 | 0.226417 | 0.249112 | 0.247111 | 0.211140 | 0.241394 | 0.304312 | 0.313245 | 0.230166 | 0.314232 | 0.527711 | 0.231738 | 0.234284 | 0.298296 | 0.463542 | 0.520064 | 1.297465 | 2.178851 | 0.861781 | 0.323474 | 0.252051 | 0.220406 | 0.214927 | 0.219348 | 0.224183 | 0.221932 | 0.483955 | 0.362882 | 0.257710 | 0.215686 | 0.247728 | 0.299542 | 0.265797 | 0.218425 | 0.220165 | 0.275723 | 0.542441 | 0.874166 | 0.552066 | 0.221217 | 0.279310 | 0.523316 | 0.357744 | 0.233806 | 0.233947 | 0.320070 | 0.916081 | 2.132770 | 3.051153 | 1.781540 | 0.681189 | 0.297326 | 0.230340 | 0.210848 | 0.212796 | 0.210988 | 0.256093 | 0.517549 | 1.022232 | 1.581949 | 2.017439 | 2.701794 | 1.569352 | 0.837350 | 0.305822 | 0.210806 | 0.257299 | 0.279165 | 0.295603 | 0.273928 | 0.301076 | 0.239910 | 0.212122 | 0.217823 | 0.229882 | 0.240458 | 0.221217 | 0.210856 | 0.238994 | 0.242266 | 0.223121 | 0.222864 | 0.223752 | 0.210889 | 0.241143 | 0.425578 | 0.597862 | 0.781222 | 0.996333 | 1.154610 | 1.199330 | 0.731568 | 0.486464 | 0.279289 | 0.213957 | 0.319512 | 0.571067 | 1.680957 | 4.791240 | 7.817375 | 4.190329 | 1.815128 | 0.932892 | 0.608978 | 0.452493 | 0.332046 | 0.328088 | 0.291281 | 0.284863 | 0.228969 | 0.210819 | 0.297461 | 0.547279 | 1.161589 | 1.517385 | 1.028551 | 0.553017 | 0.377174 | 0.252542 | 0.221614 | 0.210806 | 0.214072 | 0.215738 | 0.212625 | 0.214010 | 0.227044 | 0.256257 | 0.266672 | 0.280124 | 0.351709 | 0.291720 | 0.239737 | 0.217654 | 0.210863 | 0.210806 | 0.215124 | 0.221952 | 0.244637 | 0.278370 | 0.344277 | 0.444752 | 0.514877 | 0.503094 | 0.460487 | 0.481582 | 0.571848 | 0.709024 | 0.787674 | 0.677812 | 0.624438 | 0.655705 | 0.628506 | 0.595909 | 0.559253 | 0.640786 | 0.762029 | 0.766277 | 0.636543 | 0.582135 | 0.661393 | 0.888640 | 1.169656 | 1.885447 | 2.060493 | 1.732784 | 1.489374 | 1.350100 | 0.964834 | 0.584725 | 0.402519 | 0.418332 | 0.584498 | 0.977115 | 1.074338 | 0.825230 | 0.487063 | 0.300098 | 0.229477 | 0.213932 | 0.210814 | 0.211857 | 0.221709 | 0.278683 | 0.410999 | 0.581342 | 0.535199 | 0.375782 | 0.271821 | 0.247687 | 0.232840 | 0.228674 | 0.221834 | 0.221578 | 0.224064 | 0.219045 | 0.212569 | 0.211042 | 0.214887 | 0.222420 | 0.238922 | 0.220810 | 0.212014 | 0.211220 | 0.211007 | 0.211502 | 0.212911 | 0.215897 | 0.211478 | 0.211454 | 0.211134 | 0.211721 | 0.212921 | 0.221586 | 0.221538 | 0.220818 | 0.210929 | 0.212217 | 0.217453 | 0.242818 | 0.293788 | 0.314400 | 0.329948 | 0.279874 | 0.234431 | 0.222730 | 0.216360 | 0.228140 | 0.243723 | 0.261261 | 0.265724 | 0.291764 | 0.275181 | 0.237325 | 0.214876 | 0.211717 | 0.210844 | 0.210894 | 0.213861 | 0.227732 | 0.254136 | 0.258174 | 0.244656 | 0.243131 | 0.237613 | 0.222033 | 0.214387 | 0.211026 | 0.211499 | 0.210996 | 0.210915 | 0.211723 | 0.211697 | 0.210832 | 0.221284 | 0.286038 | 0.461843 | 0.755268 | 1.089640 | 1.428280 | 1.366544 | 0.836980 | 0.528057 | 0.366936 | 0.293408 | 0.293842 | 0.321236 | 0.406452 | 0.625370 | 0.649434 | 0.534835 | 0.469109 | 0.464671 | 0.512636 | 0.479207 | 0.355347 |
| left posterior | 0.327598 | 0.223806 | 0.218474 | 0.249876 | 0.218771 | 0.246396 | 0.335090 | 0.458317 | 0.416087 | 0.334323 | 0.304101 | 0.518938 | 0.325965 | 0.243196 | 0.216128 | 0.212079 | 0.233292 | 0.307683 | 0.597437 | 0.644827 | 0.619372 | 0.393516 | 0.407133 | 0.741929 | 1.015566 | 0.560583 | 0.406662 | 0.536969 | 0.547586 | 0.305863 | 0.228771 | 0.211735 | 0.220819 | 0.235413 | 0.210832 | 0.215377 | 0.210806 | 0.249217 | 0.312682 | 0.398149 | 0.518637 | 0.444750 | 0.269889 | 0.222180 | 0.215606 | 0.238577 | 0.284025 | 0.252274 | 0.211336 | 0.242489 | 0.257170 | 0.239963 | 0.224604 | 0.213287 | 0.213405 | 0.295339 | 0.534926 | 0.332513 | 0.216321 | 0.233638 | 0.476638 | 2.903964 | 32.052687 | 188.501181 | 443.990521 | 681.696056 | 1878.847479 | 6922.881513 | 14731.719130 | 27048.138833 | 22255.471296 | 16141.008656 | 3625.250082 | 234.581669 | 20.755341 | 4.155019 | 1.265208 | 0.497033 | 0.321264 | 0.300212 | 0.285623 | 0.326140 | 0.369707 | 0.376685 | 0.381199 | 0.349743 | 0.334937 | 0.358259 | 0.383328 | 0.431039 | 0.905892 | 3.890341 | 18.515957 | 105.955679 | 558.926324 | 588.648509 | 510.578101 | 184.796126 | 54.377310 | 28.375088 | 8.727144 | 3.126581 | 1.813953 | 1.192393 | 1.121078 | 0.925422 | 0.540555 | 0.338121 | 0.291171 | 0.253284 | 0.247261 | 0.236442 | 0.243450 | 0.334410 | 0.585282 | 0.820789 | 1.006451 | 1.185677 | 1.726711 | 2.033178 | 1.640212 | 1.258369 | 0.889992 | 0.770219 | 0.892773 | 0.674169 | 0.646153 | 0.702682 | 1.067846 | 2.113457 | 3.463305 | 3.580207 | 3.224564 | 2.354347 | 1.838346 | 1.374000 | 0.918842 | 0.749325 | 0.813398 | 0.933397 | 1.135766 | 1.468041 | 1.111530 | 1.297229 | 1.438369 | 0.770330 | 0.461211 | 0.287569 | 0.227161 | 0.236108 | 0.220519 | 0.210891 | 0.211634 | 0.210847 | 0.211169 | 0.211465 | 0.240219 | 0.248671 | 0.227243 | 0.218318 | 0.223936 | 0.223013 | 0.221160 | 0.218171 | 0.222556 | 0.230807 | 0.248165 | 0.231002 | 0.234635 | 0.231816 | 0.236488 | 0.268999 | 0.326660 | 0.394919 | 0.536674 | 0.507212 | 0.485396 | 0.445902 | 0.301833 | 0.230933 | 0.211874 | 0.210895 | 0.210845 | 0.213003 | 0.236513 | 0.350983 | 0.724210 | 1.441464 | 2.266510 | 3.447530 | 6.283477 | 5.240583 | 4.473451 | 3.774422 | 4.107915 | 2.509497 | 1.373392 | 0.619028 | 0.530896 | 0.573869 | 0.511982 | 0.386565 | 0.339654 | 0.278896 | 0.274782 | 0.279294 | 0.252234 | 0.246682 | 0.285781 | 0.288217 | 0.352387 | 0.448234 | 0.418137 | 0.367736 | 0.325738 | 0.276329 | 0.301172 | 0.259071 | 0.240960 | 0.248329 | 0.277758 | 0.317779 | 0.338638 | 0.357032 | 0.434185 | 0.403487 | 0.313242 | 0.279782 | 0.271152 | 0.279418 | 0.322004 | 0.385886 | 0.450904 | 0.533856 | 0.557310 | 0.479508 | 0.454343 | 0.402924 | 0.368861 | 0.361078 | 0.309705 | 0.288957 | 0.286752 | 0.243973 | 0.217136 | 0.210815 | 0.217445 | 0.216159 | 0.222048 | 0.238268 | 0.277740 | 0.294892 | 0.349464 | 0.398791 | 0.546239 | 0.901220 | 1.495216 | 1.614007 | 1.661119 | 1.506392 | 1.423032 | 1.002095 | 0.733959 | 0.554555 | 0.606125 | 0.669149 | 0.613264 | 0.543376 | 0.592299 | 0.588707 | 0.580965 | 0.481132 | 0.413060 | 0.409432 | 0.394540 | 0.398284 | 0.373032 | 0.363235 | 0.369946 | 0.430370 |
| right posterior | 0.250041 | 0.281871 | 0.233001 | 0.229476 | 0.239769 | 0.219975 | 0.212256 | 0.220378 | 0.215211 | 0.217557 | 0.210861 | 0.221645 | 0.223343 | 0.225146 | 0.262462 | 0.249027 | 0.218407 | 0.214222 | 0.254067 | 0.279730 | 0.255745 | 0.229158 | 0.232718 | 0.233434 | 0.229396 | 0.304769 | 0.459426 | 0.995783 | 2.499213 | 1.530569 | 0.697459 | 0.496780 | 0.294415 | 0.268309 | 0.221950 | 0.215857 | 0.214229 | 0.223737 | 0.260420 | 0.306891 | 0.403216 | 0.407613 | 0.459997 | 0.471414 | 0.306720 | 0.294008 | 0.316008 | 0.333707 | 0.339475 | 0.289339 | 0.264134 | 0.235503 | 0.210806 | 0.211504 | 0.214840 | 0.232880 | 0.310414 | 0.718024 | 2.596984 | 26.566453 | 127.042361 | 236.370232 | 877.276572 | 2049.404643 | 2179.506857 | 1939.108317 | 1761.371442 | 2398.243563 | 7942.288921 | 4597.412146 | 2732.721885 | 1982.593470 | 1343.493442 | 732.650666 | 170.694279 | 33.374322 | 19.202582 | 21.820717 | 29.836315 | 30.105028 | 26.639688 | 21.987688 | 19.288446 | 9.303708 | 3.841292 | 1.320484 | 0.687935 | 0.415505 | 0.357140 | 0.417883 | 0.580351 | 0.969669 | 1.850838 | 2.278379 | 2.546377 | 2.711574 | 1.804443 | 1.001644 | 0.704201 | 0.633056 | 0.768234 | 0.615561 | 0.363303 | 0.263745 | 0.228175 | 0.212668 | 0.216606 | 0.311122 | 0.404510 | 0.430250 | 0.430709 | 0.370528 | 0.339404 | 0.273909 | 0.211236 | 0.271035 | 0.457672 | 0.889570 | 1.245410 | 1.875663 | 2.236439 | 1.233070 | 0.696916 | 0.509653 | 0.396576 | 0.349213 | 0.321166 | 0.308309 | 0.324954 | 0.276669 | 0.289357 | 0.314553 | 0.373980 | 0.420428 | 0.459266 | 0.414583 | 0.422188 | 0.335064 | 0.284056 | 0.236508 | 0.211788 | 0.212857 | 0.221585 | 0.238559 | 0.241162 | 0.259951 | 0.261301 | 0.273421 | 0.291868 | 0.282273 | 0.275486 | 0.246977 | 0.228893 | 0.218980 | 0.212022 | 0.216206 | 0.214558 | 0.217671 | 0.210982 | 0.216022 | 0.238992 | 0.278581 | 0.272639 | 0.255217 | 0.234058 | 0.212896 | 0.211923 | 0.236435 | 0.269536 | 0.279824 | 0.267939 | 0.232933 | 0.220690 | 0.210861 | 0.234980 | 0.255740 | 0.283679 | 0.283308 | 0.256958 | 0.241773 | 0.223205 | 0.210857 | 0.211683 | 0.211132 | 0.212181 | 0.222083 | 0.248352 | 0.246874 | 0.238318 | 0.233345 | 0.219214 | 0.211726 | 0.213046 | 0.222263 | 0.223521 | 0.214759 | 0.216437 | 0.219151 | 0.218982 | 0.214769 | 0.210903 | 0.211576 | 0.222762 | 0.227311 | 0.218574 | 0.212438 | 0.210863 | 0.214837 | 0.237683 | 0.286713 | 0.342794 | 0.331840 | 0.289435 | 0.265192 | 0.233472 | 0.219466 | 0.213523 | 0.215843 | 0.224541 | 0.263250 | 0.351385 | 0.484140 | 0.646116 | 0.630239 | 0.570454 | 0.466934 | 0.474394 | 0.392864 | 0.344121 | 0.333778 | 0.346015 | 0.328178 | 0.346907 | 0.294061 | 0.283963 | 0.269068 | 0.255405 | 0.269076 | 0.345355 | 0.417283 | 0.537781 | 0.643969 | 0.777074 | 0.718072 | 0.581629 | 0.392262 | 0.305226 | 0.256577 | 0.230869 | 0.212521 | 0.210857 | 0.211006 | 0.213129 | 0.214319 | 0.218712 | 0.227451 | 0.226709 | 0.241159 | 0.257914 | 0.255651 | 0.278056 | 0.314380 | 0.324934 | 0.314327 | 0.285698 | 0.260428 | 0.250368 | 0.221216 | 0.211306 | 0.210806 | 0.210910 | 0.210974 | 0.210920 | 0.212295 | 0.219714 | 0.225507 | 0.234142 | 0.237716 | 0.243858 | 0.248516 |
| all electrodes | 0.235699 | 0.217834 | 0.264241 | 0.375264 | 0.327880 | 0.841001 | 5.964109 | 6.904817 | 2.319292 | 0.353340 | 0.245788 | 0.213860 | 0.336634 | 1.802880 | 1.006592 | 0.265926 | 0.223779 | 0.246796 | 0.266061 | 0.349504 | 0.565156 | 0.482841 | 0.260798 | 0.232432 | 0.277366 | 0.218971 | 0.239098 | 0.252436 | 0.232415 | 0.239327 | 0.337023 | 0.428189 | 0.361193 | 0.224457 | 0.211498 | 0.212156 | 0.223240 | 0.226292 | 0.212063 | 0.213137 | 0.217454 | 0.217656 | 0.211524 | 0.214575 | 0.289289 | 0.328698 | 0.407526 | 0.253891 | 0.214223 | 0.271461 | 0.404810 | 0.892474 | 0.661542 | 0.614556 | 0.572474 | 0.410260 | 0.389967 | 0.302939 | 0.254336 | 0.483290 | 0.952933 | 3.435748 | 24.267251 | 100.314583 | 269.982050 | 364.019083 | 259.125054 | 294.620466 | 64.796594 | 13.924097 | 4.969009 | 4.569689 | 5.213686 | 4.389884 | 1.330523 | 0.756163 | 0.736731 | 1.052006 | 1.305697 | 1.654754 | 1.949072 | 2.713960 | 3.492812 | 3.041686 | 2.212855 | 1.061785 | 0.345263 | 0.221027 | 0.213801 | 0.211524 | 0.226348 | 0.317337 | 0.605221 | 3.260818 | 30.421443 | 178.571110 | 30.593503 | 2.459699 | 0.851050 | 0.321229 | 0.221744 | 0.211342 | 0.217854 | 0.256502 | 0.314250 | 0.431265 | 0.382383 | 0.400894 | 0.343870 | 0.298215 | 0.270061 | 0.240245 | 0.246584 | 0.281916 | 0.329437 | 0.318856 | 0.251901 | 0.224313 | 0.233872 | 0.239917 | 0.232644 | 0.213192 | 0.217304 | 0.224660 | 0.233286 | 0.223918 | 0.213912 | 0.212608 | 0.232290 | 0.232062 | 0.214971 | 0.232460 | 0.295619 | 0.339413 | 0.310977 | 0.213761 | 0.222566 | 0.221048 | 0.242786 | 0.270283 | 0.281337 | 0.310173 | 0.312805 | 0.309435 | 0.331971 | 0.474023 | 0.635195 | 1.530032 | 2.250355 | 1.369326 | 1.837345 | 3.769872 | 5.061799 | 5.567518 | 3.230613 | 3.170433 | 3.255906 | 2.136693 | 1.624473 | 2.324408 | 4.675089 | 7.380437 | 16.784824 | 29.566259 | 53.190777 | 89.606574 | 71.722369 | 101.120619 | 159.119497 | 103.600665 | 91.679164 | 43.606979 | 27.335274 | 11.544818 | 7.824976 | 5.472865 | 5.934112 | 4.372293 | 2.787602 | 2.544401 | 1.662534 | 0.529221 | 0.308489 | 0.230009 | 0.223201 | 0.238809 | 0.222845 | 0.218109 | 0.223516 | 0.231680 | 0.264567 | 0.249191 | 0.229395 | 0.247773 | 0.241899 | 0.254463 | 0.259299 | 0.251398 | 0.243546 | 0.231607 | 0.216783 | 0.226289 | 0.218648 | 0.215761 | 0.216105 | 0.216978 | 0.234585 | 0.252320 | 0.222672 | 0.225918 | 0.215357 | 0.215048 | 0.228810 | 0.266118 | 0.301538 | 0.268247 | 0.275529 | 0.242475 | 0.211942 | 0.218296 | 0.254070 | 0.414725 | 0.712206 | 0.927381 | 1.101508 | 0.956942 | 0.857231 | 0.815010 | 0.572498 | 0.398465 | 0.413811 | 0.538406 | 0.717342 | 0.811339 | 0.791061 | 0.550768 | 0.420769 | 0.273994 | 0.214825 | 0.210806 | 0.211174 | 0.218555 | 0.223375 | 0.217698 | 0.217211 | 0.218671 | 0.211659 | 0.224598 | 0.311189 | 0.350457 | 0.369850 | 0.391082 | 0.408557 | 0.397415 | 0.266631 | 0.212684 | 0.210822 | 0.210864 | 0.210819 | 0.219978 | 0.239700 | 0.237564 | 0.215607 | 0.210847 | 0.216096 | 0.216161 | 0.216339 | 0.213140 | 0.210995 | 0.210985 | 0.226524 | 0.360535 | 0.484355 | 0.369550 | 0.254567 | 0.213510 | 0.220451 | 0.211036 | 0.238735 | 0.323800 |

Searchlight, spatiotemporal cluster permutation test

|  | start time | stop time | peak time | peak channel | cluster p | peak Cohen's d | direction |
| --- | --- | --- | --- | --- | --- | --- | --- |
| #1 | 25 | 225 | 140 | POz | 0.0175 | 1.187032 | positive |
| #2 | 455 | 735 | 605 | CP6 | 0.019 | -1.017472 | negative |

F) emoji faces (LOSO) - neutral vs angry

  
|  | time window | peak latency | cluster *p* | peak Cohen's *d* |  | | | |  |
| **all electrodes** | 70 - 180 ms | 145 ms | 0.0109 | 1.8851 |  | | | |  |
 705 - 880 ms | 850 ms | 0.02 | 0.7573 |  | | | ||  |  | | | | | | | | |

Time-resolved classification, cluster permutation tests

|  | **left hemisphere** | | | | **right hemisphere** | | | |  |
|  | time window | peak latency | cluster *p* | peak Cohen's *d* | time window | peak latency | cluster *p* | peak Cohen's *d* |  |
| **anterior** |  | | | |  | | | |  |
| **central** |  | | | | 115 - 185 ms | 145 ms | 0.0194 | 1.5595 |  |
| **posterior** | 90 - 200 ms | 145 ms | 0.0087 | 2.0116 | 85 - 225 ms | 145 ms | 0.0151 | 2.0869 |  |
 360 - 875 ms | 495 ms | 0.0001 | 0.7918 | 355 - 1195 ms | 790 ms | 0.0001 | 1.2779 |

  

Time-resolved classification, Bayesian statistics

|  | -200 | -195 | -190 | -185 | -180 | -175 | -170 | -165 | -160 | -155 | -150 | -145 | -140 | -135 | -130 | -125 | -120 | -115 | -110 | -105 | -100 | -95 | -90 | -85 | -80 | -75 | -70 | -65 | -60 | -55 | -50 | -45 | -40 | -35 | -30 | -25 | -20 | -15 | -10 | -5 | 0 | 5 | 10 | 15 | 20 | 25 | 30 | 35 | 40 | 45 | 50 | 55 | 60 | 65 | 70 | 75 | 80 | 85 | 90 | 95 | 100 | 105 | 110 | 115 | 120 | 125 | 130 | 135 | 140 | 145 | 150 | 155 | 160 | 165 | 170 | 175 | 180 | 185 | 190 | 195 | 200 | 205 | 210 | 215 | 220 | 225 | 230 | 235 | 240 | 245 | 250 | 255 | 260 | 265 | 270 | 275 | 280 | 285 | 290 | 295 | 300 | 305 | 310 | 315 | 320 | 325 | 330 | 335 | 340 | 345 | 350 | 355 | 360 | 365 | 370 | 375 | 380 | 385 | 390 | 395 | 400 | 405 | 410 | 415 | 420 | 425 | 430 | 435 | 440 | 445 | 450 | 455 | 460 | 465 | 470 | 475 | 480 | 485 | 490 | 495 | 500 | 505 | 510 | 515 | 520 | 525 | 530 | 535 | 540 | 545 | 550 | 555 | 560 | 565 | 570 | 575 | 580 | 585 | 590 | 595 | 600 | 605 | 610 | 615 | 620 | 625 | 630 | 635 | 640 | 645 | 650 | 655 | 660 | 665 | 670 | 675 | 680 | 685 | 690 | 695 | 700 | 705 | 710 | 715 | 720 | 725 | 730 | 735 | 740 | 745 | 750 | 755 | 760 | 765 | 770 | 775 | 780 | 785 | 790 | 795 | 800 | 805 | 810 | 815 | 820 | 825 | 830 | 835 | 840 | 845 | 850 | 855 | 860 | 865 | 870 | 875 | 880 | 885 | 890 | 895 | 900 | 905 | 910 | 915 | 920 | 925 | 930 | 935 | 940 | 945 | 950 | 955 | 960 | 965 | 970 | 975 | 980 | 985 | 990 | 995 | 1000 | 1005 | 1010 | 1015 | 1020 | 1025 | 1030 | 1035 | 1040 | 1045 | 1050 | 1055 | 1060 | 1065 | 1070 | 1075 | 1080 | 1085 | 1090 | 1095 | 1100 | 1105 | 1110 | 1115 | 1120 | 1125 | 1130 | 1135 | 1140 | 1145 | 1150 | 1155 | 1160 | 1165 | 1170 | 1175 | 1180 | 1185 | 1190 | 1195 |
| --- | --- | --- | --- | --- | --- | --- | --- | --- | --- | --- | --- | --- | --- | --- | --- | --- | --- | --- | --- | --- | --- | --- | --- | --- | --- | --- | --- | --- | --- | --- | --- | --- | --- | --- | --- | --- | --- | --- | --- | --- | --- | --- | --- | --- | --- | --- | --- | --- | --- | --- | --- | --- | --- | --- | --- | --- | --- | --- | --- | --- | --- | --- | --- | --- | --- | --- | --- | --- | --- | --- | --- | --- | --- | --- | --- | --- | --- | --- | --- | --- | --- | --- | --- | --- | --- | --- | --- | --- | --- | --- | --- | --- | --- | --- | --- | --- | --- | --- | --- | --- | --- | --- | --- | --- | --- | --- | --- | --- | --- | --- | --- | --- | --- | --- | --- | --- | --- | --- | --- | --- | --- | --- | --- | --- | --- | --- | --- | --- | --- | --- | --- | --- | --- | --- | --- | --- | --- | --- | --- | --- | --- | --- | --- | --- | --- | --- | --- | --- | --- | --- | --- | --- | --- | --- | --- | --- | --- | --- | --- | --- | --- | --- | --- | --- | --- | --- | --- | --- | --- | --- | --- | --- | --- | --- | --- | --- | --- | --- | --- | --- | --- | --- | --- | --- | --- | --- | --- | --- | --- | --- | --- | --- | --- | --- | --- | --- | --- | --- | --- | --- | --- | --- | --- | --- | --- | --- | --- | --- | --- | --- | --- | --- | --- | --- | --- | --- | --- | --- | --- | --- | --- | --- | --- | --- | --- | --- | --- | --- | --- | --- | --- | --- | --- | --- | --- | --- | --- | --- | --- | --- | --- | --- | --- | --- | --- | --- | --- | --- | --- | --- | --- | --- | --- | --- | --- | --- | --- | --- | --- | --- | --- | --- | --- | --- | --- | --- | --- | --- | --- | --- | --- | --- | --- | --- | --- | --- | --- | --- | --- | --- |
| left anterior | 0.289888 | 0.391559 | 0.543175 | 0.715580 | 0.892628 | 1.405833 | 2.611136 | 3.769706 | 3.947533 | 3.483531 | 4.532410 | 6.255747 | 3.180180 | 1.838532 | 1.990760 | 0.426758 | 0.214768 | 0.210991 | 0.230360 | 0.211124 | 0.257668 | 0.239042 | 0.282035 | 0.532546 | 1.300808 | 1.645784 | 2.994146 | 1.686779 | 0.965474 | 0.781683 | 0.561746 | 0.219097 | 0.241813 | 0.409730 | 0.663818 | 0.421143 | 0.276613 | 0.259069 | 0.245289 | 0.222761 | 0.243798 | 0.280351 | 0.511115 | 0.875972 | 1.147094 | 1.753598 | 2.787023 | 2.620201 | 1.736260 | 0.730692 | 0.481551 | 0.304566 | 0.222264 | 0.210806 | 0.212391 | 0.211409 | 0.212835 | 0.215623 | 0.215828 | 0.214322 | 0.240414 | 0.321948 | 0.627914 | 1.816034 | 4.497237e+00 | 1.347621e+01 | 5.357898e+01 | 1.685207e+02 | 1.482684e+02 | 1.804741e+02 | 1.939712e+02 | 3.387069e+02 | 2.356029e+02 | 3.932648e+01 | 4.438029 | 1.934124 | 1.141558 | 0.578152 | 0.349373 | 0.342424 | 0.317231 | 0.341371 | 0.262001 | 0.214497 | 0.211320 | 0.211132 | 0.212340 | 0.217923 | 0.218214 | 0.246737 | 0.285490 | 0.337857 | 0.413871 | 0.506100 | 0.594653 | 0.645287 | 0.580865 | 0.472207 | 0.329030 | 0.232403 | 0.211136 | 0.221361 | 0.265340 | 0.306773 | 0.384685 | 0.500904 | 0.410841 | 0.349619 | 0.282686 | 0.214965 | 0.210870 | 0.231171 | 0.310559 | 0.364278 | 0.526711 | 0.811547 | 1.107875 | 1.509946 | 1.055392 | 0.690734 | 0.502001 | 0.459239 | 0.503610 | 0.651641 | 0.565575 | 0.486200 | 0.497788 | 0.537905 | 0.322778 | 0.218379 | 0.213051 | 0.225150 | 0.224076 | 0.213108 | 0.212645 | 0.264901 | 0.378084 | 0.506432 | 0.759478 | 1.308064 | 1.277328 | 1.135997 | 0.495852 | 0.294676 | 0.233294 | 0.218061 | 0.211890 | 0.248099 | 0.282538 | 0.285194 | 0.261881 | 0.287437 | 0.381855 | 0.492351 | 0.412817 | 0.327988 | 0.278843 | 0.323203 | 0.309769 | 0.284140 | 0.355441 | 0.418125 | 0.437897 | 0.443622 | 0.325524 | 0.261793 | 0.221948 | 0.213232 | 0.227629 | 0.229688 | 0.225173 | 0.221776 | 0.211347 | 0.229514 | 0.297897 | 0.361909 | 0.349348 | 0.341432 | 0.388779 | 0.342348 | 0.313998 | 0.317184 | 0.315408 | 0.343021 | 0.307085 | 0.283705 | 0.263930 | 0.258717 | 0.237049 | 0.217784 | 0.212214 | 0.212529 | 0.218555 | 0.228011 | 0.221583 | 0.214713 | 0.213530 | 0.212554 | 0.211408 | 0.229513 | 0.256680 | 0.249256 | 0.234162 | 0.227574 | 0.239330 | 0.229301 | 0.227438 | 0.236985 | 0.260512 | 0.295092 | 0.274785 | 0.242200 | 0.235352 | 0.233708 | 0.219241 | 0.215982 | 0.212714 | 0.217963 | 0.250024 | 0.274790 | 0.288263 | 0.271441 | 0.269887 | 0.291695 | 0.283574 | 0.251717 | 0.242151 | 0.220292 | 0.218888 | 0.216246 | 0.211435 | 0.224069 | 0.230473 | 0.238489 | 0.237195 | 0.239716 | 0.245117 | 0.235602 | 0.218162 | 0.221076 | 0.231754 | 0.240194 | 0.236513 | 0.276022 | 0.351228 | 0.468658 | 0.494145 | 0.453514 | 0.497167 | 0.565979 | 0.460846 | 0.421280 | 0.337883 | 0.275848 | 0.230357 | 0.211163 | 0.213722 | 0.222234 | 0.244477 | 0.263700 | 0.252444 | 0.247749 | 0.225861 | 0.212576 | 0.212248 | 0.218847 | 0.225981 | 0.229899 | 0.241971 | 0.223873 | 0.212416 | 0.214835 | 0.230917 | 0.250516 | 0.329164 | 0.454911 | 0.729446 | 0.893958 | 0.974516 | 1.289845 |
| right anterior | 18.773648 | 11.281681 | 4.254413 | 1.870358 | 0.456483 | 0.237214 | 0.215560 | 0.210806 | 0.211050 | 0.212527 | 0.248568 | 0.275335 | 0.317281 | 0.623532 | 0.750523 | 0.520803 | 0.306354 | 0.214433 | 0.227951 | 0.260723 | 0.496560 | 0.542347 | 0.361453 | 0.293913 | 0.279047 | 0.221530 | 0.276376 | 0.286335 | 0.342163 | 0.237219 | 0.313294 | 0.479488 | 0.386719 | 0.326608 | 0.239403 | 0.218211 | 0.250522 | 0.242136 | 0.217422 | 0.220319 | 0.211216 | 0.226993 | 0.244336 | 0.252659 | 0.242133 | 0.267772 | 0.501061 | 0.492094 | 0.339091 | 0.311674 | 0.318223 | 0.458446 | 0.477677 | 0.276546 | 0.222822 | 0.216141 | 0.211329 | 0.238701 | 0.301157 | 0.367155 | 0.425255 | 0.489588 | 0.577155 | 0.864460 | 2.641274e+00 | 1.270890e+01 | 6.336587e+01 | 2.317495e+02 | 6.704107e+02 | 1.583015e+03 | 5.037414e+02 | 1.098390e+02 | 4.765637e+01 | 2.755746e+01 | 6.842025 | 1.777740 | 0.773035 | 0.653721 | 0.460939 | 0.335477 | 0.302382 | 0.335193 | 0.264594 | 0.214230 | 0.211942 | 0.219117 | 0.272371 | 0.414278 | 0.407346 | 0.481849 | 0.531506 | 0.571178 | 0.721479 | 0.490102 | 0.322190 | 0.345162 | 0.282137 | 0.351973 | 0.814992 | 1.334836 | 1.411440 | 3.227935 | 2.608171 | 1.363551 | 0.545731 | 0.339557 | 0.297960 | 0.288345 | 0.236419 | 0.220412 | 0.210922 | 0.212566 | 0.212919 | 0.226174 | 0.247901 | 0.236236 | 0.222952 | 0.211744 | 0.222896 | 0.239094 | 0.290384 | 0.336983 | 0.343427 | 0.293146 | 0.278683 | 0.275040 | 0.276738 | 0.272761 | 0.278819 | 0.388007 | 0.845915 | 1.594412 | 1.885115 | 2.406801 | 3.932642 | 6.172584 | 5.452925 | 2.537707 | 2.512657 | 3.041023 | 2.183092 | 1.282995 | 1.020921 | 0.616433 | 0.713284 | 0.858282 | 0.873393 | 1.133925 | 1.619785 | 1.411040 | 1.694288 | 1.674658 | 1.812428 | 1.873438 | 1.286789 | 0.719450 | 0.674353 | 0.766308 | 0.795101 | 0.555353 | 0.519889 | 0.501836 | 0.446313 | 0.342185 | 0.254427 | 0.214468 | 0.212819 | 0.236912 | 0.269216 | 0.277370 | 0.275360 | 0.252587 | 0.223228 | 0.211307 | 0.214615 | 0.218821 | 0.221727 | 0.225321 | 0.238657 | 0.247043 | 0.228519 | 0.212050 | 0.210869 | 0.210996 | 0.210878 | 0.212852 | 0.220566 | 0.214427 | 0.213852 | 0.245846 | 0.266921 | 0.324160 | 0.324699 | 0.306083 | 0.255506 | 0.227894 | 0.211144 | 0.222969 | 0.256986 | 0.286645 | 0.285798 | 0.246104 | 0.226801 | 0.216672 | 0.211180 | 0.212101 | 0.222287 | 0.229441 | 0.232622 | 0.237446 | 0.248355 | 0.250004 | 0.236690 | 0.222058 | 0.215190 | 0.211102 | 0.216267 | 0.224380 | 0.227302 | 0.225464 | 0.234060 | 0.250617 | 0.266711 | 0.275149 | 0.318274 | 0.404803 | 0.408930 | 0.330838 | 0.295402 | 0.312785 | 0.325279 | 0.324365 | 0.281436 | 0.284368 | 0.314843 | 0.316169 | 0.315829 | 0.313528 | 0.314105 | 0.341111 | 0.426451 | 0.535434 | 0.610314 | 0.478007 | 0.351726 | 0.299220 | 0.262700 | 0.257575 | 0.270897 | 0.289828 | 0.301846 | 0.308733 | 0.325141 | 0.336533 | 0.276499 | 0.264063 | 0.269287 | 0.289756 | 0.391379 | 0.516013 | 0.701143 | 1.125070 | 1.013167 | 0.849355 | 0.800822 | 0.719040 | 0.478671 | 0.364959 | 0.291209 | 0.295401 | 0.296429 | 0.300470 | 0.324118 | 0.460767 | 0.677155 | 1.047482 | 1.120560 | 1.509638 | 1.884006 | 1.639320 |
| left central | 1.710436 | 0.515337 | 0.318129 | 0.239970 | 0.211624 | 0.294089 | 0.590369 | 0.999451 | 0.961412 | 1.559143 | 2.099595 | 1.664558 | 0.843966 | 0.351618 | 0.247709 | 0.224888 | 0.220697 | 0.343640 | 0.302125 | 0.281790 | 0.238339 | 0.216616 | 0.211164 | 0.221094 | 0.219586 | 0.211162 | 0.213630 | 0.223871 | 0.217218 | 0.212202 | 0.215751 | 0.215330 | 0.213442 | 0.235713 | 0.237644 | 0.242160 | 0.256773 | 0.295934 | 0.391543 | 0.710954 | 0.599122 | 0.411214 | 0.346240 | 0.422236 | 0.385268 | 0.268499 | 0.233320 | 0.230102 | 0.254351 | 0.254082 | 0.230238 | 0.210820 | 0.225051 | 0.273459 | 0.310087 | 0.401948 | 0.390929 | 0.443552 | 0.439390 | 0.502945 | 0.610694 | 0.892997 | 0.940191 | 1.739281 | 2.779239e+00 | 4.394939e+00 | 7.066590e+00 | 1.826525e+01 | 4.686720e+01 | 9.788809e+01 | 1.251446e+02 | 1.751582e+02 | 1.777962e+02 | 6.228921e+01 | 6.006909 | 1.327242 | 0.740867 | 0.572089 | 0.635308 | 0.585673 | 0.647336 | 0.977946 | 0.925680 | 0.792982 | 0.664502 | 0.352374 | 0.274052 | 0.244067 | 0.216160 | 0.215457 | 0.213286 | 0.212375 | 0.216433 | 0.221919 | 0.221659 | 0.237977 | 0.257389 | 0.278456 | 0.259816 | 0.233055 | 0.212066 | 0.223603 | 0.338832 | 0.892587 | 4.815461 | 18.532447 | 21.219437 | 8.112060 | 3.211152 | 1.465960 | 0.568300 | 0.274174 | 0.220888 | 0.212707 | 0.213840 | 0.210806 | 0.221349 | 0.241265 | 0.257930 | 0.231003 | 0.212698 | 0.211683 | 0.221108 | 0.251038 | 0.292973 | 0.264546 | 0.227140 | 0.212148 | 0.212547 | 0.223245 | 0.241272 | 0.248609 | 0.242150 | 0.244216 | 0.226659 | 0.218311 | 0.211297 | 0.213116 | 0.213860 | 0.213471 | 0.212285 | 0.213765 | 0.217188 | 0.214119 | 0.217893 | 0.216265 | 0.211798 | 0.212854 | 0.231277 | 0.237514 | 0.224291 | 0.224077 | 0.251547 | 0.287033 | 0.256277 | 0.221226 | 0.222490 | 0.251050 | 0.248524 | 0.235011 | 0.231412 | 0.243535 | 0.265620 | 0.274540 | 0.278493 | 0.277686 | 0.307914 | 0.346716 | 0.430957 | 0.528754 | 0.575782 | 0.446715 | 0.575551 | 0.638434 | 0.575411 | 0.386771 | 0.360216 | 0.331681 | 0.393626 | 0.429266 | 0.452097 | 0.483911 | 0.574926 | 0.513801 | 0.552603 | 0.417753 | 0.286206 | 0.215636 | 0.219005 | 0.239869 | 0.243716 | 0.225192 | 0.211718 | 0.211449 | 0.218726 | 0.262102 | 0.343262 | 0.352070 | 0.312179 | 0.273942 | 0.313908 | 0.448650 | 0.434075 | 0.315145 | 0.282591 | 0.266750 | 0.283860 | 0.333644 | 0.326007 | 0.329369 | 0.390111 | 0.505240 | 0.625163 | 0.720614 | 0.639623 | 0.635526 | 0.711541 | 0.949721 | 1.241084 | 1.136319 | 0.763360 | 0.479852 | 0.316592 | 0.234968 | 0.210852 | 0.232834 | 0.259128 | 0.281464 | 0.280859 | 0.250786 | 0.237191 | 0.218130 | 0.211300 | 0.210970 | 0.213542 | 0.213704 | 0.212246 | 0.215693 | 0.212442 | 0.210967 | 0.211301 | 0.211066 | 0.218597 | 0.238770 | 0.240107 | 0.232780 | 0.222843 | 0.228756 | 0.228030 | 0.212695 | 0.211164 | 0.214721 | 0.216660 | 0.213712 | 0.218158 | 0.215387 | 0.211371 | 0.217669 | 0.224296 | 0.217877 | 0.226713 | 0.239905 | 0.292511 | 0.329190 | 0.362506 | 0.348659 | 0.372939 | 0.297703 | 0.259534 | 0.225292 | 0.214755 | 0.210806 | 0.211340 | 0.210832 | 0.216682 | 0.229543 | 0.253632 | 0.284298 | 0.313478 | 0.358069 |
| right central | 0.221655 | 0.210823 | 0.211159 | 0.211246 | 0.220531 | 0.220058 | 0.214914 | 0.213601 | 0.213887 | 0.215984 | 0.232946 | 0.296053 | 0.244893 | 0.233771 | 0.250755 | 0.317094 | 0.407335 | 0.579425 | 1.833357 | 3.569680 | 1.366547 | 0.698390 | 0.510255 | 0.518675 | 0.390009 | 0.221070 | 0.220282 | 0.211057 | 0.254790 | 0.513206 | 0.455048 | 0.531155 | 0.512960 | 0.377895 | 0.257078 | 0.214708 | 0.210825 | 0.212180 | 0.245487 | 0.265862 | 0.234519 | 0.221871 | 0.226793 | 0.224470 | 0.265490 | 0.463898 | 1.926097 | 5.236528 | 2.689535 | 0.573163 | 0.299222 | 0.211290 | 0.313791 | 0.621494 | 1.202719 | 1.264029 | 1.132152 | 1.508563 | 1.543333 | 0.767994 | 0.590054 | 0.634493 | 0.783862 | 1.387333 | 4.088521e+00 | 2.042478e+01 | 2.476952e+02 | 3.863184e+03 | 2.894527e+04 | 2.894196e+05 | 5.238958e+05 | 3.962625e+05 | 1.319556e+05 | 4.615018e+04 | 5080.152535 | 116.695006 | 11.863028 | 1.637056 | 0.856961 | 0.601693 | 0.475867 | 0.373678 | 0.325768 | 0.259014 | 0.252434 | 0.230458 | 0.230201 | 0.234206 | 0.261132 | 0.355101 | 0.446712 | 0.381730 | 0.390071 | 0.343468 | 0.338975 | 0.346917 | 0.394234 | 0.455350 | 0.438360 | 0.308295 | 0.277839 | 0.248004 | 0.261327 | 0.228930 | 0.221430 | 0.230247 | 0.261553 | 0.315785 | 0.413834 | 0.406015 | 0.564039 | 0.794240 | 1.162888 | 2.170251 | 3.382469 | 4.715456 | 7.639047 | 5.248331 | 5.351199 | 6.957006 | 9.364350 | 6.446842 | 2.250212 | 1.209455 | 0.865419 | 0.582886 | 0.360362 | 0.267025 | 0.234888 | 0.239221 | 0.217489 | 0.212042 | 0.220276 | 0.211107 | 0.223058 | 0.246413 | 0.276054 | 0.421279 | 0.899602 | 1.481227 | 2.002527 | 1.221889 | 0.639941 | 0.333380 | 0.214454 | 0.220744 | 0.241248 | 0.260030 | 0.242177 | 0.225897 | 0.211438 | 0.217072 | 0.231721 | 0.245101 | 0.259954 | 0.275152 | 0.342490 | 0.327234 | 0.267495 | 0.230014 | 0.224331 | 0.221117 | 0.214091 | 0.211222 | 0.214771 | 0.211550 | 0.213444 | 0.210815 | 0.211098 | 0.213320 | 0.216575 | 0.213687 | 0.212954 | 0.210814 | 0.211649 | 0.211230 | 0.211232 | 0.219500 | 0.219146 | 0.214195 | 0.211152 | 0.219678 | 0.269808 | 0.335354 | 0.446674 | 0.476270 | 0.455846 | 0.409173 | 0.340744 | 0.246790 | 0.224721 | 0.215053 | 0.215300 | 0.216949 | 0.222650 | 0.237354 | 0.292506 | 0.351664 | 0.441659 | 0.555879 | 0.542151 | 0.550762 | 0.549585 | 0.567279 | 0.579213 | 0.541013 | 0.587378 | 0.562353 | 0.478899 | 0.382715 | 0.348294 | 0.366684 | 0.343288 | 0.278681 | 0.289598 | 0.372956 | 0.525122 | 0.485806 | 0.364598 | 0.339184 | 0.384048 | 0.422277 | 0.450176 | 0.431023 | 0.504777 | 0.571502 | 0.577633 | 0.598559 | 0.567046 | 0.464513 | 0.349948 | 0.288024 | 0.249115 | 0.221925 | 0.220401 | 0.294998 | 0.367610 | 0.390456 | 0.344903 | 0.285314 | 0.264176 | 0.231624 | 0.229033 | 0.232212 | 0.221852 | 0.216631 | 0.215775 | 0.217684 | 0.241081 | 0.260726 | 0.244712 | 0.245227 | 0.235383 | 0.236538 | 0.251857 | 0.234425 | 0.212860 | 0.211233 | 0.218718 | 0.215270 | 0.212260 | 0.221972 | 0.229211 | 0.223423 | 0.215321 | 0.211002 | 0.212235 | 0.212272 | 0.219018 | 0.226844 | 0.259314 | 0.270162 | 0.319931 | 0.364031 | 0.318088 | 0.269708 | 0.237159 | 0.221061 | 0.216572 | 0.211178 |
| left posterior | 0.314913 | 0.364311 | 0.378841 | 0.467083 | 0.491204 | 0.494297 | 0.703751 | 1.136334 | 1.821668 | 1.573775 | 0.964722 | 0.933593 | 0.535897 | 0.355004 | 0.222551 | 0.210825 | 0.215960 | 0.224041 | 0.267732 | 0.233898 | 0.228479 | 0.213918 | 0.213215 | 0.212041 | 0.247262 | 0.530208 | 0.502292 | 0.577019 | 0.634263 | 1.129390 | 5.999332 | 8.883436 | 4.775573 | 2.985536 | 3.303807 | 2.045065 | 0.691708 | 0.296694 | 0.215023 | 0.210822 | 0.225265 | 0.215987 | 0.212945 | 0.212089 | 0.248325 | 0.260339 | 0.285713 | 0.334532 | 0.329614 | 0.326348 | 0.249447 | 0.212178 | 0.228836 | 0.230654 | 0.210827 | 0.215331 | 0.309014 | 1.196180 | 11.189305 | 36.078588 | 135.240261 | 1490.592761 | 14435.743802 | 40107.964817 | 1.070256e+05 | 3.041931e+05 | 1.459735e+06 | 3.029481e+06 | 1.062899e+07 | 2.418724e+07 | 4.865345e+07 | 7.722577e+07 | 2.749552e+07 | 1.618058e+06 | 82891.206454 | 1840.468951 | 154.643379 | 35.258599 | 15.856264 | 3.681927 | 1.394089 | 0.683462 | 0.553245 | 0.405749 | 0.312456 | 0.262163 | 0.231105 | 0.211376 | 0.211070 | 0.210852 | 0.215398 | 0.241135 | 0.297621 | 0.589515 | 2.301761 | 5.610261 | 7.077398 | 5.333586 | 3.848621 | 2.378933 | 1.640233 | 0.826921 | 0.573240 | 0.518723 | 0.583599 | 0.537177 | 0.477047 | 0.448135 | 0.459611 | 0.469200 | 0.587238 | 0.705456 | 1.303973 | 2.997419 | 8.434706 | 20.883260 | 52.173404 | 55.835290 | 63.153984 | 64.207285 | 58.334169 | 32.306823 | 13.954967 | 10.445640 | 9.812448 | 9.909467 | 8.478579 | 10.474718 | 13.265302 | 28.788548 | 34.317122 | 40.141908 | 38.189588 | 40.666554 | 44.717049 | 52.140139 | 63.828893 | 70.750688 | 59.911722 | 55.725697 | 43.048135 | 31.698001 | 22.316693 | 11.429324 | 8.717797 | 6.884941 | 5.378202 | 4.435721 | 3.553161 | 2.796019 | 3.084518 | 3.186575 | 2.633344 | 2.676266 | 3.079316 | 5.259933 | 10.889922 | 12.021703 | 8.864571 | 12.635256 | 17.646995 | 20.533900 | 21.460818 | 11.854226 | 8.297550 | 5.374441 | 3.870384 | 3.330635 | 3.476071 | 3.827798 | 6.113968 | 8.615132 | 11.997460 | 20.437019 | 30.776864 | 46.477678 | 69.777576 | 79.686456 | 63.636290 | 109.019713 | 142.212240 | 292.638957 | 389.481130 | 233.487565 | 194.828799 | 141.088919 | 45.413777 | 19.928228 | 7.988402 | 4.778394 | 4.346382 | 3.536383 | 6.298153 | 18.370894 | 95.874809 | 304.451797 | 502.924529 | 495.935474 | 831.742332 | 610.369170 | 191.232892 | 42.834481 | 19.604048 | 13.440362 | 6.583924 | 3.720364 | 1.966668 | 1.858106 | 2.591801 | 3.462077 | 4.391182 | 5.525821 | 3.254570 | 3.194475 | 2.484419 | 1.479414 | 1.035311 | 0.829433 | 0.819421 | 1.197354 | 1.255319 | 1.113132 | 1.204024 | 1.693420 | 2.551252 | 3.142663 | 3.318614 | 2.405926 | 1.962725 | 1.470148 | 0.880714 | 0.644740 | 0.503830 | 0.443013 | 0.476705 | 0.560767 | 0.735346 | 0.901014 | 0.926621 | 1.162915 | 1.580243 | 2.177849 | 2.002644 | 1.624078 | 1.476414 | 1.392249 | 1.107900 | 0.912984 | 0.929421 | 0.970557 | 1.055351 | 1.284220 | 1.443121 | 1.506966 | 1.437435 | 1.012068 | 0.749495 | 0.823353 | 0.883204 | 0.871312 | 0.837138 | 0.764984 | 0.852037 | 1.235217 | 1.250632 | 0.968594 | 0.819409 | 0.797496 | 0.803159 | 0.700917 | 0.568504 | 0.636939 | 0.807524 | 1.248447 | 1.629520 | 1.760821 | 1.878464 | 1.907877 | 1.360043 | 1.095201 |
| right posterior | 0.216418 | 0.229266 | 0.244508 | 0.253832 | 0.250527 | 0.256657 | 0.289563 | 0.317643 | 0.372822 | 0.353211 | 0.250082 | 0.214246 | 0.324996 | 1.163411 | 3.142231 | 6.671106 | 3.147667 | 1.228646 | 0.303261 | 0.213099 | 0.224525 | 0.276603 | 0.314188 | 0.258319 | 0.225510 | 0.211057 | 0.231666 | 0.230648 | 0.223435 | 0.233824 | 0.289936 | 0.293125 | 0.302624 | 0.269934 | 0.248601 | 0.231618 | 0.226204 | 0.212876 | 0.210897 | 0.211560 | 0.211197 | 0.218349 | 0.263281 | 0.256366 | 0.264881 | 0.248716 | 0.233048 | 0.227312 | 0.210806 | 0.233322 | 0.225021 | 0.240541 | 0.239823 | 0.246574 | 0.327379 | 0.579679 | 1.015461 | 5.707430 | 25.992159 | 126.866585 | 699.152710 | 4980.924952 | 59183.922513 | 601186.978936 | 1.389288e+06 | 2.321026e+06 | 5.468463e+06 | 1.154340e+07 | 2.493461e+07 | 4.784113e+07 | 6.575870e+07 | 3.866628e+07 | 6.424820e+06 | 8.253726e+05 | 32695.982896 | 1730.898302 | 54.891690 | 9.537622 | 4.645057 | 3.829673 | 2.049045 | 1.769992 | 2.442726 | 2.539574 | 2.071788 | 1.538513 | 0.892982 | 0.620912 | 0.422417 | 0.381367 | 0.437245 | 0.572438 | 1.028358 | 3.086958 | 12.221854 | 33.698627 | 65.377924 | 174.138063 | 234.185374 | 88.129710 | 13.288303 | 2.759442 | 0.738785 | 0.361904 | 0.265889 | 0.236174 | 0.233844 | 0.262402 | 0.312880 | 0.450349 | 0.679191 | 1.404225 | 4.078047 | 13.643700 | 24.428145 | 60.271538 | 172.146194 | 294.767530 | 350.762262 | 233.588191 | 89.893288 | 47.065256 | 16.872078 | 6.490968 | 4.475445 | 3.239028 | 5.143976 | 6.921036 | 9.533716 | 22.847766 | 69.957084 | 182.693431 | 353.253310 | 238.276262 | 301.985610 | 370.360744 | 431.893944 | 369.464844 | 189.318748 | 152.809914 | 172.956339 | 177.389190 | 162.237241 | 120.818281 | 47.974428 | 35.223419 | 26.201741 | 16.248455 | 11.399993 | 10.326743 | 5.981205 | 5.019498 | 3.312169 | 2.707881 | 3.866157 | 6.614804 | 8.320818 | 12.293259 | 15.260069 | 22.262119 | 22.205525 | 17.305797 | 14.106526 | 14.511918 | 21.978717 | 60.164381 | 159.546355 | 400.540614 | 828.182732 | 1078.631496 | 1102.789113 | 935.948003 | 448.243002 | 209.102579 | 185.704422 | 229.719768 | 370.987953 | 902.112884 | 1964.533477 | 5362.771117 | 16698.867370 | 25079.820675 | 13843.474949 | 9518.293967 | 5873.776084 | 4828.349853 | 2707.191896 | 1249.286714 | 809.568602 | 836.566564 | 932.786524 | 824.791256 | 759.944137 | 907.219600 | 1432.148035 | 2813.381864 | 5362.292786 | 6894.909911 | 13848.512060 | 20568.803998 | 23065.182160 | 24728.505874 | 11599.695274 | 8576.014194 | 5465.498466 | 2387.515298 | 964.010268 | 950.754719 | 1482.738890 | 2258.142348 | 3674.918328 | 5114.527335 | 4137.677553 | 4816.773597 | 4165.911499 | 3443.297326 | 1851.369183 | 827.366587 | 402.456261 | 316.193682 | 258.677499 | 276.946459 | 312.980172 | 411.483326 | 452.992663 | 647.563559 | 923.258155 | 939.237849 | 615.967699 | 326.461569 | 256.765561 | 208.524039 | 131.553938 | 71.762602 | 54.346851 | 40.585678 | 34.219733 | 28.437542 | 37.506703 | 43.677152 | 69.847998 | 96.831272 | 102.848377 | 101.882490 | 105.834904 | 126.174909 | 105.440284 | 104.965716 | 177.743605 | 190.070531 | 229.999379 | 299.634346 | 240.975460 | 333.316385 | 363.536619 | 301.203716 | 493.918033 | 832.913987 | 1127.656296 | 1331.519342 | 1314.252236 | 804.360121 | 650.941472 | 462.996370 | 300.617438 | 167.122037 | 108.802547 | 129.986722 | 182.242471 | 402.085448 | 913.917570 | 2971.053037 | 16684.824275 | 85789.245942 | 49611.404136 | 18012.612933 | 5466.474190 | 2293.430653 | 809.286117 | 350.663328 |
| all electrodes | 0.462661 | 0.584872 | 0.710293 | 0.596453 | 0.544754 | 0.638886 | 0.659676 | 0.507245 | 0.529243 | 0.405297 | 0.433416 | 0.428250 | 0.214366 | 0.393331 | 0.407742 | 0.352967 | 0.270222 | 0.275725 | 0.324344 | 0.270223 | 0.213324 | 0.212116 | 0.228296 | 0.237869 | 0.257992 | 0.226948 | 0.219956 | 0.221490 | 0.232729 | 0.231509 | 0.231092 | 0.225488 | 0.255162 | 0.275233 | 0.480695 | 0.674687 | 0.809951 | 1.448497 | 2.564246 | 1.727871 | 0.569067 | 0.273615 | 0.237170 | 0.293042 | 0.562223 | 0.636444 | 0.674823 | 0.645247 | 0.418261 | 0.301806 | 0.246813 | 0.243763 | 0.321001 | 0.744057 | 1.318186 | 1.851792 | 4.534789 | 18.408005 | 22.293734 | 68.123925 | 121.259122 | 274.244317 | 1520.036341 | 4905.524938 | 2.170143e+04 | 6.027375e+04 | 3.072469e+05 | 1.842308e+06 | 6.284041e+06 | 7.433400e+06 | 8.752520e+06 | 6.953289e+06 | 2.600039e+06 | 2.606973e+05 | 4981.969873 | 81.932546 | 4.923001 | 0.548972 | 0.262477 | 0.244938 | 0.265116 | 0.325007 | 0.413592 | 0.398021 | 0.417758 | 0.270379 | 0.219226 | 0.210920 | 0.218161 | 0.212672 | 0.210819 | 0.214556 | 0.234301 | 0.295246 | 0.379686 | 0.596501 | 0.619900 | 0.639193 | 0.624943 | 0.706791 | 0.553065 | 0.354928 | 0.230637 | 0.217560 | 0.212756 | 0.210882 | 0.210838 | 0.218088 | 0.246201 | 0.394799 | 0.552096 | 0.743107 | 0.920777 | 1.112126 | 1.398364 | 1.872605 | 1.958549 | 1.791206 | 1.657428 | 1.513888 | 1.303461 | 0.826445 | 0.584169 | 0.416997 | 0.333319 | 0.337374 | 0.426291 | 0.676472 | 1.228944 | 2.279590 | 6.739045 | 11.538589 | 13.745040 | 12.817242 | 13.280135 | 14.663384 | 18.157179 | 12.057994 | 11.529719 | 9.059965 | 6.157678 | 4.268604 | 3.075246 | 2.403134 | 2.497459 | 1.891363 | 1.500899 | 1.045174 | 1.014612 | 1.153452 | 1.054443 | 0.808986 | 0.686050 | 0.541537 | 0.547586 | 0.623789 | 0.634275 | 0.614874 | 0.674935 | 0.657402 | 0.622017 | 0.623852 | 0.481290 | 0.359990 | 0.371933 | 0.406684 | 0.472696 | 0.586756 | 0.672017 | 0.791383 | 1.280012 | 1.799967 | 1.862165 | 2.093622 | 2.527963 | 2.589667 | 1.816693 | 1.199629 | 0.821480 | 0.929264 | 1.169914 | 1.574866 | 2.223098 | 5.044862 | 10.162635 | 12.975399 | 7.356877 | 3.877099 | 2.672557 | 2.646936 | 1.785831 | 1.500272 | 1.766135 | 3.482032 | 7.773055 | 10.029393 | 10.701704 | 11.557220 | 8.892061 | 8.520105 | 5.060298 | 2.925131 | 2.540757 | 2.261185 | 3.247541 | 7.628268 | 9.516431 | 18.901028 | 22.656119 | 28.275644 | 38.020752 | 23.166077 | 11.146056 | 8.245690 | 4.337029 | 2.614118 | 1.541215 | 1.007552 | 1.041662 | 0.978737 | 0.706846 | 0.589692 | 0.701786 | 0.750543 | 1.012788 | 0.931981 | 1.001562 | 1.555407 | 3.612952 | 4.777839 | 9.639165 | 7.893808 | 5.735689 | 3.060790 | 1.543808 | 0.758563 | 0.608972 | 0.615487 | 0.761105 | 0.797096 | 1.102167 | 2.088522 | 3.934166 | 4.410119 | 3.967731 | 2.845374 | 2.699737 | 2.221429 | 1.330305 | 0.753554 | 0.587483 | 0.437375 | 0.375863 | 0.378065 | 0.371260 | 0.374254 | 0.382134 | 0.467222 | 0.647695 | 0.951178 | 1.101322 | 1.293176 | 2.222965 | 3.811406 | 4.458892 | 4.828010 | 4.956165 | 6.081705 | 5.815701 | 4.381853 | 3.393094 | 4.034898 | 4.101936 | 5.718364 | 6.959611 | 12.538650 | 14.462424 | 18.564190 | 23.333693 | 31.721480 |

Searchlight, spatiotemporal cluster permutation test

|  | start time | stop time | peak time | peak channel | cluster p | peak Cohen's d | direction |
| --- | --- | --- | --- | --- | --- | --- | --- |
| #1 | 60 | 230 | 145 | PO8 | 0.012 | 2.109067 | positive |
| #2 | 255 | 1195 | 785 | P6 | 0.0002 | 1.536356 | positive |

G) emoji faces (LOSO) - neutral vs sad

  
|  | time window | peak latency | cluster *p* | peak Cohen's *d* |  | | | |  |
| **all electrodes** | 105 - 190 ms | 155 ms | 0.0244 | 1.4999 |  | | | |  |
 720 - 920 ms | 790 ms | 0.0086 | 1.2179 |  | | | | 955 - 1070 ms | 1025 ms | 0.0456 | 0.5905 |  | | | ||  |  | | | | | | | | |

Time-resolved classification, cluster permutation tests

|  |  | **left hemisphere** | | | | **right hemisphere** | | | |
|  | time window | peak latency | cluster *p* | peak Cohen's *d* | time window | peak latency | cluster *p* | peak Cohen's *d* |  |
| **anterior** |  | | | |  | | | |  |
| **central** | 600 - 675 ms | 600 ms | 0.0416 | -0.4429 |  | | | |  |
| **posterior** | 115 - 185 ms | 150 ms | 0.0356 | 1.1776 | 105 - 185 ms | 155 ms | 0.0271 | 1.57 |  |
  | | | | 580 - 930 ms | 605 ms | 0.0027 | 0.8682 |

  

Time-resolved classification, Bayesian statistics

|  | -200 | -195 | -190 | -185 | -180 | -175 | -170 | -165 | -160 | -155 | -150 | -145 | -140 | -135 | -130 | -125 | -120 | -115 | -110 | -105 | -100 | -95 | -90 | -85 | -80 | -75 | -70 | -65 | -60 | -55 | -50 | -45 | -40 | -35 | -30 | -25 | -20 | -15 | -10 | -5 | 0 | 5 | 10 | 15 | 20 | 25 | 30 | 35 | 40 | 45 | 50 | 55 | 60 | 65 | 70 | 75 | 80 | 85 | 90 | 95 | 100 | 105 | 110 | 115 | 120 | 125 | 130 | 135 | 140 | 145 | 150 | 155 | 160 | 165 | 170 | 175 | 180 | 185 | 190 | 195 | 200 | 205 | 210 | 215 | 220 | 225 | 230 | 235 | 240 | 245 | 250 | 255 | 260 | 265 | 270 | 275 | 280 | 285 | 290 | 295 | 300 | 305 | 310 | 315 | 320 | 325 | 330 | 335 | 340 | 345 | 350 | 355 | 360 | 365 | 370 | 375 | 380 | 385 | 390 | 395 | 400 | 405 | 410 | 415 | 420 | 425 | 430 | 435 | 440 | 445 | 450 | 455 | 460 | 465 | 470 | 475 | 480 | 485 | 490 | 495 | 500 | 505 | 510 | 515 | 520 | 525 | 530 | 535 | 540 | 545 | 550 | 555 | 560 | 565 | 570 | 575 | 580 | 585 | 590 | 595 | 600 | 605 | 610 | 615 | 620 | 625 | 630 | 635 | 640 | 645 | 650 | 655 | 660 | 665 | 670 | 675 | 680 | 685 | 690 | 695 | 700 | 705 | 710 | 715 | 720 | 725 | 730 | 735 | 740 | 745 | 750 | 755 | 760 | 765 | 770 | 775 | 780 | 785 | 790 | 795 | 800 | 805 | 810 | 815 | 820 | 825 | 830 | 835 | 840 | 845 | 850 | 855 | 860 | 865 | 870 | 875 | 880 | 885 | 890 | 895 | 900 | 905 | 910 | 915 | 920 | 925 | 930 | 935 | 940 | 945 | 950 | 955 | 960 | 965 | 970 | 975 | 980 | 985 | 990 | 995 | 1000 | 1005 | 1010 | 1015 | 1020 | 1025 | 1030 | 1035 | 1040 | 1045 | 1050 | 1055 | 1060 | 1065 | 1070 | 1075 | 1080 | 1085 | 1090 | 1095 | 1100 | 1105 | 1110 | 1115 | 1120 | 1125 | 1130 | 1135 | 1140 | 1145 | 1150 | 1155 | 1160 | 1165 | 1170 | 1175 | 1180 | 1185 | 1190 | 1195 |
| --- | --- | --- | --- | --- | --- | --- | --- | --- | --- | --- | --- | --- | --- | --- | --- | --- | --- | --- | --- | --- | --- | --- | --- | --- | --- | --- | --- | --- | --- | --- | --- | --- | --- | --- | --- | --- | --- | --- | --- | --- | --- | --- | --- | --- | --- | --- | --- | --- | --- | --- | --- | --- | --- | --- | --- | --- | --- | --- | --- | --- | --- | --- | --- | --- | --- | --- | --- | --- | --- | --- | --- | --- | --- | --- | --- | --- | --- | --- | --- | --- | --- | --- | --- | --- | --- | --- | --- | --- | --- | --- | --- | --- | --- | --- | --- | --- | --- | --- | --- | --- | --- | --- | --- | --- | --- | --- | --- | --- | --- | --- | --- | --- | --- | --- | --- | --- | --- | --- | --- | --- | --- | --- | --- | --- | --- | --- | --- | --- | --- | --- | --- | --- | --- | --- | --- | --- | --- | --- | --- | --- | --- | --- | --- | --- | --- | --- | --- | --- | --- | --- | --- | --- | --- | --- | --- | --- | --- | --- | --- | --- | --- | --- | --- | --- | --- | --- | --- | --- | --- | --- | --- | --- | --- | --- | --- | --- | --- | --- | --- | --- | --- | --- | --- | --- | --- | --- | --- | --- | --- | --- | --- | --- | --- | --- | --- | --- | --- | --- | --- | --- | --- | --- | --- | --- | --- | --- | --- | --- | --- | --- | --- | --- | --- | --- | --- | --- | --- | --- | --- | --- | --- | --- | --- | --- | --- | --- | --- | --- | --- | --- | --- | --- | --- | --- | --- | --- | --- | --- | --- | --- | --- | --- | --- | --- | --- | --- | --- | --- | --- | --- | --- | --- | --- | --- | --- | --- | --- | --- | --- | --- | --- | --- | --- | --- | --- | --- | --- | --- | --- | --- | --- | --- | --- | --- | --- | --- | --- | --- | --- | --- |
| left anterior | 0.341715 | 0.546794 | 0.694739 | 0.940114 | 1.461405 | 3.049880 | 3.151093 | 3.515982 | 3.955919 | 5.019841 | 5.004625 | 2.720763 | 1.013042 | 0.347712 | 0.251403 | 0.253048 | 1.358197 | 1.814136 | 1.154411 | 0.891336 | 0.284775 | 0.224228 | 0.211959 | 0.269897 | 0.357272 | 0.440473 | 0.327380 | 0.216161 | 0.222378 | 0.222648 | 0.213605 | 0.217652 | 0.259839 | 0.357136 | 0.570107 | 1.183143 | 1.060456 | 0.496496 | 0.478604 | 0.232025 | 0.215140 | 0.240837 | 0.372044 | 0.452744 | 0.702885 | 1.034106 | 0.753516 | 0.360528 | 0.259754 | 0.217030 | 0.227791 | 0.213991 | 0.212096 | 0.211549 | 0.214517 | 0.262536 | 0.255051 | 0.213062 | 0.259088 | 0.421382 | 0.510841 | 0.553887 | 0.795645 | 0.860818 | 0.838519 | 0.840021 | 1.103046 | 1.893732 | 1.876770 | 3.372669 | 9.762125 | 39.654283 | 41.634268 | 17.918737 | 4.574462 | 2.844951 | 1.549148 | 0.840404 | 0.395436 | 0.307098 | 0.308657 | 0.359220 | 0.323213 | 0.231609 | 0.211834 | 0.222556 | 0.224391 | 0.262248 | 0.398118 | 0.489137 | 0.349631 | 0.295242 | 0.257191 | 0.238812 | 0.218749 | 0.220758 | 0.222576 | 0.223191 | 0.215001 | 0.234725 | 0.332835 | 0.425328 | 0.331861 | 0.373566 | 0.365202 | 0.333013 | 0.231172 | 0.211167 | 0.219600 | 0.223371 | 0.239681 | 0.249525 | 0.266393 | 0.269890 | 0.324546 | 0.456748 | 0.969916 | 1.701920 | 2.184584 | 1.672829 | 1.127540 | 0.821261 | 0.756959 | 0.576677 | 0.493816 | 0.428330 | 0.491961 | 0.465652 | 0.403184 | 0.293749 | 0.248815 | 0.245168 | 0.248571 | 0.250578 | 0.255909 | 0.286968 | 0.320044 | 0.336037 | 0.313265 | 0.313257 | 0.276139 | 0.223139 | 0.210806 | 0.212285 | 0.215996 | 0.217118 | 0.216611 | 0.214971 | 0.211464 | 0.211605 | 0.220592 | 0.230142 | 0.244714 | 0.262344 | 0.262886 | 0.239452 | 0.222815 | 0.217575 | 0.243375 | 0.263424 | 0.286998 | 0.275504 | 0.257964 | 0.236176 | 0.215250 | 0.219146 | 0.240844 | 0.252966 | 0.248966 | 0.236751 | 0.221915 | 0.218773 | 0.218633 | 0.216139 | 0.219125 | 0.223652 | 0.251762 | 0.298582 | 0.430280 | 0.490784 | 0.536231 | 0.517298 | 0.441137 | 0.381276 | 0.296325 | 0.246325 | 0.246099 | 0.274993 | 0.316272 | 0.402207 | 0.481705 | 0.763566 | 1.253633 | 2.280836 | 3.487834 | 4.214827 | 5.276362 | 7.372008 | 11.321011 | 9.969254 | 6.024792 | 3.160802 | 2.011722 | 1.172910 | 0.937757 | 0.774237 | 0.736262 | 0.874215 | 0.864542 | 1.046648 | 1.066136 | 0.679287 | 0.456943 | 0.343422 | 0.297509 | 0.370645 | 0.419142 | 0.480904 | 0.785693 | 1.530067 | 3.737873 | 6.706404 | 2.987878 | 2.250992 | 1.562129 | 0.779988 | 0.398978 | 0.272524 | 0.232100 | 0.232822 | 0.225062 | 0.226722 | 0.215982 | 0.213294 | 0.234176 | 0.239021 | 0.251136 | 0.275135 | 0.289071 | 0.252949 | 0.216344 | 0.218986 | 0.224588 | 0.225619 | 0.218891 | 0.213105 | 0.215264 | 0.238788 | 0.258578 | 0.280341 | 0.366095 | 0.546298 | 0.614868 | 0.534509 | 0.353077 | 0.262364 | 0.285467 | 0.287198 | 0.301610 | 0.407229 | 0.528495 | 0.615944 | 0.620505 | 0.372360 | 0.300168 | 0.271941 | 0.252458 | 0.225057 | 0.236190 | 0.247770 | 0.269343 | 0.254304 | 0.252075 | 0.255345 | 0.299144 | 0.301491 | 0.314451 | 0.320518 | 0.393110 | 0.490334 |
| right anterior | 0.216522 | 0.215596 | 0.236642 | 0.304565 | 0.583983 | 1.095538 | 0.840219 | 0.601794 | 0.411329 | 0.257561 | 0.226384 | 0.748809 | 3.616558 | 7.373053 | 22.221496 | 15.906243 | 11.795423 | 3.042422 | 0.804677 | 0.254160 | 0.263114 | 0.739784 | 3.188207 | 7.822152 | 8.377999 | 5.204286 | 3.632255 | 2.757909 | 1.476809 | 0.630908 | 0.375885 | 0.275931 | 0.216463 | 0.219968 | 0.226567 | 0.225840 | 0.233812 | 0.221816 | 0.239046 | 0.270838 | 0.306953 | 0.376952 | 0.544187 | 0.624813 | 0.513431 | 0.354060 | 0.230506 | 0.218825 | 0.230283 | 0.224430 | 0.247467 | 0.228237 | 0.220402 | 0.216253 | 0.214476 | 0.214783 | 0.213772 | 0.221247 | 0.230842 | 0.245007 | 0.303116 | 0.455531 | 0.469238 | 0.414224 | 0.460967 | 0.521636 | 0.716840 | 1.000406 | 1.125616 | 1.901651 | 2.941424 | 3.746125 | 5.680356 | 4.971802 | 3.427316 | 2.791064 | 1.817628 | 1.088369 | 0.587835 | 0.380278 | 0.301988 | 0.235764 | 0.215326 | 0.212511 | 0.254891 | 0.304948 | 0.400550 | 0.488550 | 0.458505 | 0.411907 | 0.375414 | 0.280012 | 0.244398 | 0.232563 | 0.218864 | 0.217875 | 0.219461 | 0.211054 | 0.210914 | 0.215304 | 0.244200 | 0.279861 | 0.314803 | 0.384193 | 0.438892 | 0.536795 | 0.569912 | 0.395832 | 0.301300 | 0.325747 | 0.289362 | 0.238273 | 0.233649 | 0.216048 | 0.211653 | 0.212658 | 0.213674 | 0.212183 | 0.210919 | 0.211538 | 0.211204 | 0.224646 | 0.260295 | 0.253489 | 0.227539 | 0.216073 | 0.241942 | 0.286809 | 0.445407 | 0.621509 | 0.783041 | 0.556951 | 0.292131 | 0.245636 | 0.246974 | 0.251862 | 0.354184 | 0.573953 | 0.999855 | 2.047263 | 3.471493 | 2.606278 | 1.761574 | 0.936636 | 0.567892 | 0.441940 | 0.379675 | 0.317579 | 0.282696 | 0.291159 | 0.266105 | 0.271108 | 0.262681 | 0.266202 | 0.267539 | 0.267643 | 0.245015 | 0.287639 | 0.390502 | 0.515500 | 0.368771 | 0.336738 | 0.313807 | 0.302289 | 0.256798 | 0.215515 | 0.213739 | 0.222323 | 0.211884 | 0.210867 | 0.215718 | 0.237741 | 0.269648 | 0.247243 | 0.218259 | 0.231064 | 0.266967 | 0.253377 | 0.283997 | 0.276176 | 0.238416 | 0.219306 | 0.211199 | 0.219552 | 0.235184 | 0.331921 | 0.473111 | 0.497069 | 0.393941 | 0.381729 | 0.385268 | 0.471734 | 0.712843 | 1.054956 | 2.085688 | 3.830178 | 4.557003 | 6.538705 | 10.886004 | 9.023967 | 5.997153 | 3.034987 | 2.459581 | 1.594125 | 1.049729 | 0.716268 | 0.605712 | 0.579207 | 0.876758 | 1.113062 | 1.543333 | 1.786143 | 1.529387 | 1.453663 | 1.651308 | 1.314771 | 1.408864 | 1.714372 | 1.952189 | 2.585882 | 2.589102 | 2.910972 | 3.789391 | 4.389417 | 2.787590 | 1.819265 | 1.297885 | 0.943154 | 0.787260 | 0.784217 | 0.726428 | 0.609323 | 0.432993 | 0.354866 | 0.355719 | 0.304302 | 0.246277 | 0.225470 | 0.232398 | 0.260574 | 0.270878 | 0.279195 | 0.288618 | 0.264127 | 0.230498 | 0.212033 | 0.213715 | 0.233608 | 0.275416 | 0.296597 | 0.302466 | 0.258699 | 0.253392 | 0.261774 | 0.269909 | 0.304002 | 0.367171 | 0.339860 | 0.352943 | 0.352414 | 0.340372 | 0.359617 | 0.393190 | 0.434744 | 0.502437 | 0.528468 | 0.482450 | 0.406568 | 0.307631 | 0.247639 | 0.227804 | 0.214693 | 0.212432 | 0.210831 | 0.212123 | 0.211855 | 0.211098 | 0.218562 | 0.220542 | 0.223203 |
| left central | 0.218281 | 0.213980 | 0.299446 | 0.369033 | 0.374834 | 0.494818 | 0.865477 | 0.996626 | 0.714861 | 0.540591 | 0.923990 | 0.843168 | 0.352623 | 0.257461 | 0.236892 | 0.280906 | 0.292138 | 0.234200 | 0.214449 | 0.213993 | 0.215097 | 0.237652 | 0.230210 | 0.257561 | 0.239423 | 0.257701 | 0.279098 | 0.293399 | 0.249860 | 0.266846 | 0.342077 | 1.086697 | 7.716260 | 23.089160 | 17.272509 | 11.889187 | 4.228425 | 0.716311 | 0.258884 | 0.212734 | 0.287172 | 0.366130 | 0.502292 | 0.597817 | 0.324482 | 0.216443 | 0.210998 | 0.210850 | 0.210817 | 0.212468 | 0.217498 | 0.217925 | 0.213545 | 0.232590 | 0.257334 | 0.251413 | 0.249523 | 0.218857 | 0.212066 | 0.288160 | 0.483277 | 0.792435 | 0.909628 | 0.843845 | 0.770219 | 1.198589 | 1.627543 | 4.624993 | 7.309837 | 11.662233 | 11.847749 | 10.739588 | 9.247661 | 3.451704 | 0.992739 | 0.429534 | 0.284008 | 0.233602 | 0.238226 | 0.232823 | 0.222105 | 0.212566 | 0.211862 | 0.265093 | 0.512147 | 2.937807 | 5.517419 | 5.823949 | 6.391616 | 7.060830 | 4.271011 | 2.968763 | 1.611541 | 1.226534 | 0.770598 | 0.742140 | 0.606595 | 0.372070 | 0.274877 | 0.249269 | 0.436307 | 0.819879 | 0.642461 | 0.542735 | 0.703308 | 0.985892 | 1.497927 | 0.927340 | 0.723429 | 0.540458 | 0.543259 | 0.478604 | 0.366869 | 0.314717 | 0.260876 | 0.218108 | 0.212191 | 0.216382 | 0.221659 | 0.217550 | 0.223544 | 0.210844 | 0.223384 | 0.249610 | 0.278585 | 0.244305 | 0.230175 | 0.228882 | 0.213016 | 0.211483 | 0.218910 | 0.217559 | 0.211344 | 0.211932 | 0.212281 | 0.215150 | 0.221773 | 0.234715 | 0.240567 | 0.224241 | 0.223543 | 0.238006 | 0.258969 | 0.264945 | 0.241541 | 0.217554 | 0.214661 | 0.215142 | 0.223676 | 0.249753 | 0.273448 | 0.339805 | 0.498668 | 0.696167 | 0.669200 | 0.488746 | 0.328670 | 0.375486 | 0.463952 | 0.746635 | 1.644909 | 3.909625 | 6.891547 | 15.545209 | 20.864421 | 21.741863 | 16.946111 | 18.999869 | 47.521816 | 130.014354 | 88.242171 | 78.454893 | 35.941535 | 17.670475 | 7.477178 | 3.516531 | 1.273852 | 0.873561 | 0.780273 | 1.126177 | 1.955803 | 2.076836 | 2.362680 | 3.125337 | 3.429376 | 3.779350 | 3.320899 | 2.173655 | 1.767145 | 1.285104 | 0.687226 | 0.367515 | 0.242684 | 0.213549 | 0.211820 | 0.212505 | 0.214859 | 0.211617 | 0.215829 | 0.245188 | 0.382030 | 0.820382 | 1.504594 | 2.826780 | 6.422261 | 9.709400 | 10.655033 | 6.792306 | 2.842983 | 1.488161 | 0.702107 | 0.532524 | 0.508623 | 0.368676 | 0.280555 | 0.254650 | 0.283226 | 0.391431 | 0.512794 | 0.450161 | 0.541322 | 1.072424 | 3.286703 | 4.674202 | 4.047348 | 3.001943 | 2.481276 | 1.752265 | 1.089612 | 0.485871 | 0.311086 | 0.272893 | 0.249620 | 0.266278 | 0.396601 | 0.339242 | 0.292945 | 0.280406 | 0.298499 | 0.274649 | 0.273311 | 0.246521 | 0.271546 | 0.320162 | 0.298856 | 0.259241 | 0.258043 | 0.232932 | 0.229273 | 0.220392 | 0.213463 | 0.213076 | 0.218984 | 0.216536 | 0.213543 | 0.211172 | 0.214375 | 0.221683 | 0.229037 | 0.260052 | 0.256665 | 0.232822 | 0.228054 | 0.215920 | 0.210872 | 0.211299 | 0.217564 | 0.235891 | 0.259212 | 0.316944 | 0.374108 | 0.411283 | 0.769534 | 0.586657 | 0.297587 | 0.218075 | 0.213305 | 0.224610 | 0.247645 | 0.437912 |
| right central | 1.592029 | 1.975867 | 1.159548 | 1.540437 | 2.178641 | 1.954238 | 1.629392 | 0.920486 | 0.942325 | 1.255826 | 0.514514 | 0.320939 | 0.221905 | 0.215461 | 0.216086 | 0.212063 | 0.296816 | 0.605776 | 0.951941 | 1.432054 | 0.886809 | 0.491760 | 0.367546 | 0.234452 | 0.213862 | 0.211018 | 0.212083 | 0.223906 | 0.361179 | 0.425877 | 0.334549 | 0.230366 | 0.212069 | 0.217281 | 0.222134 | 0.301596 | 0.472062 | 0.462771 | 0.354116 | 0.308727 | 0.222097 | 0.229769 | 0.211979 | 0.283230 | 0.548665 | 0.878166 | 0.599419 | 0.337689 | 0.368598 | 0.300326 | 0.219751 | 0.240780 | 0.409077 | 0.453434 | 0.356887 | 0.382074 | 0.405940 | 0.434940 | 0.675613 | 0.781731 | 0.689401 | 0.798207 | 0.893232 | 0.744301 | 0.691870 | 0.558051 | 0.941484 | 2.040354 | 3.815267 | 8.109816 | 43.454300 | 61.056320 | 69.504872 | 33.554883 | 10.667693 | 3.958143 | 0.995597 | 0.529755 | 0.299306 | 0.214662 | 0.225635 | 0.267368 | 0.359786 | 0.374924 | 0.366395 | 0.396937 | 0.498289 | 0.595616 | 0.723806 | 0.471975 | 0.389141 | 0.411871 | 0.407296 | 0.308777 | 0.244241 | 0.220254 | 0.214487 | 0.211853 | 0.221146 | 0.265210 | 0.342434 | 0.396578 | 0.446590 | 0.359131 | 0.286157 | 0.225389 | 0.222853 | 0.213753 | 0.213051 | 0.218725 | 0.237563 | 0.231598 | 0.210814 | 0.213595 | 0.216623 | 0.233511 | 0.257997 | 0.384446 | 0.480263 | 0.383455 | 0.305394 | 0.258601 | 0.230037 | 0.211435 | 0.220716 | 0.240023 | 0.269838 | 0.304130 | 0.308793 | 0.370362 | 0.420054 | 0.416088 | 0.480724 | 0.476555 | 0.508429 | 0.529665 | 0.464372 | 0.456349 | 0.423969 | 0.323953 | 0.272079 | 0.248132 | 0.238823 | 0.263105 | 0.295318 | 0.401651 | 0.586305 | 0.892767 | 1.480080 | 2.660123 | 2.782692 | 4.278824 | 8.395234 | 14.154500 | 26.801378 | 27.378826 | 19.504835 | 17.898650 | 9.626798 | 3.935158 | 1.306665 | 1.061077 | 1.172125 | 0.825493 | 0.869015 | 0.633234 | 0.480595 | 0.435428 | 0.390888 | 0.271993 | 0.251562 | 0.235697 | 0.249374 | 0.344253 | 0.459159 | 0.468075 | 0.503004 | 0.655943 | 1.025489 | 1.245217 | 0.842722 | 1.032668 | 1.337205 | 1.804119 | 2.482669 | 5.471724 | 9.778530 | 9.703209 | 7.753095 | 6.182654 | 7.541041 | 6.527469 | 2.376319 | 0.823282 | 0.666407 | 0.571578 | 0.405271 | 0.240972 | 0.214693 | 0.280657 | 0.324640 | 0.313036 | 0.293815 | 0.244201 | 0.213204 | 0.241739 | 0.394008 | 0.523411 | 0.478953 | 0.436082 | 0.376669 | 0.344069 | 0.357106 | 0.339275 | 0.306476 | 0.268895 | 0.234732 | 0.226407 | 0.236221 | 0.221913 | 0.213084 | 0.210827 | 0.212428 | 0.222410 | 0.224092 | 0.244424 | 0.233646 | 0.217765 | 0.219008 | 0.229261 | 0.241212 | 0.327829 | 0.348028 | 0.379926 | 0.573237 | 0.871083 | 1.209860 | 1.460921 | 0.951835 | 0.480832 | 0.334942 | 0.242838 | 0.221956 | 0.219750 | 0.221507 | 0.231995 | 0.277252 | 0.349351 | 0.406378 | 0.381778 | 0.307913 | 0.236444 | 0.212030 | 0.212524 | 0.229228 | 0.252361 | 0.270484 | 0.292755 | 0.236984 | 0.212495 | 0.210819 | 0.211579 | 0.211114 | 0.210818 | 0.215827 | 0.211525 | 0.212798 | 0.213957 | 0.214128 | 0.211589 | 0.221469 | 0.221016 | 0.221018 | 0.227461 | 0.239499 | 0.249623 | 0.263585 | 0.263264 | 0.276195 | 0.339946 |
| left posterior | 0.393410 | 0.529487 | 0.685898 | 0.594505 | 0.435611 | 0.361264 | 0.434142 | 0.509925 | 0.471781 | 0.386907 | 0.349007 | 0.314452 | 0.240862 | 0.229376 | 0.366167 | 0.622563 | 0.792503 | 1.040542 | 3.431654 | 1.590927 | 0.389311 | 0.212885 | 0.211707 | 0.212035 | 0.212240 | 0.218219 | 0.210839 | 0.335830 | 1.117811 | 1.576645 | 1.054461 | 0.270589 | 0.227823 | 0.361536 | 0.745699 | 1.968234 | 2.628199 | 1.421641 | 0.432053 | 0.211103 | 0.245964 | 0.308590 | 0.460347 | 0.645614 | 0.847903 | 1.159880 | 0.806950 | 0.701892 | 0.560127 | 0.680385 | 0.766407 | 1.210491 | 2.128577 | 6.672814 | 30.047235 | 43.810344 | 8.497938 | 1.533324 | 0.349671 | 0.212752 | 0.264715 | 0.528024 | 1.098529 | 2.368002 | 8.277727 | 41.325716 | 92.226162 | 268.346171 | 793.158885 | 1713.554527 | 4492.519861 | 5681.303092 | 2675.460029 | 323.236142 | 52.789001 | 11.040578 | 4.672188 | 1.862029 | 1.266848 | 0.726992 | 0.541536 | 0.342329 | 0.231227 | 0.212197 | 0.245339 | 0.305700 | 0.315925 | 0.291103 | 0.278777 | 0.267888 | 0.231114 | 0.213180 | 0.211295 | 0.210892 | 0.215149 | 0.288613 | 0.500229 | 0.634314 | 0.697847 | 0.972850 | 0.963517 | 0.766659 | 0.310851 | 0.211577 | 0.230576 | 0.300123 | 0.509943 | 0.883053 | 1.211193 | 0.862415 | 0.369660 | 0.266453 | 0.222098 | 0.213921 | 0.266665 | 0.389703 | 0.617035 | 0.601535 | 0.644303 | 0.718597 | 0.470921 | 0.253542 | 0.211568 | 0.225011 | 0.229098 | 0.228935 | 0.222667 | 0.211084 | 0.224407 | 0.256353 | 0.278401 | 0.266140 | 0.269062 | 0.263761 | 0.232886 | 0.222587 | 0.226824 | 0.234339 | 0.261113 | 0.256550 | 0.252071 | 0.241511 | 0.220120 | 0.210816 | 0.217651 | 0.251146 | 0.299980 | 0.438417 | 0.493165 | 0.337158 | 0.247055 | 0.216498 | 0.211622 | 0.211015 | 0.211640 | 0.224566 | 0.236122 | 0.230596 | 0.221597 | 0.231530 | 0.267451 | 0.314544 | 0.326192 | 0.343225 | 0.407633 | 0.589097 | 0.746910 | 0.811370 | 0.870398 | 0.831361 | 0.750991 | 0.679212 | 0.545199 | 0.476405 | 0.401945 | 0.299165 | 0.278018 | 0.236642 | 0.211761 | 0.211803 | 0.215148 | 0.213644 | 0.214961 | 0.232200 | 0.283831 | 0.424143 | 0.788107 | 1.081627 | 1.667859 | 1.653682 | 1.810361 | 2.030232 | 2.286886 | 2.223738 | 2.651925 | 2.491121 | 1.854852 | 1.270680 | 1.028710 | 0.839454 | 0.728702 | 0.626020 | 0.534116 | 0.631197 | 0.666254 | 0.566380 | 0.522895 | 0.488788 | 0.504173 | 0.473715 | 0.481449 | 0.634382 | 0.931084 | 1.186840 | 0.977756 | 0.540853 | 0.351894 | 0.267702 | 0.247333 | 0.227673 | 0.215457 | 0.214240 | 0.239480 | 0.299768 | 0.359008 | 0.354574 | 0.329615 | 0.331339 | 0.335410 | 0.311877 | 0.265474 | 0.238799 | 0.222501 | 0.217594 | 0.214724 | 0.212397 | 0.212226 | 0.212022 | 0.210845 | 0.214729 | 0.230165 | 0.248115 | 0.285416 | 0.389117 | 0.465714 | 0.423586 | 0.410293 | 0.359089 | 0.316211 | 0.270743 | 0.250976 | 0.237844 | 0.229900 | 0.215818 | 0.212259 | 0.211863 | 0.220589 | 0.238427 | 0.257411 | 0.251924 | 0.244847 | 0.233431 | 0.221096 | 0.211648 | 0.211429 | 0.216192 | 0.216791 | 0.215876 | 0.214037 | 0.212895 | 0.215080 | 0.213789 | 0.210838 | 0.218165 | 0.225961 | 0.221287 | 0.216430 | 0.226816 | 0.219875 | 0.211067 |
| right posterior | 0.451148 | 0.443192 | 0.295316 | 0.214673 | 0.242802 | 0.365549 | 0.456946 | 0.399184 | 0.294378 | 0.220126 | 0.222493 | 0.431225 | 0.557401 | 1.111107 | 2.070790 | 1.144106 | 0.606267 | 0.594178 | 0.693777 | 0.541689 | 0.300827 | 0.214735 | 0.214685 | 0.212693 | 0.248892 | 0.448292 | 1.110169 | 1.878195 | 2.790863 | 3.270453 | 1.771775 | 0.748351 | 0.349768 | 0.245095 | 0.226050 | 0.242844 | 0.272831 | 0.302797 | 0.378449 | 0.401934 | 0.347147 | 0.399291 | 0.359464 | 0.309680 | 0.260587 | 0.230239 | 0.211405 | 0.211256 | 0.223887 | 0.218018 | 0.219920 | 0.218781 | 0.214963 | 0.212269 | 0.216888 | 0.231515 | 0.212030 | 0.275486 | 0.375530 | 0.553357 | 0.729857 | 1.561501 | 4.458971 | 38.290978 | 259.460407 | 825.470222 | 1181.272259 | 3935.042156 | 12590.164503 | 60676.539377 | 186276.870081 | 322802.412090 | 451059.785965 | 178158.993063 | 15976.546996 | 650.538539 | 29.789259 | 4.027226 | 0.683250 | 0.246290 | 0.211488 | 0.223360 | 0.264764 | 0.261739 | 0.271193 | 0.251956 | 0.226620 | 0.224091 | 0.234178 | 0.217332 | 0.211379 | 0.212542 | 0.226366 | 0.234125 | 0.233379 | 0.233962 | 0.248422 | 0.279963 | 0.302789 | 0.325934 | 0.368551 | 0.466490 | 0.692400 | 0.687239 | 0.635696 | 0.499685 | 0.426264 | 0.382185 | 0.368254 | 0.305519 | 0.283757 | 0.270929 | 0.269576 | 0.290357 | 0.320363 | 0.388322 | 0.553997 | 0.795878 | 1.075062 | 1.466808 | 1.398626 | 1.342265 | 1.036537 | 0.629821 | 0.405793 | 0.345128 | 0.334425 | 0.333747 | 0.339867 | 0.332356 | 0.359441 | 0.425098 | 0.394791 | 0.402463 | 0.522717 | 0.705532 | 1.161683 | 2.632939 | 5.861027 | 17.858776 | 47.748559 | 57.723635 | 61.606820 | 61.536280 | 27.694455 | 9.562684 | 5.220766 | 3.448573 | 1.980504 | 1.312306 | 0.773076 | 0.669677 | 0.590292 | 0.513044 | 0.533566 | 0.861471 | 1.763144 | 4.823939 | 12.728032 | 41.862222 | 118.540640 | 131.564212 | 77.819784 | 42.167400 | 27.248141 | 31.679094 | 30.189525 | 29.399310 | 28.784800 | 20.361366 | 27.932492 | 35.559925 | 27.877383 | 23.447975 | 31.657093 | 43.635972 | 73.177310 | 84.512452 | 75.685410 | 49.613769 | 30.115787 | 11.754542 | 6.267869 | 4.886571 | 4.000325 | 4.527051 | 6.548304 | 10.881585 | 20.961031 | 47.322446 | 68.743708 | 66.842015 | 67.323044 | 50.093581 | 37.515542 | 38.230255 | 41.737868 | 55.174127 | 101.187735 | 119.037203 | 119.746194 | 97.270617 | 51.621033 | 16.767994 | 11.623004 | 11.438455 | 10.418035 | 14.006989 | 13.771441 | 19.566622 | 35.621102 | 46.245910 | 44.007650 | 82.478788 | 143.501078 | 288.401186 | 308.134077 | 184.994570 | 70.716597 | 27.269562 | 13.187206 | 6.665815 | 3.571694 | 2.596419 | 2.165639 | 1.698146 | 1.592839 | 1.118628 | 0.794094 | 0.603984 | 0.466651 | 0.395598 | 0.367480 | 0.338106 | 0.355151 | 0.393354 | 0.439294 | 0.506770 | 0.725815 | 1.147416 | 1.637878 | 2.175651 | 2.410471 | 2.147250 | 1.762673 | 1.537580 | 1.419021 | 1.689603 | 2.376322 | 2.980263 | 3.540633 | 4.196974 | 4.319774 | 3.497500 | 2.334138 | 1.454855 | 1.157366 | 1.240041 | 1.112329 | 0.935503 | 0.881147 | 0.877798 | 0.875589 | 1.101210 | 1.562976 | 1.748904 | 1.984830 | 2.125715 | 2.219868 | 2.972132 | 2.774177 | 2.404149 | 4.179344 | 9.479251 | 36.851377 | 120.023658 | 138.024812 | 183.290802 | 228.468205 | 246.395347 |
| all electrodes | 0.301189 | 0.263028 | 0.490151 | 0.851572 | 0.803156 | 0.990468 | 1.032245 | 1.409471 | 2.605576 | 1.970945 | 1.212180 | 0.434607 | 0.234554 | 0.231372 | 0.239963 | 0.214386 | 0.220690 | 0.259560 | 0.232444 | 0.210949 | 0.213422 | 0.254756 | 0.269987 | 0.242628 | 0.267011 | 0.229049 | 0.230385 | 0.311794 | 0.536353 | 0.738512 | 0.512016 | 0.334403 | 0.216644 | 0.239748 | 0.285851 | 0.381826 | 0.507659 | 0.616073 | 0.465945 | 0.421559 | 0.316920 | 0.249789 | 0.211140 | 0.251843 | 0.289672 | 0.373121 | 0.541604 | 0.582944 | 0.562922 | 0.815866 | 0.347036 | 0.232468 | 0.211304 | 0.210806 | 0.213167 | 0.221127 | 0.274195 | 0.344889 | 0.332776 | 0.475784 | 1.274750 | 3.641243 | 6.058062 | 11.228330 | 57.044609 | 327.795042 | 1017.429853 | 1712.163932 | 5157.588738 | 31102.527245 | 135398.357808 | 154796.356785 | 140250.972990 | 80744.382151 | 41092.895477 | 6185.591773 | 590.962995 | 61.915378 | 8.651562 | 1.277778 | 0.368790 | 0.230650 | 0.210966 | 0.261641 | 0.435724 | 0.723042 | 1.381228 | 1.665239 | 1.422439 | 1.109101 | 0.548216 | 0.279292 | 0.227975 | 0.210818 | 0.227464 | 0.276900 | 0.393749 | 0.482450 | 0.444285 | 0.397684 | 0.394713 | 0.414759 | 0.303987 | 0.238886 | 0.237635 | 0.230945 | 0.248782 | 0.246093 | 0.236696 | 0.272278 | 0.308538 | 0.273262 | 0.278938 | 0.280178 | 0.279979 | 0.294038 | 0.312105 | 0.327165 | 0.367205 | 0.353946 | 0.393234 | 0.463739 | 0.518565 | 0.319438 | 0.233482 | 0.212797 | 0.210885 | 0.214465 | 0.221205 | 0.229209 | 0.212961 | 0.216320 | 0.280032 | 0.368364 | 0.580191 | 0.939013 | 1.688019 | 1.826353 | 1.912578 | 1.255151 | 0.824398 | 0.566830 | 0.368645 | 0.289407 | 0.250401 | 0.226885 | 0.215767 | 0.210938 | 0.210885 | 0.211941 | 0.210971 | 0.211482 | 0.210966 | 0.210898 | 0.224874 | 0.254456 | 0.309685 | 0.479099 | 0.542051 | 0.384592 | 0.355157 | 0.321016 | 0.281921 | 0.280256 | 0.262271 | 0.298489 | 0.415857 | 0.490899 | 0.419164 | 0.366819 | 0.371466 | 0.431834 | 0.525741 | 0.630184 | 0.928811 | 2.805226 | 6.683387 | 6.749228 | 4.773633 | 2.662769 | 2.130036 | 1.930884 | 1.206752 | 1.118927 | 1.321902 | 1.517272 | 2.977999 | 3.583468 | 6.055677 | 11.693953 | 24.925825 | 33.794152 | 148.780474 | 403.583299 | 1293.439341 | 1456.422372 | 2435.916574 | 3210.443347 | 7082.103648 | 8484.156089 | 8586.530913 | 3993.883577 | 1988.222613 | 633.021486 | 254.208053 | 74.927389 | 23.199575 | 10.552659 | 9.168605 | 8.417321 | 8.984984 | 10.457964 | 15.393399 | 21.811569 | 27.692795 | 23.550707 | 17.171716 | 8.307159 | 5.872890 | 5.042837 | 5.144084 | 3.944227 | 3.119630 | 1.889357 | 1.491981 | 0.981534 | 0.645931 | 0.558859 | 0.553094 | 0.649348 | 1.237525 | 2.673517 | 5.149237 | 8.935925 | 9.796965 | 9.775381 | 6.078828 | 3.218139 | 2.902778 | 3.045469 | 2.694589 | 2.394854 | 3.004947 | 3.996788 | 6.068601 | 6.503531 | 7.578846 | 7.563684 | 6.469835 | 3.181713 | 2.626059 | 1.996371 | 1.849484 | 1.886875 | 1.518122 | 1.207123 | 1.158570 | 0.830232 | 0.687657 | 0.539482 | 0.419113 | 0.371227 | 0.372061 | 0.395995 | 0.447729 | 0.496203 | 0.562731 | 0.684607 | 0.931612 | 1.022058 | 1.036836 | 1.077729 | 1.093841 | 1.211346 | 1.364868 | 1.525914 | 1.839891 | 1.980661 | 1.930141 | 1.717439 |

Searchlight, spatiotemporal cluster permutation test

|  | start time | stop time | peak time | peak channel | cluster p | peak Cohen's d | direction |
| --- | --- | --- | --- | --- | --- | --- | --- |
| #1 | 85 | 200 | 150 | P4 | 0.0191 | 1.728388 | positive |
| #2 | 430 | 1195 | 885 | PO8 | 0.0058 | 0.978394 | positive |

H) real faces (LOSO) - emotion

  
|  | time window | peak latency | cluster *p* | peak Cohen's *d* |  | | | |  |
| **all electrodes** | 120 - 225 ms | 170 ms | 0.0149 | 1.1992 |  | | | |  |
 245 - 335 ms | 280 ms | 0.0326 | 0.9169 |  | | | | 345 - 430 ms | 410 ms | 0.0437 | 0.7775 |  | | | ||  |  | | | | | | | | |

Time-resolved classification, cluster permutation tests

|  |  | **left hemisphere** | | | | **right hemisphere** | | | |
|  | time window | peak latency | cluster *p* | peak Cohen's *d* | time window | peak latency | cluster *p* | peak Cohen's *d* |  |
| **anterior** |  | | | | 320 - 430 ms | 350 ms | 0.0324 | 0.7676 |  |
| **central** |  | | | | 125 - 470 ms | 270 ms | 0.0002 | 1.3182 |  |
| **posterior** | 110 - 535 ms | 150 ms | 0.0012 | 1.9544 | 90 - 665 ms | 270 ms | 0.0007 | 1.0842 |  |
 840 - 1025 ms | 890 ms | 0.022 | 0.61 | 685 - 1125 ms | 890 ms | 0.0029 | 0.9069 |

  

Time-resolved classification, Bayesian statistics

|  | -200 | -195 | -190 | -185 | -180 | -175 | -170 | -165 | -160 | -155 | -150 | -145 | -140 | -135 | -130 | -125 | -120 | -115 | -110 | -105 | -100 | -95 | -90 | -85 | -80 | -75 | -70 | -65 | -60 | -55 | -50 | -45 | -40 | -35 | -30 | -25 | -20 | -15 | -10 | -5 | 0 | 5 | 10 | 15 | 20 | 25 | 30 | 35 | 40 | 45 | 50 | 55 | 60 | 65 | 70 | 75 | 80 | 85 | 90 | 95 | 100 | 105 | 110 | 115 | 120 | 125 | 130 | 135 | 140 | 145 | 150 | 155 | 160 | 165 | 170 | 175 | 180 | 185 | 190 | 195 | 200 | 205 | 210 | 215 | 220 | 225 | 230 | 235 | 240 | 245 | 250 | 255 | 260 | 265 | 270 | 275 | 280 | 285 | 290 | 295 | 300 | 305 | 310 | 315 | 320 | 325 | 330 | 335 | 340 | 345 | 350 | 355 | 360 | 365 | 370 | 375 | 380 | 385 | 390 | 395 | 400 | 405 | 410 | 415 | 420 | 425 | 430 | 435 | 440 | 445 | 450 | 455 | 460 | 465 | 470 | 475 | 480 | 485 | 490 | 495 | 500 | 505 | 510 | 515 | 520 | 525 | 530 | 535 | 540 | 545 | 550 | 555 | 560 | 565 | 570 | 575 | 580 | 585 | 590 | 595 | 600 | 605 | 610 | 615 | 620 | 625 | 630 | 635 | 640 | 645 | 650 | 655 | 660 | 665 | 670 | 675 | 680 | 685 | 690 | 695 | 700 | 705 | 710 | 715 | 720 | 725 | 730 | 735 | 740 | 745 | 750 | 755 | 760 | 765 | 770 | 775 | 780 | 785 | 790 | 795 | 800 | 805 | 810 | 815 | 820 | 825 | 830 | 835 | 840 | 845 | 850 | 855 | 860 | 865 | 870 | 875 | 880 | 885 | 890 | 895 | 900 | 905 | 910 | 915 | 920 | 925 | 930 | 935 | 940 | 945 | 950 | 955 | 960 | 965 | 970 | 975 | 980 | 985 | 990 | 995 | 1000 | 1005 | 1010 | 1015 | 1020 | 1025 | 1030 | 1035 | 1040 | 1045 | 1050 | 1055 | 1060 | 1065 | 1070 | 1075 | 1080 | 1085 | 1090 | 1095 | 1100 | 1105 | 1110 | 1115 | 1120 | 1125 | 1130 | 1135 | 1140 | 1145 | 1150 | 1155 | 1160 | 1165 | 1170 | 1175 | 1180 | 1185 | 1190 | 1195 |
| --- | --- | --- | --- | --- | --- | --- | --- | --- | --- | --- | --- | --- | --- | --- | --- | --- | --- | --- | --- | --- | --- | --- | --- | --- | --- | --- | --- | --- | --- | --- | --- | --- | --- | --- | --- | --- | --- | --- | --- | --- | --- | --- | --- | --- | --- | --- | --- | --- | --- | --- | --- | --- | --- | --- | --- | --- | --- | --- | --- | --- | --- | --- | --- | --- | --- | --- | --- | --- | --- | --- | --- | --- | --- | --- | --- | --- | --- | --- | --- | --- | --- | --- | --- | --- | --- | --- | --- | --- | --- | --- | --- | --- | --- | --- | --- | --- | --- | --- | --- | --- | --- | --- | --- | --- | --- | --- | --- | --- | --- | --- | --- | --- | --- | --- | --- | --- | --- | --- | --- | --- | --- | --- | --- | --- | --- | --- | --- | --- | --- | --- | --- | --- | --- | --- | --- | --- | --- | --- | --- | --- | --- | --- | --- | --- | --- | --- | --- | --- | --- | --- | --- | --- | --- | --- | --- | --- | --- | --- | --- | --- | --- | --- | --- | --- | --- | --- | --- | --- | --- | --- | --- | --- | --- | --- | --- | --- | --- | --- | --- | --- | --- | --- | --- | --- | --- | --- | --- | --- | --- | --- | --- | --- | --- | --- | --- | --- | --- | --- | --- | --- | --- | --- | --- | --- | --- | --- | --- | --- | --- | --- | --- | --- | --- | --- | --- | --- | --- | --- | --- | --- | --- | --- | --- | --- | --- | --- | --- | --- | --- | --- | --- | --- | --- | --- | --- | --- | --- | --- | --- | --- | --- | --- | --- | --- | --- | --- | --- | --- | --- | --- | --- | --- | --- | --- | --- | --- | --- | --- | --- | --- | --- | --- | --- | --- | --- | --- | --- | --- | --- | --- | --- | --- | --- | --- | --- | --- | --- | --- | --- | --- |
| left anterior | 0.249447 | 0.250807 | 0.248401 | 0.267471 | 0.253214 | 0.316832 | 0.288002 | 0.226914 | 0.242739 | 0.231438 | 0.214626 | 0.220910 | 0.217228 | 0.224840 | 0.235645 | 0.246754 | 0.269874 | 0.251961 | 0.241724 | 0.215427 | 0.214665 | 0.241338 | 0.263789 | 0.352125 | 0.702053 | 2.640009 | 6.731508 | 10.020962 | 5.012675 | 3.831471 | 1.986756 | 0.712248 | 0.215614 | 0.356506 | 0.973740 | 2.528500 | 8.275667 | 13.836477 | 2.333343 | 0.591987 | 0.304299 | 0.514897 | 0.869452 | 1.157003 | 1.346315 | 1.936545 | 3.192563 | 19.851330 | 31.934163 | 18.832712 | 18.455357 | 7.011618 | 3.406243 | 2.303253 | 1.128724 | 0.477614 | 0.281194 | 0.227970 | 0.221514 | 0.224814 | 0.244587 | 0.307060 | 0.303432 | 0.260770 | 0.215252 | 0.218207 | 0.245388 | 2.791346e-01 | 4.323004e-01 | 5.298500e-01 | 5.044683e-01 | 3.506814e-01 | 0.323049 | 0.290468 | 0.316597 | 0.254217 | 0.217996 | 0.403869 | 0.928133 | 4.320102 | 12.882760 | 18.377680 | 10.730822 | 4.270157 | 1.202479 | 0.393879 | 0.236877 | 0.220183 | 0.307358 | 0.873577 | 3.569675 | 8.518581 | 6.458934 | 7.645720 | 9.012092 | 10.677854 | 5.099296 | 1.789357 | 1.541176 | 2.849167 | 2.556837 | 1.268992 | 0.663295 | 0.371743 | 0.308113 | 0.234269 | 0.214870 | 0.218870 | 0.216212 | 0.255301 | 0.424902 | 0.756040 | 1.161207 | 1.295045 | 1.094236 | 0.721977 | 0.488273 | 0.481521 | 0.411819 | 0.451306 | 0.459215 | 0.460482 | 0.412309 | 0.444729 | 0.370828 | 0.375111 | 0.345658 | 0.260414 | 0.265592 | 0.260345 | 0.242231 | 0.223130 | 0.214903 | 0.215862 | 0.216279 | 0.236117 | 0.273238 | 0.272162 | 0.273165 | 0.232244 | 0.216344 | 0.222916 | 0.245512 | 0.314134 | 0.314523 | 0.399226 | 0.332635 | 0.349014 | 0.345024 | 0.366963 | 0.294519 | 0.262834 | 0.231915 | 0.218630 | 0.227662 | 0.272394 | 0.311016 | 0.278792 | 0.235612 | 0.219128 | 0.216173 | 0.257922 | 0.309006 | 0.397833 | 0.450662 | 0.488138 | 0.575365 | 0.478598 | 0.378215 | 0.350357 | 0.278767 | 0.245506 | 0.232944 | 0.224317 | 0.227532 | 0.229022 | 0.232129 | 0.252075 | 0.296710 | 0.354332 | 0.470231 | 0.605377 | 0.645572 | 0.628342 | 0.723259 | 0.816786 | 0.814831 | 0.577141 | 0.394102 | 0.405321 | 0.410337 | 0.317938 | 0.311274 | 0.358307 | 0.497852 | 0.918798 | 1.040661 | 1.026774 | 1.214456 | 1.318834 | 0.925089 | 0.664522 | 0.453697 | 0.362053 | 0.318450 | 0.272660 | 0.237793 | 0.234070 | 0.238034 | 0.269893 | 0.311808 | 0.379527 | 0.476412 | 0.605633 | 0.712606 | 1.247625 | 1.661634 | 2.378683 | 3.412085 | 5.569663 | 6.219846 | 10.533098 | 5.970180 | 4.386873 | 3.055252 | 2.461215 | 1.330475 | 0.854978 | 0.838426 | 1.093312 | 1.063241 | 0.991478 | 0.760288 | 0.677259 | 0.652117 | 0.508326 | 0.521029 | 0.730430 | 0.763798 | 0.783159 | 0.791132 | 1.007223 | 1.616143 | 2.324154 | 1.810965 | 2.304710 | 2.900951 | 3.861002 | 3.925891 | 1.764132 | 1.194661 | 1.000365 | 1.085351 | 2.107275 | 3.588787 | 4.468265 | 3.736221 | 2.300389 | 1.869835 | 1.432092 | 0.920339 | 0.715297 | 0.637556 | 0.639028 | 0.600369 | 0.824252 | 0.734580 | 0.687120 | 0.928013 | 1.026262 | 0.787706 | 0.480916 | 0.347792 | 0.341742 | 0.355685 | 0.307228 | 0.258231 | 0.261123 | 0.333662 | 0.373024 |
| right anterior | 0.412154 | 0.407008 | 0.304850 | 0.296168 | 0.235989 | 0.231516 | 0.238225 | 0.216946 | 0.215054 | 0.223469 | 0.219170 | 0.233599 | 0.238923 | 0.381683 | 0.419601 | 0.561534 | 0.399430 | 0.303139 | 0.226679 | 0.215873 | 0.230641 | 0.281646 | 0.321018 | 0.500394 | 0.565283 | 0.610513 | 0.746082 | 2.435664 | 5.372147 | 2.131875 | 0.534810 | 0.284999 | 0.217420 | 0.264281 | 0.454816 | 1.430236 | 1.584340 | 0.691254 | 0.577099 | 0.538245 | 0.482238 | 0.773662 | 0.585674 | 0.545559 | 0.642945 | 0.754762 | 0.703273 | 0.701851 | 0.353591 | 0.295656 | 0.368805 | 0.442528 | 0.718294 | 0.974073 | 0.647772 | 0.914640 | 1.282901 | 0.677513 | 0.533147 | 0.249376 | 0.216224 | 0.216100 | 0.234126 | 0.287527 | 0.353358 | 0.550375 | 0.707928 | 1.107774e+00 | 8.776582e-01 | 9.549388e-01 | 8.133322e-01 | 9.826051e-01 | 1.041745 | 0.747626 | 0.511897 | 0.471894 | 0.324953 | 0.243559 | 0.217594 | 0.239026 | 0.255472 | 0.222625 | 0.214820 | 0.221613 | 0.265672 | 0.443324 | 0.817599 | 1.407234 | 1.888609 | 2.946020 | 4.452240 | 6.613554 | 7.579005 | 5.162498 | 3.847790 | 4.056054 | 3.002024 | 2.043803 | 1.240550 | 0.797158 | 0.779727 | 0.725581 | 0.688304 | 0.711178 | 1.559855 | 5.002813 | 12.411258 | 20.030928 | 31.340807 | 32.878376 | 34.518186 | 16.524372 | 7.073238 | 2.944436 | 2.302802 | 2.346148 | 2.465113 | 1.932016 | 2.060617 | 1.805866 | 2.888792 | 4.252568 | 4.827203 | 3.700885 | 2.342272 | 1.580289 | 1.467917 | 1.060563 | 1.086647 | 0.810101 | 0.968281 | 1.074188 | 0.833663 | 0.547688 | 0.420214 | 0.313281 | 0.284532 | 0.224828 | 0.224768 | 0.304278 | 0.347802 | 0.271140 | 0.286280 | 0.449530 | 0.589032 | 0.374752 | 0.269430 | 0.237917 | 0.301591 | 0.308651 | 0.237171 | 0.216249 | 0.218600 | 0.226230 | 0.289054 | 0.335829 | 0.398732 | 0.464923 | 0.543564 | 0.903134 | 0.998423 | 0.741796 | 0.542896 | 0.498913 | 0.509068 | 0.560070 | 0.411253 | 0.327678 | 0.257922 | 0.251425 | 0.334025 | 0.592981 | 0.755491 | 0.561563 | 0.536441 | 0.554078 | 0.483244 | 0.294161 | 0.238888 | 0.218015 | 0.218806 | 0.218049 | 0.216997 | 0.240111 | 0.313253 | 0.307176 | 0.335667 | 0.355079 | 0.330169 | 0.365686 | 0.283142 | 0.232990 | 0.223750 | 0.216617 | 0.217825 | 0.225542 | 0.275504 | 0.336069 | 0.354527 | 0.331306 | 0.250677 | 0.214648 | 0.224343 | 0.252877 | 0.254953 | 0.219165 | 0.214923 | 0.215338 | 0.222085 | 0.221690 | 0.218008 | 0.218604 | 0.215354 | 0.214775 | 0.215938 | 0.232097 | 0.283270 | 0.274982 | 0.224399 | 0.214753 | 0.260289 | 0.248313 | 0.236094 | 0.221999 | 0.215884 | 0.222662 | 0.222768 | 0.236165 | 0.225584 | 0.224970 | 0.263579 | 0.291726 | 0.398084 | 0.478764 | 0.511940 | 0.585010 | 0.473498 | 0.289991 | 0.275514 | 0.256289 | 0.256593 | 0.294185 | 0.329769 | 0.287023 | 0.262024 | 0.258974 | 0.274147 | 0.290638 | 0.318378 | 0.328155 | 0.471446 | 0.703681 | 0.770496 | 0.662893 | 0.681418 | 0.635176 | 0.726592 | 0.585595 | 0.533022 | 0.546713 | 0.612302 | 0.641331 | 0.617638 | 0.664248 | 0.824771 | 1.049812 | 0.974574 | 0.774371 | 0.472243 | 0.351563 | 0.240067 | 0.215303 | 0.223694 | 0.257875 | 0.332866 | 0.406449 | 0.452067 | 0.454425 | 0.455399 | 0.651395 |
| left central | 0.371929 | 0.394624 | 0.333935 | 0.451249 | 1.002993 | 2.881199 | 1.905098 | 0.759139 | 0.582863 | 1.067806 | 0.664774 | 0.396092 | 0.264166 | 0.355483 | 0.257562 | 0.217048 | 0.294886 | 0.370973 | 0.340255 | 0.291910 | 0.487888 | 0.357514 | 0.219357 | 0.240803 | 0.468367 | 0.612213 | 0.683179 | 0.684645 | 0.527319 | 0.247668 | 0.246149 | 0.673151 | 1.008980 | 0.819337 | 0.391231 | 0.262116 | 0.215759 | 0.224103 | 0.217578 | 0.229963 | 0.257710 | 0.245933 | 0.253355 | 0.270369 | 0.352176 | 0.780050 | 0.808415 | 0.492419 | 0.346331 | 0.255038 | 0.215902 | 0.227614 | 0.297293 | 0.347497 | 0.301848 | 0.267472 | 0.259947 | 0.234377 | 0.224958 | 0.215602 | 0.217706 | 0.226920 | 0.233960 | 0.245183 | 0.252167 | 0.319038 | 0.427370 | 5.955589e-01 | 7.523932e-01 | 1.114103e+00 | 2.234759e+00 | 1.403889e+01 | 61.205186 | 235.971824 | 525.022166 | 943.381702 | 1004.398518 | 721.659029 | 133.545898 | 32.451696 | 8.891761 | 2.389249 | 0.797910 | 0.534304 | 0.541449 | 0.880099 | 0.675762 | 0.554850 | 0.768989 | 1.340701 | 2.286563 | 2.372070 | 1.459393 | 1.674451 | 3.381191 | 4.945405 | 6.555176 | 2.516299 | 1.129310 | 0.728855 | 0.557584 | 0.322390 | 0.227842 | 0.216592 | 0.214900 | 0.214809 | 0.216929 | 0.218576 | 0.215731 | 0.231176 | 0.238295 | 0.278420 | 0.311403 | 0.369075 | 0.397349 | 0.282656 | 0.236128 | 0.222989 | 0.218520 | 0.243401 | 0.287920 | 0.347646 | 0.480269 | 0.579302 | 0.562826 | 0.466085 | 0.355500 | 0.266012 | 0.283233 | 0.318046 | 0.379372 | 0.457669 | 0.506187 | 0.511067 | 0.393050 | 0.258937 | 0.218447 | 0.215462 | 0.218139 | 0.221307 | 0.215135 | 0.217376 | 0.233424 | 0.266340 | 0.360297 | 0.657616 | 1.227275 | 0.807734 | 0.564151 | 0.435576 | 0.307440 | 0.231861 | 0.222484 | 0.220260 | 0.231013 | 0.250670 | 0.286862 | 0.343040 | 0.499496 | 0.446604 | 0.394176 | 0.357234 | 0.354913 | 0.314589 | 0.308761 | 0.314703 | 0.326307 | 0.349352 | 0.368482 | 0.324632 | 0.336392 | 0.369599 | 0.315774 | 0.312723 | 0.344899 | 0.394963 | 0.464471 | 0.549916 | 0.469290 | 0.549676 | 0.465301 | 0.308401 | 0.253906 | 0.230411 | 0.219551 | 0.216108 | 0.215596 | 0.214690 | 0.216707 | 0.216390 | 0.214626 | 0.214980 | 0.215007 | 0.217137 | 0.222133 | 0.230775 | 0.236304 | 0.228087 | 0.236146 | 0.242997 | 0.242963 | 0.224334 | 0.215751 | 0.215888 | 0.214701 | 0.216918 | 0.231072 | 0.240266 | 0.320780 | 0.542222 | 1.339969 | 2.514219 | 3.570984 | 4.334440 | 7.917663 | 7.373450 | 3.632806 | 1.525968 | 0.623517 | 0.422251 | 0.337397 | 0.275434 | 0.226668 | 0.216110 | 0.215668 | 0.217449 | 0.215665 | 0.214652 | 0.214686 | 0.214685 | 0.215724 | 0.214862 | 0.215713 | 0.222021 | 0.218973 | 0.214626 | 0.214842 | 0.214819 | 0.245075 | 0.267804 | 0.298984 | 0.285143 | 0.250005 | 0.251431 | 0.251489 | 0.231718 | 0.219973 | 0.225593 | 0.242153 | 0.248799 | 0.258374 | 0.247749 | 0.245378 | 0.234922 | 0.216095 | 0.214642 | 0.214841 | 0.214675 | 0.234932 | 0.359958 | 0.692485 | 1.103172 | 1.354470 | 1.154702 | 1.434676 | 1.529693 | 1.097369 | 0.592829 | 0.535367 | 0.462527 | 0.540436 | 0.536167 | 0.468981 | 0.472926 | 0.648815 | 0.830739 | 0.808267 | 0.798470 | 0.727770 | 0.695827 |
| right central | 0.425515 | 0.604123 | 0.814318 | 1.554150 | 2.134826 | 4.951757 | 15.664431 | 6.844277 | 11.834960 | 27.147633 | 20.669452 | 18.799101 | 4.578182 | 2.913228 | 2.415824 | 1.560095 | 1.280302 | 0.400260 | 0.216107 | 0.215545 | 0.218397 | 0.234721 | 0.232120 | 0.331670 | 0.342629 | 0.252953 | 0.226083 | 0.228907 | 0.220748 | 0.220828 | 0.262636 | 0.255572 | 0.259732 | 0.350783 | 0.276583 | 0.361560 | 0.214928 | 0.277325 | 0.418834 | 0.460779 | 0.497811 | 0.346671 | 0.285636 | 0.220031 | 0.216044 | 0.262473 | 0.256665 | 0.271393 | 0.243968 | 0.248083 | 0.247533 | 0.225291 | 0.215469 | 0.214626 | 0.216243 | 0.217131 | 0.231998 | 0.240523 | 0.251042 | 0.285423 | 0.221362 | 0.233639 | 0.340896 | 0.502680 | 0.957803 | 2.726929 | 7.033552 | 1.393574e+01 | 3.390177e+01 | 7.625893e+01 | 2.343083e+02 | 1.897242e+02 | 150.465989 | 48.191092 | 11.494188 | 4.970823 | 2.871380 | 1.983673 | 2.118559 | 4.653294 | 9.057422 | 17.796809 | 29.444867 | 47.989222 | 83.136650 | 122.412559 | 79.775285 | 137.104350 | 523.658714 | 2735.693452 | 10120.958046 | 10077.058388 | 20136.209072 | 30277.926564 | 13400.659470 | 3880.194572 | 743.613623 | 194.812883 | 110.555460 | 52.079240 | 19.839585 | 38.602387 | 60.322288 | 168.745088 | 218.256990 | 56.902060 | 63.245677 | 86.582050 | 48.978568 | 123.828687 | 122.210873 | 189.105493 | 270.605003 | 338.807577 | 314.215671 | 152.996827 | 99.991229 | 294.800533 | 356.597002 | 118.877276 | 63.606594 | 24.547561 | 10.330325 | 7.324240 | 5.395684 | 3.548266 | 2.666676 | 2.527240 | 1.750894 | 1.821979 | 1.529907 | 1.695958 | 1.586366 | 1.763922 | 1.535481 | 1.248160 | 0.704071 | 0.517808 | 0.388845 | 0.373577 | 0.286892 | 0.260644 | 0.231572 | 0.241809 | 0.250216 | 0.234714 | 0.225865 | 0.227465 | 0.225422 | 0.258607 | 0.347546 | 0.424293 | 0.478435 | 0.511820 | 0.542322 | 0.923005 | 0.851985 | 0.474943 | 0.363747 | 0.336599 | 0.373370 | 0.389562 | 0.280316 | 0.247795 | 0.219423 | 0.218316 | 0.271762 | 0.322173 | 0.374085 | 0.378899 | 0.300633 | 0.367670 | 0.343018 | 0.266893 | 0.235571 | 0.222417 | 0.233215 | 0.283954 | 0.356649 | 0.554484 | 0.998669 | 2.056490 | 2.341333 | 1.984107 | 1.248673 | 0.789312 | 0.645030 | 0.478306 | 0.348935 | 0.395756 | 0.450910 | 0.554363 | 0.879843 | 1.205078 | 1.381685 | 1.342705 | 0.857880 | 0.786625 | 0.859092 | 0.709794 | 0.782956 | 1.067817 | 1.382180 | 1.232013 | 0.783868 | 0.674774 | 0.440596 | 0.291445 | 0.240547 | 0.227843 | 0.238696 | 0.247366 | 0.247570 | 0.278553 | 0.305579 | 0.306979 | 0.295269 | 0.261704 | 0.227273 | 0.214626 | 0.227051 | 0.238557 | 0.228043 | 0.234348 | 0.227847 | 0.219004 | 0.217782 | 0.218207 | 0.222082 | 0.231239 | 0.222559 | 0.222528 | 0.214929 | 0.215084 | 0.219936 | 0.217528 | 0.214959 | 0.244809 | 0.274898 | 0.299384 | 0.268981 | 0.267565 | 0.227812 | 0.216666 | 0.215049 | 0.219747 | 0.232329 | 0.224754 | 0.217852 | 0.218121 | 0.229836 | 0.229256 | 0.223352 | 0.229619 | 0.219487 | 0.216733 | 0.225618 | 0.231983 | 0.238820 | 0.240636 | 0.247477 | 0.234730 | 0.214626 | 0.227306 | 0.256105 | 0.276260 | 0.281246 | 0.302591 | 0.313523 | 0.270220 | 0.249096 | 0.237092 | 0.254987 | 0.334557 | 0.376637 | 0.400341 | 0.419697 | 0.410376 | 0.411040 | 0.369947 |
| left posterior | 1.384911 | 0.922621 | 0.924255 | 1.844579 | 1.974061 | 1.529694 | 1.334873 | 0.594115 | 0.309858 | 0.226161 | 0.233367 | 0.308103 | 0.432762 | 0.428223 | 0.277338 | 0.214829 | 0.267722 | 1.260332 | 8.161993 | 53.091118 | 29.930599 | 4.620251 | 0.764114 | 0.287821 | 0.218808 | 0.317958 | 0.420879 | 0.307881 | 0.216333 | 0.216566 | 0.236758 | 0.306057 | 0.357911 | 0.277352 | 0.218711 | 0.250456 | 0.290700 | 0.318845 | 0.432458 | 0.579485 | 0.544543 | 0.704866 | 0.527381 | 0.301369 | 0.286463 | 0.268273 | 0.241409 | 0.237157 | 0.217608 | 0.214978 | 0.221387 | 0.266792 | 0.365116 | 0.683567 | 1.004305 | 0.981672 | 0.908114 | 0.442461 | 0.252322 | 0.215472 | 0.279757 | 0.463194 | 1.435817 | 8.833935 | 113.363800 | 5698.826677 | 165955.893156 | 1.955619e+06 | 1.047231e+07 | 9.525794e+06 | 6.649319e+06 | 3.872849e+06 | 920200.933408 | 238146.262074 | 28517.393736 | 7246.220709 | 3042.648273 | 965.407169 | 736.293823 | 2517.629443 | 12915.082553 | 7148.242370 | 577.214596 | 201.110120 | 101.272959 | 49.902761 | 38.830884 | 43.472609 | 54.944086 | 124.443979 | 158.679104 | 165.638772 | 228.440807 | 255.314388 | 182.999386 | 197.506448 | 181.915496 | 145.316886 | 113.540840 | 85.926616 | 50.754093 | 35.097679 | 30.944342 | 43.704746 | 52.977250 | 46.880771 | 34.381053 | 28.885656 | 28.923645 | 20.847433 | 12.661766 | 7.877352 | 6.563098 | 7.815452 | 8.746557 | 13.605124 | 20.091684 | 29.436170 | 58.598487 | 144.755804 | 245.865323 | 309.519695 | 234.649253 | 162.780699 | 98.683851 | 46.262279 | 29.402346 | 14.924412 | 8.065050 | 3.958396 | 3.017552 | 2.037227 | 1.743476 | 1.434508 | 1.719338 | 2.726572 | 3.594660 | 4.124740 | 5.448331 | 5.830003 | 4.954490 | 4.903706 | 3.025797 | 2.555493 | 2.285248 | 1.742586 | 1.555584 | 1.650893 | 1.297683 | 1.105106 | 1.087573 | 0.728796 | 0.578027 | 0.544621 | 0.515562 | 0.531808 | 0.575823 | 0.686012 | 1.121489 | 1.773044 | 2.079856 | 3.267687 | 4.183154 | 4.993367 | 3.528052 | 2.716883 | 1.561154 | 1.209753 | 0.952103 | 0.764684 | 0.576128 | 0.591700 | 0.561096 | 0.766149 | 0.929866 | 0.844135 | 0.941422 | 1.006744 | 1.006106 | 1.000428 | 0.849383 | 0.745612 | 0.743161 | 0.688796 | 0.674258 | 0.571497 | 0.574676 | 0.624390 | 0.607420 | 0.502523 | 0.441171 | 0.417418 | 0.527661 | 0.546061 | 0.522634 | 0.560384 | 0.661624 | 0.744675 | 0.922920 | 0.855474 | 0.941309 | 1.094620 | 1.108977 | 1.113138 | 1.027568 | 0.938202 | 0.952252 | 1.075580 | 1.341801 | 1.832168 | 2.227811 | 2.756394 | 3.150642 | 3.284502 | 3.615064 | 4.315107 | 4.626224 | 5.293329 | 6.907455 | 7.786472 | 6.847693 | 6.037876 | 5.380137 | 4.906621 | 5.166195 | 5.221986 | 5.274390 | 5.885793 | 5.860121 | 5.778020 | 5.245631 | 4.544862 | 3.819935 | 3.525328 | 3.025846 | 2.884951 | 2.343610 | 2.111941 | 1.901587 | 2.109215 | 2.645092 | 3.351914 | 3.128185 | 3.080919 | 2.013248 | 1.512539 | 1.240711 | 1.128336 | 1.337944 | 1.846590 | 1.845425 | 2.100107 | 2.666673 | 2.226065 | 1.853732 | 1.305329 | 0.817647 | 0.732458 | 0.806355 | 0.683568 | 0.674766 | 0.581033 | 0.466349 | 0.391814 | 0.347043 | 0.297593 | 0.261194 | 0.233196 | 0.222602 | 0.216886 | 0.214769 | 0.221108 | 0.239613 | 0.255825 | 0.259058 | 0.264977 | 0.268217 | 0.251523 | 0.250015 | 0.250987 |
| right posterior | 0.240519 | 0.274228 | 0.315589 | 0.420704 | 0.994840 | 1.695303 | 0.898482 | 0.423518 | 0.334316 | 0.302072 | 0.269272 | 0.222744 | 0.214986 | 0.216970 | 0.217636 | 0.233311 | 0.293007 | 0.369818 | 0.553285 | 0.517364 | 0.589684 | 0.495245 | 0.373225 | 0.230579 | 0.214723 | 0.214674 | 0.219283 | 0.233552 | 0.340542 | 0.269110 | 0.279701 | 0.363563 | 0.324147 | 0.279285 | 0.226280 | 0.265301 | 0.531495 | 1.296673 | 2.367576 | 2.936524 | 3.182697 | 2.689136 | 1.184911 | 1.153226 | 0.896528 | 0.985998 | 0.663041 | 0.285117 | 0.215042 | 0.218943 | 0.273681 | 0.287838 | 0.233103 | 0.214720 | 0.236022 | 0.363386 | 0.467612 | 0.894990 | 2.680527 | 2.536713 | 2.029634 | 2.723920 | 4.575246 | 12.106084 | 18.112414 | 32.965939 | 129.733083 | 4.001794e+02 | 1.500469e+03 | 2.956489e+03 | 5.496674e+03 | 1.604936e+04 | 19286.972786 | 9074.750033 | 1508.486018 | 164.352735 | 37.741692 | 9.255252 | 3.250725 | 1.696207 | 1.678835 | 3.608543 | 3.919113 | 5.215536 | 5.504903 | 5.707446 | 8.098835 | 13.071858 | 15.407688 | 41.538168 | 130.498883 | 348.422240 | 545.420908 | 652.207395 | 1077.366139 | 1060.675278 | 427.400064 | 163.137133 | 129.877092 | 97.428297 | 67.912750 | 31.481984 | 26.891273 | 31.738211 | 39.316061 | 56.249507 | 127.339788 | 393.140166 | 1023.211894 | 2745.618144 | 3694.150487 | 1429.111644 | 538.459732 | 284.984902 | 174.081842 | 140.415996 | 148.785998 | 233.608927 | 388.895747 | 931.589548 | 1194.179686 | 1461.000372 | 1432.586259 | 1038.420250 | 405.723330 | 307.845117 | 204.699606 | 123.809994 | 58.865271 | 27.432189 | 16.673966 | 16.808091 | 26.157964 | 49.852273 | 57.705917 | 40.798449 | 35.256654 | 30.104702 | 39.465402 | 14.914070 | 8.924480 | 8.723689 | 12.188767 | 13.604379 | 20.301463 | 15.391190 | 19.954928 | 12.861719 | 7.149079 | 4.262714 | 3.195554 | 2.645896 | 2.926156 | 2.394802 | 2.402054 | 2.966287 | 3.947394 | 5.623439 | 5.862750 | 4.710857 | 4.625004 | 5.008048 | 5.601725 | 6.059921 | 6.872051 | 7.675269 | 10.025091 | 12.128899 | 14.766359 | 11.684892 | 9.089440 | 6.440514 | 3.604821 | 2.095600 | 1.137194 | 0.991825 | 1.271842 | 2.413763 | 4.877992 | 11.062116 | 13.975334 | 14.418005 | 10.749560 | 6.781522 | 4.378483 | 3.424996 | 2.828972 | 3.276624 | 4.134726 | 4.581180 | 4.276686 | 5.344876 | 5.952293 | 6.202742 | 4.682361 | 4.950195 | 7.276649 | 9.002267 | 7.447678 | 6.557819 | 6.319025 | 8.741390 | 6.430122 | 5.042730 | 5.765495 | 4.313036 | 3.173010 | 2.598918 | 2.734313 | 3.024958 | 3.447457 | 3.639639 | 6.714832 | 11.099201 | 21.551692 | 28.807285 | 62.437849 | 109.064397 | 154.815620 | 65.637738 | 33.123931 | 23.109199 | 18.340213 | 15.443190 | 12.699795 | 9.347189 | 10.837092 | 12.855164 | 22.288032 | 24.814689 | 21.347313 | 19.965769 | 19.658249 | 16.788098 | 16.178863 | 14.538981 | 15.379023 | 19.844450 | 21.820706 | 21.884462 | 20.839197 | 20.572502 | 17.791765 | 20.265182 | 22.793328 | 24.271260 | 27.358859 | 22.384978 | 20.058187 | 15.023599 | 10.533641 | 4.946076 | 4.430022 | 3.401382 | 2.722958 | 2.473748 | 2.669310 | 2.988857 | 3.313850 | 2.701031 | 2.317786 | 2.681504 | 3.133600 | 2.986697 | 2.135783 | 1.715812 | 1.298120 | 1.101086 | 1.006291 | 0.742985 | 0.691008 | 0.812686 | 0.951546 | 1.418561 | 1.266388 | 1.049838 | 1.053439 | 1.055731 | 1.188042 | 1.244164 |
| all electrodes | 0.224083 | 0.219415 | 0.227812 | 0.273053 | 0.424627 | 0.409044 | 1.415435 | 6.882095 | 20.791558 | 2.868254 | 1.037271 | 0.504660 | 0.422723 | 0.308092 | 0.317195 | 0.351711 | 0.700328 | 1.009908 | 1.503226 | 1.397299 | 2.163940 | 1.412931 | 0.395147 | 0.214689 | 0.301631 | 0.496121 | 0.622626 | 2.198921 | 3.091124 | 0.935078 | 0.459694 | 0.263942 | 0.215002 | 0.216939 | 0.219020 | 0.221706 | 0.220885 | 0.217603 | 0.221872 | 0.237452 | 0.306977 | 0.538571 | 1.664523 | 6.367723 | 9.407828 | 10.797795 | 7.943380 | 3.006071 | 1.518219 | 0.645330 | 0.477899 | 0.385654 | 0.308486 | 0.285709 | 0.266802 | 0.243720 | 0.246743 | 0.222328 | 0.214889 | 0.223735 | 0.247990 | 0.314794 | 0.460421 | 0.750852 | 1.438890 | 3.983504 | 5.950701 | 1.048037e+01 | 3.871028e+01 | 8.568445e+01 | 2.342408e+02 | 6.167901e+02 | 2623.790953 | 2137.718770 | 3758.122138 | 5913.650459 | 9422.084915 | 15870.485217 | 7965.854998 | 3163.575674 | 1628.822934 | 109.014514 | 11.448689 | 4.005475 | 2.344078 | 1.466748 | 0.861516 | 0.543660 | 0.877050 | 2.413084 | 8.079460 | 22.302101 | 25.681140 | 41.085076 | 100.402962 | 108.576561 | 172.764646 | 171.495652 | 104.101404 | 67.474657 | 24.544441 | 13.704349 | 10.320072 | 3.447165 | 2.769553 | 1.751836 | 1.694054 | 1.441429 | 1.188344 | 2.169775 | 4.516261 | 4.007573 | 5.973729 | 5.823326 | 8.788075 | 7.765432 | 10.793130 | 13.764636 | 23.465892 | 34.436992 | 37.306613 | 34.282219 | 38.333360 | 23.295189 | 8.572661 | 3.008157 | 1.376146 | 0.902114 | 1.019172 | 1.145635 | 1.407840 | 1.831131 | 1.716040 | 1.687593 | 1.944640 | 1.025195 | 0.804712 | 0.847824 | 1.312930 | 2.695408 | 4.173129 | 5.912487 | 12.232143 | 17.283965 | 8.944024 | 2.970086 | 1.360922 | 0.711158 | 0.520550 | 0.404283 | 0.340615 | 0.279771 | 0.282359 | 0.323729 | 0.380838 | 0.405290 | 0.376087 | 0.374997 | 0.546162 | 0.732144 | 0.650519 | 0.547782 | 0.410360 | 0.422151 | 0.401673 | 0.356442 | 0.352496 | 0.356696 | 0.377473 | 0.392695 | 0.306811 | 0.233718 | 0.216852 | 0.216208 | 0.221863 | 0.217273 | 0.218877 | 0.238899 | 0.324834 | 0.381777 | 0.457483 | 0.434619 | 0.358630 | 0.298034 | 0.307748 | 0.312037 | 0.327531 | 0.306074 | 0.274041 | 0.249880 | 0.243931 | 0.250748 | 0.244079 | 0.252151 | 0.268269 | 0.367611 | 0.524912 | 0.901759 | 0.928651 | 0.969245 | 1.213769 | 1.025884 | 0.876456 | 0.934532 | 0.777119 | 0.987741 | 0.807548 | 0.534791 | 0.752252 | 0.949595 | 1.096378 | 1.094728 | 0.970470 | 1.474051 | 2.763541 | 1.900977 | 1.486676 | 0.863531 | 0.758266 | 0.771265 | 0.549284 | 0.386301 | 0.401661 | 0.475843 | 0.632173 | 0.666968 | 0.514652 | 0.449635 | 0.555898 | 0.689151 | 0.854185 | 0.932013 | 0.992046 | 1.314225 | 2.037652 | 2.349501 | 1.966698 | 1.664500 | 1.460163 | 1.558417 | 1.821431 | 1.502495 | 1.522527 | 1.477318 | 1.350717 | 1.060878 | 1.072899 | 0.843140 | 0.582839 | 0.438403 | 0.420974 | 0.435778 | 0.517730 | 0.493344 | 0.540013 | 0.786729 | 0.892343 | 0.671697 | 0.673581 | 0.805628 | 1.184652 | 1.186833 | 0.903584 | 0.718458 | 0.780748 | 0.584485 | 0.379490 | 0.249757 | 0.219185 | 0.214661 | 0.215084 | 0.220741 | 0.236159 | 0.246498 | 0.277727 | 0.360012 | 0.481868 | 0.717706 | 0.817040 | 0.796023 |

Searchlight, spatiotemporal cluster permutation test

|  | start time | stop time | peak time | peak channel | cluster p | peak Cohen's d | direction |
| --- | --- | --- | --- | --- | --- | --- | --- |
| #1 | 80 | 1145 | 150 | PO7 | 0.0001 | 1.64574 | positive |

I) real faces (LOSO) - happy vs angry

  
|  | time window | peak latency | cluster *p* | peak Cohen's *d* |  | | | |
| **all electrodes** | 145 - 215 ms | 175 ms | 0.0267 | 1.2307 |  | | | |
|  | | | | | | | | |

Time-resolved classification, cluster permutation tests

|  | **left hemisphere** | | | | **right hemisphere** | | | |
|  | time window | peak latency | cluster *p* | peak Cohen's *d* | time window | peak latency | cluster *p* | peak Cohen's *d* |
| **anterior** |  | | | |  | | | |
| **central** |  | | | |  | | | |
| **posterior** | 125 - 270 ms | 170 ms | 0.0106 | 1.0986 | 145 - 215 ms | 170 ms | 0.0475 | 1.1431 |
 320 - 500 ms | 405 ms | 0.013 | 0.7323 | 360 - 525 ms | 415 ms | 0.0284 | 0.8843 |  | | | | 650 - 1125 ms | 1005 ms | 0.0047 | 0.7603 |

  

Time-resolved classification, Bayesian statistics

|  | -200 | -195 | -190 | -185 | -180 | -175 | -170 | -165 | -160 | -155 | -150 | -145 | -140 | -135 | -130 | -125 | -120 | -115 | -110 | -105 | -100 | -95 | -90 | -85 | -80 | -75 | -70 | -65 | -60 | -55 | -50 | -45 | -40 | -35 | -30 | -25 | -20 | -15 | -10 | -5 | 0 | 5 | 10 | 15 | 20 | 25 | 30 | 35 | 40 | 45 | 50 | 55 | 60 | 65 | 70 | 75 | 80 | 85 | 90 | 95 | 100 | 105 | 110 | 115 | 120 | 125 | 130 | 135 | 140 | 145 | 150 | 155 | 160 | 165 | 170 | 175 | 180 | 185 | 190 | 195 | 200 | 205 | 210 | 215 | 220 | 225 | 230 | 235 | 240 | 245 | 250 | 255 | 260 | 265 | 270 | 275 | 280 | 285 | 290 | 295 | 300 | 305 | 310 | 315 | 320 | 325 | 330 | 335 | 340 | 345 | 350 | 355 | 360 | 365 | 370 | 375 | 380 | 385 | 390 | 395 | 400 | 405 | 410 | 415 | 420 | 425 | 430 | 435 | 440 | 445 | 450 | 455 | 460 | 465 | 470 | 475 | 480 | 485 | 490 | 495 | 500 | 505 | 510 | 515 | 520 | 525 | 530 | 535 | 540 | 545 | 550 | 555 | 560 | 565 | 570 | 575 | 580 | 585 | 590 | 595 | 600 | 605 | 610 | 615 | 620 | 625 | 630 | 635 | 640 | 645 | 650 | 655 | 660 | 665 | 670 | 675 | 680 | 685 | 690 | 695 | 700 | 705 | 710 | 715 | 720 | 725 | 730 | 735 | 740 | 745 | 750 | 755 | 760 | 765 | 770 | 775 | 780 | 785 | 790 | 795 | 800 | 805 | 810 | 815 | 820 | 825 | 830 | 835 | 840 | 845 | 850 | 855 | 860 | 865 | 870 | 875 | 880 | 885 | 890 | 895 | 900 | 905 | 910 | 915 | 920 | 925 | 930 | 935 | 940 | 945 | 950 | 955 | 960 | 965 | 970 | 975 | 980 | 985 | 990 | 995 | 1000 | 1005 | 1010 | 1015 | 1020 | 1025 | 1030 | 1035 | 1040 | 1045 | 1050 | 1055 | 1060 | 1065 | 1070 | 1075 | 1080 | 1085 | 1090 | 1095 | 1100 | 1105 | 1110 | 1115 | 1120 | 1125 | 1130 | 1135 | 1140 | 1145 | 1150 | 1155 | 1160 | 1165 | 1170 | 1175 | 1180 | 1185 | 1190 | 1195 |
| --- | --- | --- | --- | --- | --- | --- | --- | --- | --- | --- | --- | --- | --- | --- | --- | --- | --- | --- | --- | --- | --- | --- | --- | --- | --- | --- | --- | --- | --- | --- | --- | --- | --- | --- | --- | --- | --- | --- | --- | --- | --- | --- | --- | --- | --- | --- | --- | --- | --- | --- | --- | --- | --- | --- | --- | --- | --- | --- | --- | --- | --- | --- | --- | --- | --- | --- | --- | --- | --- | --- | --- | --- | --- | --- | --- | --- | --- | --- | --- | --- | --- | --- | --- | --- | --- | --- | --- | --- | --- | --- | --- | --- | --- | --- | --- | --- | --- | --- | --- | --- | --- | --- | --- | --- | --- | --- | --- | --- | --- | --- | --- | --- | --- | --- | --- | --- | --- | --- | --- | --- | --- | --- | --- | --- | --- | --- | --- | --- | --- | --- | --- | --- | --- | --- | --- | --- | --- | --- | --- | --- | --- | --- | --- | --- | --- | --- | --- | --- | --- | --- | --- | --- | --- | --- | --- | --- | --- | --- | --- | --- | --- | --- | --- | --- | --- | --- | --- | --- | --- | --- | --- | --- | --- | --- | --- | --- | --- | --- | --- | --- | --- | --- | --- | --- | --- | --- | --- | --- | --- | --- | --- | --- | --- | --- | --- | --- | --- | --- | --- | --- | --- | --- | --- | --- | --- | --- | --- | --- | --- | --- | --- | --- | --- | --- | --- | --- | --- | --- | --- | --- | --- | --- | --- | --- | --- | --- | --- | --- | --- | --- | --- | --- | --- | --- | --- | --- | --- | --- | --- | --- | --- | --- | --- | --- | --- | --- | --- | --- | --- | --- | --- | --- | --- | --- | --- | --- | --- | --- | --- | --- | --- | --- | --- | --- | --- | --- | --- | --- | --- | --- | --- | --- | --- | --- | --- | --- | --- | --- | --- | --- |
| left anterior | 0.216576 | 0.256940 | 0.528384 | 1.520630 | 2.025647 | 1.324653 | 0.482304 | 0.339386 | 0.235851 | 0.227339 | 0.270564 | 0.304879 | 0.320703 | 0.244329 | 0.216492 | 0.224831 | 0.224958 | 0.216176 | 0.229564 | 0.218382 | 0.231437 | 0.217977 | 0.216287 | 0.233798 | 0.257935 | 0.245846 | 0.511968 | 1.080099 | 1.155465 | 1.271121 | 1.381020 | 0.944026 | 0.432719 | 0.236826 | 0.217125 | 0.227111 | 0.333068 | 1.162770 | 1.907566 | 0.695936 | 0.289190 | 0.312455 | 0.396228 | 0.376873 | 0.338598 | 0.439433 | 0.430817 | 0.380169 | 0.252717 | 0.310105 | 0.352311 | 0.310642 | 0.294153 | 0.266081 | 0.353953 | 0.288949 | 0.214639 | 0.218125 | 0.222746 | 0.262107 | 0.237877 | 0.232276 | 0.214935 | 0.231064 | 0.246743 | 0.234997 | 0.251625 | 0.253767 | 0.239680 | 0.216378 | 0.215549 | 0.223403 | 0.230372 | 0.242654 | 0.286945 | 0.318156 | 0.302300 | 0.331231 | 0.324301 | 0.225918 | 0.233555 | 0.393186 | 0.675338 | 1.395096 | 4.691520 | 10.355621 | 9.531017 | 6.353362 | 5.138613 | 3.854222 | 1.369431 | 0.703304 | 0.903061 | 1.300979 | 1.812108 | 1.405626 | 1.056135 | 1.289786 | 1.767468 | 0.854584 | 0.554402 | 0.584445 | 0.470944 | 0.431574 | 0.452843 | 0.359400 | 0.401953 | 0.419176 | 0.344653 | 0.282417 | 0.224377 | 0.216377 | 0.225389 | 0.235993 | 0.269241 | 0.338155 | 0.507027 | 0.635796 | 0.643559 | 0.778257 | 0.918489 | 0.685492 | 0.452074 | 0.277014 | 0.221746 | 0.214980 | 0.214640 | 0.216344 | 0.236424 | 0.296286 | 0.294620 | 0.258574 | 0.281510 | 0.303676 | 0.324756 | 0.380270 | 0.495997 | 0.616907 | 0.788300 | 0.748468 | 0.841326 | 0.829566 | 0.492663 | 0.284283 | 0.233252 | 0.220047 | 0.215441 | 0.224769 | 0.260815 | 0.328318 | 0.343882 | 0.319702 | 0.254035 | 0.214987 | 0.279827 | 0.457766 | 0.940049 | 1.019436 | 1.131949 | 1.577889 | 2.376254 | 3.528365 | 4.505505 | 3.501552 | 4.876668 | 4.353357 | 1.876409 | 0.744404 | 0.501450 | 0.359006 | 0.287200 | 0.240902 | 0.243687 | 0.255002 | 0.275968 | 0.279010 | 0.297729 | 0.316417 | 0.319779 | 0.289382 | 0.255497 | 0.226775 | 0.217619 | 0.218642 | 0.230707 | 0.248261 | 0.262338 | 0.308827 | 0.515312 | 0.941624 | 1.237245 | 1.341827 | 1.112920 | 1.412414 | 0.969920 | 0.504704 | 0.314332 | 0.257725 | 0.254701 | 0.274228 | 0.271694 | 0.347538 | 0.351428 | 0.359014 | 0.358735 | 0.265560 | 0.240003 | 0.219745 | 0.214636 | 0.216223 | 0.223087 | 0.225336 | 0.222581 | 0.224964 | 0.241637 | 0.245512 | 0.215725 | 0.229264 | 0.262733 | 0.252400 | 0.240901 | 0.302267 | 0.303170 | 0.257059 | 0.277525 | 0.303540 | 0.379376 | 0.511634 | 0.485139 | 0.579154 | 0.627368 | 0.508522 | 0.422355 | 0.318323 | 0.268091 | 0.248156 | 0.248080 | 0.227893 | 0.218620 | 0.214716 | 0.215091 | 0.214771 | 0.215530 | 0.216431 | 0.216941 | 0.220855 | 0.240835 | 0.259064 | 0.276397 | 0.235706 | 0.214793 | 0.216906 | 0.217288 | 0.214744 | 0.214689 | 0.215456 | 0.215630 | 0.224280 | 0.232770 | 0.235755 | 0.222243 | 0.235896 | 0.250534 | 0.255029 | 0.271188 | 0.273294 | 0.247910 | 0.236013 | 0.221123 | 0.216618 | 0.215013 | 0.219396 | 0.226410 | 0.237312 | 0.233820 | 0.253278 | 0.258856 | 0.261077 | 0.253213 | 0.229055 |
| right anterior | 1.801621 | 2.205024 | 1.111110 | 0.412696 | 0.245552 | 0.218566 | 0.230847 | 0.378796 | 0.570143 | 1.169602 | 0.546999 | 0.353769 | 0.374733 | 0.462600 | 0.714085 | 0.618934 | 0.285112 | 0.347772 | 0.581122 | 0.393684 | 0.236209 | 0.219778 | 0.218179 | 0.228892 | 0.285382 | 0.380027 | 0.518154 | 0.724174 | 0.563217 | 0.340441 | 0.218771 | 0.235120 | 0.341919 | 0.645580 | 2.103285 | 4.196599 | 2.540627 | 1.605433 | 0.946930 | 0.669748 | 0.642858 | 0.595593 | 0.298768 | 0.308104 | 0.356124 | 0.560218 | 0.526683 | 0.528812 | 0.396721 | 0.559521 | 0.735354 | 0.787807 | 0.681708 | 1.167348 | 0.834429 | 0.562835 | 0.350575 | 0.337918 | 0.352464 | 0.275988 | 0.220032 | 0.215120 | 0.217487 | 0.229523 | 0.244106 | 0.246727 | 0.230347 | 0.228606 | 0.251664 | 0.260037 | 0.330175 | 0.535569 | 1.398772 | 3.209684 | 4.317153 | 1.237646 | 1.114794 | 0.453198 | 0.354873 | 0.310450 | 0.268232 | 0.230210 | 0.225852 | 0.230953 | 0.277656 | 0.450706 | 0.414588 | 0.419062 | 0.396216 | 0.486481 | 0.546018 | 0.725621 | 0.556592 | 0.741798 | 0.756323 | 0.852967 | 0.608818 | 0.470968 | 0.402867 | 0.490990 | 0.573782 | 0.672358 | 0.535507 | 0.491733 | 0.319882 | 0.256816 | 0.225160 | 0.214759 | 0.223080 | 0.256328 | 0.341491 | 0.349188 | 0.418600 | 0.479804 | 0.548458 | 0.851343 | 1.658888 | 2.253805 | 3.015780 | 1.945877 | 1.403150 | 0.947439 | 0.606038 | 0.431626 | 0.422407 | 0.461922 | 0.673509 | 0.653113 | 0.533649 | 0.340517 | 0.249829 | 0.214946 | 0.225802 | 0.250929 | 0.238118 | 0.228707 | 0.221476 | 0.226564 | 0.248478 | 0.274893 | 0.281334 | 0.264317 | 0.236441 | 0.219218 | 0.216214 | 0.218144 | 0.239482 | 0.246426 | 0.237636 | 0.243115 | 0.249215 | 0.291248 | 0.323708 | 0.263968 | 0.216356 | 0.238617 | 0.300614 | 0.365046 | 0.428962 | 0.648257 | 0.684591 | 0.478295 | 0.434879 | 0.455969 | 0.426667 | 0.587433 | 0.573311 | 0.711928 | 0.745951 | 0.464952 | 0.451135 | 0.616963 | 0.497977 | 0.446133 | 0.373918 | 0.417912 | 0.442971 | 0.386630 | 0.317645 | 0.286684 | 0.258223 | 0.221296 | 0.215235 | 0.217130 | 0.214890 | 0.214626 | 0.215089 | 0.215865 | 0.229567 | 0.246889 | 0.279768 | 0.283466 | 0.291908 | 0.279963 | 0.272169 | 0.257917 | 0.247238 | 0.217210 | 0.216809 | 0.222975 | 0.219544 | 0.214679 | 0.221156 | 0.225743 | 0.215575 | 0.225515 | 0.248030 | 0.277591 | 0.320951 | 0.352318 | 0.349386 | 0.318433 | 0.271365 | 0.283169 | 0.282705 | 0.284721 | 0.277894 | 0.289975 | 0.289111 | 0.291881 | 0.267795 | 0.269226 | 0.328874 | 0.475252 | 0.578441 | 0.712195 | 0.611867 | 0.516553 | 0.434746 | 0.336029 | 0.306855 | 0.311188 | 0.294583 | 0.321401 | 0.340636 | 0.345875 | 0.373873 | 0.314589 | 0.283695 | 0.269437 | 0.254116 | 0.246024 | 0.239985 | 0.231738 | 0.237406 | 0.231377 | 0.234765 | 0.236451 | 0.250096 | 0.286071 | 0.300046 | 0.291213 | 0.286999 | 0.274847 | 0.284148 | 0.308376 | 0.299426 | 0.271879 | 0.245879 | 0.251810 | 0.256746 | 0.257103 | 0.239327 | 0.230487 | 0.242821 | 0.277457 | 0.257971 | 0.222678 | 0.216775 | 0.259602 | 0.349225 | 0.478764 | 0.934575 | 1.075432 | 1.147414 | 0.912944 | 0.536043 | 0.448313 | 0.428460 | 0.348205 |
| left central | 0.319507 | 0.335482 | 0.600173 | 0.592295 | 0.594178 | 0.792158 | 0.432472 | 0.301600 | 0.273370 | 0.224988 | 0.222751 | 0.351656 | 1.131192 | 3.876953 | 24.194188 | 41.289116 | 77.275148 | 8.885235 | 1.258130 | 0.627702 | 0.770891 | 0.404473 | 0.291282 | 0.238915 | 0.278456 | 0.284861 | 0.256750 | 0.234180 | 0.303764 | 0.636779 | 3.505679 | 8.699433 | 5.003015 | 1.481153 | 0.580015 | 0.275373 | 0.214668 | 0.233704 | 0.288728 | 0.268587 | 0.235326 | 0.219959 | 0.225874 | 0.269224 | 0.250262 | 0.238893 | 0.228340 | 0.253345 | 0.270404 | 0.243795 | 0.239523 | 0.265534 | 0.286401 | 0.270938 | 0.293233 | 0.387308 | 0.955415 | 1.169655 | 1.227239 | 1.792432 | 4.184264 | 2.280682 | 1.107351 | 0.596817 | 0.684113 | 0.729341 | 0.726419 | 0.431054 | 0.263322 | 0.219337 | 0.218303 | 0.367909 | 0.801928 | 1.698649 | 3.282604 | 3.400126 | 1.715000 | 0.756491 | 0.330312 | 0.214634 | 0.291258 | 0.650042 | 1.299905 | 1.345208 | 0.814207 | 0.558878 | 0.465208 | 0.483416 | 0.349433 | 0.233469 | 0.215962 | 0.220048 | 0.262861 | 0.310659 | 0.293312 | 0.301388 | 0.325071 | 0.348184 | 0.351072 | 0.343211 | 0.378823 | 0.559066 | 0.953792 | 1.231703 | 1.641900 | 1.664125 | 0.806024 | 0.316421 | 0.215117 | 0.261689 | 0.325249 | 0.392038 | 0.484265 | 0.485559 | 0.443848 | 0.359936 | 0.335824 | 0.359703 | 0.438624 | 0.502773 | 0.576905 | 0.520497 | 0.464002 | 0.416750 | 0.336016 | 0.256780 | 0.220251 | 0.214775 | 0.215517 | 0.216358 | 0.218816 | 0.214659 | 0.223680 | 0.242880 | 0.226257 | 0.215090 | 0.236480 | 0.280197 | 0.332582 | 0.457797 | 0.622695 | 0.779627 | 1.107761 | 1.738142 | 3.383566 | 6.361668 | 6.154765 | 5.422302 | 2.415447 | 0.830179 | 0.381957 | 0.258918 | 0.243453 | 0.262412 | 0.247191 | 0.269242 | 0.359414 | 0.486794 | 0.475450 | 0.329807 | 0.290298 | 0.341022 | 0.426768 | 0.438566 | 0.484536 | 0.611400 | 0.859367 | 0.932723 | 0.791477 | 0.603430 | 0.496527 | 0.418665 | 0.366155 | 0.343104 | 0.325291 | 0.285987 | 0.252916 | 0.234984 | 0.220345 | 0.214893 | 0.221344 | 0.244790 | 0.276384 | 0.288373 | 0.303605 | 0.291170 | 0.288330 | 0.275336 | 0.247106 | 0.226379 | 0.225376 | 0.217574 | 0.215316 | 0.214848 | 0.225461 | 0.276059 | 0.383435 | 0.486155 | 0.659129 | 0.828409 | 1.027755 | 0.833166 | 0.651512 | 0.543819 | 0.462338 | 0.480405 | 0.424426 | 0.418422 | 0.369643 | 0.335191 | 0.354290 | 0.393833 | 0.404024 | 0.595448 | 0.859194 | 1.703205 | 4.014438 | 6.833346 | 7.167208 | 5.286507 | 3.649688 | 2.687106 | 1.493646 | 0.808651 | 0.591831 | 0.604884 | 0.680571 | 0.782155 | 0.805426 | 0.895902 | 1.111876 | 1.359709 | 1.447766 | 1.479741 | 1.253083 | 0.903386 | 0.710664 | 0.567721 | 0.458469 | 0.389731 | 0.328882 | 0.277848 | 0.264117 | 0.265030 | 0.263565 | 0.241537 | 0.220983 | 0.214657 | 0.215184 | 0.215508 | 0.216247 | 0.216778 | 0.214675 | 0.225473 | 0.274594 | 0.312744 | 0.305739 | 0.309978 | 0.312501 | 0.290953 | 0.257157 | 0.224083 | 0.215885 | 0.214634 | 0.215103 | 0.224645 | 0.235186 | 0.248877 | 0.232399 | 0.222432 | 0.219512 | 0.227158 | 0.239956 | 0.284283 | 0.334243 | 0.467534 | 0.515069 | 0.562602 | 0.619385 | 0.585883 |
| right central | 0.215889 | 0.217919 | 0.237183 | 0.249833 | 0.266361 | 0.241318 | 0.274902 | 0.424023 | 0.598792 | 0.606189 | 0.495993 | 0.779949 | 1.709943 | 3.244650 | 2.721369 | 1.360679 | 0.869405 | 0.642837 | 0.309825 | 0.218240 | 0.215038 | 0.214680 | 0.219974 | 0.233759 | 0.248855 | 0.285258 | 0.224310 | 0.248720 | 0.236131 | 0.225761 | 0.234602 | 0.215027 | 0.215659 | 0.219953 | 0.217169 | 0.215252 | 0.254069 | 0.515046 | 0.950725 | 7.568813 | 13.250295 | 18.225305 | 14.492952 | 3.314690 | 1.740137 | 1.248451 | 0.711211 | 0.576226 | 0.471679 | 0.369362 | 0.356709 | 0.373246 | 0.304133 | 0.351109 | 0.451823 | 1.450616 | 4.198254 | 5.956790 | 4.441438 | 1.569903 | 0.426518 | 0.240634 | 0.220664 | 0.245145 | 0.248631 | 0.269572 | 0.275502 | 0.319346 | 0.458327 | 1.087910 | 6.224856 | 25.838996 | 64.633395 | 76.160899 | 113.303976 | 105.771496 | 53.444595 | 27.584596 | 20.596578 | 12.714182 | 9.362729 | 2.757099 | 1.293146 | 1.143137 | 1.180724 | 1.121555 | 1.084589 | 1.868235 | 2.895468 | 3.471363 | 3.347043 | 5.326876 | 6.110481 | 6.797816 | 3.867553 | 2.855137 | 2.921665 | 2.914790 | 1.714420 | 1.097302 | 0.652491 | 0.710763 | 0.680554 | 0.722962 | 0.558875 | 0.485887 | 0.459899 | 0.415640 | 0.412685 | 0.533125 | 0.604395 | 0.768310 | 0.891912 | 1.142450 | 1.518968 | 2.030168 | 2.423517 | 4.026339 | 6.544027 | 7.167248 | 4.732492 | 2.887572 | 1.876336 | 1.014064 | 0.717699 | 0.548634 | 0.519291 | 0.572958 | 0.505353 | 0.460394 | 0.469067 | 0.410360 | 0.424408 | 0.454481 | 0.316374 | 0.308120 | 0.249929 | 0.244613 | 0.267194 | 0.286515 | 0.263660 | 0.282715 | 0.300635 | 0.406430 | 0.476728 | 0.392596 | 0.284713 | 0.272701 | 0.274546 | 0.273081 | 0.277886 | 0.288325 | 0.325627 | 0.373072 | 0.359278 | 0.398915 | 0.426390 | 0.637760 | 0.695582 | 0.778820 | 1.145411 | 1.916004 | 1.428188 | 0.813454 | 0.370927 | 0.262703 | 0.237644 | 0.234122 | 0.230875 | 0.263691 | 0.358783 | 0.525500 | 0.543797 | 0.458110 | 0.396332 | 0.404755 | 0.445327 | 0.408346 | 0.337742 | 0.396480 | 0.503643 | 0.575488 | 0.502818 | 0.379342 | 0.352525 | 0.376340 | 0.377079 | 0.360540 | 0.370833 | 0.379469 | 0.401270 | 0.507056 | 0.590424 | 0.657315 | 0.796689 | 0.719777 | 0.701044 | 0.794472 | 0.641135 | 0.576298 | 0.492599 | 0.492596 | 0.622985 | 0.808311 | 0.694382 | 0.680874 | 0.490769 | 0.430031 | 0.320196 | 0.268904 | 0.261088 | 0.292892 | 0.331846 | 0.439850 | 0.489127 | 0.554589 | 0.516666 | 0.479005 | 0.373793 | 0.298001 | 0.244937 | 0.217456 | 0.214689 | 0.216238 | 0.224579 | 0.225836 | 0.215553 | 0.215427 | 0.242196 | 0.268202 | 0.271807 | 0.287502 | 0.316079 | 0.296594 | 0.295673 | 0.284424 | 0.286029 | 0.295685 | 0.276979 | 0.249531 | 0.241043 | 0.238733 | 0.236857 | 0.232769 | 0.226235 | 0.233344 | 0.257737 | 0.300260 | 0.328272 | 0.375559 | 0.493435 | 0.756338 | 1.034918 | 1.035551 | 1.107921 | 0.972454 | 0.854419 | 0.568928 | 0.386862 | 0.307683 | 0.278852 | 0.251310 | 0.238033 | 0.219893 | 0.219732 | 0.220356 | 0.235264 | 0.244921 | 0.231822 | 0.225533 | 0.217697 | 0.214715 | 0.218105 | 0.253344 | 0.330692 | 0.355249 | 0.299896 | 0.277048 | 0.254888 | 0.244177 |
| left posterior | 25.211720 | 13.938797 | 6.859280 | 9.391477 | 5.727229 | 2.794293 | 1.327867 | 0.330611 | 0.232555 | 0.226209 | 0.214636 | 0.222288 | 0.237267 | 0.229864 | 0.263294 | 1.198874 | 2.109179 | 3.703065 | 7.873684 | 7.027637 | 22.128852 | 3.765717 | 0.275628 | 0.246860 | 0.356883 | 0.712710 | 0.660648 | 0.466139 | 0.428523 | 0.290131 | 0.214974 | 0.252662 | 0.346123 | 0.328345 | 0.255242 | 0.256105 | 0.277005 | 0.264243 | 0.281483 | 0.278958 | 0.308026 | 0.410840 | 0.594571 | 0.751506 | 0.528812 | 0.307014 | 0.237045 | 0.214664 | 0.235998 | 0.267680 | 0.270068 | 0.264377 | 0.262771 | 0.256178 | 0.233737 | 0.244428 | 0.291841 | 0.360811 | 0.362385 | 0.473866 | 0.685433 | 0.954487 | 1.062818 | 0.953981 | 1.193212 | 2.679395 | 5.124899 | 7.104403 | 15.373304 | 39.411639 | 165.327808 | 629.124025 | 715.281006 | 735.730622 | 1260.638469 | 787.916483 | 727.118393 | 518.188945 | 494.824780 | 672.051960 | 849.504674 | 524.355076 | 355.529786 | 252.372211 | 80.910433 | 21.065046 | 7.687678 | 6.495285 | 4.721406 | 6.619789 | 9.682095 | 14.716278 | 8.462313 | 5.353972 | 1.639549 | 0.960335 | 0.576963 | 0.360109 | 0.289564 | 0.273185 | 0.263231 | 0.342870 | 0.537097 | 0.985760 | 1.881901 | 3.044699 | 3.771793 | 5.415488 | 4.588530 | 3.812506 | 3.465807 | 3.881520 | 3.316057 | 3.466946 | 3.056842 | 2.779389 | 3.547648 | 6.070094 | 7.411948 | 10.334872 | 16.568539 | 23.815079 | 19.269199 | 16.344220 | 10.863717 | 9.959276 | 8.803520 | 6.210724 | 5.101429 | 5.143010 | 5.387952 | 6.113630 | 5.587491 | 4.807419 | 3.137536 | 3.067830 | 4.962979 | 5.012639 | 4.260118 | 3.545093 | 1.723547 | 1.182000 | 0.708865 | 0.465667 | 0.369507 | 0.322613 | 0.268873 | 0.259867 | 0.231147 | 0.219100 | 0.215093 | 0.218990 | 0.247770 | 0.240882 | 0.235934 | 0.217336 | 0.218997 | 0.233212 | 0.265370 | 0.332044 | 0.335665 | 0.379188 | 0.318945 | 0.281216 | 0.267614 | 0.248713 | 0.228488 | 0.227771 | 0.227545 | 0.259987 | 0.303134 | 0.375150 | 0.455836 | 0.602785 | 0.747699 | 0.865833 | 0.905369 | 0.879252 | 0.779166 | 0.810413 | 0.693511 | 0.678140 | 0.676316 | 0.714190 | 0.721299 | 0.718583 | 0.703952 | 0.786182 | 0.755353 | 0.687395 | 0.579927 | 0.527460 | 0.545547 | 0.655879 | 0.743704 | 0.983686 | 1.586168 | 2.481993 | 3.196036 | 4.141130 | 3.567347 | 3.355981 | 2.458793 | 1.459309 | 0.780342 | 0.650121 | 0.512251 | 0.416976 | 0.341895 | 0.354525 | 0.383303 | 0.510643 | 0.545418 | 0.806846 | 1.755052 | 3.727862 | 7.233095 | 15.863686 | 41.287873 | 101.469661 | 101.486979 | 44.847254 | 19.084275 | 5.439449 | 2.464825 | 0.865283 | 0.650609 | 0.589337 | 0.551499 | 0.587113 | 0.631442 | 0.667108 | 0.729186 | 0.670577 | 0.712324 | 0.884821 | 0.944060 | 1.208115 | 1.395367 | 1.683400 | 1.909000 | 1.928946 | 1.736782 | 1.410317 | 1.090849 | 0.731620 | 0.567697 | 0.421823 | 0.322625 | 0.280465 | 0.257707 | 0.237131 | 0.233648 | 0.225979 | 0.229554 | 0.251588 | 0.262186 | 0.273166 | 0.292407 | 0.334433 | 0.370870 | 0.371962 | 0.351660 | 0.326589 | 0.307710 | 0.285978 | 0.242812 | 0.230010 | 0.224531 | 0.217214 | 0.214851 | 0.214761 | 0.214940 | 0.214698 | 0.215960 | 0.219383 | 0.236783 | 0.251712 | 0.266322 | 0.326526 |
| right posterior | 0.216012 | 0.217490 | 0.215118 | 0.215376 | 0.215157 | 0.215725 | 0.225120 | 0.217054 | 0.215626 | 0.216075 | 0.230204 | 0.222530 | 0.214842 | 0.216196 | 0.214998 | 0.287907 | 0.668988 | 1.487691 | 2.454230 | 3.522202 | 3.138317 | 0.838097 | 0.244900 | 0.247017 | 0.360552 | 0.393390 | 0.304194 | 0.224933 | 0.256027 | 0.394416 | 1.226727 | 3.204340 | 2.128136 | 0.803440 | 0.272591 | 0.214892 | 0.235010 | 0.287751 | 0.326352 | 0.275217 | 0.224239 | 0.219426 | 0.252784 | 0.320691 | 0.704404 | 1.533180 | 2.370118 | 3.635419 | 5.115838 | 4.704093 | 4.465873 | 1.923092 | 0.818749 | 0.582744 | 0.302371 | 0.230506 | 0.215616 | 0.218378 | 0.251815 | 0.316271 | 0.373659 | 0.342266 | 0.335326 | 0.314035 | 0.292339 | 0.289052 | 0.306566 | 0.415522 | 1.031533 | 3.551782 | 17.557350 | 71.030418 | 338.012179 | 1364.491070 | 2048.813880 | 1645.148229 | 780.976642 | 344.161240 | 181.173989 | 84.988180 | 51.221674 | 27.245108 | 7.349858 | 2.463511 | 1.138275 | 1.197820 | 1.012279 | 1.188474 | 2.187197 | 4.385738 | 7.493107 | 9.500242 | 5.842106 | 4.836658 | 2.778388 | 1.227984 | 0.773275 | 0.618775 | 0.549727 | 0.490925 | 0.515534 | 0.701670 | 0.833340 | 1.087853 | 0.918132 | 1.032578 | 1.169541 | 1.253853 | 1.001939 | 1.062875 | 0.969685 | 1.180700 | 1.365714 | 1.584257 | 1.697628 | 2.079366 | 2.522233 | 3.708320 | 5.845389 | 10.924131 | 19.367118 | 35.999111 | 71.662301 | 121.065845 | 115.891867 | 81.781579 | 43.981590 | 17.841142 | 7.489769 | 3.172024 | 2.000899 | 1.495385 | 1.404327 | 1.355247 | 1.598529 | 1.904379 | 2.362341 | 2.722571 | 2.944986 | 2.943382 | 2.937034 | 2.936616 | 2.534348 | 2.137218 | 1.523944 | 1.442838 | 1.209778 | 1.217162 | 1.523295 | 1.546484 | 1.668050 | 2.287327 | 2.266185 | 1.978824 | 1.717392 | 1.313928 | 1.355900 | 1.543553 | 1.852575 | 2.394945 | 3.465265 | 4.493937 | 5.177860 | 4.775221 | 3.275519 | 1.943698 | 1.368242 | 1.134789 | 1.029131 | 1.174727 | 1.364629 | 1.588960 | 1.904612 | 1.809373 | 1.737668 | 2.182380 | 2.670034 | 3.748126 | 4.880997 | 6.383930 | 7.043469 | 6.879930 | 6.993467 | 5.114815 | 4.238988 | 5.450651 | 8.500932 | 11.092199 | 13.002848 | 8.540363 | 7.164924 | 6.383420 | 4.992894 | 3.524501 | 3.207369 | 2.973924 | 4.050911 | 4.844420 | 4.724420 | 3.881346 | 3.022812 | 3.039455 | 3.631105 | 3.372186 | 3.884529 | 3.989332 | 3.717688 | 4.515773 | 3.915092 | 3.107840 | 2.544624 | 1.943250 | 1.792658 | 2.107617 | 2.427262 | 3.119696 | 4.418636 | 5.937833 | 7.261617 | 7.212734 | 4.734320 | 3.515236 | 2.804576 | 2.606177 | 1.988731 | 1.383592 | 1.315945 | 1.543489 | 2.009914 | 2.233956 | 2.154183 | 2.378989 | 2.928804 | 3.418987 | 4.668021 | 6.101691 | 8.368810 | 12.484256 | 16.708836 | 23.711701 | 27.507043 | 31.956219 | 25.635044 | 25.358263 | 15.841906 | 10.312022 | 6.705590 | 4.835608 | 3.245813 | 3.111772 | 2.497473 | 2.404670 | 3.358431 | 5.260287 | 8.055230 | 12.612501 | 16.803899 | 20.145142 | 23.792245 | 25.874609 | 13.595942 | 8.968615 | 5.383672 | 3.021931 | 2.540639 | 1.598485 | 0.913147 | 0.841490 | 0.737011 | 0.607608 | 0.491881 | 0.349175 | 0.296032 | 0.262525 | 0.238649 | 0.230637 | 0.236045 | 0.234586 | 0.241302 | 0.257286 |
| all electrodes | 0.445093 | 0.454941 | 0.662132 | 0.973307 | 1.064153 | 1.024814 | 1.195497 | 2.091969 | 2.130054 | 1.160077 | 0.548315 | 0.388529 | 0.352282 | 0.326532 | 0.370416 | 0.621350 | 0.937591 | 0.883517 | 1.090541 | 2.499417 | 9.200026 | 17.401723 | 0.984298 | 0.263171 | 0.218487 | 0.270630 | 0.506717 | 0.950371 | 1.568082 | 0.510929 | 0.259298 | 0.227467 | 0.254327 | 0.257439 | 0.221249 | 0.215530 | 0.214710 | 0.214719 | 0.217999 | 0.215959 | 0.227538 | 0.334151 | 0.487204 | 0.836649 | 0.983921 | 2.341497 | 5.256516 | 6.995451 | 7.471269 | 3.937129 | 2.945156 | 1.391126 | 0.663686 | 0.495466 | 0.550576 | 0.466249 | 0.934046 | 0.724585 | 0.627523 | 0.579891 | 0.368562 | 0.223560 | 0.218702 | 0.216326 | 0.216658 | 0.216334 | 0.218505 | 0.257820 | 0.438701 | 1.405240 | 5.393450 | 25.564825 | 234.421819 | 1887.029389 | 5919.306847 | 5272.745857 | 6525.741166 | 3617.981963 | 2640.173193 | 982.012914 | 310.418659 | 58.316695 | 9.034660 | 1.364193 | 0.564884 | 0.413289 | 0.272079 | 0.251717 | 0.231870 | 0.236873 | 0.256568 | 0.248958 | 0.227346 | 0.218828 | 0.214691 | 0.219237 | 0.263825 | 0.274878 | 0.327572 | 0.425268 | 0.468896 | 0.452308 | 0.484573 | 0.376832 | 0.335848 | 0.242592 | 0.218526 | 0.214626 | 0.230991 | 0.328762 | 0.464852 | 0.376136 | 0.345646 | 0.426118 | 0.491177 | 0.503709 | 0.458923 | 0.401628 | 0.744724 | 0.878213 | 0.835813 | 0.961622 | 1.126582 | 1.174077 | 1.045258 | 0.660650 | 0.568136 | 0.378858 | 0.303187 | 0.278144 | 0.250227 | 0.257416 | 0.268659 | 0.261315 | 0.265547 | 0.240867 | 0.221946 | 0.220744 | 0.215715 | 0.214626 | 0.218228 | 0.214653 | 0.223905 | 0.239993 | 0.233333 | 0.215167 | 0.216600 | 0.215782 | 0.220399 | 0.258942 | 0.351197 | 0.423346 | 0.391513 | 0.433555 | 0.484189 | 0.588143 | 0.507901 | 0.404560 | 0.275942 | 0.225529 | 0.214727 | 0.214674 | 0.215886 | 0.217896 | 0.220460 | 0.272838 | 0.454530 | 0.648259 | 0.589380 | 0.476997 | 0.418642 | 0.332711 | 0.292372 | 0.245379 | 0.227322 | 0.217852 | 0.215653 | 0.239165 | 0.229623 | 0.236349 | 0.246585 | 0.251553 | 0.235094 | 0.216684 | 0.215191 | 0.214626 | 0.215148 | 0.223954 | 0.233545 | 0.232555 | 0.227006 | 0.246541 | 0.282714 | 0.258344 | 0.224133 | 0.216919 | 0.241269 | 0.262654 | 0.292339 | 0.311256 | 0.289609 | 0.273736 | 0.245062 | 0.241566 | 0.241838 | 0.237827 | 0.234257 | 0.245926 | 0.239197 | 0.221108 | 0.217441 | 0.217701 | 0.224284 | 0.247496 | 0.285766 | 0.354981 | 0.453199 | 0.513041 | 0.455808 | 0.460118 | 0.358008 | 0.289755 | 0.255947 | 0.288729 | 0.360526 | 0.423178 | 0.325773 | 0.299993 | 0.270667 | 0.284403 | 0.289653 | 0.286894 | 0.346349 | 0.599711 | 1.442079 | 3.629974 | 4.030143 | 4.531373 | 4.321519 | 2.280967 | 2.025842 | 1.533272 | 1.422743 | 1.961048 | 2.488257 | 2.842227 | 4.277921 | 2.936564 | 1.506568 | 0.701703 | 0.341358 | 0.246571 | 0.229062 | 0.217596 | 0.217942 | 0.229916 | 0.249601 | 0.250723 | 0.249966 | 0.236138 | 0.242150 | 0.251157 | 0.246764 | 0.230677 | 0.227656 | 0.224073 | 0.214980 | 0.215211 | 0.220927 | 0.226527 | 0.223613 | 0.230773 | 0.245053 | 0.251873 | 0.316770 | 0.372812 | 0.427915 | 0.539090 | 0.527783 | 0.521829 |

Searchlight, spatiotemporal cluster permutation test

|  | start time | stop time | peak time | peak channel | cluster p | peak Cohen's d | direction |
| --- | --- | --- | --- | --- | --- | --- | --- |
| #1 | 95 | 260 | 170 | PO7 | 0.0437 | 1.13891 | positive |
| #2 | 225 | 1155 | 425 | Oz | 0.0098 | 1.144394 | positive |

J) real faces (LOSO) - happy vs sad

  
|  | time window | peak latency | cluster *p* | peak Cohen's *d* |  | | | |
| **all electrodes** |  | | | |  | | | |
|  | | | | | | | | |

Time-resolved classification, cluster permutation tests

|  | **left hemisphere** | | | | **right hemisphere** | | | |
|  | time window | peak latency | cluster *p* | peak Cohen's *d* | time window | peak latency | cluster *p* | peak Cohen's *d* |
| **anterior** |  | | | |  | | | |
| **central** | 500 - 715 ms | 500 ms | 0.0029 | -0.515 | 160 - 300 ms | 225 ms | 0.0177 | 0.6667 |
| **posterior** |  | | | |  | | | |

  

Time-resolved classification, Bayesian statistics

|  | -200 | -195 | -190 | -185 | -180 | -175 | -170 | -165 | -160 | -155 | -150 | -145 | -140 | -135 | -130 | -125 | -120 | -115 | -110 | -105 | -100 | -95 | -90 | -85 | -80 | -75 | -70 | -65 | -60 | -55 | -50 | -45 | -40 | -35 | -30 | -25 | -20 | -15 | -10 | -5 | 0 | 5 | 10 | 15 | 20 | 25 | 30 | 35 | 40 | 45 | 50 | 55 | 60 | 65 | 70 | 75 | 80 | 85 | 90 | 95 | 100 | 105 | 110 | 115 | 120 | 125 | 130 | 135 | 140 | 145 | 150 | 155 | 160 | 165 | 170 | 175 | 180 | 185 | 190 | 195 | 200 | 205 | 210 | 215 | 220 | 225 | 230 | 235 | 240 | 245 | 250 | 255 | 260 | 265 | 270 | 275 | 280 | 285 | 290 | 295 | 300 | 305 | 310 | 315 | 320 | 325 | 330 | 335 | 340 | 345 | 350 | 355 | 360 | 365 | 370 | 375 | 380 | 385 | 390 | 395 | 400 | 405 | 410 | 415 | 420 | 425 | 430 | 435 | 440 | 445 | 450 | 455 | 460 | 465 | 470 | 475 | 480 | 485 | 490 | 495 | 500 | 505 | 510 | 515 | 520 | 525 | 530 | 535 | 540 | 545 | 550 | 555 | 560 | 565 | 570 | 575 | 580 | 585 | 590 | 595 | 600 | 605 | 610 | 615 | 620 | 625 | 630 | 635 | 640 | 645 | 650 | 655 | 660 | 665 | 670 | 675 | 680 | 685 | 690 | 695 | 700 | 705 | 710 | 715 | 720 | 725 | 730 | 735 | 740 | 745 | 750 | 755 | 760 | 765 | 770 | 775 | 780 | 785 | 790 | 795 | 800 | 805 | 810 | 815 | 820 | 825 | 830 | 835 | 840 | 845 | 850 | 855 | 860 | 865 | 870 | 875 | 880 | 885 | 890 | 895 | 900 | 905 | 910 | 915 | 920 | 925 | 930 | 935 | 940 | 945 | 950 | 955 | 960 | 965 | 970 | 975 | 980 | 985 | 990 | 995 | 1000 | 1005 | 1010 | 1015 | 1020 | 1025 | 1030 | 1035 | 1040 | 1045 | 1050 | 1055 | 1060 | 1065 | 1070 | 1075 | 1080 | 1085 | 1090 | 1095 | 1100 | 1105 | 1110 | 1115 | 1120 | 1125 | 1130 | 1135 | 1140 | 1145 | 1150 | 1155 | 1160 | 1165 | 1170 | 1175 | 1180 | 1185 | 1190 | 1195 |
| --- | --- | --- | --- | --- | --- | --- | --- | --- | --- | --- | --- | --- | --- | --- | --- | --- | --- | --- | --- | --- | --- | --- | --- | --- | --- | --- | --- | --- | --- | --- | --- | --- | --- | --- | --- | --- | --- | --- | --- | --- | --- | --- | --- | --- | --- | --- | --- | --- | --- | --- | --- | --- | --- | --- | --- | --- | --- | --- | --- | --- | --- | --- | --- | --- | --- | --- | --- | --- | --- | --- | --- | --- | --- | --- | --- | --- | --- | --- | --- | --- | --- | --- | --- | --- | --- | --- | --- | --- | --- | --- | --- | --- | --- | --- | --- | --- | --- | --- | --- | --- | --- | --- | --- | --- | --- | --- | --- | --- | --- | --- | --- | --- | --- | --- | --- | --- | --- | --- | --- | --- | --- | --- | --- | --- | --- | --- | --- | --- | --- | --- | --- | --- | --- | --- | --- | --- | --- | --- | --- | --- | --- | --- | --- | --- | --- | --- | --- | --- | --- | --- | --- | --- | --- | --- | --- | --- | --- | --- | --- | --- | --- | --- | --- | --- | --- | --- | --- | --- | --- | --- | --- | --- | --- | --- | --- | --- | --- | --- | --- | --- | --- | --- | --- | --- | --- | --- | --- | --- | --- | --- | --- | --- | --- | --- | --- | --- | --- | --- | --- | --- | --- | --- | --- | --- | --- | --- | --- | --- | --- | --- | --- | --- | --- | --- | --- | --- | --- | --- | --- | --- | --- | --- | --- | --- | --- | --- | --- | --- | --- | --- | --- | --- | --- | --- | --- | --- | --- | --- | --- | --- | --- | --- | --- | --- | --- | --- | --- | --- | --- | --- | --- | --- | --- | --- | --- | --- | --- | --- | --- | --- | --- | --- | --- | --- | --- | --- | --- | --- | --- | --- | --- | --- | --- | --- | --- | --- | --- | --- | --- | --- |
| left anterior | 0.714527 | 0.648496 | 0.520998 | 0.443707 | 0.393079 | 0.305345 | 0.224262 | 0.219633 | 0.220690 | 0.239385 | 0.316163 | 0.322113 | 0.237917 | 0.214649 | 0.230585 | 0.215914 | 0.219228 | 0.266463 | 0.231774 | 0.214837 | 0.217839 | 0.237504 | 0.363481 | 0.915534 | 1.119068 | 1.803541 | 3.066387 | 6.140526 | 12.956286 | 9.888153 | 8.871872 | 4.418439 | 0.896413 | 0.276261 | 0.216513 | 0.232568 | 0.239425 | 0.354316 | 0.262333 | 0.214626 | 0.215456 | 0.232433 | 0.234525 | 0.257838 | 0.239961 | 0.261507 | 0.234037 | 0.233772 | 0.275881 | 0.328707 | 0.309303 | 0.282615 | 0.305316 | 0.561686 | 1.045437 | 0.632153 | 0.557626 | 0.654672 | 0.594295 | 0.499626 | 0.415227 | 0.303361 | 0.228930 | 0.215835 | 0.215646 | 0.215052 | 0.218849 | 0.237791 | 0.244205 | 0.246816 | 0.279546 | 0.440195 | 1.571171 | 4.918881 | 11.333612 | 22.596395 | 11.748655 | 4.335835 | 1.109773 | 0.380906 | 0.262419 | 0.217666 | 0.219160 | 0.227384 | 0.214748 | 0.223829 | 0.234271 | 0.236075 | 0.247439 | 0.239878 | 0.220054 | 0.217604 | 0.214753 | 0.218568 | 0.215120 | 0.215789 | 0.214638 | 0.217500 | 0.215253 | 0.214678 | 0.214626 | 0.215143 | 0.220120 | 0.243371 | 0.253607 | 0.242183 | 0.256363 | 0.248791 | 0.233101 | 0.215531 | 0.215543 | 0.220598 | 0.227141 | 0.228768 | 0.258805 | 0.337872 | 0.516558 | 0.757621 | 1.212626 | 0.946761 | 0.642986 | 0.368816 | 0.249123 | 0.217053 | 0.217321 | 0.239374 | 0.261421 | 0.300719 | 0.269724 | 0.229312 | 0.217697 | 0.219229 | 0.229121 | 0.253112 | 0.285916 | 0.297406 | 0.332527 | 0.337455 | 0.318954 | 0.288251 | 0.228755 | 0.215598 | 0.236996 | 0.337854 | 0.438963 | 0.563769 | 0.967604 | 1.273932 | 1.359178 | 1.049497 | 0.602099 | 0.410600 | 0.337324 | 0.268276 | 0.233142 | 0.218251 | 0.216850 | 0.223825 | 0.227957 | 0.241443 | 0.268341 | 0.275478 | 0.302502 | 0.345976 | 0.321456 | 0.375340 | 0.317076 | 0.245743 | 0.225659 | 0.220453 | 0.215646 | 0.214702 | 0.222429 | 0.214877 | 0.222698 | 0.235934 | 0.239847 | 0.241260 | 0.240600 | 0.257835 | 0.253254 | 0.241030 | 0.239245 | 0.222459 | 0.219677 | 0.244973 | 0.246313 | 0.234937 | 0.265752 | 0.322999 | 0.461251 | 0.584606 | 0.815702 | 2.363220 | 5.280026 | 3.778191 | 1.748908 | 0.962512 | 0.726111 | 0.501327 | 0.305026 | 0.257948 | 0.238859 | 0.240972 | 0.252001 | 0.233453 | 0.220757 | 0.217371 | 0.216509 | 0.214775 | 0.220065 | 0.223151 | 0.227359 | 0.231206 | 0.217768 | 0.217388 | 0.273795 | 0.366006 | 0.520466 | 0.600516 | 0.687453 | 0.621213 | 0.472712 | 0.387606 | 0.340253 | 0.281467 | 0.265033 | 0.303188 | 0.311998 | 0.287608 | 0.261949 | 0.275804 | 0.272945 | 0.244656 | 0.220289 | 0.215376 | 0.215843 | 0.225013 | 0.217416 | 0.215747 | 0.217363 | 0.229485 | 0.246247 | 0.261687 | 0.243918 | 0.260842 | 0.336859 | 0.426871 | 0.490102 | 0.458108 | 0.490896 | 0.621971 | 0.638222 | 0.556302 | 0.565402 | 0.529185 | 0.550526 | 0.508945 | 0.424924 | 0.431642 | 0.435044 | 0.466443 | 0.459364 | 0.492297 | 0.508677 | 0.522049 | 0.439155 | 0.369636 | 0.306409 | 0.278361 | 0.258152 | 0.246009 | 0.225545 | 0.216145 | 0.214693 | 0.216411 | 0.220249 | 0.227865 | 0.245434 | 0.239915 |
| right anterior | 0.223952 | 0.258812 | 0.317278 | 0.407640 | 0.435118 | 0.543348 | 0.587091 | 0.900574 | 0.682858 | 0.519141 | 0.284868 | 0.255880 | 0.241751 | 0.255049 | 0.289042 | 0.327410 | 0.302297 | 0.288648 | 0.336261 | 0.353739 | 0.440356 | 0.479138 | 0.594038 | 1.253103 | 6.686594 | 12.176674 | 25.297425 | 15.293490 | 7.555305 | 4.274267 | 0.945464 | 0.329849 | 0.234811 | 0.228058 | 0.239355 | 0.285364 | 0.312268 | 0.319597 | 0.246439 | 0.281898 | 0.249950 | 0.238656 | 0.225250 | 0.242390 | 0.242696 | 0.243875 | 0.224390 | 0.228461 | 0.229684 | 0.233719 | 0.237634 | 0.225746 | 0.232518 | 0.221097 | 0.221902 | 0.230394 | 0.232470 | 0.219716 | 0.240599 | 0.288051 | 0.458367 | 0.760000 | 1.062479 | 1.338324 | 1.253097 | 1.386876 | 1.250391 | 0.764597 | 0.512197 | 0.520421 | 0.629399 | 0.898270 | 0.809141 | 0.799533 | 0.754604 | 0.845544 | 0.538857 | 0.301281 | 0.257861 | 0.292241 | 0.345192 | 0.478630 | 0.611004 | 1.076520 | 4.288289 | 10.129892 | 11.246283 | 11.278940 | 6.095420 | 2.770715 | 1.315883 | 1.137188 | 1.025734 | 0.811395 | 0.448269 | 0.324173 | 0.312738 | 0.319288 | 0.276312 | 0.266950 | 0.287625 | 0.353847 | 0.461445 | 0.559490 | 0.613248 | 0.865694 | 0.820745 | 0.658466 | 0.576765 | 0.375115 | 0.412774 | 0.422124 | 0.351619 | 0.414625 | 0.558260 | 0.636784 | 0.969603 | 0.711251 | 0.835372 | 1.091729 | 0.886373 | 0.583347 | 0.483246 | 0.458977 | 0.732469 | 0.649465 | 0.608084 | 0.398465 | 0.329001 | 0.292531 | 0.247956 | 0.214634 | 0.232868 | 0.301657 | 0.344528 | 0.301058 | 0.295882 | 0.299757 | 0.284543 | 0.340379 | 0.341368 | 0.303969 | 0.306585 | 0.263239 | 0.235565 | 0.214776 | 0.228350 | 0.253238 | 0.332474 | 0.359889 | 0.428953 | 0.463016 | 0.385626 | 0.342144 | 0.265716 | 0.224628 | 0.217570 | 0.216921 | 0.216247 | 0.216533 | 0.215669 | 0.215649 | 0.218787 | 0.217309 | 0.214639 | 0.217844 | 0.217998 | 0.217010 | 0.216016 | 0.226257 | 0.260752 | 0.307772 | 0.477326 | 0.711305 | 0.975408 | 1.176213 | 1.181112 | 1.003964 | 0.786212 | 0.869345 | 1.258473 | 0.993689 | 0.703859 | 0.583934 | 0.401081 | 0.306457 | 0.248983 | 0.220418 | 0.229219 | 0.260475 | 0.332518 | 0.345812 | 0.316377 | 0.381077 | 0.522526 | 0.436489 | 0.434943 | 0.434671 | 0.360663 | 0.333199 | 0.249819 | 0.218411 | 0.224504 | 0.249120 | 0.242699 | 0.215089 | 0.222707 | 0.227912 | 0.224514 | 0.242589 | 0.290155 | 0.245119 | 0.224291 | 0.218774 | 0.246959 | 0.275677 | 0.356621 | 0.303174 | 0.365116 | 0.353999 | 0.344895 | 0.315387 | 0.343704 | 0.373263 | 0.564298 | 0.774234 | 0.671983 | 0.514784 | 0.427541 | 0.387287 | 0.429428 | 0.396539 | 0.537655 | 0.828686 | 0.750270 | 0.858609 | 0.607182 | 0.358541 | 0.303683 | 0.236176 | 0.223966 | 0.279431 | 0.386897 | 0.424830 | 0.473007 | 0.483748 | 0.561117 | 0.721500 | 0.867470 | 0.948593 | 1.659648 | 1.825452 | 2.064053 | 1.331401 | 1.291513 | 1.219768 | 0.969093 | 0.674339 | 0.607201 | 0.576739 | 0.802802 | 0.753642 | 0.637266 | 0.628012 | 0.748488 | 0.735335 | 0.695317 | 0.485574 | 0.377622 | 0.304955 | 0.256906 | 0.225543 | 0.220339 | 0.216560 | 0.217321 | 0.218347 | 0.216365 | 0.216084 | 0.216059 | 0.215162 |
| left central | 0.277295 | 0.359748 | 0.391111 | 0.435193 | 0.739204 | 0.779516 | 0.643400 | 0.649237 | 0.443841 | 0.392575 | 0.492217 | 0.400956 | 0.517428 | 0.539145 | 0.428292 | 0.436699 | 0.435771 | 0.412740 | 0.563929 | 1.271728 | 2.481317 | 2.186468 | 3.078105 | 3.816739 | 12.386590 | 11.706406 | 6.631711 | 1.591319 | 0.620161 | 0.239040 | 0.226817 | 0.274881 | 0.309097 | 0.427945 | 0.253997 | 0.215613 | 0.218602 | 0.232254 | 0.223696 | 0.219750 | 0.246586 | 0.215708 | 0.222946 | 0.221671 | 0.215754 | 0.215866 | 0.220946 | 0.239303 | 0.264822 | 0.276269 | 0.282518 | 0.406031 | 0.617302 | 0.682301 | 0.916504 | 1.499292 | 2.743906 | 4.279696 | 2.345832 | 1.006324 | 0.500421 | 0.471871 | 0.278573 | 0.243014 | 0.237730 | 0.244139 | 0.319633 | 0.591374 | 0.476032 | 0.321649 | 0.240825 | 0.215487 | 0.261285 | 0.355803 | 0.537164 | 0.757389 | 0.732784 | 0.473359 | 0.330302 | 0.294371 | 0.277176 | 0.309383 | 0.258004 | 0.244298 | 0.262458 | 0.240657 | 0.222512 | 0.214626 | 0.274250 | 0.350443 | 0.524869 | 0.611163 | 0.497154 | 0.570234 | 0.624677 | 0.485077 | 0.499315 | 0.557527 | 0.984393 | 1.468903 | 1.689988 | 1.636747 | 1.603498 | 2.411690 | 2.727249 | 1.679468 | 2.600148 | 2.697063 | 3.683339 | 4.301452 | 2.481773 | 1.530219 | 1.792193 | 1.486379 | 2.408809 | 3.335683 | 1.374489 | 0.639576 | 0.423024 | 0.292029 | 0.263719 | 0.220151 | 0.220049 | 0.232664 | 0.232997 | 0.221073 | 0.214680 | 0.218484 | 0.234140 | 0.254680 | 0.304514 | 0.255726 | 0.229983 | 0.221118 | 0.214896 | 0.232405 | 0.262468 | 0.323251 | 0.340112 | 0.647298 | 2.839704 | 14.310639 | 104.564640 | 557.439021 | 1148.377419 | 1792.108972 | 242.454513 | 20.294725 | 5.164472 | 2.030123 | 1.546797 | 2.889022 | 6.462275 | 27.690441 | 188.936994 | 350.543422 | 525.161182 | 697.655933 | 769.717836 | 648.622328 | 126.811349 | 21.490127 | 7.757068 | 4.732828 | 9.000902 | 20.803903 | 46.234591 | 106.365700 | 408.840304 | 454.546240 | 166.677812 | 31.102795 | 6.418630 | 4.410631 | 15.522835 | 32.932721 | 49.817234 | 135.080189 | 128.121910 | 65.663453 | 29.587124 | 5.491152 | 1.736672 | 1.411429 | 0.872763 | 0.630538 | 0.452339 | 0.290431 | 0.246738 | 0.221291 | 0.217871 | 0.224337 | 0.227448 | 0.222716 | 0.214661 | 0.219215 | 0.221128 | 0.225169 | 0.216352 | 0.214767 | 0.216294 | 0.219772 | 0.224014 | 0.221759 | 0.221673 | 0.219063 | 0.217410 | 0.218235 | 0.218914 | 0.219181 | 0.219562 | 0.219283 | 0.216655 | 0.218747 | 0.220694 | 0.218599 | 0.215994 | 0.215090 | 0.226597 | 0.250348 | 0.278645 | 0.294565 | 0.288869 | 0.360719 | 0.447235 | 0.576291 | 0.777358 | 0.999317 | 1.279973 | 1.817105 | 1.588290 | 1.182989 | 0.645148 | 0.457598 | 0.474772 | 0.533344 | 0.648381 | 0.778754 | 0.837050 | 1.120647 | 1.682333 | 2.504876 | 2.384998 | 2.027353 | 1.487439 | 1.353556 | 1.355988 | 1.265625 | 1.151990 | 1.200612 | 1.056795 | 1.026005 | 1.056033 | 1.100194 | 0.830667 | 0.548714 | 0.432425 | 0.443123 | 0.573980 | 0.851350 | 0.861686 | 0.996898 | 0.984854 | 0.812204 | 0.843345 | 0.909986 | 0.889170 | 0.884590 | 1.105325 | 1.816289 | 4.505382 | 10.164961 | 12.653539 | 11.038744 | 14.262612 | 10.872939 | 3.885330 | 2.806653 | 1.536854 | 0.797691 |
| right central | 0.355112 | 0.300239 | 0.270743 | 0.278062 | 0.262456 | 0.247274 | 0.284506 | 0.270945 | 0.286770 | 0.291366 | 0.264966 | 0.231107 | 0.214852 | 0.223100 | 0.215151 | 0.241284 | 0.258439 | 0.216896 | 0.214642 | 0.215882 | 0.237841 | 0.228720 | 0.217597 | 0.259744 | 0.271720 | 0.229841 | 0.216492 | 0.216345 | 0.237969 | 0.280923 | 0.241694 | 0.214742 | 0.218746 | 0.218178 | 0.216133 | 0.236388 | 0.313815 | 0.602043 | 1.457445 | 3.191898 | 2.372025 | 0.747376 | 0.529099 | 0.277623 | 0.214671 | 0.219739 | 0.228345 | 0.214717 | 0.226654 | 0.219517 | 0.222071 | 0.258779 | 0.269734 | 0.325273 | 0.296583 | 0.281935 | 0.293224 | 0.378202 | 0.369335 | 0.287559 | 0.214638 | 0.236960 | 0.261139 | 0.248680 | 0.217220 | 0.225362 | 0.249424 | 0.293656 | 0.258067 | 0.218554 | 0.244133 | 0.451124 | 1.352832 | 2.382266 | 3.129139 | 4.685493 | 3.211416 | 1.594873 | 2.033020 | 3.757666 | 6.128903 | 10.007781 | 4.728776 | 4.300368 | 6.766821 | 12.125811 | 12.278365 | 9.384404 | 5.582510 | 5.978508 | 7.854881 | 15.569040 | 17.882990 | 10.203178 | 2.937311 | 2.408570 | 1.798380 | 1.769286 | 1.629946 | 1.375722 | 1.310638 | 1.258474 | 1.345066 | 1.185658 | 0.921766 | 0.604550 | 0.530102 | 0.460048 | 0.503466 | 0.368109 | 0.276679 | 0.226234 | 0.218443 | 0.215960 | 0.218038 | 0.217007 | 0.226534 | 0.257461 | 0.330662 | 0.450014 | 0.440060 | 0.483994 | 0.503565 | 0.491898 | 0.489078 | 0.444766 | 0.365463 | 0.406224 | 0.339036 | 0.321134 | 0.284470 | 0.262749 | 0.244915 | 0.229927 | 0.214663 | 0.226154 | 0.228031 | 0.220490 | 0.215619 | 0.216538 | 0.229184 | 0.250372 | 0.302456 | 0.376404 | 0.476193 | 0.585774 | 0.684148 | 0.723653 | 0.888892 | 1.377861 | 1.761338 | 1.539296 | 1.553841 | 1.021530 | 0.816589 | 0.620830 | 0.386227 | 0.279178 | 0.250596 | 0.236479 | 0.236060 | 0.236333 | 0.267977 | 0.349065 | 0.401284 | 0.483320 | 0.670320 | 0.707098 | 0.588451 | 0.526856 | 0.360956 | 0.270989 | 0.233841 | 0.215322 | 0.230635 | 0.256492 | 0.293846 | 0.280402 | 0.257850 | 0.243797 | 0.229472 | 0.218270 | 0.214669 | 0.214639 | 0.214749 | 0.215673 | 0.214830 | 0.214961 | 0.224157 | 0.248594 | 0.327267 | 0.464239 | 0.704660 | 1.151967 | 1.367576 | 1.179910 | 1.304595 | 1.582198 | 1.347217 | 0.922795 | 0.643095 | 0.572743 | 0.631905 | 0.483501 | 0.337616 | 0.285503 | 0.247305 | 0.242434 | 0.230832 | 0.221659 | 0.233067 | 0.250401 | 0.248142 | 0.259922 | 0.242346 | 0.222477 | 0.217025 | 0.214833 | 0.214631 | 0.216665 | 0.215711 | 0.221842 | 0.234549 | 0.225144 | 0.217717 | 0.216336 | 0.226248 | 0.221730 | 0.230446 | 0.238416 | 0.224828 | 0.217980 | 0.217732 | 0.220786 | 0.246603 | 0.270469 | 0.313774 | 0.361827 | 0.339053 | 0.315074 | 0.293758 | 0.244158 | 0.217155 | 0.218157 | 0.225976 | 0.223508 | 0.228566 | 0.237155 | 0.226829 | 0.214933 | 0.216836 | 0.214948 | 0.216581 | 0.220200 | 0.222164 | 0.228679 | 0.239092 | 0.265763 | 0.268488 | 0.248376 | 0.229701 | 0.218937 | 0.215189 | 0.215878 | 0.224091 | 0.247173 | 0.244415 | 0.249565 | 0.272675 | 0.301280 | 0.301668 | 0.321983 | 0.338390 | 0.430912 | 0.460963 | 0.380008 | 0.347633 | 0.354408 | 0.345179 | 0.318206 |
| left posterior | 85.060240 | 18.827916 | 7.550530 | 4.802067 | 3.678903 | 1.370789 | 0.425205 | 0.221061 | 0.218275 | 0.328996 | 0.723092 | 2.445178 | 6.320592 | 8.374790 | 2.742733 | 0.868754 | 0.247410 | 0.256545 | 1.431160 | 11.879738 | 80.103695 | 74.305262 | 31.509129 | 5.083900 | 1.819788 | 0.369361 | 0.215615 | 0.216256 | 0.220913 | 0.236511 | 0.234474 | 0.245867 | 0.260410 | 0.255180 | 0.274414 | 0.281633 | 0.238286 | 0.222236 | 0.215823 | 0.216560 | 0.228325 | 0.226803 | 0.229759 | 0.221003 | 0.223146 | 0.214889 | 0.243661 | 0.298507 | 0.346150 | 0.329241 | 0.270846 | 0.252437 | 0.219572 | 0.233963 | 0.310628 | 0.358925 | 0.391065 | 0.264104 | 0.218996 | 0.249932 | 0.478488 | 0.783494 | 0.831613 | 0.988466 | 0.621228 | 0.464713 | 0.272054 | 0.215848 | 0.216876 | 0.214972 | 0.230374 | 0.446968 | 1.293121 | 4.670368 | 26.753655 | 51.810845 | 60.259854 | 36.488242 | 9.262197 | 3.525458 | 2.024631 | 1.198515 | 1.287478 | 1.143476 | 1.142022 | 1.023321 | 0.963058 | 0.866524 | 0.665467 | 0.476932 | 0.426178 | 0.324831 | 0.306451 | 0.282948 | 0.247770 | 0.235295 | 0.261831 | 0.296710 | 0.365683 | 0.375145 | 0.397702 | 0.477714 | 0.503085 | 0.413599 | 0.406932 | 0.431659 | 0.450928 | 0.344764 | 0.261516 | 0.226812 | 0.214686 | 0.217955 | 0.233025 | 0.232701 | 0.214650 | 0.241562 | 0.297770 | 0.387011 | 0.406871 | 0.422181 | 0.387198 | 0.369652 | 0.337307 | 0.320658 | 0.356121 | 0.396831 | 0.460522 | 0.456202 | 0.359248 | 0.274889 | 0.233332 | 0.218635 | 0.215172 | 0.214752 | 0.214626 | 0.214704 | 0.217221 | 0.228775 | 0.249128 | 0.262590 | 0.253476 | 0.257909 | 0.266973 | 0.246352 | 0.230843 | 0.214777 | 0.217017 | 0.220309 | 0.222844 | 0.222618 | 0.225877 | 0.237944 | 0.252672 | 0.273446 | 0.280404 | 0.267172 | 0.271999 | 0.241218 | 0.218902 | 0.216138 | 0.222507 | 0.219461 | 0.215963 | 0.215645 | 0.216325 | 0.215470 | 0.214730 | 0.219484 | 0.217473 | 0.215614 | 0.216092 | 0.222671 | 0.233213 | 0.231432 | 0.227171 | 0.228586 | 0.230209 | 0.230242 | 0.222122 | 0.216970 | 0.214709 | 0.217566 | 0.226172 | 0.244523 | 0.269746 | 0.292895 | 0.312773 | 0.299941 | 0.322809 | 0.359490 | 0.386369 | 0.385426 | 0.348597 | 0.319490 | 0.379946 | 0.419504 | 0.436279 | 0.403480 | 0.362674 | 0.385838 | 0.425930 | 0.407060 | 0.374572 | 0.377573 | 0.471447 | 0.662500 | 0.772758 | 0.962753 | 1.036177 | 1.133930 | 1.149943 | 1.024551 | 0.842287 | 0.897319 | 1.024458 | 1.134568 | 1.169132 | 1.154540 | 1.056191 | 0.859541 | 0.593854 | 0.453171 | 0.355539 | 0.330830 | 0.320204 | 0.347412 | 0.418229 | 0.409749 | 0.414785 | 0.404549 | 0.365827 | 0.334276 | 0.280026 | 0.259174 | 0.284247 | 0.278484 | 0.296990 | 0.327484 | 0.344678 | 0.410417 | 0.429441 | 0.441315 | 0.448838 | 0.482779 | 0.509906 | 0.789932 | 1.241819 | 1.923811 | 2.702151 | 3.051905 | 2.277206 | 1.529919 | 0.797363 | 0.452565 | 0.319038 | 0.251241 | 0.225023 | 0.224114 | 0.247262 | 0.290487 | 0.359493 | 0.467610 | 0.572531 | 0.780484 | 0.925388 | 0.943930 | 0.893022 | 0.919559 | 0.891319 | 0.833632 | 0.813298 | 0.854085 | 0.706147 | 0.559787 | 0.473151 | 0.512993 | 0.685471 | 0.650525 | 0.576454 | 0.580489 |
| right posterior | 0.255614 | 0.253413 | 0.249160 | 0.252076 | 0.241081 | 0.239524 | 0.216262 | 0.226886 | 0.249181 | 0.248606 | 0.269160 | 0.259427 | 0.253890 | 0.225998 | 0.215096 | 0.224609 | 0.230637 | 0.240294 | 0.255020 | 0.240637 | 0.217235 | 0.215890 | 0.225509 | 0.289664 | 0.422642 | 0.670220 | 2.385032 | 20.925438 | 33.560808 | 4.424439 | 0.564228 | 0.295622 | 0.225768 | 0.216251 | 0.279587 | 0.351529 | 0.337382 | 0.293255 | 0.235744 | 0.229316 | 0.232056 | 0.245682 | 0.254305 | 0.290658 | 0.273555 | 0.307086 | 0.292866 | 0.274136 | 0.236348 | 0.219109 | 0.215023 | 0.223281 | 0.233855 | 0.230997 | 0.221559 | 0.220757 | 0.214635 | 0.229794 | 0.453153 | 1.435729 | 3.964664 | 9.607694 | 7.920539 | 2.145412 | 0.445804 | 0.214668 | 0.498298 | 1.807633 | 3.083319 | 1.893710 | 0.504293 | 0.215913 | 0.350725 | 0.998698 | 2.407787 | 3.484627 | 2.960504 | 1.369735 | 0.680909 | 0.367910 | 0.295275 | 0.261920 | 0.238474 | 0.235171 | 0.232090 | 0.232537 | 0.221456 | 0.214626 | 0.231241 | 0.288070 | 0.324091 | 0.318304 | 0.282477 | 0.250456 | 0.242692 | 0.228994 | 0.223930 | 0.216097 | 0.214969 | 0.214841 | 0.215904 | 0.215730 | 0.216391 | 0.224984 | 0.222924 | 0.218684 | 0.217225 | 0.221919 | 0.235390 | 0.243512 | 0.239318 | 0.244297 | 0.230330 | 0.224610 | 0.216196 | 0.214849 | 0.217377 | 0.223336 | 0.224087 | 0.214631 | 0.225592 | 0.239508 | 0.283428 | 0.346705 | 0.547394 | 0.768016 | 0.727983 | 0.661252 | 0.591522 | 0.408324 | 0.319345 | 0.261372 | 0.219528 | 0.215040 | 0.231238 | 0.252765 | 0.288414 | 0.308207 | 0.338408 | 0.304724 | 0.267086 | 0.231485 | 0.217491 | 0.215088 | 0.226469 | 0.241379 | 0.258016 | 0.279939 | 0.289733 | 0.278347 | 0.261344 | 0.233181 | 0.217707 | 0.215107 | 0.234951 | 0.264173 | 0.311383 | 0.338394 | 0.290424 | 0.241382 | 0.232384 | 0.224138 | 0.217740 | 0.214697 | 0.220111 | 0.227218 | 0.233369 | 0.251352 | 0.291384 | 0.278223 | 0.240278 | 0.219659 | 0.266210 | 0.324536 | 0.497290 | 0.758322 | 0.975200 | 1.175482 | 0.747530 | 0.583079 | 0.450040 | 0.309882 | 0.246181 | 0.215920 | 0.217372 | 0.230926 | 0.236602 | 0.271969 | 0.291279 | 0.318797 | 0.322292 | 0.331690 | 0.315565 | 0.319635 | 0.263909 | 0.246675 | 0.237649 | 0.228234 | 0.224988 | 0.222760 | 0.228061 | 0.247121 | 0.260279 | 0.253770 | 0.255705 | 0.244617 | 0.244670 | 0.256185 | 0.269401 | 0.283921 | 0.326040 | 0.352125 | 0.381412 | 0.361262 | 0.327549 | 0.306168 | 0.315692 | 0.326765 | 0.333497 | 0.323849 | 0.376504 | 0.351808 | 0.305943 | 0.238903 | 0.217771 | 0.214891 | 0.214997 | 0.219193 | 0.218346 | 0.214701 | 0.229865 | 0.258874 | 0.271315 | 0.323784 | 0.378888 | 0.417793 | 0.369554 | 0.362177 | 0.321923 | 0.317388 | 0.287023 | 0.272576 | 0.290563 | 0.315395 | 0.313924 | 0.374995 | 0.428861 | 0.537839 | 0.698298 | 0.534727 | 0.487551 | 0.506222 | 0.455426 | 0.383311 | 0.295561 | 0.245483 | 0.231857 | 0.221928 | 0.216328 | 0.214820 | 0.215434 | 0.214676 | 0.214676 | 0.214764 | 0.216460 | 0.229063 | 0.238065 | 0.280419 | 0.348028 | 0.384866 | 0.447760 | 0.439531 | 0.404569 | 0.388050 | 0.344876 | 0.276630 | 0.238619 | 0.229781 | 0.230248 | 0.218430 |
| all electrodes | 0.483593 | 0.264701 | 0.233896 | 0.269132 | 0.230812 | 0.217757 | 0.248145 | 0.358736 | 0.691957 | 0.848663 | 0.512538 | 0.237394 | 0.220207 | 0.248928 | 0.314720 | 0.382862 | 0.340379 | 0.345002 | 0.235657 | 0.251456 | 0.263386 | 0.257168 | 0.242738 | 0.226201 | 0.303207 | 0.434261 | 0.791381 | 0.953179 | 1.979127 | 1.522203 | 0.708983 | 0.428554 | 0.334421 | 0.292671 | 0.222672 | 0.229554 | 0.227140 | 0.221968 | 0.216052 | 0.215587 | 0.215260 | 0.226945 | 0.321456 | 0.382879 | 0.563395 | 0.479410 | 0.371687 | 0.322869 | 0.332102 | 0.316366 | 0.271174 | 0.235813 | 0.234324 | 0.283265 | 0.316417 | 0.292083 | 0.324888 | 0.447772 | 0.364360 | 0.253639 | 0.217775 | 0.229183 | 0.242478 | 0.231663 | 0.216085 | 0.231512 | 0.391460 | 0.359422 | 0.320500 | 0.231400 | 0.238277 | 0.441035 | 1.340316 | 6.195905 | 14.166944 | 24.644775 | 19.607238 | 10.652408 | 8.117448 | 5.201108 | 3.591515 | 4.416456 | 3.227704 | 1.873519 | 1.437752 | 1.678788 | 1.219617 | 1.047904 | 0.625471 | 0.417206 | 0.361070 | 0.361196 | 0.339039 | 0.297051 | 0.263122 | 0.248207 | 0.287619 | 0.360379 | 0.319125 | 0.252422 | 0.250731 | 0.270572 | 0.314574 | 0.299593 | 0.278113 | 0.296609 | 0.341227 | 0.309137 | 0.278542 | 0.238367 | 0.228703 | 0.216099 | 0.215346 | 0.214877 | 0.221712 | 0.255970 | 0.448076 | 0.953555 | 2.601393 | 6.317903 | 5.820467 | 3.420584 | 2.132609 | 1.222724 | 0.712496 | 0.511986 | 0.361444 | 0.268792 | 0.228340 | 0.216737 | 0.214647 | 0.215115 | 0.215141 | 0.215669 | 0.215090 | 0.220302 | 0.237137 | 0.249373 | 0.267501 | 0.268448 | 0.240437 | 0.230402 | 0.230013 | 0.217058 | 0.215340 | 0.214850 | 0.219887 | 0.214771 | 0.214626 | 0.214626 | 0.214860 | 0.214698 | 0.217510 | 0.218153 | 0.219728 | 0.215483 | 0.219841 | 0.222780 | 0.215695 | 0.224202 | 0.232681 | 0.224588 | 0.216913 | 0.218435 | 0.216284 | 0.215323 | 0.215528 | 0.217027 | 0.216133 | 0.215919 | 0.219237 | 0.223118 | 0.250725 | 0.288539 | 0.258428 | 0.246419 | 0.234189 | 0.227454 | 0.218126 | 0.216530 | 0.229367 | 0.228532 | 0.233621 | 0.225180 | 0.252231 | 0.265018 | 0.273882 | 0.270813 | 0.278669 | 0.291352 | 0.334421 | 0.378002 | 0.392250 | 0.290789 | 0.286371 | 0.361756 | 0.451462 | 0.435423 | 0.272948 | 0.230733 | 0.226140 | 0.221132 | 0.214998 | 0.214828 | 0.216671 | 0.234408 | 0.259226 | 0.308487 | 0.385135 | 0.460987 | 0.513399 | 0.499967 | 0.482345 | 0.488785 | 0.539933 | 0.494265 | 0.482725 | 0.401155 | 0.384221 | 0.473005 | 0.621700 | 0.578050 | 0.578987 | 0.398990 | 0.393256 | 0.332667 | 0.248698 | 0.229102 | 0.233410 | 0.240969 | 0.284855 | 0.316956 | 0.316558 | 0.381650 | 0.305809 | 0.242289 | 0.220482 | 0.216167 | 0.214765 | 0.214635 | 0.216863 | 0.215986 | 0.229509 | 0.266068 | 0.284864 | 0.284632 | 0.342388 | 0.478511 | 0.518886 | 0.587730 | 0.583088 | 0.416315 | 0.341545 | 0.278435 | 0.275385 | 0.271546 | 0.261953 | 0.247001 | 0.247679 | 0.259179 | 0.295798 | 0.279862 | 0.259239 | 0.250360 | 0.243914 | 0.229312 | 0.221589 | 0.214857 | 0.222416 | 0.232584 | 0.251540 | 0.286623 | 0.281749 | 0.302803 | 0.272558 | 0.239534 | 0.220680 | 0.217117 | 0.214740 | 0.217966 |

Searchlight, spatiotemporal cluster permutation test

No significant clusters observed.

K) real faces (LOSO) - neutral vs happy

  
|  | time window | peak latency | cluster *p* | peak Cohen's *d* |  | | | |  |
| **all electrodes** | 250 - 365 ms | 280 ms | 0.0271 | 0.8811 |  | | | |  |
 785 - 890 ms | 820 ms | 0.0427 | 0.5888 |  | | | ||  |  | | | | | | | | |

Time-resolved classification, cluster permutation tests

|  | **left hemisphere** | | | | **right hemisphere** | | | |  |
|  | time window | peak latency | cluster *p* | peak Cohen's *d* | time window | peak latency | cluster *p* | peak Cohen's *d* |  |
| **anterior** |  | | | | 235 - 315 ms | 280 ms | 0.0375 | 0.9373 |  |
| **central** |  | | | | 250 - 370 ms | 305 ms | 0.0209 | 0.6268 |  |
| **posterior** | 235 - 430 ms | 280 ms | 0.0084 | 1.1 | 105 - 165 ms | 135 ms | 0.0449 | 1.1806 |  |
  | | | | 245 - 495 ms | 280 ms | 0.0053 | 0.9535 |  | | | | 770 - 1005 ms | 875 ms | 0.0082 | 0.7442 |

  

Time-resolved classification, Bayesian statistics

|  | -200 | -195 | -190 | -185 | -180 | -175 | -170 | -165 | -160 | -155 | -150 | -145 | -140 | -135 | -130 | -125 | -120 | -115 | -110 | -105 | -100 | -95 | -90 | -85 | -80 | -75 | -70 | -65 | -60 | -55 | -50 | -45 | -40 | -35 | -30 | -25 | -20 | -15 | -10 | -5 | 0 | 5 | 10 | 15 | 20 | 25 | 30 | 35 | 40 | 45 | 50 | 55 | 60 | 65 | 70 | 75 | 80 | 85 | 90 | 95 | 100 | 105 | 110 | 115 | 120 | 125 | 130 | 135 | 140 | 145 | 150 | 155 | 160 | 165 | 170 | 175 | 180 | 185 | 190 | 195 | 200 | 205 | 210 | 215 | 220 | 225 | 230 | 235 | 240 | 245 | 250 | 255 | 260 | 265 | 270 | 275 | 280 | 285 | 290 | 295 | 300 | 305 | 310 | 315 | 320 | 325 | 330 | 335 | 340 | 345 | 350 | 355 | 360 | 365 | 370 | 375 | 380 | 385 | 390 | 395 | 400 | 405 | 410 | 415 | 420 | 425 | 430 | 435 | 440 | 445 | 450 | 455 | 460 | 465 | 470 | 475 | 480 | 485 | 490 | 495 | 500 | 505 | 510 | 515 | 520 | 525 | 530 | 535 | 540 | 545 | 550 | 555 | 560 | 565 | 570 | 575 | 580 | 585 | 590 | 595 | 600 | 605 | 610 | 615 | 620 | 625 | 630 | 635 | 640 | 645 | 650 | 655 | 660 | 665 | 670 | 675 | 680 | 685 | 690 | 695 | 700 | 705 | 710 | 715 | 720 | 725 | 730 | 735 | 740 | 745 | 750 | 755 | 760 | 765 | 770 | 775 | 780 | 785 | 790 | 795 | 800 | 805 | 810 | 815 | 820 | 825 | 830 | 835 | 840 | 845 | 850 | 855 | 860 | 865 | 870 | 875 | 880 | 885 | 890 | 895 | 900 | 905 | 910 | 915 | 920 | 925 | 930 | 935 | 940 | 945 | 950 | 955 | 960 | 965 | 970 | 975 | 980 | 985 | 990 | 995 | 1000 | 1005 | 1010 | 1015 | 1020 | 1025 | 1030 | 1035 | 1040 | 1045 | 1050 | 1055 | 1060 | 1065 | 1070 | 1075 | 1080 | 1085 | 1090 | 1095 | 1100 | 1105 | 1110 | 1115 | 1120 | 1125 | 1130 | 1135 | 1140 | 1145 | 1150 | 1155 | 1160 | 1165 | 1170 | 1175 | 1180 | 1185 | 1190 | 1195 |
| --- | --- | --- | --- | --- | --- | --- | --- | --- | --- | --- | --- | --- | --- | --- | --- | --- | --- | --- | --- | --- | --- | --- | --- | --- | --- | --- | --- | --- | --- | --- | --- | --- | --- | --- | --- | --- | --- | --- | --- | --- | --- | --- | --- | --- | --- | --- | --- | --- | --- | --- | --- | --- | --- | --- | --- | --- | --- | --- | --- | --- | --- | --- | --- | --- | --- | --- | --- | --- | --- | --- | --- | --- | --- | --- | --- | --- | --- | --- | --- | --- | --- | --- | --- | --- | --- | --- | --- | --- | --- | --- | --- | --- | --- | --- | --- | --- | --- | --- | --- | --- | --- | --- | --- | --- | --- | --- | --- | --- | --- | --- | --- | --- | --- | --- | --- | --- | --- | --- | --- | --- | --- | --- | --- | --- | --- | --- | --- | --- | --- | --- | --- | --- | --- | --- | --- | --- | --- | --- | --- | --- | --- | --- | --- | --- | --- | --- | --- | --- | --- | --- | --- | --- | --- | --- | --- | --- | --- | --- | --- | --- | --- | --- | --- | --- | --- | --- | --- | --- | --- | --- | --- | --- | --- | --- | --- | --- | --- | --- | --- | --- | --- | --- | --- | --- | --- | --- | --- | --- | --- | --- | --- | --- | --- | --- | --- | --- | --- | --- | --- | --- | --- | --- | --- | --- | --- | --- | --- | --- | --- | --- | --- | --- | --- | --- | --- | --- | --- | --- | --- | --- | --- | --- | --- | --- | --- | --- | --- | --- | --- | --- | --- | --- | --- | --- | --- | --- | --- | --- | --- | --- | --- | --- | --- | --- | --- | --- | --- | --- | --- | --- | --- | --- | --- | --- | --- | --- | --- | --- | --- | --- | --- | --- | --- | --- | --- | --- | --- | --- | --- | --- | --- | --- | --- | --- | --- | --- | --- | --- | --- | --- |
| left anterior | 0.216767 | 0.216048 | 0.214675 | 0.217082 | 0.217561 | 0.221085 | 0.228660 | 0.248464 | 0.277899 | 0.305419 | 0.299950 | 0.301791 | 0.324614 | 0.250937 | 0.429086 | 1.064167 | 0.571086 | 0.309689 | 0.232631 | 0.228170 | 0.214963 | 0.235057 | 0.503593 | 0.524926 | 0.268483 | 0.216058 | 0.345728 | 0.444030 | 0.622915 | 0.316967 | 0.220250 | 0.231331 | 0.761357 | 1.765477 | 4.409968 | 20.444874 | 12.622644 | 3.231314 | 1.183760 | 0.430640 | 0.435837 | 0.683222 | 0.657741 | 0.551650 | 1.090455 | 2.634807 | 5.470011 | 15.700521 | 72.366532 | 152.201998 | 281.293650 | 297.819619 | 49.018432 | 13.860173 | 2.630078 | 0.601707 | 0.508729 | 0.642211 | 0.680377 | 0.578257 | 0.465260 | 0.404872 | 0.289352 | 0.221269 | 0.221876 | 0.229914 | 0.217658 | 0.215585 | 0.232141 | 0.373036 | 0.723974 | 1.136720 | 1.674323 | 2.502587 | 5.077839 | 9.922415 | 7.344921 | 8.673078 | 16.873356 | 76.366892 | 178.096465 | 68.357381 | 13.727746 | 2.909200 | 0.763753 | 0.458889 | 0.236160 | 0.220425 | 0.312276 | 0.506499 | 1.083515 | 2.166115 | 4.732873 | 6.323379 | 8.278512 | 12.279212 | 6.802059 | 3.031392 | 1.679500 | 1.037071 | 0.727315 | 0.588931 | 0.341389 | 0.237568 | 0.221744 | 0.216352 | 0.214937 | 0.216896 | 0.219694 | 0.216611 | 0.327294 | 0.436830 | 0.399531 | 0.358325 | 0.310237 | 0.260595 | 0.232434 | 0.214701 | 0.220374 | 0.215876 | 0.215691 | 0.225025 | 0.238434 | 0.249774 | 0.283243 | 0.325742 | 0.361544 | 0.414093 | 0.501456 | 0.585338 | 0.730329 | 0.802422 | 0.810127 | 0.821328 | 0.797517 | 0.941041 | 0.802490 | 0.672214 | 0.622559 | 0.580270 | 0.435827 | 0.372602 | 0.313571 | 0.350083 | 0.348250 | 0.304705 | 0.249829 | 0.241893 | 0.221730 | 0.217485 | 0.214719 | 0.215009 | 0.215272 | 0.218425 | 0.232789 | 0.246336 | 0.238088 | 0.223420 | 0.222901 | 0.218797 | 0.216204 | 0.236022 | 0.309079 | 0.339304 | 0.350093 | 0.351839 | 0.370895 | 0.324889 | 0.302281 | 0.272749 | 0.265926 | 0.245227 | 0.233252 | 0.219976 | 0.222493 | 0.223083 | 0.222454 | 0.223517 | 0.248979 | 0.270487 | 0.297625 | 0.355351 | 0.491828 | 0.756502 | 1.171222 | 1.661094 | 3.192411 | 5.739475 | 6.184555 | 4.968234 | 3.927312 | 3.381093 | 2.088695 | 1.560371 | 1.076187 | 1.095513 | 0.941868 | 0.846464 | 0.841570 | 0.952378 | 0.772911 | 0.839308 | 0.849264 | 0.876619 | 0.925347 | 0.724794 | 0.657053 | 0.732689 | 0.788666 | 0.831421 | 0.960572 | 1.103929 | 2.007256 | 2.109300 | 2.315262 | 2.056502 | 1.395739 | 1.035159 | 0.827647 | 0.617627 | 0.620465 | 0.664601 | 0.768073 | 0.944719 | 0.947154 | 1.007570 | 1.000512 | 1.032421 | 0.767452 | 0.583394 | 0.418951 | 0.467685 | 0.376592 | 0.258843 | 0.221765 | 0.224810 | 0.240324 | 0.298989 | 0.265929 | 0.247353 | 0.276221 | 0.379707 | 0.452571 | 0.391120 | 0.281097 | 0.318148 | 0.410164 | 0.453876 | 0.454049 | 0.356610 | 0.412426 | 0.714206 | 0.961144 | 1.130113 | 1.232228 | 1.280600 | 1.556824 | 1.035590 | 0.873490 | 0.631439 | 0.598088 | 0.570266 | 0.549739 | 0.448761 | 0.459382 | 0.437961 | 0.434762 | 0.358517 | 0.326100 | 0.280538 | 0.246723 | 0.230295 | 0.221512 | 0.216990 | 0.217870 | 0.219232 | 0.217897 | 0.222879 | 0.230098 | 0.233854 |
| right anterior | 0.645642 | 0.655214 | 0.530476 | 0.526128 | 0.426506 | 0.294919 | 0.249451 | 0.216664 | 0.215169 | 0.218928 | 0.293683 | 0.481790 | 1.097306 | 1.995741 | 0.823910 | 0.336402 | 0.234858 | 0.215492 | 0.219846 | 0.224884 | 0.265146 | 0.226834 | 0.216063 | 0.216821 | 0.219664 | 0.284152 | 0.405284 | 0.646602 | 1.636684 | 2.745762 | 1.679914 | 0.459794 | 0.214643 | 0.245736 | 0.461797 | 0.799553 | 0.905588 | 0.568996 | 0.360259 | 0.225317 | 0.222579 | 0.238044 | 0.270971 | 0.374366 | 0.673314 | 0.704159 | 0.840024 | 0.682759 | 0.464073 | 0.519567 | 0.442122 | 0.251298 | 0.242731 | 0.229143 | 0.214689 | 0.222646 | 0.220265 | 0.215179 | 0.227636 | 0.238640 | 0.233120 | 0.233058 | 0.249518 | 0.216819 | 0.242389 | 0.474607 | 0.772161 | 1.071584 | 0.866827 | 0.816662 | 0.510605 | 0.395212 | 0.329320 | 0.267903 | 0.223643 | 0.225549 | 0.234500 | 0.218069 | 0.219583 | 0.255069 | 0.265451 | 0.242296 | 0.220848 | 0.214851 | 0.255504 | 0.474686 | 1.197196 | 2.899043 | 6.676461 | 19.624265 | 137.632762 | 952.353671 | 1146.460785 | 906.325433 | 359.176287 | 284.790791 | 215.843113 | 71.207978 | 16.204841 | 6.888845 | 4.152977 | 3.889742 | 2.168939 | 1.525231 | 1.264094 | 1.213244 | 1.283397 | 1.461198 | 1.233313 | 2.223732 | 2.832682 | 2.984848 | 3.465155 | 2.735954 | 1.697774 | 1.827263 | 1.269846 | 1.138509 | 1.287659 | 1.243982 | 1.297895 | 1.887013 | 1.747877 | 1.444328 | 1.321665 | 1.791382 | 1.462797 | 1.309239 | 0.987776 | 0.821529 | 1.052581 | 1.209333 | 0.889523 | 1.149753 | 1.590862 | 2.733644 | 3.486039 | 1.655674 | 0.901086 | 0.484191 | 0.358448 | 0.312337 | 0.273611 | 0.234970 | 0.229560 | 0.223706 | 0.234984 | 0.234910 | 0.223198 | 0.214963 | 0.214708 | 0.216729 | 0.225915 | 0.238497 | 0.235221 | 0.224224 | 0.218908 | 0.217684 | 0.215405 | 0.214652 | 0.220934 | 0.222612 | 0.222720 | 0.230717 | 0.251899 | 0.276755 | 0.310271 | 0.278628 | 0.287602 | 0.260025 | 0.239338 | 0.226913 | 0.222997 | 0.231248 | 0.243512 | 0.263182 | 0.287470 | 0.286851 | 0.266792 | 0.253640 | 0.247188 | 0.233397 | 0.215778 | 0.214653 | 0.214731 | 0.217191 | 0.230248 | 0.237473 | 0.324203 | 0.661826 | 1.209556 | 1.820212 | 0.916677 | 0.592183 | 0.742041 | 0.810611 | 1.065879 | 1.307281 | 1.664345 | 3.854752 | 4.829327 | 3.917699 | 2.281637 | 1.568151 | 1.562328 | 1.221333 | 1.024029 | 1.121180 | 0.889359 | 0.721647 | 0.538844 | 0.369759 | 0.323849 | 0.307267 | 0.293320 | 0.281182 | 0.309023 | 0.322619 | 0.369823 | 0.370719 | 0.314923 | 0.384090 | 0.453553 | 0.643833 | 1.211443 | 1.428823 | 1.528661 | 2.064870 | 1.478770 | 1.596783 | 1.269418 | 0.991386 | 0.969107 | 0.986723 | 0.964501 | 1.018844 | 0.703566 | 0.614071 | 0.532888 | 0.539358 | 0.667402 | 0.825023 | 1.042810 | 1.271872 | 1.508078 | 1.700239 | 2.047379 | 1.647497 | 1.510734 | 1.219781 | 1.503627 | 1.333636 | 1.211344 | 0.942525 | 1.072332 | 1.255013 | 1.665528 | 1.925483 | 2.294569 | 3.093329 | 3.722368 | 3.312551 | 3.170639 | 3.380194 | 3.180999 | 2.492295 | 1.592691 | 1.152390 | 0.829909 | 0.532146 | 0.349724 | 0.262896 | 0.237061 | 0.219996 | 0.221027 | 0.221696 | 0.222876 | 0.230933 | 0.239311 | 0.235760 |
| left central | 1.318564 | 1.114589 | 1.123223 | 0.881754 | 1.048795 | 0.408062 | 0.236186 | 0.220581 | 0.237965 | 0.248774 | 0.265538 | 0.315742 | 0.392078 | 0.289440 | 0.228706 | 0.291541 | 1.250734 | 5.357191 | 1.649258 | 0.695492 | 0.522706 | 0.593499 | 0.550629 | 0.241417 | 0.224901 | 0.233678 | 0.577488 | 0.699442 | 0.699111 | 0.933072 | 0.680394 | 0.291214 | 0.232258 | 0.524107 | 0.733747 | 0.538376 | 0.405867 | 0.445008 | 0.321378 | 0.215768 | 0.221256 | 0.223557 | 0.219344 | 0.220242 | 0.230325 | 0.226229 | 0.215015 | 0.217653 | 0.216410 | 0.220071 | 0.253471 | 0.265340 | 0.277076 | 0.236736 | 0.216049 | 0.214952 | 0.218746 | 0.224526 | 0.217091 | 0.217413 | 0.269727 | 0.309015 | 0.298877 | 0.229798 | 0.216603 | 0.284936 | 0.492679 | 1.168621 | 1.175304 | 0.804435 | 0.391951 | 0.272897 | 0.233981 | 0.226110 | 0.227846 | 0.250005 | 0.275512 | 0.342999 | 0.361219 | 0.387871 | 0.456177 | 0.405721 | 0.305705 | 0.280106 | 0.247149 | 0.272839 | 0.251281 | 0.228278 | 0.293310 | 0.407712 | 0.495168 | 0.487375 | 0.457448 | 0.590260 | 0.796023 | 0.934586 | 1.589724 | 1.948309 | 3.505103 | 5.610861 | 5.352896 | 2.863740 | 1.324148 | 0.726494 | 0.611005 | 0.334537 | 0.247095 | 0.214938 | 0.218049 | 0.219425 | 0.240238 | 0.233802 | 0.236351 | 0.220197 | 0.217305 | 0.214634 | 0.214933 | 0.214764 | 0.214890 | 0.215386 | 0.224769 | 0.238339 | 0.250465 | 0.246455 | 0.242116 | 0.227702 | 0.241286 | 0.266106 | 0.317124 | 0.379346 | 0.469931 | 0.462355 | 0.477194 | 0.390880 | 0.280603 | 0.234793 | 0.223238 | 0.221290 | 0.227483 | 0.229610 | 0.223548 | 0.237657 | 0.263604 | 0.280544 | 0.243403 | 0.235023 | 0.235941 | 0.253894 | 0.244783 | 0.240953 | 0.243537 | 0.251927 | 0.254777 | 0.252319 | 0.242321 | 0.242952 | 0.236704 | 0.232765 | 0.241198 | 0.244587 | 0.241542 | 0.244098 | 0.235739 | 0.229115 | 0.225603 | 0.221047 | 0.215154 | 0.216812 | 0.225633 | 0.237861 | 0.246604 | 0.310459 | 0.361793 | 0.322036 | 0.296337 | 0.281392 | 0.263960 | 0.255970 | 0.243410 | 0.237701 | 0.233605 | 0.215683 | 0.214633 | 0.215604 | 0.219244 | 0.226933 | 0.221969 | 0.224482 | 0.223634 | 0.228073 | 0.222839 | 0.216281 | 0.216184 | 0.233024 | 0.242316 | 0.257099 | 0.297675 | 0.405369 | 0.535659 | 0.614707 | 0.547220 | 0.614172 | 0.619883 | 0.568469 | 0.566832 | 0.595285 | 0.694729 | 0.848060 | 0.830126 | 0.735051 | 0.657048 | 0.568516 | 0.476088 | 0.375424 | 0.293090 | 0.261007 | 0.237659 | 0.228939 | 0.220005 | 0.217213 | 0.215329 | 0.214633 | 0.215502 | 0.215810 | 0.214723 | 0.218305 | 0.221038 | 0.214651 | 0.215602 | 0.214995 | 0.216474 | 0.216854 | 0.226082 | 0.260459 | 0.322018 | 0.337376 | 0.272746 | 0.219855 | 0.221583 | 0.302437 | 0.465650 | 0.528178 | 0.492190 | 0.423732 | 0.345409 | 0.325229 | 0.274851 | 0.232330 | 0.221427 | 0.218358 | 0.214636 | 0.220403 | 0.238403 | 0.255103 | 0.268841 | 0.301950 | 0.328343 | 0.378545 | 0.493301 | 0.664695 | 0.726794 | 0.665293 | 0.545979 | 0.530207 | 0.483760 | 0.340934 | 0.279261 | 0.253846 | 0.254778 | 0.263225 | 0.253395 | 0.224334 | 0.221589 | 0.247996 | 0.329816 | 0.353144 | 0.371899 | 0.405198 | 0.574226 | 1.079360 |
| right central | 0.285595 | 0.277041 | 0.226085 | 0.214883 | 0.233121 | 0.241867 | 0.224363 | 0.237430 | 0.266032 | 0.241584 | 0.217462 | 0.218372 | 0.239148 | 0.225357 | 0.215197 | 0.316347 | 0.534433 | 0.382652 | 0.300739 | 0.375834 | 0.369650 | 0.310805 | 0.236669 | 0.215084 | 0.216722 | 0.225569 | 0.227523 | 0.282597 | 0.433272 | 0.461292 | 0.391550 | 0.348275 | 0.230923 | 0.227179 | 0.219989 | 0.217904 | 0.224479 | 0.228884 | 0.221027 | 0.218567 | 0.214927 | 0.220291 | 0.252058 | 0.297163 | 0.349036 | 0.413459 | 0.576385 | 0.551862 | 0.397229 | 0.316927 | 0.256674 | 0.222497 | 0.215206 | 0.234851 | 0.253706 | 0.260530 | 0.291471 | 0.292768 | 0.298158 | 0.393331 | 0.503367 | 0.490964 | 0.356379 | 0.219292 | 0.238316 | 0.334527 | 0.659216 | 1.096005 | 1.757719 | 1.288528 | 0.524166 | 0.337690 | 0.284639 | 0.258362 | 0.235518 | 0.224964 | 0.218023 | 0.217382 | 0.223769 | 0.243913 | 0.274541 | 0.338982 | 0.472846 | 0.912854 | 1.151636 | 0.903604 | 0.593314 | 0.538523 | 0.775447 | 1.231674 | 1.906242 | 2.974987 | 4.164521 | 6.228066 | 5.929230 | 6.317544 | 4.268603 | 2.599250 | 2.619868 | 3.422141 | 5.199055 | 8.139022 | 6.555481 | 4.239122 | 4.105535 | 3.984956 | 3.118757 | 2.754313 | 2.043026 | 2.693261 | 4.304583 | 4.211563 | 2.977702 | 2.551656 | 1.827701 | 1.137830 | 0.713096 | 0.637886 | 0.524000 | 0.591363 | 0.591799 | 0.536631 | 0.630339 | 0.670233 | 0.583498 | 0.425279 | 0.277228 | 0.235346 | 0.227332 | 0.219710 | 0.225190 | 0.238902 | 0.255225 | 0.289491 | 0.303340 | 0.289275 | 0.293928 | 0.290166 | 0.256755 | 0.248528 | 0.237281 | 0.223970 | 0.221203 | 0.222427 | 0.218899 | 0.215606 | 0.214861 | 0.216301 | 0.218198 | 0.229175 | 0.228002 | 0.229649 | 0.219174 | 0.215684 | 0.214771 | 0.214711 | 0.219645 | 0.260784 | 0.324047 | 0.367548 | 0.345507 | 0.343472 | 0.395400 | 0.444615 | 0.328161 | 0.267660 | 0.222018 | 0.217358 | 0.217742 | 0.238898 | 0.292203 | 0.344290 | 0.421173 | 0.524517 | 0.565930 | 0.527344 | 0.393881 | 0.351168 | 0.347494 | 0.314375 | 0.241475 | 0.223833 | 0.214725 | 0.215268 | 0.220288 | 0.214867 | 0.218329 | 0.257460 | 0.329628 | 0.354741 | 0.297681 | 0.241193 | 0.216761 | 0.226627 | 0.228401 | 0.220354 | 0.214989 | 0.214709 | 0.217343 | 0.227392 | 0.244175 | 0.297872 | 0.369736 | 0.377219 | 0.394802 | 0.432530 | 0.353229 | 0.266039 | 0.219628 | 0.214730 | 0.218877 | 0.237252 | 0.285295 | 0.281510 | 0.248258 | 0.231014 | 0.238419 | 0.235472 | 0.227497 | 0.217379 | 0.214967 | 0.214800 | 0.217209 | 0.242059 | 0.285705 | 0.322088 | 0.302202 | 0.286979 | 0.263489 | 0.241522 | 0.231928 | 0.233706 | 0.240413 | 0.230854 | 0.218380 | 0.214786 | 0.222257 | 0.264351 | 0.357164 | 0.391460 | 0.342632 | 0.288065 | 0.230180 | 0.215288 | 0.221699 | 0.251248 | 0.252563 | 0.241448 | 0.242028 | 0.250496 | 0.265539 | 0.280060 | 0.255074 | 0.236109 | 0.227985 | 0.219916 | 0.214704 | 0.216065 | 0.214635 | 0.230769 | 0.297013 | 0.403416 | 0.428778 | 0.330469 | 0.304784 | 0.240062 | 0.214640 | 0.218326 | 0.221786 | 0.214784 | 0.221144 | 0.221011 | 0.215098 | 0.216033 | 0.252429 | 0.335622 | 0.531246 | 0.737556 | 0.850132 | 0.880434 |
| left posterior | 0.368959 | 0.333695 | 0.474280 | 0.859855 | 1.317804 | 2.449029 | 1.051951 | 0.616205 | 0.461977 | 0.236702 | 0.226149 | 0.308238 | 0.385528 | 0.355998 | 0.331183 | 0.276276 | 0.226004 | 0.216077 | 0.338665 | 0.525668 | 1.125577 | 1.045502 | 0.327895 | 0.221554 | 0.224895 | 0.324832 | 0.685619 | 1.449417 | 1.765422 | 1.080037 | 0.785075 | 0.520027 | 0.352678 | 0.314369 | 0.295969 | 0.367949 | 0.399793 | 0.310541 | 0.289715 | 0.436821 | 0.735871 | 1.342652 | 1.509190 | 1.521156 | 2.580500 | 4.265208 | 3.390632 | 1.982434 | 0.744277 | 0.345082 | 0.250623 | 0.217799 | 0.242456 | 0.329880 | 0.448948 | 0.517787 | 0.543102 | 0.443439 | 0.414846 | 0.362969 | 0.272451 | 0.217398 | 0.623133 | 12.871802 | 288.830232 | 1963.042420 | 6478.828177 | 9900.398024 | 10261.675715 | 4461.456655 | 547.999727 | 40.693809 | 6.169385 | 1.280018 | 0.529255 | 0.331501 | 0.240116 | 0.223965 | 0.220462 | 0.228385 | 0.224493 | 0.216871 | 0.214635 | 0.219077 | 0.252586 | 0.453222 | 1.245792 | 4.455287 | 14.244519 | 46.124308 | 138.254949 | 276.701739 | 510.823746 | 995.853322 | 1359.615943 | 1782.749851 | 1280.708093 | 596.440046 | 263.322892 | 137.918689 | 71.190460 | 59.450612 | 39.771798 | 41.996450 | 57.684719 | 58.094988 | 50.091277 | 74.644403 | 87.279350 | 166.574029 | 212.955175 | 110.105504 | 69.231584 | 27.573199 | 9.197374 | 4.701743 | 2.424326 | 1.557084 | 1.689973 | 1.489199 | 1.445557 | 1.619151 | 2.045167 | 2.395422 | 1.862382 | 1.499942 | 1.417973 | 1.113358 | 0.760617 | 0.682622 | 0.544067 | 0.409685 | 0.312020 | 0.261899 | 0.267699 | 0.273405 | 0.257088 | 0.251221 | 0.259367 | 0.294481 | 0.398592 | 0.460057 | 0.464822 | 0.519542 | 0.572066 | 0.752169 | 0.875210 | 0.799793 | 0.711238 | 0.849164 | 0.867193 | 0.737869 | 0.611931 | 0.598780 | 0.666128 | 0.841448 | 0.826318 | 0.870298 | 0.887917 | 0.872812 | 0.836423 | 0.767708 | 0.825258 | 0.813897 | 0.657364 | 0.605424 | 0.601445 | 0.532150 | 0.472686 | 0.385930 | 0.376236 | 0.377406 | 0.371484 | 0.367268 | 0.434824 | 0.639061 | 0.906341 | 1.314961 | 2.037290 | 2.483863 | 2.092227 | 1.687052 | 1.324893 | 1.117615 | 0.817631 | 0.625912 | 0.496863 | 0.518276 | 0.437509 | 0.378589 | 0.363448 | 0.381196 | 0.387507 | 0.427185 | 0.385807 | 0.422377 | 0.492813 | 0.546749 | 0.657570 | 0.650413 | 0.705892 | 0.686749 | 0.799278 | 0.844316 | 0.831143 | 0.700571 | 0.640197 | 0.584388 | 0.715787 | 0.712776 | 0.762639 | 0.772018 | 0.792645 | 1.211856 | 1.817778 | 1.815559 | 1.577659 | 1.169303 | 0.884853 | 0.678097 | 0.420163 | 0.330709 | 0.345235 | 0.374510 | 0.455421 | 0.649872 | 0.941574 | 1.235390 | 1.650826 | 1.812190 | 1.776659 | 1.524965 | 0.947393 | 0.773307 | 0.663242 | 0.505775 | 0.387207 | 0.333664 | 0.289865 | 0.338682 | 0.380810 | 0.386433 | 0.377661 | 0.368819 | 0.308840 | 0.286851 | 0.253067 | 0.232315 | 0.237485 | 0.238016 | 0.222939 | 0.220314 | 0.219474 | 0.219528 | 0.214894 | 0.216793 | 0.221576 | 0.218570 | 0.214656 | 0.215162 | 0.215201 | 0.224682 | 0.233317 | 0.232555 | 0.231197 | 0.220340 | 0.216558 | 0.214706 | 0.215014 | 0.216204 | 0.220563 | 0.225788 | 0.226336 | 0.224086 | 0.220711 | 0.218864 | 0.222046 | 0.218144 | 0.215575 | 0.216334 |
| right posterior | 0.272185 | 0.264554 | 0.241370 | 0.216965 | 0.237048 | 0.268617 | 0.316584 | 0.283575 | 0.227685 | 0.215168 | 0.251681 | 0.327718 | 0.297459 | 0.382706 | 0.431722 | 0.469617 | 0.385083 | 0.285547 | 0.299568 | 0.357830 | 0.219744 | 0.215138 | 0.218535 | 0.219790 | 0.246147 | 0.218121 | 0.282094 | 0.216720 | 0.234372 | 0.222047 | 0.219718 | 0.253821 | 0.252704 | 0.225848 | 0.231333 | 0.216567 | 0.302083 | 0.745315 | 2.170881 | 10.589218 | 29.373579 | 55.185243 | 48.549190 | 47.737560 | 78.988144 | 124.423674 | 105.028986 | 33.102945 | 4.056110 | 0.785024 | 0.342369 | 0.243395 | 0.228535 | 0.244332 | 0.232632 | 0.252769 | 0.254556 | 0.303530 | 0.371715 | 0.433696 | 0.548501 | 2.392759 | 23.511031 | 442.926210 | 2990.107005 | 7174.718551 | 8632.904415 | 3075.954492 | 1070.045069 | 359.773228 | 157.313545 | 43.679250 | 8.701344 | 2.408646 | 1.088240 | 0.542191 | 0.270933 | 0.216073 | 0.214668 | 0.214668 | 0.219055 | 0.232169 | 0.261677 | 0.286758 | 0.333325 | 0.365892 | 0.499846 | 0.708755 | 1.103937 | 2.114110 | 7.810922 | 18.618382 | 32.853893 | 59.799706 | 138.164158 | 226.974421 | 257.744495 | 206.742067 | 187.697916 | 163.970323 | 155.870337 | 126.510178 | 174.559650 | 225.943261 | 362.646475 | 187.797258 | 126.570769 | 124.847352 | 127.136409 | 107.962914 | 97.329176 | 61.141616 | 59.425951 | 65.346961 | 58.827545 | 45.226500 | 35.020712 | 43.041887 | 58.095806 | 59.099491 | 52.189285 | 54.846134 | 51.207157 | 25.430831 | 7.004355 | 3.685940 | 3.665744 | 4.158287 | 2.524063 | 2.375484 | 2.546350 | 3.511637 | 3.829570 | 3.068473 | 2.194338 | 1.922155 | 1.640252 | 1.457367 | 1.589464 | 1.387932 | 0.916333 | 1.024210 | 1.402935 | 1.761485 | 2.100804 | 2.637725 | 3.705926 | 4.878808 | 2.835113 | 1.692719 | 1.219847 | 1.151154 | 1.003053 | 0.758040 | 0.691801 | 0.905413 | 1.274614 | 2.189422 | 2.357722 | 1.522285 | 1.500910 | 1.478328 | 1.148041 | 0.834385 | 0.657828 | 0.478510 | 0.424249 | 0.380251 | 0.333033 | 0.327458 | 0.331345 | 0.278510 | 0.270419 | 0.262485 | 0.260843 | 0.272968 | 0.263524 | 0.267025 | 0.303009 | 0.355745 | 0.412626 | 0.464454 | 0.461926 | 0.566108 | 0.669395 | 0.687292 | 0.684741 | 0.805669 | 0.957336 | 0.964905 | 0.718405 | 0.752619 | 0.935501 | 1.048759 | 1.415588 | 1.997592 | 3.427377 | 5.827244 | 3.660500 | 2.766100 | 2.682076 | 3.211801 | 4.468154 | 5.904692 | 8.080226 | 14.110263 | 17.144961 | 16.615136 | 12.046873 | 9.678022 | 7.408610 | 7.465310 | 9.147708 | 11.040786 | 17.913267 | 26.986326 | 31.616579 | 40.341783 | 34.620180 | 20.369874 | 13.108273 | 11.980277 | 11.956380 | 11.941102 | 14.086749 | 18.125575 | 28.337529 | 34.539171 | 22.519482 | 17.321519 | 17.600462 | 13.158980 | 10.993193 | 7.091622 | 4.381396 | 4.111429 | 3.862155 | 3.273456 | 2.697266 | 2.334300 | 1.751801 | 1.580049 | 1.277717 | 1.119066 | 1.114361 | 1.141613 | 1.077958 | 1.325364 | 1.718032 | 3.311366 | 3.970825 | 4.017678 | 4.070857 | 5.063816 | 3.146375 | 1.870778 | 1.142289 | 0.923145 | 0.911708 | 0.939146 | 0.942611 | 1.073581 | 1.361565 | 1.348923 | 1.448740 | 1.063148 | 0.638210 | 0.444974 | 0.401107 | 0.326996 | 0.312186 | 0.269666 | 0.289286 | 0.376705 | 0.399102 | 0.357269 | 0.379534 | 0.345429 | 0.382704 | 0.383810 |
| all electrodes | 0.315174 | 0.716610 | 1.077069 | 0.610526 | 0.569301 | 1.317011 | 0.709663 | 0.399778 | 0.267448 | 0.260568 | 0.261968 | 0.239004 | 0.215889 | 0.223377 | 0.218414 | 0.233734 | 0.215138 | 0.215583 | 0.300936 | 0.492024 | 0.369762 | 0.256493 | 0.243243 | 0.361708 | 1.065030 | 0.915607 | 1.326110 | 3.303804 | 4.747596 | 2.793814 | 0.585175 | 0.226566 | 0.250658 | 0.393034 | 0.469506 | 0.668887 | 0.550582 | 0.331053 | 0.257253 | 0.220517 | 0.257552 | 0.247853 | 0.270028 | 0.221562 | 0.231697 | 0.234888 | 0.261463 | 0.340894 | 0.584303 | 0.805513 | 1.018254 | 1.151694 | 3.253419 | 3.148844 | 1.996832 | 0.738414 | 0.481823 | 0.355757 | 0.274094 | 0.237656 | 0.222304 | 0.216769 | 0.233445 | 0.393642 | 1.678261 | 4.974541 | 21.415569 | 37.714438 | 47.726420 | 24.028333 | 6.798832 | 1.227227 | 0.465859 | 0.233092 | 0.222696 | 0.282186 | 0.362668 | 0.405828 | 0.384128 | 0.295755 | 0.227913 | 0.220092 | 0.230620 | 0.293137 | 0.333694 | 0.293297 | 0.254855 | 0.224574 | 0.214935 | 0.313121 | 1.484920 | 9.629940 | 24.688679 | 47.592368 | 56.570845 | 102.730877 | 116.933251 | 115.872608 | 105.288244 | 50.166733 | 28.454592 | 23.814716 | 6.100209 | 2.455097 | 1.881158 | 1.569406 | 2.267433 | 2.676737 | 1.943465 | 2.740712 | 3.954214 | 5.236018 | 4.229265 | 2.005404 | 1.135214 | 0.786729 | 0.596271 | 0.619182 | 0.596108 | 0.732348 | 0.800019 | 0.713288 | 0.787643 | 0.657314 | 0.487122 | 0.329676 | 0.246374 | 0.221826 | 0.220838 | 0.224280 | 0.247660 | 0.287145 | 0.330166 | 0.363232 | 0.482342 | 0.465732 | 0.397439 | 0.342491 | 0.257232 | 0.264119 | 0.262026 | 0.264754 | 0.306216 | 0.367910 | 0.364589 | 0.465983 | 0.470017 | 0.405544 | 0.281364 | 0.236177 | 0.223010 | 0.215434 | 0.215044 | 0.217006 | 0.224865 | 0.249513 | 0.258206 | 0.249610 | 0.260773 | 0.255721 | 0.239211 | 0.217286 | 0.217079 | 0.218195 | 0.221351 | 0.220536 | 0.217871 | 0.215374 | 0.225091 | 0.234455 | 0.224119 | 0.216315 | 0.214936 | 0.223915 | 0.241118 | 0.262686 | 0.255140 | 0.261766 | 0.238990 | 0.215237 | 0.232962 | 0.256411 | 0.278175 | 0.310959 | 0.354164 | 0.378768 | 0.338595 | 0.323381 | 0.363124 | 0.368253 | 0.324302 | 0.330593 | 0.376042 | 0.451727 | 0.421820 | 0.435431 | 0.683256 | 1.441454 | 2.268993 | 2.556788 | 3.752869 | 5.461761 | 5.994730 | 6.165587 | 5.625145 | 5.841095 | 7.163827 | 4.785802 | 4.947039 | 4.161023 | 2.460857 | 2.352603 | 2.837207 | 2.344567 | 2.526492 | 2.687762 | 2.274407 | 1.885237 | 1.409379 | 0.935235 | 1.209946 | 1.342874 | 0.957728 | 1.073320 | 1.898655 | 3.504425 | 6.210922 | 4.284563 | 2.751769 | 2.705882 | 1.987268 | 1.222439 | 0.756178 | 0.522482 | 0.423525 | 0.538578 | 0.646431 | 0.739663 | 0.636853 | 0.605405 | 0.710355 | 0.990537 | 0.872208 | 0.692824 | 0.560370 | 0.664912 | 0.650148 | 0.523325 | 0.374743 | 0.328058 | 0.362582 | 0.547867 | 0.873579 | 0.912903 | 0.819428 | 0.752229 | 0.580708 | 0.397506 | 0.274147 | 0.235844 | 0.263572 | 0.289167 | 0.318888 | 0.351410 | 0.333537 | 0.277500 | 0.233285 | 0.218672 | 0.221842 | 0.219931 | 0.219553 | 0.228788 | 0.253968 | 0.256636 | 0.225348 | 0.215151 | 0.214856 | 0.216119 | 0.218914 | 0.220593 |

Searchlight, spatiotemporal cluster permutation test

|  | start time | stop time | peak time | peak channel | cluster p | peak Cohen's d | direction |
| --- | --- | --- | --- | --- | --- | --- | --- |
| #1 | 215 | 1145 | 290 | P8 | 0.0005 | 1.164651 | positive |

L) real faces (LOSO) - angry vs sad

  
|  | time window | peak latency | cluster *p* | peak Cohen's *d* |  | | | |  |
| **all electrodes** |  | | | |  | | | |  |
|  | | | | | | | | |  |

Time-resolved classification, cluster permutation tests

|  | **left hemisphere** | | | | **right hemisphere** | | | |  |
|  | time window | peak latency | cluster *p* | peak Cohen's *d* | time window | peak latency | cluster *p* | peak Cohen's *d* |  |
| **anterior** |  | | | |  | | | |  |
| **central** | 185 - 290 ms | 255 ms | 0.0179 | 0.9377 | 200 - 290 ms | 235 ms | 0.025 | 0.9198 |  |
 340 - 495 ms | 365 ms | 0.0061 | 0.8882 | 315 - 445 ms | 415 ms | 0.0079 | 0.9321 |  | | | | -200 - -115 ms | -115 ms | 0.0402 | -0.567 ||  | **posterior** |  | | | | 200 - 295 ms | 265 ms | 0.0107 | 1.1306 |
  | | | | 335 - 415 ms | 390 ms | 0.0414 | 0.7291 |

  

Time-resolved classification, Bayesian statistics

|  | -200 | -195 | -190 | -185 | -180 | -175 | -170 | -165 | -160 | -155 | -150 | -145 | -140 | -135 | -130 | -125 | -120 | -115 | -110 | -105 | -100 | -95 | -90 | -85 | -80 | -75 | -70 | -65 | -60 | -55 | -50 | -45 | -40 | -35 | -30 | -25 | -20 | -15 | -10 | -5 | 0 | 5 | 10 | 15 | 20 | 25 | 30 | 35 | 40 | 45 | 50 | 55 | 60 | 65 | 70 | 75 | 80 | 85 | 90 | 95 | 100 | 105 | 110 | 115 | 120 | 125 | 130 | 135 | 140 | 145 | 150 | 155 | 160 | 165 | 170 | 175 | 180 | 185 | 190 | 195 | 200 | 205 | 210 | 215 | 220 | 225 | 230 | 235 | 240 | 245 | 250 | 255 | 260 | 265 | 270 | 275 | 280 | 285 | 290 | 295 | 300 | 305 | 310 | 315 | 320 | 325 | 330 | 335 | 340 | 345 | 350 | 355 | 360 | 365 | 370 | 375 | 380 | 385 | 390 | 395 | 400 | 405 | 410 | 415 | 420 | 425 | 430 | 435 | 440 | 445 | 450 | 455 | 460 | 465 | 470 | 475 | 480 | 485 | 490 | 495 | 500 | 505 | 510 | 515 | 520 | 525 | 530 | 535 | 540 | 545 | 550 | 555 | 560 | 565 | 570 | 575 | 580 | 585 | 590 | 595 | 600 | 605 | 610 | 615 | 620 | 625 | 630 | 635 | 640 | 645 | 650 | 655 | 660 | 665 | 670 | 675 | 680 | 685 | 690 | 695 | 700 | 705 | 710 | 715 | 720 | 725 | 730 | 735 | 740 | 745 | 750 | 755 | 760 | 765 | 770 | 775 | 780 | 785 | 790 | 795 | 800 | 805 | 810 | 815 | 820 | 825 | 830 | 835 | 840 | 845 | 850 | 855 | 860 | 865 | 870 | 875 | 880 | 885 | 890 | 895 | 900 | 905 | 910 | 915 | 920 | 925 | 930 | 935 | 940 | 945 | 950 | 955 | 960 | 965 | 970 | 975 | 980 | 985 | 990 | 995 | 1000 | 1005 | 1010 | 1015 | 1020 | 1025 | 1030 | 1035 | 1040 | 1045 | 1050 | 1055 | 1060 | 1065 | 1070 | 1075 | 1080 | 1085 | 1090 | 1095 | 1100 | 1105 | 1110 | 1115 | 1120 | 1125 | 1130 | 1135 | 1140 | 1145 | 1150 | 1155 | 1160 | 1165 | 1170 | 1175 | 1180 | 1185 | 1190 | 1195 |
| --- | --- | --- | --- | --- | --- | --- | --- | --- | --- | --- | --- | --- | --- | --- | --- | --- | --- | --- | --- | --- | --- | --- | --- | --- | --- | --- | --- | --- | --- | --- | --- | --- | --- | --- | --- | --- | --- | --- | --- | --- | --- | --- | --- | --- | --- | --- | --- | --- | --- | --- | --- | --- | --- | --- | --- | --- | --- | --- | --- | --- | --- | --- | --- | --- | --- | --- | --- | --- | --- | --- | --- | --- | --- | --- | --- | --- | --- | --- | --- | --- | --- | --- | --- | --- | --- | --- | --- | --- | --- | --- | --- | --- | --- | --- | --- | --- | --- | --- | --- | --- | --- | --- | --- | --- | --- | --- | --- | --- | --- | --- | --- | --- | --- | --- | --- | --- | --- | --- | --- | --- | --- | --- | --- | --- | --- | --- | --- | --- | --- | --- | --- | --- | --- | --- | --- | --- | --- | --- | --- | --- | --- | --- | --- | --- | --- | --- | --- | --- | --- | --- | --- | --- | --- | --- | --- | --- | --- | --- | --- | --- | --- | --- | --- | --- | --- | --- | --- | --- | --- | --- | --- | --- | --- | --- | --- | --- | --- | --- | --- | --- | --- | --- | --- | --- | --- | --- | --- | --- | --- | --- | --- | --- | --- | --- | --- | --- | --- | --- | --- | --- | --- | --- | --- | --- | --- | --- | --- | --- | --- | --- | --- | --- | --- | --- | --- | --- | --- | --- | --- | --- | --- | --- | --- | --- | --- | --- | --- | --- | --- | --- | --- | --- | --- | --- | --- | --- | --- | --- | --- | --- | --- | --- | --- | --- | --- | --- | --- | --- | --- | --- | --- | --- | --- | --- | --- | --- | --- | --- | --- | --- | --- | --- | --- | --- | --- | --- | --- | --- | --- | --- | --- | --- | --- | --- | --- | --- | --- | --- | --- | --- |
| left anterior | 0.514354 | 1.548651 | 3.641402 | 2.943323 | 5.547288 | 3.683580 | 1.248408 | 0.571438 | 0.326779 | 0.214642 | 0.391438 | 2.074665 | 19.900674 | 64.158224 | 252.807578 | 869.067739 | 38.789110 | 3.073882 | 0.634800 | 0.240698 | 0.229289 | 0.597970 | 2.261057 | 2.193280 | 2.330149 | 1.902783 | 2.158478 | 1.173551 | 0.417795 | 0.267812 | 0.269931 | 0.246703 | 0.214841 | 0.238887 | 0.248906 | 0.307534 | 0.317727 | 0.416674 | 0.486906 | 0.450223 | 0.307534 | 0.308139 | 0.267737 | 0.279855 | 0.249682 | 0.231769 | 0.219000 | 0.243578 | 0.258917 | 0.245433 | 0.215877 | 0.216414 | 0.218089 | 0.224823 | 0.233189 | 0.231262 | 0.231658 | 0.314866 | 0.394390 | 0.495602 | 0.659142 | 0.642066 | 0.609215 | 0.643221 | 0.472737 | 0.464764 | 0.496025 | 0.922257 | 0.836617 | 0.743319 | 0.789110 | 1.782704 | 2.750190 | 3.887647 | 1.855323 | 2.047225 | 2.803127 | 2.748582 | 0.976893 | 0.753236 | 0.867892 | 1.153199 | 1.600336 | 0.987975 | 0.590774 | 0.571728 | 0.460956 | 0.303007 | 0.246560 | 0.214636 | 0.215217 | 0.215647 | 0.214626 | 0.219560 | 0.218213 | 0.218455 | 0.214698 | 0.224747 | 0.251615 | 0.401472 | 1.120184 | 1.948224 | 2.774975 | 3.241717 | 2.563755 | 1.212901 | 0.660083 | 0.428667 | 0.362062 | 0.284953 | 0.258814 | 0.220609 | 0.217611 | 0.260303 | 0.281698 | 0.312402 | 0.372473 | 0.372692 | 0.307650 | 0.248560 | 0.234425 | 0.258235 | 0.315643 | 0.346919 | 0.372497 | 0.463909 | 0.588327 | 0.655765 | 0.612160 | 0.381141 | 0.253287 | 0.215979 | 0.229180 | 0.276581 | 0.393571 | 0.759039 | 1.009106 | 1.139334 | 1.589655 | 1.282013 | 0.740076 | 0.392217 | 0.273785 | 0.265828 | 0.318390 | 0.297397 | 0.332291 | 0.305354 | 0.289411 | 0.234824 | 0.214906 | 0.215680 | 0.217034 | 0.214752 | 0.214975 | 0.218453 | 0.230880 | 0.248946 | 0.219891 | 0.214638 | 0.217432 | 0.215634 | 0.225775 | 0.264128 | 0.302905 | 0.306360 | 0.290291 | 0.277262 | 0.257355 | 0.248216 | 0.238862 | 0.239767 | 0.253731 | 0.312885 | 0.416392 | 0.532026 | 0.510727 | 0.465807 | 0.434052 | 0.428997 | 0.389097 | 0.391951 | 0.398084 | 0.462279 | 0.344819 | 0.258608 | 0.217307 | 0.222736 | 0.237134 | 0.229436 | 0.240434 | 0.232804 | 0.225428 | 0.215470 | 0.214640 | 0.214983 | 0.220124 | 0.219089 | 0.216018 | 0.219673 | 0.224594 | 0.214933 | 0.220872 | 0.256750 | 0.335031 | 0.506961 | 0.805345 | 1.117370 | 1.341604 | 2.029984 | 1.773770 | 0.758234 | 0.606682 | 0.354865 | 0.263490 | 0.218291 | 0.215025 | 0.237590 | 0.227029 | 0.249238 | 0.225586 | 0.217742 | 0.218169 | 0.222835 | 0.228389 | 0.274267 | 0.321199 | 0.381717 | 0.387709 | 0.348033 | 0.290080 | 0.240973 | 0.215687 | 0.217342 | 0.224980 | 0.228522 | 0.232992 | 0.218175 | 0.214747 | 0.220643 | 0.231698 | 0.256281 | 0.282158 | 0.386178 | 0.386705 | 0.419627 | 0.343310 | 0.300733 | 0.289887 | 0.281732 | 0.249570 | 0.249862 | 0.248797 | 0.254239 | 0.245424 | 0.225369 | 0.216618 | 0.215180 | 0.219415 | 0.221042 | 0.217314 | 0.214995 | 0.225631 | 0.267861 | 0.330937 | 0.381529 | 0.402869 | 0.400637 | 0.320862 | 0.257839 | 0.215190 | 0.226442 | 0.285656 | 0.295553 | 0.350348 | 0.337616 | 0.303335 | 0.261681 | 0.228590 | 0.217573 |
| right anterior | 0.215176 | 0.216653 | 0.228413 | 0.241736 | 0.321563 | 0.513678 | 0.358881 | 0.348656 | 0.279443 | 0.214777 | 0.228182 | 0.260275 | 0.234465 | 0.268283 | 0.707938 | 4.465869 | 128.662150 | 35.470948 | 4.278459 | 0.791003 | 0.216408 | 0.265179 | 0.602213 | 1.009462 | 0.679926 | 0.473057 | 0.345551 | 0.310516 | 0.420684 | 0.312868 | 0.265015 | 0.354946 | 0.287651 | 0.288722 | 0.224691 | 0.214691 | 0.216520 | 0.220311 | 0.215398 | 0.230319 | 0.304329 | 0.288341 | 0.254425 | 0.266520 | 0.214711 | 0.223166 | 0.315465 | 0.424430 | 0.314626 | 0.238911 | 0.221578 | 0.218771 | 0.254412 | 0.348727 | 0.388867 | 0.318946 | 0.290437 | 0.270849 | 0.283725 | 0.299304 | 0.301364 | 0.278319 | 0.280624 | 0.274156 | 0.368815 | 0.395546 | 0.361312 | 0.336081 | 0.335480 | 0.274731 | 0.332602 | 0.297919 | 0.290441 | 0.318562 | 0.371281 | 0.543851 | 1.222686 | 1.180981 | 1.803758 | 2.191733 | 1.851580 | 1.205694 | 0.648502 | 0.358601 | 0.268949 | 0.216117 | 0.235279 | 0.352233 | 0.367838 | 0.328666 | 0.298009 | 0.295078 | 0.236224 | 0.216603 | 0.219838 | 0.219010 | 0.214696 | 0.214971 | 0.222400 | 0.217873 | 0.215759 | 0.218353 | 0.220319 | 0.214688 | 0.222891 | 0.250390 | 0.258632 | 0.248774 | 0.258542 | 0.297694 | 0.327423 | 0.426651 | 0.732118 | 1.414648 | 2.814701 | 4.893546 | 5.017912 | 4.377918 | 2.984235 | 1.346566 | 1.848757 | 3.472950 | 5.876915 | 3.359450 | 1.661530 | 1.316320 | 1.603375 | 0.989574 | 0.699890 | 0.351348 | 0.289880 | 0.306053 | 0.307370 | 0.244474 | 0.216883 | 0.236052 | 0.263070 | 0.272162 | 0.302010 | 0.353745 | 0.380584 | 0.424553 | 0.476371 | 0.672262 | 1.410801 | 3.268826 | 9.210612 | 13.739707 | 7.602029 | 1.199419 | 0.458368 | 0.323423 | 0.435171 | 0.631982 | 0.850115 | 0.623891 | 0.738219 | 0.605673 | 0.527463 | 0.366564 | 0.294457 | 0.248834 | 0.234418 | 0.251048 | 0.326719 | 0.425031 | 0.417382 | 0.339074 | 0.318786 | 0.367587 | 0.346542 | 0.403765 | 0.374817 | 0.360763 | 0.469549 | 0.502475 | 0.433935 | 0.427450 | 0.353283 | 0.384039 | 0.399341 | 0.296877 | 0.261955 | 0.249597 | 0.243241 | 0.243307 | 0.216102 | 0.215504 | 0.218539 | 0.216838 | 0.215908 | 0.215186 | 0.243325 | 0.253015 | 0.316877 | 0.297485 | 0.305595 | 0.312187 | 0.263956 | 0.219760 | 0.226203 | 0.369231 | 0.412583 | 0.495437 | 0.422611 | 0.387635 | 0.420282 | 0.357796 | 0.347862 | 0.444769 | 0.441819 | 0.307512 | 0.247424 | 0.215010 | 0.230884 | 0.260013 | 0.318025 | 0.344289 | 0.266295 | 0.247404 | 0.231107 | 0.220005 | 0.214626 | 0.215341 | 0.215542 | 0.220910 | 0.230569 | 0.236134 | 0.228296 | 0.258643 | 0.266679 | 0.235486 | 0.214754 | 0.227222 | 0.230803 | 0.230008 | 0.230515 | 0.234789 | 0.238432 | 0.217754 | 0.216275 | 0.226129 | 0.271381 | 0.329108 | 0.342050 | 0.386563 | 0.400356 | 0.414332 | 0.386703 | 0.342061 | 0.289718 | 0.252024 | 0.244550 | 0.226240 | 0.215622 | 0.215419 | 0.216010 | 0.214759 | 0.216975 | 0.215214 | 0.221250 | 0.236304 | 0.332983 | 0.354250 | 0.343503 | 0.367864 | 0.428279 | 0.360797 | 0.327516 | 0.234586 | 0.214714 | 0.235850 | 0.293408 | 0.419365 | 0.961497 | 3.260805 | 10.570611 | 13.903520 | 8.737627 | 10.548578 |
| left central | 0.327054 | 0.306972 | 0.311596 | 0.281534 | 0.236713 | 0.241929 | 0.268787 | 0.378254 | 0.515527 | 0.410606 | 0.368719 | 0.460207 | 0.223990 | 0.214653 | 0.232867 | 0.260320 | 0.317330 | 0.420727 | 0.526123 | 0.221208 | 0.217786 | 0.215492 | 0.296034 | 0.834952 | 1.283951 | 1.565391 | 0.486506 | 0.232182 | 0.215642 | 0.264803 | 0.555323 | 1.073390 | 0.510451 | 0.333551 | 0.233430 | 0.307891 | 0.333074 | 0.294545 | 0.342517 | 0.234354 | 0.229339 | 0.214831 | 0.231983 | 0.272001 | 0.263673 | 0.247361 | 0.225859 | 0.233024 | 0.265716 | 0.279846 | 0.243222 | 0.215461 | 0.214835 | 0.218734 | 0.216380 | 0.215343 | 0.214986 | 0.231774 | 0.295987 | 0.307748 | 0.295953 | 0.239412 | 0.214674 | 0.215205 | 0.217744 | 0.218729 | 0.214922 | 0.223649 | 0.245226 | 0.371274 | 0.300394 | 0.274316 | 0.239458 | 0.217819 | 0.228870 | 0.317350 | 0.989617 | 3.396611 | 12.696556 | 15.939520 | 9.157921 | 5.261803 | 3.907085 | 5.469060 | 5.251754 | 5.765414 | 5.064508 | 6.864800 | 9.546046 | 21.596973 | 52.818073 | 216.814918 | 87.683679 | 61.302200 | 55.920262 | 23.682639 | 8.980301 | 5.238033 | 2.121257 | 1.285438 | 0.774112 | 0.408189 | 0.291606 | 0.290396 | 0.288619 | 0.277567 | 0.409403 | 1.182782 | 6.756257 | 58.983710 | 142.744743 | 315.341730 | 204.644817 | 126.236544 | 69.068867 | 39.494109 | 10.013555 | 5.780588 | 2.855277 | 3.968633 | 4.677244 | 3.826205 | 4.199329 | 3.809391 | 3.248652 | 2.844922 | 3.658505 | 4.158862 | 5.245631 | 4.949813 | 5.727796 | 12.621091 | 18.294205 | 21.498670 | 11.962788 | 6.133726 | 3.506047 | 3.739542 | 2.418062 | 1.633905 | 0.562133 | 0.422421 | 0.311614 | 0.240558 | 0.216046 | 0.226037 | 0.229240 | 0.215220 | 0.263064 | 0.513544 | 1.722984 | 3.537923 | 10.670681 | 17.202258 | 11.300282 | 9.614898 | 4.303068 | 1.892321 | 0.851334 | 0.439611 | 0.389128 | 0.303663 | 0.261234 | 0.277396 | 0.268374 | 0.275284 | 0.292655 | 0.311141 | 0.336552 | 0.367836 | 0.387775 | 0.420397 | 0.565170 | 0.556384 | 0.488691 | 0.426281 | 0.363638 | 0.308945 | 0.303761 | 0.258888 | 0.235962 | 0.220607 | 0.218615 | 0.276776 | 0.328363 | 0.407505 | 0.391043 | 0.398956 | 0.376700 | 0.264181 | 0.219984 | 0.218815 | 0.227452 | 0.255015 | 0.281047 | 0.277561 | 0.300754 | 0.292944 | 0.293351 | 0.250469 | 0.255981 | 0.288990 | 0.366898 | 0.386178 | 0.373990 | 0.299484 | 0.272196 | 0.219997 | 0.224939 | 0.252846 | 0.286369 | 0.279268 | 0.309490 | 0.355893 | 0.379647 | 0.360217 | 0.462064 | 0.614801 | 0.803925 | 0.690169 | 0.704215 | 0.729909 | 0.547211 | 0.443017 | 0.343915 | 0.359669 | 0.350935 | 0.344058 | 0.321431 | 0.315894 | 0.311603 | 0.314516 | 0.293969 | 0.308875 | 0.305184 | 0.300041 | 0.315214 | 0.337681 | 0.365111 | 0.361208 | 0.343011 | 0.324004 | 0.312124 | 0.272370 | 0.230045 | 0.222818 | 0.217561 | 0.216014 | 0.234948 | 0.252466 | 0.262994 | 0.249385 | 0.277906 | 0.298792 | 0.301984 | 0.340997 | 0.404746 | 0.525340 | 0.772296 | 0.900839 | 1.099688 | 1.015291 | 0.694253 | 0.481959 | 0.381955 | 0.332431 | 0.300416 | 0.243982 | 0.224822 | 0.226411 | 0.234047 | 0.242526 | 0.260849 | 0.308615 | 0.366987 | 0.573038 | 0.988150 | 1.280335 | 1.399034 | 1.257671 |
| right central | 2.260404 | 4.017851 | 2.828956 | 5.315430 | 12.608693 | 6.271697 | 7.134813 | 2.915838 | 2.301458 | 4.830671 | 6.011451 | 10.268513 | 30.113436 | 47.230339 | 32.226276 | 10.856677 | 7.121609 | 4.573623 | 1.125510 | 0.390752 | 0.233508 | 0.218632 | 0.217676 | 0.296538 | 0.610184 | 0.356147 | 0.243934 | 0.233546 | 0.388188 | 0.310733 | 0.333192 | 0.265025 | 0.223782 | 0.221213 | 0.237317 | 0.227416 | 0.224640 | 0.226525 | 0.214661 | 0.216474 | 0.226727 | 0.224052 | 0.214639 | 0.214770 | 0.216082 | 0.215461 | 0.216933 | 0.214812 | 0.215843 | 0.223887 | 0.220460 | 0.224401 | 0.215300 | 0.217870 | 0.219546 | 0.227267 | 0.288239 | 0.386725 | 0.569536 | 0.617244 | 0.388561 | 0.252748 | 0.218111 | 0.218898 | 0.262607 | 0.443160 | 0.606748 | 0.569240 | 0.298347 | 0.219649 | 0.217876 | 0.250075 | 0.273589 | 0.263923 | 0.264873 | 0.218273 | 0.230813 | 0.279717 | 0.543066 | 1.155672 | 1.936003 | 4.708702 | 9.018440 | 16.256369 | 47.970854 | 109.694270 | 215.637441 | 178.301131 | 92.472201 | 45.492974 | 23.018143 | 15.542684 | 14.569377 | 10.997587 | 7.876610 | 4.178425 | 2.219628 | 1.707676 | 1.335384 | 0.926618 | 0.740842 | 0.646488 | 1.089425 | 1.730278 | 2.065214 | 2.142991 | 2.647114 | 3.653084 | 4.879288 | 5.794344 | 3.825657 | 5.794743 | 7.818706 | 7.661552 | 12.537168 | 14.963283 | 19.310646 | 104.179402 | 96.660441 | 85.993309 | 85.820674 | 198.680262 | 309.647151 | 203.945521 | 63.841255 | 24.102075 | 15.270437 | 12.632669 | 4.410485 | 1.415167 | 0.955755 | 0.824333 | 0.711489 | 0.639984 | 0.462639 | 0.426269 | 0.499841 | 0.422872 | 0.279432 | 0.228362 | 0.215129 | 0.235039 | 0.283173 | 0.336544 | 0.299542 | 0.297698 | 0.270359 | 0.263933 | 0.240292 | 0.220340 | 0.214916 | 0.218357 | 0.252120 | 0.293438 | 0.367617 | 0.351119 | 0.330245 | 0.287870 | 0.258257 | 0.222257 | 0.216974 | 0.264880 | 0.403753 | 0.518168 | 0.896689 | 0.961785 | 0.696268 | 0.391800 | 0.262556 | 0.215559 | 0.222717 | 0.297677 | 0.411845 | 0.517652 | 0.495654 | 0.423444 | 0.367981 | 0.344886 | 0.280194 | 0.247593 | 0.230617 | 0.230033 | 0.251307 | 0.274380 | 0.288970 | 0.315836 | 0.336591 | 0.297050 | 0.245443 | 0.218540 | 0.217334 | 0.222267 | 0.247404 | 0.309070 | 0.424355 | 0.491377 | 0.408898 | 0.354680 | 0.252337 | 0.216039 | 0.228718 | 0.227305 | 0.218720 | 0.215606 | 0.220877 | 0.220347 | 0.220096 | 0.215833 | 0.215348 | 0.217594 | 0.216246 | 0.216384 | 0.215397 | 0.215076 | 0.221119 | 0.220548 | 0.218976 | 0.216947 | 0.216897 | 0.215591 | 0.215329 | 0.216325 | 0.220416 | 0.255133 | 0.292940 | 0.284245 | 0.263607 | 0.239035 | 0.218007 | 0.304108 | 0.626533 | 1.116916 | 1.274173 | 1.352785 | 0.984320 | 0.448414 | 0.294459 | 0.222148 | 0.219867 | 0.229980 | 0.232244 | 0.218015 | 0.217150 | 0.214897 | 0.215845 | 0.222822 | 0.223763 | 0.236561 | 0.262182 | 0.312296 | 0.365464 | 0.375032 | 0.383206 | 0.468816 | 0.435007 | 0.348678 | 0.360765 | 0.351720 | 0.465057 | 0.655808 | 0.561418 | 0.325445 | 0.286409 | 0.239122 | 0.252061 | 0.237814 | 0.222429 | 0.216557 | 0.249254 | 0.324700 | 0.476342 | 0.595995 | 0.960375 | 1.353837 | 1.420907 | 1.178217 | 0.774148 | 0.549901 | 0.438920 | 0.344765 |
| left posterior | 3.404823 | 2.065821 | 2.040336 | 1.996443 | 0.872257 | 0.576653 | 0.370980 | 0.284457 | 0.266495 | 0.241260 | 0.215597 | 0.277100 | 0.608360 | 0.566875 | 0.458427 | 0.267086 | 0.215425 | 0.214650 | 0.214626 | 0.215365 | 0.221299 | 0.227658 | 0.496151 | 2.279725 | 4.081081 | 2.904369 | 1.313072 | 0.716466 | 0.389854 | 0.252242 | 0.228170 | 0.249349 | 0.279789 | 0.328238 | 0.446414 | 0.828537 | 1.009084 | 0.881261 | 0.373720 | 0.214879 | 0.298551 | 0.450202 | 0.885040 | 1.518409 | 3.538240 | 2.957368 | 3.532364 | 1.480060 | 0.929713 | 0.587347 | 0.483013 | 0.508898 | 0.418613 | 0.408648 | 0.592902 | 0.472981 | 0.408124 | 0.273699 | 0.219658 | 0.220849 | 0.214790 | 0.236196 | 0.246779 | 0.258170 | 0.262932 | 0.285889 | 0.407348 | 0.621307 | 0.724489 | 0.761258 | 0.660604 | 0.520271 | 0.376634 | 0.253262 | 0.227395 | 0.238613 | 0.265351 | 0.348340 | 0.575152 | 1.556520 | 5.326442 | 5.493119 | 3.581035 | 2.857042 | 3.161779 | 3.356863 | 1.750986 | 1.069284 | 0.743689 | 0.594414 | 0.477535 | 0.371793 | 0.292800 | 0.296352 | 0.291412 | 0.348519 | 0.494752 | 0.668176 | 0.703146 | 0.705493 | 0.517710 | 0.385720 | 0.275656 | 0.233156 | 0.216238 | 0.218936 | 0.248490 | 0.250888 | 0.243371 | 0.216707 | 0.216903 | 0.241190 | 0.338678 | 0.482793 | 0.776489 | 1.300993 | 1.677677 | 1.950126 | 2.345042 | 1.861509 | 1.891751 | 1.297038 | 0.637539 | 0.418572 | 0.291584 | 0.228210 | 0.237573 | 0.248412 | 0.258627 | 0.266280 | 0.264495 | 0.268455 | 0.275376 | 0.243234 | 0.257441 | 0.276385 | 0.309989 | 0.317031 | 0.320537 | 0.291146 | 0.251378 | 0.224439 | 0.214668 | 0.217620 | 0.221556 | 0.241756 | 0.254174 | 0.236984 | 0.237172 | 0.227023 | 0.230884 | 0.252232 | 0.251889 | 0.251577 | 0.276041 | 0.292830 | 0.295817 | 0.289144 | 0.271119 | 0.276087 | 0.291359 | 0.292128 | 0.326302 | 0.410511 | 0.497001 | 0.572472 | 0.700549 | 0.542678 | 0.360312 | 0.290788 | 0.262218 | 0.250606 | 0.242472 | 0.226067 | 0.216507 | 0.218006 | 0.217749 | 0.223207 | 0.225058 | 0.228245 | 0.258015 | 0.414543 | 0.627173 | 1.167362 | 1.258281 | 1.303263 | 1.590898 | 1.286512 | 0.878097 | 0.876777 | 1.072876 | 1.054653 | 1.399664 | 2.223371 | 4.262987 | 6.353421 | 4.833114 | 2.808106 | 3.839572 | 3.872798 | 3.906378 | 2.765078 | 2.394244 | 2.077345 | 1.514029 | 1.286785 | 1.094086 | 0.777823 | 0.704209 | 0.608259 | 0.580675 | 0.723787 | 0.798842 | 0.775502 | 0.565115 | 0.383940 | 0.289751 | 0.245361 | 0.221359 | 0.214782 | 0.217766 | 0.218622 | 0.215408 | 0.218697 | 0.221481 | 0.222232 | 0.222599 | 0.216830 | 0.215987 | 0.214862 | 0.219295 | 0.231175 | 0.238705 | 0.239004 | 0.230744 | 0.226020 | 0.225259 | 0.220317 | 0.214649 | 0.216387 | 0.218617 | 0.219839 | 0.221638 | 0.234479 | 0.258662 | 0.275634 | 0.330447 | 0.517615 | 0.638856 | 0.647603 | 0.518958 | 0.443268 | 0.400894 | 0.388416 | 0.342257 | 0.403164 | 0.435419 | 0.493906 | 0.536307 | 0.635682 | 0.563375 | 0.538359 | 0.410939 | 0.398105 | 0.355102 | 0.364514 | 0.413923 | 0.490058 | 0.486202 | 0.546348 | 0.759339 | 2.059592 | 7.497794 | 17.864180 | 32.336479 | 90.527246 | 246.292184 | 289.188773 | 255.616166 | 167.956910 |
| right posterior | 5.367903 | 1.806557 | 1.397761 | 1.355958 | 1.415938 | 0.809186 | 0.657817 | 0.343839 | 0.373727 | 0.374780 | 0.487575 | 0.400215 | 0.246101 | 0.253144 | 0.263984 | 0.233830 | 0.224973 | 0.268272 | 0.347724 | 0.345621 | 0.248796 | 0.223007 | 0.263323 | 0.288481 | 0.263371 | 0.230525 | 0.221155 | 0.215630 | 0.220684 | 0.217530 | 0.222247 | 0.232206 | 0.227430 | 0.237105 | 0.263425 | 0.219570 | 0.218680 | 0.230264 | 0.226829 | 0.215332 | 0.226780 | 0.264050 | 0.254967 | 0.261271 | 0.380330 | 0.426354 | 0.283152 | 0.223287 | 0.216096 | 0.219878 | 0.226736 | 0.246774 | 0.265992 | 0.254652 | 0.219292 | 0.216499 | 0.221286 | 0.219393 | 0.221978 | 0.229761 | 0.253718 | 0.224865 | 0.219326 | 0.227959 | 0.252230 | 0.276055 | 0.322221 | 0.350067 | 0.372298 | 0.320531 | 0.256150 | 0.232783 | 0.223121 | 0.214635 | 0.216460 | 0.216341 | 0.216845 | 0.233857 | 0.280157 | 0.472712 | 1.678607 | 4.407377 | 15.713474 | 98.592228 | 224.059578 | 166.073653 | 310.193277 | 183.791459 | 240.916552 | 537.756676 | 2183.655761 | 6745.413608 | 4495.920026 | 1787.935001 | 434.770458 | 133.417407 | 29.977234 | 9.507732 | 5.672747 | 2.011037 | 0.581612 | 0.377221 | 0.368122 | 0.410902 | 0.523444 | 0.748418 | 1.268565 | 2.109559 | 2.537013 | 2.671941 | 3.828130 | 5.489415 | 6.183561 | 5.068255 | 5.659033 | 11.227186 | 17.108711 | 22.225879 | 23.033109 | 23.311882 | 19.806437 | 11.005961 | 3.944874 | 1.620345 | 0.745352 | 0.466842 | 0.380935 | 0.284397 | 0.313381 | 0.372962 | 0.464679 | 0.536926 | 0.612901 | 0.673085 | 0.765006 | 0.660788 | 0.557279 | 0.482855 | 0.398318 | 0.313078 | 0.263087 | 0.240671 | 0.222274 | 0.214774 | 0.224201 | 0.222488 | 0.215839 | 0.222347 | 0.257853 | 0.347000 | 0.457754 | 0.560892 | 0.770748 | 1.169782 | 1.304434 | 1.355870 | 1.047411 | 1.668899 | 3.009006 | 2.550167 | 1.414773 | 0.928685 | 0.784337 | 0.810317 | 0.582466 | 0.517479 | 0.711682 | 1.686760 | 3.043704 | 3.083948 | 2.324956 | 2.282208 | 2.248523 | 2.563301 | 1.965800 | 2.146440 | 3.349607 | 7.600681 | 15.139814 | 12.698172 | 6.151791 | 2.940650 | 1.238094 | 0.658842 | 0.332579 | 0.252068 | 0.222419 | 0.215214 | 0.220923 | 0.220482 | 0.222336 | 0.216272 | 0.216442 | 0.214755 | 0.220014 | 0.242187 | 0.281438 | 0.311946 | 0.424427 | 0.708218 | 0.996682 | 1.180354 | 0.944957 | 0.579319 | 0.501971 | 0.414110 | 0.295250 | 0.235510 | 0.215977 | 0.215415 | 0.216913 | 0.226064 | 0.249821 | 0.246139 | 0.230666 | 0.214679 | 0.270190 | 0.366843 | 0.512609 | 0.751441 | 0.712931 | 0.559915 | 0.423800 | 0.272646 | 0.246359 | 0.233876 | 0.227406 | 0.253852 | 0.303642 | 0.365844 | 0.483195 | 0.496332 | 0.514751 | 0.581197 | 0.572420 | 0.658731 | 0.734879 | 0.691763 | 0.830082 | 0.917735 | 0.601544 | 0.369136 | 0.256498 | 0.217798 | 0.215184 | 0.229088 | 0.239071 | 0.231046 | 0.223754 | 0.220900 | 0.214637 | 0.217260 | 0.225653 | 0.227959 | 0.225422 | 0.220166 | 0.216359 | 0.216058 | 0.234974 | 0.251044 | 0.278863 | 0.301472 | 0.312178 | 0.324085 | 0.292772 | 0.259792 | 0.248251 | 0.220807 | 0.217081 | 0.233001 | 0.242082 | 0.246409 | 0.251006 | 0.237752 | 0.214626 | 0.250142 | 0.422044 | 0.448274 | 0.568395 | 1.120785 |
| all electrodes | 0.912496 | 1.131293 | 1.961515 | 10.621556 | 45.639460 | 27.523664 | 12.499969 | 7.849056 | 2.782822 | 0.687988 | 0.255683 | 0.222804 | 0.242006 | 0.248825 | 0.239858 | 0.215506 | 0.249440 | 0.282724 | 0.322433 | 0.529805 | 0.630638 | 0.279061 | 0.214646 | 0.323481 | 0.497951 | 0.775472 | 1.192234 | 1.230557 | 0.496914 | 0.332711 | 0.240856 | 0.219665 | 0.214761 | 0.214626 | 0.220766 | 0.221769 | 0.214999 | 0.214643 | 0.214768 | 0.215467 | 0.220965 | 0.227901 | 0.236984 | 0.274819 | 0.308258 | 0.262818 | 0.255764 | 0.244944 | 0.218015 | 0.216655 | 0.214626 | 0.216087 | 0.214781 | 0.215296 | 0.214897 | 0.214731 | 0.224267 | 0.259796 | 0.314505 | 0.277425 | 0.296499 | 0.216083 | 0.220082 | 0.249541 | 0.313114 | 0.315514 | 0.285396 | 0.281121 | 0.222503 | 0.215559 | 0.248074 | 0.331876 | 0.357206 | 0.347869 | 0.276737 | 0.222736 | 0.217476 | 0.229907 | 0.357038 | 0.685695 | 1.306468 | 1.278093 | 0.822488 | 1.106890 | 1.324160 | 1.543306 | 1.354872 | 0.943154 | 0.822253 | 1.023643 | 1.162700 | 2.421455 | 2.575250 | 3.094155 | 2.249628 | 1.209554 | 0.900355 | 0.681609 | 0.568805 | 0.554594 | 0.587047 | 0.655456 | 0.880848 | 1.016906 | 1.093491 | 0.886899 | 0.805143 | 0.651345 | 0.491109 | 0.416507 | 0.391317 | 0.408694 | 0.389178 | 0.413121 | 0.424557 | 0.439576 | 0.449786 | 0.535396 | 0.428722 | 0.465622 | 0.504655 | 0.494075 | 0.363589 | 0.304666 | 0.264696 | 0.273744 | 0.292355 | 0.333875 | 0.423562 | 0.655425 | 0.818037 | 0.839568 | 0.742140 | 0.576388 | 0.372583 | 0.290721 | 0.273598 | 0.237363 | 0.222513 | 0.248963 | 0.256476 | 0.251209 | 0.251773 | 0.256573 | 0.234119 | 0.215456 | 0.217320 | 0.220315 | 0.249959 | 0.292162 | 0.316030 | 0.336519 | 0.415891 | 0.555708 | 0.958435 | 1.303126 | 1.882032 | 2.493708 | 3.486926 | 3.006231 | 1.985882 | 0.984548 | 0.658992 | 0.494265 | 0.394873 | 0.306595 | 0.339152 | 0.479884 | 0.715819 | 0.959389 | 1.236801 | 1.585558 | 1.337237 | 1.110402 | 0.749932 | 0.566268 | 0.591453 | 0.591707 | 0.406576 | 0.397322 | 0.292547 | 0.256773 | 0.235491 | 0.214686 | 0.222125 | 0.222084 | 0.224751 | 0.228773 | 0.226518 | 0.223978 | 0.215802 | 0.214805 | 0.222592 | 0.220639 | 0.216960 | 0.223663 | 0.240603 | 0.269062 | 0.293057 | 0.249942 | 0.241210 | 0.253629 | 0.259960 | 0.255890 | 0.243969 | 0.258917 | 0.375148 | 0.736216 | 0.765605 | 0.686177 | 0.722149 | 0.947882 | 0.706633 | 0.462519 | 0.281006 | 0.230598 | 0.215487 | 0.216749 | 0.220620 | 0.220983 | 0.225544 | 0.220198 | 0.216304 | 0.254572 | 0.340049 | 0.379622 | 0.374938 | 0.376143 | 0.347448 | 0.278359 | 0.231138 | 0.214654 | 0.216146 | 0.240786 | 0.309737 | 0.362819 | 0.377344 | 0.408398 | 0.374242 | 0.590781 | 0.802285 | 1.067094 | 1.118552 | 0.864862 | 0.736307 | 0.818685 | 0.624484 | 0.235572 | 0.217450 | 0.223023 | 0.218602 | 0.220358 | 0.225422 | 0.220019 | 0.217046 | 0.224482 | 0.218670 | 0.216524 | 0.214761 | 0.214701 | 0.214635 | 0.214704 | 0.218300 | 0.218086 | 0.217646 | 0.215971 | 0.224390 | 0.231866 | 0.237306 | 0.250332 | 0.259371 | 0.302659 | 0.369644 | 0.477355 | 0.700601 | 1.174933 | 2.170968 | 3.094263 | 4.320070 | 3.881237 |

Searchlight, spatiotemporal cluster permutation test

|  | start time | stop time | peak time | peak channel | cluster p | peak Cohen's d | direction |
| --- | --- | --- | --- | --- | --- | --- | --- |
| #1 | 140 | 500 | 260 | CP2 | 0.0002 | 1.146939 | positive |
| #2 | -200 | -80 | -120 | CP1 | 0.0278 | -0.94798 | negative |

M) real faces (LOSO) - neutral vs angry

  
|  | time window | peak latency | cluster *p* | peak Cohen's *d* |  | | | |
| **all electrodes** | 100 - 665 ms | 160 ms | 0.0001 | 2.106 |  | | | |
|  | | | | | | | | |

Time-resolved classification, cluster permutation tests

|  | **left hemisphere** | | | | **right hemisphere** | | | |
|  | time window | peak latency | cluster *p* | peak Cohen's *d* | time window | peak latency | cluster *p* | peak Cohen's *d* |
| **anterior** |  | | | |  | | | |
| **central** | 240 - 310 ms | 280 ms | 0.0379 | 1.0941 | 105 - 465 ms | 285 ms | 0.0001 | 1.5973 |
| **posterior** | 110 - 740 ms | 155 ms | 0.0001 | 1.7768 | 80 - 1195 ms | 150 ms | 0.0001 | 1.7856 |
 860 - 1070 ms | 900 ms | 0.0158 | 0.5718 |  | | | |

  

Time-resolved classification, Bayesian statistics

|  | -200 | -195 | -190 | -185 | -180 | -175 | -170 | -165 | -160 | -155 | -150 | -145 | -140 | -135 | -130 | -125 | -120 | -115 | -110 | -105 | -100 | -95 | -90 | -85 | -80 | -75 | -70 | -65 | -60 | -55 | -50 | -45 | -40 | -35 | -30 | -25 | -20 | -15 | -10 | -5 | 0 | 5 | 10 | 15 | 20 | 25 | 30 | 35 | 40 | 45 | 50 | 55 | 60 | 65 | 70 | 75 | 80 | 85 | 90 | 95 | 100 | 105 | 110 | 115 | 120 | 125 | 130 | 135 | 140 | 145 | 150 | 155 | 160 | 165 | 170 | 175 | 180 | 185 | 190 | 195 | 200 | 205 | 210 | 215 | 220 | 225 | 230 | 235 | 240 | 245 | 250 | 255 | 260 | 265 | 270 | 275 | 280 | 285 | 290 | 295 | 300 | 305 | 310 | 315 | 320 | 325 | 330 | 335 | 340 | 345 | 350 | 355 | 360 | 365 | 370 | 375 | 380 | 385 | 390 | 395 | 400 | 405 | 410 | 415 | 420 | 425 | 430 | 435 | 440 | 445 | 450 | 455 | 460 | 465 | 470 | 475 | 480 | 485 | 490 | 495 | 500 | 505 | 510 | 515 | 520 | 525 | 530 | 535 | 540 | 545 | 550 | 555 | 560 | 565 | 570 | 575 | 580 | 585 | 590 | 595 | 600 | 605 | 610 | 615 | 620 | 625 | 630 | 635 | 640 | 645 | 650 | 655 | 660 | 665 | 670 | 675 | 680 | 685 | 690 | 695 | 700 | 705 | 710 | 715 | 720 | 725 | 730 | 735 | 740 | 745 | 750 | 755 | 760 | 765 | 770 | 775 | 780 | 785 | 790 | 795 | 800 | 805 | 810 | 815 | 820 | 825 | 830 | 835 | 840 | 845 | 850 | 855 | 860 | 865 | 870 | 875 | 880 | 885 | 890 | 895 | 900 | 905 | 910 | 915 | 920 | 925 | 930 | 935 | 940 | 945 | 950 | 955 | 960 | 965 | 970 | 975 | 980 | 985 | 990 | 995 | 1000 | 1005 | 1010 | 1015 | 1020 | 1025 | 1030 | 1035 | 1040 | 1045 | 1050 | 1055 | 1060 | 1065 | 1070 | 1075 | 1080 | 1085 | 1090 | 1095 | 1100 | 1105 | 1110 | 1115 | 1120 | 1125 | 1130 | 1135 | 1140 | 1145 | 1150 | 1155 | 1160 | 1165 | 1170 | 1175 | 1180 | 1185 | 1190 | 1195 |
| --- | --- | --- | --- | --- | --- | --- | --- | --- | --- | --- | --- | --- | --- | --- | --- | --- | --- | --- | --- | --- | --- | --- | --- | --- | --- | --- | --- | --- | --- | --- | --- | --- | --- | --- | --- | --- | --- | --- | --- | --- | --- | --- | --- | --- | --- | --- | --- | --- | --- | --- | --- | --- | --- | --- | --- | --- | --- | --- | --- | --- | --- | --- | --- | --- | --- | --- | --- | --- | --- | --- | --- | --- | --- | --- | --- | --- | --- | --- | --- | --- | --- | --- | --- | --- | --- | --- | --- | --- | --- | --- | --- | --- | --- | --- | --- | --- | --- | --- | --- | --- | --- | --- | --- | --- | --- | --- | --- | --- | --- | --- | --- | --- | --- | --- | --- | --- | --- | --- | --- | --- | --- | --- | --- | --- | --- | --- | --- | --- | --- | --- | --- | --- | --- | --- | --- | --- | --- | --- | --- | --- | --- | --- | --- | --- | --- | --- | --- | --- | --- | --- | --- | --- | --- | --- | --- | --- | --- | --- | --- | --- | --- | --- | --- | --- | --- | --- | --- | --- | --- | --- | --- | --- | --- | --- | --- | --- | --- | --- | --- | --- | --- | --- | --- | --- | --- | --- | --- | --- | --- | --- | --- | --- | --- | --- | --- | --- | --- | --- | --- | --- | --- | --- | --- | --- | --- | --- | --- | --- | --- | --- | --- | --- | --- | --- | --- | --- | --- | --- | --- | --- | --- | --- | --- | --- | --- | --- | --- | --- | --- | --- | --- | --- | --- | --- | --- | --- | --- | --- | --- | --- | --- | --- | --- | --- | --- | --- | --- | --- | --- | --- | --- | --- | --- | --- | --- | --- | --- | --- | --- | --- | --- | --- | --- | --- | --- | --- | --- | --- | --- | --- | --- | --- | --- | --- | --- | --- | --- | --- | --- | --- |
| left anterior | 0.255454 | 0.217079 | 0.218182 | 0.218262 | 0.218584 | 0.227324 | 0.215008 | 0.231912 | 0.318261 | 0.303188 | 0.265482 | 0.226003 | 0.214949 | 0.227135 | 0.217784 | 0.234196 | 0.267055 | 0.417045 | 0.512183 | 0.623495 | 0.583249 | 0.489244 | 0.744822 | 0.477345 | 0.400015 | 0.246296 | 0.218659 | 0.226232 | 0.329458 | 0.594459 | 0.967911 | 0.729970 | 0.235720 | 0.238789 | 0.243120 | 0.355682 | 0.564143 | 0.353176 | 0.249968 | 0.214686 | 0.215279 | 0.249370 | 0.276557 | 0.265815 | 0.885375 | 1.837772 | 4.582637 | 10.823001 | 3.060657 | 1.304322 | 0.916306 | 0.322283 | 0.228408 | 0.214859 | 0.243600 | 0.392101 | 0.573462 | 0.897926 | 0.977681 | 0.613238 | 0.407961 | 0.258043 | 0.215849 | 0.214672 | 0.218259 | 0.226274 | 0.330644 | 0.596455 | 1.804118 | 4.409456 | 5.066746e+00 | 2.526319e+00 | 1.284265e+00 | 4.345392e-01 | 2.710691e-01 | 2.153509e-01 | 2.909308e-01 | 0.596962 | 0.930338 | 1.603062 | 1.904747 | 1.803483 | 1.080982 | 0.929044 | 0.597451 | 0.414767 | 0.309332 | 0.224823 | 0.226094 | 0.352544 | 0.797494 | 1.920047 | 4.093204 | 7.662892 | 10.573414 | 8.111573 | 3.597022 | 2.548979 | 1.704488 | 1.129643 | 0.662035 | 0.441150 | 0.347600 | 0.282419 | 0.228795 | 0.220700 | 0.221163 | 0.221622 | 0.217376 | 0.216945 | 0.216760 | 0.226483 | 0.272548 | 0.337854 | 0.492321 | 0.626550 | 0.900663 | 1.769179 | 2.239752 | 2.380178 | 2.739096 | 5.652530 | 10.294981 | 16.558701 | 36.086990 | 49.026660 | 68.275407 | 45.153454 | 14.889843 | 6.846248 | 2.709578 | 1.014495 | 0.573553 | 0.374179 | 0.314931 | 0.273842 | 0.259276 | 0.248170 | 0.271865 | 0.290238 | 0.318156 | 0.503107 | 0.567744 | 0.551614 | 0.627840 | 0.514896 | 0.490803 | 0.416204 | 0.275107 | 0.246229 | 0.224691 | 0.217276 | 0.218762 | 0.224716 | 0.230414 | 0.230492 | 0.221014 | 0.214816 | 0.243290 | 0.320546 | 0.472525 | 0.806257 | 1.029968 | 1.302569 | 1.083199 | 0.864171 | 0.658709 | 0.673673 | 0.527481 | 0.395771 | 0.280507 | 0.255142 | 0.234274 | 0.221512 | 0.214762 | 0.214635 | 0.217168 | 0.240457 | 0.254342 | 0.307795 | 0.340173 | 0.408992 | 0.453171 | 0.535115 | 0.533376 | 0.512598 | 0.373957 | 0.349671 | 0.321753 | 0.380689 | 0.401034 | 0.366307 | 0.373361 | 0.467563 | 0.449256 | 0.418273 | 0.286479 | 0.242240 | 0.247745 | 0.265037 | 0.246984 | 0.237467 | 0.226978 | 0.220104 | 0.215834 | 0.214687 | 0.215365 | 0.214698 | 0.216671 | 0.219976 | 0.215019 | 0.214998 | 0.215672 | 0.214739 | 0.214690 | 0.219894 | 0.238716 | 0.234198 | 0.230286 | 0.234243 | 0.222474 | 0.218299 | 0.215123 | 0.217953 | 0.233070 | 0.234644 | 0.220652 | 0.214696 | 0.214917 | 0.215131 | 0.216392 | 0.223915 | 0.240003 | 0.247224 | 0.239835 | 0.235745 | 0.217826 | 0.214819 | 0.221732 | 0.282862 | 0.496362 | 0.751700 | 0.731709 | 0.657944 | 0.670555 | 0.549242 | 0.358322 | 0.288908 | 0.267862 | 0.307872 | 0.328612 | 0.324351 | 0.333324 | 0.415925 | 0.428904 | 0.563452 | 0.551190 | 0.435596 | 0.478628 | 0.587314 | 0.739139 | 0.935729 | 0.741598 | 0.591877 | 0.665226 | 0.592314 | 0.492399 | 0.450464 | 0.538635 | 0.761375 | 0.866563 | 0.640423 | 0.730154 | 0.841241 | 0.900640 | 0.776127 | 0.998017 | 1.108858 | 2.980148 | 3.273234 |
| right anterior | 0.268811 | 0.405814 | 0.283100 | 0.240153 | 0.215737 | 0.236256 | 0.245229 | 0.253524 | 0.327830 | 0.290454 | 0.236682 | 0.251634 | 0.302469 | 0.305882 | 0.252012 | 0.254838 | 0.329982 | 0.434268 | 0.312423 | 0.252635 | 0.242071 | 0.267512 | 0.231745 | 0.217402 | 0.214989 | 0.238774 | 0.232046 | 0.240631 | 0.233434 | 0.236143 | 0.242305 | 0.237467 | 0.236365 | 0.334415 | 0.510807 | 0.960432 | 1.307898 | 1.028519 | 0.399594 | 0.297833 | 0.260684 | 0.231354 | 0.224826 | 0.246608 | 0.446090 | 1.261710 | 1.824302 | 1.087075 | 0.550534 | 0.345119 | 0.278783 | 0.225329 | 0.219335 | 0.218830 | 0.216990 | 0.220013 | 0.214728 | 0.217975 | 0.214750 | 0.215126 | 0.214641 | 0.215598 | 0.214689 | 0.214641 | 0.215743 | 0.273584 | 0.520017 | 1.596796 | 5.701872 | 20.802488 | 2.227218e+01 | 2.422820e+01 | 2.449620e+01 | 1.207870e+01 | 7.078966e+00 | 2.966598e+00 | 1.597558e+00 | 1.279218 | 0.734957 | 0.404131 | 0.287798 | 0.280871 | 0.279160 | 0.265471 | 0.278616 | 0.282543 | 0.291751 | 0.317745 | 0.419193 | 0.569592 | 0.940229 | 1.393705 | 2.552037 | 4.224877 | 8.176006 | 10.944306 | 22.412261 | 16.740198 | 9.106510 | 5.522178 | 5.143339 | 3.642378 | 1.183443 | 0.601422 | 0.573741 | 0.635286 | 0.591941 | 0.368539 | 0.241424 | 0.237468 | 0.261726 | 0.270813 | 0.297185 | 0.267213 | 0.332104 | 0.568275 | 1.292589 | 1.672974 | 1.986459 | 3.181568 | 7.428148 | 16.115745 | 43.493004 | 47.285049 | 33.404806 | 29.993232 | 14.634476 | 6.579706 | 5.011565 | 3.986475 | 4.172854 | 4.270974 | 1.958828 | 0.763546 | 0.385399 | 0.266712 | 0.215745 | 0.233469 | 0.303156 | 0.348520 | 0.295321 | 0.233510 | 0.227410 | 0.235982 | 0.261641 | 0.282794 | 0.275049 | 0.271530 | 0.294918 | 0.274209 | 0.243513 | 0.235440 | 0.243714 | 0.314966 | 0.512066 | 1.020320 | 2.441082 | 4.709524 | 2.640049 | 1.112957 | 0.551891 | 0.363269 | 0.240354 | 0.214736 | 0.223430 | 0.240366 | 0.246118 | 0.276667 | 0.312499 | 0.291696 | 0.265542 | 0.243618 | 0.219595 | 0.214829 | 0.216642 | 0.221996 | 0.233294 | 0.240555 | 0.254404 | 0.247080 | 0.238484 | 0.237056 | 0.237934 | 0.234814 | 0.239574 | 0.228737 | 0.226688 | 0.222259 | 0.217812 | 0.217725 | 0.220709 | 0.219030 | 0.215729 | 0.214681 | 0.218114 | 0.221448 | 0.230493 | 0.231487 | 0.228098 | 0.230460 | 0.229923 | 0.222740 | 0.217674 | 0.214633 | 0.214652 | 0.214724 | 0.221050 | 0.249082 | 0.261399 | 0.249214 | 0.227159 | 0.216001 | 0.214631 | 0.216560 | 0.220536 | 0.217631 | 0.214701 | 0.215800 | 0.231953 | 0.268190 | 0.315511 | 0.339694 | 0.280352 | 0.258071 | 0.240252 | 0.224947 | 0.225513 | 0.221074 | 0.214744 | 0.214626 | 0.218616 | 0.218978 | 0.222503 | 0.227677 | 0.223046 | 0.214659 | 0.223871 | 0.233241 | 0.232290 | 0.234384 | 0.225892 | 0.214999 | 0.219125 | 0.234244 | 0.229599 | 0.234108 | 0.243280 | 0.248789 | 0.238064 | 0.225095 | 0.221711 | 0.223295 | 0.220370 | 0.218721 | 0.222054 | 0.231956 | 0.258118 | 0.279234 | 0.288681 | 0.287818 | 0.286929 | 0.301213 | 0.337016 | 0.324866 | 0.328463 | 0.294362 | 0.267094 | 0.230793 | 0.216925 | 0.216996 | 0.218823 | 0.231681 | 0.242518 | 0.263605 | 0.248946 | 0.242428 | 0.233643 | 0.252335 | 0.273241 | 0.295794 |
| left central | 0.281898 | 0.353672 | 0.305579 | 0.268737 | 0.339814 | 0.438547 | 0.732228 | 0.496545 | 0.388050 | 0.627678 | 0.789522 | 0.383773 | 0.241919 | 0.218714 | 0.217014 | 0.299753 | 0.879720 | 4.337636 | 4.014514 | 1.498483 | 0.746288 | 0.464678 | 0.237563 | 0.235419 | 0.364474 | 0.427061 | 0.430428 | 0.438177 | 0.402579 | 0.314696 | 0.216154 | 0.223035 | 0.232514 | 0.215042 | 0.218879 | 0.238152 | 0.280898 | 0.401110 | 0.997744 | 3.993762 | 7.497592 | 6.739417 | 4.669469 | 3.258431 | 6.033209 | 5.726258 | 2.343889 | 0.624272 | 0.301940 | 0.259029 | 0.245026 | 0.228764 | 0.232987 | 0.260880 | 0.334781 | 0.577425 | 0.918365 | 1.418317 | 1.902288 | 0.891613 | 0.372940 | 0.299970 | 0.299927 | 0.292576 | 0.311958 | 0.478120 | 0.934216 | 3.213884 | 11.668009 | 23.575847 | 7.650001e+01 | 1.268161e+02 | 1.407781e+02 | 1.699922e+02 | 6.501168e+01 | 2.130945e+01 | 6.901023e+00 | 3.126679 | 2.247291 | 1.004557 | 0.548924 | 0.437192 | 0.355256 | 0.350173 | 0.348290 | 0.326146 | 0.428615 | 0.714665 | 1.705606 | 4.460417 | 16.408738 | 46.739237 | 112.960925 | 385.732527 | 466.122527 | 577.765659 | 1200.554334 | 798.112119 | 615.439646 | 164.125354 | 38.183206 | 15.209090 | 4.096925 | 0.861429 | 0.370943 | 0.230838 | 0.214667 | 0.221742 | 0.221647 | 0.225557 | 0.218510 | 0.214844 | 0.214853 | 0.222185 | 0.235499 | 0.234179 | 0.239265 | 0.232884 | 0.246678 | 0.306570 | 0.352531 | 0.439181 | 0.591677 | 0.657854 | 0.902853 | 0.859558 | 0.656018 | 0.507979 | 0.480783 | 0.427767 | 0.330026 | 0.239609 | 0.226138 | 0.214626 | 0.221194 | 0.235036 | 0.242903 | 0.219133 | 0.227257 | 0.260099 | 0.354659 | 0.478922 | 0.383012 | 0.281375 | 0.225328 | 0.214676 | 0.224686 | 0.226748 | 0.229877 | 0.214634 | 0.228065 | 0.316685 | 0.371966 | 0.396782 | 0.338462 | 0.293430 | 0.243093 | 0.224093 | 0.214652 | 0.216631 | 0.217755 | 0.222561 | 0.226997 | 0.230273 | 0.223222 | 0.218365 | 0.216077 | 0.214897 | 0.215448 | 0.219627 | 0.219146 | 0.231691 | 0.224179 | 0.221162 | 0.216989 | 0.215425 | 0.238220 | 0.257002 | 0.321408 | 0.267452 | 0.251983 | 0.244684 | 0.236678 | 0.228236 | 0.231244 | 0.223031 | 0.236225 | 0.241815 | 0.226943 | 0.218646 | 0.215802 | 0.214912 | 0.216191 | 0.223894 | 0.233592 | 0.219802 | 0.214707 | 0.216211 | 0.223453 | 0.227675 | 0.226952 | 0.216573 | 0.224428 | 0.276785 | 0.323950 | 0.405879 | 0.485671 | 0.511314 | 0.483571 | 0.338361 | 0.298062 | 0.285748 | 0.248348 | 0.223936 | 0.214718 | 0.225721 | 0.230708 | 0.254822 | 0.236335 | 0.214904 | 0.232565 | 0.290749 | 0.408428 | 0.531448 | 0.813974 | 1.038908 | 1.605716 | 2.599056 | 3.625188 | 5.602719 | 7.082975 | 4.922767 | 2.688243 | 2.306790 | 1.493123 | 1.293503 | 1.108599 | 0.892194 | 0.594923 | 0.394222 | 0.248854 | 0.245623 | 0.234963 | 0.226190 | 0.231862 | 0.247694 | 0.249209 | 0.285078 | 0.342893 | 0.432028 | 0.544212 | 0.723313 | 1.118010 | 1.429470 | 0.997319 | 0.653966 | 0.516821 | 0.499843 | 0.418677 | 0.394565 | 0.473069 | 0.682917 | 0.815384 | 1.402085 | 1.684038 | 1.867168 | 1.688204 | 1.389955 | 1.309812 | 1.621072 | 1.634442 | 1.678524 | 1.396112 | 1.152783 | 0.973112 | 0.954909 | 0.853206 | 0.706375 | 0.575514 | 0.529673 |
| right central | 0.224454 | 0.224354 | 0.244150 | 0.283165 | 0.357065 | 0.444247 | 0.625814 | 1.097812 | 1.908188 | 2.360155 | 1.742563 | 0.941776 | 0.700287 | 0.458761 | 0.286589 | 0.224048 | 0.270621 | 0.268598 | 0.224867 | 0.215914 | 0.227066 | 0.234860 | 0.217994 | 0.215088 | 0.223885 | 0.257058 | 0.348412 | 0.766566 | 2.505410 | 1.809337 | 0.522188 | 0.253511 | 0.220468 | 0.224759 | 0.297572 | 0.429137 | 0.331191 | 0.242426 | 0.214640 | 0.214640 | 0.215233 | 0.223711 | 0.249109 | 0.372850 | 0.585222 | 0.949543 | 0.838898 | 0.502185 | 0.366900 | 0.289795 | 0.288047 | 0.255649 | 0.217127 | 0.214665 | 0.216350 | 0.219898 | 0.233792 | 0.255733 | 0.294300 | 0.427479 | 1.002069 | 2.513927 | 8.523328 | 12.663167 | 17.915739 | 39.372934 | 113.493753 | 213.036860 | 891.258981 | 2203.571882 | 1.100668e+04 | 4.459917e+04 | 5.670639e+04 | 3.547170e+04 | 1.269882e+04 | 1.361013e+03 | 1.849957e+02 | 50.360014 | 25.859473 | 24.133057 | 29.784976 | 40.025861 | 79.153975 | 214.305750 | 224.317175 | 327.191104 | 337.997437 | 288.546791 | 371.730457 | 1564.992640 | 2187.472257 | 9132.736564 | 11491.296883 | 20256.434726 | 73770.189927 | 146378.598393 | 135237.997703 | 232670.451962 | 198952.331576 | 106543.065007 | 51163.485300 | 20578.377674 | 8754.087838 | 2439.959306 | 532.859534 | 103.762678 | 45.719573 | 26.858435 | 14.377222 | 14.417020 | 27.163534 | 52.355088 | 119.784963 | 374.798282 | 545.550378 | 560.522239 | 326.274449 | 249.279962 | 165.542922 | 86.854155 | 96.406417 | 64.014941 | 34.836685 | 29.797291 | 18.993175 | 16.164651 | 22.040129 | 16.542843 | 10.083733 | 6.078324 | 4.952815 | 6.286106 | 3.221677 | 1.927140 | 1.073037 | 0.738040 | 0.495084 | 0.436743 | 0.435359 | 0.478684 | 0.592629 | 0.824460 | 1.395164 | 1.770911 | 2.852423 | 1.340440 | 0.889541 | 0.570665 | 0.375302 | 0.284384 | 0.295003 | 0.256545 | 0.276731 | 0.282651 | 0.289162 | 0.417824 | 0.651579 | 0.837260 | 1.248532 | 1.662599 | 2.869527 | 3.696374 | 2.616512 | 2.031928 | 2.003612 | 2.158607 | 3.514080 | 3.946938 | 4.157042 | 4.591414 | 7.577420 | 6.324112 | 3.991226 | 1.805898 | 0.775002 | 0.453453 | 0.394283 | 0.338693 | 0.353711 | 0.394100 | 0.550541 | 0.915802 | 1.528823 | 1.748414 | 2.402540 | 2.639771 | 2.552698 | 1.821536 | 1.463163 | 1.129862 | 1.109168 | 0.724207 | 0.472800 | 0.423829 | 0.381151 | 0.304675 | 0.284238 | 0.294500 | 0.344216 | 0.434692 | 0.459588 | 0.432925 | 0.388476 | 0.364195 | 0.325825 | 0.307142 | 0.277986 | 0.263416 | 0.273109 | 0.266382 | 0.242664 | 0.236439 | 0.221597 | 0.220453 | 0.218179 | 0.222693 | 0.242821 | 0.302595 | 0.340939 | 0.335758 | 0.294871 | 0.294372 | 0.273825 | 0.264517 | 0.262807 | 0.250364 | 0.232400 | 0.222036 | 0.215613 | 0.214687 | 0.214684 | 0.214827 | 0.216503 | 0.236444 | 0.247425 | 0.286209 | 0.336872 | 0.334281 | 0.338169 | 0.342926 | 0.302934 | 0.309164 | 0.268226 | 0.238181 | 0.231681 | 0.220267 | 0.216408 | 0.238404 | 0.292193 | 0.365893 | 0.363825 | 0.360376 | 0.347330 | 0.318321 | 0.285191 | 0.229214 | 0.215865 | 0.214812 | 0.214667 | 0.214873 | 0.216089 | 0.217030 | 0.218235 | 0.236370 | 0.272932 | 0.303071 | 0.388544 | 0.414211 | 0.405818 | 0.300733 | 0.256960 | 0.230122 | 0.214749 | 0.223071 | 0.238564 | 0.266005 | 0.269421 | 0.290772 | 0.302753 | 0.298167 |
| left posterior | 0.771006 | 0.477475 | 0.481650 | 0.446531 | 0.413387 | 0.299898 | 0.219513 | 0.324963 | 0.298162 | 0.327592 | 0.334591 | 0.284895 | 0.271960 | 0.233610 | 0.214840 | 0.217602 | 0.331930 | 0.692513 | 2.973616 | 8.621072 | 31.548759 | 21.873162 | 4.684011 | 0.341226 | 0.262438 | 0.700224 | 1.288507 | 1.268334 | 0.483988 | 0.256146 | 0.214626 | 0.217872 | 0.217723 | 0.214702 | 0.245915 | 0.391012 | 0.621700 | 0.668688 | 0.622627 | 0.732625 | 0.991367 | 2.454603 | 4.637980 | 4.036016 | 3.568537 | 2.916251 | 2.223995 | 1.250555 | 0.341287 | 0.227245 | 0.218822 | 0.258974 | 0.371234 | 0.452091 | 0.374005 | 0.239904 | 0.222499 | 0.214876 | 0.223526 | 0.283937 | 0.363535 | 0.640271 | 1.385365 | 3.723072 | 14.625793 | 108.914829 | 720.339281 | 8177.083137 | 41828.793881 | 190592.521978 | 5.291192e+05 | 1.310877e+06 | 1.020844e+06 | 5.506678e+05 | 1.085433e+05 | 1.619290e+04 | 5.156790e+03 | 3042.769305 | 1277.338479 | 708.164871 | 357.574904 | 139.797099 | 70.028111 | 42.357088 | 32.050225 | 41.357357 | 59.001981 | 73.337409 | 124.204245 | 172.482097 | 184.145667 | 172.942151 | 156.289327 | 202.217642 | 333.700027 | 545.610929 | 615.523472 | 349.730248 | 139.135741 | 120.950456 | 55.029139 | 27.531426 | 12.407444 | 8.629692 | 8.404424 | 9.688188 | 7.363170 | 7.276649 | 8.171563 | 15.553681 | 22.718803 | 23.273128 | 29.569961 | 49.435206 | 113.269759 | 220.609558 | 410.106849 | 852.378390 | 2181.204468 | 5467.123370 | 7696.464703 | 9105.066311 | 10487.596584 | 6372.890079 | 3631.639297 | 2207.901312 | 996.123060 | 400.206682 | 129.431458 | 41.385603 | 22.539559 | 15.739133 | 16.241071 | 16.471343 | 21.137931 | 31.495876 | 43.067678 | 47.015719 | 49.859565 | 65.631887 | 81.153974 | 90.681238 | 112.931223 | 153.495128 | 239.255185 | 159.236331 | 69.214629 | 30.497171 | 22.520002 | 15.054106 | 11.224248 | 7.619039 | 6.815786 | 6.633072 | 11.097319 | 14.518114 | 21.411534 | 44.705964 | 106.008478 | 124.838682 | 91.264720 | 51.473508 | 40.463353 | 38.164421 | 20.486433 | 8.982289 | 9.481938 | 8.260843 | 10.109377 | 9.309492 | 9.309492 | 9.466526 | 11.404923 | 9.029515 | 8.438633 | 5.989386 | 4.943075 | 4.292695 | 4.075144 | 3.695103 | 3.229163 | 3.338793 | 4.006764 | 3.461114 | 2.863251 | 2.586132 | 2.603138 | 3.052412 | 2.268105 | 1.127140 | 0.815841 | 0.765411 | 0.864089 | 0.763687 | 0.756771 | 0.849185 | 1.199078 | 1.940079 | 2.842754 | 2.832121 | 2.566589 | 2.340577 | 2.332141 | 2.192674 | 1.472011 | 1.033638 | 0.796822 | 0.885669 | 0.993324 | 0.910841 | 0.984947 | 1.144337 | 1.622160 | 2.022431 | 2.127535 | 2.194367 | 2.701616 | 3.507161 | 4.841858 | 5.143869 | 4.787447 | 3.798345 | 3.373468 | 3.199085 | 2.641254 | 1.857539 | 1.539860 | 1.433726 | 1.750059 | 1.653837 | 1.698837 | 2.253978 | 2.882144 | 3.058024 | 3.815157 | 2.745492 | 3.133100 | 2.887428 | 3.442789 | 4.529502 | 6.726612 | 9.872798 | 18.515909 | 15.453279 | 14.052195 | 7.373387 | 5.455800 | 3.642950 | 2.242317 | 1.889470 | 1.339825 | 1.352618 | 1.553654 | 1.594873 | 1.440865 | 1.241765 | 1.353569 | 1.905794 | 2.023139 | 2.163803 | 2.182772 | 1.795426 | 1.490056 | 1.032962 | 0.639065 | 0.409914 | 0.291465 | 0.237202 | 0.218958 | 0.215078 | 0.217092 | 0.214859 | 0.215655 | 0.226716 | 0.250536 | 0.305815 | 0.365754 | 0.411094 | 0.442210 | 0.498468 |
| right posterior | 0.245564 | 0.272596 | 0.293707 | 0.554282 | 1.753030 | 4.380208 | 3.176719 | 0.823700 | 0.441164 | 0.369170 | 0.280578 | 0.243673 | 0.215178 | 0.215171 | 0.224968 | 0.311740 | 0.377291 | 0.438167 | 0.500344 | 0.663139 | 0.465186 | 0.244254 | 0.228025 | 0.302985 | 0.465517 | 0.544126 | 0.462768 | 0.243497 | 0.259741 | 0.320829 | 0.374609 | 0.373524 | 0.365493 | 0.296748 | 0.231334 | 0.248001 | 0.336724 | 0.417129 | 0.441881 | 0.832694 | 1.228708 | 1.667881 | 1.350052 | 0.948380 | 0.747731 | 0.830291 | 0.781540 | 0.479356 | 0.274288 | 0.217507 | 0.216570 | 0.220782 | 0.216377 | 0.215027 | 0.259285 | 0.582120 | 1.885438 | 5.389997 | 9.363246 | 7.510958 | 6.250082 | 9.685433 | 15.051152 | 26.174932 | 51.509959 | 117.529718 | 583.292127 | 7314.125083 | 67392.531921 | 526007.236104 | 1.422956e+06 | 2.246230e+06 | 2.242892e+06 | 2.958337e+06 | 9.045002e+05 | 2.510705e+05 | 5.614510e+04 | 22315.343067 | 14517.740845 | 18359.058634 | 5938.912264 | 1283.314508 | 233.088328 | 102.745667 | 85.436465 | 73.968619 | 99.090262 | 218.228846 | 697.180876 | 1550.005652 | 1520.326185 | 1791.660798 | 2861.401896 | 3169.689782 | 4898.145940 | 7311.802066 | 8170.579699 | 7553.476050 | 7110.090926 | 6902.065862 | 7316.364991 | 3471.544801 | 1237.460898 | 416.073745 | 284.485350 | 308.003440 | 235.582800 | 163.923937 | 129.978224 | 104.107117 | 99.025198 | 95.139009 | 136.009048 | 224.591629 | 423.571680 | 819.177409 | 2925.443150 | 7017.323349 | 12909.108044 | 9200.282535 | 4519.442466 | 4464.578533 | 3567.139568 | 2627.470617 | 1949.768733 | 988.692131 | 372.778282 | 256.944720 | 126.155530 | 101.305145 | 93.817094 | 73.387693 | 95.802546 | 174.250448 | 255.364643 | 301.709992 | 308.211067 | 355.251513 | 767.363076 | 1031.604416 | 1693.884934 | 2584.721604 | 3710.728250 | 3738.317372 | 3805.251683 | 2673.527410 | 1494.359285 | 593.338055 | 196.521177 | 95.003787 | 78.883971 | 60.237496 | 84.677726 | 132.360122 | 267.598323 | 613.543757 | 817.225002 | 944.277661 | 923.872086 | 664.751081 | 411.068278 | 238.209791 | 164.482200 | 134.878163 | 140.071065 | 186.480398 | 202.983315 | 186.529651 | 190.489451 | 154.896083 | 189.398306 | 160.830258 | 147.282556 | 154.684361 | 155.702848 | 205.846209 | 390.152989 | 558.257722 | 761.360678 | 549.452154 | 403.440389 | 334.802586 | 275.189499 | 169.053694 | 117.414878 | 120.443175 | 169.355380 | 214.508408 | 345.781254 | 299.722243 | 360.867344 | 304.762592 | 250.710277 | 121.975152 | 51.647348 | 22.689041 | 12.542782 | 6.472661 | 5.388035 | 4.232740 | 5.637363 | 8.251038 | 10.050422 | 15.631774 | 26.503707 | 27.764339 | 25.518947 | 22.513247 | 23.958136 | 38.328449 | 61.647009 | 90.242841 | 162.883504 | 389.284408 | 541.642561 | 1169.544498 | 2148.402129 | 2440.437924 | 2881.360167 | 1303.258824 | 702.133315 | 456.576569 | 248.904552 | 136.990707 | 77.756743 | 48.828147 | 65.151086 | 62.464193 | 61.613941 | 65.998720 | 70.095186 | 101.447232 | 113.251211 | 123.380737 | 101.616284 | 84.504156 | 111.260407 | 153.704242 | 151.454690 | 155.349758 | 116.199368 | 93.202618 | 84.614127 | 75.823077 | 83.813934 | 107.215881 | 137.099008 | 171.719058 | 134.557741 | 107.911102 | 95.648239 | 62.870138 | 38.312367 | 25.858719 | 26.548274 | 38.621819 | 66.767609 | 104.509164 | 306.559057 | 834.466178 | 1503.541995 | 3269.381163 | 6932.503624 | 10313.121998 | 6466.193092 | 1164.782933 | 229.041117 | 88.697409 | 44.505575 | 30.857140 | 27.425697 | 18.247858 | 22.487581 | 20.089407 | 18.900132 | 23.513592 | 26.506031 | 24.894760 | 26.075495 | 20.199594 |
| all electrodes | 0.815922 | 1.174110 | 0.923803 | 0.441720 | 0.271620 | 0.218255 | 0.229170 | 0.252533 | 0.301253 | 0.395410 | 0.238866 | 0.224838 | 0.221271 | 0.252521 | 0.234919 | 0.216651 | 0.363766 | 2.078795 | 5.634517 | 5.863572 | 4.354474 | 3.706805 | 0.707738 | 0.244221 | 0.239747 | 0.378732 | 0.450037 | 0.298024 | 0.275570 | 0.216761 | 0.218440 | 0.222128 | 0.217168 | 0.244252 | 0.325907 | 0.644091 | 0.968278 | 0.816633 | 0.693014 | 0.440386 | 0.312503 | 0.284731 | 0.289133 | 0.265929 | 0.260920 | 0.248148 | 0.238063 | 0.218404 | 0.214637 | 0.225488 | 0.228632 | 0.233820 | 0.217223 | 0.214754 | 0.257881 | 0.344528 | 0.553520 | 0.969617 | 1.259204 | 1.241775 | 1.403337 | 1.338300 | 1.496555 | 2.297223 | 5.093497 | 22.995620 | 186.667450 | 1614.485041 | 10847.399426 | 65020.539629 | 7.351689e+05 | 7.849094e+06 | 2.488721e+07 | 2.521965e+07 | 1.463823e+07 | 3.940995e+06 | 2.251075e+06 | 502908.604901 | 92718.992549 | 49830.940743 | 25098.706818 | 6487.334064 | 2189.008585 | 275.972160 | 157.661866 | 110.278560 | 108.115724 | 79.429489 | 72.779872 | 52.966688 | 102.884857 | 92.418871 | 130.669298 | 477.648857 | 2281.607841 | 4267.364159 | 3853.717387 | 2941.410808 | 4164.313362 | 4219.396852 | 1349.790806 | 650.410564 | 720.225149 | 568.791911 | 459.976336 | 258.501057 | 241.506077 | 206.797735 | 173.795329 | 197.450746 | 466.822005 | 859.465052 | 1575.889706 | 1071.530473 | 944.556637 | 1074.487324 | 657.430717 | 740.322735 | 712.474455 | 678.706781 | 836.153171 | 1136.126797 | 2937.706814 | 5509.181982 | 3899.036378 | 2143.011626 | 1150.123582 | 1364.761461 | 1286.456071 | 681.114451 | 797.983972 | 1076.039879 | 1218.730869 | 1033.349651 | 322.162557 | 136.377715 | 99.361620 | 68.660357 | 95.821853 | 223.798936 | 582.788946 | 1573.986987 | 1137.146539 | 312.354472 | 126.778137 | 43.889089 | 15.240369 | 5.185776 | 4.006861 | 5.799200 | 14.512527 | 42.236484 | 111.659297 | 223.259159 | 244.269938 | 318.592486 | 206.802749 | 97.996855 | 39.694905 | 25.519870 | 12.585705 | 12.223441 | 9.409057 | 9.670978 | 6.800322 | 6.174160 | 6.324567 | 10.070599 | 8.675247 | 6.447491 | 3.716363 | 4.481314 | 3.214788 | 1.392880 | 0.884395 | 0.959165 | 1.111701 | 1.244884 | 1.031552 | 1.186825 | 1.653632 | 2.270576 | 1.468094 | 1.255782 | 1.496128 | 1.297468 | 1.191095 | 1.569714 | 1.286693 | 0.942935 | 0.670042 | 0.672716 | 0.891071 | 0.973101 | 0.723703 | 0.597045 | 0.755005 | 0.979611 | 0.915134 | 0.693256 | 0.538071 | 0.507525 | 0.567050 | 0.604344 | 0.542743 | 0.454588 | 0.513132 | 0.428396 | 0.415584 | 0.350776 | 0.289402 | 0.282761 | 0.299612 | 0.294837 | 0.359551 | 0.401273 | 0.488789 | 0.621066 | 0.548188 | 0.460442 | 0.352444 | 0.282089 | 0.281094 | 0.312716 | 0.326580 | 0.310939 | 0.262310 | 0.253191 | 0.260061 | 0.234161 | 0.214793 | 0.218525 | 0.215229 | 0.214781 | 0.218125 | 0.228610 | 0.254446 | 0.281713 | 0.309184 | 0.331391 | 0.411101 | 0.474374 | 0.477983 | 0.393781 | 0.295106 | 0.269974 | 0.255271 | 0.245079 | 0.217737 | 0.217714 | 0.225624 | 0.223050 | 0.226517 | 0.224686 | 0.221397 | 0.217856 | 0.217668 | 0.215206 | 0.221392 | 0.246773 | 0.287375 | 0.302717 | 0.356214 | 0.373143 | 0.385744 | 0.279032 | 0.228905 | 0.215599 | 0.216393 | 0.228232 | 0.224602 | 0.225647 | 0.218171 | 0.214945 | 0.215030 | 0.219893 | 0.235447 | 0.237944 | 0.240418 | 0.250392 |

Searchlight, spatiotemporal cluster permutation test

|  | start time | stop time | peak time | peak channel | cluster p | peak Cohen's d | direction |
| --- | --- | --- | --- | --- | --- | --- | --- |
| #1 | 80 | 1195 | 150 | POz | 0.0001 | 1.691478 | positive |

N) real faces (LOSO) - neutral vs sad

  
|  | time window | peak latency | cluster *p* | peak Cohen's *d* |  | | | |  |
| **all electrodes** | 115 - 205 ms | 160 ms | 0.0313 | 1.0432 |  | | | |  |
 840 - 1105 ms | 865 ms | 0.0062 | 0.7572 |  | | | ||  |  | | | | | | | | |

Time-resolved classification, cluster permutation tests

|  | **left hemisphere** | | | | **right hemisphere** | | | |  |
|  | time window | peak latency | cluster *p* | peak Cohen's *d* | time window | peak latency | cluster *p* | peak Cohen's *d* |  |
| **anterior** |  | | | | 210 - 295 ms | 250 ms | 0.0435 | 0.7589 |  |
| **central** | 90 - 195 ms | 165 ms | 0.0217 | 0.8846 | 205 - 320 ms | 255 ms | 0.0122 | 0.6973 |  |
| **posterior** | 120 - 450 ms | 155 ms | 0.0064 | 1.304 | 235 - 550 ms | 265 ms | 0.0029 | 0.8002 |  |
 805 - 1015 ms | 935 ms | 0.0268 | 0.6426 |  | | | |

  

Time-resolved classification, Bayesian statistics

|  | -200 | -195 | -190 | -185 | -180 | -175 | -170 | -165 | -160 | -155 | -150 | -145 | -140 | -135 | -130 | -125 | -120 | -115 | -110 | -105 | -100 | -95 | -90 | -85 | -80 | -75 | -70 | -65 | -60 | -55 | -50 | -45 | -40 | -35 | -30 | -25 | -20 | -15 | -10 | -5 | 0 | 5 | 10 | 15 | 20 | 25 | 30 | 35 | 40 | 45 | 50 | 55 | 60 | 65 | 70 | 75 | 80 | 85 | 90 | 95 | 100 | 105 | 110 | 115 | 120 | 125 | 130 | 135 | 140 | 145 | 150 | 155 | 160 | 165 | 170 | 175 | 180 | 185 | 190 | 195 | 200 | 205 | 210 | 215 | 220 | 225 | 230 | 235 | 240 | 245 | 250 | 255 | 260 | 265 | 270 | 275 | 280 | 285 | 290 | 295 | 300 | 305 | 310 | 315 | 320 | 325 | 330 | 335 | 340 | 345 | 350 | 355 | 360 | 365 | 370 | 375 | 380 | 385 | 390 | 395 | 400 | 405 | 410 | 415 | 420 | 425 | 430 | 435 | 440 | 445 | 450 | 455 | 460 | 465 | 470 | 475 | 480 | 485 | 490 | 495 | 500 | 505 | 510 | 515 | 520 | 525 | 530 | 535 | 540 | 545 | 550 | 555 | 560 | 565 | 570 | 575 | 580 | 585 | 590 | 595 | 600 | 605 | 610 | 615 | 620 | 625 | 630 | 635 | 640 | 645 | 650 | 655 | 660 | 665 | 670 | 675 | 680 | 685 | 690 | 695 | 700 | 705 | 710 | 715 | 720 | 725 | 730 | 735 | 740 | 745 | 750 | 755 | 760 | 765 | 770 | 775 | 780 | 785 | 790 | 795 | 800 | 805 | 810 | 815 | 820 | 825 | 830 | 835 | 840 | 845 | 850 | 855 | 860 | 865 | 870 | 875 | 880 | 885 | 890 | 895 | 900 | 905 | 910 | 915 | 920 | 925 | 930 | 935 | 940 | 945 | 950 | 955 | 960 | 965 | 970 | 975 | 980 | 985 | 990 | 995 | 1000 | 1005 | 1010 | 1015 | 1020 | 1025 | 1030 | 1035 | 1040 | 1045 | 1050 | 1055 | 1060 | 1065 | 1070 | 1075 | 1080 | 1085 | 1090 | 1095 | 1100 | 1105 | 1110 | 1115 | 1120 | 1125 | 1130 | 1135 | 1140 | 1145 | 1150 | 1155 | 1160 | 1165 | 1170 | 1175 | 1180 | 1185 | 1190 | 1195 |
| --- | --- | --- | --- | --- | --- | --- | --- | --- | --- | --- | --- | --- | --- | --- | --- | --- | --- | --- | --- | --- | --- | --- | --- | --- | --- | --- | --- | --- | --- | --- | --- | --- | --- | --- | --- | --- | --- | --- | --- | --- | --- | --- | --- | --- | --- | --- | --- | --- | --- | --- | --- | --- | --- | --- | --- | --- | --- | --- | --- | --- | --- | --- | --- | --- | --- | --- | --- | --- | --- | --- | --- | --- | --- | --- | --- | --- | --- | --- | --- | --- | --- | --- | --- | --- | --- | --- | --- | --- | --- | --- | --- | --- | --- | --- | --- | --- | --- | --- | --- | --- | --- | --- | --- | --- | --- | --- | --- | --- | --- | --- | --- | --- | --- | --- | --- | --- | --- | --- | --- | --- | --- | --- | --- | --- | --- | --- | --- | --- | --- | --- | --- | --- | --- | --- | --- | --- | --- | --- | --- | --- | --- | --- | --- | --- | --- | --- | --- | --- | --- | --- | --- | --- | --- | --- | --- | --- | --- | --- | --- | --- | --- | --- | --- | --- | --- | --- | --- | --- | --- | --- | --- | --- | --- | --- | --- | --- | --- | --- | --- | --- | --- | --- | --- | --- | --- | --- | --- | --- | --- | --- | --- | --- | --- | --- | --- | --- | --- | --- | --- | --- | --- | --- | --- | --- | --- | --- | --- | --- | --- | --- | --- | --- | --- | --- | --- | --- | --- | --- | --- | --- | --- | --- | --- | --- | --- | --- | --- | --- | --- | --- | --- | --- | --- | --- | --- | --- | --- | --- | --- | --- | --- | --- | --- | --- | --- | --- | --- | --- | --- | --- | --- | --- | --- | --- | --- | --- | --- | --- | --- | --- | --- | --- | --- | --- | --- | --- | --- | --- | --- | --- | --- | --- | --- | --- | --- | --- | --- | --- | --- | --- |
| left anterior | 0.280217 | 0.237386 | 0.229923 | 0.275436 | 0.326198 | 0.333391 | 0.216648 | 0.214692 | 0.220342 | 0.224654 | 0.216812 | 0.215302 | 0.245728 | 0.356752 | 0.319003 | 0.300233 | 0.309994 | 0.298510 | 0.256259 | 0.216925 | 0.214654 | 0.270394 | 0.329114 | 0.309925 | 0.443053 | 0.872232 | 0.694203 | 0.512184 | 0.379891 | 0.290770 | 0.243969 | 0.215784 | 0.231273 | 0.281962 | 0.369480 | 0.783934 | 2.481190 | 1.994389 | 8.069053 | 33.290597 | 47.002320 | 16.818522 | 9.485562 | 7.868650 | 15.399618 | 3.994753 | 2.679111 | 6.332094 | 10.553273 | 23.018940 | 27.034744 | 8.704303 | 5.970735 | 2.871355 | 1.632964 | 0.774385 | 0.378832 | 0.246613 | 0.227101 | 0.216538 | 0.217989 | 0.215229 | 0.224476 | 0.279624 | 0.320198 | 0.400628 | 0.610947 | 1.609406 | 5.365967 | 9.162666 | 8.940070 | 5.749931 | 3.430395 | 1.682589 | 0.746145 | 0.403465 | 0.232538 | 0.220858 | 0.267698 | 0.339835 | 0.331841 | 0.329355 | 0.326037 | 0.256009 | 0.215061 | 0.224522 | 0.261545 | 0.346270 | 0.590152 | 0.853884 | 1.680388 | 2.339972 | 3.927289 | 5.616948 | 5.434385 | 4.723541 | 5.460607 | 4.058161 | 4.403926 | 3.306548 | 1.923935 | 1.474380 | 1.330012 | 0.897874 | 0.891421 | 0.628355 | 0.476496 | 0.354453 | 0.290963 | 0.244355 | 0.237945 | 0.228590 | 0.236046 | 0.277804 | 0.364225 | 0.357638 | 0.353757 | 0.352877 | 0.329861 | 0.315983 | 0.255252 | 0.230155 | 0.233385 | 0.278610 | 0.317383 | 0.309723 | 0.262229 | 0.249927 | 0.233799 | 0.219971 | 0.226455 | 0.311055 | 0.417576 | 0.391129 | 0.348786 | 0.321485 | 0.316423 | 0.293016 | 0.253106 | 0.217819 | 0.214626 | 0.215051 | 0.214626 | 0.219788 | 0.249160 | 0.233944 | 0.215416 | 0.227393 | 0.249486 | 0.241634 | 0.224025 | 0.217029 | 0.228771 | 0.281413 | 0.350250 | 0.553337 | 0.805201 | 1.051219 | 1.219040 | 0.899448 | 0.989253 | 1.024866 | 1.069626 | 1.208755 | 1.042765 | 0.869940 | 0.834227 | 0.632174 | 0.513056 | 0.454125 | 0.354239 | 0.316033 | 0.286897 | 0.263639 | 0.252427 | 0.258664 | 0.273175 | 0.297291 | 0.338955 | 0.440355 | 0.683543 | 0.955686 | 0.802313 | 0.692646 | 0.715749 | 0.688395 | 0.982097 | 1.083955 | 1.269379 | 2.145374 | 1.292396 | 0.619692 | 0.385893 | 0.282029 | 0.308125 | 0.408270 | 0.480169 | 0.766092 | 1.438255 | 2.772857 | 3.696064 | 1.674335 | 0.743675 | 0.466417 | 0.323027 | 0.266668 | 0.250655 | 0.254986 | 0.348593 | 0.515571 | 0.693202 | 0.836675 | 1.171650 | 1.663779 | 1.460886 | 0.814882 | 0.906608 | 0.840919 | 0.820403 | 0.934007 | 0.812766 | 0.751455 | 0.678071 | 0.480672 | 0.431146 | 0.491593 | 0.493049 | 0.673558 | 0.909816 | 1.248876 | 1.389875 | 1.181493 | 0.952567 | 0.778695 | 0.751204 | 0.969810 | 1.109280 | 1.119283 | 1.121432 | 1.173209 | 1.309293 | 0.902184 | 0.616147 | 0.503043 | 0.463350 | 0.486424 | 0.434848 | 0.390063 | 0.413651 | 0.390909 | 0.395376 | 0.391504 | 0.381785 | 0.374695 | 0.368271 | 0.349235 | 0.346438 | 0.306751 | 0.287042 | 0.254745 | 0.260508 | 0.251134 | 0.233169 | 0.229122 | 0.244218 | 0.263478 | 0.297807 | 0.320321 | 0.368438 | 0.563948 | 0.794416 | 0.780134 | 0.695872 | 0.719923 | 0.776847 | 0.724615 | 0.589073 | 0.510934 | 0.562154 | 0.791704 |
| right anterior | 0.296820 | 0.256162 | 0.241051 | 0.235282 | 0.229147 | 0.259363 | 0.230563 | 0.234839 | 0.285486 | 0.449434 | 0.678890 | 0.515804 | 0.224787 | 0.217852 | 0.220130 | 0.227202 | 0.259338 | 0.332833 | 0.250531 | 0.217972 | 0.302320 | 0.449057 | 0.438495 | 0.535376 | 0.501151 | 0.548672 | 0.559303 | 0.552104 | 0.303008 | 0.247015 | 0.215793 | 0.215519 | 0.231547 | 0.239176 | 0.268187 | 0.227888 | 0.233058 | 0.233906 | 0.239746 | 0.246501 | 0.374943 | 0.376423 | 0.370163 | 0.262065 | 0.224625 | 0.225442 | 0.225129 | 0.214803 | 0.218086 | 0.221685 | 0.239062 | 0.238409 | 0.215497 | 0.215365 | 0.220249 | 0.252926 | 0.323367 | 0.439044 | 0.617703 | 0.699067 | 0.748403 | 0.594160 | 0.399914 | 0.388363 | 0.445900 | 0.636665 | 1.252731 | 3.046256 | 7.012693 | 17.331338 | 21.869755 | 20.039144 | 10.693399 | 3.013452 | 1.287118 | 0.897383 | 0.603074 | 0.515278 | 0.415895 | 0.421769 | 0.529561 | 0.900436 | 1.531274 | 4.338163 | 16.995360 | 82.127631 | 159.074025 | 171.667022 | 134.088989 | 69.374941 | 31.512632 | 16.131023 | 12.813863 | 10.567889 | 11.907465 | 10.839653 | 7.078016 | 4.389340 | 2.380338 | 1.347875 | 0.899891 | 0.635567 | 0.465417 | 0.548559 | 0.842839 | 1.519075 | 4.217506 | 10.694373 | 12.143725 | 12.148402 | 7.513514 | 4.283928 | 2.926227 | 1.454865 | 0.967357 | 0.846220 | 0.629525 | 0.446968 | 0.364652 | 0.327070 | 0.373277 | 0.439322 | 0.383405 | 0.302726 | 0.239192 | 0.215120 | 0.230289 | 0.283160 | 0.352767 | 0.281209 | 0.227734 | 0.221730 | 0.236976 | 0.232443 | 0.220833 | 0.217954 | 0.214861 | 0.248660 | 0.437455 | 0.642263 | 0.662723 | 0.869607 | 1.741835 | 3.663066 | 5.341013 | 3.910494 | 3.764737 | 8.764399 | 3.177925 | 0.834155 | 0.383437 | 0.247983 | 0.233864 | 0.270921 | 0.272854 | 0.262559 | 0.255004 | 0.290356 | 0.420940 | 0.573271 | 0.468294 | 0.318735 | 0.264016 | 0.248056 | 0.225733 | 0.214660 | 0.225124 | 0.257577 | 0.269831 | 0.267745 | 0.241279 | 0.219815 | 0.214739 | 0.215284 | 0.217117 | 0.219392 | 0.223172 | 0.221881 | 0.221767 | 0.219270 | 0.217552 | 0.215042 | 0.219447 | 0.219984 | 0.218793 | 0.217733 | 0.214928 | 0.215699 | 0.214633 | 0.216794 | 0.214976 | 0.234115 | 0.297122 | 0.359961 | 0.339836 | 0.299727 | 0.281407 | 0.253320 | 0.224940 | 0.214873 | 0.215866 | 0.220300 | 0.215565 | 0.216760 | 0.214844 | 0.214663 | 0.215736 | 0.214847 | 0.214626 | 0.226877 | 0.250181 | 0.266220 | 0.268299 | 0.250341 | 0.234922 | 0.238826 | 0.218491 | 0.217540 | 0.231690 | 0.239873 | 0.284013 | 0.279612 | 0.268240 | 0.261776 | 0.252524 | 0.278759 | 0.339617 | 0.361562 | 0.363567 | 0.285239 | 0.275350 | 0.235634 | 0.214636 | 0.225562 | 0.232924 | 0.217354 | 0.236545 | 0.246551 | 0.271749 | 0.335423 | 0.326109 | 0.246463 | 0.215146 | 0.232962 | 0.230410 | 0.215639 | 0.218350 | 0.215443 | 0.215594 | 0.216321 | 0.220680 | 0.225026 | 0.241388 | 0.231191 | 0.241216 | 0.234907 | 0.227224 | 0.219456 | 0.222032 | 0.224247 | 0.222964 | 0.220932 | 0.216549 | 0.214998 | 0.215972 | 0.217942 | 0.223619 | 0.241521 | 0.263470 | 0.271025 | 0.257258 | 0.259497 | 0.275036 | 0.281395 | 0.277444 | 0.269471 | 0.268606 | 0.325983 | 0.346927 | 0.303357 |
| left central | 0.349728 | 0.478791 | 0.507831 | 0.451507 | 0.987583 | 3.780547 | 4.194017 | 1.070827 | 0.556498 | 0.512942 | 0.296418 | 0.267957 | 0.218025 | 0.229000 | 0.234836 | 0.226486 | 0.239211 | 0.296665 | 0.264157 | 0.299624 | 0.313182 | 0.300700 | 0.394255 | 0.547707 | 1.288038 | 2.527277 | 2.979360 | 1.694836 | 0.965709 | 0.433759 | 0.232602 | 0.399488 | 0.823037 | 1.059276 | 0.530730 | 0.245262 | 0.228058 | 0.223991 | 0.215943 | 0.221136 | 0.225803 | 0.217405 | 0.221848 | 0.215038 | 0.215270 | 0.219391 | 0.221383 | 0.223711 | 0.256029 | 0.340813 | 0.319999 | 0.282530 | 0.246743 | 0.253403 | 0.242558 | 0.250579 | 0.307483 | 0.557709 | 1.566281 | 5.221706 | 9.070476 | 10.168542 | 9.091959 | 8.013484 | 6.219577 | 3.843920 | 2.944332 | 4.195899 | 17.342584 | 43.256837 | 59.923841 | 43.614997 | 51.417078 | 121.420320 | 169.538950 | 81.908504 | 33.583269 | 13.035433 | 5.614637 | 2.014321 | 1.198072 | 0.878160 | 0.750680 | 0.646406 | 0.726461 | 1.107005 | 1.206778 | 1.081325 | 1.085591 | 1.385836 | 1.687722 | 1.776694 | 1.536987 | 1.413152 | 1.255111 | 1.098019 | 0.784417 | 0.549382 | 0.423213 | 0.390517 | 0.421765 | 0.407080 | 0.394468 | 0.414345 | 0.474570 | 0.540741 | 0.417239 | 0.299903 | 0.265282 | 0.244795 | 0.232989 | 0.225677 | 0.214678 | 0.215714 | 0.230259 | 0.217556 | 0.229917 | 0.261834 | 0.269952 | 0.231339 | 0.219039 | 0.224309 | 0.227935 | 0.215572 | 0.215347 | 0.227658 | 0.260651 | 0.350386 | 0.393251 | 0.329880 | 0.283089 | 0.240699 | 0.230624 | 0.270441 | 0.304341 | 0.278469 | 0.320383 | 0.330792 | 0.380506 | 0.325818 | 0.237861 | 0.217213 | 0.222303 | 0.215219 | 0.214637 | 0.215159 | 0.221068 | 0.248473 | 0.318389 | 0.343857 | 0.370330 | 0.282460 | 0.236898 | 0.231996 | 0.221643 | 0.223705 | 0.238675 | 0.278972 | 0.383207 | 0.437405 | 0.321230 | 0.293742 | 0.241478 | 0.235766 | 0.231332 | 0.225260 | 0.218487 | 0.217357 | 0.216541 | 0.226718 | 0.249682 | 0.316406 | 0.410192 | 0.628755 | 1.097417 | 1.734773 | 2.158064 | 1.229976 | 0.509297 | 0.403300 | 0.338306 | 0.302257 | 0.262201 | 0.250615 | 0.250909 | 0.243876 | 0.237623 | 0.226685 | 0.214896 | 0.214626 | 0.215402 | 0.218996 | 0.266106 | 0.375903 | 0.480141 | 0.735221 | 1.010165 | 0.849410 | 0.554100 | 0.350868 | 0.298303 | 0.327928 | 0.402425 | 0.451814 | 0.499656 | 0.443016 | 0.396133 | 0.346678 | 0.272179 | 0.229952 | 0.214885 | 0.235510 | 0.301311 | 0.507871 | 1.093852 | 1.450640 | 1.464668 | 1.104575 | 0.912613 | 0.740030 | 0.549647 | 0.526948 | 0.507414 | 0.556131 | 0.671304 | 0.711123 | 0.587502 | 0.412838 | 0.275970 | 0.216505 | 0.217639 | 0.231888 | 0.244086 | 0.247419 | 0.241323 | 0.235888 | 0.215230 | 0.214887 | 0.227753 | 0.248669 | 0.269851 | 0.283306 | 0.256294 | 0.222770 | 0.219590 | 0.222706 | 0.232862 | 0.249956 | 0.259712 | 0.316245 | 0.413323 | 0.429949 | 0.369785 | 0.253143 | 0.224433 | 0.227374 | 0.233345 | 0.223663 | 0.215501 | 0.216686 | 0.215134 | 0.216071 | 0.225782 | 0.242412 | 0.225743 | 0.216247 | 0.216962 | 0.223460 | 0.226251 | 0.248404 | 0.282060 | 0.257309 | 0.249319 | 0.246082 | 0.233559 | 0.220314 | 0.216798 | 0.215125 | 0.221874 | 0.235403 |
| right central | 0.232822 | 0.222336 | 0.226837 | 0.431079 | 0.498696 | 0.960303 | 2.479289 | 3.321681 | 13.672045 | 13.489302 | 0.993524 | 1.590285 | 1.469755 | 1.002295 | 0.542548 | 0.328924 | 0.271496 | 0.350878 | 0.228455 | 0.239430 | 0.320139 | 0.439939 | 0.332970 | 0.297023 | 0.310533 | 0.245907 | 0.214904 | 0.256831 | 0.495434 | 1.226214 | 0.536662 | 0.273364 | 0.214903 | 0.216925 | 0.272665 | 0.245990 | 0.249370 | 0.261664 | 0.241266 | 0.243260 | 0.288543 | 0.239755 | 0.221998 | 0.220645 | 0.217779 | 0.236370 | 0.250466 | 0.257757 | 0.227669 | 0.217089 | 0.262758 | 0.288970 | 0.328183 | 0.282808 | 0.266283 | 0.247666 | 0.217676 | 0.216934 | 0.215100 | 0.215935 | 0.248124 | 0.349218 | 0.434498 | 0.629829 | 1.705538 | 4.474673 | 15.072882 | 30.139159 | 40.595161 | 80.124079 | 165.961981 | 164.220775 | 167.971599 | 118.854928 | 41.656424 | 11.780666 | 5.054394 | 1.928451 | 1.154640 | 0.921626 | 1.134832 | 2.150659 | 3.771676 | 4.011294 | 6.325744 | 8.317174 | 14.227365 | 14.238047 | 9.156149 | 8.600064 | 11.989619 | 16.566819 | 20.450578 | 35.729738 | 94.267625 | 254.395332 | 388.980764 | 201.560627 | 37.568636 | 41.865033 | 32.979182 | 17.059080 | 11.752499 | 6.244193 | 1.471825 | 1.060958 | 0.798929 | 0.865457 | 0.520020 | 0.346314 | 0.363755 | 0.520131 | 0.555284 | 0.461340 | 0.301341 | 0.261722 | 0.226605 | 0.223117 | 0.255744 | 0.251423 | 0.233336 | 0.220545 | 0.221026 | 0.214838 | 0.217462 | 0.215392 | 0.214720 | 0.216885 | 0.231535 | 0.227427 | 0.234478 | 0.224721 | 0.219429 | 0.214626 | 0.227970 | 0.242743 | 0.253706 | 0.256442 | 0.266001 | 0.309041 | 0.272908 | 0.242090 | 0.235856 | 0.226959 | 0.218354 | 0.215643 | 0.235591 | 0.257517 | 0.273386 | 0.254981 | 0.230105 | 0.218041 | 0.214740 | 0.214742 | 0.215007 | 0.215617 | 0.214765 | 0.214710 | 0.240254 | 0.280028 | 0.442051 | 1.014654 | 1.929440 | 4.038286 | 10.290596 | 8.906254 | 14.344735 | 19.413749 | 11.610095 | 2.343701 | 0.388412 | 0.214647 | 0.424385 | 1.100754 | 1.354831 | 0.944528 | 0.429787 | 0.261777 | 0.242608 | 0.216218 | 0.215056 | 0.215549 | 0.217386 | 0.235574 | 0.291657 | 0.275122 | 0.299026 | 0.355270 | 0.433209 | 0.318644 | 0.244129 | 0.217159 | 0.214859 | 0.216462 | 0.214694 | 0.219721 | 0.243463 | 0.321589 | 0.438910 | 0.435385 | 0.308787 | 0.237976 | 0.219677 | 0.218416 | 0.215513 | 0.216023 | 0.235983 | 0.293765 | 0.367078 | 0.411195 | 0.396130 | 0.373526 | 0.314853 | 0.270378 | 0.261499 | 0.246045 | 0.237258 | 0.244399 | 0.260310 | 0.313430 | 0.347385 | 0.404903 | 0.347624 | 0.278121 | 0.247403 | 0.247044 | 0.256622 | 0.362679 | 0.376134 | 0.618599 | 0.816516 | 0.792066 | 0.756228 | 0.396511 | 0.244512 | 0.227213 | 0.229175 | 0.253628 | 0.310866 | 0.389367 | 0.469648 | 0.462822 | 0.376142 | 0.352323 | 0.331135 | 0.324452 | 0.309189 | 0.332332 | 0.429499 | 0.662929 | 0.624995 | 0.621183 | 0.513718 | 0.466417 | 0.494848 | 0.439415 | 0.401812 | 0.423947 | 0.490623 | 0.555493 | 0.424986 | 0.440693 | 0.602558 | 0.790923 | 0.889278 | 0.679910 | 0.568500 | 0.601832 | 0.490828 | 0.386279 | 0.311415 | 0.299382 | 0.295940 | 0.298903 | 0.313847 | 0.385769 | 0.667348 | 0.859677 | 1.042467 | 1.324855 |
| left posterior | 0.340836 | 0.583696 | 1.000669 | 1.105780 | 1.534666 | 1.625647 | 0.778638 | 0.308587 | 0.217004 | 0.256126 | 0.385834 | 0.574157 | 0.634362 | 0.574000 | 0.426794 | 0.322325 | 0.228742 | 0.221763 | 0.288012 | 0.321890 | 0.343332 | 0.340768 | 0.274812 | 0.283518 | 0.282865 | 0.262115 | 0.428463 | 0.527302 | 2.400617 | 8.748947 | 18.033484 | 22.534098 | 29.259201 | 7.317767 | 1.871195 | 0.519544 | 0.304750 | 0.222990 | 0.221277 | 0.308278 | 0.412507 | 0.418118 | 0.443198 | 0.338433 | 0.259781 | 0.225169 | 0.216779 | 0.253127 | 0.376980 | 0.560825 | 1.039806 | 2.290266 | 10.144037 | 34.509683 | 153.038393 | 430.325762 | 610.551715 | 87.094532 | 13.366945 | 3.227631 | 1.683003 | 0.596992 | 0.215042 | 0.561258 | 4.181063 | 42.999236 | 130.164051 | 505.819042 | 1940.535954 | 5993.127561 | 8280.876312 | 11522.148237 | 7474.081633 | 5721.618841 | 6664.682016 | 1843.851587 | 526.335277 | 148.940501 | 41.077597 | 9.232650 | 3.072899 | 1.671962 | 1.508707 | 2.121304 | 2.474984 | 2.606588 | 3.823990 | 6.753024 | 7.813043 | 9.346308 | 9.035134 | 9.315689 | 10.479395 | 11.655950 | 11.227295 | 13.797846 | 11.014449 | 8.235184 | 6.457412 | 7.658770 | 9.306018 | 11.133214 | 14.096419 | 23.341375 | 29.764722 | 36.486437 | 25.498328 | 11.646586 | 6.907211 | 4.628231 | 4.508303 | 5.451851 | 5.326256 | 6.480582 | 12.914235 | 25.384285 | 50.127508 | 51.874385 | 54.633399 | 76.948836 | 77.111954 | 59.618268 | 44.456419 | 40.873707 | 44.247058 | 25.031277 | 11.320293 | 5.769684 | 4.233847 | 3.055844 | 1.704365 | 1.291950 | 1.453851 | 1.788934 | 2.289816 | 2.418223 | 2.221161 | 2.981070 | 4.185133 | 5.149721 | 5.857319 | 5.644886 | 6.113201 | 6.356005 | 4.598448 | 3.303677 | 2.221685 | 1.466650 | 1.134865 | 0.766048 | 0.607578 | 0.541276 | 0.465621 | 0.477049 | 0.533596 | 0.590869 | 0.758708 | 0.957074 | 1.317937 | 1.946291 | 2.952976 | 5.147837 | 8.448902 | 14.744253 | 17.223485 | 15.470616 | 14.925464 | 8.764556 | 5.197724 | 2.999782 | 1.879988 | 1.395827 | 0.918664 | 0.820846 | 0.924718 | 0.937167 | 0.960816 | 1.405852 | 1.949226 | 2.686872 | 4.738190 | 7.503516 | 8.660927 | 8.866892 | 4.920615 | 3.263418 | 2.337917 | 1.233616 | 0.775925 | 0.623655 | 0.551774 | 0.478948 | 0.442248 | 0.437086 | 0.500667 | 0.499429 | 0.464658 | 0.518640 | 0.641284 | 0.929634 | 1.152254 | 1.364439 | 1.827748 | 2.439978 | 2.998913 | 3.643092 | 3.028708 | 2.842569 | 2.621417 | 1.820649 | 1.817639 | 2.018726 | 1.886948 | 1.941092 | 2.224051 | 2.090135 | 2.330495 | 2.259626 | 1.620686 | 1.639790 | 1.634471 | 2.320088 | 3.356739 | 3.948279 | 4.421476 | 7.457415 | 8.615166 | 9.524291 | 7.679386 | 6.541988 | 5.545607 | 4.837868 | 4.045295 | 5.618595 | 6.298667 | 8.721454 | 8.841883 | 8.078948 | 7.236766 | 5.407198 | 3.603680 | 2.869116 | 1.835332 | 1.502986 | 1.266725 | 0.945232 | 0.901330 | 0.945768 | 1.144305 | 1.228677 | 1.152742 | 1.207958 | 1.521878 | 1.806545 | 1.619353 | 1.443886 | 1.970174 | 2.715106 | 3.866635 | 3.667690 | 2.920334 | 2.554912 | 2.331778 | 1.580432 | 1.365605 | 1.192820 | 1.111132 | 1.037745 | 1.083447 | 0.936333 | 0.878018 | 0.793817 | 0.724361 | 0.687140 | 0.739492 | 0.655026 | 0.605893 | 0.647320 | 0.692698 | 0.613750 |
| right posterior | 0.396125 | 0.379644 | 0.360093 | 0.464013 | 0.530163 | 0.377683 | 0.256513 | 0.216478 | 0.218708 | 0.226429 | 0.218710 | 0.253567 | 0.239435 | 0.248355 | 0.237885 | 0.294219 | 0.270312 | 0.216923 | 0.234416 | 0.214908 | 0.216019 | 0.219380 | 0.258943 | 0.233685 | 0.214826 | 0.224844 | 0.317485 | 0.384988 | 0.643049 | 0.853169 | 0.760753 | 0.639910 | 0.478029 | 0.285372 | 0.411797 | 0.363306 | 0.285460 | 0.308278 | 0.312892 | 0.255118 | 0.228240 | 0.214677 | 0.216536 | 0.224524 | 0.230233 | 0.220531 | 0.219981 | 0.214626 | 0.220975 | 0.217724 | 0.230540 | 0.232100 | 0.216053 | 0.215528 | 0.229235 | 0.271926 | 0.309633 | 0.444561 | 0.991102 | 2.200289 | 1.064167 | 0.640663 | 0.496253 | 0.539754 | 0.739794 | 1.423568 | 3.252859 | 11.106136 | 38.510615 | 150.212481 | 333.233699 | 547.061851 | 427.234380 | 415.119194 | 305.023874 | 255.176547 | 119.698421 | 57.694548 | 26.941349 | 7.142488 | 1.895046 | 1.281297 | 0.954372 | 0.659931 | 0.574075 | 0.473955 | 0.689262 | 1.332138 | 1.391416 | 2.345632 | 6.045751 | 17.239047 | 36.608655 | 48.866379 | 58.271106 | 62.459195 | 61.712615 | 41.284765 | 26.468161 | 20.191245 | 17.063810 | 10.771672 | 9.263522 | 7.271307 | 6.558178 | 4.441298 | 3.688844 | 3.272841 | 2.489263 | 2.566045 | 4.728627 | 5.443548 | 5.856799 | 6.157118 | 6.090375 | 9.469930 | 13.245324 | 12.189501 | 11.198233 | 7.095662 | 5.108470 | 6.636845 | 7.735413 | 8.020833 | 5.988790 | 6.022178 | 9.811215 | 16.322325 | 24.422162 | 23.764455 | 18.655960 | 16.519876 | 17.829990 | 20.224690 | 21.862898 | 12.334312 | 12.913809 | 19.023139 | 30.110558 | 23.338865 | 25.313903 | 24.093370 | 26.502201 | 27.638556 | 21.854818 | 15.882756 | 15.750320 | 9.467636 | 3.834005 | 2.092823 | 1.310751 | 0.940737 | 0.694451 | 0.509860 | 0.515174 | 0.574734 | 0.712375 | 0.994376 | 1.881668 | 2.842723 | 3.757854 | 3.575602 | 4.494258 | 5.821211 | 3.769395 | 2.068216 | 1.545292 | 1.542552 | 1.239405 | 0.813277 | 0.570331 | 0.467502 | 0.416699 | 0.338099 | 0.284719 | 0.320273 | 0.411747 | 0.466575 | 0.540500 | 0.541607 | 0.636425 | 0.763720 | 0.600518 | 0.558180 | 0.554117 | 0.564461 | 0.795833 | 1.045656 | 1.340970 | 1.200984 | 0.701793 | 0.438602 | 0.331474 | 0.297164 | 0.288812 | 0.269015 | 0.237480 | 0.223000 | 0.225706 | 0.244911 | 0.274229 | 0.313767 | 0.345727 | 0.447234 | 0.787150 | 1.051877 | 1.285139 | 1.222448 | 1.105527 | 0.922023 | 1.129057 | 1.234440 | 1.498676 | 1.295697 | 1.239823 | 1.181383 | 1.510130 | 1.286598 | 1.034369 | 0.837513 | 0.759226 | 0.644888 | 0.573897 | 0.437803 | 0.407415 | 0.388394 | 0.493746 | 0.534089 | 0.596641 | 0.554434 | 0.494779 | 0.476244 | 0.537736 | 0.496833 | 0.477801 | 0.398746 | 0.398586 | 0.463685 | 0.509373 | 0.412469 | 0.327846 | 0.279738 | 0.270747 | 0.256489 | 0.244552 | 0.251566 | 0.279409 | 0.323066 | 0.362207 | 0.418091 | 0.438921 | 0.539095 | 0.591905 | 0.621896 | 0.471799 | 0.457024 | 0.394834 | 0.375176 | 0.284705 | 0.236091 | 0.215819 | 0.214656 | 0.217489 | 0.240352 | 0.276222 | 0.306883 | 0.322139 | 0.280384 | 0.237401 | 0.220453 | 0.218853 | 0.242607 | 0.287794 | 0.336686 | 0.325995 | 0.356162 | 0.456600 | 0.454811 | 0.524152 | 0.599456 |
| all electrodes | 0.284030 | 0.239891 | 0.220515 | 0.218935 | 0.293475 | 0.303217 | 0.217302 | 0.215045 | 0.215518 | 0.238440 | 0.226942 | 0.232386 | 0.304448 | 0.290645 | 0.247271 | 0.218145 | 0.231113 | 0.261061 | 0.256849 | 0.214693 | 0.221033 | 0.215857 | 0.227771 | 0.233315 | 0.214848 | 0.224015 | 0.220221 | 0.216270 | 0.225002 | 0.319325 | 0.460710 | 0.473316 | 0.462082 | 0.554800 | 0.783394 | 0.768622 | 0.333147 | 0.278417 | 0.460212 | 0.710819 | 0.912700 | 0.831558 | 0.903514 | 1.248669 | 1.237385 | 1.275510 | 1.091560 | 1.014050 | 0.886735 | 0.642327 | 0.442032 | 0.331787 | 0.247016 | 0.224559 | 0.216483 | 0.215597 | 0.214928 | 0.219140 | 0.248829 | 0.316408 | 0.378688 | 0.474672 | 0.867344 | 1.632395 | 3.487179 | 8.278737 | 23.096423 | 96.661446 | 285.294632 | 445.608762 | 595.366124 | 870.619307 | 687.983651 | 524.654301 | 400.689569 | 411.321187 | 276.746951 | 102.479008 | 19.391307 | 5.082371 | 2.344372 | 1.359318 | 0.944476 | 0.843968 | 0.917519 | 1.200680 | 1.428074 | 2.068987 | 2.963754 | 5.063611 | 11.285323 | 28.929747 | 70.688042 | 217.128207 | 587.209835 | 576.477547 | 245.283130 | 59.851331 | 13.957206 | 3.814271 | 1.649927 | 0.815422 | 0.591527 | 0.465174 | 0.444408 | 0.459235 | 0.471611 | 0.375416 | 0.365201 | 0.440543 | 0.777193 | 1.270582 | 2.147361 | 3.438401 | 9.901817 | 14.812484 | 17.346116 | 12.547200 | 7.211020 | 6.688196 | 5.260933 | 2.654433 | 1.838944 | 1.656476 | 1.206275 | 0.900015 | 0.777781 | 0.679802 | 0.782357 | 0.921001 | 1.012652 | 1.064814 | 1.047586 | 0.835403 | 0.939573 | 0.923883 | 0.895488 | 0.758854 | 0.873623 | 1.219023 | 1.792850 | 1.708502 | 1.713912 | 1.735287 | 1.987746 | 1.606486 | 0.901215 | 0.528496 | 0.412005 | 0.365031 | 0.348072 | 0.323458 | 0.328334 | 0.382168 | 0.443726 | 0.509832 | 0.470027 | 0.543678 | 0.692598 | 0.691825 | 0.732096 | 0.805966 | 0.972475 | 1.220746 | 1.104516 | 0.766400 | 0.777728 | 0.809512 | 0.912517 | 0.786410 | 0.526111 | 0.362756 | 0.354212 | 0.386001 | 0.422099 | 0.456511 | 0.621088 | 1.015191 | 1.580762 | 1.612854 | 1.362203 | 1.442841 | 1.249146 | 0.850187 | 0.717699 | 0.807027 | 0.990882 | 1.258745 | 1.833456 | 2.879107 | 3.029120 | 2.394249 | 1.469473 | 1.136663 | 1.088733 | 1.055995 | 1.222208 | 1.703385 | 2.423727 | 2.845084 | 3.878551 | 3.050588 | 2.264764 | 1.673348 | 1.036923 | 0.875169 | 0.975176 | 1.036455 | 2.017380 | 4.119017 | 9.036535 | 20.240149 | 25.313722 | 30.924203 | 25.764543 | 11.289799 | 5.642569 | 4.627158 | 6.996970 | 10.149385 | 8.516214 | 7.991153 | 9.018689 | 9.669036 | 6.390867 | 3.206240 | 1.830712 | 1.879598 | 2.703271 | 3.202403 | 3.350778 | 4.561869 | 5.080399 | 4.698131 | 5.329602 | 5.245571 | 4.219547 | 3.914649 | 3.847301 | 4.445370 | 8.599808 | 18.762189 | 35.239279 | 45.119724 | 25.708504 | 22.912822 | 26.394438 | 22.760145 | 9.529558 | 8.101834 | 8.120664 | 12.926976 | 13.924972 | 12.608871 | 14.167176 | 19.689670 | 14.823721 | 9.462785 | 7.383060 | 5.920814 | 4.340194 | 1.643869 | 0.626029 | 0.391470 | 0.299794 | 0.300942 | 0.284573 | 0.257178 | 0.273104 | 0.378732 | 0.511691 | 0.570463 | 0.357530 | 0.305231 | 0.288249 | 0.292784 | 0.267099 | 0.240515 | 0.237962 | 0.278279 |

Searchlight, spatiotemporal cluster permutation test

|  | start time | stop time | peak time | peak channel | cluster p | peak Cohen's d | direction |
| --- | --- | --- | --- | --- | --- | --- | --- |
| #1 | 75 | 1195 | 155 | POz | 0.0003 | 1.695485 | positive |

O) real to emoji faces - emotion

  
|  | time window | peak latency | cluster *p* | peak Cohen's *d* |  | | | |  |
| **all electrodes** | 115 - 180 ms | 150 ms | 0.0316 | 1.3432 |  | | | |  |
 355 - 495 ms | 405 ms | 0.0206 | 0.7662 |  | | | | 605 - 775 ms | 735 ms | 0.0136 | 0.8115 |  | | | ||  |  | | | | | | | | |

Time-resolved classification, cluster permutation tests

|  |  | **left hemisphere** | | | | **right hemisphere** | | | |
|  | time window | peak latency | cluster *p* | peak Cohen's *d* | time window | peak latency | cluster *p* | peak Cohen's *d* |  |
| **anterior** |  | | | |  | | | |  |
| **central** |  | | | |  | | | |  |
| **posterior** | 115 - 180 ms | 150 ms | 0.018 | 1.4989 | 110 - 185 ms | 150 ms | 0.0204 | 1.4804 |  |
 410 - 535 ms | 440 ms | 0.0164 | 0.9326 | 365 - 475 ms | 455 ms | 0.0499 | 0.4879 | 660 - 930 ms | 735 ms | 0.0012 | 0.8956 |  | | | | 955 - 1050 ms | 995 ms | 0.0431 | 0.6147 |  | | | |

  

Time-resolved classification, Bayesian statistics

|  | -200 | -195 | -190 | -185 | -180 | -175 | -170 | -165 | -160 | -155 | -150 | -145 | -140 | -135 | -130 | -125 | -120 | -115 | -110 | -105 | -100 | -95 | -90 | -85 | -80 | -75 | -70 | -65 | -60 | -55 | -50 | -45 | -40 | -35 | -30 | -25 | -20 | -15 | -10 | -5 | 0 | 5 | 10 | 15 | 20 | 25 | 30 | 35 | 40 | 45 | 50 | 55 | 60 | 65 | 70 | 75 | 80 | 85 | 90 | 95 | 100 | 105 | 110 | 115 | 120 | 125 | 130 | 135 | 140 | 145 | 150 | 155 | 160 | 165 | 170 | 175 | 180 | 185 | 190 | 195 | 200 | 205 | 210 | 215 | 220 | 225 | 230 | 235 | 240 | 245 | 250 | 255 | 260 | 265 | 270 | 275 | 280 | 285 | 290 | 295 | 300 | 305 | 310 | 315 | 320 | 325 | 330 | 335 | 340 | 345 | 350 | 355 | 360 | 365 | 370 | 375 | 380 | 385 | 390 | 395 | 400 | 405 | 410 | 415 | 420 | 425 | 430 | 435 | 440 | 445 | 450 | 455 | 460 | 465 | 470 | 475 | 480 | 485 | 490 | 495 | 500 | 505 | 510 | 515 | 520 | 525 | 530 | 535 | 540 | 545 | 550 | 555 | 560 | 565 | 570 | 575 | 580 | 585 | 590 | 595 | 600 | 605 | 610 | 615 | 620 | 625 | 630 | 635 | 640 | 645 | 650 | 655 | 660 | 665 | 670 | 675 | 680 | 685 | 690 | 695 | 700 | 705 | 710 | 715 | 720 | 725 | 730 | 735 | 740 | 745 | 750 | 755 | 760 | 765 | 770 | 775 | 780 | 785 | 790 | 795 | 800 | 805 | 810 | 815 | 820 | 825 | 830 | 835 | 840 | 845 | 850 | 855 | 860 | 865 | 870 | 875 | 880 | 885 | 890 | 895 | 900 | 905 | 910 | 915 | 920 | 925 | 930 | 935 | 940 | 945 | 950 | 955 | 960 | 965 | 970 | 975 | 980 | 985 | 990 | 995 | 1000 | 1005 | 1010 | 1015 | 1020 | 1025 | 1030 | 1035 | 1040 | 1045 | 1050 | 1055 | 1060 | 1065 | 1070 | 1075 | 1080 | 1085 | 1090 | 1095 | 1100 | 1105 | 1110 | 1115 | 1120 | 1125 | 1130 | 1135 | 1140 | 1145 | 1150 | 1155 | 1160 | 1165 | 1170 | 1175 | 1180 | 1185 | 1190 | 1195 |
| --- | --- | --- | --- | --- | --- | --- | --- | --- | --- | --- | --- | --- | --- | --- | --- | --- | --- | --- | --- | --- | --- | --- | --- | --- | --- | --- | --- | --- | --- | --- | --- | --- | --- | --- | --- | --- | --- | --- | --- | --- | --- | --- | --- | --- | --- | --- | --- | --- | --- | --- | --- | --- | --- | --- | --- | --- | --- | --- | --- | --- | --- | --- | --- | --- | --- | --- | --- | --- | --- | --- | --- | --- | --- | --- | --- | --- | --- | --- | --- | --- | --- | --- | --- | --- | --- | --- | --- | --- | --- | --- | --- | --- | --- | --- | --- | --- | --- | --- | --- | --- | --- | --- | --- | --- | --- | --- | --- | --- | --- | --- | --- | --- | --- | --- | --- | --- | --- | --- | --- | --- | --- | --- | --- | --- | --- | --- | --- | --- | --- | --- | --- | --- | --- | --- | --- | --- | --- | --- | --- | --- | --- | --- | --- | --- | --- | --- | --- | --- | --- | --- | --- | --- | --- | --- | --- | --- | --- | --- | --- | --- | --- | --- | --- | --- | --- | --- | --- | --- | --- | --- | --- | --- | --- | --- | --- | --- | --- | --- | --- | --- | --- | --- | --- | --- | --- | --- | --- | --- | --- | --- | --- | --- | --- | --- | --- | --- | --- | --- | --- | --- | --- | --- | --- | --- | --- | --- | --- | --- | --- | --- | --- | --- | --- | --- | --- | --- | --- | --- | --- | --- | --- | --- | --- | --- | --- | --- | --- | --- | --- | --- | --- | --- | --- | --- | --- | --- | --- | --- | --- | --- | --- | --- | --- | --- | --- | --- | --- | --- | --- | --- | --- | --- | --- | --- | --- | --- | --- | --- | --- | --- | --- | --- | --- | --- | --- | --- | --- | --- | --- | --- | --- | --- | --- | --- | --- | --- | --- | --- | --- | --- |
| left anterior | 2.355516 | 1.477234 | 0.792604 | 0.444903 | 0.436573 | 0.231216 | 0.259521 | 0.615972 | 0.848235 | 0.561142 | 0.277652 | 0.263065 | 0.218492 | 0.254843 | 0.573871 | 0.844207 | 0.553493 | 0.267370 | 0.238857 | 0.219182 | 0.210824 | 0.245465 | 0.288499 | 0.278340 | 0.271303 | 0.509413 | 0.693556 | 0.713990 | 0.296218 | 0.215082 | 0.224837 | 0.210881 | 0.219504 | 0.242997 | 0.226462 | 0.211823 | 0.214504 | 0.240002 | 0.220095 | 0.264710 | 0.354225 | 0.376925 | 0.270980 | 0.212288 | 0.212717 | 0.213622 | 0.211354 | 0.210823 | 0.217595 | 0.257692 | 0.270636 | 0.317997 | 0.365247 | 0.400606 | 0.367479 | 0.270655 | 0.220403 | 0.217627 | 0.214970 | 0.218519 | 0.216293 | 0.217735 | 0.234058 | 0.231519 | 0.214986 | 0.216440 | 0.262668 | 0.572108 | 1.466410 | 2.989475 | 3.637534 | 2.406888 | 0.910751 | 0.784509 | 0.450152 | 0.339253 | 0.293307 | 0.257449 | 0.292509 | 0.378277 | 0.254003 | 0.228757 | 0.212006 | 0.231967 | 0.325107 | 1.224251 | 7.098443 | 1.846277 | 1.164893 | 0.555032 | 0.374189 | 0.228461 | 0.213941 | 0.262049 | 0.324747 | 0.312819 | 0.243772 | 0.219515 | 0.211028 | 0.210841 | 0.211865 | 0.210948 | 0.212556 | 0.230020 | 0.284210 | 0.322537 | 0.311496 | 0.263078 | 0.236975 | 0.222170 | 0.216083 | 0.216966 | 0.243544 | 0.271057 | 0.320643 | 0.468361 | 0.703860 | 0.956544 | 1.096781 | 1.071073 | 0.845426 | 0.627627 | 0.526295 | 0.554863 | 0.547928 | 0.446783 | 0.353913 | 0.296793 | 0.323800 | 0.305203 | 0.270020 | 0.247972 | 0.240910 | 0.233418 | 0.221056 | 0.210846 | 0.214692 | 0.218399 | 0.224600 | 0.247493 | 0.302529 | 0.312060 | 0.294847 | 0.304135 | 0.312930 | 0.310015 | 0.308151 | 0.282010 | 0.270006 | 0.305103 | 0.298243 | 0.295367 | 0.293719 | 0.314484 | 0.361577 | 0.366081 | 0.347786 | 0.320601 | 0.340288 | 0.352781 | 0.303019 | 0.275375 | 0.247606 | 0.218447 | 0.219035 | 0.219060 | 0.233514 | 0.247724 | 0.252241 | 0.285126 | 0.351441 | 0.419318 | 0.406385 | 0.302114 | 0.239817 | 0.218233 | 0.216574 | 0.211455 | 0.212402 | 0.212416 | 0.221170 | 0.233621 | 0.245142 | 0.222800 | 0.226583 | 0.212357 | 0.212335 | 0.226973 | 0.245461 | 0.231567 | 0.214360 | 0.211108 | 0.210922 | 0.215406 | 0.232092 | 0.308460 | 0.337755 | 0.298024 | 0.281435 | 0.293201 | 0.294643 | 0.302053 | 0.287276 | 0.317203 | 0.518902 | 0.761830 | 1.119939 | 0.871483 | 0.880371 | 0.770951 | 0.758274 | 0.585389 | 0.457483 | 0.341844 | 0.401451 | 0.336016 | 0.309062 | 0.280192 | 0.257264 | 0.240105 | 0.236413 | 0.220703 | 0.220039 | 0.216718 | 0.218718 | 0.213468 | 0.218795 | 0.239850 | 0.249857 | 0.268558 | 0.254053 | 0.223618 | 0.227984 | 0.237735 | 0.220259 | 0.237320 | 0.241786 | 0.255917 | 0.319837 | 0.364063 | 0.313027 | 0.294321 | 0.238980 | 0.216130 | 0.210886 | 0.217459 | 0.247821 | 0.257058 | 0.230878 | 0.213807 | 0.212632 | 0.227501 | 0.249445 | 0.276672 | 0.278066 | 0.231818 | 0.214356 | 0.211740 | 0.229586 | 0.274299 | 0.330659 | 0.357266 | 0.318960 | 0.278828 | 0.248702 | 0.225461 | 0.211503 | 0.216301 | 0.233971 | 0.239496 | 0.225966 | 0.212676 | 0.211328 | 0.211189 | 0.210862 | 0.211563 | 0.211096 | 0.212256 | 0.218248 | 0.221637 |
| right anterior | 0.238741 | 0.296808 | 0.309855 | 0.366912 | 0.744189 | 0.746773 | 1.629901 | 1.520489 | 1.266619 | 2.077336 | 1.379524 | 0.323568 | 0.234703 | 0.219507 | 0.225178 | 0.226821 | 0.331058 | 0.521397 | 0.263628 | 0.214073 | 0.220448 | 0.228558 | 0.221665 | 0.226805 | 0.248048 | 0.221179 | 0.212515 | 0.282147 | 0.368148 | 0.368444 | 0.368652 | 0.357603 | 0.443542 | 0.731339 | 0.633551 | 0.708193 | 0.572997 | 0.431638 | 0.437019 | 0.315037 | 0.228815 | 0.212933 | 0.210859 | 0.212773 | 0.218178 | 0.238049 | 0.266582 | 0.305284 | 0.490954 | 0.828726 | 0.923899 | 0.779925 | 0.566552 | 0.432852 | 0.443319 | 0.551777 | 0.546399 | 0.537150 | 0.692896 | 0.855203 | 1.079489 | 0.864189 | 1.046376 | 1.764927 | 6.519367 | 52.768629 | 212.311633 | 553.711895 | 1743.678716 | 1376.079048 | 996.711863 | 303.378774 | 43.498890 | 8.913084 | 3.159041 | 1.068738 | 0.414532 | 0.228090 | 0.224752 | 0.259361 | 0.292996 | 0.260501 | 0.215785 | 0.210834 | 0.211429 | 0.210861 | 0.216127 | 0.216461 | 0.225974 | 0.213015 | 0.211094 | 0.216283 | 0.224361 | 0.210839 | 0.216918 | 0.216434 | 0.224082 | 0.257083 | 0.265964 | 0.257073 | 0.291031 | 0.267314 | 0.285077 | 0.260181 | 0.214611 | 0.211244 | 0.218378 | 0.221587 | 0.228810 | 0.279958 | 0.338226 | 0.306260 | 0.295220 | 0.220933 | 0.226176 | 0.228147 | 0.217554 | 0.211292 | 0.217056 | 0.255607 | 0.367789 | 0.358960 | 0.308158 | 0.339917 | 0.292164 | 0.227408 | 0.215101 | 0.234551 | 0.233744 | 0.218240 | 0.228150 | 0.220020 | 0.214841 | 0.213182 | 0.224403 | 0.276972 | 0.441346 | 0.635919 | 0.779775 | 0.934153 | 1.237874 | 0.937868 | 0.665863 | 0.681113 | 0.643564 | 0.518888 | 0.506175 | 0.515432 | 0.609330 | 0.553832 | 0.368264 | 0.371385 | 0.478871 | 0.532335 | 0.431824 | 0.441454 | 0.599622 | 0.934014 | 0.754131 | 0.512950 | 0.387467 | 0.360135 | 0.318192 | 0.263396 | 0.242426 | 0.242952 | 0.226450 | 0.217140 | 0.212553 | 0.210854 | 0.210848 | 0.212979 | 0.214741 | 0.210852 | 0.211359 | 0.211832 | 0.217465 | 0.232740 | 0.248451 | 0.286740 | 0.381330 | 0.372788 | 0.417701 | 0.370287 | 0.361324 | 0.384545 | 0.371974 | 0.313765 | 0.290863 | 0.241682 | 0.228080 | 0.222092 | 0.213428 | 0.211728 | 0.231005 | 0.266484 | 0.326677 | 0.297879 | 0.268564 | 0.274502 | 0.288307 | 0.266141 | 0.256738 | 0.280021 | 0.357595 | 0.465237 | 0.568444 | 0.625728 | 0.484247 | 0.362116 | 0.340791 | 0.407913 | 0.359130 | 0.275272 | 0.233788 | 0.228133 | 0.243588 | 0.229737 | 0.214455 | 0.215513 | 0.219526 | 0.224265 | 0.229548 | 0.228580 | 0.246383 | 0.248767 | 0.235367 | 0.222956 | 0.217413 | 0.220937 | 0.232272 | 0.229684 | 0.245034 | 0.280460 | 0.358200 | 0.433603 | 0.383658 | 0.314765 | 0.332605 | 0.357002 | 0.320442 | 0.292176 | 0.266508 | 0.293840 | 0.324119 | 0.333953 | 0.314350 | 0.344260 | 0.360768 | 0.390704 | 0.356113 | 0.306998 | 0.258295 | 0.267313 | 0.252013 | 0.301016 | 0.421409 | 0.636947 | 0.858406 | 1.080342 | 1.394527 | 2.022046 | 1.315295 | 0.862956 | 0.779813 | 0.854141 | 0.687391 | 0.384257 | 0.321748 | 0.322748 | 0.301110 | 0.315610 | 0.291726 | 0.321973 | 0.329416 | 0.362687 | 0.484041 | 0.535931 | 0.392349 | 0.355769 |
| left central | 0.368923 | 0.328640 | 0.301505 | 0.387364 | 0.709251 | 0.624165 | 0.523039 | 0.362786 | 0.253420 | 0.255241 | 0.229882 | 0.213822 | 0.215793 | 0.210848 | 0.214013 | 0.211256 | 0.213328 | 0.215069 | 0.213010 | 0.214648 | 0.222590 | 0.215489 | 0.224923 | 0.227367 | 0.214260 | 0.269834 | 0.284799 | 0.239986 | 0.288561 | 0.466427 | 0.624795 | 1.601140 | 2.491981 | 4.281992 | 3.062660 | 2.671481 | 1.336680 | 1.090497 | 0.745231 | 0.260845 | 0.213131 | 0.213016 | 0.230133 | 0.251246 | 0.242137 | 0.211792 | 0.210936 | 0.212307 | 0.210913 | 0.261291 | 0.343471 | 0.679642 | 0.819882 | 0.706328 | 0.990682 | 0.846799 | 0.468541 | 0.350835 | 0.287574 | 0.267639 | 0.268165 | 0.239638 | 0.222227 | 0.214632 | 0.218110 | 0.399886 | 2.721017 | 41.351161 | 743.888623 | 4339.116480 | 11335.981998 | 4596.563346 | 1246.706873 | 753.026465 | 112.666234 | 7.803081 | 0.492999 | 0.249589 | 0.219299 | 0.210866 | 0.218565 | 0.260049 | 0.262582 | 0.223483 | 0.260374 | 0.301361 | 0.411607 | 0.395363 | 0.306240 | 0.298625 | 0.410767 | 0.281730 | 0.224889 | 0.214378 | 0.238784 | 0.416272 | 0.782375 | 1.822537 | 3.033280 | 1.868062 | 1.084404 | 0.473590 | 0.238356 | 0.215499 | 0.236812 | 0.280989 | 0.343504 | 0.483936 | 0.645777 | 0.468375 | 0.323096 | 0.305622 | 0.232209 | 0.212209 | 0.283292 | 0.465905 | 0.405131 | 0.393811 | 0.508642 | 0.690011 | 1.096006 | 1.536741 | 1.884141 | 2.935691 | 4.710923 | 3.726968 | 2.257116 | 1.547493 | 1.162049 | 0.826016 | 0.531140 | 0.368027 | 0.258411 | 0.222913 | 0.211190 | 0.213817 | 0.210814 | 0.212715 | 0.225005 | 0.265995 | 0.286412 | 0.302571 | 0.316401 | 0.314889 | 0.318771 | 0.279701 | 0.253381 | 0.223975 | 0.211968 | 0.211153 | 0.213233 | 0.231866 | 0.255328 | 0.303136 | 0.257548 | 0.227657 | 0.226272 | 0.242422 | 0.228065 | 0.210896 | 0.247638 | 0.250196 | 0.232526 | 0.256176 | 0.392741 | 0.469108 | 0.557675 | 0.730975 | 0.948075 | 1.253209 | 0.932605 | 0.670801 | 0.586319 | 0.474238 | 0.299132 | 0.293406 | 0.287411 | 0.287784 | 0.264403 | 0.231125 | 0.221954 | 0.246851 | 0.246367 | 0.237476 | 0.231678 | 0.231633 | 0.244741 | 0.236858 | 0.222428 | 0.221072 | 0.228021 | 0.227009 | 0.215767 | 0.210900 | 0.210900 | 0.211084 | 0.215036 | 0.222619 | 0.217883 | 0.210806 | 0.214063 | 0.221545 | 0.221801 | 0.222013 | 0.221321 | 0.216973 | 0.212070 | 0.216044 | 0.219881 | 0.257869 | 0.379852 | 0.685751 | 1.483196 | 1.849619 | 1.316584 | 0.944093 | 0.579316 | 0.398388 | 0.299096 | 0.238790 | 0.224229 | 0.228874 | 0.225346 | 0.227679 | 0.221272 | 0.217072 | 0.216224 | 0.214510 | 0.214841 | 0.219325 | 0.221030 | 0.227630 | 0.236538 | 0.252586 | 0.295377 | 0.328144 | 0.336601 | 0.384026 | 0.467886 | 0.567286 | 0.578283 | 0.532129 | 0.525512 | 0.542097 | 0.570958 | 0.501045 | 0.453368 | 0.467210 | 0.466579 | 0.455250 | 0.450370 | 0.448204 | 0.448253 | 0.430447 | 0.388413 | 0.325106 | 0.285206 | 0.277341 | 0.255704 | 0.237882 | 0.234761 | 0.219731 | 0.217569 | 0.224861 | 0.231801 | 0.240207 | 0.269038 | 0.286673 | 0.384385 | 0.541390 | 0.556889 | 0.487941 | 0.502713 | 0.526688 | 0.571435 | 0.395544 | 0.314035 | 0.320136 | 0.346342 | 0.308889 |
| right central | 0.361201 | 0.408508 | 0.344431 | 0.278445 | 0.254618 | 0.216659 | 0.219121 | 0.252809 | 0.255630 | 0.244378 | 0.241633 | 0.239078 | 0.213875 | 0.255920 | 0.230088 | 0.224006 | 0.315460 | 0.258345 | 0.226633 | 0.224234 | 0.221505 | 0.215769 | 0.383901 | 0.736027 | 0.566602 | 0.601704 | 0.378327 | 0.293392 | 0.276252 | 0.327369 | 0.467710 | 1.043404 | 1.184935 | 2.144681 | 2.308158 | 0.957212 | 0.850275 | 0.434623 | 0.251014 | 0.239541 | 0.281233 | 0.267888 | 0.267002 | 0.221118 | 0.219794 | 0.224166 | 0.212341 | 0.226074 | 0.256463 | 0.286467 | 0.236302 | 0.212961 | 0.212145 | 0.213287 | 0.235772 | 0.302723 | 0.461031 | 0.839280 | 1.860694 | 2.035427 | 3.978122 | 1.119750 | 0.326442 | 0.213138 | 0.284012 | 1.134896 | 5.436366 | 36.555246 | 140.718187 | 644.135035 | 703.657813 | 642.443113 | 163.043954 | 62.907095 | 7.479801 | 2.357925 | 0.732926 | 0.260675 | 0.212612 | 0.264610 | 0.354916 | 0.321312 | 0.361943 | 0.338811 | 0.308941 | 0.292948 | 0.315285 | 0.271465 | 0.241204 | 0.220426 | 0.215378 | 0.211657 | 0.245757 | 0.351679 | 0.432641 | 0.331758 | 0.346705 | 0.344115 | 0.374641 | 0.335696 | 0.323920 | 0.274766 | 0.283398 | 0.263737 | 0.428429 | 0.638603 | 0.859815 | 0.882171 | 1.027011 | 1.461309 | 2.664242 | 1.587290 | 2.478297 | 2.530538 | 2.195375 | 1.591428 | 0.847314 | 0.652777 | 0.666609 | 0.587089 | 0.619098 | 0.801382 | 1.185466 | 2.729823 | 3.073837 | 2.491915 | 1.793315 | 1.546766 | 1.364036 | 0.869678 | 0.544553 | 0.498318 | 0.400701 | 0.347692 | 0.231970 | 0.217796 | 0.221833 | 0.227655 | 0.240259 | 0.243232 | 0.232093 | 0.255966 | 0.223626 | 0.213524 | 0.210829 | 0.213062 | 0.211188 | 0.210896 | 0.212513 | 0.211036 | 0.210831 | 0.212444 | 0.216374 | 0.212431 | 0.214818 | 0.225076 | 0.222092 | 0.229374 | 0.234423 | 0.231869 | 0.271258 | 0.345218 | 0.322915 | 0.345542 | 0.345242 | 0.260033 | 0.261632 | 0.282090 | 0.238202 | 0.244634 | 0.271101 | 0.265156 | 0.379917 | 0.320720 | 0.273689 | 0.248534 | 0.229396 | 0.223131 | 0.233173 | 0.217957 | 0.227953 | 0.217441 | 0.225352 | 0.217437 | 0.212261 | 0.212269 | 0.211298 | 0.212099 | 0.212018 | 0.211161 | 0.213618 | 0.210867 | 0.211051 | 0.211267 | 0.222441 | 0.225757 | 0.249593 | 0.321212 | 0.337369 | 0.342726 | 0.395639 | 0.474879 | 0.430545 | 0.304731 | 0.241020 | 0.211841 | 0.212240 | 0.221799 | 0.223882 | 0.216161 | 0.210880 | 0.212065 | 0.213147 | 0.228064 | 0.241707 | 0.291634 | 0.339136 | 0.424659 | 0.435800 | 0.315255 | 0.241408 | 0.223678 | 0.211474 | 0.239928 | 0.315141 | 0.420735 | 0.462826 | 0.525039 | 0.997507 | 1.014429 | 1.011218 | 0.508493 | 0.363699 | 0.265184 | 0.239910 | 0.216247 | 0.212953 | 0.220982 | 0.234327 | 0.277605 | 0.288902 | 0.281738 | 0.348605 | 0.443008 | 0.306673 | 0.255130 | 0.220445 | 0.231656 | 0.268856 | 0.246147 | 0.220784 | 0.217899 | 0.221478 | 0.224070 | 0.230657 | 0.233903 | 0.229612 | 0.241161 | 0.238130 | 0.239287 | 0.261271 | 0.258428 | 0.234801 | 0.226466 | 0.240731 | 0.297157 | 0.348944 | 0.408416 | 0.384751 | 0.422995 | 0.459246 | 0.401287 | 0.287458 | 0.236251 | 0.215017 | 0.210813 | 0.212485 | 0.215509 | 0.222888 | 0.212491 |
| left posterior | 0.223336 | 0.211631 | 0.213429 | 0.273217 | 0.374882 | 0.365649 | 0.405744 | 0.240106 | 0.234823 | 0.319128 | 0.470641 | 1.234012 | 1.752271 | 0.685181 | 0.231460 | 0.217614 | 0.216251 | 0.211605 | 0.230671 | 0.262789 | 0.308164 | 0.348655 | 0.271554 | 0.236693 | 0.216569 | 0.213997 | 0.210874 | 0.212055 | 0.237865 | 0.327297 | 0.297224 | 0.278576 | 0.684027 | 0.657749 | 0.460377 | 0.280377 | 0.237835 | 0.219638 | 0.212701 | 0.243713 | 0.269987 | 0.354656 | 0.401964 | 0.517355 | 0.481582 | 0.330824 | 0.249680 | 0.253663 | 0.268941 | 0.290862 | 0.288390 | 0.278414 | 0.336882 | 0.434748 | 0.688299 | 0.923895 | 0.939460 | 1.409045 | 1.859231 | 2.491508 | 1.794832 | 0.270766 | 0.348818 | 1.803676 | 19.204139 | 457.548407 | 10294.084580 | 20560.685770 | 26647.647906 | 55595.489474 | 153177.670081 | 169175.091407 | 27794.807738 | 2459.667217 | 191.352227 | 22.824269 | 2.443818 | 0.602933 | 0.289816 | 0.215633 | 0.210884 | 0.214540 | 0.257502 | 0.263035 | 0.237921 | 0.223267 | 0.211145 | 0.223255 | 0.243480 | 0.250066 | 0.260753 | 0.292925 | 0.388741 | 0.662616 | 0.882500 | 1.716444 | 2.539901 | 2.617138 | 1.481962 | 1.388871 | 1.024148 | 1.061607 | 0.646245 | 0.538516 | 0.498364 | 0.438562 | 0.285793 | 0.252321 | 0.220491 | 0.221843 | 0.233794 | 0.235524 | 0.252392 | 0.318299 | 0.310592 | 0.340878 | 0.295011 | 0.261468 | 0.269804 | 0.328796 | 0.368176 | 0.626685 | 1.350616 | 4.051408 | 20.057814 | 81.025805 | 142.878685 | 463.057205 | 274.215832 | 48.136827 | 13.566000 | 4.169045 | 2.427428 | 2.396570 | 2.053938 | 1.809015 | 2.833149 | 4.871648 | 7.467447 | 7.415709 | 4.979734 | 5.214576 | 3.917442 | 3.711044 | 4.966417 | 3.929237 | 3.026547 | 2.495850 | 1.272317 | 1.499977 | 1.498719 | 1.229651 | 1.336529 | 1.266281 | 1.349795 | 3.065344 | 2.915848 | 1.942158 | 1.671100 | 1.266501 | 1.188503 | 0.895957 | 0.588320 | 0.541108 | 0.557893 | 0.503375 | 0.493025 | 0.529072 | 0.767432 | 0.897109 | 1.008138 | 1.080616 | 1.444060 | 3.000091 | 5.670379 | 8.583701 | 12.672690 | 15.907642 | 13.207376 | 8.428222 | 4.899932 | 7.763977 | 11.377981 | 19.560331 | 31.765911 | 83.441822 | 174.016437 | 179.646610 | 90.685803 | 44.841801 | 25.811236 | 13.314374 | 10.372092 | 7.872405 | 15.107663 | 23.722029 | 71.642207 | 77.925847 | 128.618196 | 176.229429 | 294.904194 | 307.452113 | 135.964203 | 65.083678 | 44.706629 | 30.002869 | 21.531394 | 11.666481 | 6.415641 | 4.907722 | 5.463171 | 4.534621 | 3.117135 | 2.729893 | 3.167451 | 4.884199 | 8.063102 | 8.676368 | 8.970137 | 9.684332 | 10.410603 | 7.516895 | 5.040891 | 3.671999 | 2.332599 | 1.617889 | 1.468134 | 1.200158 | 1.014094 | 1.089824 | 1.243997 | 1.654140 | 1.813750 | 1.901201 | 2.117000 | 2.833995 | 3.590490 | 3.979160 | 4.373373 | 8.305205 | 12.659865 | 10.911608 | 8.858646 | 5.189073 | 5.860429 | 6.708120 | 3.451745 | 2.697153 | 2.060844 | 1.736088 | 1.398683 | 0.703987 | 0.407662 | 0.301803 | 0.244021 | 0.221880 | 0.222810 | 0.229975 | 0.260036 | 0.336818 | 0.435194 | 0.491073 | 0.671267 | 0.740118 | 0.654933 | 0.536580 | 0.413497 | 0.368591 | 0.331011 | 0.334894 | 0.351807 | 0.392531 | 0.426557 | 0.446733 | 0.425727 | 0.524155 | 0.545002 | 0.507141 | 0.460180 | 0.464545 |
| right posterior | 0.277409 | 0.253417 | 0.243006 | 0.247822 | 0.290575 | 0.400361 | 0.399184 | 0.452627 | 0.367709 | 0.302842 | 0.256147 | 0.228414 | 0.215227 | 0.211113 | 0.216778 | 0.217500 | 0.214267 | 0.210935 | 0.211492 | 0.231636 | 0.351357 | 0.499878 | 0.430748 | 0.364392 | 0.329051 | 0.376832 | 0.363020 | 0.325392 | 0.283866 | 0.254110 | 0.257331 | 0.313911 | 0.241579 | 0.213862 | 0.211220 | 0.218842 | 0.222067 | 0.210846 | 0.213324 | 0.212491 | 0.228239 | 0.219586 | 0.212167 | 0.210927 | 0.211092 | 0.217286 | 0.255513 | 0.330554 | 0.313823 | 0.253043 | 0.217086 | 0.211588 | 0.210991 | 0.210836 | 0.211373 | 0.210916 | 0.227311 | 0.276855 | 0.289381 | 0.314289 | 0.523005 | 0.978284 | 1.969107 | 5.886532 | 37.996853 | 622.956637 | 3107.563329 | 7659.799232 | 26852.971649 | 67849.102146 | 125862.294945 | 78581.299163 | 32369.629427 | 9389.283984 | 2706.435486 | 354.180371 | 42.154176 | 2.837732 | 0.465888 | 0.225772 | 0.219316 | 0.291755 | 0.397281 | 0.382627 | 0.395093 | 0.366156 | 0.320423 | 0.272801 | 0.262420 | 0.220639 | 0.213833 | 0.211951 | 0.214799 | 0.285595 | 0.426201 | 1.054358 | 1.181379 | 1.412645 | 1.956134 | 1.652862 | 1.373836 | 0.951826 | 0.658110 | 0.580145 | 0.473499 | 0.510202 | 0.514730 | 0.408358 | 0.415903 | 0.403082 | 0.418656 | 0.648290 | 1.062289 | 1.752011 | 2.378835 | 1.985902 | 1.843503 | 2.085795 | 2.226264 | 2.325253 | 3.299571 | 3.418566 | 3.969246 | 5.378645 | 4.164958 | 4.989973 | 4.150644 | 2.938686 | 2.865203 | 3.126155 | 2.742654 | 2.438782 | 1.843474 | 1.822817 | 1.525561 | 1.347703 | 0.986841 | 0.910343 | 0.966049 | 0.985693 | 1.212498 | 1.477029 | 1.773244 | 1.927796 | 1.281743 | 1.012174 | 0.853012 | 0.557322 | 0.412017 | 0.324757 | 0.282795 | 0.268979 | 0.257349 | 0.254956 | 0.252904 | 0.269736 | 0.273578 | 0.312431 | 0.326088 | 0.351062 | 0.419690 | 0.597607 | 0.676217 | 0.783704 | 0.756801 | 0.712578 | 0.741085 | 0.647128 | 0.714896 | 0.745235 | 0.636641 | 0.463046 | 0.383989 | 0.352156 | 0.371455 | 0.366714 | 0.397163 | 0.513754 | 0.537576 | 0.720500 | 0.862966 | 0.956781 | 0.885482 | 0.690687 | 0.628587 | 0.753239 | 0.879722 | 1.178163 | 1.363005 | 1.470836 | 1.794890 | 2.234511 | 4.173605 | 7.624172 | 10.189432 | 9.925681 | 8.125416 | 5.599384 | 5.129232 | 2.893220 | 1.934033 | 1.396458 | 1.022778 | 0.878029 | 0.846828 | 0.819118 | 0.700130 | 0.659194 | 0.869863 | 1.141068 | 1.581778 | 2.143475 | 1.848252 | 1.594373 | 1.192223 | 0.725419 | 0.497637 | 0.392173 | 0.328867 | 0.291796 | 0.300567 | 0.315737 | 0.396603 | 0.492998 | 0.608506 | 0.707404 | 0.876162 | 0.860416 | 0.810744 | 0.624679 | 0.602274 | 0.529149 | 0.515411 | 0.468908 | 0.319481 | 0.280410 | 0.245102 | 0.221755 | 0.213546 | 0.210855 | 0.211899 | 0.223539 | 0.252570 | 0.289966 | 0.307189 | 0.306723 | 0.284976 | 0.252543 | 0.243722 | 0.233864 | 0.217475 | 0.212384 | 0.211624 | 0.213726 | 0.222786 | 0.257859 | 0.321218 | 0.347206 | 0.323161 | 0.251725 | 0.226868 | 0.214940 | 0.210994 | 0.211265 | 0.212582 | 0.232885 | 0.261042 | 0.268267 | 0.284291 | 0.268937 | 0.227727 | 0.214699 | 0.211925 | 0.214128 | 0.216263 | 0.224540 | 0.234982 | 0.228612 | 0.235583 | 0.253350 |
| all electrodes | 2.014669 | 1.471765 | 0.640998 | 0.641162 | 0.710377 | 0.465786 | 0.378811 | 0.367786 | 0.397781 | 0.660301 | 0.806810 | 0.751364 | 0.711918 | 0.468881 | 0.358431 | 0.227549 | 0.220828 | 0.248444 | 0.225882 | 0.212470 | 0.212571 | 0.222341 | 0.339920 | 0.613601 | 0.495983 | 0.313717 | 0.213963 | 0.225165 | 0.249829 | 0.263928 | 0.304349 | 0.267976 | 0.228629 | 0.210973 | 0.220699 | 0.221637 | 0.222671 | 0.226874 | 0.228838 | 0.233806 | 0.276584 | 0.291845 | 0.271567 | 0.223378 | 0.210843 | 0.220459 | 0.226082 | 0.238060 | 0.221781 | 0.212515 | 0.216941 | 0.255836 | 0.322846 | 0.570618 | 0.865389 | 0.564434 | 0.323713 | 0.222595 | 0.218446 | 0.211805 | 0.217381 | 0.284344 | 0.503849 | 1.419330 | 18.734946 | 784.328963 | 4328.860873 | 12286.438798 | 17259.014747 | 23232.855109 | 28527.374817 | 13546.529663 | 3991.125121 | 912.689307 | 160.687661 | 31.481442 | 5.214890 | 0.749710 | 0.320171 | 0.236282 | 0.218638 | 0.220926 | 0.231032 | 0.252843 | 0.304817 | 0.320130 | 0.300902 | 0.259721 | 0.246484 | 0.253601 | 0.270887 | 0.368004 | 0.632883 | 1.455328 | 3.962411 | 8.250140 | 5.361407 | 3.474308 | 1.915270 | 1.494015 | 1.397262 | 1.364640 | 1.181506 | 1.292544 | 1.293974 | 1.895396 | 2.252548 | 1.310089 | 1.061432 | 0.917088 | 1.022948 | 1.383105 | 1.317344 | 1.752524 | 5.386066 | 10.129537 | 18.400712 | 19.350689 | 22.886694 | 20.790990 | 21.913783 | 41.929321 | 32.941696 | 34.776987 | 45.335003 | 47.940542 | 43.477232 | 32.717991 | 8.400910 | 6.346104 | 3.811867 | 3.471271 | 3.067659 | 3.444975 | 2.985077 | 2.918829 | 3.880444 | 2.994377 | 2.214636 | 1.455627 | 0.975278 | 0.888088 | 1.176529 | 0.806491 | 0.460116 | 0.347198 | 0.311573 | 0.305541 | 0.297285 | 0.289052 | 0.307773 | 0.360127 | 0.345870 | 0.311462 | 0.273645 | 0.246446 | 0.235556 | 0.234033 | 0.264962 | 0.375875 | 0.687747 | 1.316492 | 2.658567 | 4.893813 | 9.392009 | 12.567877 | 20.549729 | 21.139441 | 17.007521 | 8.286115 | 7.016429 | 9.560930 | 9.802521 | 3.812739 | 2.746969 | 2.290453 | 4.384593 | 3.641917 | 2.874914 | 1.790754 | 3.087423 | 7.070628 | 17.684286 | 40.592116 | 54.522986 | 46.284115 | 72.031355 | 69.474220 | 56.603892 | 31.333842 | 12.373186 | 9.792813 | 8.844606 | 4.251572 | 2.397025 | 1.346401 | 0.811950 | 0.639728 | 0.501695 | 0.453067 | 0.557577 | 0.645546 | 0.884567 | 1.106608 | 1.308826 | 1.247429 | 1.076528 | 0.760541 | 0.865554 | 0.985288 | 1.186923 | 1.091940 | 0.975908 | 1.189017 | 1.224110 | 0.904165 | 0.790784 | 0.515891 | 0.415159 | 0.408948 | 0.356330 | 0.360113 | 0.548980 | 0.807872 | 1.232015 | 1.373160 | 1.318452 | 1.416085 | 1.162217 | 0.751765 | 0.528207 | 0.520002 | 0.587350 | 0.678059 | 0.727756 | 0.886360 | 0.874056 | 0.880236 | 0.900161 | 0.842819 | 0.599443 | 0.442354 | 0.454735 | 0.483321 | 0.521549 | 0.624965 | 0.803174 | 1.345769 | 1.592085 | 1.220260 | 1.172287 | 1.028807 | 0.749048 | 0.536655 | 0.395574 | 0.332527 | 0.349591 | 0.350390 | 0.373779 | 0.437692 | 0.636993 | 0.829084 | 1.036019 | 0.916122 | 0.649136 | 0.415707 | 0.301927 | 0.251832 | 0.239933 | 0.241522 | 0.248506 | 0.262963 | 0.322749 | 0.389004 | 0.397271 | 0.410633 | 0.453213 | 0.491649 | 0.547043 | 0.542794 |

Searchlight, spatiotemporal cluster permutation test

|  | start time | stop time | peak time | peak channel | cluster p | peak Cohen's d | direction |
| --- | --- | --- | --- | --- | --- | --- | --- |
| #1 | 90 | 200 | 160 | P8 | 0.0079 | 1.762895 | positive |
| #2 | 265 | 1065 | 430 | O2 | 0.0011 | 1.15742 | positive |

P) real to emoji faces - happy vs angry

  
|  | time window | peak latency | cluster *p* | peak Cohen's *d* |  | | | |
| **all electrodes** | 145 - 230 ms | 165 ms | 0.0392 | 1.0239 |  | | | |
|  | | | | | | | | |

Time-resolved classification, cluster permutation tests

|  | **left hemisphere** | | | | **right hemisphere** | | | |
|  | time window | peak latency | cluster *p* | peak Cohen's *d* | time window | peak latency | cluster *p* | peak Cohen's *d* |
| **anterior** |  | | | |  | | | |
| **central** |  | | | |  | | | |
| **posterior** | 140 - 215 ms | 160 ms | 0.0393 | 1.1014 | 415 - 530 ms | 495 ms | 0.0381 | 0.75 |
 410 - 530 ms | 435 ms | 0.0318 | 0.6273 |  | | | |

  

Time-resolved classification, Bayesian statistics

|  | -200 | -195 | -190 | -185 | -180 | -175 | -170 | -165 | -160 | -155 | -150 | -145 | -140 | -135 | -130 | -125 | -120 | -115 | -110 | -105 | -100 | -95 | -90 | -85 | -80 | -75 | -70 | -65 | -60 | -55 | -50 | -45 | -40 | -35 | -30 | -25 | -20 | -15 | -10 | -5 | 0 | 5 | 10 | 15 | 20 | 25 | 30 | 35 | 40 | 45 | 50 | 55 | 60 | 65 | 70 | 75 | 80 | 85 | 90 | 95 | 100 | 105 | 110 | 115 | 120 | 125 | 130 | 135 | 140 | 145 | 150 | 155 | 160 | 165 | 170 | 175 | 180 | 185 | 190 | 195 | 200 | 205 | 210 | 215 | 220 | 225 | 230 | 235 | 240 | 245 | 250 | 255 | 260 | 265 | 270 | 275 | 280 | 285 | 290 | 295 | 300 | 305 | 310 | 315 | 320 | 325 | 330 | 335 | 340 | 345 | 350 | 355 | 360 | 365 | 370 | 375 | 380 | 385 | 390 | 395 | 400 | 405 | 410 | 415 | 420 | 425 | 430 | 435 | 440 | 445 | 450 | 455 | 460 | 465 | 470 | 475 | 480 | 485 | 490 | 495 | 500 | 505 | 510 | 515 | 520 | 525 | 530 | 535 | 540 | 545 | 550 | 555 | 560 | 565 | 570 | 575 | 580 | 585 | 590 | 595 | 600 | 605 | 610 | 615 | 620 | 625 | 630 | 635 | 640 | 645 | 650 | 655 | 660 | 665 | 670 | 675 | 680 | 685 | 690 | 695 | 700 | 705 | 710 | 715 | 720 | 725 | 730 | 735 | 740 | 745 | 750 | 755 | 760 | 765 | 770 | 775 | 780 | 785 | 790 | 795 | 800 | 805 | 810 | 815 | 820 | 825 | 830 | 835 | 840 | 845 | 850 | 855 | 860 | 865 | 870 | 875 | 880 | 885 | 890 | 895 | 900 | 905 | 910 | 915 | 920 | 925 | 930 | 935 | 940 | 945 | 950 | 955 | 960 | 965 | 970 | 975 | 980 | 985 | 990 | 995 | 1000 | 1005 | 1010 | 1015 | 1020 | 1025 | 1030 | 1035 | 1040 | 1045 | 1050 | 1055 | 1060 | 1065 | 1070 | 1075 | 1080 | 1085 | 1090 | 1095 | 1100 | 1105 | 1110 | 1115 | 1120 | 1125 | 1130 | 1135 | 1140 | 1145 | 1150 | 1155 | 1160 | 1165 | 1170 | 1175 | 1180 | 1185 | 1190 | 1195 |
| --- | --- | --- | --- | --- | --- | --- | --- | --- | --- | --- | --- | --- | --- | --- | --- | --- | --- | --- | --- | --- | --- | --- | --- | --- | --- | --- | --- | --- | --- | --- | --- | --- | --- | --- | --- | --- | --- | --- | --- | --- | --- | --- | --- | --- | --- | --- | --- | --- | --- | --- | --- | --- | --- | --- | --- | --- | --- | --- | --- | --- | --- | --- | --- | --- | --- | --- | --- | --- | --- | --- | --- | --- | --- | --- | --- | --- | --- | --- | --- | --- | --- | --- | --- | --- | --- | --- | --- | --- | --- | --- | --- | --- | --- | --- | --- | --- | --- | --- | --- | --- | --- | --- | --- | --- | --- | --- | --- | --- | --- | --- | --- | --- | --- | --- | --- | --- | --- | --- | --- | --- | --- | --- | --- | --- | --- | --- | --- | --- | --- | --- | --- | --- | --- | --- | --- | --- | --- | --- | --- | --- | --- | --- | --- | --- | --- | --- | --- | --- | --- | --- | --- | --- | --- | --- | --- | --- | --- | --- | --- | --- | --- | --- | --- | --- | --- | --- | --- | --- | --- | --- | --- | --- | --- | --- | --- | --- | --- | --- | --- | --- | --- | --- | --- | --- | --- | --- | --- | --- | --- | --- | --- | --- | --- | --- | --- | --- | --- | --- | --- | --- | --- | --- | --- | --- | --- | --- | --- | --- | --- | --- | --- | --- | --- | --- | --- | --- | --- | --- | --- | --- | --- | --- | --- | --- | --- | --- | --- | --- | --- | --- | --- | --- | --- | --- | --- | --- | --- | --- | --- | --- | --- | --- | --- | --- | --- | --- | --- | --- | --- | --- | --- | --- | --- | --- | --- | --- | --- | --- | --- | --- | --- | --- | --- | --- | --- | --- | --- | --- | --- | --- | --- | --- | --- | --- | --- | --- | --- | --- | --- | --- |
| left anterior | 0.436840 | 0.410768 | 0.291090 | 0.302821 | 0.314021 | 0.250388 | 0.223240 | 0.210959 | 0.213999 | 0.220556 | 0.300209 | 0.723652 | 0.617887 | 0.406452 | 0.384366 | 0.390159 | 0.297184 | 0.240360 | 0.211355 | 0.222827 | 0.339344 | 0.278490 | 0.274822 | 0.401493 | 0.669548 | 2.611016 | 4.303357 | 1.352842 | 1.057116 | 0.822270 | 0.413651 | 0.235740 | 0.210996 | 0.219175 | 0.219846 | 0.244596 | 0.273491 | 0.274627 | 0.278563 | 0.384750 | 0.687247 | 0.790389 | 0.584046 | 0.518573 | 0.605135 | 0.428491 | 0.293750 | 0.227056 | 0.215063 | 0.258747 | 0.289604 | 0.273878 | 0.397901 | 0.558128 | 1.472220 | 2.205179 | 0.807144 | 0.365126 | 0.339203 | 0.357375 | 0.339481 | 0.278304 | 0.326908 | 0.353106 | 0.438955 | 0.575701 | 0.380594 | 0.244730 | 0.214678 | 0.295601 | 0.442110 | 0.984911 | 3.486358 | 9.695028 | 9.043476 | 2.489866 | 0.428989 | 0.220614 | 0.211148 | 0.216008 | 0.216793 | 0.210919 | 0.219945 | 0.223684 | 0.211895 | 0.217053 | 0.221581 | 0.230474 | 0.259536 | 0.278856 | 0.245643 | 0.210858 | 0.210911 | 0.212848 | 0.211606 | 0.222430 | 0.225023 | 0.226521 | 0.231373 | 0.273738 | 0.270258 | 0.265059 | 0.231284 | 0.228966 | 0.223663 | 0.212477 | 0.211680 | 0.213721 | 0.210872 | 0.212285 | 0.220888 | 0.237607 | 0.245133 | 0.240252 | 0.231962 | 0.235295 | 0.216767 | 0.214116 | 0.211538 | 0.211458 | 0.221390 | 0.230958 | 0.236421 | 0.239929 | 0.251362 | 0.290207 | 0.296020 | 0.250933 | 0.216280 | 0.210988 | 0.211282 | 0.213825 | 0.211602 | 0.220666 | 0.229036 | 0.231281 | 0.250769 | 0.341557 | 0.448989 | 0.486677 | 0.490695 | 0.456661 | 0.417612 | 0.419195 | 0.375466 | 0.351439 | 0.316787 | 0.355733 | 0.468469 | 0.531728 | 0.457235 | 0.411611 | 0.414130 | 0.635365 | 0.816537 | 0.946320 | 0.795922 | 0.626759 | 0.448937 | 0.346376 | 0.294297 | 0.243645 | 0.222444 | 0.217649 | 0.215583 | 0.226160 | 0.231705 | 0.225748 | 0.217612 | 0.223330 | 0.241810 | 0.462726 | 0.753213 | 1.405465 | 2.196765 | 3.302586 | 2.931422 | 1.711769 | 0.943130 | 0.771177 | 0.784336 | 0.600429 | 0.693655 | 0.944672 | 1.830834 | 2.229887 | 2.174010 | 1.863685 | 3.592495 | 4.314204 | 6.134582 | 3.906409 | 2.726511 | 3.021768 | 2.548206 | 1.514832 | 1.291825 | 0.986090 | 1.010189 | 1.103674 | 0.915892 | 1.310846 | 1.942259 | 2.026484 | 1.959471 | 1.616009 | 1.405130 | 1.504426 | 1.408668 | 1.231363 | 1.343652 | 1.113822 | 0.910584 | 0.861300 | 0.875648 | 0.790922 | 0.660890 | 0.530650 | 0.488407 | 0.520340 | 0.471335 | 0.426400 | 0.429134 | 0.509000 | 0.603183 | 0.710559 | 0.759319 | 0.850307 | 0.684876 | 0.601688 | 0.461809 | 0.381040 | 0.327477 | 0.297669 | 0.252627 | 0.259444 | 0.247392 | 0.235870 | 0.219544 | 0.212123 | 0.211328 | 0.213233 | 0.211014 | 0.217341 | 0.215553 | 0.210806 | 0.215902 | 0.216558 | 0.214956 | 0.225959 | 0.290616 | 0.361766 | 0.350200 | 0.316838 | 0.297812 | 0.269593 | 0.247454 | 0.222897 | 0.216890 | 0.217492 | 0.214652 | 0.212688 | 0.213465 | 0.224093 | 0.222210 | 0.225072 | 0.221853 | 0.250945 | 0.291778 | 0.327477 | 0.269375 | 0.241186 | 0.225100 | 0.222449 | 0.210968 | 0.214248 | 0.213273 | 0.211732 | 0.213150 | 0.216804 |
| right anterior | 0.210984 | 0.219911 | 0.226389 | 0.252328 | 0.315804 | 0.444514 | 0.299432 | 0.273815 | 0.235834 | 0.213783 | 0.216451 | 0.264887 | 0.508418 | 0.881251 | 0.698590 | 0.409790 | 0.395928 | 0.362521 | 0.287529 | 0.241540 | 0.220690 | 0.221607 | 0.264073 | 0.324678 | 0.334721 | 0.389439 | 0.311911 | 0.346271 | 0.256546 | 0.210887 | 0.213200 | 0.228730 | 0.278248 | 0.266680 | 0.231527 | 0.220420 | 0.212382 | 0.214286 | 0.211397 | 0.215750 | 0.238682 | 0.297724 | 0.293735 | 0.278544 | 0.257322 | 0.218717 | 0.231314 | 0.403475 | 0.944765 | 2.768181 | 4.443503 | 6.620774 | 2.831810 | 1.432022 | 0.748393 | 0.559748 | 0.398237 | 0.365504 | 0.317476 | 0.256993 | 0.211498 | 0.211601 | 0.211851 | 0.211604 | 0.210977 | 0.224109 | 0.218805 | 0.214393 | 0.255067 | 0.313013 | 0.290547 | 0.296501 | 0.365924 | 0.389277 | 0.336591 | 0.297690 | 0.252371 | 0.227786 | 0.247803 | 0.308109 | 0.591314 | 1.305613 | 1.663936 | 1.471197 | 2.099947 | 2.123593 | 1.259266 | 0.409669 | 0.248809 | 0.221569 | 0.218842 | 0.211518 | 0.211028 | 0.210857 | 0.211557 | 0.215437 | 0.212899 | 0.211383 | 0.211786 | 0.211609 | 0.223008 | 0.219931 | 0.226043 | 0.236821 | 0.266154 | 0.279793 | 0.326346 | 0.372639 | 0.530957 | 0.724575 | 0.827805 | 0.813974 | 0.692184 | 0.898887 | 0.687988 | 0.549757 | 0.440126 | 0.337708 | 0.253862 | 0.216953 | 0.241003 | 0.323903 | 0.347725 | 0.360176 | 0.369685 | 0.342036 | 0.281638 | 0.260526 | 0.262526 | 0.247768 | 0.245512 | 0.234179 | 0.224068 | 0.217484 | 0.210814 | 0.219055 | 0.228347 | 0.259911 | 0.278779 | 0.296279 | 0.315807 | 0.308477 | 0.376705 | 0.436962 | 0.449632 | 0.427982 | 0.340846 | 0.319795 | 0.313186 | 0.276923 | 0.280977 | 0.288499 | 0.286164 | 0.336506 | 0.376531 | 0.596318 | 0.859379 | 0.930750 | 0.792872 | 0.730935 | 0.600003 | 0.503980 | 0.378591 | 0.267060 | 0.217648 | 0.214595 | 0.221809 | 0.237254 | 0.230638 | 0.212815 | 0.212293 | 0.227679 | 0.252476 | 0.277412 | 0.284941 | 0.267650 | 0.243739 | 0.232016 | 0.222774 | 0.220681 | 0.216558 | 0.216851 | 0.211610 | 0.220359 | 0.253502 | 0.278586 | 0.266741 | 0.255335 | 0.235138 | 0.245732 | 0.244012 | 0.220385 | 0.213782 | 0.224662 | 0.244365 | 0.295666 | 0.302447 | 0.262048 | 0.239688 | 0.236719 | 0.221060 | 0.224559 | 0.218944 | 0.225315 | 0.271518 | 0.339455 | 0.401907 | 0.440606 | 0.422605 | 0.397985 | 0.357627 | 0.289761 | 0.273861 | 0.239440 | 0.236018 | 0.246756 | 0.263016 | 0.275367 | 0.282278 | 0.286313 | 0.289500 | 0.284530 | 0.257966 | 0.228752 | 0.214929 | 0.210990 | 0.222377 | 0.236258 | 0.245679 | 0.238493 | 0.219088 | 0.210851 | 0.214842 | 0.228665 | 0.263221 | 0.267997 | 0.268941 | 0.288084 | 0.289310 | 0.307589 | 0.295832 | 0.279014 | 0.296105 | 0.292142 | 0.263127 | 0.242056 | 0.234984 | 0.228539 | 0.219537 | 0.214439 | 0.213300 | 0.212351 | 0.210812 | 0.211806 | 0.210934 | 0.219554 | 0.230763 | 0.226735 | 0.223622 | 0.250299 | 0.285153 | 0.277404 | 0.250528 | 0.246201 | 0.252969 | 0.283472 | 0.274267 | 0.258805 | 0.261028 | 0.262728 | 0.252646 | 0.262487 | 0.251129 | 0.241979 | 0.254229 | 0.258102 | 0.291092 | 0.291196 | 0.278584 | 0.292126 |
| left central | 0.213703 | 0.215227 | 0.220692 | 0.218256 | 0.233437 | 0.214826 | 0.218338 | 0.236350 | 0.231742 | 0.234809 | 0.219919 | 0.211982 | 0.230974 | 0.247757 | 0.282558 | 0.227526 | 0.231445 | 0.253923 | 0.295719 | 0.338733 | 0.314614 | 0.251121 | 0.366358 | 0.355340 | 0.210833 | 0.280765 | 0.240991 | 0.226616 | 0.223889 | 0.243776 | 0.258726 | 0.218225 | 0.211333 | 0.270345 | 0.410020 | 0.552060 | 0.702357 | 0.753070 | 1.039209 | 0.792830 | 0.502876 | 0.389575 | 0.303816 | 0.238835 | 0.229730 | 0.214145 | 0.210862 | 0.210820 | 0.225069 | 0.264708 | 0.271459 | 0.279847 | 0.283682 | 0.279668 | 0.331327 | 0.286167 | 0.234229 | 0.211814 | 0.212418 | 0.246471 | 0.373403 | 0.512973 | 0.800330 | 0.655144 | 0.628733 | 0.856385 | 1.197542 | 1.528700 | 3.766956 | 6.546670 | 13.107386 | 15.371791 | 13.758779 | 16.270054 | 17.825011 | 6.470299 | 1.385457 | 0.582170 | 0.319514 | 0.276017 | 0.227984 | 0.213836 | 0.210820 | 0.210921 | 0.210818 | 0.211242 | 0.210904 | 0.211042 | 0.213053 | 0.212098 | 0.210816 | 0.216778 | 0.223676 | 0.245213 | 0.284357 | 0.292339 | 0.280534 | 0.283545 | 0.243997 | 0.214333 | 0.212457 | 0.226061 | 0.214596 | 0.211039 | 0.213147 | 0.212779 | 0.210938 | 0.211000 | 0.210817 | 0.214372 | 0.236073 | 0.219801 | 0.222338 | 0.235100 | 0.239320 | 0.246638 | 0.253298 | 0.254364 | 0.348369 | 0.568010 | 0.953212 | 1.656898 | 2.000885 | 2.517804 | 3.425342 | 2.984353 | 2.510403 | 2.242678 | 1.587218 | 1.225415 | 1.133938 | 1.056983 | 0.876240 | 0.471958 | 0.296679 | 0.243114 | 0.241898 | 0.225348 | 0.217768 | 0.222929 | 0.235463 | 0.234174 | 0.226843 | 0.222449 | 0.223909 | 0.219878 | 0.212232 | 0.216758 | 0.223777 | 0.219163 | 0.215365 | 0.217654 | 0.217525 | 0.217363 | 0.215463 | 0.222557 | 0.246057 | 0.285873 | 0.277164 | 0.287109 | 0.261763 | 0.237529 | 0.219162 | 0.211077 | 0.216953 | 0.225460 | 0.224455 | 0.228522 | 0.232696 | 0.235713 | 0.246950 | 0.258819 | 0.313907 | 0.417765 | 0.390714 | 0.402750 | 0.426026 | 0.413067 | 0.431082 | 0.312978 | 0.257897 | 0.248461 | 0.243578 | 0.235162 | 0.232996 | 0.214740 | 0.213153 | 0.213591 | 0.214988 | 0.212320 | 0.211582 | 0.211053 | 0.212623 | 0.216802 | 0.219711 | 0.219995 | 0.221679 | 0.222301 | 0.235593 | 0.241452 | 0.229570 | 0.220032 | 0.212710 | 0.212520 | 0.215221 | 0.216870 | 0.226735 | 0.245160 | 0.270795 | 0.319858 | 0.366047 | 0.367296 | 0.328494 | 0.258708 | 0.233383 | 0.219586 | 0.215840 | 0.212592 | 0.213719 | 0.217249 | 0.228520 | 0.230018 | 0.234297 | 0.227718 | 0.225655 | 0.216022 | 0.211749 | 0.211427 | 0.211527 | 0.210812 | 0.211175 | 0.211082 | 0.211167 | 0.211281 | 0.210813 | 0.211388 | 0.211448 | 0.213998 | 0.230712 | 0.244519 | 0.255072 | 0.231217 | 0.221101 | 0.219809 | 0.216950 | 0.216579 | 0.212767 | 0.211474 | 0.216517 | 0.217274 | 0.215461 | 0.212134 | 0.211036 | 0.212472 | 0.210864 | 0.210864 | 0.210863 | 0.210806 | 0.210971 | 0.211047 | 0.210867 | 0.210814 | 0.210815 | 0.210815 | 0.211255 | 0.212532 | 0.214523 | 0.219042 | 0.213866 | 0.211646 | 0.210813 | 0.210960 | 0.211428 | 0.211459 | 0.218453 | 0.215194 | 0.213303 | 0.212335 | 0.211281 | 0.210835 |
| right central | 0.211741 | 0.215186 | 0.241911 | 0.328818 | 0.406183 | 0.708078 | 1.122420 | 1.289234 | 1.069605 | 0.774651 | 0.467097 | 0.371628 | 0.265069 | 0.225434 | 0.220826 | 0.228753 | 0.241717 | 0.262151 | 0.246383 | 0.241818 | 0.211687 | 0.210948 | 0.219247 | 0.229677 | 0.229357 | 0.256814 | 0.230943 | 0.211574 | 0.212926 | 0.211135 | 0.211539 | 0.213438 | 0.237891 | 0.272088 | 0.303478 | 0.293136 | 0.307896 | 0.261812 | 0.234121 | 0.221537 | 0.212202 | 0.222942 | 0.273064 | 0.277891 | 0.215680 | 0.213583 | 0.212044 | 0.275725 | 0.449328 | 0.774221 | 0.824211 | 0.596277 | 0.709310 | 0.975364 | 0.385095 | 0.267800 | 0.252028 | 0.229589 | 0.219020 | 0.211886 | 0.234943 | 0.213092 | 0.219913 | 0.238537 | 0.241974 | 0.211275 | 0.320861 | 2.773165 | 28.983259 | 330.423380 | 255.689188 | 154.225671 | 72.403582 | 35.168547 | 5.125569 | 1.799162 | 0.865945 | 0.776010 | 0.702747 | 0.699367 | 0.695549 | 0.719145 | 0.705625 | 0.878343 | 0.652932 | 0.544232 | 0.378328 | 0.308884 | 0.277564 | 0.263155 | 0.235320 | 0.218898 | 0.212476 | 0.213258 | 0.214252 | 0.220874 | 0.238897 | 0.259910 | 0.308139 | 0.358585 | 0.400747 | 0.399207 | 0.437611 | 0.357650 | 0.367304 | 0.367698 | 0.422776 | 0.512394 | 0.745900 | 0.939763 | 1.232161 | 0.782363 | 0.608213 | 0.443330 | 0.330931 | 0.306655 | 0.283338 | 0.305670 | 0.505979 | 0.804165 | 1.217681 | 1.862638 | 1.933555 | 2.602063 | 3.757480 | 3.175075 | 2.303860 | 2.148471 | 2.516191 | 2.536821 | 3.076948 | 3.691475 | 4.047513 | 3.215064 | 1.264418 | 0.691147 | 0.599724 | 0.471948 | 0.427008 | 0.385310 | 0.436231 | 0.540119 | 0.521169 | 0.464305 | 0.440105 | 0.528961 | 0.658047 | 0.556701 | 0.457679 | 0.421284 | 0.338964 | 0.312578 | 0.266851 | 0.236817 | 0.235409 | 0.253670 | 0.251978 | 0.301982 | 0.458814 | 0.600530 | 0.635550 | 0.951918 | 1.173921 | 1.642175 | 2.057003 | 1.987619 | 2.455500 | 2.565812 | 1.728276 | 0.890764 | 0.529126 | 0.379899 | 0.321028 | 0.265900 | 0.269403 | 0.278193 | 0.341013 | 0.429674 | 0.514198 | 0.458792 | 0.569947 | 0.607362 | 0.598529 | 0.445183 | 0.402442 | 0.428579 | 0.506898 | 0.576945 | 0.664620 | 0.623777 | 0.557849 | 0.553083 | 0.511675 | 0.536602 | 0.458675 | 0.344335 | 0.323067 | 0.303774 | 0.309197 | 0.316945 | 0.284327 | 0.237442 | 0.220627 | 0.216984 | 0.218379 | 0.211403 | 0.210839 | 0.210872 | 0.211239 | 0.215988 | 0.214274 | 0.216329 | 0.225819 | 0.229107 | 0.226705 | 0.233402 | 0.228215 | 0.253158 | 0.258489 | 0.229960 | 0.211087 | 0.222415 | 0.241674 | 0.255246 | 0.309943 | 0.366713 | 0.413060 | 0.340700 | 0.319380 | 0.288840 | 0.264715 | 0.219470 | 0.213481 | 0.257485 | 0.264114 | 0.319465 | 0.336892 | 0.527962 | 0.622907 | 0.581042 | 0.507606 | 0.578747 | 0.703942 | 0.923604 | 0.739892 | 0.584146 | 0.547255 | 0.551388 | 0.638252 | 0.575772 | 0.501321 | 0.419396 | 0.453164 | 0.437243 | 0.495036 | 0.551417 | 0.602669 | 0.635387 | 0.854805 | 0.995126 | 1.201281 | 0.930787 | 0.573587 | 0.408957 | 0.378153 | 0.358944 | 0.359755 | 0.388317 | 0.404192 | 0.558162 | 0.757461 | 0.978577 | 1.157207 | 0.774363 | 0.546003 | 0.548768 | 0.561044 | 0.562490 | 0.487843 | 0.427408 |
| left posterior | 0.220710 | 0.215186 | 0.214733 | 0.237001 | 0.215028 | 0.214001 | 0.210822 | 0.219571 | 0.234057 | 0.239004 | 0.295585 | 0.371338 | 0.443736 | 0.362716 | 0.323944 | 0.440064 | 0.801562 | 1.082417 | 0.685009 | 0.558762 | 0.435345 | 0.233700 | 0.219600 | 0.279792 | 0.331614 | 0.324485 | 0.348391 | 0.303371 | 0.477084 | 0.832430 | 0.876817 | 0.582205 | 0.451183 | 0.336588 | 0.447190 | 0.255020 | 0.216450 | 0.210912 | 0.213119 | 0.231866 | 0.224221 | 0.234161 | 0.258290 | 0.270408 | 0.249541 | 0.243535 | 0.224024 | 0.243448 | 0.329990 | 0.488038 | 0.747053 | 0.900253 | 1.010053 | 1.197165 | 1.300508 | 0.870013 | 0.976623 | 2.491463 | 5.912120 | 16.768076 | 38.372615 | 20.063124 | 6.484410 | 1.095869 | 0.422992 | 0.266717 | 0.213285 | 0.395790 | 1.365153 | 8.020938 | 47.227853 | 393.877961 | 1891.601577 | 2325.892622 | 1371.351107 | 511.594741 | 324.721959 | 89.027585 | 28.779812 | 18.742488 | 12.123728 | 7.415445 | 2.945574 | 1.302773 | 0.743709 | 0.563622 | 0.464439 | 0.531057 | 0.655708 | 0.897195 | 0.875129 | 0.640437 | 0.440219 | 0.379061 | 0.272201 | 0.238565 | 0.217506 | 0.211125 | 0.212154 | 0.212587 | 0.228475 | 0.288656 | 0.357214 | 0.445754 | 0.670396 | 0.646213 | 0.486354 | 0.408924 | 0.308505 | 0.271235 | 0.251317 | 0.245400 | 0.234917 | 0.242546 | 0.254175 | 0.290742 | 0.313232 | 0.348163 | 0.350944 | 0.509118 | 0.700140 | 1.039022 | 2.111860 | 3.834534 | 5.876636 | 7.623097 | 8.078648 | 9.460091 | 10.240285 | 6.621590 | 5.170149 | 4.339171 | 3.322085 | 3.593150 | 3.480744 | 2.835697 | 2.287645 | 2.958114 | 4.077913 | 5.138567 | 5.234608 | 4.669602 | 5.758494 | 6.344982 | 4.142964 | 2.470499 | 1.356365 | 0.650008 | 0.383006 | 0.282918 | 0.250990 | 0.242268 | 0.243888 | 0.266711 | 0.321696 | 0.592219 | 1.344704 | 2.157495 | 2.347256 | 2.494506 | 1.994912 | 2.155607 | 1.543803 | 1.055045 | 1.103793 | 1.172692 | 1.196021 | 1.608989 | 1.980599 | 2.226732 | 2.144933 | 1.411044 | 1.272626 | 1.049560 | 1.132039 | 1.573221 | 2.382786 | 2.690492 | 3.281626 | 2.872160 | 2.425307 | 2.004003 | 1.745937 | 1.709120 | 2.086949 | 2.810912 | 3.955534 | 3.893470 | 2.963661 | 2.646432 | 1.984150 | 1.444611 | 1.226043 | 0.895080 | 0.915421 | 1.242985 | 1.384479 | 1.488650 | 1.675600 | 1.698524 | 1.752750 | 1.509206 | 0.915174 | 0.637982 | 0.508131 | 0.509550 | 0.465410 | 0.444108 | 0.463096 | 0.550946 | 0.782926 | 1.155644 | 1.225678 | 1.167553 | 0.942188 | 0.721887 | 0.567567 | 0.351747 | 0.271918 | 0.240724 | 0.232281 | 0.227593 | 0.237317 | 0.247373 | 0.291713 | 0.324660 | 0.370130 | 0.397228 | 0.393970 | 0.381712 | 0.421960 | 0.395277 | 0.403341 | 0.459353 | 0.591615 | 0.693338 | 0.791415 | 0.779256 | 1.307798 | 1.988675 | 1.932786 | 1.603807 | 1.618503 | 1.591379 | 1.928123 | 1.765718 | 1.885987 | 2.005918 | 1.634226 | 1.294395 | 1.469306 | 0.982839 | 0.640137 | 0.379617 | 0.265069 | 0.236077 | 0.223514 | 0.212104 | 0.212835 | 0.216377 | 0.224359 | 0.257836 | 0.281838 | 0.290663 | 0.305963 | 0.304653 | 0.304480 | 0.301914 | 0.294526 | 0.358136 | 0.549761 | 0.706657 | 0.735048 | 0.637104 | 0.625446 | 0.660824 | 0.853101 | 0.647571 | 0.572730 | 0.594691 |
| right posterior | 0.210943 | 0.210913 | 0.210806 | 0.212635 | 0.224427 | 0.215445 | 0.212004 | 0.212058 | 0.214897 | 0.228988 | 0.222795 | 0.230125 | 0.237222 | 0.269419 | 0.309367 | 0.331399 | 0.318688 | 0.313298 | 0.244830 | 0.220740 | 0.219820 | 0.212357 | 0.213751 | 0.214165 | 0.234594 | 0.284038 | 0.485010 | 1.017148 | 3.205769 | 2.330647 | 0.929665 | 0.642295 | 0.502697 | 0.359794 | 0.238337 | 0.216067 | 0.225420 | 0.247042 | 0.222045 | 0.213685 | 0.217641 | 0.216775 | 0.210850 | 0.213113 | 0.217143 | 0.220390 | 0.211540 | 0.213893 | 0.224359 | 0.278300 | 0.323465 | 0.317828 | 0.324990 | 0.298853 | 0.300723 | 0.291501 | 0.211826 | 0.211708 | 0.224944 | 0.255204 | 0.290601 | 0.343774 | 0.589829 | 1.301724 | 0.831392 | 0.325581 | 0.212865 | 0.305406 | 1.015487 | 6.049602 | 40.571463 | 151.403748 | 366.871190 | 511.973929 | 415.126983 | 167.047742 | 32.621444 | 8.975927 | 2.690754 | 0.739291 | 0.360514 | 0.247289 | 0.215454 | 0.212675 | 0.220233 | 0.232631 | 0.228164 | 0.215100 | 0.211081 | 0.218488 | 0.258797 | 0.275668 | 0.289267 | 0.271620 | 0.262068 | 0.251451 | 0.249115 | 0.256304 | 0.258483 | 0.269061 | 0.292767 | 0.316942 | 0.360388 | 0.395191 | 0.400675 | 0.583746 | 0.716789 | 0.809837 | 1.044479 | 1.100911 | 0.982002 | 1.060003 | 1.120175 | 1.456058 | 1.573313 | 1.116723 | 0.909924 | 0.776880 | 0.797394 | 0.709968 | 0.670356 | 0.763796 | 1.017279 | 1.349474 | 2.391245 | 4.072732 | 5.049629 | 5.226934 | 4.961266 | 4.481540 | 3.906146 | 3.151779 | 2.384113 | 2.683775 | 4.523330 | 10.060627 | 20.483517 | 34.389047 | 40.857149 | 35.073990 | 24.290110 | 12.212164 | 7.998372 | 5.098121 | 3.857904 | 2.775724 | 1.795587 | 1.021198 | 0.636632 | 0.430638 | 0.327695 | 0.278278 | 0.250149 | 0.262114 | 0.263431 | 0.275421 | 0.308499 | 0.353914 | 0.442184 | 0.603193 | 0.799984 | 1.520701 | 2.442226 | 2.676106 | 3.066325 | 2.769682 | 2.706067 | 3.205116 | 3.569388 | 4.085599 | 3.979763 | 2.646490 | 1.764309 | 1.438766 | 1.006952 | 0.917799 | 0.912799 | 0.871846 | 0.825465 | 0.707057 | 0.501525 | 0.426046 | 0.349817 | 0.313019 | 0.280068 | 0.280020 | 0.331874 | 0.450041 | 0.498250 | 0.568647 | 0.670337 | 0.790769 | 0.994406 | 0.969182 | 0.755907 | 0.669955 | 0.540360 | 0.423992 | 0.370116 | 0.331080 | 0.289903 | 0.260825 | 0.243127 | 0.237388 | 0.247577 | 0.297527 | 0.342413 | 0.445365 | 0.714093 | 0.884190 | 1.138114 | 1.192822 | 0.780973 | 0.545027 | 0.393662 | 0.319788 | 0.292794 | 0.276120 | 0.261051 | 0.248150 | 0.231500 | 0.218417 | 0.212254 | 0.210951 | 0.211484 | 0.216554 | 0.231182 | 0.243870 | 0.230334 | 0.216752 | 0.211156 | 0.213339 | 0.220345 | 0.236603 | 0.259673 | 0.260707 | 0.240720 | 0.224059 | 0.211799 | 0.212177 | 0.213834 | 0.213775 | 0.212134 | 0.211662 | 0.212111 | 0.220804 | 0.218637 | 0.215134 | 0.216357 | 0.216078 | 0.215965 | 0.217603 | 0.210812 | 0.213675 | 0.224905 | 0.248070 | 0.280698 | 0.311544 | 0.330343 | 0.314233 | 0.305358 | 0.312494 | 0.296417 | 0.300982 | 0.328942 | 0.367645 | 0.326658 | 0.294605 | 0.252718 | 0.230102 | 0.211673 | 0.220444 | 0.239018 | 0.249989 | 0.250692 | 0.248506 | 0.253740 | 0.243691 | 0.240002 | 0.260685 |
| all electrodes | 0.214748 | 0.212925 | 0.248367 | 0.286631 | 0.302247 | 0.334049 | 0.267717 | 0.223806 | 0.215526 | 0.265371 | 0.455521 | 0.611111 | 0.801719 | 0.792097 | 0.710814 | 0.309118 | 0.238180 | 0.212603 | 0.211335 | 0.210825 | 0.213047 | 0.229850 | 0.217400 | 0.215727 | 0.213615 | 0.212185 | 0.212207 | 0.218459 | 0.241933 | 0.253593 | 0.211745 | 0.221564 | 0.218891 | 0.211206 | 0.212643 | 0.221379 | 0.229538 | 0.222765 | 0.221120 | 0.212108 | 0.212247 | 0.213089 | 0.233638 | 0.238971 | 0.225624 | 0.237633 | 0.217634 | 0.211564 | 0.211190 | 0.216570 | 0.224822 | 0.221326 | 0.264199 | 0.228963 | 0.215204 | 0.212272 | 0.211095 | 0.231461 | 0.364080 | 0.668218 | 0.893162 | 1.365947 | 1.501450 | 1.096366 | 0.571817 | 0.294892 | 0.214535 | 0.255099 | 0.729857 | 5.333802 | 27.337100 | 73.458648 | 243.569651 | 779.611786 | 1744.017157 | 729.005807 | 124.249696 | 21.644138 | 5.892158 | 4.021768 | 3.418448 | 3.257938 | 3.279621 | 4.553500 | 5.244418 | 3.137871 | 1.403058 | 0.831524 | 0.394517 | 0.374489 | 0.291789 | 0.239312 | 0.219405 | 0.217976 | 0.225302 | 0.236133 | 0.220317 | 0.216588 | 0.227504 | 0.255392 | 0.283439 | 0.337303 | 0.355674 | 0.541079 | 1.127179 | 1.704462 | 1.809951 | 2.046665 | 1.265002 | 1.799288 | 1.650888 | 1.007053 | 0.800675 | 0.725732 | 0.762751 | 0.892463 | 0.622763 | 0.685782 | 0.817037 | 1.103935 | 1.896118 | 3.261945 | 4.285370 | 6.140257 | 6.448956 | 8.386135 | 6.879304 | 6.868386 | 6.225568 | 6.033182 | 5.372013 | 3.016489 | 1.252654 | 0.608568 | 0.335598 | 0.253926 | 0.231560 | 0.226512 | 0.214254 | 0.211889 | 0.219309 | 0.236003 | 0.228019 | 0.231080 | 0.225678 | 0.271002 | 0.357864 | 0.376932 | 0.347891 | 0.360145 | 0.288766 | 0.287009 | 0.233227 | 0.213252 | 0.213102 | 0.218446 | 0.215892 | 0.211373 | 0.241799 | 0.450909 | 1.056440 | 2.833487 | 11.937243 | 24.983368 | 25.254592 | 35.727107 | 40.200920 | 35.784106 | 23.698616 | 18.908429 | 11.047348 | 12.025739 | 8.675337 | 3.723396 | 2.392883 | 1.950052 | 1.006636 | 0.910297 | 0.739454 | 0.590553 | 0.707308 | 0.780147 | 1.146534 | 1.791076 | 2.986779 | 6.040609 | 7.435307 | 8.764084 | 15.272950 | 10.338323 | 5.471175 | 2.314078 | 1.481622 | 1.485787 | 1.323231 | 0.837389 | 0.810974 | 0.701528 | 0.840558 | 1.309228 | 1.449543 | 1.407536 | 2.092477 | 2.358262 | 2.515315 | 2.340617 | 2.146139 | 2.360049 | 3.783953 | 5.461244 | 6.451031 | 11.444632 | 10.910436 | 5.244353 | 3.188079 | 1.847780 | 1.314388 | 1.029714 | 0.805005 | 0.753039 | 0.663548 | 0.681591 | 0.640218 | 0.529411 | 0.424889 | 0.375923 | 0.345302 | 0.397783 | 0.381722 | 0.409487 | 0.476505 | 0.767746 | 1.010980 | 1.054645 | 0.848035 | 0.961143 | 1.149105 | 1.138480 | 0.833732 | 0.829884 | 1.107362 | 1.507996 | 1.638845 | 1.299415 | 1.407367 | 2.438000 | 3.201458 | 2.711360 | 2.025789 | 1.829660 | 1.432457 | 0.951726 | 0.618412 | 0.445016 | 0.359575 | 0.336594 | 0.307839 | 0.323890 | 0.317608 | 0.301777 | 0.364136 | 0.440913 | 0.522244 | 0.642041 | 0.638238 | 0.853482 | 0.884336 | 0.772670 | 0.854045 | 0.874128 | 0.875574 | 1.173311 | 1.482559 | 3.497764 | 6.515651 | 7.783311 | 13.577991 | 14.797864 | 14.250555 | 8.414309 |

Searchlight, spatiotemporal cluster permutation test

No significant clusters observed.

Q) real to emoji faces - happy vs sad

  
|  | time window | peak latency | cluster *p* | peak Cohen's *d* |  | | | |
| **all electrodes** | 135 - 225 ms | 170 ms | 0.0329 | 0.9814 |  | | | |
|  | | | | | | | | |

Time-resolved classification, cluster permutation tests

|  | **left hemisphere** | | | | **right hemisphere** | | | |
|  | time window | peak latency | cluster *p* | peak Cohen's *d* | time window | peak latency | cluster *p* | peak Cohen's *d* |
| **anterior** |  | | | |  | | | |
| **central** |  | | | |  | | | |
| **posterior** |  | | | | 105 - 205 ms | 175 ms | 0.028 | 1.0135 |

  

Time-resolved classification, Bayesian statistics

|  | -200 | -195 | -190 | -185 | -180 | -175 | -170 | -165 | -160 | -155 | -150 | -145 | -140 | -135 | -130 | -125 | -120 | -115 | -110 | -105 | -100 | -95 | -90 | -85 | -80 | -75 | -70 | -65 | -60 | -55 | -50 | -45 | -40 | -35 | -30 | -25 | -20 | -15 | -10 | -5 | 0 | 5 | 10 | 15 | 20 | 25 | 30 | 35 | 40 | 45 | 50 | 55 | 60 | 65 | 70 | 75 | 80 | 85 | 90 | 95 | 100 | 105 | 110 | 115 | 120 | 125 | 130 | 135 | 140 | 145 | 150 | 155 | 160 | 165 | 170 | 175 | 180 | 185 | 190 | 195 | 200 | 205 | 210 | 215 | 220 | 225 | 230 | 235 | 240 | 245 | 250 | 255 | 260 | 265 | 270 | 275 | 280 | 285 | 290 | 295 | 300 | 305 | 310 | 315 | 320 | 325 | 330 | 335 | 340 | 345 | 350 | 355 | 360 | 365 | 370 | 375 | 380 | 385 | 390 | 395 | 400 | 405 | 410 | 415 | 420 | 425 | 430 | 435 | 440 | 445 | 450 | 455 | 460 | 465 | 470 | 475 | 480 | 485 | 490 | 495 | 500 | 505 | 510 | 515 | 520 | 525 | 530 | 535 | 540 | 545 | 550 | 555 | 560 | 565 | 570 | 575 | 580 | 585 | 590 | 595 | 600 | 605 | 610 | 615 | 620 | 625 | 630 | 635 | 640 | 645 | 650 | 655 | 660 | 665 | 670 | 675 | 680 | 685 | 690 | 695 | 700 | 705 | 710 | 715 | 720 | 725 | 730 | 735 | 740 | 745 | 750 | 755 | 760 | 765 | 770 | 775 | 780 | 785 | 790 | 795 | 800 | 805 | 810 | 815 | 820 | 825 | 830 | 835 | 840 | 845 | 850 | 855 | 860 | 865 | 870 | 875 | 880 | 885 | 890 | 895 | 900 | 905 | 910 | 915 | 920 | 925 | 930 | 935 | 940 | 945 | 950 | 955 | 960 | 965 | 970 | 975 | 980 | 985 | 990 | 995 | 1000 | 1005 | 1010 | 1015 | 1020 | 1025 | 1030 | 1035 | 1040 | 1045 | 1050 | 1055 | 1060 | 1065 | 1070 | 1075 | 1080 | 1085 | 1090 | 1095 | 1100 | 1105 | 1110 | 1115 | 1120 | 1125 | 1130 | 1135 | 1140 | 1145 | 1150 | 1155 | 1160 | 1165 | 1170 | 1175 | 1180 | 1185 | 1190 | 1195 |
| --- | --- | --- | --- | --- | --- | --- | --- | --- | --- | --- | --- | --- | --- | --- | --- | --- | --- | --- | --- | --- | --- | --- | --- | --- | --- | --- | --- | --- | --- | --- | --- | --- | --- | --- | --- | --- | --- | --- | --- | --- | --- | --- | --- | --- | --- | --- | --- | --- | --- | --- | --- | --- | --- | --- | --- | --- | --- | --- | --- | --- | --- | --- | --- | --- | --- | --- | --- | --- | --- | --- | --- | --- | --- | --- | --- | --- | --- | --- | --- | --- | --- | --- | --- | --- | --- | --- | --- | --- | --- | --- | --- | --- | --- | --- | --- | --- | --- | --- | --- | --- | --- | --- | --- | --- | --- | --- | --- | --- | --- | --- | --- | --- | --- | --- | --- | --- | --- | --- | --- | --- | --- | --- | --- | --- | --- | --- | --- | --- | --- | --- | --- | --- | --- | --- | --- | --- | --- | --- | --- | --- | --- | --- | --- | --- | --- | --- | --- | --- | --- | --- | --- | --- | --- | --- | --- | --- | --- | --- | --- | --- | --- | --- | --- | --- | --- | --- | --- | --- | --- | --- | --- | --- | --- | --- | --- | --- | --- | --- | --- | --- | --- | --- | --- | --- | --- | --- | --- | --- | --- | --- | --- | --- | --- | --- | --- | --- | --- | --- | --- | --- | --- | --- | --- | --- | --- | --- | --- | --- | --- | --- | --- | --- | --- | --- | --- | --- | --- | --- | --- | --- | --- | --- | --- | --- | --- | --- | --- | --- | --- | --- | --- | --- | --- | --- | --- | --- | --- | --- | --- | --- | --- | --- | --- | --- | --- | --- | --- | --- | --- | --- | --- | --- | --- | --- | --- | --- | --- | --- | --- | --- | --- | --- | --- | --- | --- | --- | --- | --- | --- | --- | --- | --- | --- | --- | --- | --- | --- | --- | --- | --- |
| left anterior | 0.290833 | 0.328564 | 0.301826 | 0.345338 | 0.642838 | 0.419800 | 0.231650 | 0.229149 | 0.346073 | 0.456272 | 1.208701 | 1.305376 | 0.621981 | 0.366146 | 0.289745 | 0.236106 | 0.212192 | 0.216813 | 0.256350 | 0.290663 | 0.266668 | 0.258256 | 0.217841 | 0.243409 | 0.349543 | 0.628264 | 0.701910 | 0.357356 | 0.218394 | 0.234327 | 0.482342 | 0.633457 | 0.643159 | 0.519814 | 0.280179 | 0.217521 | 0.211036 | 0.215071 | 0.210828 | 0.236604 | 0.282226 | 0.453972 | 0.470379 | 0.340132 | 0.362687 | 0.344431 | 0.366849 | 0.400468 | 0.407380 | 0.268932 | 0.257399 | 0.228748 | 0.211253 | 0.211135 | 0.214818 | 0.249063 | 0.228168 | 0.249345 | 0.270619 | 0.248370 | 0.266497 | 0.299540 | 0.290734 | 0.264701 | 0.250387 | 0.266442 | 0.270255 | 0.358490 | 0.467175 | 0.570115 | 0.484058 | 0.373641 | 0.315179 | 0.540829 | 0.961837 | 1.677556 | 3.920437 | 16.843314 | 57.805755 | 83.040824 | 17.292189 | 2.727935 | 0.792379 | 0.364540 | 0.250611 | 0.226964 | 0.227033 | 0.244178 | 0.226731 | 0.221541 | 0.229694 | 0.257349 | 0.374192 | 0.471173 | 0.575250 | 0.883244 | 1.451087 | 1.177729 | 0.794653 | 0.444051 | 0.408177 | 0.346818 | 0.344054 | 0.229986 | 0.216522 | 0.210973 | 0.210893 | 0.212458 | 0.213210 | 0.214865 | 0.211617 | 0.210806 | 0.220212 | 0.229483 | 0.242913 | 0.239909 | 0.226735 | 0.218702 | 0.214329 | 0.212928 | 0.213340 | 0.217452 | 0.239001 | 0.297883 | 0.421261 | 0.447284 | 0.471797 | 0.425124 | 0.334870 | 0.284240 | 0.235919 | 0.217306 | 0.216265 | 0.214550 | 0.210951 | 0.222771 | 0.288337 | 0.428118 | 0.805521 | 1.653743 | 2.827401 | 4.051363 | 2.834347 | 2.798557 | 2.337114 | 1.243721 | 0.631217 | 0.469690 | 0.364763 | 0.357345 | 0.260806 | 0.224008 | 0.221775 | 0.242499 | 0.259618 | 0.233557 | 0.220969 | 0.213319 | 0.211007 | 0.214346 | 0.247396 | 0.329280 | 0.286414 | 0.260101 | 0.244308 | 0.228744 | 0.234155 | 0.232741 | 0.232078 | 0.282993 | 0.396378 | 0.417350 | 0.375486 | 0.322862 | 0.289809 | 0.286887 | 0.254511 | 0.242177 | 0.256820 | 0.292298 | 0.305574 | 0.341881 | 0.351559 | 0.461507 | 0.487164 | 0.421327 | 0.469594 | 0.624608 | 0.735946 | 0.787971 | 0.523762 | 0.436279 | 0.467373 | 0.347280 | 0.264500 | 0.213378 | 0.218899 | 0.214477 | 0.213535 | 0.211353 | 0.211326 | 0.213380 | 0.221280 | 0.219832 | 0.211204 | 0.212275 | 0.218404 | 0.219659 | 0.218188 | 0.220877 | 0.219405 | 0.230050 | 0.235745 | 0.231828 | 0.234859 | 0.229895 | 0.217997 | 0.214452 | 0.210814 | 0.210884 | 0.210806 | 0.212438 | 0.221866 | 0.238240 | 0.243454 | 0.244885 | 0.220598 | 0.220370 | 0.213084 | 0.210860 | 0.220430 | 0.236509 | 0.269066 | 0.328093 | 0.343706 | 0.312639 | 0.327771 | 0.298568 | 0.258502 | 0.227831 | 0.210806 | 0.212138 | 0.210985 | 0.213914 | 0.221622 | 0.214449 | 0.211060 | 0.226799 | 0.229869 | 0.219452 | 0.211767 | 0.213066 | 0.216751 | 0.211550 | 0.210916 | 0.224802 | 0.258073 | 0.366455 | 0.404621 | 0.412632 | 0.499770 | 0.500560 | 0.383497 | 0.385352 | 0.319921 | 0.284693 | 0.264865 | 0.234519 | 0.217096 | 0.218858 | 0.221565 | 0.224895 | 0.232044 | 0.229838 | 0.248355 | 0.261752 | 0.225777 | 0.215504 | 0.211584 | 0.210843 |
| right anterior | 0.211212 | 0.214284 | 0.220910 | 0.232007 | 0.233825 | 0.263063 | 0.387897 | 0.363720 | 0.317672 | 0.248867 | 0.214208 | 0.212828 | 0.233436 | 0.262083 | 0.282927 | 0.264517 | 0.296718 | 0.381127 | 0.790957 | 0.598597 | 0.607380 | 0.499894 | 0.450592 | 0.250153 | 0.222499 | 0.214869 | 0.222860 | 0.210821 | 0.212565 | 0.221324 | 0.257949 | 0.290892 | 0.357882 | 0.527499 | 0.381835 | 0.344132 | 0.360902 | 0.546644 | 0.510479 | 0.353664 | 0.279454 | 0.252675 | 0.250695 | 0.248556 | 0.250858 | 0.308204 | 0.395518 | 0.559281 | 0.925785 | 1.872430 | 2.438187 | 2.102431 | 0.942330 | 0.567876 | 0.352689 | 0.300174 | 0.269225 | 0.314393 | 0.408314 | 0.513321 | 0.687964 | 0.615246 | 0.570166 | 0.505759 | 0.435189 | 0.394720 | 0.400665 | 0.349060 | 0.497883 | 0.386267 | 0.294934 | 0.237958 | 0.217632 | 0.219684 | 0.233467 | 0.256038 | 0.304336 | 0.434672 | 0.560624 | 0.589529 | 0.472605 | 0.399735 | 0.292759 | 0.264570 | 0.274537 | 0.375150 | 0.452296 | 0.478158 | 0.408019 | 0.384345 | 0.435095 | 0.432517 | 0.411738 | 0.599965 | 0.731014 | 0.831901 | 0.825431 | 0.525270 | 0.312073 | 0.249970 | 0.210821 | 0.217070 | 0.251382 | 0.261958 | 0.285677 | 0.247873 | 0.235748 | 0.225777 | 0.214571 | 0.211633 | 0.214329 | 0.210884 | 0.212151 | 0.225667 | 0.223688 | 0.217012 | 0.211724 | 0.211710 | 0.212720 | 0.234434 | 0.275630 | 0.310808 | 0.345811 | 0.359261 | 0.435466 | 0.410977 | 0.272254 | 0.273591 | 0.278377 | 0.224556 | 0.210822 | 0.217707 | 0.222808 | 0.212349 | 0.214759 | 0.242743 | 0.276749 | 0.337057 | 0.456883 | 0.482164 | 0.538262 | 0.559045 | 0.767145 | 0.939238 | 0.862890 | 0.685878 | 0.531799 | 0.414385 | 0.432190 | 0.378906 | 0.336782 | 0.320393 | 0.319967 | 0.387406 | 0.528121 | 0.717357 | 0.723107 | 0.762050 | 0.596672 | 0.494228 | 0.385927 | 0.362407 | 0.361995 | 0.338559 | 0.303019 | 0.289653 | 0.280028 | 0.292432 | 0.248734 | 0.219520 | 0.210806 | 0.213282 | 0.214981 | 0.211697 | 0.211126 | 0.211654 | 0.211329 | 0.211496 | 0.211399 | 0.213718 | 0.212369 | 0.210900 | 0.211178 | 0.212167 | 0.212707 | 0.212286 | 0.212017 | 0.210985 | 0.214245 | 0.215240 | 0.225450 | 0.249336 | 0.287476 | 0.313960 | 0.328669 | 0.356980 | 0.510975 | 0.763244 | 0.951276 | 0.633444 | 0.378807 | 0.283098 | 0.232752 | 0.210815 | 0.256036 | 0.410006 | 0.616510 | 0.783131 | 0.633722 | 0.590706 | 0.525835 | 0.440146 | 0.425035 | 0.339392 | 0.298608 | 0.281384 | 0.244424 | 0.224168 | 0.213730 | 0.211278 | 0.213240 | 0.221593 | 0.231438 | 0.250628 | 0.295344 | 0.369298 | 0.393905 | 0.374937 | 0.331624 | 0.259665 | 0.217313 | 0.222579 | 0.315545 | 0.713461 | 2.052393 | 4.349860 | 3.997624 | 4.728146 | 4.647263 | 2.668344 | 1.149929 | 0.453320 | 0.285053 | 0.254850 | 0.239428 | 0.234316 | 0.230713 | 0.245413 | 0.293882 | 0.456541 | 0.636683 | 0.629855 | 0.743348 | 0.872091 | 0.779747 | 0.666682 | 0.492424 | 0.432186 | 0.374409 | 0.340662 | 0.327912 | 0.344490 | 0.318998 | 0.320961 | 0.383484 | 0.440759 | 0.440201 | 0.310878 | 0.264723 | 0.261481 | 0.281587 | 0.288871 | 0.259435 | 0.217418 | 0.211536 | 0.210977 | 0.210816 | 0.216635 | 0.256665 | 0.305181 |
| left central | 1.053513 | 0.731941 | 0.560789 | 0.390132 | 0.306668 | 0.231846 | 0.220580 | 0.210995 | 0.210966 | 0.211587 | 0.225006 | 0.235701 | 0.266433 | 0.248127 | 0.241028 | 0.289984 | 0.347065 | 0.328939 | 0.443454 | 0.719886 | 1.328960 | 2.001598 | 2.773948 | 1.185213 | 0.729885 | 0.315576 | 0.222396 | 0.212114 | 0.212635 | 0.231488 | 0.230688 | 0.242791 | 0.213762 | 0.210919 | 0.210910 | 0.214879 | 0.212352 | 0.211964 | 0.224612 | 0.256875 | 0.279479 | 0.331754 | 0.395422 | 0.522576 | 0.547857 | 0.407358 | 0.297539 | 0.314694 | 0.356467 | 0.385087 | 0.339697 | 0.337607 | 0.577418 | 1.038611 | 1.183228 | 0.674809 | 0.426017 | 0.466915 | 0.565363 | 0.459821 | 0.419548 | 0.368898 | 0.399686 | 0.415547 | 0.292128 | 0.232528 | 0.210820 | 0.271130 | 0.448408 | 0.459483 | 0.406866 | 0.324829 | 0.316065 | 0.372591 | 0.429926 | 0.419051 | 0.398967 | 0.338432 | 0.264564 | 0.221907 | 0.217853 | 0.301668 | 0.419213 | 0.450247 | 0.382437 | 0.355076 | 0.343997 | 0.346505 | 0.335813 | 0.333973 | 0.346809 | 0.332361 | 0.267660 | 0.230702 | 0.213691 | 0.257128 | 0.297284 | 0.307173 | 0.287301 | 0.231037 | 0.211087 | 0.234763 | 0.303988 | 0.326903 | 0.298903 | 0.299288 | 0.292935 | 0.328417 | 0.378132 | 0.377595 | 0.370811 | 0.371281 | 0.367012 | 0.315619 | 0.259089 | 0.214067 | 0.225169 | 0.269825 | 0.360539 | 0.558603 | 0.645061 | 0.630676 | 0.484086 | 0.349725 | 0.335126 | 0.307405 | 0.252618 | 0.212457 | 0.210817 | 0.210883 | 0.212901 | 0.226945 | 0.264541 | 0.293757 | 0.301927 | 0.328986 | 0.350155 | 0.342656 | 0.290421 | 0.283147 | 0.277799 | 0.290976 | 0.283275 | 0.311669 | 0.300023 | 0.289044 | 0.257824 | 0.265198 | 0.224256 | 0.212376 | 0.211308 | 0.211615 | 0.211462 | 0.211446 | 0.220472 | 0.218377 | 0.211176 | 0.217116 | 0.246027 | 0.297469 | 0.352232 | 0.446720 | 0.464393 | 0.393624 | 0.403480 | 0.379572 | 0.310405 | 0.256227 | 0.222179 | 0.211447 | 0.210956 | 0.211095 | 0.215473 | 0.241623 | 0.270310 | 0.295427 | 0.311213 | 0.288845 | 0.264953 | 0.231110 | 0.217826 | 0.216034 | 0.214041 | 0.223827 | 0.234842 | 0.223296 | 0.218712 | 0.211821 | 0.211002 | 0.210837 | 0.211176 | 0.210814 | 0.211911 | 0.216971 | 0.224827 | 0.244665 | 0.258340 | 0.258502 | 0.249980 | 0.254616 | 0.242181 | 0.250521 | 0.262823 | 0.241877 | 0.245528 | 0.241616 | 0.233351 | 0.229410 | 0.212958 | 0.211200 | 0.212838 | 0.215390 | 0.214524 | 0.212389 | 0.210895 | 0.213512 | 0.216921 | 0.214860 | 0.213410 | 0.214433 | 0.214856 | 0.210885 | 0.210815 | 0.211314 | 0.210814 | 0.211566 | 0.211449 | 0.210814 | 0.210840 | 0.211624 | 0.216270 | 0.225398 | 0.252355 | 0.283580 | 0.305084 | 0.335464 | 0.383233 | 0.349285 | 0.363316 | 0.331379 | 0.347493 | 0.331124 | 0.293910 | 0.277083 | 0.287295 | 0.282334 | 0.339677 | 0.352024 | 0.494100 | 0.622663 | 0.546041 | 0.580378 | 0.524409 | 0.403874 | 0.353493 | 0.249347 | 0.229565 | 0.240631 | 0.237231 | 0.233964 | 0.237968 | 0.242099 | 0.268884 | 0.255643 | 0.251153 | 0.273573 | 0.337393 | 0.385046 | 0.498365 | 0.622566 | 0.561290 | 0.493268 | 0.345335 | 0.251414 | 0.222983 | 0.218805 | 0.211008 | 0.210824 | 0.212850 | 0.211120 |
| right central | 0.367407 | 0.418839 | 0.414505 | 0.452231 | 0.346720 | 0.229371 | 0.220565 | 0.290523 | 0.303950 | 0.353172 | 0.398161 | 0.396361 | 0.305960 | 0.246595 | 0.290730 | 0.471113 | 0.395214 | 0.429121 | 0.315024 | 0.279414 | 0.223064 | 0.217594 | 0.238605 | 0.249151 | 0.218189 | 0.252014 | 0.384375 | 0.515214 | 0.480810 | 0.441319 | 0.564483 | 0.366572 | 0.213516 | 0.248523 | 0.230533 | 0.211616 | 0.220076 | 0.210829 | 0.212963 | 0.220171 | 0.212000 | 0.217173 | 0.235305 | 0.306543 | 0.294494 | 0.241824 | 0.229177 | 0.229531 | 0.210874 | 0.224494 | 0.288366 | 0.329310 | 0.341371 | 0.314174 | 0.311769 | 0.261920 | 0.217281 | 0.217256 | 0.241569 | 0.314232 | 0.326631 | 0.259411 | 0.221716 | 0.211217 | 0.231673 | 0.318945 | 0.521927 | 1.151124 | 1.591084 | 1.127179 | 0.629112 | 0.552590 | 0.597896 | 0.820908 | 1.092980 | 1.124065 | 1.051492 | 1.138560 | 0.738337 | 0.515272 | 0.325475 | 0.247987 | 0.246647 | 0.235529 | 0.223895 | 0.217099 | 0.214287 | 0.236015 | 0.263404 | 0.345072 | 0.575848 | 0.819445 | 0.848986 | 0.624342 | 0.501908 | 0.533833 | 0.503908 | 0.356517 | 0.340139 | 0.329763 | 0.282999 | 0.288823 | 0.300986 | 0.295766 | 0.234109 | 0.213319 | 0.210818 | 0.211303 | 0.219920 | 0.244500 | 0.231924 | 0.217132 | 0.220408 | 0.216253 | 0.211362 | 0.212942 | 0.211459 | 0.217913 | 0.224571 | 0.226171 | 0.252626 | 0.349666 | 0.392333 | 0.393639 | 0.371155 | 0.296498 | 0.254488 | 0.233211 | 0.213234 | 0.212486 | 0.213252 | 0.227997 | 0.236422 | 0.253629 | 0.311078 | 0.321290 | 0.276020 | 0.306155 | 0.263004 | 0.232874 | 0.213240 | 0.213166 | 0.214743 | 0.213314 | 0.216699 | 0.213473 | 0.215849 | 0.218599 | 0.214919 | 0.212667 | 0.210814 | 0.212521 | 0.219532 | 0.246822 | 0.306566 | 0.357439 | 0.389814 | 0.307893 | 0.275401 | 0.224585 | 0.211261 | 0.211723 | 0.214375 | 0.223187 | 0.234221 | 0.271266 | 0.273654 | 0.322171 | 0.358785 | 0.464335 | 0.549055 | 0.443120 | 0.329108 | 0.236111 | 0.210806 | 0.210914 | 0.219651 | 0.218582 | 0.227217 | 0.223763 | 0.243531 | 0.262175 | 0.247083 | 0.227550 | 0.212621 | 0.210819 | 0.212422 | 0.213772 | 0.212569 | 0.212486 | 0.212881 | 0.221791 | 0.236038 | 0.236980 | 0.237129 | 0.234260 | 0.222914 | 0.212590 | 0.211107 | 0.211302 | 0.211406 | 0.212499 | 0.212584 | 0.211982 | 0.250295 | 0.300120 | 0.351861 | 0.371109 | 0.378213 | 0.362045 | 0.325193 | 0.269274 | 0.281964 | 0.279745 | 0.282291 | 0.300739 | 0.330297 | 0.343738 | 0.366157 | 0.323399 | 0.290945 | 0.264068 | 0.220255 | 0.212211 | 0.214420 | 0.210879 | 0.211575 | 0.222280 | 0.227482 | 0.234894 | 0.255775 | 0.269110 | 0.360570 | 0.508488 | 0.646213 | 1.032757 | 1.709453 | 2.077380 | 1.981245 | 1.371330 | 0.798066 | 0.488746 | 0.329220 | 0.293891 | 0.282888 | 0.296626 | 0.312153 | 0.364613 | 0.558453 | 0.633886 | 0.511275 | 0.419811 | 0.400468 | 0.360371 | 0.365415 | 0.330296 | 0.296928 | 0.265026 | 0.246035 | 0.251743 | 0.249923 | 0.238438 | 0.225300 | 0.250546 | 0.374211 | 0.920627 | 1.799136 | 2.374455 | 1.786585 | 1.478129 | 1.092114 | 0.976504 | 0.733479 | 0.467334 | 0.357156 | 0.342037 | 0.346206 | 0.327165 | 0.259656 | 0.234240 |
| left posterior | 0.223093 | 0.212152 | 0.210806 | 0.212191 | 0.218556 | 0.236649 | 0.218320 | 0.210860 | 0.212316 | 0.217131 | 0.221620 | 0.220607 | 0.211704 | 0.211889 | 0.300561 | 0.547525 | 0.342144 | 0.271794 | 0.217053 | 0.228520 | 0.310153 | 0.306908 | 0.296351 | 0.210827 | 0.237112 | 0.367512 | 1.184226 | 5.316230 | 3.264730 | 4.025526 | 2.185873 | 2.150417 | 1.483862 | 0.743581 | 0.388152 | 0.274767 | 0.243629 | 0.224207 | 0.217208 | 0.219467 | 0.218775 | 0.213039 | 0.224763 | 0.224884 | 0.254063 | 0.232372 | 0.241146 | 0.231009 | 0.236309 | 0.226788 | 0.214609 | 0.211158 | 0.211152 | 0.243946 | 0.313836 | 0.338678 | 0.339160 | 0.302561 | 0.255180 | 0.214292 | 0.222766 | 0.366297 | 0.594273 | 0.776290 | 0.779769 | 0.519390 | 0.527519 | 0.657042 | 0.569231 | 0.554008 | 0.565124 | 0.692289 | 1.569063 | 2.326308 | 3.070608 | 3.868601 | 2.928161 | 1.709581 | 1.102885 | 0.707559 | 0.439430 | 0.344123 | 0.306727 | 0.268222 | 0.257226 | 0.222345 | 0.211865 | 0.211402 | 0.239886 | 0.360948 | 0.505499 | 0.628407 | 0.491930 | 0.326033 | 0.258152 | 0.211353 | 0.238861 | 0.306515 | 0.355710 | 0.424412 | 0.492148 | 0.599477 | 0.532391 | 0.482199 | 0.505055 | 0.567641 | 0.472159 | 0.347334 | 0.262886 | 0.252192 | 0.235263 | 0.219543 | 0.226513 | 0.223064 | 0.220344 | 0.241845 | 0.240447 | 0.247122 | 0.244911 | 0.228984 | 0.231909 | 0.258101 | 0.269656 | 0.293024 | 0.305990 | 0.308344 | 0.300201 | 0.307251 | 0.284768 | 0.232780 | 0.211100 | 0.248469 | 0.258529 | 0.255191 | 0.236132 | 0.239423 | 0.239177 | 0.224286 | 0.210849 | 0.213158 | 0.216140 | 0.212050 | 0.213166 | 0.219712 | 0.223410 | 0.253076 | 0.288054 | 0.301116 | 0.287257 | 0.286637 | 0.339382 | 0.394463 | 0.388739 | 0.330633 | 0.271371 | 0.254008 | 0.237015 | 0.214349 | 0.211555 | 0.220074 | 0.223118 | 0.214375 | 0.210884 | 0.213144 | 0.225071 | 0.248382 | 0.276801 | 0.345127 | 0.386212 | 0.344761 | 0.319138 | 0.293170 | 0.301029 | 0.327511 | 0.315484 | 0.328178 | 0.390934 | 0.471857 | 0.536966 | 0.526538 | 0.491547 | 0.585684 | 0.648213 | 0.660113 | 0.712525 | 0.878912 | 0.852883 | 0.890065 | 0.698577 | 0.560977 | 0.503181 | 0.453415 | 0.384922 | 0.341919 | 0.304137 | 0.265438 | 0.252110 | 0.246737 | 0.261978 | 0.284359 | 0.324138 | 0.368500 | 0.452115 | 0.552787 | 0.598581 | 0.639429 | 0.678933 | 0.649110 | 0.640173 | 0.697725 | 0.820072 | 1.048178 | 0.996754 | 0.843598 | 0.856549 | 0.762154 | 0.668791 | 0.553793 | 0.500413 | 0.416046 | 0.372810 | 0.338951 | 0.334964 | 0.293256 | 0.283807 | 0.293539 | 0.334432 | 0.336416 | 0.337991 | 0.361222 | 0.485566 | 0.735253 | 0.883279 | 0.913425 | 1.042269 | 1.133107 | 1.403720 | 1.591432 | 1.735196 | 1.711411 | 1.280942 | 0.868261 | 0.710049 | 0.446117 | 0.341615 | 0.262892 | 0.235739 | 0.247759 | 0.268679 | 0.264445 | 0.255641 | 0.228611 | 0.214780 | 0.212264 | 0.227073 | 0.259106 | 0.269191 | 0.286245 | 0.266567 | 0.240591 | 0.221970 | 0.217220 | 0.211260 | 0.212803 | 0.212188 | 0.213779 | 0.214924 | 0.212211 | 0.210840 | 0.211361 | 0.220198 | 0.231097 | 0.250379 | 0.250488 | 0.236740 | 0.223561 | 0.216011 | 0.212607 | 0.211041 | 0.212270 |
| right posterior | 0.271472 | 0.283896 | 0.281617 | 0.282622 | 0.319789 | 0.335172 | 0.286203 | 0.251137 | 0.227281 | 0.226270 | 0.227452 | 0.221938 | 0.229921 | 0.254238 | 0.292104 | 0.466112 | 0.406447 | 0.366510 | 0.397566 | 0.253364 | 0.212873 | 0.228547 | 0.293730 | 0.340675 | 0.291444 | 0.280639 | 0.217708 | 0.211659 | 0.213540 | 0.212369 | 0.230073 | 0.246517 | 0.251195 | 0.277850 | 0.256267 | 0.341700 | 0.345398 | 0.276337 | 0.235937 | 0.220574 | 0.211032 | 0.211059 | 0.210960 | 0.221462 | 0.227945 | 0.232926 | 0.240378 | 0.250932 | 0.255322 | 0.226888 | 0.217005 | 0.223437 | 0.255363 | 0.277859 | 0.272599 | 0.252536 | 0.223431 | 0.220159 | 0.284052 | 0.486945 | 0.900797 | 2.137121 | 3.051521 | 6.813112 | 5.404947 | 4.254923 | 4.641797 | 8.840201 | 12.075163 | 13.101621 | 5.777256 | 4.540233 | 13.751942 | 51.531323 | 259.691230 | 691.926905 | 1658.540029 | 1527.050424 | 1009.454092 | 255.469989 | 36.715825 | 2.581049 | 0.523026 | 0.241866 | 0.212447 | 0.269721 | 0.390251 | 0.664247 | 1.100829 | 2.844951 | 5.186113 | 4.756773 | 2.126171 | 1.173911 | 0.592742 | 0.346784 | 0.264041 | 0.257506 | 0.275524 | 0.317324 | 0.401220 | 0.592539 | 0.918261 | 1.115962 | 0.977872 | 0.706951 | 0.669440 | 0.532312 | 0.339106 | 0.297112 | 0.290359 | 0.321462 | 0.271693 | 0.232543 | 0.217475 | 0.215422 | 0.211565 | 0.211150 | 0.222294 | 0.230601 | 0.234143 | 0.234737 | 0.241324 | 0.247409 | 0.258920 | 0.282851 | 0.316793 | 0.380105 | 0.377808 | 0.347756 | 0.293841 | 0.246956 | 0.217116 | 0.210806 | 0.212963 | 0.213462 | 0.210876 | 0.213261 | 0.221837 | 0.236605 | 0.257983 | 0.249941 | 0.234635 | 0.210817 | 0.216102 | 0.227345 | 0.224287 | 0.224766 | 0.232978 | 0.238932 | 0.223932 | 0.216491 | 0.214395 | 0.222500 | 0.231457 | 0.227629 | 0.211810 | 0.211091 | 0.212887 | 0.219939 | 0.243106 | 0.258461 | 0.301971 | 0.323664 | 0.370797 | 0.432865 | 0.347933 | 0.260018 | 0.249381 | 0.226015 | 0.212053 | 0.212290 | 0.228489 | 0.226075 | 0.215905 | 0.214786 | 0.211064 | 0.217798 | 0.246725 | 0.314574 | 0.465791 | 0.764456 | 1.202395 | 1.226154 | 0.961664 | 0.659889 | 0.497819 | 0.420116 | 0.375709 | 0.301204 | 0.248015 | 0.242334 | 0.254889 | 0.250139 | 0.243248 | 0.252215 | 0.260664 | 0.288216 | 0.313754 | 0.284835 | 0.294589 | 0.319249 | 0.320816 | 0.338975 | 0.339775 | 0.357961 | 0.455848 | 0.575987 | 0.611114 | 0.572933 | 0.599559 | 0.691825 | 0.707599 | 0.565140 | 0.452905 | 0.367041 | 0.323363 | 0.274153 | 0.254590 | 0.231306 | 0.231315 | 0.235358 | 0.253858 | 0.277291 | 0.281187 | 0.259501 | 0.245032 | 0.225546 | 0.223569 | 0.217410 | 0.213121 | 0.214810 | 0.216383 | 0.220632 | 0.226933 | 0.224035 | 0.219666 | 0.214852 | 0.211120 | 0.211250 | 0.210898 | 0.210988 | 0.211346 | 0.210806 | 0.211251 | 0.214009 | 0.214858 | 0.218419 | 0.224085 | 0.227149 | 0.223740 | 0.223834 | 0.221445 | 0.215098 | 0.210810 | 0.214344 | 0.227655 | 0.248010 | 0.262165 | 0.264950 | 0.244092 | 0.223937 | 0.215475 | 0.210948 | 0.211058 | 0.210842 | 0.210965 | 0.211701 | 0.218068 | 0.222434 | 0.227836 | 0.227887 | 0.213962 | 0.212955 | 0.211321 | 0.210832 | 0.210806 | 0.210839 | 0.211334 | 0.211807 |
| all electrodes | 0.262343 | 0.249568 | 0.246553 | 0.295556 | 0.313839 | 0.280347 | 0.307967 | 0.282992 | 0.349667 | 0.314785 | 0.271298 | 0.247733 | 0.262890 | 0.250992 | 0.244254 | 0.232779 | 0.229696 | 0.210894 | 0.211052 | 0.211141 | 0.213037 | 0.227897 | 0.241094 | 0.259615 | 0.291101 | 0.267533 | 0.232656 | 0.211902 | 0.211201 | 0.215201 | 0.223718 | 0.319535 | 0.545213 | 0.590322 | 0.733659 | 0.447897 | 0.228846 | 0.212007 | 0.244976 | 0.293622 | 0.279085 | 0.264638 | 0.254674 | 0.247071 | 0.238688 | 0.222424 | 0.210897 | 0.217567 | 0.240513 | 0.312453 | 0.610165 | 2.832128 | 3.304524 | 1.390266 | 1.042165 | 0.473390 | 0.247753 | 0.214125 | 0.260738 | 0.391959 | 0.493640 | 0.709954 | 0.437238 | 0.362578 | 0.316181 | 0.453525 | 0.605025 | 2.247454 | 16.705738 | 219.986195 | 1072.205396 | 592.069651 | 724.159145 | 621.696283 | 478.871114 | 217.506982 | 80.412739 | 35.936591 | 15.955723 | 6.440754 | 4.778348 | 2.030626 | 1.632368 | 2.978247 | 2.499194 | 1.954051 | 0.825653 | 0.416560 | 0.271812 | 0.211326 | 0.237903 | 0.246728 | 0.222763 | 0.212783 | 0.211248 | 0.221439 | 0.244870 | 0.242428 | 0.230580 | 0.232347 | 0.257764 | 0.284462 | 0.289950 | 0.285990 | 0.256826 | 0.255547 | 0.226627 | 0.213039 | 0.213062 | 0.246501 | 0.255761 | 0.216987 | 0.211385 | 0.213843 | 0.211098 | 0.211907 | 0.216498 | 0.211181 | 0.212245 | 0.210893 | 0.217047 | 0.248129 | 0.287116 | 0.351625 | 0.453389 | 0.473307 | 0.332687 | 0.279479 | 0.212558 | 0.219584 | 0.259658 | 0.296701 | 0.299433 | 0.257833 | 0.261629 | 0.259458 | 0.284083 | 0.297219 | 0.288087 | 0.304138 | 0.287836 | 0.257947 | 0.240502 | 0.230440 | 0.224565 | 0.220894 | 0.213767 | 0.214951 | 0.217460 | 0.214867 | 0.211775 | 0.210856 | 0.214536 | 0.238239 | 0.240403 | 0.234657 | 0.233551 | 0.261850 | 0.261735 | 0.219486 | 0.212259 | 0.214252 | 0.220478 | 0.250057 | 0.310446 | 0.362114 | 0.368618 | 0.388981 | 0.397622 | 0.423800 | 0.377832 | 0.350288 | 0.293492 | 0.247936 | 0.231970 | 0.231517 | 0.255470 | 0.278916 | 0.327451 | 0.420934 | 0.583577 | 1.000850 | 2.125890 | 3.016414 | 5.512368 | 9.082317 | 19.712465 | 17.060933 | 6.148055 | 2.333803 | 1.258535 | 0.954181 | 0.626110 | 0.453129 | 0.354951 | 0.303219 | 0.291810 | 0.305419 | 0.299557 | 0.356044 | 0.363944 | 0.376933 | 0.398084 | 0.374105 | 0.303879 | 0.352990 | 0.383247 | 0.476497 | 0.676448 | 0.733801 | 1.088354 | 2.512766 | 2.148840 | 1.748338 | 1.382563 | 0.935142 | 0.897463 | 0.640038 | 0.398103 | 0.282352 | 0.220255 | 0.212234 | 0.212110 | 0.213369 | 0.218297 | 0.219481 | 0.219958 | 0.242898 | 0.262739 | 0.291802 | 0.288747 | 0.315813 | 0.380867 | 0.470837 | 0.501208 | 0.592429 | 0.746948 | 0.910580 | 0.832209 | 0.748881 | 0.864852 | 0.688778 | 0.623039 | 0.616992 | 0.769806 | 0.855505 | 0.851874 | 0.779594 | 0.868936 | 0.859273 | 0.832394 | 0.659053 | 0.574394 | 0.468848 | 0.380181 | 0.346618 | 0.304828 | 0.304615 | 0.307166 | 0.309062 | 0.318907 | 0.326159 | 0.307599 | 0.295420 | 0.248653 | 0.224636 | 0.219570 | 0.217287 | 0.216599 | 0.224171 | 0.245726 | 0.268041 | 0.270132 | 0.248117 | 0.242855 | 0.237432 | 0.225218 | 0.214251 | 0.211965 | 0.211837 |

Searchlight, spatiotemporal cluster permutation test

No significant clusters observed.

R) real to emoji faces - neutral vs happy

  
|  | time window | peak latency | cluster *p* | peak Cohen's *d* |  | | | |
| **all electrodes** | 95 - 160 ms | 130 ms | 0.0197 | 1.5061 |  | | | |
|  | | | | | | | | |

Time-resolved classification, cluster permutation tests

|  | **left hemisphere** | | | | **right hemisphere** | | | |
|  | time window | peak latency | cluster *p* | peak Cohen's *d* | time window | peak latency | cluster *p* | peak Cohen's *d* |
| **anterior** |  | | | |  | | | |
| **central** |  | | | |  | | | |
| **posterior** |  | | | | 95 - 175 ms | 130 ms | 0.019 | 1.3655 |

  

Time-resolved classification, Bayesian statistics

|  | -200 | -195 | -190 | -185 | -180 | -175 | -170 | -165 | -160 | -155 | -150 | -145 | -140 | -135 | -130 | -125 | -120 | -115 | -110 | -105 | -100 | -95 | -90 | -85 | -80 | -75 | -70 | -65 | -60 | -55 | -50 | -45 | -40 | -35 | -30 | -25 | -20 | -15 | -10 | -5 | 0 | 5 | 10 | 15 | 20 | 25 | 30 | 35 | 40 | 45 | 50 | 55 | 60 | 65 | 70 | 75 | 80 | 85 | 90 | 95 | 100 | 105 | 110 | 115 | 120 | 125 | 130 | 135 | 140 | 145 | 150 | 155 | 160 | 165 | 170 | 175 | 180 | 185 | 190 | 195 | 200 | 205 | 210 | 215 | 220 | 225 | 230 | 235 | 240 | 245 | 250 | 255 | 260 | 265 | 270 | 275 | 280 | 285 | 290 | 295 | 300 | 305 | 310 | 315 | 320 | 325 | 330 | 335 | 340 | 345 | 350 | 355 | 360 | 365 | 370 | 375 | 380 | 385 | 390 | 395 | 400 | 405 | 410 | 415 | 420 | 425 | 430 | 435 | 440 | 445 | 450 | 455 | 460 | 465 | 470 | 475 | 480 | 485 | 490 | 495 | 500 | 505 | 510 | 515 | 520 | 525 | 530 | 535 | 540 | 545 | 550 | 555 | 560 | 565 | 570 | 575 | 580 | 585 | 590 | 595 | 600 | 605 | 610 | 615 | 620 | 625 | 630 | 635 | 640 | 645 | 650 | 655 | 660 | 665 | 670 | 675 | 680 | 685 | 690 | 695 | 700 | 705 | 710 | 715 | 720 | 725 | 730 | 735 | 740 | 745 | 750 | 755 | 760 | 765 | 770 | 775 | 780 | 785 | 790 | 795 | 800 | 805 | 810 | 815 | 820 | 825 | 830 | 835 | 840 | 845 | 850 | 855 | 860 | 865 | 870 | 875 | 880 | 885 | 890 | 895 | 900 | 905 | 910 | 915 | 920 | 925 | 930 | 935 | 940 | 945 | 950 | 955 | 960 | 965 | 970 | 975 | 980 | 985 | 990 | 995 | 1000 | 1005 | 1010 | 1015 | 1020 | 1025 | 1030 | 1035 | 1040 | 1045 | 1050 | 1055 | 1060 | 1065 | 1070 | 1075 | 1080 | 1085 | 1090 | 1095 | 1100 | 1105 | 1110 | 1115 | 1120 | 1125 | 1130 | 1135 | 1140 | 1145 | 1150 | 1155 | 1160 | 1165 | 1170 | 1175 | 1180 | 1185 | 1190 | 1195 |
| --- | --- | --- | --- | --- | --- | --- | --- | --- | --- | --- | --- | --- | --- | --- | --- | --- | --- | --- | --- | --- | --- | --- | --- | --- | --- | --- | --- | --- | --- | --- | --- | --- | --- | --- | --- | --- | --- | --- | --- | --- | --- | --- | --- | --- | --- | --- | --- | --- | --- | --- | --- | --- | --- | --- | --- | --- | --- | --- | --- | --- | --- | --- | --- | --- | --- | --- | --- | --- | --- | --- | --- | --- | --- | --- | --- | --- | --- | --- | --- | --- | --- | --- | --- | --- | --- | --- | --- | --- | --- | --- | --- | --- | --- | --- | --- | --- | --- | --- | --- | --- | --- | --- | --- | --- | --- | --- | --- | --- | --- | --- | --- | --- | --- | --- | --- | --- | --- | --- | --- | --- | --- | --- | --- | --- | --- | --- | --- | --- | --- | --- | --- | --- | --- | --- | --- | --- | --- | --- | --- | --- | --- | --- | --- | --- | --- | --- | --- | --- | --- | --- | --- | --- | --- | --- | --- | --- | --- | --- | --- | --- | --- | --- | --- | --- | --- | --- | --- | --- | --- | --- | --- | --- | --- | --- | --- | --- | --- | --- | --- | --- | --- | --- | --- | --- | --- | --- | --- | --- | --- | --- | --- | --- | --- | --- | --- | --- | --- | --- | --- | --- | --- | --- | --- | --- | --- | --- | --- | --- | --- | --- | --- | --- | --- | --- | --- | --- | --- | --- | --- | --- | --- | --- | --- | --- | --- | --- | --- | --- | --- | --- | --- | --- | --- | --- | --- | --- | --- | --- | --- | --- | --- | --- | --- | --- | --- | --- | --- | --- | --- | --- | --- | --- | --- | --- | --- | --- | --- | --- | --- | --- | --- | --- | --- | --- | --- | --- | --- | --- | --- | --- | --- | --- | --- | --- | --- | --- | --- | --- | --- | --- |
| left anterior | 3.629892 | 9.172674 | 7.955237 | 3.738027 | 1.215408 | 0.486462 | 0.244867 | 0.210919 | 0.210920 | 0.211847 | 0.244024 | 0.254569 | 0.376907 | 0.790181 | 0.971088 | 0.436958 | 0.238542 | 0.230468 | 0.218856 | 0.227698 | 0.218346 | 0.211188 | 0.223838 | 0.237521 | 0.293498 | 0.438912 | 0.343979 | 0.335666 | 0.769776 | 0.991214 | 2.383813 | 7.943917 | 2.475306 | 3.235570 | 2.747475 | 0.899270 | 0.582738 | 0.513582 | 0.401288 | 0.472172 | 0.643530 | 0.454158 | 0.284538 | 0.226736 | 0.211103 | 0.218765 | 0.225588 | 0.236613 | 0.210963 | 0.222654 | 0.236163 | 0.254596 | 0.310651 | 0.368763 | 0.272518 | 0.223535 | 0.212610 | 0.216458 | 0.250689 | 0.295484 | 0.247906 | 0.260195 | 0.231110 | 0.215720 | 0.332529 | 1.049781 | 3.491359 | 7.575001 | 8.412024 | 7.309123 | 2.631246 | 0.397312 | 0.210909 | 0.325948 | 0.611841 | 0.403280 | 0.324685 | 0.273599 | 0.218931 | 0.213274 | 0.211386 | 0.222614 | 0.242207 | 0.252354 | 0.242178 | 0.254255 | 0.224281 | 0.222588 | 0.364767 | 0.573256 | 0.928428 | 1.870748 | 3.261822 | 2.840210 | 2.255359 | 2.331742 | 1.432800 | 0.780491 | 0.444117 | 0.381682 | 0.342344 | 0.287434 | 0.274493 | 0.268652 | 0.253009 | 0.239274 | 0.221090 | 0.210894 | 0.211150 | 0.214415 | 0.212045 | 0.255629 | 0.346679 | 0.333752 | 0.544157 | 0.681894 | 0.727028 | 0.545346 | 0.435264 | 0.410037 | 0.387038 | 0.299623 | 0.308008 | 0.337003 | 0.381243 | 0.394051 | 0.348024 | 0.309384 | 0.327616 | 0.291417 | 0.258207 | 0.233042 | 0.232992 | 0.233321 | 0.243877 | 0.223229 | 0.213555 | 0.216133 | 0.239585 | 0.260506 | 0.314663 | 0.365046 | 0.457214 | 0.600662 | 0.602043 | 0.546152 | 0.484666 | 0.383823 | 0.286419 | 0.238148 | 0.212761 | 0.212717 | 0.218437 | 0.219157 | 0.221417 | 0.226825 | 0.239561 | 0.233503 | 0.227162 | 0.230177 | 0.252285 | 0.255977 | 0.256820 | 0.246665 | 0.268696 | 0.321030 | 0.501600 | 0.513195 | 0.587570 | 0.569203 | 0.703232 | 0.778844 | 0.812730 | 0.737843 | 0.872823 | 0.893267 | 0.676063 | 0.476536 | 0.480313 | 0.501886 | 0.548988 | 0.479005 | 0.444935 | 0.480246 | 0.440873 | 0.350873 | 0.295817 | 0.262455 | 0.273674 | 0.292250 | 0.276062 | 0.268122 | 0.239798 | 0.214084 | 0.211411 | 0.213403 | 0.210817 | 0.211959 | 0.212504 | 0.210923 | 0.223582 | 0.240617 | 0.250409 | 0.283994 | 0.417737 | 0.706150 | 0.847234 | 0.794680 | 0.654300 | 0.561332 | 0.506028 | 0.473084 | 0.372852 | 0.288869 | 0.250964 | 0.222834 | 0.210806 | 0.212363 | 0.219846 | 0.220862 | 0.214743 | 0.211861 | 0.210946 | 0.213204 | 0.214224 | 0.215005 | 0.212883 | 0.223915 | 0.227536 | 0.239214 | 0.268943 | 0.282101 | 0.278308 | 0.284567 | 0.253017 | 0.281200 | 0.296378 | 0.273439 | 0.248675 | 0.232658 | 0.236251 | 0.251204 | 0.236512 | 0.218209 | 0.210889 | 0.211049 | 0.212003 | 0.213764 | 0.214032 | 0.212846 | 0.212294 | 0.223854 | 0.232945 | 0.228314 | 0.218929 | 0.210915 | 0.211628 | 0.212178 | 0.220969 | 0.254123 | 0.295021 | 0.345385 | 0.381123 | 0.416449 | 0.420867 | 0.401237 | 0.284158 | 0.237742 | 0.222852 | 0.221093 | 0.218065 | 0.216068 | 0.212578 | 0.222062 | 0.225935 | 0.219500 | 0.213588 | 0.211377 | 0.210822 | 0.211166 |
| right anterior | 5.221504 | 16.946111 | 24.009686 | 39.170744 | 15.742887 | 9.060615 | 30.218258 | 23.888616 | 7.994281 | 2.823435 | 1.448656 | 0.582665 | 0.272640 | 0.216386 | 0.212734 | 0.213227 | 0.211500 | 0.222845 | 0.212101 | 0.350113 | 0.478315 | 0.621123 | 0.574260 | 0.821516 | 1.191418 | 0.938888 | 0.584760 | 0.456946 | 0.469899 | 0.559092 | 0.356582 | 0.335934 | 0.346716 | 0.276755 | 0.288891 | 0.222342 | 0.210820 | 0.210806 | 0.214527 | 0.212920 | 0.217622 | 0.219544 | 0.210885 | 0.211122 | 0.211634 | 0.212086 | 0.210872 | 0.210806 | 0.228248 | 0.260416 | 0.287045 | 0.351136 | 0.334343 | 0.349288 | 0.330818 | 0.382922 | 0.495064 | 0.585381 | 0.569105 | 0.952540 | 1.144582 | 2.100833 | 3.424317 | 6.470332 | 12.049786 | 19.651916 | 52.857637 | 70.352117 | 32.141441 | 41.900083 | 66.812464 | 78.095558 | 44.959958 | 5.411725 | 3.792780 | 2.749659 | 0.837673 | 0.450066 | 0.490936 | 0.359220 | 0.257726 | 0.210806 | 0.238254 | 0.266341 | 0.275243 | 0.288619 | 0.250529 | 0.212113 | 0.253530 | 0.418553 | 0.589750 | 0.456388 | 0.282723 | 0.239463 | 0.228858 | 0.216117 | 0.211068 | 0.211493 | 0.212553 | 0.214354 | 0.213098 | 0.212651 | 0.212923 | 0.215588 | 0.221935 | 0.227034 | 0.230723 | 0.242379 | 0.257466 | 0.285296 | 0.328877 | 0.310563 | 0.287031 | 0.285956 | 0.270762 | 0.242653 | 0.221353 | 0.211778 | 0.211447 | 0.210806 | 0.212478 | 0.212662 | 0.212600 | 0.222299 | 0.239318 | 0.221211 | 0.215276 | 0.212303 | 0.213663 | 0.217235 | 0.211324 | 0.210814 | 0.219168 | 0.220380 | 0.216796 | 0.211943 | 0.213966 | 0.217667 | 0.231501 | 0.252369 | 0.220313 | 0.217652 | 0.212534 | 0.210816 | 0.211762 | 0.211733 | 0.221309 | 0.233960 | 0.232409 | 0.223053 | 0.217581 | 0.217733 | 0.223665 | 0.228560 | 0.231300 | 0.242026 | 0.309609 | 0.391362 | 0.422761 | 0.398918 | 0.381555 | 0.390125 | 0.412478 | 0.348463 | 0.318195 | 0.289554 | 0.300808 | 0.319062 | 0.294292 | 0.242561 | 0.221544 | 0.213166 | 0.210876 | 0.212044 | 0.217065 | 0.214362 | 0.216118 | 0.215481 | 0.215223 | 0.217351 | 0.211971 | 0.212277 | 0.214786 | 0.218010 | 0.238290 | 0.275608 | 0.307940 | 0.365423 | 0.430068 | 0.494948 | 0.494322 | 0.461091 | 0.450894 | 0.477779 | 0.620472 | 0.787324 | 0.694343 | 0.546949 | 0.422499 | 0.349632 | 0.378898 | 0.438792 | 0.452127 | 0.610566 | 1.115385 | 2.155475 | 3.879342 | 2.869246 | 1.779849 | 1.180872 | 0.848987 | 0.738596 | 0.644991 | 0.573016 | 0.701052 | 0.985770 | 1.428456 | 1.462273 | 0.873827 | 0.574570 | 0.475759 | 0.390377 | 0.332930 | 0.306288 | 0.282793 | 0.316975 | 0.390690 | 0.447682 | 0.460152 | 0.468236 | 0.440135 | 0.421506 | 0.399218 | 0.402948 | 0.409037 | 0.503446 | 0.587677 | 0.595452 | 0.723700 | 0.792032 | 0.613267 | 0.561693 | 0.482345 | 0.417812 | 0.435673 | 0.423846 | 0.399319 | 0.415208 | 0.414496 | 0.431395 | 0.446077 | 0.424564 | 0.405589 | 0.396105 | 0.370274 | 0.372199 | 0.393595 | 0.435732 | 0.548774 | 0.611597 | 0.635768 | 0.718507 | 0.741121 | 0.756439 | 0.737320 | 0.766817 | 0.648643 | 0.595077 | 0.631365 | 0.643877 | 0.585805 | 0.559038 | 0.440438 | 0.379056 | 0.335386 | 0.300079 | 0.295690 | 0.290689 | 0.266130 | 0.251822 |
| left central | 0.254719 | 0.210853 | 0.211990 | 0.229636 | 0.465991 | 0.583887 | 0.434499 | 0.745850 | 2.992806 | 5.956310 | 8.680065 | 3.225504 | 2.935906 | 1.509979 | 0.632953 | 0.311109 | 0.219957 | 0.231132 | 0.211914 | 0.215582 | 0.212391 | 0.246640 | 0.231486 | 0.211661 | 0.294500 | 0.480226 | 0.625071 | 0.669256 | 0.303956 | 0.244160 | 0.240541 | 0.210829 | 0.211417 | 0.215668 | 0.224038 | 0.220690 | 0.237099 | 0.257887 | 0.269719 | 0.412742 | 0.547426 | 0.515223 | 0.393739 | 0.293629 | 0.248429 | 0.234359 | 0.243509 | 0.240179 | 0.241547 | 0.301762 | 0.440792 | 0.839428 | 2.305024 | 2.325289 | 2.802243 | 2.139804 | 1.187825 | 0.651216 | 0.530466 | 0.384438 | 0.278990 | 0.221731 | 0.215571 | 0.269611 | 0.423531 | 0.741065 | 1.786832 | 3.327454 | 3.831202 | 2.966703 | 2.033860 | 0.736355 | 0.309996 | 0.215098 | 0.246762 | 0.555649 | 1.988051 | 3.854627 | 2.604669 | 1.754186 | 1.431346 | 0.910874 | 0.527799 | 0.305396 | 0.241837 | 0.253341 | 0.224516 | 0.211922 | 0.222500 | 0.285557 | 0.290428 | 0.268421 | 0.282189 | 0.274797 | 0.265621 | 0.246582 | 0.223003 | 0.228167 | 0.242031 | 0.220749 | 0.211697 | 0.220746 | 0.235269 | 0.228801 | 0.239872 | 0.321659 | 0.360760 | 0.320525 | 0.254917 | 0.236592 | 0.230768 | 0.229013 | 0.210828 | 0.219615 | 0.291598 | 0.308487 | 0.392187 | 0.524957 | 0.848371 | 1.090397 | 1.412211 | 0.892958 | 0.709324 | 0.492296 | 0.418553 | 0.440802 | 0.411584 | 0.382315 | 0.329035 | 0.262157 | 0.226502 | 0.210806 | 0.229205 | 0.247006 | 0.264327 | 0.240461 | 0.223277 | 0.212294 | 0.242099 | 0.300924 | 0.267862 | 0.258181 | 0.252677 | 0.239802 | 0.228911 | 0.215063 | 0.211627 | 0.212921 | 0.233154 | 0.246580 | 0.245578 | 0.243026 | 0.267571 | 0.291148 | 0.302810 | 0.297836 | 0.333810 | 0.396178 | 0.390930 | 0.379639 | 0.273661 | 0.228624 | 0.211247 | 0.211562 | 0.220294 | 0.214277 | 0.211033 | 0.213377 | 0.223864 | 0.226495 | 0.233721 | 0.253100 | 0.232635 | 0.219447 | 0.212471 | 0.210817 | 0.211744 | 0.216426 | 0.221625 | 0.249682 | 0.302053 | 0.329110 | 0.379601 | 0.424372 | 0.356587 | 0.320365 | 0.256530 | 0.225120 | 0.213490 | 0.212580 | 0.224142 | 0.227296 | 0.213048 | 0.219339 | 0.237064 | 0.302868 | 0.431739 | 0.657490 | 0.848815 | 0.597057 | 0.449785 | 0.369256 | 0.272059 | 0.239289 | 0.215524 | 0.211064 | 0.210905 | 0.210989 | 0.219298 | 0.238008 | 0.286047 | 0.344399 | 0.414206 | 0.403518 | 0.446463 | 0.415938 | 0.318674 | 0.250006 | 0.217258 | 0.210857 | 0.210806 | 0.211718 | 0.211320 | 0.211197 | 0.210870 | 0.215726 | 0.228435 | 0.218341 | 0.215568 | 0.215889 | 0.227997 | 0.221263 | 0.213119 | 0.210957 | 0.212984 | 0.213537 | 0.219031 | 0.220475 | 0.210977 | 0.217303 | 0.243835 | 0.265240 | 0.241197 | 0.244800 | 0.259607 | 0.242501 | 0.219157 | 0.213540 | 0.217834 | 0.230282 | 0.232738 | 0.228765 | 0.232843 | 0.278749 | 0.296444 | 0.258678 | 0.247764 | 0.264283 | 0.258153 | 0.248054 | 0.221886 | 0.213061 | 0.211010 | 0.210986 | 0.219664 | 0.231900 | 0.232480 | 0.229434 | 0.230268 | 0.220224 | 0.215835 | 0.212970 | 0.214240 | 0.211523 | 0.211779 | 0.210806 | 0.211272 | 0.212692 | 0.216724 | 0.229403 |
| right central | 0.639881 | 0.494783 | 0.410596 | 0.310730 | 0.251596 | 0.211495 | 0.228916 | 0.222933 | 0.224218 | 0.222396 | 0.216060 | 0.213646 | 0.210806 | 0.210929 | 0.250303 | 0.324959 | 0.404072 | 0.415877 | 0.505317 | 0.386858 | 0.319098 | 0.269491 | 0.237026 | 0.217143 | 0.221516 | 0.213450 | 0.211735 | 0.214145 | 0.210824 | 0.213350 | 0.210860 | 0.210920 | 0.211029 | 0.224343 | 0.248099 | 0.239712 | 0.250069 | 0.235793 | 0.249757 | 0.239237 | 0.237743 | 0.244004 | 0.250318 | 0.247462 | 0.296001 | 0.328964 | 0.283231 | 0.232511 | 0.230900 | 0.224880 | 0.227727 | 0.243166 | 0.274590 | 0.515969 | 0.943261 | 0.971240 | 1.262156 | 1.058657 | 0.649719 | 0.424400 | 0.292028 | 0.252342 | 0.212703 | 0.265069 | 0.549239 | 1.361548 | 3.417670 | 6.334594 | 14.302963 | 9.176147 | 3.549135 | 1.134859 | 0.418262 | 0.241905 | 0.211187 | 0.235094 | 0.280823 | 0.369472 | 0.512041 | 0.560505 | 0.510686 | 0.479448 | 0.435914 | 0.329897 | 0.309560 | 0.298968 | 0.300492 | 0.241715 | 0.215251 | 0.211887 | 0.223095 | 0.334721 | 0.529391 | 0.571144 | 0.405519 | 0.329505 | 0.290271 | 0.302774 | 0.290618 | 0.287135 | 0.305996 | 0.337682 | 0.374542 | 0.393840 | 0.441923 | 0.482510 | 0.556603 | 0.432505 | 0.360489 | 0.401447 | 0.412343 | 0.377166 | 0.331028 | 0.310959 | 0.379834 | 0.411295 | 0.279917 | 0.229788 | 0.224936 | 0.226901 | 0.212967 | 0.213583 | 0.214916 | 0.222153 | 0.239057 | 0.240319 | 0.229544 | 0.230291 | 0.211001 | 0.219294 | 0.239086 | 0.292082 | 0.522186 | 0.687660 | 0.718237 | 0.529102 | 0.384954 | 0.300417 | 0.249090 | 0.211468 | 0.226321 | 0.255283 | 0.243445 | 0.240227 | 0.247669 | 0.261780 | 0.231143 | 0.217773 | 0.210937 | 0.212060 | 0.211159 | 0.218137 | 0.254946 | 0.300474 | 0.296309 | 0.277671 | 0.284328 | 0.281355 | 0.256018 | 0.229403 | 0.212619 | 0.213822 | 0.217228 | 0.216390 | 0.213685 | 0.235074 | 0.242967 | 0.240139 | 0.223917 | 0.217118 | 0.213323 | 0.215014 | 0.212356 | 0.218393 | 0.235768 | 0.263930 | 0.239054 | 0.224474 | 0.211181 | 0.211602 | 0.224776 | 0.232796 | 0.226698 | 0.211176 | 0.226833 | 0.236949 | 0.242478 | 0.244363 | 0.241591 | 0.221022 | 0.211220 | 0.216821 | 0.211252 | 0.214509 | 0.228886 | 0.252354 | 0.315448 | 0.530354 | 0.713405 | 0.634105 | 0.670576 | 0.698367 | 0.799475 | 0.620889 | 0.465113 | 0.371818 | 0.361951 | 0.353508 | 0.375791 | 0.328354 | 0.332805 | 0.357405 | 0.398955 | 0.370797 | 0.327555 | 0.242125 | 0.222026 | 0.216200 | 0.210806 | 0.211013 | 0.217309 | 0.242002 | 0.293332 | 0.368481 | 0.426257 | 0.675414 | 1.359984 | 2.644552 | 2.902781 | 4.001992 | 3.585537 | 2.396022 | 0.850717 | 0.440272 | 0.369930 | 0.404268 | 0.363346 | 0.328697 | 0.373257 | 0.422132 | 0.469909 | 0.594530 | 0.573366 | 0.555378 | 0.431391 | 0.394865 | 0.401324 | 0.341611 | 0.314105 | 0.416001 | 0.667110 | 1.340018 | 2.610350 | 8.601966 | 13.448839 | 7.085317 | 3.749143 | 2.355294 | 1.800785 | 1.265902 | 0.691973 | 0.498605 | 0.446718 | 0.468915 | 0.492085 | 0.435888 | 0.278426 | 0.224442 | 0.215810 | 0.216605 | 0.211050 | 0.210816 | 0.216699 | 0.307687 | 0.532085 | 0.630512 | 0.571270 | 0.586211 | 0.651352 | 0.503817 |
| left posterior | 0.239734 | 0.258506 | 0.265628 | 0.234005 | 0.226362 | 0.225248 | 0.214111 | 0.216358 | 0.212807 | 0.211590 | 0.226076 | 0.316488 | 0.448159 | 0.397342 | 0.379278 | 0.291065 | 0.296582 | 0.283160 | 0.279812 | 0.216202 | 0.212231 | 0.247834 | 0.334674 | 0.314945 | 0.281688 | 0.286549 | 0.330870 | 0.341472 | 0.290329 | 0.219945 | 0.212280 | 0.221469 | 0.259820 | 0.224484 | 0.210819 | 0.214099 | 0.228602 | 0.251731 | 0.282785 | 0.323373 | 0.318954 | 0.365792 | 0.264054 | 0.229910 | 0.213897 | 0.212393 | 0.211113 | 0.211363 | 0.211903 | 0.210891 | 0.211566 | 0.221450 | 0.253140 | 0.328501 | 0.460946 | 0.583711 | 0.701243 | 0.608962 | 0.389194 | 0.210853 | 0.461795 | 4.856331 | 76.084264 | 1939.574582 | 16334.329191 | 16607.178123 | 1722.291013 | 374.632821 | 118.708656 | 46.342772 | 13.098001 | 3.516349 | 1.188985 | 0.420021 | 0.226561 | 0.217965 | 0.354778 | 0.632906 | 1.013984 | 1.860467 | 1.958849 | 2.039884 | 2.105167 | 1.406048 | 1.007473 | 0.532451 | 0.284327 | 0.220768 | 0.227971 | 0.332794 | 0.635167 | 1.367278 | 2.647535 | 3.608907 | 5.495105 | 5.696860 | 9.378962 | 8.273844 | 6.676961 | 4.144047 | 2.571433 | 1.470774 | 0.953649 | 0.603147 | 0.541829 | 0.393555 | 0.325865 | 0.287000 | 0.280719 | 0.288766 | 0.301968 | 0.266950 | 0.289216 | 0.312577 | 0.346887 | 0.352538 | 0.289090 | 0.273520 | 0.314460 | 0.347644 | 0.353912 | 0.345214 | 0.343525 | 0.390326 | 0.375293 | 0.353998 | 0.406287 | 0.457996 | 0.523623 | 0.560507 | 0.608755 | 0.672010 | 0.689220 | 0.576309 | 0.610064 | 0.648477 | 0.642303 | 0.543488 | 0.565301 | 0.619140 | 0.589752 | 0.515525 | 0.385153 | 0.422891 | 0.516336 | 0.678152 | 0.748199 | 0.822666 | 0.764180 | 0.966907 | 1.126103 | 1.171156 | 1.071304 | 0.881013 | 0.648269 | 0.795878 | 0.743284 | 0.484449 | 0.347598 | 0.293752 | 0.261672 | 0.278119 | 0.247918 | 0.236437 | 0.230563 | 0.228999 | 0.222034 | 0.216755 | 0.214614 | 0.212910 | 0.222489 | 0.247399 | 0.259645 | 0.266228 | 0.279931 | 0.272403 | 0.264417 | 0.238680 | 0.216687 | 0.211630 | 0.211045 | 0.210914 | 0.210813 | 0.214501 | 0.218677 | 0.224206 | 0.249676 | 0.299453 | 0.396449 | 0.442174 | 0.446529 | 0.431879 | 0.414037 | 0.306684 | 0.253745 | 0.232702 | 0.228466 | 0.222741 | 0.221670 | 0.227998 | 0.277450 | 0.341197 | 0.379601 | 0.501179 | 0.739689 | 1.168079 | 1.310505 | 1.284271 | 1.092052 | 1.124371 | 0.828863 | 0.755952 | 0.823746 | 1.200471 | 1.988301 | 3.855529 | 7.129537 | 14.443341 | 13.873518 | 6.808018 | 2.988166 | 1.716936 | 0.985489 | 0.689269 | 0.483958 | 0.409597 | 0.372037 | 0.336549 | 0.309219 | 0.326563 | 0.343684 | 0.410575 | 0.454771 | 0.497273 | 0.536534 | 0.650221 | 0.639172 | 0.671284 | 0.674640 | 0.946534 | 0.946366 | 1.186059 | 1.193287 | 1.208149 | 0.936146 | 0.673865 | 0.476290 | 0.431576 | 0.388741 | 0.326764 | 0.365319 | 0.440140 | 0.408738 | 0.384173 | 0.504784 | 0.659133 | 0.934785 | 0.942912 | 0.912987 | 1.373402 | 1.299660 | 0.921466 | 0.693867 | 0.546978 | 0.599209 | 0.565795 | 0.452439 | 0.384913 | 0.357067 | 0.457422 | 0.840384 | 1.364050 | 2.420212 | 3.998824 | 5.708138 | 5.184403 | 2.960270 | 2.267419 | 1.559900 | 1.287519 |
| right posterior | 0.236079 | 0.217483 | 0.210903 | 0.212039 | 0.245327 | 0.303181 | 0.292922 | 0.262761 | 0.241883 | 0.220128 | 0.211533 | 0.214323 | 0.234211 | 0.289132 | 0.444619 | 0.900862 | 0.686626 | 0.473773 | 0.326351 | 0.276052 | 0.276212 | 0.217226 | 0.213085 | 0.301740 | 0.380171 | 0.420647 | 0.457406 | 0.559973 | 0.824986 | 0.585819 | 0.332358 | 0.258184 | 0.216193 | 0.214518 | 0.264650 | 0.401091 | 0.417647 | 0.361707 | 0.350417 | 0.316371 | 0.249857 | 0.251083 | 0.246447 | 0.281146 | 0.329933 | 0.424530 | 0.397857 | 0.335956 | 0.252555 | 0.216170 | 0.210900 | 0.222973 | 0.314758 | 0.410831 | 0.458326 | 0.339506 | 0.225313 | 0.218866 | 0.338226 | 1.335120 | 11.487051 | 134.960871 | 1389.688142 | 10476.024925 | 39217.118296 | 65776.052577 | 36409.075402 | 4575.402282 | 982.502253 | 446.181557 | 181.176252 | 44.435207 | 9.828370 | 3.221338 | 1.936420 | 1.558297 | 0.745634 | 0.283572 | 0.214114 | 0.211475 | 0.221196 | 0.239150 | 0.263359 | 0.270541 | 0.235320 | 0.213578 | 0.210843 | 0.214926 | 0.234945 | 0.267380 | 0.289891 | 0.328361 | 0.380155 | 0.574421 | 0.971003 | 1.357424 | 1.371659 | 1.603595 | 1.681454 | 1.631702 | 1.541369 | 1.426121 | 1.108358 | 0.891350 | 0.582436 | 0.420353 | 0.345977 | 0.301919 | 0.264675 | 0.247297 | 0.239204 | 0.261540 | 0.297613 | 0.358859 | 0.361206 | 0.328141 | 0.340143 | 0.393729 | 0.486072 | 0.504801 | 0.492229 | 0.423472 | 0.367371 | 0.295814 | 0.259459 | 0.245812 | 0.240644 | 0.231955 | 0.237526 | 0.257893 | 0.312008 | 0.359733 | 0.393234 | 0.428923 | 0.454600 | 0.597878 | 0.762405 | 0.861798 | 0.695059 | 0.454445 | 0.387190 | 0.390876 | 0.333013 | 0.321430 | 0.308024 | 0.301844 | 0.309841 | 0.280858 | 0.267242 | 0.341172 | 0.339816 | 0.339684 | 0.409487 | 0.497424 | 0.744043 | 0.969696 | 1.185885 | 1.408999 | 0.861154 | 0.480072 | 0.369895 | 0.291077 | 0.251492 | 0.217704 | 0.211903 | 0.212487 | 0.228946 | 0.225337 | 0.226242 | 0.224028 | 0.233080 | 0.238593 | 0.252362 | 0.252289 | 0.262807 | 0.274646 | 0.296340 | 0.307180 | 0.313027 | 0.299926 | 0.284513 | 0.291664 | 0.277765 | 0.256528 | 0.241778 | 0.247199 | 0.241154 | 0.240616 | 0.231105 | 0.233675 | 0.253816 | 0.263706 | 0.262511 | 0.266820 | 0.282723 | 0.354063 | 0.398380 | 0.328814 | 0.284641 | 0.251369 | 0.257497 | 0.246942 | 0.231096 | 0.222193 | 0.221571 | 0.217660 | 0.219508 | 0.223717 | 0.224197 | 0.220747 | 0.218806 | 0.217018 | 0.236593 | 0.274621 | 0.308185 | 0.359993 | 0.493749 | 0.703922 | 1.323509 | 1.152591 | 0.893996 | 0.699783 | 0.616246 | 0.501838 | 0.403497 | 0.309885 | 0.279867 | 0.256393 | 0.241176 | 0.234605 | 0.240300 | 0.258958 | 0.286009 | 0.315767 | 0.342569 | 0.387595 | 0.334148 | 0.268795 | 0.227903 | 0.220080 | 0.245849 | 0.266918 | 0.290319 | 0.393671 | 0.575172 | 0.550994 | 0.372757 | 0.252237 | 0.231325 | 0.211986 | 0.216715 | 0.227296 | 0.222675 | 0.210932 | 0.210806 | 0.213723 | 0.225535 | 0.227092 | 0.235888 | 0.235368 | 0.246756 | 0.246603 | 0.235972 | 0.227892 | 0.234517 | 0.249568 | 0.253356 | 0.232901 | 0.228565 | 0.214657 | 0.213031 | 0.230232 | 0.252932 | 0.287812 | 0.286578 | 0.279304 | 0.257689 | 0.244580 | 0.237707 | 0.238910 |
| all electrodes | 2.588068 | 1.743544 | 1.121650 | 1.037380 | 0.589129 | 0.416763 | 0.271413 | 0.271738 | 0.309356 | 0.361559 | 0.388534 | 0.392865 | 0.372133 | 0.401793 | 0.419845 | 0.326029 | 0.221786 | 0.214303 | 0.217043 | 0.210839 | 0.222890 | 0.225173 | 0.232229 | 0.348366 | 0.518719 | 0.562292 | 0.424436 | 0.435377 | 0.508020 | 0.358358 | 0.231501 | 0.217000 | 0.212463 | 0.213009 | 0.214814 | 0.276637 | 0.252377 | 0.213735 | 0.211278 | 0.222465 | 0.226647 | 0.284284 | 0.366452 | 0.286434 | 0.235894 | 0.237439 | 0.213124 | 0.217980 | 0.212015 | 0.211926 | 0.210814 | 0.212861 | 0.211233 | 0.219388 | 0.223160 | 0.210930 | 0.237127 | 0.393490 | 0.708195 | 2.278347 | 18.813191 | 242.621308 | 2279.026773 | 47920.356491 | 211282.660804 | 414270.718664 | 165275.756368 | 32196.412925 | 6440.828545 | 1590.709843 | 161.151159 | 13.994405 | 2.083739 | 0.474657 | 0.246409 | 0.212359 | 0.239847 | 0.295913 | 0.333426 | 0.322469 | 0.282671 | 0.239842 | 0.227971 | 0.227245 | 0.223374 | 0.215950 | 0.216711 | 0.211527 | 0.212901 | 0.218924 | 0.263920 | 0.399435 | 0.487496 | 0.545981 | 0.512046 | 0.545873 | 0.589941 | 0.493685 | 0.474667 | 0.477031 | 0.443768 | 0.508623 | 0.424917 | 0.356314 | 0.380019 | 0.364665 | 0.319054 | 0.325606 | 0.287399 | 0.253006 | 0.241453 | 0.225704 | 0.218763 | 0.226934 | 0.239975 | 0.256821 | 0.305375 | 0.427701 | 0.714562 | 1.109333 | 1.133180 | 0.838381 | 0.556944 | 0.594419 | 0.534558 | 0.398188 | 0.331617 | 0.344371 | 0.322049 | 0.371332 | 0.315273 | 0.312416 | 0.298137 | 0.326343 | 0.304754 | 0.358854 | 0.388897 | 0.347797 | 0.341924 | 0.456459 | 0.362245 | 0.379977 | 0.331679 | 0.299540 | 0.303270 | 0.305663 | 0.270542 | 0.276020 | 0.267866 | 0.267345 | 0.265275 | 0.251404 | 0.226135 | 0.212581 | 0.211107 | 0.210909 | 0.213180 | 0.232023 | 0.252977 | 0.242084 | 0.225224 | 0.211953 | 0.215294 | 0.219139 | 0.214461 | 0.215290 | 0.216023 | 0.216453 | 0.228119 | 0.226217 | 0.213400 | 0.213530 | 0.211465 | 0.214993 | 0.225959 | 0.225261 | 0.219084 | 0.233319 | 0.224679 | 0.210843 | 0.240186 | 0.331857 | 0.388685 | 0.386776 | 0.387819 | 0.329132 | 0.262798 | 0.220624 | 0.210957 | 0.210890 | 0.215217 | 0.243615 | 0.285317 | 0.317062 | 0.328851 | 0.378578 | 0.356126 | 0.269702 | 0.241321 | 0.257252 | 0.254033 | 0.244554 | 0.229558 | 0.215735 | 0.219058 | 0.219719 | 0.225690 | 0.237563 | 0.286916 | 0.388856 | 0.532828 | 0.584788 | 0.685575 | 0.824058 | 1.218320 | 1.372005 | 1.218016 | 1.234619 | 1.107068 | 0.996663 | 0.814624 | 0.700810 | 0.704153 | 0.859869 | 0.743563 | 0.673367 | 0.532526 | 0.441310 | 0.359293 | 0.294895 | 0.268316 | 0.286107 | 0.318383 | 0.344745 | 0.330201 | 0.305935 | 0.296959 | 0.250342 | 0.232503 | 0.228644 | 0.239075 | 0.260449 | 0.289676 | 0.294064 | 0.323094 | 0.357726 | 0.387454 | 0.404470 | 0.425554 | 0.432129 | 0.438850 | 0.418872 | 0.405109 | 0.441977 | 0.426186 | 0.450222 | 0.484479 | 0.651399 | 0.937648 | 1.260343 | 1.105554 | 1.092593 | 0.940876 | 0.677277 | 0.511468 | 0.410148 | 0.363602 | 0.339183 | 0.334231 | 0.309325 | 0.328067 | 0.320227 | 0.287258 | 0.280310 | 0.285226 | 0.259683 | 0.271597 | 0.275537 | 0.283485 | 0.337831 |

Searchlight, spatiotemporal cluster permutation test

|  | start time | stop time | peak time | peak channel | cluster p | peak Cohen's d | direction |
| --- | --- | --- | --- | --- | --- | --- | --- |
| #1 | 85 | 180 | 140 | P8 | 0.0385 | 1.454276 | positive |

S) real to emoji faces - angry vs sad

  
|  | time window | peak latency | cluster *p* | peak Cohen's *d* |  | | | |
| **all electrodes** |  | | | |  | | | |
|  | | | | | | | | |

Time-resolved classification, cluster permutation tests

|  | **left hemisphere** | | | | **right hemisphere** | | | |
|  | time window | peak latency | cluster *p* | peak Cohen's *d* | time window | peak latency | cluster *p* | peak Cohen's *d* |
| **anterior** |  | | | |  | | | |
| **central** |  | | | |  | | | |
| **posterior** |  | | | |  | | | |

  

Time-resolved classification, Bayesian statistics

|  | -200 | -195 | -190 | -185 | -180 | -175 | -170 | -165 | -160 | -155 | -150 | -145 | -140 | -135 | -130 | -125 | -120 | -115 | -110 | -105 | -100 | -95 | -90 | -85 | -80 | -75 | -70 | -65 | -60 | -55 | -50 | -45 | -40 | -35 | -30 | -25 | -20 | -15 | -10 | -5 | 0 | 5 | 10 | 15 | 20 | 25 | 30 | 35 | 40 | 45 | 50 | 55 | 60 | 65 | 70 | 75 | 80 | 85 | 90 | 95 | 100 | 105 | 110 | 115 | 120 | 125 | 130 | 135 | 140 | 145 | 150 | 155 | 160 | 165 | 170 | 175 | 180 | 185 | 190 | 195 | 200 | 205 | 210 | 215 | 220 | 225 | 230 | 235 | 240 | 245 | 250 | 255 | 260 | 265 | 270 | 275 | 280 | 285 | 290 | 295 | 300 | 305 | 310 | 315 | 320 | 325 | 330 | 335 | 340 | 345 | 350 | 355 | 360 | 365 | 370 | 375 | 380 | 385 | 390 | 395 | 400 | 405 | 410 | 415 | 420 | 425 | 430 | 435 | 440 | 445 | 450 | 455 | 460 | 465 | 470 | 475 | 480 | 485 | 490 | 495 | 500 | 505 | 510 | 515 | 520 | 525 | 530 | 535 | 540 | 545 | 550 | 555 | 560 | 565 | 570 | 575 | 580 | 585 | 590 | 595 | 600 | 605 | 610 | 615 | 620 | 625 | 630 | 635 | 640 | 645 | 650 | 655 | 660 | 665 | 670 | 675 | 680 | 685 | 690 | 695 | 700 | 705 | 710 | 715 | 720 | 725 | 730 | 735 | 740 | 745 | 750 | 755 | 760 | 765 | 770 | 775 | 780 | 785 | 790 | 795 | 800 | 805 | 810 | 815 | 820 | 825 | 830 | 835 | 840 | 845 | 850 | 855 | 860 | 865 | 870 | 875 | 880 | 885 | 890 | 895 | 900 | 905 | 910 | 915 | 920 | 925 | 930 | 935 | 940 | 945 | 950 | 955 | 960 | 965 | 970 | 975 | 980 | 985 | 990 | 995 | 1000 | 1005 | 1010 | 1015 | 1020 | 1025 | 1030 | 1035 | 1040 | 1045 | 1050 | 1055 | 1060 | 1065 | 1070 | 1075 | 1080 | 1085 | 1090 | 1095 | 1100 | 1105 | 1110 | 1115 | 1120 | 1125 | 1130 | 1135 | 1140 | 1145 | 1150 | 1155 | 1160 | 1165 | 1170 | 1175 | 1180 | 1185 | 1190 | 1195 |
| --- | --- | --- | --- | --- | --- | --- | --- | --- | --- | --- | --- | --- | --- | --- | --- | --- | --- | --- | --- | --- | --- | --- | --- | --- | --- | --- | --- | --- | --- | --- | --- | --- | --- | --- | --- | --- | --- | --- | --- | --- | --- | --- | --- | --- | --- | --- | --- | --- | --- | --- | --- | --- | --- | --- | --- | --- | --- | --- | --- | --- | --- | --- | --- | --- | --- | --- | --- | --- | --- | --- | --- | --- | --- | --- | --- | --- | --- | --- | --- | --- | --- | --- | --- | --- | --- | --- | --- | --- | --- | --- | --- | --- | --- | --- | --- | --- | --- | --- | --- | --- | --- | --- | --- | --- | --- | --- | --- | --- | --- | --- | --- | --- | --- | --- | --- | --- | --- | --- | --- | --- | --- | --- | --- | --- | --- | --- | --- | --- | --- | --- | --- | --- | --- | --- | --- | --- | --- | --- | --- | --- | --- | --- | --- | --- | --- | --- | --- | --- | --- | --- | --- | --- | --- | --- | --- | --- | --- | --- | --- | --- | --- | --- | --- | --- | --- | --- | --- | --- | --- | --- | --- | --- | --- | --- | --- | --- | --- | --- | --- | --- | --- | --- | --- | --- | --- | --- | --- | --- | --- | --- | --- | --- | --- | --- | --- | --- | --- | --- | --- | --- | --- | --- | --- | --- | --- | --- | --- | --- | --- | --- | --- | --- | --- | --- | --- | --- | --- | --- | --- | --- | --- | --- | --- | --- | --- | --- | --- | --- | --- | --- | --- | --- | --- | --- | --- | --- | --- | --- | --- | --- | --- | --- | --- | --- | --- | --- | --- | --- | --- | --- | --- | --- | --- | --- | --- | --- | --- | --- | --- | --- | --- | --- | --- | --- | --- | --- | --- | --- | --- | --- | --- | --- | --- | --- | --- | --- | --- | --- | --- | --- |
| left anterior | 0.242699 | 0.222593 | 0.229675 | 0.225467 | 0.235987 | 0.215154 | 0.221398 | 0.282246 | 0.333005 | 0.619194 | 0.575266 | 0.394119 | 0.263632 | 0.210918 | 0.260623 | 0.455977 | 2.925283 | 5.172306 | 7.716823 | 2.764553 | 0.531090 | 0.359533 | 0.320541 | 0.231428 | 0.210824 | 0.211451 | 0.225673 | 0.236642 | 0.227528 | 0.238390 | 0.227976 | 0.212447 | 0.216088 | 0.212041 | 0.211139 | 0.210827 | 0.229205 | 0.294634 | 0.507303 | 0.963666 | 1.841848 | 3.899877 | 3.338906 | 0.965628 | 0.673876 | 0.361985 | 0.262316 | 0.217718 | 0.224020 | 0.296673 | 0.274482 | 0.381253 | 0.287106 | 0.238364 | 0.223556 | 0.226733 | 0.221128 | 0.237237 | 0.237549 | 0.280709 | 0.347929 | 0.408694 | 0.280610 | 0.229397 | 0.211957 | 0.286527 | 0.412968 | 0.297059 | 0.251545 | 0.211371 | 0.214524 | 0.240674 | 0.247318 | 0.271543 | 0.212414 | 0.220092 | 0.233002 | 0.213951 | 0.210806 | 0.228178 | 0.250026 | 0.287104 | 0.301385 | 0.266308 | 0.223924 | 0.211677 | 0.242311 | 0.238434 | 0.210806 | 0.241184 | 0.321861 | 0.492854 | 0.806343 | 1.534007 | 2.040747 | 1.275651 | 0.764676 | 0.566603 | 0.321184 | 0.252718 | 0.234612 | 0.235040 | 0.232185 | 0.234539 | 0.231020 | 0.243047 | 0.222516 | 0.213678 | 0.297051 | 0.413515 | 0.572370 | 0.779378 | 0.846456 | 0.605113 | 0.467631 | 0.339860 | 0.247729 | 0.240361 | 0.233492 | 0.235907 | 0.256525 | 0.226607 | 0.213424 | 0.212509 | 0.211721 | 0.212497 | 0.211308 | 0.211384 | 0.211499 | 0.214455 | 0.222087 | 0.227662 | 0.257371 | 0.267899 | 0.287026 | 0.363381 | 0.273631 | 0.253510 | 0.227186 | 0.215667 | 0.211759 | 0.211051 | 0.212379 | 0.213356 | 0.227645 | 0.252248 | 0.275096 | 0.281794 | 0.291279 | 0.318772 | 0.293317 | 0.243653 | 0.214112 | 0.212099 | 0.210813 | 0.211076 | 0.210945 | 0.213304 | 0.215830 | 0.215003 | 0.211819 | 0.217218 | 0.221362 | 0.226269 | 0.223623 | 0.228418 | 0.239013 | 0.253427 | 0.303793 | 0.331230 | 0.362655 | 0.340624 | 0.280126 | 0.264020 | 0.266036 | 0.239039 | 0.232275 | 0.235816 | 0.236601 | 0.244208 | 0.264552 | 0.292119 | 0.302572 | 0.314584 | 0.305414 | 0.409347 | 0.528200 | 0.547370 | 0.491482 | 0.494084 | 0.502168 | 0.615452 | 0.531143 | 0.418076 | 0.332601 | 0.245029 | 0.211104 | 0.232643 | 0.301432 | 0.300749 | 0.285004 | 0.279904 | 0.249699 | 0.236511 | 0.244000 | 0.257595 | 0.282177 | 0.291224 | 0.303803 | 0.305274 | 0.312922 | 0.283989 | 0.285997 | 0.294007 | 0.315934 | 0.267892 | 0.246119 | 0.238489 | 0.242930 | 0.251921 | 0.235507 | 0.215429 | 0.215264 | 0.211302 | 0.210833 | 0.210830 | 0.210859 | 0.211769 | 0.213230 | 0.218644 | 0.237105 | 0.248217 | 0.246273 | 0.245266 | 0.231797 | 0.246783 | 0.241100 | 0.237522 | 0.241655 | 0.240342 | 0.224344 | 0.214607 | 0.228817 | 0.269615 | 0.306621 | 0.335664 | 0.377546 | 0.410204 | 0.336740 | 0.229257 | 0.212044 | 0.211718 | 0.210990 | 0.212710 | 0.212157 | 0.220700 | 0.212311 | 0.212206 | 0.219245 | 0.237803 | 0.240872 | 0.218304 | 0.211739 | 0.216613 | 0.223321 | 0.219653 | 0.217352 | 0.217727 | 0.225692 | 0.236305 | 0.219515 | 0.212123 | 0.210896 | 0.213482 | 0.219569 | 0.231140 | 0.237240 | 0.232847 | 0.238389 | 0.249436 |
| right anterior | 0.213280 | 0.214730 | 0.219446 | 0.222609 | 0.218887 | 0.213607 | 0.211058 | 0.210877 | 0.222034 | 0.324528 | 0.499978 | 0.569667 | 0.434051 | 0.295349 | 0.227620 | 0.225965 | 0.222772 | 0.241583 | 0.239017 | 0.210832 | 0.213195 | 0.219971 | 0.211102 | 0.210961 | 0.221190 | 0.254947 | 0.432623 | 0.407664 | 0.658953 | 1.118482 | 0.782874 | 0.477470 | 0.372953 | 0.542042 | 1.402294 | 0.761245 | 0.364727 | 0.277441 | 0.241053 | 0.212638 | 0.228648 | 0.235112 | 0.221151 | 0.211894 | 0.228908 | 0.298611 | 0.588634 | 0.901519 | 1.261989 | 1.216297 | 0.627603 | 0.407644 | 0.280987 | 0.222587 | 0.215846 | 0.217646 | 0.224050 | 0.309520 | 0.576565 | 0.797476 | 1.448976 | 1.431154 | 0.748865 | 0.529746 | 0.460512 | 0.381263 | 0.407645 | 0.470353 | 0.679355 | 1.150329 | 2.945009 | 1.467962 | 0.548647 | 0.273829 | 0.218171 | 0.216345 | 0.307047 | 0.639339 | 0.527338 | 0.351113 | 0.231042 | 0.241066 | 0.300221 | 0.272799 | 0.264110 | 0.274071 | 0.290010 | 0.428055 | 0.446005 | 0.316692 | 0.234504 | 0.212567 | 0.211600 | 0.222204 | 0.229706 | 0.239004 | 0.246771 | 0.223713 | 0.214303 | 0.210969 | 0.226730 | 0.226113 | 0.220916 | 0.224564 | 0.216788 | 0.229996 | 0.258363 | 0.295088 | 0.350397 | 0.394587 | 0.511439 | 0.711032 | 0.854870 | 0.911543 | 0.741524 | 0.675727 | 0.941089 | 1.002196 | 0.842191 | 0.522150 | 0.293054 | 0.240329 | 0.216142 | 0.211362 | 0.220317 | 0.220643 | 0.212965 | 0.210903 | 0.212481 | 0.212441 | 0.212414 | 0.224405 | 0.217365 | 0.220296 | 0.238961 | 0.251778 | 0.267164 | 0.299775 | 0.298132 | 0.360766 | 0.320855 | 0.299926 | 0.319273 | 0.318110 | 0.259946 | 0.274179 | 0.252724 | 0.257263 | 0.246265 | 0.227737 | 0.211070 | 0.211686 | 0.211604 | 0.213848 | 0.242092 | 0.261945 | 0.297266 | 0.372828 | 0.349725 | 0.273727 | 0.234903 | 0.218908 | 0.249551 | 0.248429 | 0.274320 | 0.283489 | 0.339634 | 0.472070 | 0.651586 | 0.486903 | 0.814402 | 0.713516 | 0.714894 | 0.581979 | 0.365279 | 0.308255 | 0.300457 | 0.249698 | 0.240759 | 0.251903 | 0.276141 | 0.370156 | 0.460871 | 0.434235 | 0.462719 | 0.470223 | 0.413491 | 0.371473 | 0.311647 | 0.243532 | 0.229454 | 0.216528 | 0.210970 | 0.210833 | 0.210814 | 0.211613 | 0.211119 | 0.214066 | 0.221538 | 0.217871 | 0.228449 | 0.222248 | 0.224332 | 0.228171 | 0.227379 | 0.227818 | 0.246615 | 0.249738 | 0.279923 | 0.262450 | 0.256153 | 0.242735 | 0.231152 | 0.222414 | 0.220444 | 0.238118 | 0.257032 | 0.266240 | 0.242081 | 0.227862 | 0.212438 | 0.211290 | 0.228994 | 0.264892 | 0.302976 | 0.373951 | 0.418828 | 0.424773 | 0.387299 | 0.320252 | 0.274492 | 0.236811 | 0.211832 | 0.212495 | 0.218697 | 0.213864 | 0.210965 | 0.210816 | 0.211259 | 0.215482 | 0.218016 | 0.217045 | 0.218166 | 0.225221 | 0.257882 | 0.313609 | 0.354375 | 0.430193 | 0.511496 | 0.531843 | 0.490447 | 0.415561 | 0.351563 | 0.319925 | 0.288214 | 0.262314 | 0.237922 | 0.233383 | 0.236009 | 0.248124 | 0.256502 | 0.304360 | 0.319501 | 0.339816 | 0.377691 | 0.411179 | 0.371455 | 0.362558 | 0.283285 | 0.291763 | 0.308770 | 0.319191 | 0.320261 | 0.389542 | 0.408993 | 0.441919 | 0.422970 | 0.422868 | 0.381432 | 0.345096 |
| left central | 0.212410 | 0.210898 | 0.213621 | 0.215632 | 0.257115 | 0.273447 | 0.270611 | 0.235634 | 0.238434 | 0.244556 | 0.272986 | 0.224460 | 0.224973 | 0.227691 | 0.227006 | 0.219827 | 0.211089 | 0.211212 | 0.210806 | 0.210885 | 0.210980 | 0.228134 | 0.267249 | 0.240803 | 0.220817 | 0.215771 | 0.211669 | 0.216604 | 0.226655 | 0.277985 | 0.353887 | 0.384582 | 0.318998 | 0.310054 | 0.338902 | 0.602791 | 1.039092 | 0.757589 | 0.919029 | 0.832274 | 0.822869 | 0.838940 | 0.665010 | 0.276245 | 0.216715 | 0.212274 | 0.227309 | 0.225567 | 0.243897 | 0.259018 | 0.213360 | 0.219599 | 0.221988 | 0.245313 | 0.287294 | 0.562605 | 0.494484 | 0.479834 | 0.471191 | 0.590161 | 0.529168 | 0.378223 | 0.249000 | 0.257548 | 0.331191 | 0.653020 | 1.178579 | 2.973742 | 6.176081 | 7.989727 | 4.421968 | 1.792587 | 0.579179 | 0.348115 | 0.238477 | 0.231033 | 0.348177 | 0.381715 | 0.395175 | 0.420251 | 0.529630 | 0.568459 | 0.454106 | 0.423653 | 0.481569 | 0.529408 | 0.549171 | 0.518425 | 0.400661 | 0.293377 | 0.229110 | 0.210842 | 0.225678 | 0.275043 | 0.387478 | 0.543488 | 0.678138 | 0.819608 | 0.527074 | 0.308082 | 0.213059 | 0.284745 | 0.558624 | 0.758263 | 1.036684 | 0.875876 | 0.692961 | 0.597025 | 0.566025 | 0.912242 | 1.017113 | 1.282897 | 0.843108 | 0.577257 | 0.404282 | 0.289242 | 0.244323 | 0.245694 | 0.250795 | 0.247942 | 0.247883 | 0.238161 | 0.227086 | 0.212792 | 0.210846 | 0.213863 | 0.210806 | 0.210852 | 0.211779 | 0.210858 | 0.211580 | 0.219674 | 0.226329 | 0.236362 | 0.274473 | 0.362955 | 0.379076 | 0.416608 | 0.418403 | 0.332412 | 0.290096 | 0.270080 | 0.248130 | 0.239732 | 0.235435 | 0.247645 | 0.316681 | 0.304836 | 0.352074 | 0.342633 | 0.315784 | 0.349237 | 0.279877 | 0.225731 | 0.216313 | 0.212920 | 0.217245 | 0.213262 | 0.213708 | 0.213513 | 0.211139 | 0.237740 | 0.271879 | 0.265671 | 0.233271 | 0.214616 | 0.210806 | 0.212620 | 0.231527 | 0.266198 | 0.290982 | 0.270094 | 0.255580 | 0.244193 | 0.230797 | 0.234728 | 0.221845 | 0.223177 | 0.219446 | 0.213165 | 0.213930 | 0.225572 | 0.218578 | 0.214774 | 0.215482 | 0.210897 | 0.210955 | 0.211784 | 0.223581 | 0.229071 | 0.246634 | 0.319005 | 0.315107 | 0.347668 | 0.361558 | 0.401539 | 0.404416 | 0.325871 | 0.258175 | 0.233867 | 0.227894 | 0.232545 | 0.221193 | 0.230080 | 0.248184 | 0.265426 | 0.292130 | 0.304425 | 0.289271 | 0.287279 | 0.261303 | 0.270399 | 0.298032 | 0.322404 | 0.398985 | 0.575735 | 0.761749 | 0.719928 | 0.459389 | 0.324392 | 0.259856 | 0.230956 | 0.221510 | 0.221981 | 0.234248 | 0.260586 | 0.311433 | 0.339857 | 0.325901 | 0.317308 | 0.305246 | 0.318120 | 0.346720 | 0.346438 | 0.369317 | 0.489355 | 0.497869 | 0.422958 | 0.336631 | 0.271680 | 0.237008 | 0.229000 | 0.213833 | 0.210831 | 0.213647 | 0.214442 | 0.214013 | 0.213781 | 0.213788 | 0.211793 | 0.211037 | 0.214199 | 0.218391 | 0.220807 | 0.229700 | 0.227833 | 0.223358 | 0.211703 | 0.210890 | 0.213744 | 0.218713 | 0.220833 | 0.218269 | 0.215965 | 0.224413 | 0.221730 | 0.210806 | 0.215893 | 0.229797 | 0.248205 | 0.255381 | 0.236087 | 0.239414 | 0.248698 | 0.262597 | 0.268206 | 0.248135 | 0.244087 | 0.268718 | 0.272119 |
| right central | 0.230869 | 0.238651 | 0.254269 | 0.325172 | 0.272471 | 0.251667 | 0.237374 | 0.240262 | 0.224709 | 0.210806 | 0.258096 | 0.274355 | 0.234545 | 0.221858 | 0.231756 | 0.249240 | 0.235028 | 0.210984 | 0.225212 | 0.230154 | 0.238962 | 0.307134 | 0.367779 | 0.405148 | 0.275846 | 0.239407 | 0.238159 | 0.261710 | 0.266853 | 0.317934 | 0.682205 | 1.376147 | 1.871160 | 0.934810 | 0.704244 | 0.447091 | 0.329811 | 0.232543 | 0.212337 | 0.225712 | 0.211098 | 0.211151 | 0.217861 | 0.213600 | 0.211573 | 0.215088 | 0.211403 | 0.225837 | 0.269348 | 0.460495 | 0.315880 | 0.287874 | 0.263314 | 0.236804 | 0.216801 | 0.211394 | 0.252891 | 0.241184 | 0.236435 | 0.264197 | 0.264242 | 0.238247 | 0.211085 | 0.225274 | 0.231476 | 0.284455 | 0.434746 | 0.765005 | 1.133465 | 1.018320 | 0.660841 | 0.586039 | 0.413254 | 0.311349 | 0.251897 | 0.214298 | 0.218370 | 0.251464 | 0.279418 | 0.309265 | 0.381424 | 0.547239 | 0.548501 | 0.433125 | 0.334060 | 0.288811 | 0.261646 | 0.213928 | 0.223401 | 0.281625 | 0.359539 | 0.424983 | 0.578292 | 0.797334 | 0.960209 | 1.043104 | 1.083048 | 1.515440 | 1.817001 | 1.008061 | 0.569052 | 0.316811 | 0.220309 | 0.248056 | 0.422548 | 0.328100 | 0.219163 | 0.227290 | 0.273187 | 0.394936 | 0.550874 | 0.618575 | 0.535494 | 0.406963 | 0.299064 | 0.279312 | 0.277771 | 0.283756 | 0.303245 | 0.298744 | 0.365741 | 0.470544 | 0.653903 | 0.857517 | 1.499979 | 1.918274 | 1.549345 | 1.925221 | 2.474853 | 3.041692 | 3.555054 | 4.656548 | 4.179557 | 2.249689 | 0.718207 | 0.464869 | 0.397029 | 0.456091 | 0.627292 | 0.765503 | 0.986291 | 1.276698 | 1.051509 | 0.605870 | 0.387976 | 0.279533 | 0.255103 | 0.240825 | 0.233560 | 0.224493 | 0.233176 | 0.220876 | 0.215622 | 0.211276 | 0.213046 | 0.219765 | 0.247377 | 0.283220 | 0.394355 | 0.411106 | 0.468945 | 0.583733 | 0.592520 | 0.551443 | 0.422516 | 0.317719 | 0.314985 | 0.277773 | 0.228599 | 0.214173 | 0.210815 | 0.215325 | 0.215508 | 0.217498 | 0.217967 | 0.212455 | 0.213999 | 0.213817 | 0.211330 | 0.210839 | 0.211363 | 0.212072 | 0.210892 | 0.212468 | 0.210806 | 0.212015 | 0.218091 | 0.224078 | 0.216310 | 0.216285 | 0.220735 | 0.216981 | 0.218267 | 0.216390 | 0.215069 | 0.218611 | 0.223591 | 0.232260 | 0.237592 | 0.224175 | 0.218931 | 0.213478 | 0.215851 | 0.216283 | 0.213989 | 0.216275 | 0.220654 | 0.230774 | 0.225303 | 0.222213 | 0.215991 | 0.212425 | 0.210850 | 0.211131 | 0.212553 | 0.210806 | 0.210902 | 0.211027 | 0.213239 | 0.213290 | 0.213684 | 0.220156 | 0.230964 | 0.229494 | 0.241270 | 0.220681 | 0.214425 | 0.210909 | 0.212980 | 0.212656 | 0.212198 | 0.212446 | 0.211396 | 0.218858 | 0.246397 | 0.259530 | 0.256638 | 0.256050 | 0.231245 | 0.215985 | 0.210986 | 0.214647 | 0.211808 | 0.211339 | 0.211648 | 0.211249 | 0.210816 | 0.216809 | 0.229055 | 0.257695 | 0.277953 | 0.348279 | 0.304835 | 0.274806 | 0.243876 | 0.240823 | 0.242662 | 0.235042 | 0.230504 | 0.235218 | 0.235799 | 0.241714 | 0.232043 | 0.218608 | 0.221352 | 0.210846 | 0.212927 | 0.221444 | 0.241620 | 0.235862 | 0.222217 | 0.214218 | 0.212108 | 0.222077 | 0.252248 | 0.275957 | 0.265736 | 0.259977 | 0.235710 | 0.219463 |
| left posterior | 0.212070 | 0.222130 | 0.254321 | 0.274459 | 0.232101 | 0.213827 | 0.257893 | 0.349981 | 1.072258 | 2.478237 | 2.377960 | 1.255852 | 0.540706 | 0.360304 | 0.255808 | 0.218044 | 0.247061 | 0.229904 | 0.227380 | 0.261405 | 0.241377 | 0.219678 | 0.227971 | 0.231367 | 0.215467 | 0.217148 | 0.261096 | 0.213583 | 0.235200 | 0.302540 | 0.279156 | 0.267550 | 0.231171 | 0.214090 | 0.212940 | 0.221103 | 0.226572 | 0.243450 | 0.284644 | 0.255574 | 0.252983 | 0.281735 | 0.348154 | 0.400816 | 0.311513 | 0.259095 | 0.218891 | 0.213291 | 0.218327 | 0.255391 | 0.269208 | 0.245915 | 0.250848 | 0.329940 | 0.371018 | 0.299324 | 0.231899 | 0.212180 | 0.211606 | 0.225682 | 0.323070 | 1.095477 | 3.394933 | 5.991167 | 5.552080 | 5.027973 | 5.829023 | 4.408994 | 2.655288 | 2.629139 | 2.704323 | 3.103035 | 2.674287 | 1.640508 | 1.115624 | 0.611576 | 0.396294 | 0.285270 | 0.233348 | 0.213998 | 0.210832 | 0.216932 | 0.231126 | 0.249732 | 0.258671 | 0.268420 | 0.257007 | 0.273031 | 0.299211 | 0.296887 | 0.376626 | 0.418403 | 0.325737 | 0.263052 | 0.212337 | 0.217521 | 0.250248 | 0.269199 | 0.240107 | 0.212575 | 0.218071 | 0.250073 | 0.285075 | 0.387113 | 0.401367 | 0.337564 | 0.290578 | 0.291376 | 0.286610 | 0.307694 | 0.316277 | 0.332078 | 0.328053 | 0.353034 | 0.356310 | 0.407657 | 0.463812 | 0.426571 | 0.366190 | 0.376162 | 0.329098 | 0.259863 | 0.214323 | 0.226175 | 0.265753 | 0.322304 | 0.307307 | 0.316541 | 0.306845 | 0.322612 | 0.252502 | 0.235197 | 0.218081 | 0.227337 | 0.222630 | 0.219198 | 0.211829 | 0.212104 | 0.212456 | 0.211714 | 0.212576 | 0.221177 | 0.249422 | 0.290228 | 0.330988 | 0.486181 | 0.789101 | 0.824396 | 0.846903 | 0.681528 | 0.687546 | 0.753080 | 0.619420 | 0.478907 | 0.402434 | 0.365035 | 0.355878 | 0.373705 | 0.351834 | 0.328338 | 0.411280 | 0.588728 | 0.663961 | 0.625462 | 0.509484 | 0.458354 | 0.424105 | 0.295168 | 0.231193 | 0.210928 | 0.214867 | 0.220223 | 0.237413 | 0.251856 | 0.253819 | 0.238872 | 0.231876 | 0.231179 | 0.253108 | 0.237590 | 0.217467 | 0.210847 | 0.218914 | 0.234112 | 0.254471 | 0.288063 | 0.243763 | 0.214778 | 0.211557 | 0.224214 | 0.231292 | 0.249195 | 0.262834 | 0.247267 | 0.229077 | 0.221628 | 0.217547 | 0.216619 | 0.222195 | 0.228750 | 0.222535 | 0.216864 | 0.210889 | 0.216550 | 0.224467 | 0.245086 | 0.262749 | 0.264497 | 0.271395 | 0.284003 | 0.277355 | 0.278916 | 0.260452 | 0.257181 | 0.249291 | 0.226887 | 0.211607 | 0.212640 | 0.221288 | 0.216202 | 0.211176 | 0.211331 | 0.220148 | 0.244938 | 0.321990 | 0.435656 | 0.480274 | 0.463416 | 0.468184 | 0.466326 | 0.384275 | 0.339575 | 0.332077 | 0.343182 | 0.381158 | 0.357217 | 0.330310 | 0.393346 | 0.360919 | 0.330519 | 0.280795 | 0.253215 | 0.264058 | 0.323008 | 0.350376 | 0.465634 | 0.741179 | 0.989795 | 1.170994 | 1.395424 | 1.531946 | 1.746360 | 1.590409 | 1.444643 | 1.512122 | 1.481495 | 1.302773 | 0.994229 | 0.911898 | 0.904654 | 0.861221 | 0.923415 | 0.841379 | 0.758014 | 0.641162 | 0.639542 | 0.818269 | 0.764291 | 0.644172 | 0.490833 | 0.392254 | 0.335938 | 0.289787 | 0.225427 | 0.211698 | 0.213241 | 0.211760 | 0.212695 | 0.213150 | 0.222444 |
| right posterior | 0.224643 | 0.213486 | 0.221923 | 0.252557 | 0.295849 | 0.353781 | 0.371480 | 0.434271 | 0.601791 | 0.518257 | 0.359010 | 0.265722 | 0.245313 | 0.247604 | 0.228974 | 0.214231 | 0.211738 | 0.216117 | 0.212338 | 0.212682 | 0.211048 | 0.214019 | 0.210866 | 0.211678 | 0.227490 | 0.228980 | 0.224557 | 0.212845 | 0.215043 | 0.256474 | 0.280158 | 0.279090 | 0.344756 | 0.473957 | 0.464796 | 0.344669 | 0.276098 | 0.256374 | 0.228180 | 0.226965 | 0.244830 | 0.279129 | 0.311365 | 0.375408 | 0.454670 | 0.434097 | 0.371984 | 0.324733 | 0.341938 | 0.290626 | 0.290394 | 0.256481 | 0.255327 | 0.265550 | 0.235770 | 0.216558 | 0.212397 | 0.228979 | 0.297183 | 0.435959 | 0.990218 | 2.417261 | 5.055698 | 7.889433 | 11.516285 | 19.112096 | 17.317151 | 9.421629 | 4.076797 | 2.105289 | 1.313240 | 0.989232 | 0.552771 | 0.312032 | 0.233498 | 0.212602 | 0.263672 | 0.404492 | 0.838639 | 1.428445 | 1.342801 | 1.050358 | 0.757725 | 0.565116 | 0.378643 | 0.285606 | 0.234001 | 0.217093 | 0.211928 | 0.213000 | 0.226996 | 0.249257 | 0.318645 | 0.510156 | 0.802648 | 1.221843 | 1.105497 | 0.682521 | 0.371939 | 0.240291 | 0.215273 | 0.279511 | 0.361826 | 0.386305 | 0.361826 | 0.310518 | 0.234514 | 0.210845 | 0.226515 | 0.245494 | 0.278663 | 0.291835 | 0.315013 | 0.325009 | 0.328098 | 0.353505 | 0.396324 | 0.441134 | 0.425346 | 0.398628 | 0.312095 | 0.263623 | 0.236676 | 0.231362 | 0.221736 | 0.234365 | 0.229420 | 0.242935 | 0.261246 | 0.294970 | 0.282021 | 0.282807 | 0.259631 | 0.261184 | 0.245533 | 0.233776 | 0.214255 | 0.211227 | 0.212269 | 0.216818 | 0.226371 | 0.238662 | 0.237607 | 0.236320 | 0.226450 | 0.210818 | 0.237607 | 0.296486 | 0.332797 | 0.415870 | 0.445280 | 0.526666 | 0.762423 | 0.892138 | 0.911047 | 1.492490 | 1.533793 | 1.616921 | 1.277821 | 0.455802 | 0.252880 | 0.210968 | 0.226438 | 0.285093 | 0.341713 | 0.357984 | 0.261642 | 0.223371 | 0.211391 | 0.214149 | 0.240891 | 0.259106 | 0.294013 | 0.255829 | 0.233462 | 0.221280 | 0.219610 | 0.213873 | 0.214454 | 0.211640 | 0.216943 | 0.235701 | 0.253502 | 0.293816 | 0.283681 | 0.287257 | 0.256198 | 0.230990 | 0.220622 | 0.213536 | 0.210843 | 0.211201 | 0.210854 | 0.210855 | 0.212803 | 0.219378 | 0.216658 | 0.214558 | 0.216133 | 0.222569 | 0.213977 | 0.210844 | 0.216165 | 0.244209 | 0.272999 | 0.289721 | 0.302312 | 0.274423 | 0.256499 | 0.255666 | 0.229401 | 0.219138 | 0.221810 | 0.219356 | 0.224192 | 0.242392 | 0.257807 | 0.296489 | 0.312411 | 0.302010 | 0.329855 | 0.309710 | 0.284906 | 0.262946 | 0.240809 | 0.241786 | 0.245486 | 0.229637 | 0.227540 | 0.216409 | 0.215102 | 0.219654 | 0.225506 | 0.250994 | 0.318440 | 0.386358 | 0.465812 | 0.412934 | 0.332812 | 0.296198 | 0.253329 | 0.219608 | 0.213470 | 0.211204 | 0.214266 | 0.236666 | 0.285331 | 0.338815 | 0.471464 | 0.547965 | 0.701957 | 0.932490 | 0.894269 | 0.727588 | 0.780127 | 0.870797 | 0.836590 | 0.992623 | 0.698775 | 0.505745 | 0.491301 | 0.379278 | 0.303478 | 0.329887 | 0.310573 | 0.368662 | 0.435535 | 0.473962 | 0.612953 | 0.789610 | 1.069146 | 2.239718 | 3.765651 | 4.833100 | 3.440583 | 1.816956 | 0.988358 | 0.695920 | 0.375404 | 0.254942 |
| all electrodes | 0.240075 | 0.218058 | 0.216892 | 0.222047 | 0.215208 | 0.212858 | 0.214922 | 0.222622 | 0.294227 | 0.873704 | 1.204347 | 0.999577 | 1.111542 | 0.631789 | 0.469989 | 0.265131 | 0.212319 | 0.215800 | 0.218458 | 0.211473 | 0.213016 | 0.216277 | 0.226879 | 0.263285 | 0.277544 | 0.240595 | 0.214804 | 0.233517 | 0.244782 | 0.247734 | 0.214498 | 0.215130 | 0.303363 | 0.625082 | 0.671088 | 0.424700 | 0.339073 | 0.259786 | 0.230363 | 0.233200 | 0.231473 | 0.231728 | 0.313542 | 0.349504 | 0.268756 | 0.238429 | 0.210820 | 0.215301 | 0.225018 | 0.273160 | 0.252255 | 0.224397 | 0.211482 | 0.239923 | 0.233850 | 0.216183 | 0.212755 | 0.210930 | 0.220310 | 0.248422 | 0.407077 | 0.647758 | 0.760229 | 0.883635 | 1.605649 | 1.915570 | 0.830059 | 0.805759 | 0.534363 | 0.425370 | 0.376204 | 0.288810 | 0.215950 | 0.220337 | 0.243856 | 0.437701 | 0.670223 | 0.956279 | 1.275860 | 1.094951 | 1.214447 | 0.708212 | 0.457569 | 0.393683 | 0.358047 | 0.316723 | 0.288499 | 0.262169 | 0.239169 | 0.210987 | 0.238904 | 0.338557 | 0.638273 | 1.307587 | 1.630958 | 1.331494 | 0.815674 | 0.560856 | 0.476648 | 0.338200 | 0.243296 | 0.228779 | 0.217524 | 0.211119 | 0.226585 | 0.265297 | 0.269423 | 0.234854 | 0.230548 | 0.230734 | 0.225015 | 0.217034 | 0.211201 | 0.211734 | 0.210874 | 0.212890 | 0.210875 | 0.211097 | 0.210844 | 0.210817 | 0.212100 | 0.210806 | 0.211203 | 0.212620 | 0.210864 | 0.211006 | 0.211336 | 0.216718 | 0.224370 | 0.211535 | 0.211276 | 0.210816 | 0.211185 | 0.215955 | 0.220963 | 0.234827 | 0.266726 | 0.274008 | 0.354276 | 0.390051 | 0.399369 | 0.354026 | 0.323905 | 0.359160 | 0.422534 | 0.416361 | 0.514064 | 0.611858 | 0.768737 | 0.592451 | 0.500554 | 0.453190 | 0.469621 | 0.455334 | 0.481768 | 0.537189 | 0.629770 | 0.680981 | 0.542925 | 0.411979 | 0.352300 | 0.245533 | 0.211055 | 0.213802 | 0.217308 | 0.230095 | 0.246882 | 0.280555 | 0.292522 | 0.268723 | 0.250649 | 0.262481 | 0.251614 | 0.231450 | 0.214514 | 0.211087 | 0.213214 | 0.230362 | 0.249578 | 0.259934 | 0.281723 | 0.295651 | 0.287825 | 0.259052 | 0.260429 | 0.256316 | 0.246570 | 0.231874 | 0.228576 | 0.236079 | 0.245982 | 0.245445 | 0.229016 | 0.217387 | 0.216950 | 0.214813 | 0.210939 | 0.219902 | 0.240782 | 0.254190 | 0.276213 | 0.290719 | 0.288538 | 0.289614 | 0.289159 | 0.284262 | 0.248625 | 0.220609 | 0.212426 | 0.211897 | 0.211154 | 0.210935 | 0.210811 | 0.213511 | 0.237284 | 0.257849 | 0.254638 | 0.251576 | 0.263779 | 0.258123 | 0.249230 | 0.227774 | 0.219092 | 0.214202 | 0.210853 | 0.210806 | 0.214564 | 0.220347 | 0.223158 | 0.231083 | 0.235813 | 0.260641 | 0.309906 | 0.329893 | 0.354278 | 0.403299 | 0.455838 | 0.637079 | 0.794035 | 0.781709 | 0.835472 | 0.736876 | 0.545359 | 0.481451 | 0.391960 | 0.330239 | 0.320917 | 0.275419 | 0.240674 | 0.237416 | 0.237994 | 0.249101 | 0.274854 | 0.294922 | 0.376141 | 0.613446 | 0.906494 | 1.248233 | 0.784258 | 0.476485 | 0.335901 | 0.277392 | 0.243238 | 0.266061 | 0.324185 | 0.502651 | 0.632270 | 0.481446 | 0.285839 | 0.251674 | 0.222249 | 0.211288 | 0.210827 | 0.212386 | 0.228950 | 0.255554 | 0.288337 | 0.295208 | 0.288022 | 0.274892 |

Searchlight, spatiotemporal cluster permutation test

No significant clusters observed.

T) real to emoji faces - neutral vs angry

  
|  | time window | peak latency | cluster *p* | peak Cohen's *d* |  | | | |  |
| **all electrodes** | 120 - 180 ms | 150 ms | 0.0496 | 1.2391 |  | | | |  |
 330 - 880 ms | 405 ms | 0.0009 | 0.646 |  | | | ||  |  | | | | | | | | |

Time-resolved classification, cluster permutation tests

|  | **left hemisphere** | | | | **right hemisphere** | | | |  |
|  | time window | peak latency | cluster *p* | peak Cohen's *d* | time window | peak latency | cluster *p* | peak Cohen's *d* |  |
| **anterior** |  | | | |  | | | |  |
| **central** |  | | | | 125 - 180 ms | 150 ms | 0.0215 | 1.3931 |  |
  | | | | 350 - 465 ms | 380 ms | 0.0205 | 0.579 ||  | **posterior** | 120 - 180 ms | 150 ms | 0.0298 | 1.7727 | 120 - 180 ms | 150 ms | 0.0448 | 1.5384 |
 255 - 1195 ms | 790 ms | 0.0001 | 1.6548 | 340 - 575 ms | 465 ms | 0.0112 | 0.9051 |  | | | | 595 - 885 ms | 780 ms | 0.0073 | 0.8757 |

  

Time-resolved classification, Bayesian statistics

|  | -200 | -195 | -190 | -185 | -180 | -175 | -170 | -165 | -160 | -155 | -150 | -145 | -140 | -135 | -130 | -125 | -120 | -115 | -110 | -105 | -100 | -95 | -90 | -85 | -80 | -75 | -70 | -65 | -60 | -55 | -50 | -45 | -40 | -35 | -30 | -25 | -20 | -15 | -10 | -5 | 0 | 5 | 10 | 15 | 20 | 25 | 30 | 35 | 40 | 45 | 50 | 55 | 60 | 65 | 70 | 75 | 80 | 85 | 90 | 95 | 100 | 105 | 110 | 115 | 120 | 125 | 130 | 135 | 140 | 145 | 150 | 155 | 160 | 165 | 170 | 175 | 180 | 185 | 190 | 195 | 200 | 205 | 210 | 215 | 220 | 225 | 230 | 235 | 240 | 245 | 250 | 255 | 260 | 265 | 270 | 275 | 280 | 285 | 290 | 295 | 300 | 305 | 310 | 315 | 320 | 325 | 330 | 335 | 340 | 345 | 350 | 355 | 360 | 365 | 370 | 375 | 380 | 385 | 390 | 395 | 400 | 405 | 410 | 415 | 420 | 425 | 430 | 435 | 440 | 445 | 450 | 455 | 460 | 465 | 470 | 475 | 480 | 485 | 490 | 495 | 500 | 505 | 510 | 515 | 520 | 525 | 530 | 535 | 540 | 545 | 550 | 555 | 560 | 565 | 570 | 575 | 580 | 585 | 590 | 595 | 600 | 605 | 610 | 615 | 620 | 625 | 630 | 635 | 640 | 645 | 650 | 655 | 660 | 665 | 670 | 675 | 680 | 685 | 690 | 695 | 700 | 705 | 710 | 715 | 720 | 725 | 730 | 735 | 740 | 745 | 750 | 755 | 760 | 765 | 770 | 775 | 780 | 785 | 790 | 795 | 800 | 805 | 810 | 815 | 820 | 825 | 830 | 835 | 840 | 845 | 850 | 855 | 860 | 865 | 870 | 875 | 880 | 885 | 890 | 895 | 900 | 905 | 910 | 915 | 920 | 925 | 930 | 935 | 940 | 945 | 950 | 955 | 960 | 965 | 970 | 975 | 980 | 985 | 990 | 995 | 1000 | 1005 | 1010 | 1015 | 1020 | 1025 | 1030 | 1035 | 1040 | 1045 | 1050 | 1055 | 1060 | 1065 | 1070 | 1075 | 1080 | 1085 | 1090 | 1095 | 1100 | 1105 | 1110 | 1115 | 1120 | 1125 | 1130 | 1135 | 1140 | 1145 | 1150 | 1155 | 1160 | 1165 | 1170 | 1175 | 1180 | 1185 | 1190 | 1195 |
| --- | --- | --- | --- | --- | --- | --- | --- | --- | --- | --- | --- | --- | --- | --- | --- | --- | --- | --- | --- | --- | --- | --- | --- | --- | --- | --- | --- | --- | --- | --- | --- | --- | --- | --- | --- | --- | --- | --- | --- | --- | --- | --- | --- | --- | --- | --- | --- | --- | --- | --- | --- | --- | --- | --- | --- | --- | --- | --- | --- | --- | --- | --- | --- | --- | --- | --- | --- | --- | --- | --- | --- | --- | --- | --- | --- | --- | --- | --- | --- | --- | --- | --- | --- | --- | --- | --- | --- | --- | --- | --- | --- | --- | --- | --- | --- | --- | --- | --- | --- | --- | --- | --- | --- | --- | --- | --- | --- | --- | --- | --- | --- | --- | --- | --- | --- | --- | --- | --- | --- | --- | --- | --- | --- | --- | --- | --- | --- | --- | --- | --- | --- | --- | --- | --- | --- | --- | --- | --- | --- | --- | --- | --- | --- | --- | --- | --- | --- | --- | --- | --- | --- | --- | --- | --- | --- | --- | --- | --- | --- | --- | --- | --- | --- | --- | --- | --- | --- | --- | --- | --- | --- | --- | --- | --- | --- | --- | --- | --- | --- | --- | --- | --- | --- | --- | --- | --- | --- | --- | --- | --- | --- | --- | --- | --- | --- | --- | --- | --- | --- | --- | --- | --- | --- | --- | --- | --- | --- | --- | --- | --- | --- | --- | --- | --- | --- | --- | --- | --- | --- | --- | --- | --- | --- | --- | --- | --- | --- | --- | --- | --- | --- | --- | --- | --- | --- | --- | --- | --- | --- | --- | --- | --- | --- | --- | --- | --- | --- | --- | --- | --- | --- | --- | --- | --- | --- | --- | --- | --- | --- | --- | --- | --- | --- | --- | --- | --- | --- | --- | --- | --- | --- | --- | --- | --- | --- | --- | --- | --- | --- | --- |
| left anterior | 1.053918 | 1.362059 | 1.296333 | 1.355939 | 1.940925 | 2.332696 | 2.321661 | 1.761980 | 0.800079 | 0.339765 | 0.234182 | 0.210953 | 0.231365 | 0.244239 | 0.345308 | 0.414002 | 0.210806 | 0.216979 | 0.210835 | 0.213301 | 0.216646 | 0.213156 | 0.211643 | 0.228675 | 0.237193 | 0.212437 | 0.269251 | 0.266559 | 0.280946 | 0.339275 | 0.551925 | 0.525741 | 0.757197 | 0.422483 | 0.262005 | 0.212297 | 0.217836 | 0.260664 | 0.232374 | 0.283015 | 0.368248 | 0.423067 | 0.431037 | 0.424474 | 0.383175 | 0.441033 | 0.424173 | 0.683922 | 0.559441 | 0.448585 | 0.359800 | 0.307061 | 0.342288 | 0.383407 | 0.332315 | 0.381151 | 0.319276 | 0.356263 | 1.145332 | 1.979286 | 2.889669 | 4.460690 | 2.674910 | 2.748249 | 0.848460 | 0.229184 | 0.244726 | 0.457874 | 1.344634 | 4.397398e+00 | 1.371166e+01 | 2.368668e+01 | 15.918363 | 7.222761 | 1.827205 | 0.576726 | 0.259800 | 0.216767 | 0.256883 | 0.270813 | 0.290279 | 0.284880 | 0.299817 | 0.357044 | 0.336853 | 0.329893 | 0.363457 | 0.277773 | 0.230546 | 0.211129 | 0.225943 | 0.264174 | 0.405876 | 0.951216 | 1.272101 | 1.580438 | 1.381435 | 1.038224 | 0.798238 | 0.607163 | 0.459951 | 0.406734 | 0.315115 | 0.291528 | 0.317694 | 0.298052 | 0.304624 | 0.295338 | 0.298193 | 0.359694 | 0.366156 | 0.357064 | 0.401831 | 0.413641 | 0.541293 | 0.679372 | 0.516231 | 0.435750 | 0.344993 | 0.339183 | 0.358378 | 0.345942 | 0.330933 | 0.360894 | 0.375879 | 0.394926 | 0.377406 | 0.300942 | 0.269956 | 0.282073 | 0.264273 | 0.243313 | 0.225628 | 0.213678 | 0.211017 | 0.212011 | 0.224447 | 0.228954 | 0.212551 | 0.213008 | 0.219086 | 0.239386 | 0.260738 | 0.250337 | 0.231028 | 0.214811 | 0.210936 | 0.213266 | 0.225384 | 0.249425 | 0.245823 | 0.232529 | 0.230247 | 0.216026 | 0.210956 | 0.210816 | 0.210900 | 0.210912 | 0.210856 | 0.212645 | 0.211116 | 0.210916 | 0.214545 | 0.228417 | 0.217179 | 0.213915 | 0.213438 | 0.246565 | 0.292615 | 0.371832 | 0.386975 | 0.311199 | 0.316267 | 0.274866 | 0.225555 | 0.210907 | 0.212524 | 0.214239 | 0.218731 | 0.218879 | 0.227707 | 0.223541 | 0.211177 | 0.210846 | 0.212999 | 0.216952 | 0.248630 | 0.257999 | 0.284420 | 0.355500 | 0.374088 | 0.313493 | 0.278842 | 2.398671e-01 | 2.284888e-01 | 2.164939e-01 | 0.214720 | 2.320351e-01 | 0.253094 | 0.264860 | 0.283376 | 0.327983 | 0.335760 | 0.310974 | 0.263479 | 0.230062 | 0.224302 | 0.231190 | 0.216324 | 0.210835 | 0.210813 | 0.210978 | 0.210974 | 0.212952 | 0.224563 | 0.228821 | 0.222782 | 0.226592 | 0.224922 | 0.218920 | 0.211062 | 0.210863 | 0.213850 | 0.211130 | 0.210833 | 0.212557 | 0.210837 | 0.214597 | 0.220829 | 0.221374 | 0.256787 | 0.282041 | 0.324537 | 0.304234 | 0.296176 | 0.275796 | 0.268255 | 0.261963 | 0.264720 | 0.256865 | 0.246881 | 0.227358 | 0.217499 | 0.211337 | 0.214865 | 0.221890 | 0.228046 | 0.220616 | 0.222886 | 0.225136 | 0.234063 | 0.226588 | 0.219953 | 0.223200 | 0.234533 | 0.235253 | 0.228096 | 0.223021 | 0.225626 | 0.242443 | 0.265964 | 0.306340 | 0.337899 | 0.349122 | 0.334046 | 0.281556 | 0.258344 | 0.226935 | 0.215847 | 0.210858 | 0.210859 | 0.211176 | 0.212093 | 0.211085 | 0.210894 | 0.211262 | 0.213066 | 0.217983 | 0.229541 | 0.237291 |
| right anterior | 1.031096 | 0.782658 | 0.821230 | 0.789748 | 0.586607 | 0.277876 | 0.238518 | 0.213410 | 0.218560 | 0.260972 | 0.314709 | 0.351082 | 0.338947 | 0.341696 | 0.321219 | 0.247399 | 0.221637 | 0.218547 | 0.235866 | 0.224887 | 0.214601 | 0.213568 | 0.211619 | 0.210928 | 0.220568 | 0.232715 | 0.274637 | 0.413796 | 0.589014 | 0.502042 | 0.633638 | 0.689154 | 0.639673 | 0.413645 | 0.266077 | 0.300346 | 0.326314 | 0.217199 | 0.213847 | 0.226932 | 0.304235 | 0.425360 | 0.508427 | 0.565364 | 0.480422 | 0.589650 | 0.514184 | 0.352889 | 0.218841 | 0.211177 | 0.229650 | 0.275932 | 0.522969 | 0.738819 | 1.350801 | 1.395028 | 1.352620 | 0.846431 | 0.778561 | 0.554073 | 0.329747 | 0.227823 | 0.217001 | 0.256560 | 0.466254 | 1.223448 | 5.916881 | 60.536181 | 329.016729 | 3.409308e+02 | 2.210886e+02 | 8.714219e+01 | 37.321751 | 21.612148 | 4.791589 | 1.978661 | 0.861166 | 0.401962 | 0.345064 | 0.305918 | 0.238881 | 0.215842 | 0.214203 | 0.238293 | 0.260217 | 0.266865 | 0.234255 | 0.210819 | 0.245178 | 0.409225 | 0.940691 | 1.464455 | 2.369002 | 3.530693 | 2.137105 | 1.473057 | 1.149272 | 0.783900 | 0.821030 | 0.899256 | 0.491829 | 0.350247 | 0.281705 | 0.282423 | 0.314214 | 0.350121 | 0.335539 | 0.363172 | 0.575160 | 1.306398 | 2.092039 | 2.983150 | 2.729604 | 3.226978 | 3.468132 | 3.298070 | 2.299149 | 1.958023 | 1.419485 | 1.089780 | 0.918394 | 0.910770 | 0.610135 | 0.440439 | 0.358138 | 0.296571 | 0.278651 | 0.259048 | 0.252158 | 0.248190 | 0.245598 | 0.248737 | 0.265728 | 0.260583 | 0.255298 | 0.225042 | 0.224101 | 0.228951 | 0.217691 | 0.210865 | 0.210819 | 0.211073 | 0.215946 | 0.219744 | 0.216254 | 0.212694 | 0.212266 | 0.211174 | 0.210890 | 0.213809 | 0.225481 | 0.239375 | 0.228300 | 0.237294 | 0.252709 | 0.259126 | 0.255242 | 0.269757 | 0.272124 | 0.277450 | 0.264678 | 0.251772 | 0.259631 | 0.257159 | 0.237439 | 0.217303 | 0.214209 | 0.211153 | 0.211995 | 0.211580 | 0.214471 | 0.220293 | 0.216893 | 0.218562 | 0.235116 | 0.306955 | 0.379225 | 0.470414 | 0.525049 | 0.681788 | 0.817028 | 1.012932 | 1.194964 | 0.911502 | 0.868044 | 0.955740 | 0.886076 | 0.817468 | 0.515459 | 0.348856 | 0.312968 | 0.255392 | 0.238347 | 2.281184e-01 | 2.304288e-01 | 2.519471e-01 | 0.281145 | 2.887063e-01 | 0.359412 | 0.397530 | 0.475004 | 0.410228 | 0.390842 | 0.331225 | 0.293952 | 0.251277 | 0.223470 | 0.211990 | 0.213301 | 0.220660 | 0.248871 | 0.288728 | 0.333402 | 0.360950 | 0.375813 | 0.320218 | 0.264811 | 0.226357 | 0.213119 | 0.211757 | 0.212484 | 0.214382 | 0.210832 | 0.214725 | 0.232239 | 0.255116 | 0.278016 | 0.275444 | 0.257661 | 0.225052 | 0.212892 | 0.211610 | 0.211107 | 0.217090 | 0.216491 | 0.222807 | 0.221299 | 0.225008 | 0.223669 | 0.222784 | 0.220890 | 0.228510 | 0.236389 | 0.281198 | 0.323029 | 0.353290 | 0.466939 | 0.516427 | 0.519971 | 0.427443 | 0.326256 | 0.256907 | 0.229125 | 0.220122 | 0.217147 | 0.253324 | 0.357346 | 0.434791 | 0.607047 | 1.002147 | 1.301070 | 1.773270 | 1.441242 | 1.392817 | 1.826042 | 2.109765 | 1.262041 | 0.640832 | 0.471255 | 0.471693 | 0.451738 | 0.535519 | 0.550344 | 0.644250 | 0.777272 | 0.742376 | 0.807267 | 0.771337 | 0.470686 | 0.372992 |
| left central | 0.220455 | 0.244259 | 0.248389 | 0.211804 | 0.234879 | 0.295829 | 0.289586 | 0.277762 | 0.409733 | 0.810137 | 0.610877 | 0.359132 | 0.338810 | 0.337317 | 0.354579 | 0.235343 | 0.214862 | 0.290204 | 0.216305 | 0.223034 | 0.219747 | 0.211597 | 0.211947 | 0.217061 | 0.211431 | 0.260703 | 0.304715 | 0.335071 | 0.354142 | 0.283249 | 0.264092 | 0.388013 | 0.458174 | 0.666009 | 0.534636 | 0.812383 | 0.993034 | 0.712647 | 0.454507 | 0.250808 | 0.218205 | 0.242801 | 0.252473 | 0.259911 | 0.234935 | 0.211536 | 0.234072 | 0.328954 | 0.588026 | 0.977582 | 1.173364 | 1.006298 | 0.787244 | 0.379849 | 0.286586 | 0.223580 | 0.214974 | 0.234861 | 0.224198 | 0.214413 | 0.230194 | 0.243702 | 0.230929 | 0.210950 | 0.226978 | 0.433109 | 1.841048 | 20.448156 | 221.234537 | 1.970936e+03 | 1.801860e+04 | 4.266473e+04 | 9584.913326 | 851.801923 | 37.759684 | 3.971549 | 0.592033 | 0.278757 | 0.218534 | 0.211397 | 0.227753 | 0.240316 | 0.278935 | 0.289377 | 0.313748 | 0.309046 | 0.270198 | 0.228513 | 0.212363 | 0.280885 | 0.444815 | 1.001913 | 1.955982 | 3.637986 | 4.713527 | 4.222298 | 3.578948 | 3.507195 | 2.673608 | 1.652131 | 0.811332 | 0.463612 | 0.293632 | 0.237172 | 0.214872 | 0.217989 | 0.233328 | 0.240062 | 0.251932 | 0.290141 | 0.351974 | 0.637852 | 0.845525 | 0.775421 | 0.832103 | 1.006614 | 0.844639 | 0.705719 | 0.441702 | 0.333222 | 0.350822 | 0.414035 | 0.487236 | 0.631905 | 0.782960 | 1.028289 | 1.437464 | 1.788996 | 1.711995 | 1.395394 | 1.122208 | 1.110271 | 0.896575 | 0.651863 | 0.557568 | 0.563984 | 0.541020 | 0.593680 | 0.574994 | 0.602291 | 0.536566 | 0.467476 | 0.424374 | 0.366564 | 0.324218 | 0.295360 | 0.252739 | 0.229682 | 0.215906 | 0.215052 | 0.217488 | 0.218048 | 0.215743 | 0.214533 | 0.240915 | 0.305507 | 0.337957 | 0.377723 | 0.338127 | 0.323921 | 0.367294 | 0.385784 | 0.299598 | 0.279887 | 0.305033 | 0.436758 | 0.490961 | 0.716533 | 0.831141 | 1.348662 | 1.255176 | 0.812378 | 0.604308 | 0.732751 | 0.528069 | 0.487433 | 0.530333 | 0.729632 | 0.917637 | 0.739826 | 0.469528 | 0.391778 | 0.313405 | 0.241985 | 0.226206 | 0.212502 | 0.211119 | 0.211009 | 0.216622 | 0.241696 | 0.289953 | 0.282809 | 0.292255 | 3.236826e-01 | 2.864557e-01 | 2.680845e-01 | 0.233521 | 2.336838e-01 | 0.236869 | 0.242209 | 0.239160 | 0.234301 | 0.229555 | 0.226950 | 0.215654 | 0.217398 | 0.215499 | 0.220545 | 0.240570 | 0.256073 | 0.289220 | 0.345872 | 0.305723 | 0.288042 | 0.273094 | 0.274584 | 0.245465 | 0.221012 | 0.210904 | 0.214933 | 0.222970 | 0.235461 | 0.252356 | 0.247954 | 0.241438 | 0.215997 | 0.210870 | 0.210813 | 0.211150 | 0.211056 | 0.213721 | 0.211858 | 0.211416 | 0.217793 | 0.215753 | 0.214050 | 0.210899 | 0.210812 | 0.210945 | 0.213169 | 0.211290 | 0.211024 | 0.213260 | 0.229700 | 0.224863 | 0.229457 | 0.237399 | 0.234879 | 0.240625 | 0.228829 | 0.224713 | 0.231019 | 0.229601 | 0.223088 | 0.222945 | 0.220100 | 0.233005 | 0.233596 | 0.265133 | 0.314277 | 0.337555 | 0.409258 | 0.593818 | 0.635155 | 0.718274 | 0.533321 | 0.396986 | 0.372537 | 0.299769 | 0.251606 | 0.236623 | 0.224252 | 0.215207 | 0.213145 | 0.213288 | 0.215517 | 0.216274 | 0.215483 | 0.212734 | 0.215422 |
| right central | 0.251609 | 0.228965 | 0.258214 | 0.292501 | 0.251847 | 0.237109 | 0.210967 | 0.213902 | 0.212362 | 0.230533 | 0.233410 | 0.211231 | 0.217092 | 0.217731 | 0.211181 | 0.222833 | 0.234842 | 0.211921 | 0.213441 | 0.223122 | 0.278465 | 0.235584 | 0.231260 | 0.266988 | 0.567015 | 4.021093 | 24.767606 | 36.690657 | 20.758283 | 5.167319 | 7.273132 | 0.744780 | 0.319624 | 0.284845 | 0.327301 | 0.442555 | 0.583476 | 0.580920 | 0.789646 | 1.898777 | 2.292640 | 1.844263 | 1.029102 | 0.749575 | 0.618542 | 0.662947 | 0.614433 | 0.348891 | 0.255700 | 0.303355 | 0.491590 | 0.700600 | 0.702427 | 0.790814 | 1.044828 | 1.270139 | 0.800334 | 0.409020 | 0.381175 | 0.317171 | 0.295498 | 0.228027 | 0.211890 | 0.246230 | 0.833125 | 8.850297 | 57.425369 | 1155.276198 | 7531.996586 | 2.663676e+04 | 4.920402e+04 | 3.322062e+04 | 12790.708696 | 2863.338543 | 211.365597 | 17.862606 | 2.411497 | 0.739389 | 0.375663 | 0.273219 | 0.222464 | 0.211524 | 0.233545 | 0.319632 | 0.388170 | 0.412941 | 0.520853 | 0.418044 | 0.286180 | 0.221313 | 0.217155 | 0.274844 | 0.377509 | 0.688131 | 1.229095 | 2.182030 | 4.252357 | 4.852034 | 3.626375 | 1.544108 | 0.918512 | 0.675964 | 0.528731 | 0.420649 | 0.403673 | 0.416472 | 0.532847 | 0.732236 | 0.903775 | 1.212625 | 2.273470 | 3.128211 | 4.687531 | 5.386009 | 4.996183 | 6.289901 | 5.799068 | 3.612775 | 3.517714 | 3.238920 | 3.392043 | 3.703532 | 3.437363 | 3.562520 | 4.026342 | 3.557006 | 2.891059 | 2.482593 | 2.799270 | 2.439786 | 2.380899 | 2.522982 | 2.080391 | 1.661253 | 1.275666 | 0.765775 | 0.666863 | 0.785799 | 0.699781 | 0.632403 | 0.525582 | 0.411890 | 0.388273 | 0.336742 | 0.282334 | 0.247741 | 0.244880 | 0.241193 | 0.245238 | 0.299255 | 0.324525 | 0.400826 | 0.423053 | 0.382187 | 0.507033 | 0.608447 | 0.376454 | 0.295177 | 0.238197 | 0.234624 | 0.253289 | 0.247884 | 0.235146 | 0.237155 | 0.247111 | 0.284296 | 0.274321 | 0.263126 | 0.275119 | 0.392439 | 0.770613 | 1.328293 | 1.359172 | 3.123059 | 4.316995 | 2.973905 | 1.492329 | 0.730872 | 0.498427 | 0.505043 | 0.454084 | 0.385779 | 0.434037 | 0.605407 | 1.008867 | 2.421580 | 3.295062 | 2.706638 | 1.756914 | 0.975450 | 0.490061 | 0.302134 | 0.223750 | 2.118538e-01 | 2.108717e-01 | 2.110549e-01 | 0.210806 | 2.108673e-01 | 0.210806 | 0.221106 | 0.232769 | 0.225551 | 0.221490 | 0.233148 | 0.222095 | 0.212567 | 0.212476 | 0.232612 | 0.248690 | 0.239305 | 0.225944 | 0.211805 | 0.219145 | 0.232728 | 0.237225 | 0.253401 | 0.246667 | 0.230832 | 0.223720 | 0.213797 | 0.212305 | 0.225755 | 0.230187 | 0.225807 | 0.211696 | 0.213516 | 0.224707 | 0.272944 | 0.366558 | 0.479796 | 0.610142 | 0.512611 | 0.409425 | 0.355990 | 0.249850 | 0.228949 | 0.216978 | 0.211228 | 0.211221 | 0.212291 | 0.214217 | 0.211120 | 0.215454 | 0.220978 | 0.221177 | 0.219930 | 0.214095 | 0.211124 | 0.210884 | 0.214136 | 0.232891 | 0.231135 | 0.226799 | 0.217588 | 0.210817 | 0.217325 | 0.227208 | 0.235081 | 0.235174 | 0.227222 | 0.220487 | 0.214139 | 0.210806 | 0.217386 | 0.216986 | 0.222591 | 0.232225 | 0.225126 | 0.216897 | 0.216136 | 0.211756 | 0.211754 | 0.211155 | 0.213067 | 0.212236 | 0.210816 | 0.210956 | 0.211113 | 0.213503 | 0.220215 |
| left posterior | 0.310833 | 0.389343 | 0.317290 | 0.269650 | 0.230230 | 0.211869 | 0.211429 | 0.212576 | 0.210982 | 0.222477 | 0.258654 | 0.532683 | 0.672635 | 0.453264 | 0.364071 | 0.330229 | 0.330977 | 0.354201 | 0.298698 | 0.353858 | 0.371700 | 0.320301 | 0.323472 | 0.310332 | 0.263167 | 0.252001 | 0.242983 | 0.251768 | 0.234946 | 0.232295 | 0.220892 | 0.213103 | 0.210952 | 0.228667 | 0.298333 | 0.282844 | 0.240761 | 0.218892 | 0.210823 | 0.228580 | 0.315005 | 1.185135 | 3.020008 | 2.073343 | 0.836434 | 0.399534 | 0.306020 | 0.273849 | 0.274276 | 0.241446 | 0.231973 | 0.224076 | 0.237557 | 0.237937 | 0.269322 | 0.242890 | 0.254835 | 0.353610 | 0.767954 | 1.035989 | 0.705087 | 0.311416 | 0.212052 | 0.343376 | 3.037085 | 107.591652 | 4090.863831 | 73143.505954 | 670896.375514 | 1.867506e+06 | 2.508243e+06 | 1.693566e+06 | 415565.319282 | 40170.512355 | 2152.957302 | 168.776375 | 8.454582 | 0.851890 | 0.244093 | 0.226432 | 0.356734 | 0.446906 | 0.579700 | 0.680333 | 0.594641 | 0.439645 | 0.331180 | 0.253967 | 0.212013 | 0.228180 | 0.546608 | 3.248828 | 38.816312 | 403.964824 | 3353.004137 | 8263.518942 | 10036.156790 | 2784.219098 | 834.525058 | 308.348759 | 130.608369 | 49.316695 | 34.670704 | 24.980053 | 35.503682 | 30.453682 | 23.258991 | 16.589177 | 9.503811 | 6.681599 | 10.770107 | 15.192804 | 18.509162 | 36.097795 | 31.455516 | 34.487403 | 33.455395 | 28.391459 | 38.281019 | 39.083882 | 23.651027 | 27.870329 | 42.190581 | 55.983556 | 88.492800 | 137.126622 | 143.809704 | 165.363729 | 267.461682 | 330.326869 | 662.438384 | 522.876381 | 346.203884 | 505.729979 | 542.715663 | 699.140123 | 631.496048 | 616.925230 | 836.829610 | 810.201143 | 620.653038 | 751.472739 | 1269.804436 | 3327.048564 | 5774.249889 | 5837.296621 | 9302.957085 | 5071.776618 | 2639.720082 | 838.421373 | 330.699794 | 170.657047 | 88.262238 | 43.173638 | 46.760848 | 57.985798 | 106.202709 | 162.521707 | 127.740880 | 79.820242 | 83.810192 | 63.595362 | 47.374054 | 44.181679 | 49.446185 | 66.511539 | 137.726096 | 141.073993 | 148.204203 | 146.086780 | 169.406140 | 181.762780 | 195.864556 | 153.317471 | 204.055570 | 274.233948 | 339.896873 | 298.836436 | 485.275366 | 711.100944 | 2971.376062 | 14274.825888 | 39459.661422 | 51582.441244 | 92382.879037 | 61340.785720 | 55809.812904 | 40848.258099 | 18439.482058 | 13941.021140 | 23561.685147 | 62135.810438 | 255546.281654 | 1.217911e+06 | 2.304423e+06 | 1.388232e+06 | 752935.828779 | 1.049783e+06 | 772170.848702 | 317449.886883 | 75521.378605 | 35457.168704 | 19755.126623 | 15065.915726 | 7234.071091 | 1672.308297 | 827.404574 | 1071.116171 | 1505.764180 | 2591.471652 | 2913.606177 | 3360.290633 | 1659.653112 | 1107.229241 | 723.349673 | 387.612307 | 84.707750 | 28.959835 | 13.437506 | 7.893599 | 5.291787 | 4.780242 | 5.484040 | 9.018196 | 14.923544 | 18.832398 | 20.823703 | 19.492639 | 16.616051 | 13.757873 | 13.149404 | 8.309558 | 5.388180 | 4.890764 | 4.777224 | 5.487739 | 4.330146 | 4.929377 | 5.449268 | 9.519772 | 14.635897 | 19.058109 | 22.853245 | 32.145317 | 31.708792 | 48.348430 | 124.112758 | 579.302946 | 1297.778519 | 2001.444538 | 1958.606058 | 1521.929020 | 353.316042 | 47.650318 | 6.947935 | 2.782451 | 1.888009 | 1.634054 | 2.189807 | 4.264594 | 19.514175 | 100.179161 | 234.046725 | 320.087586 | 260.295788 | 129.242468 | 60.019446 | 18.217941 | 7.250867 | 6.547542 | 5.912605 | 5.364987 | 4.168378 | 2.922551 | 2.706620 | 3.468518 | 4.215031 | 3.122591 | 2.295043 | 2.415856 |
| right posterior | 0.436877 | 0.630402 | 0.900298 | 1.491628 | 1.439004 | 1.310868 | 0.793520 | 0.474048 | 0.328882 | 0.238401 | 0.217839 | 0.257477 | 0.287172 | 0.283452 | 0.311785 | 0.293666 | 0.213159 | 0.217967 | 0.243683 | 0.210948 | 0.226125 | 0.212193 | 0.212595 | 0.273921 | 0.252860 | 0.270179 | 0.275479 | 0.288852 | 0.275944 | 0.229960 | 0.214035 | 0.212766 | 0.210870 | 0.211932 | 0.210826 | 0.227837 | 0.290105 | 0.297135 | 0.288842 | 0.355116 | 0.341191 | 0.308546 | 0.267223 | 0.256350 | 0.242106 | 0.241678 | 0.223469 | 0.220600 | 0.210816 | 0.211946 | 0.224896 | 0.277503 | 0.277714 | 0.274925 | 0.284323 | 0.336293 | 0.365196 | 0.332324 | 0.293021 | 0.320065 | 0.268893 | 0.211050 | 0.300963 | 0.958463 | 5.540537 | 77.014200 | 1268.066323 | 11414.643973 | 65724.513273 | 1.779910e+05 | 2.321403e+05 | 1.149028e+05 | 22096.172126 | 4261.321891 | 533.507335 | 39.119367 | 2.980659 | 0.551407 | 0.225992 | 0.227463 | 0.357853 | 0.560998 | 0.545282 | 0.495476 | 0.494371 | 0.474911 | 0.449727 | 0.320653 | 0.240915 | 0.211653 | 0.229208 | 0.337750 | 0.912267 | 3.016256 | 8.370119 | 14.883179 | 12.291913 | 6.224341 | 3.232289 | 1.318710 | 0.735796 | 0.421857 | 0.331200 | 0.292458 | 0.316717 | 0.429862 | 0.633492 | 0.918275 | 1.613448 | 2.180517 | 2.956551 | 4.328772 | 5.200272 | 6.720741 | 9.685631 | 15.783951 | 41.705334 | 71.052202 | 51.834070 | 41.359008 | 50.329567 | 42.183835 | 20.087342 | 11.269859 | 8.875799 | 13.216280 | 20.524148 | 29.116926 | 37.572883 | 57.827421 | 123.499729 | 214.366145 | 228.872448 | 200.152180 | 122.744816 | 111.298124 | 69.213964 | 25.435496 | 10.932831 | 4.465893 | 2.837777 | 2.516446 | 2.185881 | 1.834672 | 2.073448 | 2.548026 | 3.312141 | 3.786415 | 4.225790 | 4.934452 | 6.600876 | 5.611991 | 3.020126 | 1.975836 | 1.347089 | 1.299190 | 1.227933 | 1.060439 | 0.993412 | 1.402888 | 2.168332 | 2.678327 | 2.772108 | 2.687540 | 2.764185 | 3.098324 | 3.980379 | 4.048358 | 4.412383 | 3.703230 | 4.807662 | 5.236758 | 6.288661 | 5.707816 | 7.064939 | 7.644030 | 8.991088 | 9.290990 | 12.094445 | 13.973360 | 25.880869 | 38.822075 | 67.923472 | 98.111786 | 101.219002 | 117.578799 | 90.562978 | 50.075983 | 33.677670 | 31.738877 | 31.906686 | 38.032693 | 42.664305 | 9.282431e+01 | 1.619287e+02 | 1.491399e+02 | 143.247100 | 1.349909e+02 | 128.450935 | 139.030667 | 80.533321 | 44.800374 | 29.196406 | 20.425164 | 15.519919 | 10.526954 | 7.407679 | 8.533719 | 10.899318 | 12.514576 | 14.082050 | 14.784284 | 11.420785 | 7.162504 | 4.038308 | 3.122979 | 2.768847 | 1.966781 | 1.268572 | 0.900237 | 1.005081 | 1.443408 | 1.720275 | 2.107045 | 2.097919 | 2.436886 | 3.426815 | 2.894065 | 2.180874 | 1.702334 | 1.265851 | 1.284244 | 1.240451 | 0.806935 | 0.636668 | 0.558095 | 0.538137 | 0.535651 | 0.605857 | 0.745885 | 1.219417 | 2.097211 | 2.993428 | 3.412953 | 3.492380 | 2.843887 | 2.705166 | 2.206327 | 1.956753 | 1.574030 | 1.278387 | 1.150277 | 0.977584 | 0.748482 | 0.620985 | 0.549783 | 0.561116 | 0.534768 | 0.556907 | 0.640527 | 0.712262 | 0.815124 | 0.679163 | 0.527353 | 0.434994 | 0.357774 | 0.305520 | 0.295452 | 0.307185 | 0.400642 | 0.675105 | 1.244979 | 2.096086 | 2.424408 | 2.317642 | 1.974791 | 1.470529 | 1.056182 | 0.804402 | 0.618146 |
| all electrodes | 0.212021 | 0.225246 | 0.256504 | 0.258027 | 0.241590 | 0.223940 | 0.228106 | 0.216346 | 0.213376 | 0.225828 | 0.272025 | 0.257253 | 0.221582 | 0.231375 | 0.224603 | 0.218578 | 0.212341 | 0.213018 | 0.219648 | 0.216572 | 0.216473 | 0.230135 | 0.217650 | 0.234551 | 0.253902 | 0.282638 | 0.358234 | 0.973160 | 1.069569 | 0.869922 | 0.767201 | 0.690622 | 0.439686 | 0.258839 | 0.211239 | 0.231518 | 0.273182 | 0.320038 | 0.351912 | 0.482065 | 0.872322 | 1.110393 | 1.484751 | 1.061606 | 1.142631 | 0.714284 | 0.386373 | 0.272788 | 0.271440 | 0.300814 | 0.342599 | 0.437548 | 0.644460 | 1.160304 | 1.856058 | 0.927037 | 0.559890 | 0.540246 | 0.440660 | 0.474961 | 0.416335 | 0.245726 | 0.214094 | 0.480324 | 4.556030 | 105.345058 | 2416.153264 | 11330.730879 | 16914.834585 | 1.448884e+04 | 8.978950e+03 | 5.261208e+03 | 1844.180336 | 514.324115 | 99.303653 | 14.559082 | 2.908815 | 0.671738 | 0.281031 | 0.210824 | 0.231084 | 0.286250 | 0.333476 | 0.411935 | 0.390672 | 0.348470 | 0.286625 | 0.247265 | 0.220063 | 0.210813 | 0.233284 | 0.337949 | 0.539725 | 0.993773 | 2.148877 | 6.003892 | 10.520274 | 9.463378 | 4.751361 | 2.575079 | 1.121501 | 0.551273 | 0.382099 | 0.371225 | 0.454360 | 0.677898 | 1.377425 | 3.385822 | 7.968513 | 24.891445 | 65.472811 | 75.105491 | 48.116751 | 37.032230 | 43.885614 | 75.281994 | 101.732894 | 68.188403 | 43.186750 | 39.926603 | 21.354966 | 11.483203 | 5.819659 | 4.833777 | 3.666605 | 3.597770 | 3.623922 | 4.578937 | 6.325874 | 8.422902 | 8.133752 | 11.404939 | 10.007977 | 10.679975 | 9.876811 | 11.641873 | 10.803418 | 9.056873 | 9.465683 | 10.657500 | 9.197533 | 6.526382 | 6.228243 | 6.037503 | 5.563165 | 5.058907 | 4.666670 | 5.041534 | 5.888344 | 6.390735 | 8.897376 | 9.213318 | 9.133759 | 9.333592 | 8.406971 | 9.145608 | 12.388077 | 14.740788 | 11.452461 | 7.738674 | 7.490247 | 9.957587 | 14.645436 | 13.416092 | 11.153619 | 18.224051 | 55.696835 | 108.678450 | 102.582037 | 69.001046 | 67.521057 | 178.833376 | 261.027946 | 260.538149 | 239.426545 | 135.339524 | 122.346116 | 74.812738 | 23.235959 | 13.240260 | 11.945133 | 18.679353 | 48.663627 | 87.208146 | 146.271454 | 224.863653 | 230.842259 | 126.114880 | 45.076510 | 27.941999 | 27.973208 | 29.427698 | 33.196992 | 3.432008e+01 | 4.345144e+01 | 6.081942e+01 | 60.345823 | 5.000946e+01 | 46.525172 | 37.509501 | 28.847049 | 12.465068 | 7.024118 | 5.460190 | 3.817541 | 2.786862 | 2.566852 | 2.157866 | 3.538515 | 5.116376 | 7.446925 | 7.628356 | 6.558079 | 6.107521 | 4.359855 | 2.007536 | 1.328519 | 0.992575 | 0.908563 | 1.105870 | 0.869083 | 0.863681 | 1.067561 | 1.456213 | 1.504690 | 1.215837 | 0.769336 | 0.657580 | 0.639296 | 0.605157 | 0.575128 | 0.511291 | 0.639862 | 0.775313 | 0.853898 | 0.982428 | 0.930525 | 0.924755 | 1.370004 | 1.124662 | 0.949459 | 1.051647 | 1.151229 | 1.167529 | 1.383148 | 1.474888 | 1.847697 | 3.212884 | 4.083032 | 3.510917 | 3.524023 | 2.966855 | 1.994316 | 1.304120 | 0.833848 | 0.686207 | 0.590547 | 0.591307 | 0.675265 | 0.850361 | 1.188900 | 1.809685 | 1.617096 | 1.171505 | 0.935571 | 0.575134 | 0.428302 | 0.359241 | 0.285970 | 0.294698 | 0.349239 | 0.396963 | 0.548023 | 0.660113 | 0.813340 | 1.119313 | 1.340926 | 1.342962 | 1.489025 | 1.460982 |

Searchlight, spatiotemporal cluster permutation test

|  | start time | stop time | peak time | peak channel | cluster p | peak Cohen's d | direction |
| --- | --- | --- | --- | --- | --- | --- | --- |
| #1 | 110 | 190 | 150 | P8 | 0.0167 | 1.927037 | positive |
| #2 | 220 | 1195 | 800 | P8 | 0.0001 | 1.338393 | positive |

U) real to emoji faces - neutral vs sad

  
|  | time window | peak latency | cluster *p* | peak Cohen's *d* |  | | | |  |
| **all electrodes** | 130 - 190 ms | 155 ms | 0.0192 | 1.6776 |  | | | |  |
 355 - 455 ms | 425 ms | 0.0432 | 0.7233 |  | | | | 655 - 785 ms | 725 ms | 0.0143 | 1.018 |  | | | ||  |  | | | | | | | | |

Time-resolved classification, cluster permutation tests

|  |  | **left hemisphere** | | | | **right hemisphere** | | | |
|  | time window | peak latency | cluster *p* | peak Cohen's *d* | time window | peak latency | cluster *p* | peak Cohen's *d* |  |
| **anterior** |  | | | |  | | | |  |
| **central** |  | | | |  | | | |  |
| **posterior** | 130 - 185 ms | 155 ms | 0.0335 | 1.791 | 130 - 190 ms | 155 ms | 0.0378 | 1.5489 |  |
 710 - 820 ms | 735 ms | 0.0439 | 0.7453 |  | | | |

  

Time-resolved classification, Bayesian statistics

|  | -200 | -195 | -190 | -185 | -180 | -175 | -170 | -165 | -160 | -155 | -150 | -145 | -140 | -135 | -130 | -125 | -120 | -115 | -110 | -105 | -100 | -95 | -90 | -85 | -80 | -75 | -70 | -65 | -60 | -55 | -50 | -45 | -40 | -35 | -30 | -25 | -20 | -15 | -10 | -5 | 0 | 5 | 10 | 15 | 20 | 25 | 30 | 35 | 40 | 45 | 50 | 55 | 60 | 65 | 70 | 75 | 80 | 85 | 90 | 95 | 100 | 105 | 110 | 115 | 120 | 125 | 130 | 135 | 140 | 145 | 150 | 155 | 160 | 165 | 170 | 175 | 180 | 185 | 190 | 195 | 200 | 205 | 210 | 215 | 220 | 225 | 230 | 235 | 240 | 245 | 250 | 255 | 260 | 265 | 270 | 275 | 280 | 285 | 290 | 295 | 300 | 305 | 310 | 315 | 320 | 325 | 330 | 335 | 340 | 345 | 350 | 355 | 360 | 365 | 370 | 375 | 380 | 385 | 390 | 395 | 400 | 405 | 410 | 415 | 420 | 425 | 430 | 435 | 440 | 445 | 450 | 455 | 460 | 465 | 470 | 475 | 480 | 485 | 490 | 495 | 500 | 505 | 510 | 515 | 520 | 525 | 530 | 535 | 540 | 545 | 550 | 555 | 560 | 565 | 570 | 575 | 580 | 585 | 590 | 595 | 600 | 605 | 610 | 615 | 620 | 625 | 630 | 635 | 640 | 645 | 650 | 655 | 660 | 665 | 670 | 675 | 680 | 685 | 690 | 695 | 700 | 705 | 710 | 715 | 720 | 725 | 730 | 735 | 740 | 745 | 750 | 755 | 760 | 765 | 770 | 775 | 780 | 785 | 790 | 795 | 800 | 805 | 810 | 815 | 820 | 825 | 830 | 835 | 840 | 845 | 850 | 855 | 860 | 865 | 870 | 875 | 880 | 885 | 890 | 895 | 900 | 905 | 910 | 915 | 920 | 925 | 930 | 935 | 940 | 945 | 950 | 955 | 960 | 965 | 970 | 975 | 980 | 985 | 990 | 995 | 1000 | 1005 | 1010 | 1015 | 1020 | 1025 | 1030 | 1035 | 1040 | 1045 | 1050 | 1055 | 1060 | 1065 | 1070 | 1075 | 1080 | 1085 | 1090 | 1095 | 1100 | 1105 | 1110 | 1115 | 1120 | 1125 | 1130 | 1135 | 1140 | 1145 | 1150 | 1155 | 1160 | 1165 | 1170 | 1175 | 1180 | 1185 | 1190 | 1195 |
| --- | --- | --- | --- | --- | --- | --- | --- | --- | --- | --- | --- | --- | --- | --- | --- | --- | --- | --- | --- | --- | --- | --- | --- | --- | --- | --- | --- | --- | --- | --- | --- | --- | --- | --- | --- | --- | --- | --- | --- | --- | --- | --- | --- | --- | --- | --- | --- | --- | --- | --- | --- | --- | --- | --- | --- | --- | --- | --- | --- | --- | --- | --- | --- | --- | --- | --- | --- | --- | --- | --- | --- | --- | --- | --- | --- | --- | --- | --- | --- | --- | --- | --- | --- | --- | --- | --- | --- | --- | --- | --- | --- | --- | --- | --- | --- | --- | --- | --- | --- | --- | --- | --- | --- | --- | --- | --- | --- | --- | --- | --- | --- | --- | --- | --- | --- | --- | --- | --- | --- | --- | --- | --- | --- | --- | --- | --- | --- | --- | --- | --- | --- | --- | --- | --- | --- | --- | --- | --- | --- | --- | --- | --- | --- | --- | --- | --- | --- | --- | --- | --- | --- | --- | --- | --- | --- | --- | --- | --- | --- | --- | --- | --- | --- | --- | --- | --- | --- | --- | --- | --- | --- | --- | --- | --- | --- | --- | --- | --- | --- | --- | --- | --- | --- | --- | --- | --- | --- | --- | --- | --- | --- | --- | --- | --- | --- | --- | --- | --- | --- | --- | --- | --- | --- | --- | --- | --- | --- | --- | --- | --- | --- | --- | --- | --- | --- | --- | --- | --- | --- | --- | --- | --- | --- | --- | --- | --- | --- | --- | --- | --- | --- | --- | --- | --- | --- | --- | --- | --- | --- | --- | --- | --- | --- | --- | --- | --- | --- | --- | --- | --- | --- | --- | --- | --- | --- | --- | --- | --- | --- | --- | --- | --- | --- | --- | --- | --- | --- | --- | --- | --- | --- | --- | --- | --- | --- | --- | --- | --- | --- | --- |
| left anterior | 0.223816 | 0.248502 | 0.236115 | 0.244948 | 0.235450 | 0.328445 | 0.519482 | 0.403469 | 0.255830 | 0.234240 | 0.212578 | 0.216779 | 0.212414 | 0.246442 | 0.247766 | 0.231928 | 0.251274 | 0.228789 | 0.309762 | 0.429735 | 0.374982 | 0.242098 | 0.211170 | 0.276139 | 0.312598 | 0.541932 | 0.790443 | 0.606158 | 0.494285 | 0.531409 | 0.456986 | 0.409302 | 0.428117 | 0.524303 | 0.807682 | 0.731934 | 0.316743 | 0.309001 | 0.852268 | 0.851215 | 0.452635 | 0.293650 | 0.234015 | 0.330092 | 0.332234 | 0.226030 | 0.217183 | 0.212791 | 0.213466 | 0.212090 | 0.231121 | 0.251811 | 0.244002 | 0.248777 | 0.238865 | 0.218732 | 0.211013 | 0.211492 | 0.214564 | 0.213149 | 0.211076 | 0.210905 | 0.210820 | 0.210986 | 0.273899 | 0.823912 | 3.255443 | 6.901479 | 11.596567 | 24.652189 | 4.002731e+01 | 1.376583e+01 | 4.540611e+00 | 1.698295e+00 | 1.536741e+00 | 1.283956 | 1.053248 | 0.769275 | 0.747109 | 0.756267 | 0.789570 | 0.670251 | 0.472134 | 0.335311 | 0.241712 | 0.218002 | 0.210867 | 0.228143 | 0.278082 | 0.269224 | 0.327950 | 0.421025 | 0.635907 | 0.987660 | 0.821047 | 0.868533 | 1.047486 | 1.374914 | 0.878458 | 0.538174 | 0.326709 | 0.249128 | 0.214556 | 0.212607 | 0.248438 | 0.262751 | 0.258629 | 0.241919 | 0.235592 | 0.225245 | 0.220229 | 0.227847 | 0.253271 | 0.299137 | 0.341907 | 0.381301 | 0.420568 | 0.632765 | 0.846045 | 0.804065 | 0.718917 | 0.778187 | 0.677458 | 0.811959 | 0.612857 | 0.392704 | 0.345561 | 0.335339 | 0.294355 | 0.290326 | 0.285595 | 0.292681 | 0.335064 | 0.311632 | 0.267100 | 0.240160 | 0.221232 | 0.225208 | 0.231907 | 0.231608 | 0.223937 | 0.223961 | 0.214093 | 0.211903 | 0.213380 | 0.220966 | 0.243142 | 0.245595 | 0.239578 | 0.219127 | 0.213380 | 0.211086 | 0.211492 | 0.211247 | 0.216534 | 0.219100 | 0.226283 | 0.239822 | 0.227364 | 0.228094 | 0.252942 | 0.242060 | 0.230912 | 0.216891 | 0.213330 | 0.212578 | 0.210812 | 0.214799 | 0.221996 | 0.229078 | 0.220844 | 0.235097 | 0.221754 | 0.211533 | 0.218941 | 0.245180 | 0.254366 | 0.250080 | 0.247164 | 0.239505 | 0.236590 | 0.250596 | 0.260603 | 0.289514 | 0.303965 | 0.334866 | 0.368979 | 0.361900 | 0.351256 | 0.326581 | 0.351578 | 0.398483 | 0.443238 | 0.426359 | 0.458641 | 0.485268 | 0.531661 | 0.448055 | 0.363663 | 0.325234 | 0.304756 | 0.262623 | 0.232219 | 0.221334 | 0.222704 | 0.238588 | 0.254534 | 0.273696 | 0.282313 | 0.349286 | 0.462641 | 0.482518 | 0.421286 | 0.382971 | 0.335018 | 0.325219 | 0.258745 | 0.217958 | 0.213799 | 0.214314 | 0.210920 | 0.210813 | 0.211386 | 0.210812 | 0.214996 | 0.224484 | 0.232264 | 0.246919 | 0.267644 | 0.324008 | 0.389429 | 0.380889 | 0.287953 | 0.241822 | 0.240766 | 0.239104 | 0.241507 | 0.243410 | 0.242858 | 0.245997 | 0.252058 | 0.251617 | 0.251690 | 0.241084 | 0.221039 | 0.218027 | 0.222394 | 0.224939 | 0.228940 | 0.244940 | 0.263106 | 0.309160 | 0.379353 | 0.484040 | 0.623777 | 0.578309 | 0.476247 | 0.457913 | 0.566744 | 0.560467 | 0.452260 | 0.361148 | 0.320985 | 0.317782 | 0.307405 | 0.266455 | 0.230602 | 0.227264 | 0.240145 | 0.256560 | 0.274654 | 0.285441 | 0.310204 | 0.329143 | 0.313270 | 0.302845 | 0.313936 | 0.313948 | 0.290235 | 0.250851 |
| right anterior | 0.211753 | 0.214275 | 0.216804 | 0.219902 | 0.240711 | 0.251632 | 0.219467 | 0.223396 | 0.219836 | 0.236107 | 0.244482 | 0.237551 | 0.260098 | 0.332126 | 0.280884 | 0.230955 | 0.210829 | 0.225583 | 0.236466 | 0.226037 | 0.214323 | 0.211301 | 0.282817 | 0.284264 | 0.256890 | 0.234451 | 0.211226 | 0.239286 | 0.363178 | 1.032756 | 0.728442 | 0.558889 | 0.566779 | 0.748112 | 0.681655 | 0.404512 | 0.254918 | 0.227714 | 0.210930 | 0.221039 | 0.231667 | 0.225207 | 0.234346 | 0.212318 | 0.225723 | 0.273288 | 0.367647 | 0.345996 | 0.303778 | 0.318429 | 0.268161 | 0.212095 | 0.214786 | 0.229549 | 0.216329 | 0.211185 | 0.223433 | 0.230591 | 0.268416 | 0.278505 | 0.251426 | 0.226457 | 0.212430 | 0.210806 | 0.221093 | 0.263685 | 0.452228 | 1.055046 | 2.892521 | 6.381438 | 5.435607e+00 | 3.903822e+00 | 2.453818e+00 | 1.553317e+00 | 1.501537e+00 | 0.869818 | 0.534433 | 0.532858 | 0.448253 | 0.345405 | 0.264579 | 0.217385 | 0.213063 | 0.214939 | 0.217195 | 0.228128 | 0.225393 | 0.218658 | 0.222062 | 0.217068 | 0.214486 | 0.211476 | 0.234897 | 0.251440 | 0.295136 | 0.341040 | 0.331981 | 0.345188 | 0.314690 | 0.264841 | 0.236216 | 0.216723 | 0.210806 | 0.214130 | 0.227585 | 0.274948 | 0.314910 | 0.339021 | 0.416627 | 0.524233 | 0.602877 | 0.544976 | 0.400170 | 0.436665 | 0.464472 | 0.518727 | 0.550560 | 0.561277 | 0.702731 | 1.208730 | 1.255341 | 1.500000 | 1.248746 | 0.992668 | 0.914094 | 0.570115 | 0.354761 | 0.288630 | 0.247412 | 0.239243 | 0.231368 | 0.230383 | 0.244087 | 0.274219 | 0.250821 | 0.223334 | 0.211465 | 0.218354 | 0.250440 | 0.314848 | 0.371354 | 0.310955 | 0.298710 | 0.284123 | 0.318938 | 0.355735 | 0.330176 | 0.345842 | 0.320944 | 0.268301 | 0.241359 | 0.213052 | 0.223401 | 0.267979 | 0.273031 | 0.233794 | 0.213321 | 0.210806 | 0.228348 | 0.293369 | 0.303733 | 0.299890 | 0.300213 | 0.296094 | 0.309393 | 0.313172 | 0.302565 | 0.356251 | 0.383826 | 0.324699 | 0.318233 | 0.332160 | 0.360638 | 0.401810 | 0.425933 | 0.259450 | 0.219563 | 0.210896 | 0.232023 | 0.301051 | 0.447352 | 0.534120 | 0.494285 | 0.444066 | 0.533419 | 0.426804 | 0.341823 | 0.265698 | 0.254018 | 0.220650 | 0.211036 | 0.228816 | 0.244650 | 0.240898 | 0.227204 | 0.215161 | 0.211868 | 0.211809 | 0.210817 | 0.211963 | 0.214740 | 0.218667 | 0.220531 | 0.242797 | 0.287601 | 0.319810 | 0.336999 | 0.315823 | 0.298491 | 0.286531 | 0.287653 | 0.335778 | 0.300562 | 0.288710 | 0.313169 | 0.339261 | 0.269208 | 0.225700 | 0.211509 | 0.212965 | 0.221420 | 0.224073 | 0.213942 | 0.243649 | 0.310405 | 0.339317 | 0.328209 | 0.332378 | 0.343969 | 0.359068 | 0.302303 | 0.255615 | 0.249082 | 0.243690 | 0.230216 | 0.221227 | 0.217348 | 0.210838 | 0.211617 | 0.211997 | 0.211655 | 0.220439 | 0.223013 | 0.215832 | 0.212395 | 0.223989 | 0.235831 | 0.237297 | 0.268422 | 0.253849 | 0.235571 | 0.218183 | 0.212413 | 0.211172 | 0.211938 | 0.221793 | 0.221771 | 0.218132 | 0.211803 | 0.210849 | 0.210907 | 0.212809 | 0.210960 | 0.213288 | 0.225471 | 0.237446 | 0.241377 | 0.253749 | 0.245485 | 0.214822 | 0.211280 | 0.215564 | 0.222990 | 0.221496 | 0.211594 | 0.212032 | 0.224551 | 0.237913 | 0.260014 | 0.363114 |
| left central | 0.226249 | 0.220299 | 0.232680 | 0.239143 | 0.267732 | 0.241852 | 0.234140 | 0.264686 | 0.350935 | 0.264752 | 0.227493 | 0.210978 | 0.216398 | 0.261319 | 0.254117 | 0.295590 | 0.389826 | 0.426408 | 0.408486 | 0.386844 | 0.270084 | 0.215514 | 0.243624 | 0.235509 | 0.212019 | 0.231136 | 0.245677 | 0.234270 | 0.300730 | 0.473314 | 0.946988 | 1.547017 | 1.509932 | 2.180171 | 3.611522 | 2.947486 | 0.330533 | 0.212481 | 0.245610 | 0.308682 | 0.293166 | 0.294413 | 0.295822 | 0.218015 | 0.213348 | 0.227098 | 0.230905 | 0.233794 | 0.262126 | 0.333456 | 0.785784 | 2.111428 | 2.298978 | 1.587533 | 1.531102 | 1.082576 | 0.491566 | 0.271486 | 0.214450 | 0.211399 | 0.214191 | 0.240220 | 0.243844 | 0.262607 | 0.234356 | 0.211741 | 0.243335 | 0.516500 | 2.296212 | 18.586933 | 1.079904e+02 | 7.424751e+02 | 1.158433e+03 | 5.073409e+02 | 4.338685e+01 | 6.061544 | 1.490745 | 0.730549 | 0.476133 | 0.429366 | 0.390034 | 0.378690 | 0.403429 | 0.582246 | 0.638304 | 0.291038 | 0.211873 | 0.274958 | 0.366376 | 0.413935 | 0.787718 | 1.208391 | 1.340219 | 1.033416 | 0.536726 | 0.306610 | 0.279807 | 0.237422 | 0.210886 | 0.232045 | 0.246292 | 0.259535 | 0.243711 | 0.237615 | 0.232295 | 0.211976 | 0.218805 | 0.219625 | 0.219365 | 0.215282 | 0.214254 | 0.211191 | 0.220026 | 0.229575 | 0.289524 | 0.352603 | 0.355473 | 0.331782 | 0.330299 | 0.310118 | 0.410447 | 0.368244 | 0.300199 | 0.276965 | 0.322000 | 0.393417 | 0.400403 | 0.311416 | 0.239962 | 0.227452 | 0.212832 | 0.222064 | 0.276148 | 0.341938 | 0.356949 | 0.283215 | 0.238183 | 0.211566 | 0.215922 | 0.251696 | 0.280244 | 0.262464 | 0.276602 | 0.244316 | 0.225515 | 0.214775 | 0.211625 | 0.218265 | 0.224641 | 0.228636 | 0.232403 | 0.251836 | 0.257036 | 0.248833 | 0.244732 | 0.232821 | 0.221844 | 0.212748 | 0.210942 | 0.223234 | 0.263074 | 0.250418 | 0.229234 | 0.216713 | 0.211304 | 0.210929 | 0.213033 | 0.229861 | 0.233649 | 0.247947 | 0.267339 | 0.287269 | 0.372417 | 0.461837 | 0.521570 | 0.490478 | 0.411198 | 0.402084 | 0.364963 | 0.267064 | 0.221309 | 0.211733 | 0.212468 | 0.213650 | 0.212696 | 0.211934 | 0.214128 | 0.223106 | 0.223687 | 0.214722 | 0.215543 | 0.219936 | 0.222493 | 0.221528 | 0.217281 | 0.242530 | 0.318168 | 0.350766 | 0.371307 | 0.382273 | 0.383843 | 0.512825 | 0.393526 | 0.300776 | 0.267999 | 0.236599 | 0.248788 | 0.295929 | 0.266387 | 0.251266 | 0.223561 | 0.212088 | 0.212282 | 0.210946 | 0.214622 | 0.221397 | 0.245975 | 0.265048 | 0.273599 | 0.281823 | 0.294921 | 0.255693 | 0.241171 | 0.228479 | 0.228601 | 0.229275 | 0.220170 | 0.214466 | 0.212049 | 0.211199 | 0.211776 | 0.214064 | 0.215315 | 0.217639 | 0.212923 | 0.213177 | 0.217837 | 0.214613 | 0.213791 | 0.211199 | 0.210842 | 0.210843 | 0.210884 | 0.221428 | 0.226489 | 0.229768 | 0.225055 | 0.225472 | 0.223564 | 0.227945 | 0.219613 | 0.222957 | 0.231598 | 0.224308 | 0.212271 | 0.213092 | 0.215517 | 0.213400 | 0.212106 | 0.216387 | 0.220678 | 0.219008 | 0.212799 | 0.212885 | 0.214635 | 0.226184 | 0.229287 | 0.216379 | 0.211131 | 0.212011 | 0.211798 | 0.214791 | 0.211161 | 0.214018 | 0.213606 | 0.212387 | 0.211972 | 0.212329 | 0.217467 | 0.215781 |
| right central | 0.676151 | 0.978796 | 0.953855 | 0.511898 | 0.253267 | 0.212496 | 0.321597 | 0.592206 | 0.836108 | 1.323937 | 0.708788 | 0.494152 | 0.344859 | 0.245781 | 0.214139 | 0.266604 | 0.299028 | 0.326223 | 0.268161 | 0.219526 | 0.210942 | 0.210975 | 0.270400 | 0.372994 | 0.570393 | 0.602562 | 0.295144 | 0.235172 | 0.229586 | 0.211394 | 0.252197 | 0.389528 | 0.469732 | 0.276075 | 0.218683 | 0.217358 | 0.217514 | 0.210819 | 0.213493 | 0.213235 | 0.211605 | 0.216104 | 0.225173 | 0.226623 | 0.215713 | 0.215302 | 0.217499 | 0.218420 | 0.219303 | 0.231883 | 0.268901 | 0.332524 | 0.425969 | 0.580517 | 0.802797 | 1.049234 | 0.828064 | 0.625974 | 0.602925 | 0.451360 | 0.525836 | 0.579093 | 0.470860 | 0.314232 | 0.214351 | 0.250602 | 0.685941 | 6.072086 | 31.880376 | 96.991801 | 1.054773e+02 | 1.166150e+02 | 1.580005e+02 | 1.872133e+02 | 1.340732e+02 | 49.878905 | 20.660141 | 4.395357 | 1.012470 | 0.466050 | 0.252739 | 0.210908 | 0.241235 | 0.337594 | 0.318588 | 0.288974 | 0.267146 | 0.307454 | 0.300307 | 0.264602 | 0.232503 | 0.215991 | 0.229315 | 0.251879 | 0.247288 | 0.284675 | 0.326630 | 0.369382 | 0.411350 | 0.385943 | 0.311333 | 0.266030 | 0.236196 | 0.214948 | 0.222896 | 0.286129 | 0.391330 | 0.428344 | 0.405777 | 0.409160 | 0.546010 | 0.645836 | 0.675996 | 0.564238 | 0.561364 | 0.503045 | 0.432947 | 0.325986 | 0.285510 | 0.306885 | 0.350126 | 0.365172 | 0.476714 | 0.841678 | 0.896367 | 0.849890 | 0.557277 | 0.375814 | 0.360123 | 0.357812 | 0.290548 | 0.267769 | 0.230902 | 0.220500 | 0.221705 | 0.214104 | 0.214003 | 0.218093 | 0.224330 | 0.239128 | 0.270429 | 0.278236 | 0.303294 | 0.305337 | 0.315386 | 0.324748 | 0.275130 | 0.234906 | 0.225978 | 0.216124 | 0.216593 | 0.219640 | 0.212225 | 0.212080 | 0.217683 | 0.218979 | 0.217264 | 0.214614 | 0.210974 | 0.214205 | 0.223262 | 0.229417 | 0.246376 | 0.278515 | 0.301371 | 0.337311 | 0.285581 | 0.253524 | 0.233487 | 0.214003 | 0.213241 | 0.263002 | 0.315938 | 0.308512 | 0.333953 | 0.392805 | 0.423131 | 0.405995 | 0.367585 | 0.319280 | 0.277042 | 0.263329 | 0.254454 | 0.236542 | 0.252344 | 0.250509 | 0.273091 | 0.264036 | 0.234571 | 0.216806 | 0.210968 | 0.222209 | 0.251302 | 0.317633 | 0.323146 | 0.355601 | 0.398924 | 0.448229 | 0.494265 | 0.492313 | 0.393004 | 0.435988 | 0.378004 | 0.334869 | 0.271649 | 0.231274 | 0.213867 | 0.210968 | 0.213829 | 0.214842 | 0.216416 | 0.219848 | 0.216025 | 0.211209 | 0.211859 | 0.221952 | 0.231913 | 0.254066 | 0.354039 | 0.368173 | 0.406961 | 0.536090 | 0.469243 | 0.436494 | 0.359965 | 0.298036 | 0.296189 | 0.274456 | 0.247267 | 0.217011 | 0.212161 | 0.218498 | 0.213711 | 0.211344 | 0.213249 | 0.235147 | 0.250088 | 0.316640 | 0.383727 | 0.371855 | 0.365075 | 0.278108 | 0.232843 | 0.212941 | 0.211834 | 0.210902 | 0.214250 | 0.220174 | 0.217089 | 0.211659 | 0.225515 | 0.261946 | 0.340198 | 0.405910 | 0.405054 | 0.316923 | 0.274164 | 0.244545 | 0.221931 | 0.215752 | 0.211587 | 0.218620 | 0.253610 | 0.260580 | 0.242137 | 0.265672 | 0.262095 | 0.270295 | 0.268891 | 0.267592 | 0.258341 | 0.257128 | 0.218549 | 0.217473 | 0.214941 | 0.213257 | 0.211674 | 0.210848 | 0.211146 | 0.216759 |
| left posterior | 0.334470 | 0.230282 | 0.210907 | 0.279356 | 0.467638 | 0.843497 | 1.225418 | 0.682368 | 0.357901 | 0.239133 | 0.213632 | 0.252595 | 0.308352 | 0.266109 | 0.221512 | 0.217388 | 0.216614 | 0.234379 | 0.291840 | 0.411636 | 0.690193 | 1.396110 | 1.398530 | 1.125244 | 0.663770 | 0.572682 | 0.227376 | 0.215887 | 0.436088 | 0.840780 | 0.922431 | 0.517374 | 0.662837 | 0.318539 | 0.232485 | 0.241983 | 0.274494 | 0.268198 | 0.237342 | 0.223993 | 0.215374 | 0.214971 | 0.216366 | 0.220513 | 0.218990 | 0.221381 | 0.277135 | 0.411921 | 0.537541 | 0.740144 | 0.813098 | 0.786707 | 1.016838 | 0.757070 | 0.684117 | 1.117104 | 0.631601 | 0.666553 | 0.858434 | 1.012340 | 1.316222 | 1.224907 | 1.229812 | 0.635643 | 0.226832 | 0.425814 | 4.067432 | 51.812885 | 385.854990 | 4853.563759 | 1.125526e+05 | 3.001659e+06 | 9.078780e+06 | 1.737571e+07 | 3.469978e+06 | 33256.343711 | 189.549635 | 5.678997 | 0.836300 | 0.303906 | 0.230397 | 0.213751 | 0.219894 | 0.234322 | 0.262103 | 0.356345 | 0.436649 | 0.454752 | 0.397162 | 0.288168 | 0.234816 | 0.212842 | 0.231361 | 0.231830 | 0.218954 | 0.210863 | 0.217012 | 0.217575 | 0.220108 | 0.237670 | 0.234237 | 0.221391 | 0.213017 | 0.214403 | 0.216846 | 0.213256 | 0.211153 | 0.216306 | 0.213252 | 0.211077 | 0.210925 | 0.215039 | 0.237122 | 0.292664 | 0.375890 | 0.375511 | 0.392894 | 0.372763 | 0.344654 | 0.302993 | 0.278049 | 0.266010 | 0.274615 | 0.340896 | 0.405955 | 0.423293 | 0.375362 | 0.381844 | 0.558443 | 0.781523 | 0.735620 | 0.725784 | 0.833268 | 1.044291 | 1.127791 | 0.769722 | 0.554421 | 0.530334 | 0.478550 | 0.485679 | 0.518151 | 0.722839 | 0.770333 | 0.814078 | 0.737215 | 0.790997 | 0.783005 | 0.662915 | 0.507422 | 0.584294 | 0.761702 | 0.923089 | 0.854932 | 0.690580 | 0.672032 | 0.677501 | 0.669580 | 0.474409 | 0.436976 | 0.466497 | 0.503758 | 0.607024 | 0.567143 | 0.510160 | 0.529720 | 0.510256 | 0.483835 | 0.509537 | 0.551885 | 0.800116 | 1.090042 | 1.521864 | 1.577569 | 2.044414 | 2.052445 | 1.740737 | 1.738115 | 1.441909 | 1.361504 | 1.239759 | 0.924440 | 1.275988 | 1.804552 | 2.918682 | 5.839469 | 6.733824 | 13.089668 | 33.313550 | 29.733896 | 25.940310 | 17.353728 | 12.527035 | 12.861138 | 9.828834 | 8.210920 | 7.423817 | 5.494590 | 3.249567 | 3.014018 | 3.106719 | 2.090761 | 1.590275 | 1.492889 | 1.601958 | 1.819620 | 1.254392 | 0.979868 | 1.165951 | 1.398350 | 1.377137 | 1.314370 | 1.407520 | 2.185584 | 2.575521 | 2.775403 | 2.645814 | 2.449376 | 2.381598 | 1.803553 | 1.182103 | 0.913019 | 0.676852 | 0.613995 | 0.676742 | 0.833333 | 1.017075 | 1.178444 | 1.735895 | 3.106076 | 3.236058 | 3.111538 | 2.923879 | 2.594729 | 1.812500 | 1.215383 | 0.787816 | 0.655333 | 0.615018 | 0.631015 | 0.841355 | 0.953144 | 1.090729 | 0.957924 | 1.001033 | 1.027104 | 1.021510 | 0.851614 | 1.044081 | 0.966616 | 0.749041 | 0.504174 | 0.302843 | 0.240408 | 0.213770 | 0.213325 | 0.226066 | 0.222840 | 0.213653 | 0.211757 | 0.224156 | 0.234711 | 0.238363 | 0.242212 | 0.234474 | 0.229687 | 0.235163 | 0.227366 | 0.229526 | 0.245937 | 0.280817 | 0.396891 | 0.506771 | 0.596809 | 0.686572 | 0.762771 | 0.700306 | 0.554176 | 0.454925 | 0.388704 | 0.327293 |
| right posterior | 0.297381 | 0.269716 | 0.278462 | 0.318034 | 0.359072 | 0.481870 | 0.377620 | 0.277020 | 0.252472 | 0.237085 | 0.225083 | 0.231414 | 0.213867 | 0.222680 | 0.272532 | 0.335447 | 0.528116 | 0.564047 | 0.598857 | 1.049992 | 1.639841 | 1.970639 | 2.820140 | 1.558528 | 0.808276 | 0.488971 | 0.331784 | 0.254153 | 0.245112 | 0.223813 | 0.212654 | 0.213045 | 0.211246 | 0.214319 | 0.210817 | 0.211009 | 0.212618 | 0.212361 | 0.217153 | 0.245489 | 0.292333 | 0.287963 | 0.281673 | 0.244024 | 0.289182 | 0.478455 | 0.597061 | 0.774180 | 0.718044 | 0.405217 | 0.325707 | 0.265142 | 0.218980 | 0.212425 | 0.210806 | 0.210889 | 0.211912 | 0.228931 | 0.400298 | 1.105872 | 2.517253 | 3.637047 | 2.663524 | 0.762709 | 0.250905 | 0.324222 | 3.410558 | 53.054585 | 1619.637601 | 13067.165206 | 6.133386e+04 | 2.590651e+05 | 5.090989e+05 | 3.321407e+05 | 3.689122e+05 | 225319.005919 | 11491.485042 | 230.044646 | 4.245156 | 0.620998 | 0.291628 | 0.232349 | 0.219479 | 0.225229 | 0.233867 | 0.248743 | 0.265971 | 0.275228 | 0.269927 | 0.298552 | 0.269156 | 0.242428 | 0.235889 | 0.237295 | 0.242987 | 0.242157 | 0.226438 | 0.227096 | 0.234574 | 0.292342 | 0.314049 | 0.296614 | 0.292835 | 0.308611 | 0.319750 | 0.346451 | 0.284776 | 0.256535 | 0.241966 | 0.253120 | 0.270194 | 0.324062 | 0.330877 | 0.397663 | 0.593000 | 0.834532 | 0.912352 | 1.080604 | 1.357838 | 1.620609 | 1.598541 | 1.138504 | 0.848016 | 0.803435 | 0.564512 | 0.347463 | 0.273701 | 0.243359 | 0.254102 | 0.301485 | 0.351492 | 0.416113 | 0.456526 | 0.516191 | 0.559633 | 0.540904 | 0.473215 | 0.440075 | 0.366206 | 0.367115 | 0.377696 | 0.355386 | 0.364844 | 0.389984 | 0.404359 | 0.514300 | 0.582883 | 0.546165 | 0.660567 | 0.675178 | 0.733473 | 0.875394 | 0.898956 | 0.764733 | 0.810375 | 0.699324 | 0.663506 | 0.573784 | 0.463573 | 0.390244 | 0.356601 | 0.333485 | 0.326897 | 0.303185 | 0.296498 | 0.279092 | 0.275651 | 0.278861 | 0.268994 | 0.267683 | 0.293046 | 0.331042 | 0.420151 | 0.527571 | 0.651435 | 0.768220 | 0.968732 | 1.143095 | 1.188873 | 0.967445 | 0.911466 | 1.036502 | 1.441629 | 1.999616 | 2.962476 | 5.939266 | 14.988167 | 29.645250 | 44.943791 | 38.741998 | 24.266523 | 15.429893 | 8.841834 | 4.501070 | 2.716341 | 1.731555 | 1.541093 | 1.427129 | 1.168747 | 1.016196 | 1.206997 | 1.222463 | 1.582765 | 1.702516 | 2.259893 | 3.656974 | 4.374967 | 4.395023 | 4.452146 | 3.727329 | 4.586675 | 5.050357 | 4.907237 | 4.830975 | 4.120137 | 3.276866 | 2.627511 | 1.788381 | 1.210459 | 0.774887 | 0.621616 | 0.613759 | 0.646128 | 0.681029 | 0.712579 | 0.769491 | 0.910758 | 0.884424 | 0.738149 | 0.676747 | 0.536404 | 0.423106 | 0.353546 | 0.336758 | 0.320371 | 0.309301 | 0.283107 | 0.295621 | 0.321530 | 0.372712 | 0.402667 | 0.464087 | 0.550509 | 0.725931 | 0.677379 | 0.574239 | 0.494390 | 0.449841 | 0.394989 | 0.322588 | 0.247971 | 0.225523 | 0.222307 | 0.220172 | 0.213002 | 0.211639 | 0.226796 | 0.300575 | 0.469137 | 0.643607 | 0.843527 | 1.006285 | 1.129101 | 0.919767 | 0.722742 | 0.426665 | 0.323498 | 0.282387 | 0.289969 | 0.346516 | 0.438397 | 0.507656 | 0.934812 | 1.355144 | 1.332152 | 1.041733 | 0.617530 | 0.460187 | 0.385486 | 0.289798 |
| all electrodes | 0.210826 | 0.211450 | 0.211038 | 0.215535 | 0.211011 | 0.210955 | 0.211621 | 0.215793 | 0.231626 | 0.347907 | 0.354897 | 0.233180 | 0.226781 | 0.231981 | 0.215122 | 0.211602 | 0.262085 | 0.245992 | 0.227327 | 0.214843 | 0.211290 | 0.264366 | 0.436117 | 0.793259 | 0.761718 | 0.676826 | 0.426903 | 0.275845 | 0.210875 | 0.260717 | 0.294044 | 0.289426 | 0.332774 | 0.257858 | 0.215137 | 0.210891 | 0.227141 | 0.214885 | 0.210871 | 0.215243 | 0.210875 | 0.210826 | 0.211413 | 0.281126 | 0.445005 | 0.652740 | 1.239728 | 0.579846 | 0.422356 | 0.276523 | 0.210913 | 0.283021 | 0.521080 | 0.735984 | 0.393349 | 0.253029 | 0.213653 | 0.227988 | 0.242916 | 0.231314 | 0.216879 | 0.210886 | 0.212230 | 0.210846 | 0.274220 | 1.206651 | 39.613121 | 1476.607252 | 44967.431587 | 740486.311529 | 1.828402e+06 | 9.726659e+05 | 7.360551e+05 | 6.748848e+05 | 1.513096e+05 | 18831.337225 | 643.187725 | 22.241585 | 3.356952 | 1.047022 | 0.456185 | 0.288444 | 0.288444 | 0.382144 | 0.440248 | 0.505633 | 0.385842 | 0.372613 | 0.418775 | 0.348784 | 0.255825 | 0.221005 | 0.211605 | 0.211667 | 0.210857 | 0.211619 | 0.213500 | 0.214297 | 0.217337 | 0.259374 | 0.316875 | 0.347163 | 0.384684 | 0.416042 | 0.484362 | 0.459392 | 0.575327 | 0.642739 | 0.647870 | 0.690092 | 0.990608 | 1.728205 | 3.593973 | 4.823239 | 6.161032 | 6.309931 | 8.579664 | 7.083395 | 3.139628 | 1.922962 | 1.455988 | 1.645528 | 3.006572 | 5.804894 | 11.364968 | 26.201076 | 19.663749 | 15.132186 | 8.742450 | 5.957764 | 3.328714 | 1.965569 | 1.268983 | 1.138143 | 1.344078 | 1.896890 | 2.183451 | 2.352741 | 3.731559 | 4.105916 | 5.379794 | 3.546980 | 1.948272 | 1.142739 | 1.052509 | 0.853770 | 0.805230 | 0.708509 | 0.649197 | 0.793176 | 1.286576 | 1.338659 | 0.992402 | 0.880797 | 0.758727 | 0.887487 | 1.082748 | 1.056909 | 0.787812 | 0.756909 | 0.700029 | 0.607270 | 0.460728 | 0.386079 | 0.372095 | 0.427718 | 0.463921 | 0.436377 | 0.502928 | 0.617895 | 0.952540 | 1.778053 | 3.254802 | 4.257167 | 7.037043 | 9.339927 | 9.311093 | 7.645712 | 3.565066 | 4.226418 | 8.048696 | 28.036902 | 86.190811 | 188.885900 | 406.134781 | 728.194313 | 646.117593 | 393.772840 | 209.616851 | 110.279669 | 41.455308 | 17.103650 | 9.378170 | 6.227124 | 3.461296 | 2.471654 | 1.805073 | 1.472921 | 1.173258 | 1.072669 | 0.922124 | 0.892469 | 0.939929 | 1.034625 | 1.251420 | 1.434744 | 1.378417 | 1.706603 | 1.873930 | 1.803020 | 2.006737 | 2.637572 | 2.791964 | 2.714519 | 1.434016 | 0.800723 | 0.545071 | 0.462044 | 0.401191 | 0.365937 | 0.293179 | 0.293045 | 0.317070 | 0.347354 | 0.370001 | 0.374844 | 0.384812 | 0.448682 | 0.409937 | 0.385842 | 0.339817 | 0.322009 | 0.291483 | 0.285504 | 0.257451 | 0.255567 | 0.245101 | 0.239574 | 0.229730 | 0.223484 | 0.219107 | 0.225013 | 0.241680 | 0.296525 | 0.352849 | 0.400907 | 0.519497 | 0.636292 | 0.663757 | 0.486234 | 0.300068 | 0.238746 | 0.220424 | 0.210845 | 0.213266 | 0.216097 | 0.212469 | 0.215691 | 0.260688 | 0.305363 | 0.355096 | 0.347280 | 0.379294 | 0.366733 | 0.283665 | 0.228064 | 0.214112 | 0.212480 | 0.212717 | 0.212144 | 0.220665 | 0.269770 | 0.343603 | 0.362067 | 0.355992 | 0.287470 | 0.238998 | 0.225488 | 0.212898 | 0.211419 |

Searchlight, spatiotemporal cluster permutation test

|  | start time | stop time | peak time | peak channel | cluster p | peak Cohen's d | direction |
| --- | --- | --- | --- | --- | --- | --- | --- |
| #1 | 105 | 235 | 155 | PO8 | 0.0174 | 1.991536 | positive |
| #2 | 340 | 1055 | 880 | PO4 | 0.0064 | 0.843058 | positive |

V) emoji to real faces - emotion

  
|  | time window | peak latency | cluster *p* | peak Cohen's *d* |  | | | |  |
| **all electrodes** | 115 - 200 ms | 155 ms | 0.0162 | 1.5551 |  | | | |  |
 365 - 525 ms | 420 ms | 0.0068 | 0.9585 |  | | | ||  |  | | | | | | | | |

Time-resolved classification, cluster permutation tests

|  | **left hemisphere** | | | | **right hemisphere** | | | |  |
|  | time window | peak latency | cluster *p* | peak Cohen's *d* | time window | peak latency | cluster *p* | peak Cohen's *d* |  |
| **anterior** |  | | | |  | | | |  |
| **central** |  | | | | 130 - 185 ms | 150 ms | 0.0443 | 1.1636 |  |
  | | | | 350 - 500 ms | 425 ms | 0.0154 | 0.7565 ||  | **posterior** | 120 - 180 ms | 150 ms | 0.0279 | 1.3565 | 120 - 190 ms | 150 ms | 0.0149 | 2.0479 |
 725 - 880 ms | 805 ms | 0.0122 | 0.6582 |  | | | | 935 - 1060 ms | 980 ms | 0.0155 | 0.9365 |  | | | |

  

Time-resolved classification, Bayesian statistics

|  | -200 | -195 | -190 | -185 | -180 | -175 | -170 | -165 | -160 | -155 | -150 | -145 | -140 | -135 | -130 | -125 | -120 | -115 | -110 | -105 | -100 | -95 | -90 | -85 | -80 | -75 | -70 | -65 | -60 | -55 | -50 | -45 | -40 | -35 | -30 | -25 | -20 | -15 | -10 | -5 | 0 | 5 | 10 | 15 | 20 | 25 | 30 | 35 | 40 | 45 | 50 | 55 | 60 | 65 | 70 | 75 | 80 | 85 | 90 | 95 | 100 | 105 | 110 | 115 | 120 | 125 | 130 | 135 | 140 | 145 | 150 | 155 | 160 | 165 | 170 | 175 | 180 | 185 | 190 | 195 | 200 | 205 | 210 | 215 | 220 | 225 | 230 | 235 | 240 | 245 | 250 | 255 | 260 | 265 | 270 | 275 | 280 | 285 | 290 | 295 | 300 | 305 | 310 | 315 | 320 | 325 | 330 | 335 | 340 | 345 | 350 | 355 | 360 | 365 | 370 | 375 | 380 | 385 | 390 | 395 | 400 | 405 | 410 | 415 | 420 | 425 | 430 | 435 | 440 | 445 | 450 | 455 | 460 | 465 | 470 | 475 | 480 | 485 | 490 | 495 | 500 | 505 | 510 | 515 | 520 | 525 | 530 | 535 | 540 | 545 | 550 | 555 | 560 | 565 | 570 | 575 | 580 | 585 | 590 | 595 | 600 | 605 | 610 | 615 | 620 | 625 | 630 | 635 | 640 | 645 | 650 | 655 | 660 | 665 | 670 | 675 | 680 | 685 | 690 | 695 | 700 | 705 | 710 | 715 | 720 | 725 | 730 | 735 | 740 | 745 | 750 | 755 | 760 | 765 | 770 | 775 | 780 | 785 | 790 | 795 | 800 | 805 | 810 | 815 | 820 | 825 | 830 | 835 | 840 | 845 | 850 | 855 | 860 | 865 | 870 | 875 | 880 | 885 | 890 | 895 | 900 | 905 | 910 | 915 | 920 | 925 | 930 | 935 | 940 | 945 | 950 | 955 | 960 | 965 | 970 | 975 | 980 | 985 | 990 | 995 | 1000 | 1005 | 1010 | 1015 | 1020 | 1025 | 1030 | 1035 | 1040 | 1045 | 1050 | 1055 | 1060 | 1065 | 1070 | 1075 | 1080 | 1085 | 1090 | 1095 | 1100 | 1105 | 1110 | 1115 | 1120 | 1125 | 1130 | 1135 | 1140 | 1145 | 1150 | 1155 | 1160 | 1165 | 1170 | 1175 | 1180 | 1185 | 1190 | 1195 |
| --- | --- | --- | --- | --- | --- | --- | --- | --- | --- | --- | --- | --- | --- | --- | --- | --- | --- | --- | --- | --- | --- | --- | --- | --- | --- | --- | --- | --- | --- | --- | --- | --- | --- | --- | --- | --- | --- | --- | --- | --- | --- | --- | --- | --- | --- | --- | --- | --- | --- | --- | --- | --- | --- | --- | --- | --- | --- | --- | --- | --- | --- | --- | --- | --- | --- | --- | --- | --- | --- | --- | --- | --- | --- | --- | --- | --- | --- | --- | --- | --- | --- | --- | --- | --- | --- | --- | --- | --- | --- | --- | --- | --- | --- | --- | --- | --- | --- | --- | --- | --- | --- | --- | --- | --- | --- | --- | --- | --- | --- | --- | --- | --- | --- | --- | --- | --- | --- | --- | --- | --- | --- | --- | --- | --- | --- | --- | --- | --- | --- | --- | --- | --- | --- | --- | --- | --- | --- | --- | --- | --- | --- | --- | --- | --- | --- | --- | --- | --- | --- | --- | --- | --- | --- | --- | --- | --- | --- | --- | --- | --- | --- | --- | --- | --- | --- | --- | --- | --- | --- | --- | --- | --- | --- | --- | --- | --- | --- | --- | --- | --- | --- | --- | --- | --- | --- | --- | --- | --- | --- | --- | --- | --- | --- | --- | --- | --- | --- | --- | --- | --- | --- | --- | --- | --- | --- | --- | --- | --- | --- | --- | --- | --- | --- | --- | --- | --- | --- | --- | --- | --- | --- | --- | --- | --- | --- | --- | --- | --- | --- | --- | --- | --- | --- | --- | --- | --- | --- | --- | --- | --- | --- | --- | --- | --- | --- | --- | --- | --- | --- | --- | --- | --- | --- | --- | --- | --- | --- | --- | --- | --- | --- | --- | --- | --- | --- | --- | --- | --- | --- | --- | --- | --- | --- | --- | --- | --- | --- | --- | --- | --- |
| left anterior | 0.399383 | 0.370013 | 0.375238 | 0.266063 | 0.244788 | 0.224522 | 0.264183 | 0.611840 | 1.448445 | 1.274286 | 0.530182 | 0.463369 | 0.228436 | 0.226104 | 0.221669 | 0.233665 | 0.223262 | 0.233539 | 0.264177 | 0.226041 | 0.221453 | 0.222224 | 0.338513 | 0.518653 | 0.805193 | 1.509733 | 0.970592 | 0.957711 | 0.700390 | 0.350139 | 0.253802 | 0.223577 | 0.232986 | 0.233590 | 0.278189 | 0.267035 | 0.283452 | 0.429079 | 0.467565 | 0.308378 | 0.372799 | 0.434351 | 0.281074 | 0.226429 | 0.225626 | 0.237381 | 0.217517 | 0.217952 | 0.223932 | 0.260147 | 0.309935 | 0.375894 | 0.394624 | 0.391638 | 0.378675 | 0.326852 | 0.258700 | 0.222375 | 0.216157 | 0.215694 | 0.220118 | 0.216839 | 0.230620 | 0.242457 | 0.291700 | 0.327351 | 0.427243 | 0.962558 | 1.559650 | 3.581295e+00 | 3.191901e+00 | 1.749505e+00 | 2.068383e+00 | 2.338278e+00 | 2.488747 | 2.728043 | 2.476656 | 2.523796 | 2.618551 | 2.144159 | 1.468841 | 0.576110 | 0.573579 | 0.382505 | 0.260081 | 0.228659 | 0.216260 | 0.215422 | 0.223409 | 0.221680 | 0.220407 | 0.252017 | 0.371001 | 0.465108 | 0.613670 | 0.516001 | 0.360450 | 0.342733 | 0.287920 | 0.244085 | 0.243591 | 0.223484 | 0.220015 | 0.215399 | 0.214633 | 0.218221 | 0.237234 | 0.304893 | 0.285931 | 0.277051 | 0.264468 | 0.286010 | 0.284962 | 0.268702 | 0.227019 | 0.226963 | 0.221336 | 0.218036 | 0.214889 | 0.214666 | 0.218873 | 0.240898 | 0.230142 | 0.222321 | 0.215673 | 0.218419 | 0.216106 | 0.218845 | 0.218610 | 0.225917 | 0.276909 | 0.407229 | 0.381523 | 0.469670 | 0.321603 | 0.282503 | 0.306657 | 0.286401 | 0.260238 | 0.266156 | 0.253179 | 0.276822 | 0.297843 | 0.239480 | 0.215707 | 0.214728 | 0.214626 | 0.214653 | 0.222600 | 0.228964 | 0.270151 | 0.359216 | 0.355818 | 0.302783 | 0.252816 | 0.221647 | 0.224663 | 0.219145 | 0.221591 | 0.214781 | 0.214910 | 0.215358 | 0.214737 | 0.220351 | 0.226987 | 0.248135 | 0.246079 | 0.220193 | 0.226679 | 0.247117 | 0.251061 | 0.225171 | 0.214864 | 0.215540 | 0.214797 | 0.215748 | 0.218363 | 0.223745 | 0.219140 | 0.216157 | 0.214845 | 0.226149 | 0.255276 | 0.263103 | 0.239706 | 0.269523 | 0.288111 | 0.256412 | 0.241440 | 0.267906 | 0.384541 | 0.546054 | 0.555740 | 0.809394 | 0.814404 | 0.465059 | 0.263070 | 0.214635 | 0.218202 | 0.243563 | 0.287619 | 0.243582 | 0.223970 | 0.221127 | 0.220968 | 0.246517 | 0.241036 | 0.302559 | 0.386330 | 0.572234 | 0.617206 | 0.683591 | 0.492154 | 0.440904 | 0.315730 | 0.294539 | 0.243914 | 0.232615 | 0.217609 | 0.217388 | 0.219441 | 0.215254 | 0.215678 | 0.229992 | 0.271381 | 0.344524 | 0.381743 | 0.353009 | 0.428907 | 0.396415 | 0.354419 | 0.288757 | 0.235046 | 0.218205 | 0.215712 | 0.221818 | 0.231801 | 0.226868 | 0.218995 | 0.215440 | 0.216085 | 0.220695 | 0.216995 | 0.216489 | 0.219323 | 0.219276 | 0.220189 | 0.218875 | 0.214646 | 0.216972 | 0.223950 | 0.234201 | 0.240470 | 0.247824 | 0.296702 | 0.305822 | 0.274295 | 0.272810 | 0.334138 | 0.355861 | 0.337758 | 0.279084 | 0.245811 | 0.221727 | 0.223506 | 0.214960 | 0.221704 | 0.239385 | 0.254180 | 0.247409 | 0.224371 | 0.221359 | 0.214964 | 0.225551 | 0.262537 | 0.274667 | 0.270618 | 0.275509 | 0.251809 | 0.226635 |
| right anterior | 0.215103 | 0.216118 | 0.223357 | 0.216321 | 0.217176 | 0.214908 | 0.218368 | 0.216716 | 0.214634 | 0.219163 | 0.234633 | 0.219178 | 0.215572 | 0.215054 | 0.218514 | 0.218049 | 0.215751 | 0.241125 | 0.259294 | 0.223364 | 0.216048 | 0.216437 | 0.221468 | 0.269540 | 0.265519 | 0.218984 | 0.227729 | 0.244177 | 0.243524 | 0.275682 | 0.296591 | 0.301189 | 0.412151 | 0.402914 | 0.413429 | 0.764097 | 0.585667 | 0.505904 | 0.279114 | 0.216341 | 0.260606 | 0.343112 | 0.483831 | 0.356088 | 0.252983 | 0.215686 | 0.255329 | 0.375408 | 1.001671 | 1.000365 | 0.773032 | 0.416582 | 0.309908 | 0.284048 | 0.418005 | 0.551980 | 1.080370 | 2.098170 | 4.939217 | 6.603986 | 5.007346 | 2.576925 | 1.498716 | 1.216620 | 1.007142 | 0.925795 | 1.737296 | 5.843686 | 17.279147 | 3.721131e+01 | 4.671734e+01 | 1.955876e+01 | 8.211156e+00 | 6.965791e+00 | 4.197390 | 2.644383 | 0.504288 | 0.335076 | 0.342480 | 0.334454 | 0.271611 | 0.217764 | 0.238640 | 0.237273 | 0.259019 | 0.263055 | 0.230326 | 0.216681 | 0.216707 | 0.217826 | 0.224366 | 0.229067 | 0.217874 | 0.214634 | 0.216005 | 0.214744 | 0.222659 | 0.234928 | 0.299412 | 0.327265 | 0.334925 | 0.367781 | 0.385616 | 0.367857 | 0.312587 | 0.231746 | 0.219632 | 0.218216 | 0.215494 | 0.219660 | 0.220274 | 0.221887 | 0.229073 | 0.234444 | 0.229798 | 0.215658 | 0.216784 | 0.250163 | 0.309504 | 0.538692 | 0.678973 | 0.920183 | 0.692746 | 0.662134 | 0.503584 | 0.469559 | 0.409727 | 0.307216 | 0.241613 | 0.241402 | 0.252873 | 0.243350 | 0.232918 | 0.232379 | 0.278130 | 0.304654 | 0.265096 | 0.276547 | 0.267549 | 0.251378 | 0.261685 | 0.274525 | 0.330830 | 0.350832 | 0.298323 | 0.290080 | 0.301694 | 0.264505 | 0.250559 | 0.229793 | 0.224517 | 0.221632 | 0.224343 | 0.227595 | 0.222932 | 0.218881 | 0.218887 | 0.233029 | 0.234663 | 0.238042 | 0.248958 | 0.248136 | 0.281911 | 0.308547 | 0.264014 | 0.256991 | 0.246537 | 0.232627 | 0.224463 | 0.227425 | 0.218143 | 0.219133 | 0.224062 | 0.227422 | 0.235596 | 0.284556 | 0.300072 | 0.311433 | 0.323106 | 0.322045 | 0.337710 | 0.354318 | 0.320138 | 0.276321 | 0.293725 | 0.336074 | 0.397278 | 0.394383 | 0.349105 | 0.396583 | 0.468912 | 0.710335 | 0.721017 | 0.629479 | 0.607468 | 0.589551 | 0.445539 | 0.394308 | 0.311652 | 0.306942 | 0.368386 | 0.541504 | 0.845668 | 1.308182 | 1.081388 | 0.811727 | 0.557063 | 0.454358 | 0.348653 | 0.279916 | 0.278954 | 0.368035 | 0.438336 | 0.497077 | 0.390805 | 0.332376 | 0.304553 | 0.284748 | 0.243097 | 0.232304 | 0.226301 | 0.236547 | 0.245792 | 0.261841 | 0.252039 | 0.295042 | 0.382339 | 0.467000 | 0.445093 | 0.440581 | 0.427298 | 0.341716 | 0.262277 | 0.235746 | 0.231767 | 0.250262 | 0.249437 | 0.237088 | 0.253647 | 0.271074 | 0.278230 | 0.274339 | 0.276804 | 0.299731 | 0.345876 | 0.351897 | 0.345300 | 0.314762 | 0.336104 | 0.316013 | 0.338593 | 0.346197 | 0.320381 | 0.291625 | 0.308911 | 0.294081 | 0.301555 | 0.320973 | 0.465944 | 0.817777 | 1.666415 | 2.179202 | 2.487063 | 2.328588 | 1.363455 | 0.560572 | 0.349398 | 0.285271 | 0.268219 | 0.251227 | 0.259097 | 0.295241 | 0.479898 | 0.652114 | 0.580289 | 0.660755 | 0.663077 | 0.627253 | 0.580559 | 0.375528 |
| left central | 0.334888 | 0.375222 | 0.370268 | 0.312445 | 0.268119 | 0.235284 | 0.216144 | 0.230548 | 0.236261 | 0.217415 | 0.216225 | 0.303078 | 0.495406 | 0.572500 | 0.614780 | 0.551605 | 0.477720 | 0.551216 | 0.484408 | 0.584914 | 0.781737 | 0.487021 | 0.648978 | 0.714563 | 0.333101 | 0.214737 | 0.259026 | 0.261062 | 0.225702 | 0.329977 | 0.384729 | 0.485921 | 0.324769 | 0.306871 | 0.360469 | 0.360700 | 0.275909 | 0.273308 | 0.220115 | 0.216225 | 0.247187 | 0.272891 | 0.274270 | 0.250809 | 0.291520 | 0.316861 | 0.276632 | 0.256759 | 0.271851 | 0.307836 | 0.440349 | 0.399575 | 0.398124 | 0.458410 | 0.402745 | 0.265285 | 0.219550 | 0.219662 | 0.218664 | 0.219068 | 0.215680 | 0.217398 | 0.228460 | 0.216200 | 0.223789 | 0.286367 | 0.514989 | 1.435233 | 7.159941 | 2.699167e+01 | 5.563887e+01 | 4.644247e+01 | 7.644103e+01 | 7.483607e+01 | 8.254714 | 1.070169 | 0.465014 | 0.287481 | 0.229889 | 0.217520 | 0.234373 | 0.282552 | 0.268119 | 0.311170 | 0.323190 | 0.375220 | 0.408985 | 0.394891 | 0.236946 | 0.217538 | 0.214705 | 0.218437 | 0.219734 | 0.278921 | 0.412141 | 0.476322 | 0.705556 | 1.495868 | 1.967075 | 2.453116 | 2.059875 | 1.125276 | 0.540787 | 0.396201 | 0.295528 | 0.310428 | 0.265932 | 0.216391 | 0.218204 | 0.246340 | 0.367997 | 0.437148 | 0.429462 | 0.373296 | 0.270603 | 0.237093 | 0.218008 | 0.214870 | 0.224929 | 0.242048 | 0.253481 | 0.269782 | 0.253763 | 0.273319 | 0.300588 | 0.309127 | 0.255780 | 0.263762 | 0.234704 | 0.223971 | 0.215381 | 0.214688 | 0.224738 | 0.223780 | 0.225918 | 0.229991 | 0.225744 | 0.223412 | 0.225909 | 0.216349 | 0.215243 | 0.216584 | 0.215967 | 0.219546 | 0.228859 | 0.228291 | 0.214655 | 0.249392 | 0.278764 | 0.238148 | 0.254753 | 0.264655 | 0.300046 | 0.299877 | 0.270962 | 0.323119 | 0.436756 | 0.366879 | 0.308423 | 0.234849 | 0.217735 | 0.215371 | 0.219669 | 0.222436 | 0.223453 | 0.223517 | 0.221708 | 0.221483 | 0.218030 | 0.218020 | 0.214832 | 0.215954 | 0.226839 | 0.232598 | 0.222859 | 0.216292 | 0.214885 | 0.225351 | 0.249200 | 0.270172 | 0.249539 | 0.220361 | 0.215597 | 0.232528 | 0.296612 | 0.402501 | 0.520235 | 0.422008 | 0.281907 | 0.229099 | 0.216271 | 0.215936 | 0.222512 | 0.222975 | 0.215024 | 0.216437 | 0.228233 | 0.249817 | 0.256969 | 0.267197 | 0.261811 | 0.258469 | 0.242094 | 0.250605 | 0.256443 | 0.305551 | 0.319455 | 0.294143 | 0.248794 | 0.247282 | 0.219932 | 0.215882 | 0.229801 | 0.227811 | 0.217243 | 0.216084 | 0.216701 | 0.216085 | 0.214715 | 0.215097 | 0.219114 | 0.225591 | 0.231930 | 0.231895 | 0.233401 | 0.246063 | 0.271760 | 0.260954 | 0.243009 | 0.250448 | 0.245417 | 0.243602 | 0.242717 | 0.239422 | 0.229378 | 0.225057 | 0.215863 | 0.218559 | 0.220138 | 0.215490 | 0.218883 | 0.221469 | 0.222972 | 0.215656 | 0.214630 | 0.214660 | 0.214660 | 0.214641 | 0.216434 | 0.222239 | 0.236352 | 0.287031 | 0.320007 | 0.342153 | 0.332418 | 0.286909 | 0.274310 | 0.276016 | 0.286005 | 0.276804 | 0.288551 | 0.273059 | 0.275988 | 0.261332 | 0.248376 | 0.234462 | 0.239149 | 0.225448 | 0.236399 | 0.254811 | 0.287513 | 0.317467 | 0.291238 | 0.264150 | 0.243797 | 0.216369 | 0.221422 | 0.244853 | 0.281721 | 0.292906 |
| right central | 0.214736 | 0.221421 | 0.222231 | 0.237111 | 0.217710 | 0.217516 | 0.216941 | 0.222515 | 0.243015 | 0.282002 | 0.329764 | 0.331017 | 0.261806 | 0.370099 | 0.384468 | 0.518967 | 0.671142 | 1.152377 | 1.073280 | 0.744650 | 0.419791 | 0.246484 | 0.229522 | 0.214882 | 0.233528 | 0.245196 | 0.218154 | 0.222706 | 0.229506 | 0.248934 | 0.237528 | 0.233243 | 0.306575 | 0.420374 | 0.404293 | 0.466703 | 0.329137 | 0.302981 | 0.246687 | 0.244708 | 0.273865 | 0.336647 | 0.324130 | 0.541197 | 0.761968 | 2.366679 | 3.023244 | 1.811482 | 0.678866 | 0.543020 | 0.632271 | 1.157148 | 0.843826 | 0.539053 | 0.506302 | 0.504753 | 0.433530 | 0.318584 | 0.253475 | 0.236168 | 0.239450 | 0.223565 | 0.214632 | 0.231302 | 0.320940 | 0.593778 | 2.135628 | 11.569076 | 63.125484 | 5.127631e+02 | 2.557572e+03 | 1.071215e+04 | 7.818170e+03 | 7.171568e+03 | 1900.120685 | 281.772966 | 22.985157 | 2.727614 | 0.619023 | 0.447762 | 0.272300 | 0.216064 | 0.230048 | 0.239390 | 0.260429 | 0.269634 | 0.310202 | 0.258019 | 0.214626 | 0.257463 | 0.365559 | 0.614085 | 1.129721 | 2.074959 | 2.562652 | 1.317420 | 0.457004 | 0.251669 | 0.224482 | 0.229393 | 0.248154 | 0.288589 | 0.313312 | 0.386991 | 0.543344 | 0.618824 | 0.789924 | 0.852141 | 0.737091 | 0.786327 | 1.832607 | 4.154339 | 11.577765 | 7.890728 | 9.006221 | 9.135919 | 5.235899 | 2.127804 | 1.977932 | 2.195054 | 5.019680 | 7.713622 | 12.223374 | 22.014653 | 45.893867 | 30.711765 | 30.793805 | 26.811251 | 19.854207 | 10.290745 | 4.827030 | 2.556289 | 2.312319 | 3.160681 | 1.847971 | 1.787199 | 1.865820 | 1.981459 | 2.377067 | 2.046712 | 1.323293 | 1.000747 | 0.942419 | 0.657840 | 0.623022 | 0.663053 | 0.976937 | 1.382960 | 1.804328 | 1.342826 | 2.053562 | 2.759717 | 2.100360 | 1.336638 | 0.774219 | 0.777709 | 1.005388 | 0.779912 | 0.859383 | 1.521334 | 1.332502 | 1.093817 | 0.841173 | 0.725899 | 0.756144 | 0.620565 | 0.499145 | 0.452678 | 0.402458 | 0.416169 | 0.375753 | 0.308964 | 0.272082 | 0.236200 | 0.243920 | 0.299752 | 0.319289 | 0.357575 | 0.380091 | 0.409170 | 0.516369 | 0.523933 | 0.310562 | 0.240365 | 0.217629 | 0.216799 | 0.218287 | 0.215476 | 0.215038 | 0.252515 | 0.281942 | 0.328720 | 0.308510 | 0.273346 | 0.249357 | 0.235678 | 0.224732 | 0.221182 | 0.214683 | 0.217238 | 0.223679 | 0.225670 | 0.229443 | 0.236865 | 0.241784 | 0.229452 | 0.221372 | 0.216922 | 0.216117 | 0.215271 | 0.216637 | 0.214684 | 0.216789 | 0.222194 | 0.218935 | 0.217407 | 0.221130 | 0.245468 | 0.278371 | 0.275944 | 0.251198 | 0.245884 | 0.234940 | 0.220826 | 0.214626 | 0.225629 | 0.235902 | 0.246495 | 0.239712 | 0.226263 | 0.216196 | 0.216236 | 0.223393 | 0.227243 | 0.244332 | 0.279121 | 0.295428 | 0.279233 | 0.257103 | 0.263003 | 0.285971 | 0.288350 | 0.251672 | 0.220976 | 0.215643 | 0.217305 | 0.226700 | 0.238724 | 0.235706 | 0.227702 | 0.218964 | 0.219577 | 0.215008 | 0.215032 | 0.215212 | 0.216636 | 0.214675 | 0.220362 | 0.226253 | 0.232073 | 0.233470 | 0.244853 | 0.245769 | 0.230370 | 0.239442 | 0.267557 | 0.315130 | 0.318765 | 0.349053 | 0.385676 | 0.521699 | 0.512947 | 0.445751 | 0.400780 | 0.435340 | 0.440071 | 0.487166 | 0.424799 | 0.382628 | 0.399744 |
| left posterior | 0.338580 | 0.384341 | 0.313331 | 0.270891 | 0.271166 | 0.247713 | 0.226885 | 0.248790 | 0.273440 | 0.400097 | 0.665028 | 1.414726 | 2.350637 | 4.041948 | 1.569238 | 1.045230 | 0.810576 | 1.148257 | 0.459355 | 0.300742 | 0.358570 | 0.837472 | 1.190158 | 0.933552 | 0.296377 | 0.262216 | 0.222884 | 0.222810 | 0.355792 | 1.093007 | 2.029952 | 1.873031 | 1.231547 | 0.991669 | 1.016793 | 0.511765 | 0.248668 | 0.214633 | 0.221489 | 0.235013 | 0.288912 | 0.335710 | 0.280551 | 0.222872 | 0.214636 | 0.226261 | 0.245036 | 0.251854 | 0.237564 | 0.215459 | 0.230244 | 0.289699 | 0.470553 | 1.220530 | 1.934330 | 2.556179 | 1.631603 | 1.529303 | 1.488220 | 1.297831 | 0.507855 | 0.271609 | 0.219874 | 0.393601 | 2.226385 | 15.919162 | 244.584687 | 2505.468852 | 9971.079126 | 2.218263e+04 | 2.003793e+04 | 7.304913e+03 | 3.935132e+03 | 9.807277e+02 | 195.679554 | 27.676005 | 3.661902 | 0.707977 | 0.309429 | 0.226966 | 0.215932 | 0.243277 | 0.270531 | 0.322522 | 0.339778 | 0.358307 | 0.363531 | 0.314965 | 0.225125 | 0.239947 | 0.450488 | 0.836539 | 1.846158 | 2.668446 | 10.225957 | 12.519669 | 8.276493 | 7.306755 | 6.525622 | 5.406586 | 5.875100 | 1.165962 | 0.558152 | 0.425491 | 0.319845 | 0.280064 | 0.233307 | 0.247000 | 0.329840 | 0.363047 | 0.370386 | 0.442674 | 0.495755 | 0.679066 | 0.829439 | 0.707179 | 0.573498 | 0.458825 | 0.356578 | 0.408490 | 0.563051 | 0.817811 | 1.280682 | 1.665079 | 2.585485 | 3.528873 | 4.332539 | 3.332520 | 2.223777 | 2.059316 | 2.075938 | 1.690153 | 1.523753 | 1.212973 | 1.049280 | 0.891586 | 0.920958 | 1.149984 | 1.202598 | 1.507028 | 1.484481 | 1.639109 | 3.175416 | 3.985919 | 4.726061 | 7.128015 | 7.108365 | 6.898010 | 5.898247 | 3.302146 | 2.349016 | 1.308875 | 1.120162 | 0.932872 | 0.802333 | 0.745963 | 0.711907 | 0.720229 | 0.818158 | 0.800532 | 0.859619 | 0.808282 | 0.812923 | 0.754035 | 0.878535 | 0.984073 | 0.966856 | 1.127180 | 1.527844 | 1.487831 | 2.196328 | 3.026330 | 4.068939 | 8.186049 | 12.630828 | 15.947375 | 16.522710 | 13.496341 | 5.789467 | 2.488313 | 1.773015 | 1.770329 | 1.459190 | 1.279932 | 1.227086 | 1.857613 | 4.274402 | 8.311015 | 9.297778 | 10.046469 | 13.334370 | 15.834936 | 15.799708 | 11.835297 | 7.439327 | 7.537801 | 6.540525 | 8.898102 | 10.650650 | 12.108760 | 11.430186 | 11.134357 | 8.127333 | 8.222901 | 6.479405 | 7.129784 | 6.746155 | 6.668686 | 9.865990 | 16.464637 | 16.309307 | 8.152253 | 3.419600 | 2.555719 | 1.872991 | 1.752376 | 1.607235 | 1.197613 | 1.324562 | 1.359639 | 1.297403 | 1.764996 | 1.780812 | 1.409118 | 1.314161 | 1.118693 | 1.287643 | 2.017913 | 2.598641 | 3.240122 | 8.068897 | 21.632392 | 63.865252 | 117.570659 | 122.197683 | 201.896256 | 213.821629 | 186.476446 | 222.582694 | 319.969219 | 196.597088 | 67.568431 | 31.261572 | 19.608354 | 5.963630 | 2.523523 | 1.616542 | 1.307849 | 1.753544 | 2.399436 | 1.934732 | 1.620288 | 1.364052 | 0.827918 | 0.626219 | 0.409310 | 0.403469 | 0.488698 | 0.705034 | 0.994259 | 1.354719 | 1.894242 | 2.528364 | 1.414832 | 1.382868 | 0.845107 | 0.538309 | 0.427743 | 0.334430 | 0.359902 | 0.461776 | 0.642475 | 0.947208 | 1.297015 | 1.662752 | 2.338734 | 3.233124 | 3.227233 | 2.297778 | 1.617608 |
| right posterior | 0.215051 | 0.214725 | 0.215774 | 0.219737 | 0.215757 | 0.224341 | 0.305404 | 0.261157 | 0.226010 | 0.217564 | 0.218535 | 0.224621 | 0.286581 | 0.387896 | 0.282873 | 0.239252 | 0.219738 | 0.214803 | 0.221397 | 0.223623 | 0.215467 | 0.286690 | 0.305668 | 0.265863 | 0.281583 | 0.357772 | 0.456727 | 0.472442 | 0.475675 | 0.529300 | 0.450595 | 0.337830 | 0.311656 | 0.256265 | 0.215928 | 0.249610 | 0.324736 | 0.319406 | 0.302297 | 0.301934 | 0.255404 | 0.215424 | 0.235180 | 0.292397 | 0.303611 | 0.321095 | 0.315007 | 0.305217 | 0.241699 | 0.229063 | 0.245434 | 0.257884 | 0.254267 | 0.250802 | 0.239082 | 0.247532 | 0.229055 | 0.214654 | 0.217373 | 0.214783 | 0.215105 | 0.225441 | 0.302457 | 0.584363 | 1.657536 | 8.811224 | 111.713102 | 2675.821001 | 114170.801547 | 2.200017e+06 | 1.512359e+07 | 4.976508e+07 | 2.147234e+07 | 3.634800e+06 | 767939.340609 | 121246.032521 | 6888.359895 | 22.768833 | 1.304007 | 0.342332 | 0.214656 | 0.337357 | 0.602574 | 0.673631 | 0.490834 | 0.575039 | 0.599145 | 0.504842 | 0.352150 | 0.306080 | 0.250422 | 0.223405 | 0.216459 | 0.222220 | 0.232115 | 0.234885 | 0.243836 | 0.234555 | 0.221775 | 0.214719 | 0.215229 | 0.215108 | 0.218104 | 0.227246 | 0.219912 | 0.215206 | 0.231710 | 0.246307 | 0.261610 | 0.287925 | 0.331510 | 0.364332 | 0.344804 | 0.345388 | 0.402813 | 0.447628 | 0.635007 | 0.899634 | 1.649149 | 2.789059 | 3.300755 | 2.988817 | 2.313623 | 2.306444 | 2.128244 | 1.192222 | 0.765914 | 0.652353 | 0.605813 | 0.562270 | 0.479766 | 0.490634 | 0.547212 | 0.768541 | 1.083833 | 1.685373 | 2.690865 | 4.341424 | 5.363008 | 7.815673 | 5.775174 | 5.258942 | 4.435169 | 2.425767 | 1.406876 | 0.677527 | 0.372652 | 0.276593 | 0.229572 | 0.214956 | 0.234132 | 0.281815 | 0.270704 | 0.251965 | 0.236894 | 0.222966 | 0.215679 | 0.217250 | 0.236250 | 0.281018 | 0.398409 | 0.526705 | 0.542191 | 0.698238 | 0.778182 | 0.740610 | 0.522698 | 0.367135 | 0.287415 | 0.276751 | 0.250842 | 0.234971 | 0.243444 | 0.270356 | 0.279840 | 0.320031 | 0.372580 | 0.381715 | 0.404176 | 0.323579 | 0.294413 | 0.315380 | 0.340597 | 0.396258 | 0.467173 | 0.605061 | 0.891288 | 1.191456 | 1.003124 | 0.955999 | 0.707910 | 0.521766 | 0.425246 | 0.416042 | 0.498434 | 0.789828 | 1.058504 | 1.067600 | 1.199078 | 1.236718 | 0.866145 | 0.638694 | 0.510836 | 0.436068 | 0.429708 | 0.422645 | 0.368635 | 0.334694 | 0.289039 | 0.270614 | 0.260177 | 0.245280 | 0.268106 | 0.290418 | 0.299540 | 0.295955 | 0.272944 | 0.290534 | 0.311823 | 0.263804 | 0.237624 | 0.232465 | 0.235303 | 0.248320 | 0.228596 | 0.214965 | 0.218152 | 0.221823 | 0.239570 | 0.265558 | 0.271331 | 0.254099 | 0.236134 | 0.236968 | 0.243675 | 0.237672 | 0.235683 | 0.240188 | 0.254197 | 0.246017 | 0.219035 | 0.214696 | 0.215294 | 0.215706 | 0.215148 | 0.217175 | 0.226072 | 0.248483 | 0.238108 | 0.230704 | 0.228736 | 0.221181 | 0.236609 | 0.249039 | 0.249399 | 0.296552 | 0.366257 | 0.363552 | 0.355918 | 0.277348 | 0.242914 | 0.239079 | 0.224147 | 0.219486 | 0.224104 | 0.237163 | 0.239059 | 0.236309 | 0.239543 | 0.242941 | 0.243206 | 0.226050 | 0.215789 | 0.214834 | 0.214779 | 0.218392 | 0.222933 | 0.230346 | 0.226069 | 0.223300 |
| all electrodes | 0.410586 | 0.316266 | 0.310221 | 0.259747 | 0.226008 | 0.234067 | 0.229988 | 0.232074 | 0.214637 | 0.215362 | 0.268445 | 0.422858 | 0.622170 | 0.358901 | 0.250039 | 0.219959 | 0.214626 | 0.235322 | 0.309173 | 0.271019 | 0.222755 | 0.340043 | 0.284113 | 0.297866 | 0.305183 | 0.280491 | 0.225901 | 0.215152 | 0.217110 | 0.221613 | 0.255588 | 0.284116 | 0.264710 | 0.272219 | 0.551268 | 1.126942 | 0.509411 | 0.421275 | 0.303662 | 0.282046 | 0.247365 | 0.227222 | 0.345688 | 0.334850 | 0.295932 | 0.258933 | 0.270759 | 0.287913 | 0.258167 | 0.228706 | 0.229631 | 0.260804 | 0.264291 | 0.301923 | 0.275566 | 0.219667 | 0.220049 | 0.228664 | 0.327452 | 0.353430 | 0.553137 | 0.853106 | 0.879128 | 2.270687 | 3.995122 | 10.315951 | 25.739949 | 160.700234 | 1024.272484 | 7.374418e+03 | 4.986997e+04 | 1.530330e+05 | 1.191669e+05 | 7.508343e+04 | 9729.978027 | 2194.794029 | 309.407455 | 44.478836 | 14.828330 | 4.081899 | 2.164733 | 0.766964 | 0.565733 | 0.535903 | 0.387495 | 0.352576 | 0.317177 | 0.273495 | 0.287029 | 0.396050 | 0.315886 | 0.360050 | 0.389538 | 0.599063 | 1.126284 | 2.427261 | 2.775933 | 10.855531 | 8.582784 | 6.160950 | 3.424730 | 2.720591 | 1.617155 | 1.320093 | 1.117453 | 1.466524 | 1.000205 | 1.012770 | 0.969600 | 1.165996 | 1.234189 | 1.098700 | 1.070577 | 1.778710 | 2.339909 | 3.407830 | 4.683759 | 5.449075 | 10.193720 | 26.859497 | 43.062894 | 84.181642 | 154.211549 | 272.294227 | 272.201542 | 160.120854 | 169.636219 | 251.169935 | 126.738444 | 59.510122 | 46.741875 | 57.865257 | 77.886738 | 21.921233 | 8.123934 | 5.456980 | 3.509155 | 4.479799 | 3.485875 | 3.233023 | 3.675731 | 4.503683 | 5.389002 | 8.006020 | 4.072890 | 2.012325 | 0.901369 | 0.550634 | 0.469985 | 0.445171 | 0.337308 | 0.235202 | 0.216973 | 0.221631 | 0.241230 | 0.298990 | 0.405327 | 0.588957 | 1.092798 | 1.356924 | 2.056821 | 2.586904 | 1.273875 | 0.797699 | 0.883318 | 1.056530 | 1.188158 | 0.592743 | 0.401216 | 0.402577 | 0.352873 | 0.286296 | 0.292132 | 0.324754 | 0.429889 | 0.470971 | 0.491237 | 0.480268 | 0.453809 | 0.356756 | 0.363082 | 0.335114 | 0.342466 | 0.370961 | 0.391603 | 0.621900 | 1.075450 | 1.334179 | 2.086908 | 1.961460 | 1.419238 | 1.350516 | 0.948560 | 0.762739 | 0.626822 | 0.453140 | 0.431684 | 0.420323 | 0.394621 | 0.315335 | 0.246665 | 0.223350 | 0.230004 | 0.243616 | 0.248320 | 0.240392 | 0.238611 | 0.250921 | 0.267557 | 0.289120 | 0.310299 | 0.342263 | 0.376776 | 0.376062 | 0.403996 | 0.426525 | 0.333768 | 0.261800 | 0.232258 | 0.229783 | 0.263431 | 0.277269 | 0.258605 | 0.285318 | 0.367659 | 0.361207 | 0.323766 | 0.283754 | 0.344094 | 0.422487 | 0.423664 | 0.358720 | 0.359162 | 0.413058 | 0.429383 | 0.401745 | 0.400500 | 0.391592 | 0.406555 | 0.459882 | 0.459334 | 0.589367 | 0.652693 | 0.710616 | 0.756749 | 0.688676 | 0.778409 | 0.681672 | 0.693056 | 0.681133 | 0.604067 | 0.686756 | 0.519321 | 0.392914 | 0.358515 | 0.331290 | 0.277214 | 0.265254 | 0.244322 | 0.283717 | 0.342604 | 0.355172 | 0.293591 | 0.279681 | 0.244814 | 0.230163 | 0.221166 | 0.214789 | 0.225908 | 0.224102 | 0.225175 | 0.214732 | 0.217481 | 0.218594 | 0.226608 | 0.248190 | 0.251744 | 0.266583 | 0.242487 | 0.238723 |

Searchlight, spatiotemporal cluster permutation test

|  | start time | stop time | peak time | peak channel | cluster p | peak Cohen's d | direction |
| --- | --- | --- | --- | --- | --- | --- | --- |
| #1 | 75 | 215 | 155 | O2 | 0.0065 | 1.532777 | positive |
| #2 | 235 | 555 | 295 | TP7 | 0.0173 | 1.023348 | positive |
| #3 | 600 | 1060 | 685 | PO3 | 0.0229 | 0.98815 | positive |

W) emoji to real faces - happy vs angry

  
|  | time window | peak latency | cluster *p* | peak Cohen's *d* |  | | | |
| **all electrodes** |  | | | |  | | | |
|  | | | | | | | | |

Time-resolved classification, cluster permutation tests

|  | **left hemisphere** | | | | **right hemisphere** | | | |
|  | time window | peak latency | cluster *p* | peak Cohen's *d* | time window | peak latency | cluster *p* | peak Cohen's *d* |
| **anterior** |  | | | |  | | | |
| **central** |  | | | | 135 - 210 ms | 165 ms | 0.0418 | 1.0786 |
| **posterior** |  | | | | 145 - 200 ms | 170 ms | 0.0422 | 1.1285 |
  | | | | 590 - 780 ms | 645 ms | 0.0054 | 0.8484 |

  

Time-resolved classification, Bayesian statistics

|  | -200 | -195 | -190 | -185 | -180 | -175 | -170 | -165 | -160 | -155 | -150 | -145 | -140 | -135 | -130 | -125 | -120 | -115 | -110 | -105 | -100 | -95 | -90 | -85 | -80 | -75 | -70 | -65 | -60 | -55 | -50 | -45 | -40 | -35 | -30 | -25 | -20 | -15 | -10 | -5 | 0 | 5 | 10 | 15 | 20 | 25 | 30 | 35 | 40 | 45 | 50 | 55 | 60 | 65 | 70 | 75 | 80 | 85 | 90 | 95 | 100 | 105 | 110 | 115 | 120 | 125 | 130 | 135 | 140 | 145 | 150 | 155 | 160 | 165 | 170 | 175 | 180 | 185 | 190 | 195 | 200 | 205 | 210 | 215 | 220 | 225 | 230 | 235 | 240 | 245 | 250 | 255 | 260 | 265 | 270 | 275 | 280 | 285 | 290 | 295 | 300 | 305 | 310 | 315 | 320 | 325 | 330 | 335 | 340 | 345 | 350 | 355 | 360 | 365 | 370 | 375 | 380 | 385 | 390 | 395 | 400 | 405 | 410 | 415 | 420 | 425 | 430 | 435 | 440 | 445 | 450 | 455 | 460 | 465 | 470 | 475 | 480 | 485 | 490 | 495 | 500 | 505 | 510 | 515 | 520 | 525 | 530 | 535 | 540 | 545 | 550 | 555 | 560 | 565 | 570 | 575 | 580 | 585 | 590 | 595 | 600 | 605 | 610 | 615 | 620 | 625 | 630 | 635 | 640 | 645 | 650 | 655 | 660 | 665 | 670 | 675 | 680 | 685 | 690 | 695 | 700 | 705 | 710 | 715 | 720 | 725 | 730 | 735 | 740 | 745 | 750 | 755 | 760 | 765 | 770 | 775 | 780 | 785 | 790 | 795 | 800 | 805 | 810 | 815 | 820 | 825 | 830 | 835 | 840 | 845 | 850 | 855 | 860 | 865 | 870 | 875 | 880 | 885 | 890 | 895 | 900 | 905 | 910 | 915 | 920 | 925 | 930 | 935 | 940 | 945 | 950 | 955 | 960 | 965 | 970 | 975 | 980 | 985 | 990 | 995 | 1000 | 1005 | 1010 | 1015 | 1020 | 1025 | 1030 | 1035 | 1040 | 1045 | 1050 | 1055 | 1060 | 1065 | 1070 | 1075 | 1080 | 1085 | 1090 | 1095 | 1100 | 1105 | 1110 | 1115 | 1120 | 1125 | 1130 | 1135 | 1140 | 1145 | 1150 | 1155 | 1160 | 1165 | 1170 | 1175 | 1180 | 1185 | 1190 | 1195 |
| --- | --- | --- | --- | --- | --- | --- | --- | --- | --- | --- | --- | --- | --- | --- | --- | --- | --- | --- | --- | --- | --- | --- | --- | --- | --- | --- | --- | --- | --- | --- | --- | --- | --- | --- | --- | --- | --- | --- | --- | --- | --- | --- | --- | --- | --- | --- | --- | --- | --- | --- | --- | --- | --- | --- | --- | --- | --- | --- | --- | --- | --- | --- | --- | --- | --- | --- | --- | --- | --- | --- | --- | --- | --- | --- | --- | --- | --- | --- | --- | --- | --- | --- | --- | --- | --- | --- | --- | --- | --- | --- | --- | --- | --- | --- | --- | --- | --- | --- | --- | --- | --- | --- | --- | --- | --- | --- | --- | --- | --- | --- | --- | --- | --- | --- | --- | --- | --- | --- | --- | --- | --- | --- | --- | --- | --- | --- | --- | --- | --- | --- | --- | --- | --- | --- | --- | --- | --- | --- | --- | --- | --- | --- | --- | --- | --- | --- | --- | --- | --- | --- | --- | --- | --- | --- | --- | --- | --- | --- | --- | --- | --- | --- | --- | --- | --- | --- | --- | --- | --- | --- | --- | --- | --- | --- | --- | --- | --- | --- | --- | --- | --- | --- | --- | --- | --- | --- | --- | --- | --- | --- | --- | --- | --- | --- | --- | --- | --- | --- | --- | --- | --- | --- | --- | --- | --- | --- | --- | --- | --- | --- | --- | --- | --- | --- | --- | --- | --- | --- | --- | --- | --- | --- | --- | --- | --- | --- | --- | --- | --- | --- | --- | --- | --- | --- | --- | --- | --- | --- | --- | --- | --- | --- | --- | --- | --- | --- | --- | --- | --- | --- | --- | --- | --- | --- | --- | --- | --- | --- | --- | --- | --- | --- | --- | --- | --- | --- | --- | --- | --- | --- | --- | --- | --- | --- | --- | --- | --- | --- | --- | --- |
| left anterior | 0.547347 | 0.439564 | 0.415649 | 0.429963 | 0.442646 | 0.290236 | 0.245824 | 0.214665 | 0.217485 | 0.231570 | 0.274331 | 0.381967 | 0.327642 | 0.396342 | 0.442429 | 0.446048 | 0.479057 | 0.270853 | 0.224146 | 0.231012 | 0.241885 | 0.214659 | 0.228510 | 0.214654 | 0.265289 | 0.444128 | 0.490575 | 0.389640 | 0.691137 | 0.920536 | 0.445802 | 0.258889 | 0.215211 | 0.230391 | 0.215646 | 0.221291 | 0.220827 | 0.249776 | 0.252712 | 0.332790 | 0.478359 | 0.935745 | 0.906352 | 0.519785 | 0.430675 | 0.308329 | 0.245558 | 0.221169 | 0.215174 | 0.222560 | 0.254470 | 0.327643 | 0.529998 | 0.556665 | 0.620239 | 0.503565 | 0.344168 | 0.313685 | 0.262620 | 0.285997 | 0.319759 | 0.321204 | 0.489373 | 0.611377 | 0.413266 | 0.426386 | 0.277907 | 0.225793 | 0.233737 | 0.439356 | 0.898484 | 1.137582 | 1.964295 | 1.736355 | 0.910001 | 0.583754 | 0.484321 | 0.312292 | 0.282412 | 0.313811 | 0.328036 | 0.298987 | 0.302891 | 0.250206 | 0.233392 | 0.214865 | 0.237519 | 0.272901 | 0.312931 | 0.488174 | 0.469972 | 0.412142 | 0.259206 | 0.251403 | 0.221378 | 0.217069 | 0.216425 | 0.219224 | 0.216736 | 0.251993 | 0.234788 | 0.233129 | 0.220608 | 0.216624 | 0.214912 | 0.214822 | 0.216897 | 0.214634 | 0.215713 | 0.215476 | 0.214766 | 0.216150 | 0.216726 | 0.215126 | 0.216058 | 0.217904 | 0.220599 | 0.223520 | 0.226502 | 0.240940 | 0.244591 | 0.250321 | 0.246950 | 0.266030 | 0.309308 | 0.385935 | 0.396806 | 0.364883 | 0.304907 | 0.268703 | 0.254452 | 0.229460 | 0.221163 | 0.228840 | 0.220435 | 0.214975 | 0.229551 | 0.282347 | 0.343962 | 0.353847 | 0.378950 | 0.267523 | 0.218699 | 0.230541 | 0.312534 | 0.503783 | 0.640150 | 0.424335 | 0.260106 | 0.224212 | 0.215708 | 0.216845 | 0.227684 | 0.251531 | 0.295469 | 0.302753 | 0.309568 | 0.353883 | 0.322519 | 0.308095 | 0.240599 | 0.214702 | 0.216550 | 0.217373 | 0.214947 | 0.219443 | 0.217640 | 0.216908 | 0.215436 | 0.216711 | 0.235708 | 0.242764 | 0.253634 | 0.228359 | 0.217400 | 0.237584 | 0.246176 | 0.246998 | 0.236429 | 0.244145 | 0.245065 | 0.239793 | 0.261786 | 0.282078 | 0.286835 | 0.246602 | 0.243612 | 0.231427 | 0.227471 | 0.218476 | 0.229107 | 0.227823 | 0.223493 | 0.217348 | 0.221072 | 0.227561 | 0.242224 | 0.239912 | 0.260960 | 0.357495 | 0.408992 | 0.394531 | 0.430460 | 0.473183 | 0.517538 | 0.553056 | 0.402308 | 0.398613 | 0.571396 | 0.634208 | 0.730895 | 0.839123 | 0.925364 | 1.193817 | 1.451740 | 1.222125 | 0.990714 | 0.691171 | 0.584670 | 0.561672 | 0.426261 | 0.394999 | 0.485702 | 0.647861 | 1.075765 | 1.804906 | 1.822427 | 2.926186 | 2.422088 | 1.115916 | 0.845241 | 0.587568 | 0.364819 | 0.311999 | 0.289608 | 0.295573 | 0.343484 | 0.338643 | 0.298232 | 0.271598 | 0.251442 | 0.238043 | 0.218863 | 0.219474 | 0.225553 | 0.255981 | 0.311289 | 0.343013 | 0.398344 | 0.514691 | 0.529483 | 0.615282 | 0.615175 | 0.611540 | 0.914718 | 0.902157 | 0.863275 | 0.771678 | 0.406139 | 0.266407 | 0.236105 | 0.220313 | 0.229151 | 0.240080 | 0.224959 | 0.223605 | 0.238981 | 0.237076 | 0.252280 | 0.244860 | 0.230495 | 0.221082 | 0.214890 | 0.214637 | 0.215820 | 0.220076 | 0.226117 | 0.230128 | 0.221301 | 0.214801 |
| right anterior | 0.399753 | 0.411396 | 0.316837 | 0.270020 | 0.230118 | 0.216229 | 0.216296 | 0.229252 | 0.235997 | 0.222619 | 0.215503 | 0.239663 | 0.346997 | 0.463872 | 0.435503 | 0.314432 | 0.297071 | 0.375968 | 0.331594 | 0.278687 | 0.268945 | 0.278430 | 0.478310 | 0.675276 | 0.390412 | 0.337794 | 0.261388 | 0.234295 | 0.216634 | 0.214663 | 0.216204 | 0.214742 | 0.217008 | 0.214685 | 0.217712 | 0.245434 | 0.229167 | 0.217471 | 0.215187 | 0.216459 | 0.214767 | 0.220364 | 0.215969 | 0.218777 | 0.217313 | 0.227386 | 0.277617 | 0.423554 | 0.697698 | 0.498807 | 0.333743 | 0.413182 | 0.330608 | 0.294043 | 0.267618 | 0.256711 | 0.270030 | 0.334120 | 0.272153 | 0.282992 | 0.248733 | 0.226928 | 0.219506 | 0.219874 | 0.215362 | 0.217119 | 0.216342 | 0.216154 | 0.233678 | 0.288521 | 0.289289 | 0.346819 | 0.519686 | 0.786484 | 1.090682 | 0.877816 | 0.499364 | 0.483570 | 0.522877 | 0.351163 | 0.328039 | 0.310017 | 0.305748 | 0.312305 | 0.374734 | 0.336142 | 0.338521 | 0.262152 | 0.220423 | 0.219336 | 0.271671 | 0.331824 | 0.368232 | 0.339059 | 0.296856 | 0.246826 | 0.224019 | 0.215606 | 0.231821 | 0.251614 | 0.243647 | 0.233924 | 0.225511 | 0.215366 | 0.217016 | 0.225232 | 0.230150 | 0.220199 | 0.215834 | 0.217488 | 0.246168 | 0.304297 | 0.426556 | 0.569080 | 0.720478 | 0.823759 | 0.942558 | 0.918537 | 0.886842 | 0.834446 | 0.723801 | 0.619502 | 0.429359 | 0.276883 | 0.256719 | 0.262839 | 0.259748 | 0.304247 | 0.310220 | 0.347243 | 0.470158 | 0.436931 | 0.399639 | 0.421563 | 0.450872 | 0.564695 | 0.819411 | 0.915922 | 0.935250 | 0.940311 | 0.619633 | 0.471407 | 0.393913 | 0.304579 | 0.264237 | 0.259583 | 0.270312 | 0.292560 | 0.322296 | 0.318150 | 0.341950 | 0.361232 | 0.328145 | 0.292190 | 0.305250 | 0.267759 | 0.250008 | 0.224959 | 0.218102 | 0.218249 | 0.216484 | 0.216202 | 0.220978 | 0.215009 | 0.214762 | 0.215058 | 0.214659 | 0.216710 | 0.221581 | 0.226989 | 0.214854 | 0.214662 | 0.215660 | 0.216353 | 0.228720 | 0.226471 | 0.219673 | 0.218283 | 0.214863 | 0.214632 | 0.214632 | 0.214780 | 0.214632 | 0.215559 | 0.217934 | 0.215465 | 0.215072 | 0.214718 | 0.214984 | 0.214719 | 0.217915 | 0.220630 | 0.226445 | 0.224772 | 0.222240 | 0.229300 | 0.226468 | 0.218643 | 0.221242 | 0.217107 | 0.216535 | 0.219693 | 0.220113 | 0.223263 | 0.223773 | 0.217975 | 0.215139 | 0.214652 | 0.215411 | 0.222166 | 0.224562 | 0.215064 | 0.216120 | 0.220623 | 0.219173 | 0.243947 | 0.327743 | 0.387992 | 0.378409 | 0.458287 | 0.656377 | 0.756277 | 0.568992 | 0.373476 | 0.320317 | 0.297510 | 0.263092 | 0.238406 | 0.232073 | 0.228499 | 0.226362 | 0.216562 | 0.217454 | 0.217169 | 0.215777 | 0.214929 | 0.214720 | 0.214650 | 0.214728 | 0.215268 | 0.218633 | 0.235520 | 0.237527 | 0.243142 | 0.258204 | 0.233615 | 0.222040 | 0.221197 | 0.215703 | 0.215076 | 0.215194 | 0.216016 | 0.225578 | 0.250069 | 0.248362 | 0.241917 | 0.224760 | 0.221816 | 0.214635 | 0.221565 | 0.247776 | 0.264798 | 0.257907 | 0.220307 | 0.218168 | 0.216876 | 0.215420 | 0.214648 | 0.214890 | 0.214676 | 0.217821 | 0.232713 | 0.237640 | 0.249705 | 0.257574 | 0.251751 | 0.229436 | 0.224514 | 0.216843 | 0.214979 |
| left central | 0.367436 | 0.456967 | 0.420685 | 0.388364 | 0.338570 | 0.255902 | 0.214689 | 0.256027 | 0.269698 | 0.344659 | 0.436417 | 0.418724 | 0.490118 | 0.385239 | 0.265229 | 0.268319 | 0.216927 | 0.223525 | 0.223869 | 0.253299 | 0.281146 | 0.360335 | 0.725121 | 0.707164 | 0.288945 | 0.259472 | 0.219226 | 0.224642 | 0.214899 | 0.240403 | 0.339377 | 0.300447 | 0.365415 | 0.405616 | 1.129179 | 1.719828 | 5.054130 | 14.343769 | 16.990950 | 2.694050 | 1.057786 | 0.310100 | 0.229810 | 0.214981 | 0.251368 | 0.244851 | 0.214888 | 0.216050 | 0.218877 | 0.225534 | 0.235134 | 0.218934 | 0.226424 | 0.257258 | 0.264114 | 0.230231 | 0.214661 | 0.218875 | 0.220035 | 0.234760 | 0.241623 | 0.248932 | 0.258170 | 0.255988 | 0.254253 | 0.286485 | 0.411950 | 0.744299 | 2.025877 | 7.422149 | 38.973721 | 67.333532 | 43.177350 | 22.775075 | 11.684416 | 4.375259 | 1.340183 | 0.492755 | 0.300646 | 0.238369 | 0.217913 | 0.214794 | 0.219255 | 0.227949 | 0.236267 | 0.240728 | 0.233938 | 0.224137 | 0.215352 | 0.214970 | 0.215180 | 0.219423 | 0.219579 | 0.216763 | 0.214831 | 0.222537 | 0.220942 | 0.215485 | 0.219205 | 0.221461 | 0.221202 | 0.217892 | 0.219272 | 0.215544 | 0.215133 | 0.214813 | 0.217876 | 0.217694 | 0.220482 | 0.216515 | 0.215577 | 0.214747 | 0.214626 | 0.216694 | 0.215144 | 0.218671 | 0.217030 | 0.218496 | 0.220792 | 0.215072 | 0.216819 | 0.233190 | 0.272930 | 0.331451 | 0.450794 | 0.669139 | 0.805277 | 1.335465 | 2.101455 | 2.526134 | 2.108816 | 1.072389 | 0.605163 | 0.450536 | 0.328926 | 0.260212 | 0.235112 | 0.220802 | 0.224753 | 0.239160 | 0.243907 | 0.250224 | 0.253075 | 0.243878 | 0.248981 | 0.247289 | 0.240333 | 0.215949 | 0.214626 | 0.216004 | 0.224927 | 0.234140 | 0.224738 | 0.224338 | 0.217699 | 0.218309 | 0.222558 | 0.215082 | 0.215376 | 0.216432 | 0.218830 | 0.226653 | 0.243817 | 0.292294 | 0.342485 | 0.398175 | 0.403108 | 0.391276 | 0.404040 | 0.391369 | 0.358376 | 0.406099 | 0.467539 | 0.590580 | 0.759297 | 0.986256 | 0.823257 | 0.874269 | 0.609577 | 0.488338 | 0.379809 | 0.301984 | 0.272523 | 0.292299 | 0.284025 | 0.287373 | 0.265196 | 0.279246 | 0.278834 | 0.290154 | 0.288034 | 0.279549 | 0.265480 | 0.278909 | 0.272551 | 0.285190 | 0.262390 | 0.251696 | 0.254756 | 0.272216 | 0.271343 | 0.278781 | 0.253579 | 0.238262 | 0.239158 | 0.235635 | 0.236683 | 0.237002 | 0.242554 | 0.272545 | 0.317817 | 0.305020 | 0.277502 | 0.259287 | 0.272545 | 0.297939 | 0.309248 | 0.371042 | 0.542372 | 0.838166 | 1.189241 | 1.253554 | 1.211941 | 1.214958 | 0.829276 | 0.609460 | 0.449552 | 0.352095 | 0.301348 | 0.259460 | 0.242491 | 0.237604 | 0.242533 | 0.237426 | 0.232196 | 0.231680 | 0.247978 | 0.279678 | 0.329131 | 0.363426 | 0.438763 | 0.554003 | 0.587055 | 0.526061 | 0.358435 | 0.313047 | 0.317216 | 0.306238 | 0.299571 | 0.300757 | 0.293035 | 0.346233 | 0.354439 | 0.317443 | 0.355569 | 0.381378 | 0.361609 | 0.341745 | 0.324312 | 0.326804 | 0.399604 | 0.350382 | 0.327321 | 0.350921 | 0.379194 | 0.404805 | 0.435933 | 0.388756 | 0.440238 | 0.505831 | 0.394853 | 0.314331 | 0.275208 | 0.248423 | 0.245285 | 0.230968 | 0.222283 | 0.219086 | 0.224436 | 0.231018 |
| right central | 0.214756 | 0.220420 | 0.232833 | 0.235238 | 0.267560 | 0.351124 | 0.291693 | 0.249207 | 0.258281 | 0.269156 | 0.240670 | 0.219750 | 0.215176 | 0.224002 | 0.254354 | 0.227791 | 0.226480 | 0.323213 | 0.481460 | 0.580047 | 0.399272 | 0.252657 | 0.247686 | 0.221017 | 0.229794 | 0.276900 | 0.333207 | 0.521497 | 0.493100 | 0.704614 | 1.040661 | 0.347120 | 0.272089 | 0.264650 | 0.214705 | 0.221069 | 0.216161 | 0.216337 | 0.226800 | 0.306323 | 0.378624 | 0.429471 | 0.356199 | 0.285199 | 0.289616 | 0.268480 | 0.220113 | 0.214831 | 0.233150 | 0.264222 | 0.237071 | 0.251917 | 0.325782 | 0.417070 | 0.420397 | 0.283986 | 0.238130 | 0.244331 | 0.241924 | 0.260646 | 0.250763 | 0.268311 | 0.370394 | 0.520850 | 0.642624 | 0.779275 | 0.947658 | 2.659144 | 9.846860 | 40.808661 | 333.294158 | 1421.721580 | 1254.632954 | 1013.908979 | 460.580058 | 277.344780 | 111.997735 | 24.353825 | 9.863876 | 20.189318 | 18.927610 | 5.503960 | 1.770353 | 1.023065 | 0.950916 | 0.732123 | 0.357965 | 0.271416 | 0.279733 | 0.279492 | 0.264755 | 0.232589 | 0.218656 | 0.229578 | 0.265776 | 0.281962 | 0.277569 | 0.308447 | 0.484346 | 1.031253 | 1.629123 | 1.875479 | 2.488214 | 4.644572 | 3.715497 | 2.881753 | 2.020226 | 1.371520 | 1.230365 | 0.915114 | 0.644650 | 0.673092 | 0.547496 | 0.458133 | 0.495389 | 0.570318 | 0.702807 | 0.698851 | 0.674853 | 0.762768 | 1.456746 | 2.166129 | 2.240996 | 2.119407 | 2.953821 | 2.707527 | 2.406572 | 2.323655 | 2.436279 | 2.980460 | 2.274571 | 1.983037 | 2.152363 | 2.388303 | 1.594688 | 1.263035 | 0.795292 | 0.796080 | 0.569963 | 0.495370 | 0.538716 | 0.686380 | 0.884339 | 1.131694 | 1.045154 | 1.276111 | 1.443733 | 1.150533 | 0.808667 | 0.457609 | 0.325883 | 0.332464 | 0.339956 | 0.332485 | 0.321090 | 0.351369 | 0.652905 | 1.740288 | 3.897716 | 7.483443 | 7.756595 | 14.294912 | 22.016596 | 12.461373 | 6.054615 | 2.910364 | 1.386499 | 0.782803 | 0.505222 | 0.420416 | 0.452976 | 0.531191 | 0.718277 | 0.828341 | 1.393823 | 1.913997 | 2.187376 | 1.859621 | 1.415150 | 1.102814 | 1.006473 | 0.694284 | 0.474826 | 0.320167 | 0.235888 | 0.232002 | 0.233089 | 0.230184 | 0.226304 | 0.222035 | 0.215493 | 0.215948 | 0.214751 | 0.217018 | 0.220546 | 0.240401 | 0.259846 | 0.241878 | 0.234393 | 0.221646 | 0.222826 | 0.235077 | 0.226613 | 0.222441 | 0.226948 | 0.235020 | 0.269278 | 0.303173 | 0.317996 | 0.330069 | 0.336909 | 0.325737 | 0.300875 | 0.315353 | 0.345944 | 0.416332 | 0.419150 | 0.379826 | 0.323458 | 0.319071 | 0.301947 | 0.309227 | 0.270629 | 0.265064 | 0.291962 | 0.389135 | 0.454397 | 0.498470 | 0.395974 | 0.356312 | 0.294258 | 0.235408 | 0.216525 | 0.246752 | 0.322008 | 0.304579 | 0.281656 | 0.277201 | 0.270517 | 0.246971 | 0.227583 | 0.223282 | 0.229592 | 0.234738 | 0.222396 | 0.214626 | 0.216597 | 0.216497 | 0.215008 | 0.215477 | 0.218080 | 0.225445 | 0.233648 | 0.247756 | 0.265291 | 0.270030 | 0.258713 | 0.265842 | 0.267368 | 0.316735 | 0.325873 | 0.289706 | 0.245570 | 0.224234 | 0.219910 | 0.219122 | 0.222498 | 0.228320 | 0.238752 | 0.255490 | 0.287086 | 0.289773 | 0.314604 | 0.313437 | 0.311050 | 0.306322 | 0.332411 | 0.359265 | 0.385212 | 0.380135 |
| left posterior | 0.327553 | 0.404425 | 0.353536 | 0.256347 | 0.263668 | 0.255682 | 0.221243 | 0.214649 | 0.216869 | 0.217567 | 0.291173 | 0.403506 | 0.433025 | 0.491329 | 0.311232 | 0.248867 | 0.218874 | 0.215256 | 0.237273 | 0.353442 | 0.538107 | 0.330913 | 0.350112 | 0.402908 | 0.610453 | 0.716010 | 0.677576 | 0.508222 | 0.873824 | 1.176015 | 1.124390 | 0.721745 | 0.520174 | 0.559066 | 0.691386 | 0.437073 | 0.270863 | 0.233885 | 0.219860 | 0.218350 | 0.214636 | 0.214636 | 0.214637 | 0.215794 | 0.219681 | 0.214626 | 0.214734 | 0.223292 | 0.268151 | 0.329979 | 0.339385 | 0.474933 | 0.709741 | 1.714802 | 1.938093 | 1.469234 | 1.447139 | 2.252178 | 2.506836 | 4.989153 | 5.603730 | 6.552801 | 6.057007 | 4.674321 | 3.286733 | 1.552785 | 0.539232 | 0.237888 | 0.248753 | 0.560741 | 2.341874 | 22.190351 | 108.633447 | 198.031322 | 178.368046 | 221.973704 | 188.588973 | 137.025849 | 60.442037 | 16.939734 | 6.834711 | 3.032352 | 1.437447 | 0.653278 | 0.402905 | 0.256093 | 0.224917 | 0.214626 | 0.216175 | 0.230611 | 0.276711 | 0.382818 | 0.697492 | 0.523490 | 0.499781 | 0.421728 | 0.334777 | 0.275651 | 0.250740 | 0.250175 | 0.274600 | 0.284392 | 0.268935 | 0.240763 | 0.245694 | 0.248304 | 0.244428 | 0.234396 | 0.222426 | 0.219526 | 0.218949 | 0.221130 | 0.221364 | 0.219916 | 0.224421 | 0.230539 | 0.232313 | 0.234614 | 0.235550 | 0.250802 | 0.307123 | 0.383851 | 0.555797 | 0.787030 | 1.332809 | 2.080772 | 2.602615 | 2.744471 | 2.338053 | 1.506346 | 1.160194 | 0.978841 | 0.686217 | 0.545145 | 0.436774 | 0.415862 | 0.391650 | 0.370323 | 0.343067 | 0.352205 | 0.381520 | 0.394559 | 0.423846 | 0.454036 | 0.494276 | 0.523481 | 0.535066 | 0.442677 | 0.433114 | 0.374446 | 0.301033 | 0.257917 | 0.240976 | 0.236237 | 0.249612 | 0.257483 | 0.276982 | 0.384517 | 0.651818 | 1.307751 | 2.774523 | 3.202349 | 2.823651 | 1.889178 | 1.269241 | 1.016511 | 0.831884 | 0.892208 | 1.016343 | 1.339655 | 1.734328 | 2.132034 | 1.789820 | 1.540301 | 1.188744 | 1.307465 | 1.500438 | 1.425731 | 1.251284 | 1.005632 | 1.002074 | 0.969081 | 0.842986 | 0.681017 | 0.632127 | 0.550328 | 0.653654 | 0.695158 | 0.636916 | 0.555220 | 0.511392 | 0.458767 | 0.532822 | 0.565312 | 0.605005 | 0.607108 | 0.669825 | 0.651119 | 0.720225 | 0.926084 | 1.059544 | 0.965021 | 1.024450 | 0.889922 | 1.083935 | 1.447886 | 1.222143 | 0.887544 | 0.818129 | 0.797909 | 0.952720 | 0.655130 | 0.470081 | 0.415084 | 0.457448 | 0.489229 | 0.453353 | 0.395894 | 0.416844 | 0.410541 | 0.382416 | 0.327556 | 0.266720 | 0.256105 | 0.262111 | 0.257513 | 0.269871 | 0.310138 | 0.370732 | 0.582623 | 0.699493 | 0.853085 | 1.077878 | 1.296442 | 1.148586 | 1.108819 | 1.265721 | 1.699765 | 2.094080 | 2.827140 | 3.545748 | 6.122743 | 8.747702 | 9.027071 | 7.465461 | 5.230539 | 5.209870 | 4.035213 | 2.313084 | 1.750004 | 0.852594 | 0.548421 | 0.401699 | 0.304017 | 0.250154 | 0.240273 | 0.241942 | 0.293809 | 0.348617 | 0.400293 | 0.375485 | 0.350828 | 0.295766 | 0.268570 | 0.262672 | 0.301783 | 0.331844 | 0.405055 | 0.496445 | 0.702595 | 0.909200 | 0.896313 | 0.665028 | 0.675260 | 0.692794 | 0.877253 | 1.092715 | 0.976175 | 1.088007 | 1.635235 |
| right posterior | 0.215203 | 0.217931 | 0.215177 | 0.214729 | 0.215777 | 0.215543 | 0.216521 | 0.222949 | 0.238931 | 0.227624 | 0.226265 | 0.215949 | 0.214948 | 0.214965 | 0.215876 | 0.216491 | 0.228799 | 0.239835 | 0.320944 | 0.421954 | 0.643150 | 0.620029 | 0.394318 | 0.285001 | 0.222756 | 0.218145 | 0.258487 | 0.373934 | 0.445042 | 0.376850 | 0.412655 | 0.326525 | 0.299082 | 0.262881 | 0.218040 | 0.217122 | 0.227449 | 0.251028 | 0.310805 | 0.361581 | 0.604771 | 0.347775 | 0.265493 | 0.238576 | 0.233420 | 0.226902 | 0.233568 | 0.231401 | 0.244497 | 0.222747 | 0.218224 | 0.216473 | 0.215211 | 0.217296 | 0.216730 | 0.221198 | 0.224411 | 0.229663 | 0.220992 | 0.217277 | 0.216820 | 0.244080 | 0.337817 | 0.372235 | 0.475108 | 0.374067 | 0.283751 | 0.215953 | 0.347130 | 1.971884 | 19.645997 | 173.465677 | 920.307557 | 1332.256832 | 1746.770792 | 1318.271129 | 1254.071455 | 512.570873 | 163.125270 | 17.529411 | 2.196060 | 0.441675 | 0.249738 | 0.214787 | 0.214726 | 0.214633 | 0.221341 | 0.229520 | 0.243929 | 0.231299 | 0.215532 | 0.215910 | 0.215619 | 0.224745 | 0.218945 | 0.214861 | 0.216313 | 0.220592 | 0.226168 | 0.216807 | 0.215342 | 0.217877 | 0.234393 | 0.259919 | 0.327340 | 0.434129 | 0.572654 | 0.779042 | 1.364439 | 2.068612 | 1.960426 | 1.354580 | 1.121578 | 0.693113 | 0.549428 | 0.499358 | 0.526343 | 0.755523 | 1.298150 | 2.023594 | 3.520284 | 6.481935 | 6.312432 | 5.117943 | 4.816314 | 3.823174 | 3.638090 | 3.470485 | 2.589077 | 2.291166 | 1.889307 | 1.610443 | 1.436793 | 1.043299 | 0.976428 | 1.122369 | 1.985802 | 3.868626 | 7.420848 | 14.159112 | 17.798712 | 24.546358 | 23.381276 | 13.063227 | 6.972981 | 2.479277 | 1.086472 | 0.771130 | 0.460388 | 0.376168 | 0.362803 | 0.327399 | 0.338056 | 0.381924 | 0.385660 | 0.503612 | 0.766165 | 1.034550 | 1.638438 | 2.869145 | 5.662094 | 9.503882 | 10.959228 | 12.326044 | 11.837012 | 13.037300 | 15.716605 | 20.629338 | 39.189617 | 81.995405 | 114.585538 | 176.396992 | 302.361305 | 245.385492 | 131.273713 | 72.191512 | 36.255216 | 23.332168 | 20.221285 | 10.250292 | 10.046103 | 8.887462 | 5.898100 | 4.070833 | 3.393896 | 3.089528 | 4.853041 | 7.857086 | 10.268698 | 11.777720 | 17.039878 | 14.984047 | 11.191888 | 8.653824 | 4.657736 | 2.362384 | 1.362026 | 0.831242 | 0.829900 | 0.824748 | 0.746971 | 0.804900 | 0.861341 | 0.786485 | 0.716866 | 0.490339 | 0.377776 | 0.327158 | 0.293356 | 0.262252 | 0.251383 | 0.250238 | 0.312614 | 0.380183 | 0.479272 | 0.773166 | 1.444719 | 2.170900 | 2.390664 | 1.736182 | 1.351222 | 0.912453 | 0.572678 | 0.372671 | 0.294268 | 0.254371 | 0.232206 | 0.235062 | 0.247448 | 0.258151 | 0.326281 | 0.430809 | 0.619776 | 0.973909 | 1.205477 | 1.406544 | 1.618551 | 1.290707 | 1.242871 | 1.160421 | 0.879374 | 0.624585 | 0.560483 | 0.525739 | 0.686047 | 0.709938 | 0.738531 | 0.714288 | 0.710753 | 0.622970 | 0.454719 | 0.373444 | 0.319125 | 0.271939 | 0.249085 | 0.222300 | 0.214909 | 0.215710 | 0.226399 | 0.228840 | 0.218680 | 0.214921 | 0.235944 | 0.267418 | 0.316402 | 0.404455 | 0.643555 | 1.189385 | 1.484364 | 1.510889 | 1.458971 | 1.595381 | 1.601195 | 1.607746 | 1.451874 | 2.018504 | 2.712480 | 2.818218 | 2.921866 | 3.245570 |
| all electrodes | 0.234664 | 0.220635 | 0.220050 | 0.217410 | 0.214762 | 0.264782 | 0.275580 | 0.255742 | 0.226575 | 0.222020 | 0.214626 | 0.218676 | 0.246847 | 0.216002 | 0.219790 | 0.215314 | 0.239783 | 0.382690 | 0.377592 | 0.314414 | 0.309430 | 0.240050 | 0.312007 | 0.261120 | 0.282122 | 0.514130 | 0.503596 | 0.294840 | 0.331202 | 0.317140 | 0.374821 | 0.229822 | 0.245486 | 0.216706 | 0.224639 | 0.239840 | 0.230670 | 0.215067 | 0.217280 | 0.219071 | 0.214637 | 0.221836 | 0.225444 | 0.218793 | 0.215941 | 0.218887 | 0.216282 | 0.218462 | 0.222248 | 0.245385 | 0.290101 | 0.295862 | 0.261848 | 0.334932 | 0.348377 | 0.249761 | 0.234813 | 0.219801 | 0.217193 | 0.224518 | 0.216830 | 0.219928 | 0.261124 | 0.254443 | 0.239202 | 0.233107 | 0.217487 | 0.222763 | 0.295873 | 0.559263 | 1.729382 | 7.920755 | 72.722795 | 244.110513 | 592.140087 | 759.196367 | 389.008093 | 149.970119 | 252.935015 | 121.195371 | 191.773902 | 82.199670 | 29.878329 | 15.628755 | 5.196392 | 1.198569 | 0.512343 | 0.278312 | 0.238906 | 0.227239 | 0.218981 | 0.214680 | 0.216844 | 0.215452 | 0.214877 | 0.214626 | 0.218327 | 0.221345 | 0.214638 | 0.226494 | 0.250227 | 0.276270 | 0.257388 | 0.276437 | 0.330912 | 0.290729 | 0.262507 | 0.234418 | 0.240688 | 0.281194 | 0.283686 | 0.268394 | 0.285259 | 0.304733 | 0.323550 | 0.325499 | 0.340990 | 0.418256 | 0.584253 | 0.927187 | 1.292730 | 1.362974 | 1.303124 | 1.410239 | 1.831776 | 3.120907 | 5.340172 | 7.202327 | 7.825978 | 9.268819 | 11.898399 | 7.828402 | 3.570925 | 1.399165 | 0.947637 | 0.880609 | 0.667673 | 0.558814 | 0.475571 | 0.469820 | 0.585181 | 0.771442 | 0.624397 | 0.523861 | 0.446425 | 0.459147 | 0.420855 | 0.321015 | 0.238919 | 0.234115 | 0.235391 | 0.218067 | 0.216103 | 0.222975 | 0.223238 | 0.214803 | 0.233435 | 0.288912 | 0.443362 | 0.948993 | 2.666887 | 7.809414 | 6.784340 | 4.574021 | 3.758781 | 5.258311 | 3.071803 | 1.079373 | 0.589481 | 0.380627 | 0.294986 | 0.292456 | 0.254431 | 0.259545 | 0.286004 | 0.330960 | 0.383822 | 0.379481 | 0.301053 | 0.328371 | 0.383079 | 0.449534 | 0.457893 | 0.409561 | 0.402706 | 0.423809 | 0.395888 | 0.416994 | 0.404971 | 0.375851 | 0.385999 | 0.404513 | 0.425561 | 0.404987 | 0.290225 | 0.247638 | 0.246965 | 0.261476 | 0.270200 | 0.267445 | 0.270757 | 0.291130 | 0.314003 | 0.311980 | 0.309221 | 0.306072 | 0.301013 | 0.288554 | 0.274095 | 0.265072 | 0.264808 | 0.285289 | 0.307438 | 0.361194 | 0.414507 | 0.555233 | 0.760955 | 0.755417 | 0.688553 | 0.639630 | 0.632220 | 0.670401 | 0.507191 | 0.334661 | 0.260726 | 0.224321 | 0.216700 | 0.218061 | 0.216877 | 0.219885 | 0.242176 | 0.279975 | 0.355014 | 0.418647 | 0.394321 | 0.417374 | 0.396232 | 0.357125 | 0.340583 | 0.288843 | 0.279258 | 0.272639 | 0.270745 | 0.283753 | 0.285265 | 0.290777 | 0.324787 | 0.369280 | 0.368245 | 0.386974 | 0.434373 | 0.457879 | 0.448045 | 0.452684 | 0.382538 | 0.417987 | 0.359515 | 0.314880 | 0.304622 | 0.354970 | 0.407858 | 0.374106 | 0.383892 | 0.445255 | 0.485378 | 0.543247 | 0.481116 | 0.587730 | 0.728161 | 0.692822 | 0.640715 | 0.704941 | 0.836424 | 0.998619 | 1.033078 | 1.379546 | 1.411615 | 1.711816 | 1.846786 | 1.523222 |

Searchlight, spatiotemporal cluster permutation test

|  | start time | stop time | peak time | peak channel | cluster p | peak Cohen's d | direction |
| --- | --- | --- | --- | --- | --- | --- | --- |
| #1 | 130 | 230 | 170 | O2 | 0.0453 | 1.298038 | positive |

X) emoji to real faces - happy vs sad

  
|  | time window | peak latency | cluster *p* | peak Cohen's *d* |  | | | |
| **all electrodes** |  | | | |  | | | |
|  | | | | | | | | |

Time-resolved classification, cluster permutation tests

|  | **left hemisphere** | | | | **right hemisphere** | | | |
|  | time window | peak latency | cluster *p* | peak Cohen's *d* | time window | peak latency | cluster *p* | peak Cohen's *d* |
| **anterior** |  | | | |  | | | |
| **central** |  | | | |  | | | |
| **posterior** |  | | | |  | | | |

  

Time-resolved classification, Bayesian statistics

|  | -200 | -195 | -190 | -185 | -180 | -175 | -170 | -165 | -160 | -155 | -150 | -145 | -140 | -135 | -130 | -125 | -120 | -115 | -110 | -105 | -100 | -95 | -90 | -85 | -80 | -75 | -70 | -65 | -60 | -55 | -50 | -45 | -40 | -35 | -30 | -25 | -20 | -15 | -10 | -5 | 0 | 5 | 10 | 15 | 20 | 25 | 30 | 35 | 40 | 45 | 50 | 55 | 60 | 65 | 70 | 75 | 80 | 85 | 90 | 95 | 100 | 105 | 110 | 115 | 120 | 125 | 130 | 135 | 140 | 145 | 150 | 155 | 160 | 165 | 170 | 175 | 180 | 185 | 190 | 195 | 200 | 205 | 210 | 215 | 220 | 225 | 230 | 235 | 240 | 245 | 250 | 255 | 260 | 265 | 270 | 275 | 280 | 285 | 290 | 295 | 300 | 305 | 310 | 315 | 320 | 325 | 330 | 335 | 340 | 345 | 350 | 355 | 360 | 365 | 370 | 375 | 380 | 385 | 390 | 395 | 400 | 405 | 410 | 415 | 420 | 425 | 430 | 435 | 440 | 445 | 450 | 455 | 460 | 465 | 470 | 475 | 480 | 485 | 490 | 495 | 500 | 505 | 510 | 515 | 520 | 525 | 530 | 535 | 540 | 545 | 550 | 555 | 560 | 565 | 570 | 575 | 580 | 585 | 590 | 595 | 600 | 605 | 610 | 615 | 620 | 625 | 630 | 635 | 640 | 645 | 650 | 655 | 660 | 665 | 670 | 675 | 680 | 685 | 690 | 695 | 700 | 705 | 710 | 715 | 720 | 725 | 730 | 735 | 740 | 745 | 750 | 755 | 760 | 765 | 770 | 775 | 780 | 785 | 790 | 795 | 800 | 805 | 810 | 815 | 820 | 825 | 830 | 835 | 840 | 845 | 850 | 855 | 860 | 865 | 870 | 875 | 880 | 885 | 890 | 895 | 900 | 905 | 910 | 915 | 920 | 925 | 930 | 935 | 940 | 945 | 950 | 955 | 960 | 965 | 970 | 975 | 980 | 985 | 990 | 995 | 1000 | 1005 | 1010 | 1015 | 1020 | 1025 | 1030 | 1035 | 1040 | 1045 | 1050 | 1055 | 1060 | 1065 | 1070 | 1075 | 1080 | 1085 | 1090 | 1095 | 1100 | 1105 | 1110 | 1115 | 1120 | 1125 | 1130 | 1135 | 1140 | 1145 | 1150 | 1155 | 1160 | 1165 | 1170 | 1175 | 1180 | 1185 | 1190 | 1195 |
| --- | --- | --- | --- | --- | --- | --- | --- | --- | --- | --- | --- | --- | --- | --- | --- | --- | --- | --- | --- | --- | --- | --- | --- | --- | --- | --- | --- | --- | --- | --- | --- | --- | --- | --- | --- | --- | --- | --- | --- | --- | --- | --- | --- | --- | --- | --- | --- | --- | --- | --- | --- | --- | --- | --- | --- | --- | --- | --- | --- | --- | --- | --- | --- | --- | --- | --- | --- | --- | --- | --- | --- | --- | --- | --- | --- | --- | --- | --- | --- | --- | --- | --- | --- | --- | --- | --- | --- | --- | --- | --- | --- | --- | --- | --- | --- | --- | --- | --- | --- | --- | --- | --- | --- | --- | --- | --- | --- | --- | --- | --- | --- | --- | --- | --- | --- | --- | --- | --- | --- | --- | --- | --- | --- | --- | --- | --- | --- | --- | --- | --- | --- | --- | --- | --- | --- | --- | --- | --- | --- | --- | --- | --- | --- | --- | --- | --- | --- | --- | --- | --- | --- | --- | --- | --- | --- | --- | --- | --- | --- | --- | --- | --- | --- | --- | --- | --- | --- | --- | --- | --- | --- | --- | --- | --- | --- | --- | --- | --- | --- | --- | --- | --- | --- | --- | --- | --- | --- | --- | --- | --- | --- | --- | --- | --- | --- | --- | --- | --- | --- | --- | --- | --- | --- | --- | --- | --- | --- | --- | --- | --- | --- | --- | --- | --- | --- | --- | --- | --- | --- | --- | --- | --- | --- | --- | --- | --- | --- | --- | --- | --- | --- | --- | --- | --- | --- | --- | --- | --- | --- | --- | --- | --- | --- | --- | --- | --- | --- | --- | --- | --- | --- | --- | --- | --- | --- | --- | --- | --- | --- | --- | --- | --- | --- | --- | --- | --- | --- | --- | --- | --- | --- | --- | --- | --- | --- | --- | --- | --- | --- | --- |
| left anterior | 0.462079 | 0.432946 | 0.359158 | 0.322445 | 0.558075 | 0.501464 | 0.265466 | 0.214703 | 0.228157 | 0.234059 | 0.257528 | 0.472094 | 0.399868 | 0.305303 | 0.238391 | 0.217049 | 0.220465 | 0.245476 | 0.264446 | 0.229604 | 0.238215 | 0.216133 | 0.214704 | 0.222993 | 0.333857 | 0.341757 | 0.275011 | 0.244404 | 0.225486 | 0.215639 | 0.218115 | 0.354758 | 0.553386 | 0.417540 | 0.320025 | 0.217849 | 0.218675 | 0.216828 | 0.227710 | 0.221196 | 0.258068 | 0.281865 | 0.387401 | 0.276987 | 0.269422 | 0.251323 | 0.319214 | 0.307556 | 0.287282 | 0.320986 | 0.381282 | 0.303306 | 0.268907 | 0.266421 | 0.271126 | 0.244414 | 0.214880 | 0.240450 | 0.228900 | 0.227579 | 0.278296 | 0.355510 | 0.295053 | 0.254165 | 0.228735 | 0.236342 | 0.268361 | 0.256722 | 0.293006 | 0.305824 | 0.281365 | 0.275527 | 0.303649 | 0.374772 | 0.874393 | 0.939874 | 1.467823 | 1.940206 | 1.368701 | 0.882319 | 0.525512 | 0.345831 | 0.287386 | 0.284256 | 0.265506 | 0.248073 | 0.258050 | 0.269069 | 0.274644 | 0.244880 | 0.222317 | 0.219574 | 0.251845 | 0.259599 | 0.249974 | 0.226295 | 0.221111 | 0.216321 | 0.228404 | 0.377793 | 0.470147 | 0.533504 | 0.472274 | 0.500531 | 0.505333 | 0.508290 | 0.428654 | 0.475859 | 0.499306 | 0.630674 | 0.816198 | 1.122212 | 1.005950 | 0.791554 | 0.462179 | 0.295796 | 0.231652 | 0.216815 | 0.215185 | 0.215354 | 0.214672 | 0.214637 | 0.222089 | 0.260814 | 0.284073 | 0.270367 | 0.260492 | 0.321098 | 0.405585 | 0.451188 | 0.558310 | 0.643497 | 0.743727 | 1.052621 | 1.216937 | 1.514002 | 2.198830 | 3.104839 | 6.551898 | 8.522189 | 11.241577 | 14.960757 | 18.835381 | 9.353013 | 4.575659 | 2.103059 | 1.468330 | 1.569679 | 1.862168 | 1.689026 | 1.274964 | 1.060579 | 1.160094 | 1.181777 | 0.889314 | 0.498204 | 0.468245 | 0.656915 | 0.815305 | 0.704706 | 0.544463 | 0.468538 | 0.569044 | 0.449203 | 0.301765 | 0.251961 | 0.230559 | 0.234650 | 0.232154 | 0.221138 | 0.215674 | 0.222155 | 0.245803 | 0.282435 | 0.235977 | 0.218630 | 0.217394 | 0.215022 | 0.220298 | 0.237025 | 0.241684 | 0.228829 | 0.218781 | 0.216776 | 0.214635 | 0.219895 | 0.215800 | 0.226535 | 0.242681 | 0.257865 | 0.276511 | 0.293154 | 0.271894 | 0.251003 | 0.218370 | 0.214633 | 0.215089 | 0.218400 | 0.224925 | 0.222121 | 0.224970 | 0.215478 | 0.215021 | 0.215018 | 0.214634 | 0.215397 | 0.219150 | 0.227187 | 0.250400 | 0.268573 | 0.299120 | 0.335281 | 0.333044 | 0.294727 | 0.246950 | 0.221550 | 0.218419 | 0.215706 | 0.216686 | 0.218019 | 0.228712 | 0.257841 | 0.333373 | 0.417580 | 0.553828 | 0.684352 | 0.750435 | 0.571791 | 0.410440 | 0.303822 | 0.248460 | 0.232920 | 0.230132 | 0.216032 | 0.216151 | 0.214700 | 0.214698 | 0.215864 | 0.216910 | 0.219489 | 0.214960 | 0.215885 | 0.215682 | 0.218473 | 0.230529 | 0.224370 | 0.215807 | 0.215763 | 0.220634 | 0.229657 | 0.235485 | 0.248791 | 0.241757 | 0.243257 | 0.238403 | 0.242745 | 0.238369 | 0.255005 | 0.280105 | 0.324870 | 0.354216 | 0.333478 | 0.298562 | 0.302663 | 0.292337 | 0.295244 | 0.341840 | 0.316318 | 0.313109 | 0.331559 | 0.352703 | 0.361824 | 0.340936 | 0.254804 | 0.237499 | 0.225005 | 0.215600 | 0.214743 | 0.217078 | 0.226957 |
| right anterior | 0.366760 | 0.380053 | 0.427401 | 0.432963 | 0.300835 | 0.255301 | 0.278516 | 0.337221 | 0.286544 | 0.237405 | 0.227209 | 0.219189 | 0.215298 | 0.226940 | 0.312241 | 0.241649 | 0.223616 | 0.215441 | 0.216684 | 0.240180 | 0.297825 | 0.354467 | 0.363509 | 0.443498 | 0.347225 | 0.222298 | 0.256721 | 0.300418 | 0.279514 | 0.362483 | 0.370980 | 0.338004 | 0.269561 | 0.237587 | 0.235466 | 0.349579 | 0.436790 | 0.638192 | 0.872942 | 0.422832 | 0.242852 | 0.225133 | 0.232922 | 0.231869 | 0.217597 | 0.235938 | 0.327914 | 0.770776 | 1.400567 | 5.116247 | 3.976407 | 1.345824 | 0.634289 | 0.301988 | 0.284787 | 0.241397 | 0.216506 | 0.217032 | 0.230457 | 0.241283 | 0.304667 | 0.300556 | 0.399651 | 0.440995 | 0.419265 | 0.531609 | 1.335842 | 1.943978 | 2.487824 | 1.646630 | 1.752353 | 1.445198 | 0.670913 | 0.359208 | 0.473771 | 0.493793 | 0.423194 | 0.358974 | 0.363997 | 0.478236 | 0.466749 | 0.293136 | 0.250764 | 0.263750 | 0.292651 | 0.433145 | 0.583577 | 0.597511 | 0.486452 | 0.404783 | 0.378731 | 0.367588 | 0.338842 | 0.311103 | 0.351088 | 0.393567 | 0.331935 | 0.233092 | 0.215306 | 0.227144 | 0.229565 | 0.238686 | 0.231213 | 0.223145 | 0.215851 | 0.228239 | 0.237491 | 0.241124 | 0.257481 | 0.297393 | 0.323628 | 0.291519 | 0.276497 | 0.267734 | 0.251461 | 0.298408 | 0.355962 | 0.441000 | 0.682929 | 0.940406 | 1.325607 | 1.817621 | 1.311906 | 0.925660 | 0.795454 | 0.622645 | 0.493186 | 0.356167 | 0.292646 | 0.266824 | 0.243032 | 0.225775 | 0.224406 | 0.230980 | 0.240104 | 0.245988 | 0.230617 | 0.229706 | 0.229833 | 0.230248 | 0.238767 | 0.249060 | 0.294651 | 0.348616 | 0.349576 | 0.347440 | 0.339739 | 0.325357 | 0.325008 | 0.282121 | 0.263997 | 0.246118 | 0.232946 | 0.234013 | 0.227537 | 0.221946 | 0.215147 | 0.214626 | 0.217849 | 0.226604 | 0.223687 | 0.223357 | 0.218148 | 0.218737 | 0.216250 | 0.214764 | 0.215874 | 0.216346 | 0.217321 | 0.220017 | 0.225164 | 0.233553 | 0.230356 | 0.216035 | 0.215026 | 0.215289 | 0.218705 | 0.233330 | 0.251963 | 0.321227 | 0.480148 | 0.574891 | 0.744680 | 0.716909 | 0.622757 | 0.646840 | 0.482518 | 0.392086 | 0.389681 | 0.358994 | 0.379305 | 0.446099 | 0.548202 | 0.619388 | 0.539608 | 0.397981 | 0.261223 | 0.224729 | 0.216493 | 0.215060 | 0.216008 | 0.227257 | 0.255350 | 0.330692 | 0.443785 | 0.602164 | 0.837871 | 0.743111 | 0.521006 | 0.372406 | 0.306337 | 0.303605 | 0.282121 | 0.270021 | 0.239661 | 0.219817 | 0.215076 | 0.214652 | 0.216172 | 0.218244 | 0.220010 | 0.216236 | 0.215060 | 0.214871 | 0.218960 | 0.231545 | 0.262211 | 0.264041 | 0.255128 | 0.274736 | 0.310638 | 0.280920 | 0.277138 | 0.267283 | 0.281886 | 0.331784 | 0.317233 | 0.275872 | 0.311480 | 0.292978 | 0.261225 | 0.240810 | 0.221518 | 0.221521 | 0.237082 | 0.231750 | 0.226244 | 0.222492 | 0.231645 | 0.270318 | 0.327414 | 0.383320 | 0.470282 | 0.521139 | 0.561488 | 0.529696 | 0.485716 | 0.464532 | 0.397906 | 0.400402 | 0.495777 | 0.530757 | 0.655717 | 0.546054 | 0.566295 | 0.439982 | 0.338466 | 0.293875 | 0.310512 | 0.294364 | 0.268683 | 0.228050 | 0.249562 | 0.249511 | 0.237112 | 0.230028 | 0.224243 | 0.223587 | 0.233446 | 0.231823 |
| left central | 0.755492 | 0.526587 | 0.458225 | 0.442896 | 0.290369 | 0.267884 | 0.228691 | 0.214740 | 0.214677 | 0.214968 | 0.214690 | 0.216666 | 0.214724 | 0.228904 | 0.216956 | 0.226588 | 0.221264 | 0.233315 | 0.266496 | 0.417383 | 0.642733 | 0.488600 | 0.472348 | 0.459280 | 0.317373 | 0.237541 | 0.215585 | 0.214701 | 0.214906 | 0.214756 | 0.214873 | 0.216965 | 0.216932 | 0.218825 | 0.214684 | 0.220281 | 0.235792 | 0.262629 | 0.239860 | 0.216707 | 0.273090 | 0.477880 | 0.564523 | 0.566432 | 1.127652 | 1.004132 | 0.642789 | 0.363554 | 0.350748 | 0.303783 | 0.395965 | 0.446635 | 0.526253 | 0.807371 | 1.297938 | 0.332087 | 0.253698 | 0.224521 | 0.227685 | 0.217214 | 0.216556 | 0.214789 | 0.219064 | 0.215897 | 0.215250 | 0.216606 | 0.225943 | 0.229716 | 0.250125 | 0.258750 | 0.250935 | 0.253189 | 0.298147 | 0.390546 | 0.592328 | 0.804001 | 0.664682 | 0.479212 | 0.314710 | 0.229354 | 0.214758 | 0.265076 | 0.370777 | 0.395154 | 0.448136 | 0.559096 | 0.637240 | 0.614760 | 0.498870 | 0.549888 | 0.716735 | 0.729869 | 0.407930 | 0.276605 | 0.215679 | 0.230720 | 0.244109 | 0.230119 | 0.220350 | 0.217620 | 0.240467 | 0.279259 | 0.360462 | 0.339267 | 0.281661 | 0.260976 | 0.246567 | 0.256832 | 0.307226 | 0.308481 | 0.315722 | 0.313684 | 0.284490 | 0.328287 | 0.247835 | 0.217878 | 0.215480 | 0.215175 | 0.214626 | 0.215088 | 0.216142 | 0.217830 | 0.238393 | 0.266653 | 0.270717 | 0.331441 | 0.474329 | 0.629350 | 0.616468 | 0.647184 | 0.552490 | 0.649013 | 0.602898 | 0.454175 | 0.537493 | 0.531561 | 0.446021 | 0.354855 | 0.319636 | 0.319435 | 0.312101 | 0.291294 | 0.305327 | 0.262905 | 0.292178 | 0.286521 | 0.288956 | 0.280168 | 0.260463 | 0.244803 | 0.275081 | 0.314314 | 0.382880 | 0.369526 | 0.417827 | 0.488793 | 0.526950 | 0.618391 | 0.526146 | 0.450757 | 0.494423 | 0.519336 | 0.488580 | 0.423779 | 0.309794 | 0.265624 | 0.245398 | 0.219542 | 0.216848 | 0.221619 | 0.229110 | 0.279036 | 0.350419 | 0.376691 | 0.355064 | 0.312986 | 0.285150 | 0.266996 | 0.224082 | 0.215321 | 0.214633 | 0.218737 | 0.238095 | 0.237451 | 0.247241 | 0.274219 | 0.289466 | 0.305132 | 0.268423 | 0.244500 | 0.240820 | 0.243375 | 0.249660 | 0.261613 | 0.270785 | 0.307215 | 0.329233 | 0.351996 | 0.309907 | 0.288766 | 0.276427 | 0.295170 | 0.317121 | 0.323531 | 0.364248 | 0.524433 | 0.759729 | 0.735249 | 0.665285 | 0.663527 | 0.668231 | 0.694730 | 0.557747 | 0.478959 | 0.546674 | 0.525403 | 0.435720 | 0.427589 | 0.422048 | 0.470280 | 0.476664 | 0.442747 | 0.436241 | 0.474135 | 0.475802 | 0.489918 | 0.468030 | 0.454671 | 0.447271 | 0.357174 | 0.281200 | 0.261065 | 0.234110 | 0.220706 | 0.215583 | 0.214898 | 0.222866 | 0.239453 | 0.239120 | 0.246609 | 0.272017 | 0.269885 | 0.260588 | 0.229692 | 0.225172 | 0.229502 | 0.224440 | 0.214626 | 0.222115 | 0.238645 | 0.242639 | 0.277493 | 0.351773 | 0.393672 | 0.341856 | 0.296339 | 0.287397 | 0.322838 | 0.298313 | 0.284014 | 0.270079 | 0.278866 | 0.272280 | 0.243013 | 0.217088 | 0.215452 | 0.215446 | 0.215244 | 0.217918 | 0.225661 | 0.248784 | 0.240617 | 0.224277 | 0.214736 | 0.215065 | 0.214651 | 0.216387 | 0.225581 | 0.226990 | 0.219837 |
| right central | 0.334063 | 0.350512 | 0.254809 | 0.220685 | 0.216315 | 0.215262 | 0.217656 | 0.224188 | 0.236436 | 0.220411 | 0.221501 | 0.222535 | 0.222996 | 0.234637 | 0.258807 | 0.287077 | 0.296451 | 0.224617 | 0.216305 | 0.217908 | 0.221716 | 0.232941 | 0.235664 | 0.256028 | 0.222689 | 0.249968 | 0.501653 | 0.439444 | 0.440664 | 0.479845 | 0.512748 | 0.306814 | 0.214687 | 0.253942 | 0.256119 | 0.297096 | 0.310396 | 0.244866 | 0.215260 | 0.214626 | 0.226651 | 0.301235 | 0.377714 | 0.524140 | 0.660846 | 0.706393 | 0.623272 | 0.427620 | 0.274419 | 0.215907 | 0.216013 | 0.216886 | 0.219618 | 0.216004 | 0.215682 | 0.215878 | 0.223033 | 0.226677 | 0.223092 | 0.234521 | 0.225020 | 0.228406 | 0.221202 | 0.215203 | 0.216469 | 0.215205 | 0.216734 | 0.232371 | 0.273792 | 0.388282 | 0.589395 | 1.201703 | 1.806364 | 2.511333 | 3.999899 | 4.204917 | 3.549591 | 1.694431 | 0.712402 | 0.451940 | 0.334422 | 0.276884 | 0.279155 | 0.247927 | 0.251303 | 0.221657 | 0.214776 | 0.216035 | 0.216778 | 0.224975 | 0.244996 | 0.267745 | 0.273246 | 0.339205 | 0.381694 | 0.472999 | 0.781596 | 0.747152 | 1.019284 | 0.943508 | 0.782018 | 0.735091 | 0.589895 | 0.275987 | 0.231536 | 0.216659 | 0.216726 | 0.217057 | 0.226993 | 0.233710 | 0.241723 | 0.313441 | 0.333243 | 0.311379 | 0.320141 | 0.346250 | 0.311597 | 0.314220 | 0.289831 | 0.297037 | 0.320758 | 0.309623 | 0.352780 | 0.486035 | 0.663492 | 0.919751 | 1.040718 | 0.876325 | 0.699041 | 0.496128 | 0.474774 | 0.364839 | 0.310336 | 0.324999 | 0.335094 | 0.365813 | 0.420732 | 0.389505 | 0.473771 | 0.583267 | 0.647860 | 0.758727 | 0.650165 | 0.593607 | 0.731075 | 0.686051 | 0.816318 | 1.091215 | 1.109750 | 2.030852 | 2.854606 | 2.140458 | 1.377186 | 0.775565 | 0.476999 | 0.458498 | 0.367798 | 0.380410 | 0.473498 | 0.573689 | 0.593064 | 0.502418 | 0.467275 | 0.532685 | 0.504956 | 0.464271 | 0.390172 | 0.327434 | 0.298987 | 0.274247 | 0.237095 | 0.222527 | 0.221372 | 0.259006 | 0.315448 | 0.284401 | 0.233690 | 0.218049 | 0.215812 | 0.220073 | 0.261172 | 0.319737 | 0.368709 | 0.340041 | 0.291774 | 0.263820 | 0.246683 | 0.227847 | 0.229685 | 0.223490 | 0.223479 | 0.231861 | 0.245725 | 0.269625 | 0.302331 | 0.269803 | 0.270684 | 0.272374 | 0.282651 | 0.291602 | 0.265307 | 0.222001 | 0.219035 | 0.222260 | 0.217362 | 0.216663 | 0.242343 | 0.251327 | 0.233767 | 0.223825 | 0.242448 | 0.244075 | 0.221895 | 0.215369 | 0.216558 | 0.218638 | 0.217786 | 0.220115 | 0.238181 | 0.238014 | 0.233200 | 0.225814 | 0.222143 | 0.217818 | 0.215985 | 0.217343 | 0.215098 | 0.217914 | 0.241813 | 0.287570 | 0.341521 | 0.445552 | 0.643737 | 0.722336 | 0.805766 | 0.849241 | 0.667762 | 0.556184 | 0.419987 | 0.322795 | 0.335138 | 0.323369 | 0.284669 | 0.348428 | 0.409409 | 0.542943 | 0.798854 | 0.940605 | 0.766713 | 0.712340 | 0.567385 | 0.427759 | 0.506288 | 0.712782 | 1.035624 | 1.249012 | 1.281713 | 1.193156 | 1.226539 | 0.903138 | 0.657603 | 0.470743 | 0.455762 | 0.470323 | 0.530063 | 0.647484 | 0.707848 | 0.646874 | 0.614809 | 0.699031 | 0.908616 | 1.093712 | 0.942954 | 1.084550 | 1.100171 | 0.991746 | 0.791193 | 0.594792 | 0.445556 | 0.395620 |
| left posterior | 0.232618 | 0.215824 | 0.239013 | 0.341578 | 0.414498 | 0.869080 | 0.593936 | 0.409191 | 0.295226 | 0.215476 | 0.238643 | 0.257469 | 0.229758 | 0.215352 | 0.223512 | 0.317795 | 0.506754 | 0.467548 | 0.316793 | 0.228271 | 0.233275 | 0.291721 | 0.295224 | 0.270808 | 0.231717 | 0.225706 | 0.219777 | 0.214645 | 0.223345 | 0.281274 | 0.299397 | 0.262303 | 0.246258 | 0.236716 | 0.341672 | 0.280433 | 0.216141 | 0.220207 | 0.224862 | 0.222372 | 0.230492 | 0.274275 | 0.318467 | 0.391232 | 0.597989 | 0.471691 | 0.493399 | 0.471093 | 0.490000 | 0.507446 | 0.467696 | 0.524103 | 0.852870 | 0.857139 | 1.161542 | 0.873783 | 0.522060 | 0.433614 | 0.279539 | 0.234308 | 0.226873 | 0.443139 | 1.153660 | 2.696044 | 3.032029 | 1.666535 | 1.260429 | 0.632409 | 0.408793 | 0.289446 | 0.251332 | 0.250575 | 0.280650 | 0.313916 | 0.310566 | 0.280833 | 0.260126 | 0.231955 | 0.214626 | 0.219772 | 0.237501 | 0.248569 | 0.247127 | 0.244704 | 0.248379 | 0.237502 | 0.243838 | 0.260834 | 0.271689 | 0.284060 | 0.252042 | 0.239720 | 0.227327 | 0.220050 | 0.214842 | 0.218632 | 0.229130 | 0.224675 | 0.225194 | 0.225281 | 0.236526 | 0.237172 | 0.241969 | 0.242999 | 0.255538 | 0.259076 | 0.280122 | 0.298852 | 0.300694 | 0.298921 | 0.272672 | 0.264669 | 0.273072 | 0.260744 | 0.233313 | 0.220609 | 0.214913 | 0.214802 | 0.218732 | 0.220542 | 0.222114 | 0.215947 | 0.216526 | 0.239232 | 0.264876 | 0.284895 | 0.264271 | 0.251890 | 0.264293 | 0.234307 | 0.214849 | 0.214682 | 0.215388 | 0.214928 | 0.214714 | 0.223482 | 0.234581 | 0.249466 | 0.259751 | 0.264289 | 0.306078 | 0.315549 | 0.346514 | 0.360385 | 0.321751 | 0.305702 | 0.302047 | 0.253249 | 0.253123 | 0.248113 | 0.243063 | 0.246739 | 0.245970 | 0.268037 | 0.326262 | 0.371968 | 0.350803 | 0.341832 | 0.357035 | 0.378420 | 0.331698 | 0.285129 | 0.257997 | 0.255961 | 0.261452 | 0.252096 | 0.244321 | 0.240002 | 0.243715 | 0.242309 | 0.228392 | 0.220318 | 0.218237 | 0.217223 | 0.215048 | 0.214798 | 0.217388 | 0.222543 | 0.222152 | 0.227466 | 0.232624 | 0.221807 | 0.215963 | 0.214639 | 0.214630 | 0.214684 | 0.214720 | 0.215024 | 0.219236 | 0.228763 | 0.227587 | 0.221014 | 0.215910 | 0.215649 | 0.221196 | 0.226297 | 0.238915 | 0.231996 | 0.226222 | 0.217374 | 0.216371 | 0.220045 | 0.222357 | 0.227175 | 0.230798 | 0.239192 | 0.241851 | 0.232777 | 0.239245 | 0.246689 | 0.264341 | 0.272040 | 0.273523 | 0.311648 | 0.390466 | 0.421839 | 0.472280 | 0.368309 | 0.313220 | 0.292928 | 0.255751 | 0.220082 | 0.214866 | 0.223505 | 0.220290 | 0.218333 | 0.218561 | 0.214718 | 0.222554 | 0.254551 | 0.375355 | 0.622960 | 0.834687 | 1.246789 | 1.171127 | 1.169855 | 1.398770 | 1.442042 | 1.314512 | 1.245142 | 0.687746 | 0.432862 | 0.274581 | 0.214835 | 0.228247 | 0.269863 | 0.269647 | 0.237357 | 0.216420 | 0.218356 | 0.233657 | 0.251063 | 0.304411 | 0.307693 | 0.305055 | 0.271083 | 0.248645 | 0.267536 | 0.271430 | 0.260707 | 0.249720 | 0.252865 | 0.272005 | 0.285929 | 0.268733 | 0.258633 | 0.239269 | 0.234827 | 0.221399 | 0.214785 | 0.214894 | 0.214693 | 0.215365 | 0.214801 | 0.215150 | 0.215763 | 0.214626 | 0.214950 | 0.214952 | 0.223891 |
| right posterior | 0.239852 | 0.226898 | 0.221262 | 0.223150 | 0.237264 | 0.246098 | 0.278706 | 0.315356 | 0.365668 | 0.417691 | 0.490905 | 0.421763 | 0.454954 | 0.445740 | 0.422504 | 0.354672 | 0.408541 | 0.319712 | 0.263207 | 0.231049 | 0.214902 | 0.216075 | 0.216593 | 0.227874 | 0.219002 | 0.214784 | 0.220053 | 0.227376 | 0.225295 | 0.214649 | 0.216003 | 0.215004 | 0.215163 | 0.215669 | 0.238102 | 0.315994 | 0.638433 | 1.304532 | 0.687183 | 0.623339 | 0.490566 | 0.325051 | 0.271689 | 0.223998 | 0.214811 | 0.214672 | 0.229059 | 0.262665 | 0.268783 | 0.303691 | 0.341324 | 0.327067 | 0.349620 | 0.291901 | 0.245578 | 0.235162 | 0.215040 | 0.239754 | 0.315250 | 0.402685 | 0.467246 | 0.478666 | 0.562767 | 0.599057 | 0.482384 | 0.352804 | 0.316881 | 0.321079 | 0.368086 | 0.470638 | 0.585910 | 0.849143 | 1.611013 | 4.072569 | 4.519895 | 2.699032 | 1.094674 | 0.476414 | 0.286628 | 0.243228 | 0.221890 | 0.215928 | 0.216915 | 0.215616 | 0.215499 | 0.217171 | 0.219309 | 0.240993 | 0.264705 | 0.285355 | 0.271981 | 0.274281 | 0.265915 | 0.236672 | 0.232321 | 0.233369 | 0.237632 | 0.258515 | 0.285682 | 0.356228 | 0.610866 | 1.034268 | 1.550125 | 1.981566 | 1.644192 | 0.943134 | 0.765284 | 0.638146 | 0.611760 | 0.598002 | 0.525452 | 0.390860 | 0.347140 | 0.308829 | 0.269575 | 0.257772 | 0.237287 | 0.223970 | 0.223491 | 0.221174 | 0.221941 | 0.225640 | 0.214626 | 0.221770 | 0.225527 | 0.217600 | 0.214671 | 0.215387 | 0.214673 | 0.225068 | 0.237266 | 0.250458 | 0.262977 | 0.257218 | 0.251112 | 0.267451 | 0.262084 | 0.264245 | 0.247930 | 0.225976 | 0.228993 | 0.242430 | 0.246743 | 0.271689 | 0.332004 | 0.441886 | 0.550053 | 0.609811 | 0.721925 | 0.767666 | 0.808999 | 0.827593 | 0.696196 | 0.622164 | 0.604833 | 0.523841 | 0.483010 | 0.371553 | 0.324408 | 0.290998 | 0.298348 | 0.296900 | 0.308411 | 0.307885 | 0.369217 | 0.387928 | 0.473227 | 0.474853 | 0.411692 | 0.422852 | 0.462616 | 0.430645 | 0.389749 | 0.329365 | 0.283384 | 0.262968 | 0.243038 | 0.223374 | 0.221261 | 0.220604 | 0.217498 | 0.218445 | 0.219116 | 0.217342 | 0.217114 | 0.214626 | 0.216162 | 0.217626 | 0.216990 | 0.217975 | 0.216461 | 0.214695 | 0.216027 | 0.225385 | 0.248972 | 0.257774 | 0.277076 | 0.268608 | 0.269880 | 0.282238 | 0.265877 | 0.239386 | 0.222551 | 0.220255 | 0.218737 | 0.218585 | 0.215705 | 0.219026 | 0.222133 | 0.233629 | 0.240159 | 0.248637 | 0.234938 | 0.228545 | 0.224233 | 0.227503 | 0.224910 | 0.219667 | 0.217952 | 0.221273 | 0.234854 | 0.245308 | 0.257601 | 0.267553 | 0.264136 | 0.267305 | 0.256085 | 0.238478 | 0.227312 | 0.220411 | 0.215774 | 0.215367 | 0.214749 | 0.214626 | 0.216046 | 0.220527 | 0.223573 | 0.227105 | 0.244728 | 0.309680 | 0.403498 | 0.469762 | 0.437832 | 0.407590 | 0.400571 | 0.377302 | 0.326219 | 0.299484 | 0.280690 | 0.277956 | 0.281113 | 0.296415 | 0.297719 | 0.290760 | 0.275222 | 0.260548 | 0.254941 | 0.249603 | 0.241615 | 0.235277 | 0.222155 | 0.218084 | 0.215785 | 0.215033 | 0.214797 | 0.215807 | 0.217470 | 0.223628 | 0.234176 | 0.249908 | 0.253104 | 0.247662 | 0.234611 | 0.223510 | 0.215978 | 0.214962 | 0.218380 | 0.220066 | 0.224441 | 0.227482 |
| all electrodes | 0.241537 | 0.248557 | 0.238329 | 0.217122 | 0.224177 | 0.219607 | 0.251997 | 0.239941 | 0.247581 | 0.251813 | 0.240085 | 0.223222 | 0.222554 | 0.214696 | 0.221943 | 0.243251 | 0.232255 | 0.216486 | 0.214692 | 0.216267 | 0.221165 | 0.216474 | 0.228031 | 0.241589 | 0.235614 | 0.217783 | 0.218622 | 0.214749 | 0.215914 | 0.214626 | 0.242145 | 0.244238 | 0.215640 | 0.235670 | 0.218180 | 0.396101 | 0.786424 | 1.162409 | 1.259130 | 1.183760 | 0.838627 | 0.307889 | 0.214641 | 0.239841 | 0.306606 | 0.369311 | 0.448891 | 0.485104 | 0.411239 | 0.317834 | 0.289136 | 0.305271 | 0.287570 | 0.312889 | 0.273939 | 0.222487 | 0.223218 | 0.247514 | 0.344285 | 0.471523 | 0.939492 | 0.888157 | 0.957800 | 0.613035 | 0.500859 | 0.412991 | 0.446693 | 0.571797 | 0.828010 | 0.785762 | 0.861190 | 1.242147 | 1.992680 | 2.248643 | 3.725118 | 3.913249 | 2.154658 | 1.500551 | 1.553676 | 1.590376 | 1.731874 | 0.749846 | 0.461174 | 0.516659 | 0.418247 | 0.290799 | 0.220144 | 0.223914 | 0.258023 | 0.268825 | 0.309106 | 0.337358 | 0.420571 | 0.394741 | 0.331146 | 0.290083 | 0.293539 | 0.264834 | 0.251183 | 0.218919 | 0.221485 | 0.218651 | 0.227153 | 0.218451 | 0.215909 | 0.222117 | 0.243770 | 0.269170 | 0.302258 | 0.288043 | 0.305094 | 0.336402 | 0.264911 | 0.226190 | 0.215300 | 0.238262 | 0.270015 | 0.281916 | 0.334695 | 0.368664 | 0.477360 | 0.637405 | 0.618162 | 0.647161 | 0.828782 | 1.009769 | 0.679147 | 0.410624 | 0.292195 | 0.241084 | 0.219240 | 0.214993 | 0.215219 | 0.214634 | 0.214866 | 0.214787 | 0.216812 | 0.215422 | 0.215735 | 0.215203 | 0.214654 | 0.216155 | 0.216399 | 0.216891 | 0.217089 | 0.214661 | 0.219390 | 0.231666 | 0.227743 | 0.223365 | 0.215082 | 0.222140 | 0.273028 | 0.308217 | 0.347206 | 0.383715 | 0.563196 | 0.782528 | 0.466771 | 0.297524 | 0.249199 | 0.225665 | 0.225650 | 0.214635 | 0.223824 | 0.221188 | 0.214827 | 0.215063 | 0.216052 | 0.215783 | 0.214841 | 0.223707 | 0.248685 | 0.230548 | 0.251163 | 0.251949 | 0.256014 | 0.240343 | 0.238031 | 0.220835 | 0.214636 | 0.220688 | 0.227829 | 0.232041 | 0.247592 | 0.280082 | 0.296419 | 0.290595 | 0.282962 | 0.269481 | 0.248646 | 0.261570 | 0.295122 | 0.276290 | 0.235795 | 0.217200 | 0.215056 | 0.216423 | 0.216172 | 0.216150 | 0.216904 | 0.215199 | 0.220688 | 0.227979 | 0.243218 | 0.241986 | 0.219765 | 0.214706 | 0.217646 | 0.217470 | 0.221335 | 0.237201 | 0.241880 | 0.222911 | 0.214686 | 0.218350 | 0.217518 | 0.217142 | 0.215843 | 0.214679 | 0.214679 | 0.214995 | 0.220537 | 0.234132 | 0.246064 | 0.230503 | 0.228624 | 0.220454 | 0.215099 | 0.215185 | 0.223420 | 0.225368 | 0.220010 | 0.222798 | 0.221992 | 0.224982 | 0.242123 | 0.260176 | 0.322810 | 0.331379 | 0.299569 | 0.271897 | 0.242568 | 0.220036 | 0.216282 | 0.216521 | 0.219616 | 0.214664 | 0.220998 | 0.233822 | 0.277941 | 0.319616 | 0.346795 | 0.289419 | 0.259597 | 0.235941 | 0.230030 | 0.216298 | 0.215118 | 0.217225 | 0.215038 | 0.216387 | 0.219172 | 0.237589 | 0.237437 | 0.220623 | 0.215593 | 0.214694 | 0.214690 | 0.214741 | 0.215403 | 0.217129 | 0.215569 | 0.214626 | 0.218013 | 0.238908 | 0.305234 | 0.342010 | 0.376030 | 0.445152 |

Searchlight, spatiotemporal cluster permutation test

No significant clusters observed.

Y) emoji to real faces - neutral vs happy

  
|  | time window | peak latency | cluster *p* | peak Cohen's *d* |  | | | |
| **all electrodes** | 105 - 160 ms | 135 ms | 0.0468 | 1.1402 |  | | | |
|  | | | | | | | | |

Time-resolved classification, cluster permutation tests

|  | **left hemisphere** | | | | **right hemisphere** | | | |
|  | time window | peak latency | cluster *p* | peak Cohen's *d* | time window | peak latency | cluster *p* | peak Cohen's *d* |
| **anterior** |  | | | |  | | | |
| **central** |  | | | |  | | | |
| **posterior** | 110 - 160 ms | 130 ms | 0.0341 | 1.4281 | 100 - 170 ms | 130 ms | 0.0226 | 1.3913 |
 250 - 315 ms | 290 ms | 0.0441 | 0.8275 |  | | | | 710 - 910 ms | 825 ms | 0.0104 | 0.5813 |  | | | | 925 - 1145 ms | 1120 ms | 0.007 | 0.6887 |  | | | |

  

Time-resolved classification, Bayesian statistics

|  | -200 | -195 | -190 | -185 | -180 | -175 | -170 | -165 | -160 | -155 | -150 | -145 | -140 | -135 | -130 | -125 | -120 | -115 | -110 | -105 | -100 | -95 | -90 | -85 | -80 | -75 | -70 | -65 | -60 | -55 | -50 | -45 | -40 | -35 | -30 | -25 | -20 | -15 | -10 | -5 | 0 | 5 | 10 | 15 | 20 | 25 | 30 | 35 | 40 | 45 | 50 | 55 | 60 | 65 | 70 | 75 | 80 | 85 | 90 | 95 | 100 | 105 | 110 | 115 | 120 | 125 | 130 | 135 | 140 | 145 | 150 | 155 | 160 | 165 | 170 | 175 | 180 | 185 | 190 | 195 | 200 | 205 | 210 | 215 | 220 | 225 | 230 | 235 | 240 | 245 | 250 | 255 | 260 | 265 | 270 | 275 | 280 | 285 | 290 | 295 | 300 | 305 | 310 | 315 | 320 | 325 | 330 | 335 | 340 | 345 | 350 | 355 | 360 | 365 | 370 | 375 | 380 | 385 | 390 | 395 | 400 | 405 | 410 | 415 | 420 | 425 | 430 | 435 | 440 | 445 | 450 | 455 | 460 | 465 | 470 | 475 | 480 | 485 | 490 | 495 | 500 | 505 | 510 | 515 | 520 | 525 | 530 | 535 | 540 | 545 | 550 | 555 | 560 | 565 | 570 | 575 | 580 | 585 | 590 | 595 | 600 | 605 | 610 | 615 | 620 | 625 | 630 | 635 | 640 | 645 | 650 | 655 | 660 | 665 | 670 | 675 | 680 | 685 | 690 | 695 | 700 | 705 | 710 | 715 | 720 | 725 | 730 | 735 | 740 | 745 | 750 | 755 | 760 | 765 | 770 | 775 | 780 | 785 | 790 | 795 | 800 | 805 | 810 | 815 | 820 | 825 | 830 | 835 | 840 | 845 | 850 | 855 | 860 | 865 | 870 | 875 | 880 | 885 | 890 | 895 | 900 | 905 | 910 | 915 | 920 | 925 | 930 | 935 | 940 | 945 | 950 | 955 | 960 | 965 | 970 | 975 | 980 | 985 | 990 | 995 | 1000 | 1005 | 1010 | 1015 | 1020 | 1025 | 1030 | 1035 | 1040 | 1045 | 1050 | 1055 | 1060 | 1065 | 1070 | 1075 | 1080 | 1085 | 1090 | 1095 | 1100 | 1105 | 1110 | 1115 | 1120 | 1125 | 1130 | 1135 | 1140 | 1145 | 1150 | 1155 | 1160 | 1165 | 1170 | 1175 | 1180 | 1185 | 1190 | 1195 |
| --- | --- | --- | --- | --- | --- | --- | --- | --- | --- | --- | --- | --- | --- | --- | --- | --- | --- | --- | --- | --- | --- | --- | --- | --- | --- | --- | --- | --- | --- | --- | --- | --- | --- | --- | --- | --- | --- | --- | --- | --- | --- | --- | --- | --- | --- | --- | --- | --- | --- | --- | --- | --- | --- | --- | --- | --- | --- | --- | --- | --- | --- | --- | --- | --- | --- | --- | --- | --- | --- | --- | --- | --- | --- | --- | --- | --- | --- | --- | --- | --- | --- | --- | --- | --- | --- | --- | --- | --- | --- | --- | --- | --- | --- | --- | --- | --- | --- | --- | --- | --- | --- | --- | --- | --- | --- | --- | --- | --- | --- | --- | --- | --- | --- | --- | --- | --- | --- | --- | --- | --- | --- | --- | --- | --- | --- | --- | --- | --- | --- | --- | --- | --- | --- | --- | --- | --- | --- | --- | --- | --- | --- | --- | --- | --- | --- | --- | --- | --- | --- | --- | --- | --- | --- | --- | --- | --- | --- | --- | --- | --- | --- | --- | --- | --- | --- | --- | --- | --- | --- | --- | --- | --- | --- | --- | --- | --- | --- | --- | --- | --- | --- | --- | --- | --- | --- | --- | --- | --- | --- | --- | --- | --- | --- | --- | --- | --- | --- | --- | --- | --- | --- | --- | --- | --- | --- | --- | --- | --- | --- | --- | --- | --- | --- | --- | --- | --- | --- | --- | --- | --- | --- | --- | --- | --- | --- | --- | --- | --- | --- | --- | --- | --- | --- | --- | --- | --- | --- | --- | --- | --- | --- | --- | --- | --- | --- | --- | --- | --- | --- | --- | --- | --- | --- | --- | --- | --- | --- | --- | --- | --- | --- | --- | --- | --- | --- | --- | --- | --- | --- | --- | --- | --- | --- | --- | --- | --- | --- | --- | --- | --- |
| left anterior | 0.827286 | 0.745638 | 0.702332 | 0.611228 | 0.420465 | 0.268484 | 0.215345 | 0.216796 | 0.215026 | 0.239907 | 0.297962 | 0.332170 | 0.611535 | 1.914461 | 3.205248 | 1.263843 | 0.332959 | 0.217475 | 0.218528 | 0.214687 | 0.225592 | 0.235485 | 0.215548 | 0.215794 | 0.237055 | 0.283869 | 0.523574 | 0.595545 | 0.751608 | 0.405569 | 0.586061 | 1.682537 | 1.437284 | 0.506728 | 0.423013 | 0.315541 | 0.551792 | 0.902162 | 0.482464 | 0.348643 | 0.566757 | 0.662535 | 0.588446 | 0.289807 | 0.227750 | 0.231206 | 0.265289 | 0.243266 | 0.284553 | 0.439306 | 0.651291 | 0.725267 | 0.567197 | 0.638255 | 0.630205 | 0.375467 | 0.278673 | 0.247845 | 0.276146 | 0.308906 | 0.274453 | 0.242233 | 0.227122 | 0.224161 | 0.404750 | 1.913978 | 4.395897 | 5.070277 | 3.661260 | 2.256859 | 0.873742 | 0.334791 | 0.221691 | 0.227334 | 0.293955 | 0.372691 | 0.360053 | 0.296121 | 0.271700 | 0.224606 | 0.215686 | 0.215731 | 0.219141 | 0.224345 | 0.228228 | 0.221056 | 0.215576 | 0.215277 | 0.243327 | 0.419764 | 0.691208 | 0.864699 | 1.567574 | 2.796303 | 3.047101 | 3.318367 | 1.669301 | 0.739308 | 0.573050 | 0.351619 | 0.258416 | 0.214953 | 0.232493 | 0.240643 | 0.233217 | 0.249370 | 0.252758 | 0.249743 | 0.216673 | 0.221022 | 0.248133 | 0.293278 | 0.332609 | 0.341753 | 0.323404 | 0.337397 | 0.317949 | 0.273700 | 0.241067 | 0.221819 | 0.217769 | 0.220609 | 0.225729 | 0.222915 | 0.223771 | 0.216763 | 0.214667 | 0.218189 | 0.228632 | 0.248768 | 0.289883 | 0.403505 | 0.430218 | 0.367305 | 0.306537 | 0.273761 | 0.300115 | 0.309555 | 0.286236 | 0.280766 | 0.283716 | 0.284455 | 0.290707 | 0.249831 | 0.220088 | 0.214952 | 0.214633 | 0.214653 | 0.215290 | 0.214726 | 0.215814 | 0.214781 | 0.216582 | 0.217599 | 0.215823 | 0.216702 | 0.214799 | 0.230091 | 0.245563 | 0.260593 | 0.258458 | 0.254422 | 0.290094 | 0.312684 | 0.351044 | 0.485269 | 0.675517 | 0.849427 | 0.600860 | 0.413884 | 0.349452 | 0.395732 | 0.434565 | 0.465205 | 0.502969 | 0.590122 | 0.673005 | 0.927626 | 0.931818 | 0.811566 | 0.737397 | 0.592971 | 0.546433 | 0.534305 | 0.511317 | 0.558004 | 0.508010 | 0.564381 | 0.549390 | 0.496891 | 0.428588 | 0.298035 | 0.246487 | 0.243091 | 0.251011 | 0.298749 | 0.318765 | 0.295979 | 0.280417 | 0.283307 | 0.263498 | 0.241587 | 0.227794 | 0.238982 | 0.262060 | 0.331658 | 0.369554 | 0.418744 | 0.402353 | 0.434765 | 0.438931 | 0.484678 | 0.504773 | 0.468472 | 0.434557 | 0.381164 | 0.348139 | 0.314232 | 0.306420 | 0.279183 | 0.269951 | 0.258579 | 0.264654 | 0.266677 | 0.299120 | 0.298512 | 0.336862 | 0.397787 | 0.474506 | 0.533246 | 0.555192 | 0.460144 | 0.420407 | 0.360280 | 0.350978 | 0.365992 | 0.417650 | 0.481752 | 0.506535 | 0.637997 | 0.651422 | 0.587659 | 0.522179 | 0.390838 | 0.326512 | 0.300002 | 0.259553 | 0.273268 | 0.281214 | 0.313324 | 0.412954 | 0.439316 | 0.397342 | 0.346580 | 0.275446 | 0.252429 | 0.229701 | 0.217229 | 0.216147 | 0.219863 | 0.215461 | 0.214838 | 0.214635 | 0.215518 | 0.222240 | 0.214833 | 0.216323 | 0.215241 | 0.214901 | 0.219098 | 0.221867 | 0.225897 | 0.236060 | 0.245338 | 0.250643 | 0.273191 | 0.271239 | 0.267505 | 0.256393 | 0.265566 |
| right anterior | 0.586476 | 0.658351 | 0.767138 | 1.101011 | 1.300791 | 0.818267 | 0.720371 | 0.890439 | 0.843219 | 0.700399 | 0.558660 | 0.396631 | 0.324797 | 0.306940 | 0.244566 | 0.214642 | 0.256213 | 0.459453 | 0.671475 | 0.447496 | 0.380297 | 0.300365 | 0.245633 | 0.224910 | 0.363989 | 0.704759 | 0.716361 | 0.796372 | 1.637811 | 1.812133 | 0.353999 | 0.246060 | 0.216778 | 0.215183 | 0.219583 | 0.248123 | 0.291817 | 0.219690 | 0.217366 | 0.216234 | 0.214878 | 0.214994 | 0.221651 | 0.238795 | 0.236163 | 0.264232 | 0.276148 | 0.350259 | 0.290916 | 0.232039 | 0.217026 | 0.215438 | 0.215813 | 0.237017 | 0.311475 | 0.336054 | 0.374292 | 0.411890 | 0.416000 | 0.551830 | 0.467883 | 0.432824 | 0.754943 | 2.135273 | 6.924135 | 29.525742 | 33.839222 | 73.632392 | 113.045016 | 56.419035 | 19.058454 | 7.742939 | 2.187184 | 1.211468 | 0.651321 | 0.391872 | 0.252523 | 0.221398 | 0.221695 | 0.246357 | 0.310869 | 0.382802 | 0.685467 | 0.816805 | 0.690849 | 0.468099 | 0.265142 | 0.219735 | 0.303199 | 0.528013 | 0.823274 | 0.801320 | 0.642218 | 0.638682 | 0.544756 | 0.456246 | 0.490036 | 0.528096 | 0.663973 | 0.500485 | 0.289468 | 0.217991 | 0.220031 | 0.249254 | 0.290866 | 0.295288 | 0.228121 | 0.222664 | 0.267693 | 0.295810 | 0.290941 | 0.257650 | 0.253302 | 0.241791 | 0.217627 | 0.214658 | 0.214635 | 0.218124 | 0.216894 | 0.214634 | 0.214761 | 0.215428 | 0.221862 | 0.229478 | 0.218730 | 0.225313 | 0.231213 | 0.224240 | 0.224945 | 0.220918 | 0.236113 | 0.300352 | 0.325439 | 0.312415 | 0.297609 | 0.274753 | 0.246616 | 0.221025 | 0.214967 | 0.216482 | 0.221708 | 0.215955 | 0.214706 | 0.217304 | 0.219975 | 0.216101 | 0.223975 | 0.236491 | 0.259039 | 0.306128 | 0.309439 | 0.282178 | 0.270886 | 0.263045 | 0.251246 | 0.216095 | 0.219990 | 0.230184 | 0.222408 | 0.219170 | 0.221809 | 0.231785 | 0.220349 | 0.214638 | 0.214724 | 0.214637 | 0.215133 | 0.216721 | 0.215326 | 0.215455 | 0.229034 | 0.223916 | 0.229685 | 0.219890 | 0.216409 | 0.214996 | 0.217821 | 0.237904 | 0.273769 | 0.297293 | 0.293787 | 0.285137 | 0.287246 | 0.390766 | 0.562471 | 0.478030 | 0.474119 | 0.464802 | 0.444328 | 0.433619 | 0.430695 | 0.474268 | 0.600128 | 0.708525 | 0.892910 | 1.507582 | 1.780820 | 1.470068 | 1.388731 | 1.903729 | 2.746662 | 3.519052 | 3.614464 | 3.490103 | 2.448067 | 1.649923 | 0.883464 | 0.679188 | 0.559718 | 0.540835 | 0.614122 | 0.752776 | 0.938564 | 1.890034 | 4.625462 | 9.789849 | 11.538876 | 9.198614 | 6.520816 | 3.502288 | 3.628735 | 2.869939 | 1.588451 | 1.556882 | 2.103696 | 3.227922 | 8.384568 | 6.991969 | 5.367708 | 6.297077 | 5.198439 | 3.647213 | 2.269077 | 1.435848 | 1.137957 | 0.857110 | 0.699460 | 0.647121 | 0.591951 | 0.602068 | 0.606054 | 0.621688 | 0.800192 | 0.851277 | 0.980459 | 1.089606 | 1.445238 | 1.732613 | 2.194159 | 2.114685 | 2.233978 | 2.316697 | 1.983737 | 1.967801 | 2.566714 | 3.571871 | 5.890572 | 9.166809 | 10.508470 | 15.801631 | 14.522009 | 9.853388 | 5.890376 | 3.635283 | 1.612156 | 1.059792 | 0.933877 | 0.729067 | 0.684558 | 0.601838 | 0.529604 | 0.604001 | 0.585849 | 0.579380 | 0.587596 | 0.514530 | 0.513635 | 0.526156 | 0.481320 | 0.437325 |
| left central | 0.214654 | 0.216396 | 0.219232 | 0.235185 | 0.261952 | 0.245939 | 0.224302 | 0.227126 | 0.253323 | 0.338980 | 0.397811 | 0.980234 | 1.116106 | 1.372528 | 1.448010 | 0.613455 | 0.293576 | 0.243237 | 0.214643 | 0.216391 | 0.215173 | 0.225080 | 0.217381 | 0.214639 | 0.224022 | 0.271906 | 0.354116 | 0.375995 | 0.308949 | 0.345731 | 0.427002 | 0.340496 | 0.265825 | 0.226144 | 0.220303 | 0.221611 | 0.351045 | 0.736283 | 1.623022 | 4.916003 | 9.826315 | 14.713355 | 7.044232 | 1.320202 | 0.644219 | 0.360123 | 0.322608 | 0.286037 | 0.327521 | 0.447598 | 0.945298 | 1.043060 | 0.736787 | 0.454947 | 0.386257 | 0.269266 | 0.241136 | 0.244917 | 0.238158 | 0.244141 | 0.224110 | 0.222922 | 0.214657 | 0.255015 | 0.524375 | 1.414149 | 3.640220 | 5.464135 | 13.116723 | 9.954847 | 2.863070 | 0.788212 | 0.342450 | 0.216462 | 0.270238 | 0.578538 | 0.939387 | 1.181358 | 0.937816 | 1.220773 | 1.093557 | 1.095519 | 0.833287 | 0.577477 | 0.371950 | 0.289786 | 0.215279 | 0.277960 | 0.777391 | 1.697168 | 2.283441 | 3.510951 | 3.248981 | 4.318587 | 3.276160 | 2.010115 | 1.663108 | 1.280414 | 1.079873 | 0.860855 | 0.641078 | 0.425588 | 0.260882 | 0.214826 | 0.247971 | 0.306356 | 0.413266 | 0.619976 | 0.838115 | 0.733528 | 0.524922 | 0.323989 | 0.225812 | 0.237495 | 0.478586 | 2.160939 | 13.007491 | 19.852311 | 14.535753 | 2.794603 | 1.240973 | 0.680838 | 0.493450 | 0.375328 | 0.366325 | 0.370914 | 0.558456 | 0.663874 | 0.837775 | 0.522601 | 0.349454 | 0.295669 | 0.263622 | 0.247551 | 0.224631 | 0.215200 | 0.214636 | 0.215552 | 0.218738 | 0.222875 | 0.222574 | 0.239844 | 0.260212 | 0.289287 | 0.257094 | 0.232929 | 0.227051 | 0.220091 | 0.214999 | 0.214757 | 0.221214 | 0.229015 | 0.253151 | 0.326436 | 0.400323 | 0.558516 | 0.617781 | 0.549926 | 0.526799 | 0.450550 | 0.351933 | 0.305415 | 0.288516 | 0.286234 | 0.286172 | 0.282157 | 0.282818 | 0.280331 | 0.293945 | 0.285610 | 0.301356 | 0.295557 | 0.294023 | 0.293731 | 0.314513 | 0.304151 | 0.313081 | 0.311096 | 0.299533 | 0.282390 | 0.281654 | 0.287303 | 0.308676 | 0.325236 | 0.337430 | 0.353225 | 0.362339 | 0.368978 | 0.355169 | 0.356900 | 0.363830 | 0.319717 | 0.297740 | 0.294701 | 0.316576 | 0.351512 | 0.395852 | 0.460054 | 0.609424 | 0.610846 | 0.645761 | 0.676248 | 0.736357 | 0.753606 | 0.622430 | 0.631326 | 0.907463 | 0.866183 | 0.668241 | 0.427097 | 0.359306 | 0.325627 | 0.295387 | 0.257768 | 0.230525 | 0.227850 | 0.233733 | 0.239482 | 0.249192 | 0.249981 | 0.269940 | 0.295690 | 0.320985 | 0.352410 | 0.320102 | 0.276812 | 0.260663 | 0.247568 | 0.240308 | 0.226346 | 0.224140 | 0.236409 | 0.253812 | 0.283480 | 0.298060 | 0.301233 | 0.309129 | 0.278685 | 0.241657 | 0.225550 | 0.215875 | 0.214657 | 0.215732 | 0.218174 | 0.219617 | 0.221864 | 0.231958 | 0.228147 | 0.225732 | 0.216974 | 0.216576 | 0.220477 | 0.222184 | 0.217110 | 0.214893 | 0.215442 | 0.215218 | 0.216603 | 0.227121 | 0.229340 | 0.223516 | 0.214675 | 0.221262 | 0.232022 | 0.239940 | 0.261026 | 0.279930 | 0.309984 | 0.278786 | 0.247827 | 0.232692 | 0.236582 | 0.228103 | 0.223873 | 0.223391 | 0.232798 | 0.244821 | 0.256527 | 0.241697 | 0.251666 |
| right central | 0.344059 | 0.940273 | 1.112361 | 0.632641 | 0.379127 | 0.278806 | 0.218886 | 0.215390 | 0.271392 | 0.333221 | 0.303066 | 0.228377 | 0.216107 | 0.216470 | 0.216707 | 0.239667 | 0.268350 | 0.259055 | 0.347290 | 0.453449 | 0.292369 | 0.231591 | 0.217635 | 0.224626 | 0.281187 | 0.458827 | 0.519265 | 0.475280 | 0.410327 | 0.667029 | 0.602720 | 0.339816 | 0.346812 | 0.473666 | 0.672951 | 0.952181 | 1.092081 | 1.807537 | 2.173747 | 2.497649 | 1.979829 | 2.345704 | 2.716369 | 2.830745 | 2.132303 | 1.883372 | 0.690332 | 0.477339 | 0.335311 | 0.247383 | 0.223122 | 0.253503 | 0.268720 | 0.474925 | 0.486314 | 0.421782 | 0.371186 | 0.351520 | 0.295720 | 0.290753 | 0.254053 | 0.230958 | 0.219817 | 0.215959 | 0.286122 | 0.890047 | 4.423085 | 15.838125 | 53.058752 | 78.785304 | 9.901125 | 1.495334 | 0.441250 | 0.252998 | 0.216131 | 0.243161 | 0.299463 | 0.254428 | 0.226179 | 0.225429 | 0.241767 | 0.237074 | 0.241659 | 0.231951 | 0.248108 | 0.263786 | 0.255942 | 0.226815 | 0.216466 | 0.216103 | 0.226721 | 0.258113 | 0.301076 | 0.293027 | 0.286442 | 0.282279 | 0.241189 | 0.219615 | 0.215553 | 0.216861 | 0.215208 | 0.214703 | 0.214687 | 0.215402 | 0.214646 | 0.217193 | 0.217583 | 0.233327 | 0.266445 | 0.241624 | 0.215945 | 0.214805 | 0.227342 | 0.236155 | 0.259772 | 0.321790 | 0.267096 | 0.229247 | 0.227615 | 0.235452 | 0.258516 | 0.364918 | 0.481916 | 0.554848 | 0.777546 | 0.675626 | 0.448629 | 0.328252 | 0.216600 | 0.239802 | 0.255415 | 0.312729 | 0.403186 | 0.461262 | 0.481661 | 0.381489 | 0.329097 | 0.358553 | 0.335296 | 0.299248 | 0.273353 | 0.271188 | 0.265879 | 0.251465 | 0.234613 | 0.220151 | 0.218980 | 0.214670 | 0.215376 | 0.214820 | 0.214928 | 0.222144 | 0.256593 | 0.259055 | 0.249134 | 0.246012 | 0.228519 | 0.243682 | 0.241635 | 0.224376 | 0.215694 | 0.224804 | 0.228269 | 0.257122 | 0.262816 | 0.269640 | 0.283200 | 0.327566 | 0.274345 | 0.261172 | 0.236219 | 0.231529 | 0.237623 | 0.247357 | 0.250833 | 0.261029 | 0.241237 | 0.237144 | 0.239802 | 0.226343 | 0.218588 | 0.216418 | 0.219688 | 0.232113 | 0.261340 | 0.275026 | 0.292932 | 0.354799 | 0.355369 | 0.384936 | 0.458963 | 0.373999 | 0.325693 | 0.312905 | 0.265199 | 0.301631 | 0.356379 | 0.381221 | 0.437015 | 0.571551 | 0.668725 | 0.731166 | 0.728221 | 0.576075 | 0.511243 | 0.431098 | 0.309335 | 0.255254 | 0.251124 | 0.243995 | 0.236657 | 0.224819 | 0.223924 | 0.252634 | 0.307958 | 0.299820 | 0.280944 | 0.241984 | 0.248875 | 0.287854 | 0.334472 | 0.415837 | 0.420616 | 0.516239 | 0.803666 | 0.755356 | 0.589962 | 0.435016 | 0.341750 | 0.369682 | 0.424886 | 0.471863 | 0.585694 | 0.586945 | 0.706111 | 0.697596 | 0.570891 | 0.439076 | 0.345886 | 0.295671 | 0.308247 | 0.276063 | 0.270957 | 0.293450 | 0.323181 | 0.497211 | 0.838954 | 1.027850 | 1.065950 | 0.982941 | 0.730798 | 0.542135 | 0.421415 | 0.362284 | 0.388859 | 0.476133 | 0.426241 | 0.399828 | 0.368715 | 0.306685 | 0.262016 | 0.234512 | 0.226607 | 0.238251 | 0.249343 | 0.249615 | 0.238817 | 0.225926 | 0.225367 | 0.232549 | 0.234354 | 0.235619 | 0.268690 | 0.354688 | 0.410874 | 0.333617 | 0.256401 | 0.238238 | 0.223539 | 0.214646 |
| left posterior | 0.272924 | 0.290434 | 0.357836 | 0.306531 | 0.259015 | 0.229575 | 0.222294 | 0.223946 | 0.232468 | 0.251237 | 0.428385 | 1.498235 | 3.288892 | 4.385808 | 2.497378 | 1.300554 | 1.178913 | 0.434309 | 0.255223 | 0.265114 | 0.271347 | 0.590315 | 2.433172 | 1.344319 | 1.740397 | 1.338738 | 0.729387 | 0.419635 | 0.249552 | 0.218801 | 0.237083 | 0.279122 | 0.257613 | 0.246797 | 0.230218 | 0.228761 | 0.217735 | 0.259930 | 0.285171 | 0.236937 | 0.248227 | 0.238141 | 0.233350 | 0.214996 | 0.234309 | 0.227470 | 0.215664 | 0.218847 | 0.225194 | 0.274914 | 0.424426 | 0.740571 | 1.137744 | 1.634474 | 3.373282 | 5.915160 | 1.305635 | 0.682815 | 0.463826 | 0.271006 | 0.232580 | 1.080853 | 28.663654 | 472.785517 | 3373.290174 | 22512.441193 | 42169.986923 | 45287.183594 | 6735.206882 | 955.895407 | 171.646507 | 27.656421 | 4.382688 | 0.771087 | 0.281664 | 0.216214 | 0.337026 | 1.329225 | 5.734861 | 22.097269 | 28.433934 | 18.211985 | 10.427676 | 4.801040 | 2.083580 | 0.931608 | 0.497486 | 0.317655 | 0.216045 | 0.554500 | 1.838345 | 5.075998 | 11.978525 | 16.174534 | 29.286310 | 33.323489 | 25.688831 | 32.962757 | 65.496251 | 201.650137 | 482.306816 | 153.562025 | 11.957116 | 1.601377 | 0.514259 | 0.295466 | 0.230237 | 0.235149 | 0.300583 | 0.338509 | 0.366611 | 0.340856 | 0.321516 | 0.363715 | 0.346979 | 0.297819 | 0.270702 | 0.250492 | 0.235044 | 0.240031 | 0.229932 | 0.229715 | 0.223219 | 0.215843 | 0.214678 | 0.214632 | 0.214717 | 0.220416 | 0.232726 | 0.255179 | 0.384847 | 0.563208 | 0.934689 | 0.911895 | 0.883130 | 0.910300 | 0.973094 | 0.736568 | 0.725839 | 0.796287 | 1.250784 | 1.383595 | 1.498690 | 1.548516 | 3.044605 | 5.438238 | 3.414432 | 1.640903 | 1.311518 | 0.927227 | 1.104915 | 0.759352 | 0.649981 | 0.502568 | 0.409887 | 0.361786 | 0.371599 | 0.309210 | 0.280050 | 0.240354 | 0.240752 | 0.237496 | 0.245692 | 0.230468 | 0.228704 | 0.224862 | 0.223381 | 0.220053 | 0.222861 | 0.224355 | 0.241409 | 0.250364 | 0.262265 | 0.301571 | 0.385858 | 0.412339 | 0.417689 | 0.459623 | 0.522678 | 0.709103 | 1.094463 | 1.187489 | 1.678435 | 1.743408 | 1.743485 | 2.637882 | 3.971958 | 3.752323 | 3.063270 | 2.412691 | 2.893928 | 2.803301 | 2.505332 | 2.243750 | 2.066872 | 2.205546 | 2.351427 | 1.848150 | 2.157531 | 2.098678 | 1.933879 | 2.089444 | 2.757397 | 3.177537 | 4.480562 | 5.238618 | 7.743118 | 10.267664 | 10.541963 | 7.395861 | 6.507753 | 5.272336 | 4.374295 | 4.249260 | 5.002699 | 5.899130 | 6.792814 | 5.911375 | 5.629165 | 4.396131 | 3.377871 | 2.463171 | 1.623222 | 1.204440 | 1.215335 | 1.336113 | 1.618682 | 3.166448 | 6.654861 | 15.845258 | 13.301775 | 8.926518 | 9.025781 | 9.876665 | 7.932263 | 7.420288 | 7.156169 | 9.418653 | 12.097872 | 10.421492 | 7.878032 | 6.516827 | 4.689439 | 3.560895 | 3.235371 | 3.319273 | 4.739109 | 5.922314 | 9.577839 | 13.044233 | 20.697001 | 22.876650 | 8.254130 | 3.033760 | 1.804120 | 1.587450 | 1.560422 | 1.405990 | 1.577962 | 2.793544 | 4.063735 | 8.249016 | 8.860633 | 12.208843 | 15.168980 | 16.978693 | 9.318597 | 4.447178 | 2.244587 | 1.317275 | 0.941828 | 0.760297 | 0.662455 | 0.816135 | 1.288322 | 1.786802 | 2.552050 | 2.304863 | 2.515017 | 3.192650 |
| right posterior | 0.336786 | 0.266517 | 0.245494 | 0.239483 | 0.237533 | 0.221476 | 0.217752 | 0.216055 | 0.214645 | 0.227574 | 0.312899 | 0.361660 | 0.398258 | 0.537496 | 0.334581 | 0.270253 | 0.219545 | 0.237253 | 0.349235 | 0.413606 | 0.322797 | 0.231230 | 0.220338 | 0.233868 | 0.241176 | 0.240722 | 0.280917 | 0.283842 | 0.279397 | 0.231101 | 0.217497 | 0.215323 | 0.215671 | 0.218721 | 0.260870 | 0.347446 | 0.451214 | 0.490277 | 0.538562 | 0.638320 | 0.545177 | 0.404380 | 0.545604 | 0.565891 | 0.668571 | 0.535950 | 0.304761 | 0.230491 | 0.214980 | 0.314777 | 0.585271 | 1.171981 | 1.642940 | 1.206651 | 0.604823 | 0.385082 | 0.231639 | 0.216159 | 0.277203 | 0.641733 | 6.325318 | 33.265441 | 193.231408 | 526.177310 | 2513.519469 | 8548.909931 | 28831.759130 | 40348.847587 | 17973.494635 | 6291.761892 | 912.961190 | 125.652389 | 19.081734 | 5.155210 | 2.420459 | 1.289774 | 0.645224 | 0.313538 | 0.237535 | 0.236889 | 0.244058 | 0.223793 | 0.214989 | 0.219049 | 0.214714 | 0.215780 | 0.217698 | 0.214640 | 0.215091 | 0.227792 | 0.349627 | 0.636490 | 1.112479 | 2.030615 | 5.681720 | 13.029436 | 14.409945 | 7.617597 | 4.637063 | 3.569631 | 2.180925 | 1.229393 | 0.551163 | 0.331507 | 0.291508 | 0.250260 | 0.228004 | 0.221289 | 0.218246 | 0.215080 | 0.215075 | 0.216937 | 0.220600 | 0.221535 | 0.221083 | 0.217693 | 0.214646 | 0.225746 | 0.276249 | 0.362490 | 0.454777 | 0.393929 | 0.317359 | 0.266017 | 0.227993 | 0.214631 | 0.217417 | 0.222746 | 0.215813 | 0.216407 | 0.229008 | 0.249889 | 0.283332 | 0.295607 | 0.299867 | 0.302943 | 0.310804 | 0.321493 | 0.355478 | 0.356913 | 0.389607 | 0.456492 | 0.482534 | 0.568623 | 0.547332 | 0.467971 | 0.389527 | 0.311881 | 0.249466 | 0.233701 | 0.215934 | 0.214631 | 0.214809 | 0.217789 | 0.227487 | 0.258047 | 0.284557 | 0.350363 | 0.356275 | 0.322533 | 0.299375 | 0.265917 | 0.246738 | 0.240818 | 0.248724 | 0.263905 | 0.254901 | 0.243421 | 0.235126 | 0.220899 | 0.219651 | 0.215180 | 0.215566 | 0.216595 | 0.214637 | 0.220713 | 0.249197 | 0.304493 | 0.386329 | 0.390560 | 0.382558 | 0.359802 | 0.388895 | 0.434394 | 0.430947 | 0.419126 | 0.479222 | 0.650239 | 0.499571 | 0.428330 | 0.341314 | 0.303072 | 0.267736 | 0.252723 | 0.230885 | 0.238593 | 0.224939 | 0.219976 | 0.214766 | 0.216333 | 0.228286 | 0.236319 | 0.234550 | 0.225562 | 0.232162 | 0.234167 | 0.248466 | 0.245845 | 0.254962 | 0.275594 | 0.301643 | 0.262695 | 0.225673 | 0.215173 | 0.232506 | 0.289220 | 0.377608 | 0.571242 | 0.800037 | 0.752521 | 0.602569 | 0.468251 | 0.384371 | 0.326133 | 0.267442 | 0.231809 | 0.220144 | 0.214987 | 0.215372 | 0.218363 | 0.222676 | 0.227291 | 0.232372 | 0.236253 | 0.237150 | 0.250324 | 0.267966 | 0.273206 | 0.287768 | 0.275298 | 0.264332 | 0.264356 | 0.234691 | 0.237261 | 0.253475 | 0.265729 | 0.290499 | 0.337559 | 0.350458 | 0.414872 | 0.380767 | 0.302259 | 0.273215 | 0.271874 | 0.270556 | 0.294398 | 0.316797 | 0.331491 | 0.356992 | 0.361761 | 0.388563 | 0.466103 | 0.602306 | 0.815920 | 1.295198 | 2.241582 | 2.246600 | 1.363041 | 0.789248 | 0.494743 | 0.413703 | 0.375177 | 0.320633 | 0.345435 | 0.460606 | 0.489136 | 0.506335 | 0.457640 | 0.409805 | 0.426148 |
| all electrodes | 1.570175 | 1.466972 | 1.809756 | 0.934883 | 0.512467 | 0.305114 | 0.246877 | 0.235374 | 0.268538 | 0.263933 | 0.405974 | 0.418324 | 0.397307 | 0.552070 | 0.457875 | 0.280942 | 0.220620 | 0.255194 | 0.272786 | 0.295352 | 0.423640 | 0.342842 | 0.232433 | 0.248105 | 0.533939 | 0.765894 | 0.720699 | 1.016370 | 1.101844 | 0.640573 | 0.269097 | 0.217322 | 0.216313 | 0.221667 | 0.270722 | 0.424798 | 0.554959 | 0.371231 | 0.312840 | 0.297936 | 0.306623 | 0.234302 | 0.215801 | 0.267431 | 0.239956 | 0.245352 | 0.330621 | 0.445776 | 0.373597 | 0.234362 | 0.224105 | 0.219112 | 0.216573 | 0.224702 | 0.228140 | 0.227445 | 0.216927 | 0.216100 | 0.231242 | 0.287682 | 0.607373 | 1.882209 | 18.378781 | 287.381800 | 1183.366779 | 3313.321001 | 3121.852635 | 1984.163090 | 1011.564585 | 266.472811 | 59.133459 | 12.553108 | 1.402309 | 0.352364 | 0.215274 | 0.254581 | 0.534969 | 0.652246 | 0.475511 | 0.279597 | 0.239916 | 0.218666 | 0.218266 | 0.217332 | 0.222586 | 0.231237 | 0.235091 | 0.230475 | 0.227759 | 0.216303 | 0.220568 | 0.254445 | 0.253509 | 0.243103 | 0.257277 | 0.254274 | 0.252566 | 0.238225 | 0.227471 | 0.220970 | 0.214772 | 0.238084 | 0.240193 | 0.269557 | 0.278550 | 0.247744 | 0.220402 | 0.214881 | 0.227871 | 0.239694 | 0.286472 | 0.380237 | 0.322080 | 0.275424 | 0.316540 | 0.455717 | 0.523049 | 0.537073 | 0.465797 | 0.496917 | 0.556076 | 0.461141 | 0.356672 | 0.304856 | 0.262803 | 0.253798 | 0.246448 | 0.239054 | 0.231040 | 0.224949 | 0.235508 | 0.223839 | 0.216460 | 0.217633 | 0.221740 | 0.223129 | 0.225978 | 0.235071 | 0.236379 | 0.219383 | 0.222260 | 0.222988 | 0.218178 | 0.217176 | 0.215581 | 0.220672 | 0.232923 | 0.219223 | 0.219610 | 0.219291 | 0.225592 | 0.227234 | 0.219607 | 0.214633 | 0.216563 | 0.217663 | 0.215381 | 0.216085 | 0.221593 | 0.227053 | 0.224855 | 0.221049 | 0.226089 | 0.238726 | 0.236751 | 0.263215 | 0.296860 | 0.333131 | 0.368172 | 0.361421 | 0.294439 | 0.313921 | 0.289203 | 0.242671 | 0.226142 | 0.217094 | 0.223779 | 0.270335 | 0.263649 | 0.251846 | 0.282075 | 0.262957 | 0.254416 | 0.224324 | 0.214656 | 0.215327 | 0.214947 | 0.214724 | 0.217288 | 0.222258 | 0.222613 | 0.222314 | 0.215584 | 0.217655 | 0.222076 | 0.224941 | 0.229964 | 0.234985 | 0.228215 | 0.235742 | 0.224477 | 0.224075 | 0.224558 | 0.224079 | 0.219412 | 0.216873 | 0.219912 | 0.226950 | 0.221220 | 0.214756 | 0.216552 | 0.215811 | 0.216260 | 0.236210 | 0.260597 | 0.280791 | 0.253819 | 0.238633 | 0.236351 | 0.218514 | 0.220761 | 0.228788 | 0.236076 | 0.223154 | 0.223975 | 0.220677 | 0.215404 | 0.216618 | 0.215462 | 0.215924 | 0.214912 | 0.214700 | 0.214660 | 0.218707 | 0.215076 | 0.215332 | 0.220312 | 0.221798 | 0.216008 | 0.220461 | 0.219688 | 0.214773 | 0.225309 | 0.235802 | 0.243442 | 0.235149 | 0.242095 | 0.223675 | 0.218112 | 0.219337 | 0.227455 | 0.225051 | 0.227959 | 0.238940 | 0.239727 | 0.231772 | 0.227600 | 0.229622 | 0.231641 | 0.241367 | 0.260598 | 0.314762 | 0.392032 | 0.421416 | 0.355468 | 0.346752 | 0.275659 | 0.226261 | 0.216978 | 0.216205 | 0.216228 | 0.214992 | 0.216584 | 0.220198 | 0.229728 | 0.268170 | 0.307725 | 0.323896 | 0.340967 | 0.374095 |

Searchlight, spatiotemporal cluster permutation test

|  | start time | stop time | peak time | peak channel | cluster p | peak Cohen's d | direction |
| --- | --- | --- | --- | --- | --- | --- | --- |
| #1 | 100 | 180 | 135 | POz | 0.0141 | 1.190415 | positive |
| #2 | 230 | 350 | 285 | P10 | 0.0096 | 1.05139 | positive |

Z) emoji to real faces - angry vs sad

  
|  | time window | peak latency | cluster *p* | peak Cohen's *d* |  | | | |
| **all electrodes** |  | | | |  | | | |
|  | | | | | | | | |

Time-resolved classification, cluster permutation tests

|  | **left hemisphere** | | | | **right hemisphere** | | | |
|  | time window | peak latency | cluster *p* | peak Cohen's *d* | time window | peak latency | cluster *p* | peak Cohen's *d* |
| **anterior** |  | | | |  | | | |
| **central** |  | | | | 400 - 570 ms | 445 ms | 0.0079 | 0.8726 |
| **posterior** |  | | | |  | | | |

  

Time-resolved classification, Bayesian statistics

|  | -200 | -195 | -190 | -185 | -180 | -175 | -170 | -165 | -160 | -155 | -150 | -145 | -140 | -135 | -130 | -125 | -120 | -115 | -110 | -105 | -100 | -95 | -90 | -85 | -80 | -75 | -70 | -65 | -60 | -55 | -50 | -45 | -40 | -35 | -30 | -25 | -20 | -15 | -10 | -5 | 0 | 5 | 10 | 15 | 20 | 25 | 30 | 35 | 40 | 45 | 50 | 55 | 60 | 65 | 70 | 75 | 80 | 85 | 90 | 95 | 100 | 105 | 110 | 115 | 120 | 125 | 130 | 135 | 140 | 145 | 150 | 155 | 160 | 165 | 170 | 175 | 180 | 185 | 190 | 195 | 200 | 205 | 210 | 215 | 220 | 225 | 230 | 235 | 240 | 245 | 250 | 255 | 260 | 265 | 270 | 275 | 280 | 285 | 290 | 295 | 300 | 305 | 310 | 315 | 320 | 325 | 330 | 335 | 340 | 345 | 350 | 355 | 360 | 365 | 370 | 375 | 380 | 385 | 390 | 395 | 400 | 405 | 410 | 415 | 420 | 425 | 430 | 435 | 440 | 445 | 450 | 455 | 460 | 465 | 470 | 475 | 480 | 485 | 490 | 495 | 500 | 505 | 510 | 515 | 520 | 525 | 530 | 535 | 540 | 545 | 550 | 555 | 560 | 565 | 570 | 575 | 580 | 585 | 590 | 595 | 600 | 605 | 610 | 615 | 620 | 625 | 630 | 635 | 640 | 645 | 650 | 655 | 660 | 665 | 670 | 675 | 680 | 685 | 690 | 695 | 700 | 705 | 710 | 715 | 720 | 725 | 730 | 735 | 740 | 745 | 750 | 755 | 760 | 765 | 770 | 775 | 780 | 785 | 790 | 795 | 800 | 805 | 810 | 815 | 820 | 825 | 830 | 835 | 840 | 845 | 850 | 855 | 860 | 865 | 870 | 875 | 880 | 885 | 890 | 895 | 900 | 905 | 910 | 915 | 920 | 925 | 930 | 935 | 940 | 945 | 950 | 955 | 960 | 965 | 970 | 975 | 980 | 985 | 990 | 995 | 1000 | 1005 | 1010 | 1015 | 1020 | 1025 | 1030 | 1035 | 1040 | 1045 | 1050 | 1055 | 1060 | 1065 | 1070 | 1075 | 1080 | 1085 | 1090 | 1095 | 1100 | 1105 | 1110 | 1115 | 1120 | 1125 | 1130 | 1135 | 1140 | 1145 | 1150 | 1155 | 1160 | 1165 | 1170 | 1175 | 1180 | 1185 | 1190 | 1195 |
| --- | --- | --- | --- | --- | --- | --- | --- | --- | --- | --- | --- | --- | --- | --- | --- | --- | --- | --- | --- | --- | --- | --- | --- | --- | --- | --- | --- | --- | --- | --- | --- | --- | --- | --- | --- | --- | --- | --- | --- | --- | --- | --- | --- | --- | --- | --- | --- | --- | --- | --- | --- | --- | --- | --- | --- | --- | --- | --- | --- | --- | --- | --- | --- | --- | --- | --- | --- | --- | --- | --- | --- | --- | --- | --- | --- | --- | --- | --- | --- | --- | --- | --- | --- | --- | --- | --- | --- | --- | --- | --- | --- | --- | --- | --- | --- | --- | --- | --- | --- | --- | --- | --- | --- | --- | --- | --- | --- | --- | --- | --- | --- | --- | --- | --- | --- | --- | --- | --- | --- | --- | --- | --- | --- | --- | --- | --- | --- | --- | --- | --- | --- | --- | --- | --- | --- | --- | --- | --- | --- | --- | --- | --- | --- | --- | --- | --- | --- | --- | --- | --- | --- | --- | --- | --- | --- | --- | --- | --- | --- | --- | --- | --- | --- | --- | --- | --- | --- | --- | --- | --- | --- | --- | --- | --- | --- | --- | --- | --- | --- | --- | --- | --- | --- | --- | --- | --- | --- | --- | --- | --- | --- | --- | --- | --- | --- | --- | --- | --- | --- | --- | --- | --- | --- | --- | --- | --- | --- | --- | --- | --- | --- | --- | --- | --- | --- | --- | --- | --- | --- | --- | --- | --- | --- | --- | --- | --- | --- | --- | --- | --- | --- | --- | --- | --- | --- | --- | --- | --- | --- | --- | --- | --- | --- | --- | --- | --- | --- | --- | --- | --- | --- | --- | --- | --- | --- | --- | --- | --- | --- | --- | --- | --- | --- | --- | --- | --- | --- | --- | --- | --- | --- | --- | --- | --- | --- | --- | --- | --- | --- | --- |
| left anterior | 1.355478 | 1.143311 | 0.651938 | 0.576258 | 0.723717 | 0.480956 | 0.239314 | 0.222293 | 0.378059 | 0.354342 | 0.544547 | 0.846315 | 0.974924 | 0.616115 | 0.291668 | 0.238754 | 0.558603 | 2.036973 | 2.330805 | 0.760902 | 0.712999 | 0.500825 | 0.311771 | 0.216038 | 0.258544 | 0.324276 | 0.300093 | 0.360024 | 0.353687 | 0.268574 | 0.246147 | 0.241009 | 0.216086 | 0.224932 | 0.281892 | 0.264297 | 0.280445 | 0.319241 | 0.600623 | 1.114992 | 1.405458 | 1.050565 | 0.540958 | 0.277308 | 0.249706 | 0.221445 | 0.214804 | 0.218338 | 0.219159 | 0.214961 | 0.214635 | 0.216104 | 0.214626 | 0.235477 | 0.238908 | 0.250367 | 0.242250 | 0.229271 | 0.245194 | 0.235347 | 0.218380 | 0.216974 | 0.214947 | 0.217238 | 0.235769 | 0.256959 | 0.270855 | 0.248548 | 0.226804 | 0.224438 | 0.221501 | 0.220090 | 0.218088 | 0.227020 | 0.224827 | 0.235763 | 0.214626 | 0.215004 | 0.215070 | 0.215564 | 0.218792 | 0.235347 | 0.258948 | 0.344616 | 0.394535 | 0.396071 | 0.354196 | 0.309640 | 0.299904 | 0.258346 | 0.279684 | 0.297593 | 0.316820 | 0.317758 | 0.265888 | 0.234112 | 0.237199 | 0.226673 | 0.230026 | 0.249953 | 0.262905 | 0.273336 | 0.291947 | 0.306126 | 0.371714 | 0.399002 | 0.450653 | 0.599486 | 0.874496 | 1.462650 | 1.373133 | 1.659341 | 1.149097 | 0.643557 | 0.468235 | 0.348500 | 0.269647 | 0.271697 | 0.248131 | 0.251757 | 0.254085 | 0.232364 | 0.215346 | 0.214807 | 0.225820 | 0.221528 | 0.215112 | 0.227847 | 0.263076 | 0.357481 | 0.508101 | 0.649904 | 0.523303 | 0.457020 | 0.416812 | 0.440234 | 0.529041 | 0.475478 | 0.437960 | 0.405984 | 0.506195 | 0.502796 | 0.417085 | 0.381184 | 0.368094 | 0.400250 | 0.402637 | 0.307656 | 0.292025 | 0.271639 | 0.279741 | 0.334946 | 0.362452 | 0.405265 | 0.411248 | 0.392445 | 0.405972 | 0.369897 | 0.346275 | 0.375549 | 0.338955 | 0.340762 | 0.255844 | 0.227356 | 0.218124 | 0.216323 | 0.214905 | 0.214817 | 0.215548 | 0.214902 | 0.216226 | 0.233421 | 0.235494 | 0.230347 | 0.248660 | 0.295809 | 0.292879 | 0.273681 | 0.234854 | 0.225671 | 0.216794 | 0.214881 | 0.226931 | 0.223081 | 0.215629 | 0.214661 | 0.214635 | 0.215736 | 0.214863 | 0.222549 | 0.243118 | 0.264150 | 0.244759 | 0.239255 | 0.220682 | 0.220132 | 0.298922 | 0.409816 | 0.568815 | 0.499156 | 0.394468 | 0.392657 | 0.347266 | 0.328947 | 0.353142 | 0.313431 | 0.323028 | 0.444787 | 0.535912 | 0.622940 | 0.637768 | 0.563877 | 0.431486 | 0.422232 | 0.389787 | 0.326249 | 0.283857 | 0.250752 | 0.235893 | 0.265638 | 0.278166 | 0.248406 | 0.226365 | 0.225130 | 0.223240 | 0.219907 | 0.216731 | 0.214737 | 0.216598 | 0.226296 | 0.218697 | 0.219212 | 0.224291 | 0.227689 | 0.230336 | 0.221992 | 0.222794 | 0.275817 | 0.335693 | 0.373392 | 0.331498 | 0.295810 | 0.307093 | 0.255626 | 0.218853 | 0.215713 | 0.215241 | 0.214700 | 0.216623 | 0.227358 | 0.258545 | 0.289965 | 0.271819 | 0.246150 | 0.238079 | 0.221935 | 0.214973 | 0.215252 | 0.215271 | 0.215426 | 0.215381 | 0.221749 | 0.249320 | 0.314202 | 0.331070 | 0.352260 | 0.399206 | 0.445542 | 0.489509 | 0.427646 | 0.335960 | 0.300646 | 0.272786 | 0.216106 | 0.214863 | 0.215435 | 0.217254 | 0.220898 | 0.228309 | 0.250047 |
| right anterior | 0.388063 | 0.530683 | 0.637209 | 0.389852 | 0.353882 | 0.228096 | 0.217087 | 0.224281 | 0.268408 | 0.426938 | 0.420471 | 0.322421 | 0.294497 | 0.240626 | 0.233648 | 0.232910 | 0.226250 | 0.259781 | 0.265185 | 0.303070 | 0.347922 | 0.323594 | 0.346759 | 0.469203 | 0.618126 | 0.494940 | 0.561075 | 0.359955 | 0.612779 | 1.750551 | 0.614544 | 0.326156 | 0.248451 | 0.286191 | 0.466583 | 0.459576 | 0.364789 | 0.695444 | 1.202182 | 0.657944 | 0.310875 | 0.247258 | 0.235187 | 0.245410 | 0.307136 | 0.412411 | 0.733007 | 1.015811 | 0.939958 | 0.795765 | 0.658698 | 0.386231 | 0.276023 | 0.288520 | 0.376864 | 0.773533 | 2.561267 | 6.797907 | 10.899060 | 10.663867 | 5.559534 | 2.572936 | 0.707211 | 0.416911 | 0.264340 | 0.256346 | 0.327937 | 0.350111 | 0.347167 | 0.347350 | 0.279521 | 0.275837 | 0.254841 | 0.219250 | 0.215642 | 0.215678 | 0.232111 | 0.233401 | 0.218784 | 0.214810 | 0.216974 | 0.215096 | 0.215093 | 0.215735 | 0.227966 | 0.341707 | 0.549801 | 0.624701 | 0.669340 | 0.385465 | 0.249487 | 0.217639 | 0.219509 | 0.218299 | 0.214668 | 0.222856 | 0.302997 | 0.375657 | 0.666120 | 1.193187 | 0.922500 | 0.447975 | 0.339819 | 0.311784 | 0.369709 | 0.342772 | 0.349801 | 0.357690 | 0.380397 | 0.363163 | 0.343989 | 0.296127 | 0.272322 | 0.246873 | 0.273988 | 0.294887 | 0.282275 | 0.269816 | 0.260774 | 0.246511 | 0.260747 | 0.244022 | 0.229384 | 0.226506 | 0.230656 | 0.240560 | 0.251273 | 0.268764 | 0.287707 | 0.354648 | 0.421563 | 0.375362 | 0.440514 | 0.502331 | 0.443661 | 0.450939 | 0.326516 | 0.232454 | 0.219930 | 0.217467 | 0.251047 | 0.296865 | 0.350977 | 0.352053 | 0.330471 | 0.327961 | 0.309124 | 0.296235 | 0.314969 | 0.334659 | 0.330154 | 0.275915 | 0.250262 | 0.272900 | 0.288089 | 0.310813 | 0.374161 | 0.408703 | 0.442055 | 0.477551 | 0.363551 | 0.319271 | 0.330308 | 0.321282 | 0.344610 | 0.390142 | 0.346667 | 0.335031 | 0.322597 | 0.294620 | 0.286252 | 0.253945 | 0.244296 | 0.262201 | 0.257891 | 0.248494 | 0.240851 | 0.253056 | 0.277788 | 0.307635 | 0.356848 | 0.471336 | 0.673747 | 0.850695 | 0.780113 | 0.752588 | 0.619983 | 0.590890 | 0.436406 | 0.334558 | 0.285927 | 0.264054 | 0.251784 | 0.251637 | 0.236421 | 0.234997 | 0.224844 | 0.220880 | 0.219493 | 0.217045 | 0.216421 | 0.215206 | 0.216500 | 0.215110 | 0.216158 | 0.215469 | 0.215699 | 0.214664 | 0.215080 | 0.217623 | 0.216619 | 0.222868 | 0.256902 | 0.243312 | 0.238646 | 0.273447 | 0.299126 | 0.261231 | 0.253768 | 0.249385 | 0.247306 | 0.235089 | 0.216271 | 0.215818 | 0.220107 | 0.227015 | 0.243913 | 0.263745 | 0.252767 | 0.238488 | 0.231896 | 0.219344 | 0.214966 | 0.219419 | 0.230060 | 0.222803 | 0.215479 | 0.214650 | 0.220428 | 0.238072 | 0.244836 | 0.255417 | 0.246734 | 0.234048 | 0.229474 | 0.224551 | 0.219489 | 0.225623 | 0.235537 | 0.268563 | 0.269253 | 0.253835 | 0.222733 | 0.217051 | 0.215219 | 0.216023 | 0.219723 | 0.218403 | 0.214863 | 0.218757 | 0.237742 | 0.281055 | 0.270242 | 0.255114 | 0.239050 | 0.220232 | 0.215469 | 0.227266 | 0.257096 | 0.292979 | 0.293482 | 0.308340 | 0.267625 | 0.223477 | 0.231428 | 0.227400 | 0.215168 | 0.214805 | 0.215147 | 0.214626 |
| left central | 0.416890 | 0.303037 | 0.235351 | 0.232116 | 0.219330 | 0.223397 | 0.236898 | 0.224672 | 0.221301 | 0.223548 | 0.218212 | 0.216551 | 0.215578 | 0.224209 | 0.215562 | 0.215138 | 0.218018 | 0.224554 | 0.217143 | 0.218548 | 0.255586 | 0.262569 | 0.319092 | 0.255559 | 0.216878 | 0.216729 | 0.214793 | 0.254926 | 0.343628 | 0.320189 | 0.262443 | 0.228441 | 0.230292 | 0.235886 | 0.243444 | 0.246219 | 0.327070 | 0.423862 | 0.791418 | 0.895865 | 0.847697 | 1.148958 | 0.569826 | 0.365524 | 0.235640 | 0.215133 | 0.214782 | 0.215385 | 0.215734 | 0.217681 | 0.218319 | 0.217638 | 0.218066 | 0.214715 | 0.219142 | 0.329665 | 0.408949 | 0.652908 | 1.185847 | 2.088698 | 3.385629 | 3.056567 | 1.130007 | 0.799529 | 0.665798 | 0.714256 | 0.872160 | 1.257397 | 2.599186 | 4.086168 | 2.325121 | 0.819000 | 0.672739 | 0.682686 | 0.434122 | 0.279392 | 0.245602 | 0.302409 | 0.490286 | 0.532462 | 0.349255 | 0.288525 | 0.236130 | 0.215410 | 0.262510 | 0.415511 | 0.874786 | 1.181636 | 0.462555 | 0.224404 | 0.225524 | 0.283829 | 0.335529 | 0.559787 | 1.498663 | 2.228577 | 2.967760 | 3.584188 | 2.123421 | 1.462938 | 0.656900 | 0.310903 | 0.216738 | 0.269180 | 0.454512 | 0.962382 | 1.575520 | 1.600845 | 2.412851 | 3.117576 | 2.875680 | 2.203044 | 1.374115 | 1.371955 | 1.225575 | 1.033160 | 0.903671 | 0.629078 | 0.446019 | 0.351698 | 0.295082 | 0.255122 | 0.220735 | 0.215988 | 0.219983 | 0.247947 | 0.320652 | 0.336999 | 0.329866 | 0.289837 | 0.252381 | 0.226306 | 0.216418 | 0.214634 | 0.215036 | 0.217064 | 0.224361 | 0.241605 | 0.321012 | 0.333585 | 0.334997 | 0.283788 | 0.244025 | 0.226244 | 0.228734 | 0.228748 | 0.266206 | 0.387301 | 0.500298 | 0.556866 | 0.644584 | 0.694308 | 0.668779 | 0.532339 | 0.357018 | 0.318704 | 0.374112 | 0.495233 | 0.443986 | 0.317936 | 0.312206 | 0.482101 | 0.779366 | 0.553040 | 0.307779 | 0.247464 | 0.282821 | 0.280088 | 0.227449 | 0.214964 | 0.216211 | 0.261913 | 0.364686 | 0.364262 | 0.271937 | 0.250582 | 0.254140 | 0.259795 | 0.261732 | 0.246698 | 0.223702 | 0.242301 | 0.246697 | 0.255598 | 0.258594 | 0.255659 | 0.262529 | 0.326050 | 0.332063 | 0.397036 | 0.423996 | 0.406347 | 0.373974 | 0.316703 | 0.290948 | 0.289344 | 0.272932 | 0.255337 | 0.258779 | 0.252466 | 0.258449 | 0.245742 | 0.251495 | 0.272639 | 0.283153 | 0.280886 | 0.279259 | 0.299136 | 0.326791 | 0.299036 | 0.258840 | 0.246118 | 0.253218 | 0.261495 | 0.264363 | 0.282168 | 0.322141 | 0.376257 | 0.362283 | 0.315964 | 0.277283 | 0.247642 | 0.231690 | 0.218393 | 0.217828 | 0.221986 | 0.218599 | 0.217256 | 0.216973 | 0.215330 | 0.215102 | 0.229115 | 0.242126 | 0.233567 | 0.231188 | 0.231592 | 0.219881 | 0.215336 | 0.236404 | 0.241539 | 0.242942 | 0.236566 | 0.230727 | 0.225274 | 0.218535 | 0.214632 | 0.214798 | 0.216758 | 0.218562 | 0.214822 | 0.215398 | 0.214741 | 0.214860 | 0.214839 | 0.215114 | 0.218015 | 0.215461 | 0.214859 | 0.217684 | 0.217748 | 0.222586 | 0.232902 | 0.243102 | 0.259592 | 0.284561 | 0.279354 | 0.282154 | 0.257996 | 0.239640 | 0.230855 | 0.229069 | 0.217747 | 0.215422 | 0.216448 | 0.218301 | 0.216577 | 0.215140 | 0.214731 | 0.214664 | 0.214641 |
| right central | 0.252685 | 0.235454 | 0.266935 | 0.331428 | 0.287434 | 0.244118 | 0.222462 | 0.218331 | 0.223426 | 0.214637 | 0.226939 | 0.265422 | 0.316811 | 0.309182 | 0.319983 | 0.317847 | 0.270828 | 0.237418 | 0.230763 | 0.317241 | 0.332438 | 0.462018 | 0.497971 | 0.388472 | 0.271750 | 0.225740 | 0.215817 | 0.220581 | 0.220849 | 0.221592 | 0.251425 | 0.281932 | 0.355108 | 0.408223 | 0.351166 | 0.231125 | 0.222376 | 0.303767 | 0.337185 | 0.374527 | 0.398541 | 0.353908 | 0.292508 | 0.224455 | 0.215078 | 0.242381 | 0.328766 | 0.303617 | 0.255060 | 0.270901 | 0.233657 | 0.242392 | 0.217904 | 0.224297 | 0.220814 | 0.218400 | 0.224033 | 0.214670 | 0.216343 | 0.214633 | 0.215902 | 0.215300 | 0.227056 | 0.273186 | 0.429685 | 0.450373 | 0.500247 | 0.622853 | 0.627022 | 0.724640 | 0.503485 | 0.330333 | 0.294188 | 0.305323 | 0.308549 | 0.267300 | 0.215081 | 0.222266 | 0.239242 | 0.305396 | 0.616638 | 3.631420 | 25.246549 | 92.924867 | 121.721083 | 63.371651 | 13.952101 | 1.692173 | 0.369148 | 0.215788 | 0.291771 | 0.539056 | 1.235359 | 2.301633 | 5.053086 | 7.986350 | 6.337476 | 3.235151 | 1.752892 | 1.218557 | 1.174079 | 0.764665 | 0.370030 | 0.255626 | 0.302155 | 0.402190 | 0.602716 | 0.544281 | 0.447696 | 0.451585 | 0.957745 | 0.646625 | 0.728611 | 0.574060 | 0.419717 | 0.471491 | 0.623139 | 0.476089 | 0.591032 | 0.762465 | 1.537886 | 4.234885 | 6.031701 | 14.161643 | 43.780909 | 39.185116 | 53.726194 | 72.915655 | 91.240545 | 106.566848 | 73.773447 | 42.355241 | 49.397744 | 80.524643 | 53.478014 | 60.095227 | 57.682922 | 85.259624 | 203.550911 | 197.663007 | 126.191631 | 100.224951 | 94.970265 | 35.794912 | 10.986412 | 5.264309 | 3.811526 | 2.531407 | 2.076115 | 1.636543 | 1.621869 | 1.826518 | 1.735857 | 1.681229 | 1.341185 | 1.090550 | 1.139922 | 1.030329 | 1.489537 | 2.037091 | 2.301324 | 2.499876 | 3.240926 | 3.276280 | 5.234402 | 2.913274 | 1.180229 | 0.557808 | 0.386122 | 0.302922 | 0.282927 | 0.232373 | 0.226510 | 0.231699 | 0.235720 | 0.240657 | 0.246852 | 0.242914 | 0.273406 | 0.286762 | 0.291851 | 0.302142 | 0.238742 | 0.216705 | 0.214673 | 0.218432 | 0.252364 | 0.282796 | 0.262208 | 0.217598 | 0.214784 | 0.214990 | 0.218910 | 0.214718 | 0.225692 | 0.231918 | 0.221039 | 0.216678 | 0.217161 | 0.228714 | 0.243958 | 0.243795 | 0.233892 | 0.223546 | 0.218641 | 0.215262 | 0.216802 | 0.224936 | 0.233901 | 0.224388 | 0.217334 | 0.214691 | 0.218755 | 0.222984 | 0.229517 | 0.229429 | 0.235825 | 0.256469 | 0.283224 | 0.332831 | 0.376321 | 0.396780 | 0.413091 | 0.386262 | 0.353558 | 0.307512 | 0.247337 | 0.236485 | 0.226015 | 0.219391 | 0.215520 | 0.214626 | 0.214633 | 0.226021 | 0.243622 | 0.238322 | 0.237944 | 0.240852 | 0.258490 | 0.268982 | 0.277621 | 0.264496 | 0.273378 | 0.255258 | 0.262814 | 0.234765 | 0.229186 | 0.215976 | 0.215012 | 0.215413 | 0.216855 | 0.215081 | 0.216123 | 0.214626 | 0.215184 | 0.215165 | 0.215196 | 0.215680 | 0.232523 | 0.287206 | 0.377421 | 0.490731 | 0.509276 | 0.410130 | 0.346016 | 0.271249 | 0.235915 | 0.225027 | 0.227544 | 0.236181 | 0.246235 | 0.235173 | 0.224326 | 0.221687 | 0.232666 | 0.236028 | 0.232386 | 0.239223 | 0.255686 | 0.332576 |
| left posterior | 0.214654 | 0.215777 | 0.230123 | 0.225896 | 0.215689 | 0.218297 | 0.223737 | 0.317677 | 0.548859 | 0.924101 | 0.819274 | 0.634526 | 0.470127 | 0.309825 | 0.217145 | 0.227412 | 0.272182 | 0.366980 | 0.734464 | 1.210206 | 0.832217 | 0.596825 | 0.510258 | 0.516655 | 0.658742 | 0.369945 | 0.270605 | 0.270773 | 0.333365 | 0.386798 | 0.375384 | 0.318653 | 0.296764 | 0.284823 | 0.369395 | 0.358178 | 0.268490 | 0.247272 | 0.229758 | 0.239612 | 0.243099 | 0.218710 | 0.218038 | 0.217510 | 0.239791 | 0.285616 | 0.302784 | 0.266534 | 0.234901 | 0.219360 | 0.224656 | 0.260271 | 0.399426 | 0.508887 | 0.581562 | 0.533709 | 0.321190 | 0.247548 | 0.223848 | 0.218616 | 0.234711 | 0.262600 | 0.298925 | 0.300071 | 0.327186 | 0.326677 | 0.315393 | 0.355679 | 0.473584 | 0.522826 | 0.537412 | 0.532324 | 0.429765 | 0.360651 | 0.257307 | 0.219850 | 0.221293 | 0.244317 | 0.278890 | 0.304469 | 0.341093 | 0.330935 | 0.350773 | 0.317701 | 0.377437 | 0.531151 | 0.712433 | 1.255989 | 1.852851 | 2.587195 | 2.099966 | 1.028220 | 0.593935 | 0.446180 | 0.277715 | 0.221970 | 0.215935 | 0.227220 | 0.216525 | 0.232191 | 0.272267 | 0.326156 | 0.370105 | 0.470020 | 0.497757 | 0.444596 | 0.321481 | 0.248223 | 0.238443 | 0.240366 | 0.232440 | 0.229226 | 0.226190 | 0.233713 | 0.248968 | 0.294826 | 0.448824 | 0.698778 | 1.182489 | 2.058072 | 2.353153 | 1.967712 | 1.066089 | 0.503855 | 0.314746 | 0.233906 | 0.215191 | 0.225580 | 0.239920 | 0.260618 | 0.290430 | 0.328953 | 0.353046 | 0.342228 | 0.369653 | 0.360613 | 0.397525 | 0.453391 | 0.464567 | 0.435954 | 0.397376 | 0.364178 | 0.349626 | 0.343324 | 0.328989 | 0.293744 | 0.307058 | 0.320112 | 0.334234 | 0.385687 | 0.372142 | 0.361363 | 0.368062 | 0.367458 | 0.442569 | 0.487645 | 0.431802 | 0.372523 | 0.334547 | 0.327971 | 0.298599 | 0.264897 | 0.229367 | 0.217931 | 0.219407 | 0.223505 | 0.225469 | 0.224824 | 0.222298 | 0.238667 | 0.282356 | 0.293360 | 0.315455 | 0.382078 | 0.455574 | 0.542712 | 0.480631 | 0.367764 | 0.278390 | 0.236408 | 0.214808 | 0.232949 | 0.261880 | 0.293502 | 0.260459 | 0.221073 | 0.215070 | 0.222945 | 0.252640 | 0.283987 | 0.345116 | 0.362080 | 0.374574 | 0.349527 | 0.310143 | 0.281280 | 0.253623 | 0.263258 | 0.264455 | 0.267965 | 0.259813 | 0.250680 | 0.227484 | 0.221260 | 0.214744 | 0.219056 | 0.223473 | 0.223424 | 0.218401 | 0.214813 | 0.216464 | 0.215201 | 0.214869 | 0.218504 | 0.240207 | 0.282246 | 0.374634 | 0.446829 | 0.364706 | 0.338531 | 0.391392 | 0.331193 | 0.294441 | 0.248004 | 0.225553 | 0.239198 | 0.260885 | 0.259413 | 0.290515 | 0.334875 | 0.322596 | 0.323481 | 0.296934 | 0.256645 | 0.248153 | 0.225208 | 0.214639 | 0.216153 | 0.216210 | 0.216833 | 0.220609 | 0.235088 | 0.220054 | 0.215637 | 0.230353 | 0.263227 | 0.292881 | 0.396856 | 0.533043 | 0.480036 | 0.450443 | 0.505793 | 0.427029 | 0.399398 | 0.377024 | 0.397358 | 0.528341 | 0.653214 | 0.702184 | 0.709465 | 0.495071 | 0.337362 | 0.271901 | 0.241589 | 0.228933 | 0.234969 | 0.277725 | 0.319301 | 0.400694 | 0.372416 | 0.350802 | 0.325891 | 0.301518 | 0.244610 | 0.236050 | 0.221974 | 0.224399 | 0.223901 | 0.222041 | 0.218822 |
| right posterior | 0.271875 | 0.247105 | 0.233690 | 0.271688 | 0.325287 | 0.366384 | 0.365099 | 0.485472 | 1.020953 | 1.766033 | 1.574833 | 0.775748 | 0.668449 | 0.795919 | 0.437655 | 0.301420 | 0.253709 | 0.215533 | 0.217286 | 0.216567 | 0.221684 | 0.226585 | 0.252430 | 0.257331 | 0.223863 | 0.216575 | 0.217928 | 0.221122 | 0.219712 | 0.214947 | 0.215315 | 0.216222 | 0.218905 | 0.215315 | 0.214939 | 0.215413 | 0.214716 | 0.216524 | 0.222116 | 0.243671 | 0.300290 | 0.335205 | 0.496334 | 1.541719 | 1.758041 | 1.117491 | 0.616576 | 0.300770 | 0.239212 | 0.217734 | 0.224278 | 0.260402 | 0.240561 | 0.218006 | 0.214638 | 0.214638 | 0.215816 | 0.215948 | 0.214730 | 0.239452 | 0.327003 | 0.349792 | 0.360632 | 0.412275 | 0.450633 | 0.372702 | 0.334013 | 0.344959 | 0.413746 | 0.375939 | 0.346882 | 0.317191 | 0.278164 | 0.222061 | 0.232825 | 0.374286 | 0.838369 | 1.885127 | 2.387936 | 2.038605 | 1.723764 | 1.538708 | 1.291409 | 1.076133 | 0.897842 | 0.733171 | 0.485735 | 0.303184 | 0.217244 | 0.232274 | 0.306056 | 0.555399 | 1.160120 | 2.321069 | 4.114357 | 3.005189 | 1.718138 | 1.024999 | 0.609297 | 0.380137 | 0.246854 | 0.215056 | 0.260685 | 0.370408 | 0.390325 | 0.322799 | 0.261150 | 0.219489 | 0.217512 | 0.243210 | 0.283543 | 0.304330 | 0.255335 | 0.225571 | 0.214997 | 0.216197 | 0.219128 | 0.219475 | 0.217588 | 0.214771 | 0.214680 | 0.215888 | 0.218080 | 0.219644 | 0.217644 | 0.215493 | 0.214632 | 0.214728 | 0.217346 | 0.229213 | 0.247921 | 0.275733 | 0.289564 | 0.308562 | 0.314283 | 0.292581 | 0.308089 | 0.387421 | 0.501414 | 0.656948 | 0.871058 | 0.905728 | 0.694894 | 0.294679 | 0.214675 | 0.249371 | 0.369876 | 0.518427 | 0.728534 | 1.094942 | 1.739496 | 3.876257 | 10.588756 | 27.829117 | 51.987226 | 29.203923 | 16.087778 | 8.005333 | 3.246176 | 1.764570 | 0.862942 | 0.570272 | 0.469021 | 0.336753 | 0.280365 | 0.243799 | 0.231409 | 0.241974 | 0.234926 | 0.229761 | 0.247353 | 0.266213 | 0.269109 | 0.259026 | 0.237926 | 0.248254 | 0.285697 | 0.306637 | 0.390778 | 0.520312 | 0.531401 | 0.541843 | 0.688072 | 0.612356 | 0.403521 | 0.259594 | 0.224522 | 0.216737 | 0.215780 | 0.216196 | 0.225436 | 0.226525 | 0.223357 | 0.224280 | 0.226974 | 0.233737 | 0.250518 | 0.253567 | 0.242853 | 0.239663 | 0.228403 | 0.223472 | 0.216066 | 0.216416 | 0.220166 | 0.214815 | 0.215605 | 0.220913 | 0.222481 | 0.218468 | 0.218066 | 0.216018 | 0.214634 | 0.220198 | 0.243210 | 0.305232 | 0.335170 | 0.305703 | 0.284103 | 0.294141 | 0.315289 | 0.272023 | 0.230066 | 0.215237 | 0.215646 | 0.216302 | 0.220065 | 0.219196 | 0.234132 | 0.241682 | 0.252918 | 0.265095 | 0.331229 | 0.361628 | 0.342731 | 0.314466 | 0.346629 | 0.374267 | 0.394484 | 0.329919 | 0.263698 | 0.232897 | 0.214989 | 0.227433 | 0.240555 | 0.249969 | 0.234719 | 0.217553 | 0.215241 | 0.216538 | 0.229012 | 0.274953 | 0.353728 | 0.392075 | 0.396173 | 0.524050 | 0.613190 | 0.632084 | 0.423467 | 0.293979 | 0.237045 | 0.216186 | 0.222317 | 0.233443 | 0.229506 | 0.218072 | 0.215343 | 0.223635 | 0.241159 | 0.241934 | 0.237649 | 0.227822 | 0.228189 | 0.220753 | 0.217410 | 0.218043 | 0.228065 | 0.239252 | 0.250298 | 0.250829 |
| all electrodes | 0.232562 | 0.244520 | 0.244849 | 0.233170 | 0.219940 | 0.216497 | 0.221372 | 0.221571 | 0.292858 | 0.360160 | 0.383656 | 0.432378 | 0.625766 | 0.494386 | 0.290098 | 0.214886 | 0.242858 | 0.279033 | 0.309278 | 0.312243 | 0.300248 | 0.215283 | 0.217189 | 0.218417 | 0.221743 | 0.245057 | 0.289032 | 0.309645 | 0.333759 | 0.267853 | 0.217029 | 0.255898 | 0.429529 | 0.453963 | 0.307134 | 0.217586 | 0.226564 | 0.274982 | 0.329077 | 0.472202 | 0.371817 | 0.262982 | 0.214934 | 0.215269 | 0.214626 | 0.217558 | 0.237566 | 0.312001 | 0.324006 | 0.356872 | 0.320930 | 0.303190 | 0.240491 | 0.216487 | 0.215328 | 0.215258 | 0.215320 | 0.214715 | 0.217122 | 0.242301 | 0.258365 | 0.279912 | 0.276279 | 0.309075 | 0.314050 | 0.252213 | 0.265896 | 0.362879 | 0.425137 | 0.299333 | 0.268680 | 0.268286 | 0.274151 | 0.215645 | 0.244818 | 0.339424 | 0.372221 | 0.450700 | 0.429537 | 0.387226 | 0.312269 | 0.271571 | 0.259347 | 0.236243 | 0.223223 | 0.227374 | 0.229483 | 0.217097 | 0.214859 | 0.234119 | 0.264629 | 0.300098 | 0.363119 | 0.539275 | 0.536972 | 0.579981 | 0.584473 | 0.418664 | 0.334065 | 0.304322 | 0.260135 | 0.272824 | 0.258438 | 0.234129 | 0.248321 | 0.301626 | 0.288165 | 0.293200 | 0.247943 | 0.246234 | 0.249463 | 0.233806 | 0.226770 | 0.222486 | 0.218898 | 0.234929 | 0.243342 | 0.249796 | 0.272986 | 0.269228 | 0.313884 | 0.356303 | 0.329590 | 0.333843 | 0.313754 | 0.277853 | 0.273708 | 0.250878 | 0.255413 | 0.244906 | 0.240333 | 0.259070 | 0.310410 | 0.294792 | 0.258694 | 0.226127 | 0.222548 | 0.229782 | 0.227846 | 0.220416 | 0.221100 | 0.216522 | 0.214854 | 0.214794 | 0.230424 | 0.263183 | 0.306947 | 0.379179 | 0.359865 | 0.420982 | 0.519527 | 0.524384 | 0.617647 | 0.613471 | 0.602928 | 0.502728 | 0.303685 | 0.258836 | 0.244472 | 0.237987 | 0.239617 | 0.226133 | 0.222186 | 0.230412 | 0.219063 | 0.215959 | 0.215123 | 0.219063 | 0.249701 | 0.287035 | 0.322412 | 0.300194 | 0.319790 | 0.462415 | 0.794059 | 0.785847 | 1.176550 | 1.074130 | 0.943631 | 0.666971 | 0.430537 | 0.403999 | 0.353286 | 0.274258 | 0.288080 | 0.345632 | 0.461327 | 0.431632 | 0.288639 | 0.258081 | 0.248283 | 0.220950 | 0.214744 | 0.228131 | 0.232340 | 0.230689 | 0.235437 | 0.239557 | 0.236255 | 0.242648 | 0.283547 | 0.401635 | 0.437560 | 0.529755 | 0.468922 | 0.464883 | 0.508917 | 0.472703 | 0.310709 | 0.289327 | 0.249481 | 0.284685 | 0.309623 | 0.346185 | 0.357217 | 0.502344 | 0.496398 | 0.423152 | 0.393976 | 0.398628 | 0.311112 | 0.286475 | 0.240268 | 0.226213 | 0.218296 | 0.214626 | 0.216689 | 0.216110 | 0.229743 | 0.231771 | 0.242698 | 0.232024 | 0.218596 | 0.215338 | 0.215105 | 0.214633 | 0.214691 | 0.216457 | 0.220092 | 0.218894 | 0.216405 | 0.218017 | 0.223892 | 0.216530 | 0.219975 | 0.221104 | 0.217514 | 0.214676 | 0.216249 | 0.214635 | 0.219363 | 0.239826 | 0.244925 | 0.251050 | 0.259187 | 0.257256 | 0.271836 | 0.266492 | 0.285368 | 0.260203 | 0.231020 | 0.217038 | 0.218622 | 0.224830 | 0.252687 | 0.254086 | 0.363924 | 0.403645 | 0.453910 | 0.436457 | 0.379669 | 0.378629 | 0.362467 | 0.306929 | 0.383027 | 0.391580 | 0.452732 | 0.493465 | 0.453233 | 0.518291 |

Searchlight, spatiotemporal cluster permutation test

No significant clusters observed.

AA) emoji to real faces - neutral vs angry

  
|  | time window | peak latency | cluster *p* | peak Cohen's *d* |  | | | |  |
| **all electrodes** | 125 - 185 ms | 155 ms | 0.0348 | 1.7192 |  | | | |  |
 325 - 525 ms | 415 ms | 0.0087 | 0.9491 |  | | | | 660 - 935 ms | 815 ms | 0.0054 | 0.6838 |  | | | ||  |  | | | | | | | | |

Time-resolved classification, cluster permutation tests

|  |  | **left hemisphere** | | | | **right hemisphere** | | | |
|  | time window | peak latency | cluster *p* | peak Cohen's *d* | time window | peak latency | cluster *p* | peak Cohen's *d* |  |
| **anterior** |  | | | |  | | | |  |
| **central** |  | | | | 130 - 180 ms | 155 ms | 0.0324 | 1.4996 |  |
  | | | | 255 - 305 ms | 280 ms | 0.0428 | 1.2616 |  | | | | 345 - 470 ms | 440 ms | 0.0138 | 0.7123 ||  | **posterior** | 120 - 185 ms | 155 ms | 0.0308 | 1.5717 | 125 - 185 ms | 150 ms | 0.0324 | 1.7375 |
 255 - 905 ms | 280 ms | 0.0001 | 1.1683 | 360 - 540 ms | 410 ms | 0.0155 | 0.8493 | 920 - 1135 ms | 1020 ms | 0.0122 | 0.7371 | 580 - 930 ms | 740 ms | 0.0029 | 0.8755 |

  

Time-resolved classification, Bayesian statistics

|  | -200 | -195 | -190 | -185 | -180 | -175 | -170 | -165 | -160 | -155 | -150 | -145 | -140 | -135 | -130 | -125 | -120 | -115 | -110 | -105 | -100 | -95 | -90 | -85 | -80 | -75 | -70 | -65 | -60 | -55 | -50 | -45 | -40 | -35 | -30 | -25 | -20 | -15 | -10 | -5 | 0 | 5 | 10 | 15 | 20 | 25 | 30 | 35 | 40 | 45 | 50 | 55 | 60 | 65 | 70 | 75 | 80 | 85 | 90 | 95 | 100 | 105 | 110 | 115 | 120 | 125 | 130 | 135 | 140 | 145 | 150 | 155 | 160 | 165 | 170 | 175 | 180 | 185 | 190 | 195 | 200 | 205 | 210 | 215 | 220 | 225 | 230 | 235 | 240 | 245 | 250 | 255 | 260 | 265 | 270 | 275 | 280 | 285 | 290 | 295 | 300 | 305 | 310 | 315 | 320 | 325 | 330 | 335 | 340 | 345 | 350 | 355 | 360 | 365 | 370 | 375 | 380 | 385 | 390 | 395 | 400 | 405 | 410 | 415 | 420 | 425 | 430 | 435 | 440 | 445 | 450 | 455 | 460 | 465 | 470 | 475 | 480 | 485 | 490 | 495 | 500 | 505 | 510 | 515 | 520 | 525 | 530 | 535 | 540 | 545 | 550 | 555 | 560 | 565 | 570 | 575 | 580 | 585 | 590 | 595 | 600 | 605 | 610 | 615 | 620 | 625 | 630 | 635 | 640 | 645 | 650 | 655 | 660 | 665 | 670 | 675 | 680 | 685 | 690 | 695 | 700 | 705 | 710 | 715 | 720 | 725 | 730 | 735 | 740 | 745 | 750 | 755 | 760 | 765 | 770 | 775 | 780 | 785 | 790 | 795 | 800 | 805 | 810 | 815 | 820 | 825 | 830 | 835 | 840 | 845 | 850 | 855 | 860 | 865 | 870 | 875 | 880 | 885 | 890 | 895 | 900 | 905 | 910 | 915 | 920 | 925 | 930 | 935 | 940 | 945 | 950 | 955 | 960 | 965 | 970 | 975 | 980 | 985 | 990 | 995 | 1000 | 1005 | 1010 | 1015 | 1020 | 1025 | 1030 | 1035 | 1040 | 1045 | 1050 | 1055 | 1060 | 1065 | 1070 | 1075 | 1080 | 1085 | 1090 | 1095 | 1100 | 1105 | 1110 | 1115 | 1120 | 1125 | 1130 | 1135 | 1140 | 1145 | 1150 | 1155 | 1160 | 1165 | 1170 | 1175 | 1180 | 1185 | 1190 | 1195 |
| --- | --- | --- | --- | --- | --- | --- | --- | --- | --- | --- | --- | --- | --- | --- | --- | --- | --- | --- | --- | --- | --- | --- | --- | --- | --- | --- | --- | --- | --- | --- | --- | --- | --- | --- | --- | --- | --- | --- | --- | --- | --- | --- | --- | --- | --- | --- | --- | --- | --- | --- | --- | --- | --- | --- | --- | --- | --- | --- | --- | --- | --- | --- | --- | --- | --- | --- | --- | --- | --- | --- | --- | --- | --- | --- | --- | --- | --- | --- | --- | --- | --- | --- | --- | --- | --- | --- | --- | --- | --- | --- | --- | --- | --- | --- | --- | --- | --- | --- | --- | --- | --- | --- | --- | --- | --- | --- | --- | --- | --- | --- | --- | --- | --- | --- | --- | --- | --- | --- | --- | --- | --- | --- | --- | --- | --- | --- | --- | --- | --- | --- | --- | --- | --- | --- | --- | --- | --- | --- | --- | --- | --- | --- | --- | --- | --- | --- | --- | --- | --- | --- | --- | --- | --- | --- | --- | --- | --- | --- | --- | --- | --- | --- | --- | --- | --- | --- | --- | --- | --- | --- | --- | --- | --- | --- | --- | --- | --- | --- | --- | --- | --- | --- | --- | --- | --- | --- | --- | --- | --- | --- | --- | --- | --- | --- | --- | --- | --- | --- | --- | --- | --- | --- | --- | --- | --- | --- | --- | --- | --- | --- | --- | --- | --- | --- | --- | --- | --- | --- | --- | --- | --- | --- | --- | --- | --- | --- | --- | --- | --- | --- | --- | --- | --- | --- | --- | --- | --- | --- | --- | --- | --- | --- | --- | --- | --- | --- | --- | --- | --- | --- | --- | --- | --- | --- | --- | --- | --- | --- | --- | --- | --- | --- | --- | --- | --- | --- | --- | --- | --- | --- | --- | --- | --- | --- | --- | --- | --- | --- | --- | --- |
| left anterior | 0.563919 | 0.678476 | 1.256179 | 1.661157 | 3.176944 | 10.930781 | 9.531753 | 2.053830 | 0.628940 | 0.246632 | 0.232044 | 0.409216 | 0.954568 | 0.743727 | 0.484030 | 0.443381 | 0.446881 | 0.252727 | 0.226458 | 0.249787 | 0.252617 | 0.260103 | 0.225659 | 0.215276 | 0.259100 | 0.275893 | 0.220999 | 0.277922 | 0.394923 | 0.656495 | 2.436701 | 6.098833 | 8.254808 | 8.634359 | 10.913540 | 3.360238 | 0.793193 | 0.334805 | 0.280388 | 0.238506 | 0.226726 | 0.344090 | 0.306865 | 0.249862 | 0.250845 | 0.239754 | 0.235742 | 0.223968 | 0.216374 | 0.221637 | 0.232148 | 0.219699 | 0.214673 | 0.215455 | 0.221320 | 0.227511 | 0.248735 | 0.215092 | 0.267633 | 0.522358 | 1.150115 | 1.541886 | 1.326664 | 1.031272 | 0.386167 | 0.222212 | 0.263822 | 0.613669 | 1.784636 | 3.313847 | 4.007221 | 2.138157e+00 | 1.807048e+00 | 1.093995e+00 | 0.466143 | 0.321017 | 0.269638 | 0.243334 | 0.258385 | 0.252288 | 0.274499 | 0.308186 | 0.451932 | 0.381703 | 0.281073 | 0.233213 | 0.228251 | 0.235282 | 0.327264 | 0.364841 | 0.561889 | 1.342752 | 2.886112 | 5.242476 | 5.751667 | 2.709671 | 1.170975 | 0.612102 | 0.419872 | 0.395723 | 0.312210 | 0.316827 | 0.408815 | 0.400672 | 0.461776 | 0.474868 | 0.406333 | 0.355603 | 0.330908 | 0.336442 | 0.478626 | 0.610357 | 0.758649 | 0.770687 | 0.924136 | 0.860502 | 0.901643 | 0.879184 | 0.634975 | 0.511988 | 0.446785 | 0.397676 | 0.494106 | 0.455814 | 0.389915 | 0.424200 | 0.340754 | 0.263599 | 0.224964 | 0.214797 | 0.214681 | 0.217807 | 0.225329 | 0.219493 | 0.214774 | 0.223162 | 0.229694 | 0.226975 | 0.248913 | 0.289367 | 0.295163 | 0.290914 | 0.235450 | 0.225732 | 0.221932 | 0.219886 | 0.215019 | 0.217214 | 0.223795 | 0.230775 | 0.253256 | 0.281294 | 0.290102 | 0.297365 | 0.304508 | 0.359275 | 0.307696 | 0.283730 | 0.271510 | 0.281097 | 0.287840 | 0.280033 | 0.244836 | 0.257852 | 0.258330 | 0.239166 | 0.238895 | 0.250909 | 0.241033 | 0.255430 | 0.271493 | 0.272298 | 0.266750 | 0.279473 | 0.266984 | 0.269661 | 0.234977 | 0.218736 | 0.214651 | 0.214860 | 0.215478 | 0.221055 | 0.234776 | 0.233525 | 0.222768 | 0.224525 | 0.231026 | 0.220500 | 0.220043 | 0.227846 | 0.237227 | 0.257368 | 0.251846 | 0.253189 | 0.312702 | 0.327625 | 0.285207 | 0.286162 | 0.288723 | 0.311464 | 0.360322 | 0.360723 | 0.346437 | 0.320256 | 0.299757 | 0.303314 | 0.264813 | 0.220773 | 0.215047 | 0.214626 | 0.217861 | 0.214626 | 0.217270 | 0.216249 | 0.214635 | 0.214626 | 0.215664 | 0.215273 | 0.215382 | 0.215336 | 0.214633 | 0.214745 | 0.216447 | 0.215412 | 0.215457 | 0.219269 | 0.218548 | 0.215256 | 0.218991 | 0.234023 | 0.277905 | 0.304504 | 0.282111 | 0.245865 | 0.229489 | 0.240376 | 0.245741 | 0.233809 | 0.219261 | 0.214638 | 0.215576 | 0.216273 | 0.216608 | 0.217539 | 0.230125 | 0.241095 | 0.218825 | 0.216459 | 0.219653 | 0.251473 | 0.359801 | 0.482251 | 0.580081 | 1.013206 | 1.182900 | 2.019318 | 2.084412 | 1.750698 | 1.430233 | 1.126566 | 0.840628 | 0.722870 | 0.454257 | 0.357908 | 0.304209 | 0.275921 | 0.323152 | 0.322391 | 0.297563 | 0.325937 | 0.437567 | 0.516907 | 0.470991 | 0.310037 | 0.311034 | 0.365430 | 0.370570 | 0.285259 | 0.248729 | 0.238024 |
| right anterior | 1.174841 | 0.627826 | 0.460317 | 0.346552 | 0.237913 | 0.214916 | 0.215850 | 0.234732 | 0.291323 | 0.371987 | 0.341718 | 0.268749 | 0.254348 | 0.272983 | 0.220491 | 0.214927 | 0.215323 | 0.216012 | 0.218508 | 0.243989 | 0.279253 | 0.335414 | 0.234323 | 0.216434 | 0.216523 | 0.214641 | 0.228064 | 0.229309 | 0.217159 | 0.214670 | 0.215911 | 0.215740 | 0.214956 | 0.216434 | 0.215477 | 0.226508 | 0.217808 | 0.215960 | 0.216342 | 0.214970 | 0.254903 | 0.275966 | 0.268465 | 0.223942 | 0.222492 | 0.266439 | 0.326109 | 0.259017 | 0.229100 | 0.234720 | 0.256227 | 0.234762 | 0.217952 | 0.298927 | 0.334978 | 0.330814 | 0.507812 | 0.442719 | 0.362206 | 0.258700 | 0.217319 | 0.226500 | 0.239848 | 0.283313 | 0.543481 | 1.883659 | 13.777718 | 69.909282 | 409.223334 | 654.123219 | 553.293466 | 2.365708e+02 | 8.565098e+01 | 1.995307e+01 | 5.078330 | 2.071696 | 1.176570 | 0.612146 | 0.564110 | 0.464183 | 0.471777 | 0.476237 | 0.337388 | 0.271140 | 0.254573 | 0.233518 | 0.234947 | 0.246351 | 0.295739 | 0.476675 | 0.989413 | 1.554757 | 1.438262 | 1.214480 | 1.941685 | 1.447026 | 0.768672 | 0.306453 | 0.239050 | 0.228872 | 0.241509 | 0.236354 | 0.270970 | 0.274125 | 0.340438 | 0.462344 | 0.429283 | 0.407526 | 0.394399 | 0.367227 | 0.349325 | 0.333985 | 0.251079 | 0.250806 | 0.255843 | 0.254230 | 0.236521 | 0.240820 | 0.260619 | 0.273763 | 0.265632 | 0.219034 | 0.214637 | 0.216608 | 0.214890 | 0.217619 | 0.222257 | 0.219198 | 0.230239 | 0.237242 | 0.229760 | 0.221225 | 0.214632 | 0.215187 | 0.215136 | 0.215462 | 0.214658 | 0.230332 | 0.247340 | 0.237851 | 0.233806 | 0.223330 | 0.216846 | 0.217009 | 0.268492 | 0.367999 | 0.329577 | 0.430370 | 0.517638 | 0.611857 | 0.792474 | 0.540654 | 0.486921 | 0.693711 | 0.728994 | 0.778182 | 0.662294 | 0.457311 | 0.450738 | 0.325971 | 0.247175 | 0.227349 | 0.219488 | 0.216561 | 0.229222 | 0.238995 | 0.252479 | 0.275777 | 0.275496 | 0.262065 | 0.248492 | 0.219871 | 0.216056 | 0.227511 | 0.263535 | 0.321562 | 0.388350 | 0.381120 | 0.295108 | 0.240768 | 0.221249 | 0.214711 | 0.215698 | 0.219016 | 0.221046 | 0.215037 | 0.215629 | 0.217343 | 0.223063 | 0.224075 | 0.229450 | 0.238225 | 0.220145 | 0.222444 | 0.222169 | 0.215223 | 0.214681 | 0.216208 | 0.223874 | 0.261256 | 0.302712 | 0.345754 | 0.465868 | 0.538023 | 0.442332 | 0.307932 | 0.247207 | 0.240392 | 0.224621 | 0.224693 | 0.230376 | 0.226955 | 0.249770 | 0.270272 | 0.267522 | 0.275526 | 0.256762 | 0.240772 | 0.244444 | 0.243525 | 0.257807 | 0.278383 | 0.333150 | 0.431146 | 0.665004 | 0.877236 | 0.908692 | 0.675354 | 0.473200 | 0.416736 | 0.354213 | 0.303941 | 0.254686 | 0.238713 | 0.258184 | 0.281853 | 0.249787 | 0.248663 | 0.254167 | 0.253018 | 0.269890 | 0.281940 | 0.276541 | 0.395855 | 0.502691 | 0.486820 | 0.501838 | 0.421521 | 0.412214 | 0.406549 | 0.316116 | 0.256484 | 0.233691 | 0.237876 | 0.262161 | 0.276306 | 0.353249 | 0.414557 | 0.614085 | 0.765584 | 0.798447 | 1.008596 | 1.178373 | 1.186371 | 1.095304 | 0.748151 | 0.525700 | 0.382298 | 0.283925 | 0.246246 | 0.228172 | 0.218730 | 0.219855 | 0.225766 | 0.240877 | 0.241625 | 0.240463 | 0.235875 | 0.233201 | 0.218253 |
| left central | 0.228160 | 0.218956 | 0.215501 | 0.214760 | 0.215517 | 0.214626 | 0.223764 | 0.215051 | 0.217502 | 0.236801 | 0.263302 | 0.267347 | 0.320031 | 0.401590 | 0.699371 | 0.380181 | 0.271981 | 0.238480 | 0.238058 | 0.251488 | 0.217781 | 0.246296 | 0.219833 | 0.225511 | 0.214649 | 0.254543 | 0.317618 | 0.313869 | 0.238670 | 0.245417 | 0.270302 | 0.346473 | 0.354376 | 0.427608 | 0.603300 | 0.651025 | 0.593039 | 0.666225 | 0.349964 | 0.218062 | 0.249725 | 0.297967 | 0.233654 | 0.241357 | 0.370189 | 0.495161 | 0.549091 | 0.679783 | 1.171990 | 1.790370 | 1.314579 | 0.615993 | 0.377799 | 0.259940 | 0.231989 | 0.215420 | 0.221243 | 0.220770 | 0.214701 | 0.217188 | 0.244232 | 0.277453 | 0.332813 | 0.313055 | 0.229450 | 0.226414 | 0.522838 | 3.486020 | 42.024150 | 384.999544 | 1670.268692 | 2.387794e+03 | 8.689469e+02 | 1.937886e+02 | 18.133312 | 1.068258 | 0.293206 | 0.214636 | 0.246697 | 0.278386 | 0.326659 | 0.318657 | 0.253305 | 0.232550 | 0.217774 | 0.215011 | 0.217681 | 0.245611 | 0.457668 | 0.897296 | 2.341301 | 7.525079 | 24.130174 | 52.721263 | 110.490849 | 133.149656 | 147.960059 | 110.034347 | 72.547197 | 49.966906 | 13.694246 | 5.038360 | 1.689782 | 1.390299 | 1.271472 | 0.839359 | 0.475912 | 0.485880 | 0.470939 | 0.471748 | 0.336594 | 0.293528 | 0.291350 | 0.312219 | 0.358465 | 0.415951 | 0.505331 | 0.612982 | 0.733192 | 1.155715 | 2.423395 | 3.412600 | 5.245516 | 6.299453 | 6.978585 | 4.669660 | 2.807462 | 1.403884 | 0.852612 | 0.606353 | 0.473821 | 0.428129 | 0.530223 | 0.525887 | 0.526345 | 0.592289 | 0.507454 | 0.508958 | 0.489785 | 0.436075 | 0.426879 | 0.421308 | 0.369778 | 0.330363 | 0.312415 | 0.266073 | 0.238326 | 0.214635 | 0.223071 | 0.219561 | 0.215546 | 0.217538 | 0.214802 | 0.214741 | 0.226010 | 0.243860 | 0.280076 | 0.339966 | 0.470374 | 0.542582 | 0.817362 | 1.029030 | 0.984773 | 0.655731 | 0.657315 | 0.584885 | 0.534423 | 0.332623 | 0.276948 | 0.293995 | 0.299677 | 0.246147 | 0.243093 | 0.271406 | 0.370973 | 0.450870 | 0.516754 | 0.454932 | 0.471011 | 0.394967 | 0.297446 | 0.248346 | 0.225589 | 0.214774 | 0.217636 | 0.218218 | 0.215292 | 0.215918 | 0.221904 | 0.218425 | 0.217414 | 0.219928 | 0.232819 | 0.218072 | 0.214715 | 0.216984 | 0.218164 | 0.230545 | 0.228018 | 0.252885 | 0.227827 | 0.229483 | 0.224295 | 0.219630 | 0.215964 | 0.215051 | 0.216612 | 0.215622 | 0.216115 | 0.238890 | 0.252759 | 0.255911 | 0.253394 | 0.268024 | 0.241031 | 0.261750 | 0.240226 | 0.229622 | 0.219677 | 0.215783 | 0.214688 | 0.218674 | 0.224099 | 0.232714 | 0.254753 | 0.266685 | 0.266466 | 0.275712 | 0.262772 | 0.298420 | 0.314028 | 0.267903 | 0.242725 | 0.234541 | 0.225067 | 0.220698 | 0.215381 | 0.214701 | 0.215047 | 0.214701 | 0.215336 | 0.218403 | 0.217666 | 0.216735 | 0.216589 | 0.215436 | 0.216161 | 0.220313 | 0.219782 | 0.215334 | 0.215865 | 0.215207 | 0.214626 | 0.216561 | 0.217155 | 0.216817 | 0.214662 | 0.215689 | 0.223666 | 0.229340 | 0.229131 | 0.225035 | 0.220433 | 0.218845 | 0.216472 | 0.214702 | 0.214647 | 0.215096 | 0.214735 | 0.214919 | 0.232424 | 0.267423 | 0.285857 | 0.306770 | 0.273372 | 0.257203 | 0.239319 | 0.221134 | 0.215296 | 0.214710 |
| right central | 0.237703 | 0.232798 | 0.227350 | 0.218825 | 0.220334 | 0.215110 | 0.214693 | 0.214847 | 0.232056 | 0.250750 | 0.258169 | 0.346797 | 0.452326 | 0.510280 | 0.404648 | 0.263459 | 0.225715 | 0.219292 | 0.217148 | 0.229636 | 0.237079 | 0.221087 | 0.214914 | 0.219222 | 0.214918 | 0.242028 | 0.297485 | 0.312293 | 0.265425 | 0.233027 | 0.232394 | 0.215093 | 0.216012 | 0.214762 | 0.218458 | 0.226068 | 0.235745 | 0.241546 | 0.294100 | 0.477648 | 0.809886 | 1.104950 | 1.296755 | 1.035824 | 1.167435 | 1.473482 | 0.603296 | 0.328683 | 0.315217 | 0.353561 | 0.761798 | 0.959512 | 0.967797 | 0.878261 | 2.128693 | 2.709870 | 2.282130 | 1.196747 | 1.478970 | 1.572378 | 2.953175 | 1.332257 | 0.420446 | 0.224135 | 0.271451 | 1.233418 | 9.421119 | 156.022416 | 1394.235450 | 11458.715580 | 67865.208090 | 8.758095e+04 | 2.752962e+04 | 9.269723e+03 | 1277.475643 | 90.470647 | 3.564644 | 0.400943 | 0.219507 | 0.224652 | 0.306288 | 0.480222 | 0.824581 | 0.828749 | 0.770594 | 0.892900 | 1.077766 | 0.573285 | 0.285149 | 0.216530 | 0.460122 | 3.849634 | 133.998736 | 3217.903340 | 6448.835976 | 8632.173500 | 7340.440373 | 3624.308718 | 850.687590 | 166.113315 | 19.519835 | 3.537736 | 1.194722 | 0.596937 | 0.462060 | 0.495834 | 0.626363 | 0.745495 | 1.051000 | 1.771462 | 3.247272 | 7.482018 | 9.216017 | 7.898574 | 6.903734 | 9.118684 | 13.535070 | 10.355858 | 4.763050 | 3.935541 | 2.826853 | 4.005764 | 5.474079 | 5.410331 | 5.924872 | 7.371363 | 7.277261 | 12.679149 | 19.355268 | 18.181224 | 17.620345 | 13.552235 | 7.387273 | 4.174006 | 1.686986 | 0.586961 | 0.338780 | 0.277830 | 0.258600 | 0.240470 | 0.248768 | 0.237763 | 0.263439 | 0.253978 | 0.252361 | 0.270367 | 0.309412 | 0.357745 | 0.397426 | 0.443297 | 0.470536 | 0.416258 | 0.355598 | 0.299248 | 0.244674 | 0.218758 | 0.214637 | 0.215026 | 0.216459 | 0.254308 | 0.350141 | 0.404222 | 0.583906 | 0.629040 | 0.667260 | 0.660063 | 0.346270 | 0.229249 | 0.220923 | 0.222471 | 0.215140 | 0.219030 | 0.230552 | 0.255793 | 0.300294 | 0.398299 | 0.452036 | 0.379713 | 0.295575 | 0.247545 | 0.227871 | 0.224013 | 0.215119 | 0.221279 | 0.230191 | 0.234928 | 0.262920 | 0.330149 | 0.438348 | 0.388556 | 0.460625 | 0.493931 | 0.477694 | 0.343863 | 0.284002 | 0.267427 | 0.249270 | 0.241133 | 0.236801 | 0.224795 | 0.224499 | 0.216671 | 0.214666 | 0.214867 | 0.215680 | 0.227050 | 0.254511 | 0.286429 | 0.312201 | 0.347164 | 0.397807 | 0.399678 | 0.295276 | 0.265296 | 0.234194 | 0.239677 | 0.228987 | 0.214672 | 0.223140 | 0.225689 | 0.247136 | 0.276011 | 0.346540 | 0.352697 | 0.325221 | 0.282363 | 0.259255 | 0.221994 | 0.218985 | 0.248637 | 0.282325 | 0.366215 | 0.398327 | 0.391308 | 0.300591 | 0.245196 | 0.223784 | 0.216860 | 0.214805 | 0.215120 | 0.218382 | 0.218747 | 0.217951 | 0.216994 | 0.219912 | 0.233229 | 0.247093 | 0.280019 | 0.336434 | 0.379267 | 0.338421 | 0.253990 | 0.219742 | 0.214778 | 0.238692 | 0.267745 | 0.257010 | 0.232567 | 0.215548 | 0.214972 | 0.217059 | 0.228552 | 0.227194 | 0.223416 | 0.225091 | 0.223873 | 0.227990 | 0.238841 | 0.266060 | 0.299485 | 0.315661 | 0.294643 | 0.264937 | 0.281363 | 0.296702 | 0.311265 | 0.430040 | 0.459458 | 0.588972 | 1.070376 |
| left posterior | 0.820987 | 0.618599 | 0.509285 | 0.395663 | 0.389727 | 0.287393 | 0.240122 | 0.214626 | 0.224216 | 0.228131 | 0.307816 | 0.451582 | 0.481571 | 0.493195 | 0.562620 | 0.401981 | 0.324403 | 0.261110 | 0.239949 | 0.272852 | 0.320881 | 0.410310 | 0.359050 | 0.355978 | 0.296791 | 0.250339 | 0.217993 | 0.216265 | 0.226244 | 0.235460 | 0.250182 | 0.294545 | 0.378851 | 0.537338 | 0.698202 | 0.630682 | 0.378049 | 0.238109 | 0.215692 | 0.260372 | 0.370997 | 0.503987 | 0.560364 | 0.441870 | 0.295490 | 0.242034 | 0.214938 | 0.215069 | 0.214626 | 0.219552 | 0.221849 | 0.215247 | 0.216150 | 0.268749 | 0.319637 | 0.444423 | 0.770720 | 1.921635 | 3.220881 | 4.219565 | 1.265817 | 0.433485 | 0.214632 | 0.420861 | 2.294415 | 14.199253 | 262.019221 | 4903.442016 | 44475.567431 | 99155.852361 | 136417.257752 | 1.806044e+05 | 2.676329e+05 | 1.351939e+05 | 32129.064015 | 8352.341766 | 327.828359 | 12.739203 | 0.823550 | 0.286593 | 0.214951 | 0.249034 | 0.340814 | 0.323689 | 0.302738 | 0.278488 | 0.257325 | 0.252538 | 0.215645 | 0.277226 | 0.745217 | 9.423213 | 106.424281 | 514.287322 | 1316.490653 | 3442.896740 | 2691.004959 | 1185.155009 | 369.536441 | 121.657308 | 43.016409 | 27.379443 | 14.620505 | 13.852982 | 35.098618 | 57.037067 | 50.560917 | 39.933729 | 20.247484 | 11.440832 | 6.060875 | 4.007237 | 4.691110 | 5.390918 | 6.322631 | 8.279642 | 13.086168 | 20.811892 | 30.930813 | 38.696070 | 83.626361 | 193.591522 | 319.899680 | 342.609136 | 320.255126 | 188.492702 | 128.927130 | 72.870207 | 41.237216 | 23.762578 | 16.931491 | 12.855627 | 13.207558 | 18.505387 | 24.467930 | 24.784212 | 19.453047 | 17.377444 | 21.940373 | 29.007887 | 31.208847 | 35.041346 | 44.447546 | 67.565445 | 108.780208 | 91.113382 | 53.578828 | 36.570963 | 28.675230 | 16.363671 | 10.582297 | 6.925546 | 5.798548 | 6.083329 | 7.574004 | 7.470938 | 9.348327 | 11.820446 | 13.125727 | 12.168675 | 14.357594 | 20.478344 | 27.002279 | 33.966224 | 40.813846 | 46.749029 | 55.365456 | 49.872078 | 42.983795 | 37.803117 | 40.238492 | 40.699645 | 58.762368 | 98.830086 | 132.041652 | 154.837077 | 191.107940 | 205.501291 | 201.923118 | 129.019390 | 73.354267 | 57.601221 | 48.453364 | 37.609557 | 23.277526 | 19.043156 | 19.649443 | 24.152114 | 20.695596 | 19.198281 | 10.599391 | 8.186714 | 9.678711 | 9.837213 | 9.310241 | 9.231489 | 9.230612 | 15.267647 | 22.619645 | 34.170547 | 44.634568 | 49.707782 | 65.107455 | 55.504529 | 54.853990 | 45.924343 | 23.060493 | 20.055613 | 17.106618 | 12.443424 | 19.426331 | 21.724556 | 19.335003 | 14.766456 | 8.783057 | 8.478780 | 7.785979 | 3.962022 | 2.154194 | 1.664125 | 1.447224 | 1.452925 | 1.223884 | 1.284442 | 1.622161 | 1.875266 | 2.286368 | 3.015078 | 3.209395 | 3.610802 | 4.741946 | 5.401969 | 8.145540 | 10.227602 | 10.369751 | 12.173714 | 12.044609 | 9.643649 | 12.407488 | 9.240244 | 8.785641 | 8.875029 | 13.284788 | 18.539787 | 25.054568 | 31.106877 | 65.749281 | 110.847284 | 80.096597 | 23.875273 | 10.222896 | 7.052150 | 5.511685 | 3.166074 | 1.901726 | 1.468886 | 1.444857 | 2.089166 | 2.312540 | 2.261897 | 2.879212 | 3.349014 | 4.327466 | 5.104492 | 3.460395 | 2.175802 | 1.681431 | 1.411080 | 1.067404 | 0.986320 | 0.858575 | 0.932614 | 1.137006 | 1.353101 | 1.328981 | 1.741360 | 1.639003 | 1.718530 | 1.676966 | 1.672489 |
| right posterior | 0.298246 | 0.281752 | 0.302964 | 0.259487 | 0.280011 | 0.302319 | 0.296378 | 0.283282 | 0.242287 | 0.215005 | 0.222615 | 0.246557 | 0.360900 | 0.493095 | 0.478143 | 0.315587 | 0.237713 | 0.214981 | 0.233765 | 0.283035 | 0.288966 | 0.227756 | 0.217597 | 0.214977 | 0.215448 | 0.216632 | 0.243156 | 0.227441 | 0.215265 | 0.214705 | 0.215878 | 0.215148 | 0.218867 | 0.234801 | 0.218342 | 0.226815 | 0.280849 | 0.381827 | 0.504030 | 0.963456 | 4.626423 | 5.212256 | 3.475178 | 1.558545 | 1.493059 | 1.350395 | 0.856793 | 0.390315 | 0.260831 | 0.264253 | 0.331302 | 0.360728 | 0.437631 | 0.518376 | 0.702775 | 0.925364 | 0.682784 | 0.714570 | 0.706705 | 0.547321 | 0.381305 | 0.283262 | 0.216770 | 0.266558 | 0.825605 | 4.175117 | 26.362816 | 394.845296 | 9937.967668 | 143452.687233 | 904555.891111 | 1.879613e+06 | 2.772026e+06 | 3.756174e+06 | 600179.564986 | 37427.064321 | 1183.731390 | 17.801539 | 0.893874 | 0.252175 | 0.226086 | 0.317934 | 0.349436 | 0.443508 | 0.641078 | 0.959863 | 1.060070 | 0.871184 | 0.598660 | 0.411306 | 0.218493 | 0.363204 | 1.075935 | 2.178237 | 5.311458 | 10.407222 | 23.896547 | 16.835676 | 6.184557 | 3.370083 | 1.633632 | 0.704369 | 0.347059 | 0.247044 | 0.234925 | 0.250394 | 0.266339 | 0.308089 | 0.403212 | 0.635842 | 0.881860 | 0.961882 | 1.352062 | 2.187131 | 4.505978 | 10.709995 | 19.145885 | 29.118550 | 45.623795 | 61.483465 | 72.696992 | 72.935358 | 82.817352 | 81.328087 | 83.514802 | 80.662893 | 58.004464 | 47.916110 | 43.730736 | 38.493740 | 37.277521 | 34.075409 | 32.421147 | 24.184141 | 16.532591 | 17.892903 | 14.356797 | 12.760591 | 10.615805 | 7.897301 | 6.901843 | 7.529235 | 7.967929 | 9.631017 | 8.669422 | 5.849844 | 3.402058 | 2.227552 | 1.572309 | 1.010857 | 0.701826 | 0.638993 | 0.600406 | 0.633888 | 0.759850 | 1.079564 | 1.585891 | 2.146693 | 2.395985 | 2.675552 | 3.614178 | 3.266283 | 3.098951 | 2.813023 | 2.039243 | 2.084096 | 2.636673 | 3.200600 | 7.783129 | 8.992402 | 7.103408 | 12.349915 | 10.295601 | 7.026213 | 4.510734 | 2.918023 | 2.351118 | 4.293176 | 4.368123 | 4.787496 | 5.710833 | 7.387771 | 10.877254 | 15.190627 | 13.220505 | 27.260401 | 42.172834 | 78.683654 | 109.985043 | 83.267684 | 66.124416 | 48.270431 | 29.712887 | 26.567767 | 20.885612 | 15.247369 | 14.178613 | 16.620081 | 25.295034 | 37.316619 | 43.747841 | 41.017070 | 44.814000 | 53.937102 | 49.042376 | 25.820655 | 14.021934 | 13.723363 | 16.816989 | 18.237194 | 16.238136 | 11.817930 | 12.880801 | 11.350892 | 8.298407 | 5.153189 | 5.292489 | 4.020181 | 4.277954 | 3.465869 | 2.996459 | 2.503760 | 2.925442 | 2.941074 | 2.512599 | 2.097828 | 1.816195 | 1.252136 | 1.067340 | 0.715371 | 0.539674 | 0.448704 | 0.424855 | 0.423660 | 0.506963 | 0.523046 | 0.667246 | 0.822218 | 1.397487 | 1.950886 | 3.567856 | 5.920103 | 8.986686 | 9.606706 | 11.796464 | 15.460724 | 18.378095 | 20.438388 | 17.223368 | 15.074004 | 8.835586 | 5.668270 | 2.121854 | 1.143306 | 1.116668 | 0.950905 | 0.961856 | 1.571058 | 2.533158 | 3.641803 | 3.949030 | 2.124876 | 2.324755 | 2.010996 | 1.386791 | 0.756930 | 0.470344 | 0.317923 | 0.266763 | 0.230077 | 0.227877 | 0.249851 | 0.341185 | 0.489767 | 0.562432 | 0.763220 | 1.066886 | 1.237640 | 1.070671 | 0.767300 |
| all electrodes | 0.221261 | 0.219932 | 0.216886 | 0.215674 | 0.215345 | 0.214626 | 0.214772 | 0.214691 | 0.231051 | 0.240538 | 0.315471 | 0.448902 | 0.608622 | 0.834326 | 1.534912 | 1.657482 | 2.157483 | 0.720288 | 0.421160 | 0.236584 | 0.219658 | 0.218844 | 0.215350 | 0.253724 | 0.330664 | 0.379146 | 0.296808 | 0.337392 | 0.386489 | 0.470464 | 0.404849 | 0.338589 | 0.258344 | 0.232723 | 0.227377 | 0.216987 | 0.226457 | 0.259571 | 0.278076 | 0.331617 | 0.346764 | 0.503890 | 0.483359 | 0.467061 | 0.544183 | 0.591075 | 0.510531 | 0.394581 | 0.423099 | 0.322109 | 0.297837 | 0.340712 | 0.326150 | 0.334642 | 0.459047 | 0.568990 | 0.505877 | 0.509547 | 0.667240 | 0.724945 | 0.474613 | 0.238140 | 0.216751 | 0.274401 | 0.541379 | 1.971800 | 10.733193 | 93.915771 | 1277.534287 | 16692.101228 | 199474.110330 | 7.598576e+05 | 1.034526e+06 | 2.414788e+05 | 36793.858741 | 950.033050 | 42.355694 | 2.959839 | 0.491136 | 0.222013 | 0.237185 | 0.370179 | 0.757884 | 1.381226 | 1.661319 | 3.530377 | 5.748690 | 3.721060 | 0.892556 | 0.275366 | 0.232959 | 0.439990 | 1.509228 | 5.132797 | 10.036645 | 14.543347 | 14.403280 | 9.714235 | 5.054442 | 3.157113 | 2.009623 | 1.572005 | 1.002073 | 0.875599 | 1.151241 | 1.908406 | 4.022760 | 6.948854 | 7.539273 | 10.810875 | 14.371891 | 26.915647 | 23.719509 | 14.907710 | 15.556711 | 14.136406 | 11.162538 | 10.949611 | 9.500693 | 14.677460 | 30.967139 | 96.402352 | 154.634607 | 245.640755 | 187.105538 | 91.944279 | 54.704374 | 49.349601 | 23.044609 | 12.720474 | 9.919427 | 11.205214 | 13.873484 | 28.428155 | 27.309538 | 18.970572 | 15.333436 | 15.322258 | 12.176847 | 10.624388 | 7.580546 | 6.193229 | 5.250510 | 4.549704 | 2.684018 | 1.596465 | 0.806729 | 0.500683 | 0.458999 | 0.457069 | 0.543739 | 0.711657 | 0.901362 | 1.060259 | 1.493266 | 1.840755 | 2.189940 | 2.827864 | 2.955410 | 2.442256 | 3.226597 | 3.733950 | 3.261283 | 2.920863 | 2.448045 | 2.031477 | 2.400136 | 2.007503 | 1.439955 | 1.074211 | 0.799501 | 0.901409 | 1.303067 | 1.497605 | 1.822597 | 2.285402 | 3.610234 | 5.437294 | 5.422644 | 4.824751 | 4.265422 | 3.336724 | 3.569493 | 6.186708 | 9.665478 | 14.339759 | 20.090314 | 21.494568 | 28.604102 | 47.367161 | 25.359941 | 31.301606 | 39.622293 | 23.460761 | 19.081693 | 10.195722 | 10.347995 | 14.711972 | 7.429786 | 5.059677 | 6.760961 | 8.001153 | 12.580724 | 14.431420 | 12.892996 | 8.801562 | 6.965663 | 5.526533 | 5.847156 | 6.339942 | 4.310098 | 3.907307 | 3.917355 | 3.139708 | 2.860582 | 2.850473 | 2.782361 | 3.655002 | 3.900888 | 2.517642 | 2.640835 | 2.986310 | 2.462555 | 2.345687 | 2.171574 | 1.754742 | 1.820606 | 1.815432 | 1.183816 | 0.762401 | 0.570903 | 0.533030 | 0.696008 | 0.921764 | 1.038460 | 1.189911 | 1.966420 | 1.576026 | 1.414940 | 0.961334 | 0.929218 | 0.889072 | 0.682999 | 0.585269 | 0.765108 | 1.018745 | 1.221773 | 1.293933 | 1.348975 | 1.924675 | 2.425620 | 2.232188 | 1.407577 | 1.064253 | 0.716094 | 0.578485 | 0.606444 | 0.576266 | 0.566567 | 0.625383 | 1.119190 | 1.892096 | 3.481093 | 2.268793 | 1.726448 | 1.015399 | 0.766525 | 0.471077 | 0.352083 | 0.292073 | 0.331141 | 0.368757 | 0.461273 | 0.615222 | 0.715630 | 0.856176 | 0.805550 | 0.710147 | 0.616448 | 0.469958 |

Searchlight, spatiotemporal cluster permutation test

|  | start time | stop time | peak time | peak channel | cluster p | peak Cohen's d | direction |
| --- | --- | --- | --- | --- | --- | --- | --- |
| #1 | 120 | 210 | 150 | O2 | 0.029 | 1.878939 | positive |
| #2 | 225 | 1195 | 285 | P10 | 0.0001 | 1.37338 | positive |

AB) emoji to real faces - neutral vs sad

  
|  | time window | peak latency | cluster *p* | peak Cohen's *d* |  | | | |
| **all electrodes** | 125 - 195 ms | 150 ms | 0.0251 | 1.4665 |  | | | |
|  | | | | | | | | |

Time-resolved classification, cluster permutation tests

|  | **left hemisphere** | | | | **right hemisphere** | | | |
|  | time window | peak latency | cluster *p* | peak Cohen's *d* | time window | peak latency | cluster *p* | peak Cohen's *d* |
| **anterior** |  | | | |  | | | |
| **central** |  | | | |  | | | |
| **posterior** | 125 - 180 ms | 150 ms | 0.0361 | 1.2042 | 130 - 190 ms | 155 ms | 0.0461 | 1.2253 |
 855 - 1040 ms | 950 ms | 0.0098 | 0.8465 |  | | | |

  

Time-resolved classification, Bayesian statistics

|  | -200 | -195 | -190 | -185 | -180 | -175 | -170 | -165 | -160 | -155 | -150 | -145 | -140 | -135 | -130 | -125 | -120 | -115 | -110 | -105 | -100 | -95 | -90 | -85 | -80 | -75 | -70 | -65 | -60 | -55 | -50 | -45 | -40 | -35 | -30 | -25 | -20 | -15 | -10 | -5 | 0 | 5 | 10 | 15 | 20 | 25 | 30 | 35 | 40 | 45 | 50 | 55 | 60 | 65 | 70 | 75 | 80 | 85 | 90 | 95 | 100 | 105 | 110 | 115 | 120 | 125 | 130 | 135 | 140 | 145 | 150 | 155 | 160 | 165 | 170 | 175 | 180 | 185 | 190 | 195 | 200 | 205 | 210 | 215 | 220 | 225 | 230 | 235 | 240 | 245 | 250 | 255 | 260 | 265 | 270 | 275 | 280 | 285 | 290 | 295 | 300 | 305 | 310 | 315 | 320 | 325 | 330 | 335 | 340 | 345 | 350 | 355 | 360 | 365 | 370 | 375 | 380 | 385 | 390 | 395 | 400 | 405 | 410 | 415 | 420 | 425 | 430 | 435 | 440 | 445 | 450 | 455 | 460 | 465 | 470 | 475 | 480 | 485 | 490 | 495 | 500 | 505 | 510 | 515 | 520 | 525 | 530 | 535 | 540 | 545 | 550 | 555 | 560 | 565 | 570 | 575 | 580 | 585 | 590 | 595 | 600 | 605 | 610 | 615 | 620 | 625 | 630 | 635 | 640 | 645 | 650 | 655 | 660 | 665 | 670 | 675 | 680 | 685 | 690 | 695 | 700 | 705 | 710 | 715 | 720 | 725 | 730 | 735 | 740 | 745 | 750 | 755 | 760 | 765 | 770 | 775 | 780 | 785 | 790 | 795 | 800 | 805 | 810 | 815 | 820 | 825 | 830 | 835 | 840 | 845 | 850 | 855 | 860 | 865 | 870 | 875 | 880 | 885 | 890 | 895 | 900 | 905 | 910 | 915 | 920 | 925 | 930 | 935 | 940 | 945 | 950 | 955 | 960 | 965 | 970 | 975 | 980 | 985 | 990 | 995 | 1000 | 1005 | 1010 | 1015 | 1020 | 1025 | 1030 | 1035 | 1040 | 1045 | 1050 | 1055 | 1060 | 1065 | 1070 | 1075 | 1080 | 1085 | 1090 | 1095 | 1100 | 1105 | 1110 | 1115 | 1120 | 1125 | 1130 | 1135 | 1140 | 1145 | 1150 | 1155 | 1160 | 1165 | 1170 | 1175 | 1180 | 1185 | 1190 | 1195 |
| --- | --- | --- | --- | --- | --- | --- | --- | --- | --- | --- | --- | --- | --- | --- | --- | --- | --- | --- | --- | --- | --- | --- | --- | --- | --- | --- | --- | --- | --- | --- | --- | --- | --- | --- | --- | --- | --- | --- | --- | --- | --- | --- | --- | --- | --- | --- | --- | --- | --- | --- | --- | --- | --- | --- | --- | --- | --- | --- | --- | --- | --- | --- | --- | --- | --- | --- | --- | --- | --- | --- | --- | --- | --- | --- | --- | --- | --- | --- | --- | --- | --- | --- | --- | --- | --- | --- | --- | --- | --- | --- | --- | --- | --- | --- | --- | --- | --- | --- | --- | --- | --- | --- | --- | --- | --- | --- | --- | --- | --- | --- | --- | --- | --- | --- | --- | --- | --- | --- | --- | --- | --- | --- | --- | --- | --- | --- | --- | --- | --- | --- | --- | --- | --- | --- | --- | --- | --- | --- | --- | --- | --- | --- | --- | --- | --- | --- | --- | --- | --- | --- | --- | --- | --- | --- | --- | --- | --- | --- | --- | --- | --- | --- | --- | --- | --- | --- | --- | --- | --- | --- | --- | --- | --- | --- | --- | --- | --- | --- | --- | --- | --- | --- | --- | --- | --- | --- | --- | --- | --- | --- | --- | --- | --- | --- | --- | --- | --- | --- | --- | --- | --- | --- | --- | --- | --- | --- | --- | --- | --- | --- | --- | --- | --- | --- | --- | --- | --- | --- | --- | --- | --- | --- | --- | --- | --- | --- | --- | --- | --- | --- | --- | --- | --- | --- | --- | --- | --- | --- | --- | --- | --- | --- | --- | --- | --- | --- | --- | --- | --- | --- | --- | --- | --- | --- | --- | --- | --- | --- | --- | --- | --- | --- | --- | --- | --- | --- | --- | --- | --- | --- | --- | --- | --- | --- | --- | --- | --- | --- | --- | --- |
| left anterior | 0.220721 | 0.218530 | 0.215199 | 0.215560 | 0.215640 | 0.233301 | 0.275253 | 0.276881 | 0.308381 | 0.242141 | 0.215687 | 0.224152 | 0.365223 | 0.451035 | 0.404822 | 0.520074 | 0.453103 | 0.437153 | 0.394172 | 0.272815 | 0.255489 | 0.230729 | 0.217016 | 0.244043 | 0.333552 | 0.566156 | 0.662838 | 0.765506 | 1.102905 | 1.467910 | 0.750871 | 0.292978 | 0.225610 | 0.228722 | 0.254898 | 0.239505 | 0.231782 | 0.234943 | 0.264522 | 0.269762 | 0.244519 | 0.215669 | 0.221222 | 0.267692 | 0.259785 | 0.254312 | 0.216869 | 0.215851 | 0.220115 | 0.226744 | 0.248450 | 0.332452 | 0.330280 | 0.272992 | 0.262086 | 0.259327 | 0.270312 | 0.307296 | 0.229676 | 0.238264 | 0.238130 | 0.260710 | 0.273256 | 0.239444 | 0.214661 | 0.241653 | 0.474257 | 1.073010 | 4.664502 | 22.894739 | 44.211147 | 22.402565 | 7.940190 | 2.571032 | 1.258595 | 0.634081 | 0.476739 | 0.355133 | 0.334772 | 0.337599 | 0.283507 | 0.245089 | 0.246137 | 0.262485 | 0.282422 | 0.292554 | 0.264773 | 0.226882 | 0.217638 | 0.214632 | 0.228267 | 0.240641 | 0.241129 | 0.225649 | 0.216918 | 0.223081 | 0.215248 | 0.216575 | 0.216643 | 0.226053 | 0.226284 | 0.227890 | 0.243832 | 0.282716 | 0.261680 | 0.256579 | 0.226703 | 0.218853 | 0.215572 | 0.215370 | 0.215228 | 0.214702 | 0.223764 | 0.224581 | 0.218537 | 0.214694 | 0.214633 | 0.216411 | 0.218689 | 0.233282 | 0.224665 | 0.215530 | 0.216285 | 0.214811 | 0.214687 | 0.215410 | 0.218635 | 0.219715 | 0.221333 | 0.226476 | 0.220399 | 0.214714 | 0.218246 | 0.215957 | 0.215146 | 0.223805 | 0.226106 | 0.234800 | 0.260694 | 0.270468 | 0.233022 | 0.215029 | 0.227377 | 0.227914 | 0.230142 | 0.235836 | 0.243196 | 0.227624 | 0.216019 | 0.214857 | 0.214737 | 0.215547 | 0.217745 | 0.215234 | 0.215304 | 0.214792 | 0.218294 | 0.237134 | 0.263909 | 0.315476 | 0.284274 | 0.294770 | 0.289723 | 0.251488 | 0.237382 | 0.235488 | 0.224894 | 0.227763 | 0.221170 | 0.216128 | 0.215847 | 0.241864 | 0.279342 | 0.272491 | 0.258840 | 0.237304 | 0.223065 | 0.216587 | 0.215808 | 0.216606 | 0.219329 | 0.229577 | 0.243040 | 0.241658 | 0.270537 | 0.280220 | 0.291875 | 0.247980 | 0.224285 | 0.214848 | 0.215279 | 0.214626 | 0.214971 | 0.215168 | 0.214936 | 0.214853 | 0.215514 | 0.216033 | 0.223258 | 0.239312 | 0.236083 | 0.219708 | 0.214845 | 0.219654 | 0.225775 | 0.238297 | 0.259377 | 0.267599 | 0.287355 | 0.366452 | 0.492607 | 0.696499 | 0.607708 | 0.586239 | 0.446963 | 0.266514 | 0.214663 | 0.255368 | 0.278428 | 0.321793 | 0.444863 | 0.448869 | 0.383326 | 0.335112 | 0.302998 | 0.264665 | 0.228875 | 0.218593 | 0.221321 | 0.226394 | 0.249887 | 0.278033 | 0.322720 | 0.371973 | 0.325885 | 0.260943 | 0.239947 | 0.222807 | 0.216257 | 0.226728 | 0.235897 | 0.228704 | 0.216315 | 0.215559 | 0.215169 | 0.214850 | 0.215839 | 0.224738 | 0.237743 | 0.268140 | 0.289029 | 0.304909 | 0.306375 | 0.316795 | 0.307802 | 0.272581 | 0.251172 | 0.237532 | 0.233013 | 0.218426 | 0.214783 | 0.220154 | 0.222950 | 0.237802 | 0.250292 | 0.312094 | 0.324049 | 0.353614 | 0.309566 | 0.280105 | 0.241141 | 0.221902 | 0.215435 | 0.218955 | 0.216769 | 0.214750 | 0.214816 | 0.214872 | 0.218459 | 0.229666 |
| right anterior | 0.251896 | 0.224243 | 0.224682 | 0.221525 | 0.215079 | 0.217844 | 0.224759 | 0.236715 | 0.223782 | 0.240641 | 0.271175 | 0.268485 | 0.312679 | 0.412356 | 0.529273 | 0.381093 | 0.219600 | 0.221742 | 0.239807 | 0.248482 | 0.238699 | 0.220503 | 0.254436 | 0.255397 | 0.237973 | 0.243051 | 0.236913 | 0.214727 | 0.224737 | 0.227047 | 0.241109 | 0.251493 | 0.279654 | 0.498681 | 0.910454 | 1.297269 | 2.022829 | 2.470859 | 1.877481 | 0.879053 | 0.310857 | 0.217447 | 0.215051 | 0.215738 | 0.242063 | 0.324263 | 0.322507 | 0.338119 | 0.340144 | 0.264056 | 0.232485 | 0.214636 | 0.234574 | 0.215312 | 0.252507 | 0.374257 | 0.586902 | 0.587711 | 0.529307 | 0.511080 | 0.315376 | 0.221831 | 0.217877 | 0.224906 | 0.219851 | 0.243970 | 0.343451 | 0.982868 | 2.080659 | 4.301165 | 5.729360 | 2.597006 | 1.348646 | 0.939215 | 0.448402 | 0.299378 | 0.246119 | 0.223936 | 0.220500 | 0.215444 | 0.220407 | 0.217793 | 0.221567 | 0.227455 | 0.247813 | 0.261270 | 0.282423 | 0.257723 | 0.307615 | 0.355336 | 0.359547 | 0.605936 | 0.974508 | 1.072086 | 1.205436 | 1.228496 | 1.032410 | 1.016726 | 0.620794 | 0.491934 | 0.357234 | 0.336389 | 0.254293 | 0.239616 | 0.229437 | 0.224495 | 0.221962 | 0.234206 | 0.230626 | 0.245247 | 0.265833 | 0.305053 | 0.383133 | 0.481475 | 0.371393 | 0.310928 | 0.267384 | 0.231315 | 0.216502 | 0.218093 | 0.238764 | 0.229899 | 0.219957 | 0.215524 | 0.214649 | 0.214651 | 0.217459 | 0.225064 | 0.251950 | 0.241230 | 0.227041 | 0.229091 | 0.238203 | 0.226666 | 0.216900 | 0.217201 | 0.235067 | 0.262691 | 0.283871 | 0.326982 | 0.316468 | 0.350892 | 0.296789 | 0.278383 | 0.274474 | 0.288802 | 0.281215 | 0.285821 | 0.272149 | 0.259930 | 0.273116 | 0.273043 | 0.242055 | 0.223709 | 0.227053 | 0.249389 | 0.277160 | 0.268878 | 0.255304 | 0.244044 | 0.262681 | 0.284636 | 0.283904 | 0.299566 | 0.318312 | 0.367228 | 0.381906 | 0.390187 | 0.390792 | 0.352459 | 0.323423 | 0.305060 | 0.254153 | 0.245777 | 0.222873 | 0.214967 | 0.216081 | 0.214906 | 0.215049 | 0.215204 | 0.214984 | 0.218761 | 0.238717 | 0.251990 | 0.254767 | 0.246778 | 0.257917 | 0.308801 | 0.407743 | 0.450081 | 0.430488 | 0.375264 | 0.408200 | 0.460700 | 0.418059 | 0.337169 | 0.280451 | 0.265139 | 0.276110 | 0.292439 | 0.267436 | 0.238541 | 0.233768 | 0.244945 | 0.245846 | 0.251305 | 0.228940 | 0.257052 | 0.386860 | 0.540579 | 0.770464 | 1.046307 | 0.945045 | 0.817063 | 0.548481 | 0.355801 | 0.331031 | 0.293593 | 0.294092 | 0.357443 | 0.446743 | 0.539874 | 0.508083 | 0.422804 | 0.322517 | 0.262879 | 0.243018 | 0.234563 | 0.233636 | 0.258798 | 0.279924 | 0.315432 | 0.307414 | 0.304843 | 0.304916 | 0.282098 | 0.256910 | 0.234725 | 0.222111 | 0.231770 | 0.231430 | 0.239348 | 0.236073 | 0.234630 | 0.229227 | 0.226763 | 0.218900 | 0.216623 | 0.214735 | 0.219027 | 0.238603 | 0.306848 | 0.407198 | 0.558454 | 0.683970 | 0.773944 | 0.786886 | 0.584690 | 0.456803 | 0.356671 | 0.286240 | 0.280896 | 0.261444 | 0.254640 | 0.286882 | 0.368019 | 0.577708 | 0.811377 | 0.776802 | 0.874063 | 0.680394 | 0.520421 | 0.349161 | 0.292020 | 0.314310 | 0.370785 | 0.430620 | 0.519315 | 0.617253 | 1.069435 |
| left central | 0.220260 | 0.214649 | 0.235259 | 0.258396 | 0.274869 | 0.347102 | 0.419479 | 0.613757 | 0.477281 | 0.280863 | 0.231739 | 0.214640 | 0.239934 | 0.226379 | 0.242192 | 0.215051 | 0.224578 | 0.228853 | 0.231833 | 0.276609 | 0.225078 | 0.236445 | 0.216990 | 0.216565 | 0.216947 | 0.222919 | 0.239117 | 0.260942 | 0.231309 | 0.247292 | 0.287315 | 0.279270 | 0.242651 | 0.228641 | 0.228180 | 0.233341 | 0.217043 | 0.216311 | 0.215132 | 0.217675 | 0.214741 | 0.220073 | 0.243674 | 0.243099 | 0.292609 | 0.465360 | 0.614197 | 0.646032 | 0.783277 | 0.681403 | 0.682191 | 0.581800 | 0.543301 | 0.449871 | 0.449894 | 0.343358 | 0.298048 | 0.282968 | 0.304237 | 0.313095 | 0.345188 | 0.346054 | 0.448402 | 0.496380 | 0.319519 | 0.217114 | 0.261088 | 0.728725 | 3.535291 | 18.978513 | 65.351774 | 95.527267 | 178.109366 | 102.306317 | 11.170556 | 2.977767 | 0.971863 | 0.473248 | 0.294593 | 0.226220 | 0.217918 | 0.221219 | 0.230045 | 0.266626 | 0.364987 | 0.343929 | 0.307633 | 0.268368 | 0.256451 | 0.225054 | 0.216050 | 0.253017 | 0.304323 | 0.303944 | 0.254144 | 0.217501 | 0.218827 | 0.286249 | 0.433238 | 1.271759 | 7.164049 | 10.303693 | 2.892328 | 1.549257 | 1.010961 | 0.537129 | 0.314824 | 0.224131 | 0.215574 | 0.224995 | 0.225010 | 0.243518 | 0.305535 | 0.452557 | 0.729237 | 1.138345 | 1.102428 | 1.038218 | 0.874195 | 0.768684 | 0.609131 | 0.413455 | 0.279084 | 0.248303 | 0.250976 | 0.261386 | 0.237627 | 0.219171 | 0.214788 | 0.220358 | 0.244523 | 0.257321 | 0.334881 | 0.283543 | 0.248234 | 0.231006 | 0.219158 | 0.215177 | 0.215045 | 0.216971 | 0.214918 | 0.219692 | 0.220907 | 0.214669 | 0.217140 | 0.214710 | 0.233836 | 0.245726 | 0.254083 | 0.251483 | 0.330936 | 0.462415 | 0.573645 | 0.557009 | 0.667760 | 0.825060 | 0.776114 | 0.570585 | 0.380887 | 0.303626 | 0.257858 | 0.232923 | 0.223814 | 0.223099 | 0.230190 | 0.234139 | 0.235269 | 0.253207 | 0.228990 | 0.228540 | 0.237739 | 0.283857 | 0.380353 | 0.481238 | 0.425061 | 0.600443 | 0.654921 | 0.617195 | 0.371852 | 0.286402 | 0.261559 | 0.258474 | 0.265277 | 0.262661 | 0.318098 | 0.433738 | 0.617005 | 0.633041 | 0.556472 | 0.469134 | 0.472412 | 0.404641 | 0.323400 | 0.237066 | 0.217465 | 0.225837 | 0.247887 | 0.246932 | 0.235741 | 0.227178 | 0.233817 | 0.264419 | 0.247480 | 0.238103 | 0.247656 | 0.244701 | 0.248606 | 0.255224 | 0.229923 | 0.219825 | 0.214634 | 0.230954 | 0.244580 | 0.251489 | 0.241975 | 0.221285 | 0.215243 | 0.226613 | 0.274943 | 0.327416 | 0.389210 | 0.421684 | 0.372719 | 0.333565 | 0.258944 | 0.232634 | 0.223203 | 0.214756 | 0.215132 | 0.214909 | 0.218109 | 0.249767 | 0.306280 | 0.353317 | 0.451936 | 0.380836 | 0.348853 | 0.329789 | 0.315665 | 0.263913 | 0.243091 | 0.242902 | 0.257603 | 0.268884 | 0.261991 | 0.231639 | 0.222180 | 0.215528 | 0.215787 | 0.221680 | 0.229917 | 0.255986 | 0.250995 | 0.228072 | 0.215058 | 0.262212 | 0.354053 | 0.385174 | 0.381763 | 0.353493 | 0.313626 | 0.307088 | 0.267163 | 0.257443 | 0.261576 | 0.299349 | 0.346049 | 0.392794 | 0.392169 | 0.381533 | 0.353259 | 0.327781 | 0.288675 | 0.282092 | 0.272672 | 0.299401 | 0.350088 | 0.342645 | 0.383062 | 0.451408 |
| right central | 0.271091 | 0.229053 | 0.215417 | 0.215080 | 0.227865 | 0.247839 | 0.239668 | 0.222411 | 0.217525 | 0.214906 | 0.215091 | 0.219817 | 0.247808 | 0.248174 | 0.252845 | 0.393655 | 0.567197 | 0.803073 | 1.024793 | 0.580747 | 0.335648 | 0.245790 | 0.217279 | 0.225706 | 0.230130 | 0.254836 | 0.287239 | 0.226145 | 0.222986 | 0.214759 | 0.215722 | 0.215699 | 0.216276 | 0.218065 | 0.226888 | 0.237720 | 0.257044 | 0.258399 | 0.215523 | 0.221146 | 0.245776 | 0.253062 | 0.268621 | 0.290166 | 0.253628 | 0.215283 | 0.246233 | 0.286689 | 0.343687 | 0.371545 | 0.466187 | 0.635010 | 0.538061 | 0.406892 | 0.330147 | 0.317109 | 0.329909 | 0.299248 | 0.272974 | 0.277070 | 0.286050 | 0.311842 | 0.263435 | 0.222231 | 0.224873 | 0.351597 | 1.152539 | 5.535664 | 42.071602 | 101.125127 | 120.160503 | 120.442609 | 122.779168 | 76.139188 | 52.228809 | 18.576778 | 5.386290 | 1.265978 | 0.291824 | 0.227414 | 0.214686 | 0.241263 | 0.242981 | 0.230452 | 0.221901 | 0.218073 | 0.216249 | 0.219246 | 0.215540 | 0.215245 | 0.215503 | 0.214923 | 0.215639 | 0.214733 | 0.215615 | 0.223201 | 0.296017 | 0.434562 | 0.740037 | 0.781826 | 0.519059 | 0.301041 | 0.254532 | 0.217596 | 0.226587 | 0.345677 | 0.466578 | 0.443188 | 0.357885 | 0.404747 | 0.584687 | 0.877248 | 0.980871 | 1.463468 | 1.182492 | 0.852766 | 0.631400 | 0.522971 | 0.528531 | 0.554771 | 0.456967 | 0.536246 | 0.940042 | 1.609312 | 1.092933 | 0.520061 | 0.380336 | 0.471084 | 0.434213 | 0.457834 | 0.443494 | 0.523295 | 0.867313 | 0.824549 | 0.442779 | 0.452732 | 0.364571 | 0.271729 | 0.221055 | 0.220277 | 0.238233 | 0.235106 | 0.252757 | 0.248503 | 0.231244 | 0.214674 | 0.232115 | 0.285404 | 0.332289 | 0.403279 | 0.472780 | 0.543537 | 0.423000 | 0.298198 | 0.222371 | 0.214963 | 0.214820 | 0.214945 | 0.214772 | 0.215373 | 0.218200 | 0.224789 | 0.254977 | 0.313037 | 0.391296 | 0.485960 | 0.508745 | 0.507333 | 0.497440 | 0.389830 | 0.322589 | 0.267912 | 0.234541 | 0.238942 | 0.310113 | 0.323972 | 0.326326 | 0.359776 | 0.466269 | 0.421043 | 0.276533 | 0.221374 | 0.214734 | 0.215974 | 0.218055 | 0.215323 | 0.230340 | 0.250574 | 0.237911 | 0.236972 | 0.254689 | 0.281721 | 0.302425 | 0.351481 | 0.469856 | 0.558481 | 0.670942 | 0.624395 | 0.585123 | 0.537270 | 0.528147 | 0.526376 | 0.691001 | 0.589762 | 0.481554 | 0.363737 | 0.259613 | 0.235333 | 0.229247 | 0.219211 | 0.218396 | 0.219467 | 0.224976 | 0.231746 | 0.229250 | 0.224348 | 0.216866 | 0.216745 | 0.216268 | 0.217366 | 0.220566 | 0.219473 | 0.218456 | 0.228700 | 0.233463 | 0.249323 | 0.233214 | 0.217849 | 0.215620 | 0.215071 | 0.214784 | 0.216587 | 0.220979 | 0.219307 | 0.257663 | 0.229374 | 0.214626 | 0.219193 | 0.217417 | 0.217758 | 0.218756 | 0.215401 | 0.218609 | 0.243255 | 0.283440 | 0.296213 | 0.315084 | 0.264642 | 0.249083 | 0.269625 | 0.280610 | 0.285108 | 0.305499 | 0.311544 | 0.296088 | 0.267029 | 0.249353 | 0.252571 | 0.290013 | 0.284418 | 0.261954 | 0.263741 | 0.266585 | 0.267375 | 0.284026 | 0.267703 | 0.257652 | 0.263235 | 0.265410 | 0.290545 | 0.294430 | 0.254792 | 0.238899 | 0.220999 | 0.214912 | 0.217303 | 0.217615 | 0.216942 | 0.218576 | 0.217623 |
| left posterior | 0.258971 | 0.265008 | 0.247120 | 0.223651 | 0.214909 | 0.236203 | 0.301383 | 0.273109 | 0.261951 | 0.229228 | 0.216068 | 0.240880 | 0.381721 | 0.413946 | 0.352349 | 0.298876 | 0.246879 | 0.243663 | 0.214766 | 0.214924 | 0.243838 | 0.416831 | 0.503012 | 0.463131 | 0.415034 | 0.450218 | 0.327993 | 0.228908 | 0.214916 | 0.230057 | 0.295052 | 0.330412 | 0.258227 | 0.239698 | 0.221884 | 0.231987 | 0.221387 | 0.214626 | 0.215589 | 0.226312 | 0.234394 | 0.219019 | 0.214641 | 0.214966 | 0.216062 | 0.226903 | 0.240114 | 0.255893 | 0.246458 | 0.261434 | 0.249127 | 0.240739 | 0.226224 | 0.230287 | 0.222833 | 0.223155 | 0.229889 | 0.288068 | 0.539784 | 1.065679 | 1.099013 | 0.943700 | 0.523801 | 0.262928 | 0.311934 | 3.013052 | 70.299264 | 979.558097 | 8830.377902 | 10687.429796 | 3968.052502 | 1441.465481 | 970.776839 | 780.038329 | 339.055891 | 47.717583 | 4.453736 | 0.805905 | 0.298991 | 0.223022 | 0.216456 | 0.220567 | 0.218560 | 0.214764 | 0.228779 | 0.255077 | 0.251312 | 0.225529 | 0.217050 | 0.214820 | 0.215158 | 0.215702 | 0.217420 | 0.214650 | 0.232232 | 0.265189 | 0.280695 | 0.267335 | 0.260393 | 0.266918 | 0.237713 | 0.222068 | 0.215594 | 0.221428 | 0.219177 | 0.217539 | 0.215419 | 0.230894 | 0.234005 | 0.249270 | 0.255445 | 0.296914 | 0.397420 | 0.488994 | 0.538097 | 0.689139 | 0.688080 | 0.480007 | 0.377111 | 0.335996 | 0.358245 | 0.452067 | 0.557923 | 0.766122 | 1.275407 | 1.597572 | 2.070959 | 2.402348 | 1.946098 | 2.163112 | 3.655674 | 5.521506 | 5.540853 | 4.028581 | 2.371969 | 1.194992 | 0.776802 | 0.534149 | 0.407900 | 0.335730 | 0.307676 | 0.324973 | 0.389381 | 0.399783 | 0.432943 | 0.425769 | 0.443556 | 0.511666 | 0.479603 | 0.447888 | 0.424973 | 0.413158 | 0.422761 | 0.415823 | 0.434955 | 0.511169 | 0.532631 | 0.604442 | 0.676349 | 0.881146 | 1.394889 | 1.832537 | 2.368707 | 3.023285 | 2.529763 | 1.813858 | 1.319017 | 1.006463 | 0.819434 | 1.038623 | 1.112187 | 1.418163 | 1.657399 | 1.471054 | 1.600642 | 2.091099 | 2.710225 | 4.627215 | 4.284772 | 4.775321 | 5.615579 | 5.987663 | 5.924981 | 3.749686 | 2.828565 | 2.871662 | 3.116829 | 3.252758 | 1.896752 | 1.024219 | 0.761789 | 0.605141 | 0.570598 | 0.448970 | 0.404700 | 0.459365 | 0.627824 | 0.972366 | 1.665969 | 2.573708 | 7.161608 | 6.781087 | 5.739418 | 3.710642 | 2.070239 | 1.216255 | 0.773391 | 0.543721 | 0.544577 | 0.742001 | 1.070434 | 1.590720 | 2.341976 | 2.717730 | 2.342484 | 2.129945 | 1.903508 | 1.869534 | 1.536152 | 1.412726 | 1.706167 | 2.636740 | 4.175327 | 4.702957 | 4.367586 | 6.530927 | 13.240486 | 20.095529 | 38.609678 | 59.094589 | 80.347060 | 83.485362 | 74.919179 | 47.340003 | 23.401108 | 14.225096 | 7.167292 | 4.492341 | 3.301927 | 2.789943 | 2.779960 | 2.677246 | 2.223378 | 1.647411 | 1.427317 | 1.527086 | 1.667924 | 1.666516 | 1.750960 | 1.105659 | 0.699405 | 0.469725 | 0.324651 | 0.251280 | 0.217465 | 0.233452 | 0.264669 | 0.350425 | 0.391688 | 0.414010 | 0.423221 | 0.386123 | 0.321622 | 0.396248 | 0.393908 | 0.342156 | 0.325177 | 0.311801 | 0.320797 | 0.263954 | 0.221169 | 0.218650 | 0.242079 | 0.265685 | 0.258632 | 0.245674 | 0.220133 | 0.214886 | 0.218311 | 0.239413 |
| right posterior | 0.269328 | 0.248776 | 0.236471 | 0.230058 | 0.264997 | 0.332498 | 0.269978 | 0.244777 | 0.229972 | 0.218111 | 0.225817 | 0.246835 | 0.215563 | 0.236499 | 0.231106 | 0.226687 | 0.227246 | 0.224688 | 0.244093 | 0.257250 | 0.312025 | 0.366381 | 0.335668 | 0.272408 | 0.303469 | 0.270390 | 0.226614 | 0.214641 | 0.232337 | 0.283060 | 0.402035 | 0.391558 | 0.263885 | 0.223189 | 0.220197 | 0.219778 | 0.215385 | 0.245376 | 0.247699 | 0.226606 | 0.214643 | 0.216046 | 0.216236 | 0.215608 | 0.228314 | 0.251822 | 0.281972 | 0.288216 | 0.270479 | 0.324531 | 0.263609 | 0.224233 | 0.216283 | 0.214626 | 0.214968 | 0.229779 | 0.297234 | 0.367975 | 1.007922 | 4.988695 | 25.793419 | 34.014846 | 6.165628 | 1.865850 | 0.251043 | 0.438901 | 3.765186 | 38.257709 | 259.547164 | 1012.575294 | 4171.258299 | 4976.578285 | 5636.686174 | 4629.254868 | 2369.420417 | 649.926961 | 71.718657 | 7.746575 | 1.413287 | 0.439317 | 0.248021 | 0.215016 | 0.228168 | 0.217139 | 0.214633 | 0.217079 | 0.224515 | 0.231934 | 0.253662 | 0.280963 | 0.304934 | 0.323288 | 0.304540 | 0.279060 | 0.276112 | 0.259302 | 0.247910 | 0.236602 | 0.237411 | 0.247203 | 0.260468 | 0.268807 | 0.276103 | 0.259274 | 0.267716 | 0.276661 | 0.280618 | 0.283592 | 0.275625 | 0.266362 | 0.274151 | 0.264647 | 0.274798 | 0.277867 | 0.288983 | 0.307703 | 0.320205 | 0.323245 | 0.333033 | 0.341286 | 0.329223 | 0.279467 | 0.255370 | 0.252407 | 0.266650 | 0.256467 | 0.226687 | 0.217444 | 0.224592 | 0.224644 | 0.224679 | 0.218524 | 0.218809 | 0.227237 | 0.242464 | 0.258014 | 0.279484 | 0.271688 | 0.271933 | 0.277677 | 0.271374 | 0.257652 | 0.236138 | 0.227008 | 0.225802 | 0.216179 | 0.214922 | 0.214690 | 0.215106 | 0.215039 | 0.214646 | 0.214665 | 0.215655 | 0.218899 | 0.217187 | 0.216540 | 0.220501 | 0.223135 | 0.233515 | 0.246083 | 0.257384 | 0.273053 | 0.283506 | 0.285540 | 0.314414 | 0.331471 | 0.291623 | 0.267921 | 0.253454 | 0.239059 | 0.223014 | 0.215780 | 0.214727 | 0.214632 | 0.214649 | 0.214895 | 0.215448 | 0.227674 | 0.251450 | 0.275085 | 0.290956 | 0.344710 | 0.540443 | 0.674717 | 0.633990 | 0.547603 | 0.528523 | 0.592789 | 0.653628 | 0.566355 | 0.496509 | 0.459642 | 0.514890 | 0.506245 | 0.497939 | 0.440381 | 0.499541 | 0.710008 | 0.952351 | 1.036929 | 1.194767 | 1.045274 | 1.036741 | 0.783648 | 0.583177 | 0.524425 | 0.476387 | 0.438015 | 0.492467 | 0.532164 | 0.543316 | 0.517137 | 0.482048 | 0.424523 | 0.362425 | 0.288147 | 0.258576 | 0.241868 | 0.231791 | 0.225592 | 0.225046 | 0.230835 | 0.249555 | 0.270893 | 0.285884 | 0.313157 | 0.288922 | 0.267560 | 0.255186 | 0.251657 | 0.240718 | 0.227795 | 0.216970 | 0.214765 | 0.215019 | 0.218968 | 0.222205 | 0.241246 | 0.272537 | 0.313003 | 0.379145 | 0.422973 | 0.348742 | 0.268798 | 0.235755 | 0.220917 | 0.214768 | 0.218214 | 0.222112 | 0.216709 | 0.214715 | 0.214722 | 0.214717 | 0.228223 | 0.237672 | 0.241539 | 0.237659 | 0.234077 | 0.227036 | 0.218500 | 0.226335 | 0.251911 | 0.286698 | 0.384957 | 0.435510 | 0.484682 | 0.496311 | 0.411623 | 0.404067 | 0.376750 | 0.276238 | 0.240332 | 0.221071 | 0.214633 | 0.215978 | 0.236462 | 0.284681 | 0.284712 | 0.283521 | 0.302960 |
| all electrodes | 0.232967 | 0.224727 | 0.252650 | 0.267849 | 0.238866 | 0.226949 | 0.233910 | 0.262287 | 0.298974 | 0.225494 | 0.214910 | 0.223842 | 0.227602 | 0.227626 | 0.216400 | 0.217596 | 0.217080 | 0.214886 | 0.219560 | 0.220762 | 0.220034 | 0.278769 | 0.311715 | 0.256197 | 0.240881 | 0.232898 | 0.214714 | 0.257515 | 0.281125 | 0.270193 | 0.297514 | 0.345523 | 0.403376 | 0.384330 | 0.419728 | 0.536893 | 0.983320 | 1.418588 | 1.847452 | 0.820771 | 0.467195 | 0.295468 | 0.250221 | 0.228382 | 0.240040 | 0.240236 | 0.254301 | 0.228416 | 0.224762 | 0.219230 | 0.214908 | 0.246286 | 0.328977 | 0.285099 | 0.221935 | 0.217952 | 0.220727 | 0.219376 | 0.215052 | 0.221534 | 0.276766 | 0.330410 | 0.411084 | 0.238791 | 0.282174 | 1.801841 | 21.561275 | 146.809636 | 847.313217 | 14338.443616 | 62533.459040 | 111814.714254 | 50023.244587 | 14746.693564 | 8617.322114 | 2013.428142 | 423.295193 | 53.793501 | 8.104418 | 3.297037 | 0.895838 | 0.369066 | 0.302166 | 0.297364 | 0.420105 | 0.546050 | 0.586285 | 0.688095 | 0.729366 | 0.641809 | 0.751596 | 0.961554 | 1.165206 | 0.993032 | 0.961580 | 1.477215 | 1.446912 | 0.998630 | 0.760707 | 0.714846 | 1.079156 | 1.168738 | 0.995094 | 0.973017 | 1.044816 | 0.973301 | 1.147600 | 0.962791 | 0.966785 | 1.094275 | 1.439449 | 2.139480 | 2.720553 | 2.226955 | 2.615935 | 2.249647 | 1.696508 | 1.587123 | 1.460309 | 1.377226 | 1.522006 | 1.859706 | 2.955641 | 5.611460 | 6.007065 | 5.184377 | 2.955608 | 2.069514 | 1.347121 | 0.672655 | 0.467788 | 0.367285 | 0.316126 | 0.324653 | 0.399015 | 0.454803 | 0.661005 | 0.788521 | 0.875421 | 0.838857 | 1.117644 | 0.856512 | 0.811526 | 0.886399 | 0.812242 | 0.574070 | 0.438833 | 0.364393 | 0.359091 | 0.316405 | 0.246711 | 0.219499 | 0.216753 | 0.214757 | 0.216930 | 0.220665 | 0.219090 | 0.214646 | 0.218311 | 0.225479 | 0.250286 | 0.299262 | 0.336595 | 0.298046 | 0.249833 | 0.228352 | 0.219165 | 0.215968 | 0.214729 | 0.214633 | 0.220999 | 0.256057 | 0.301781 | 0.355906 | 0.308332 | 0.260913 | 0.238654 | 0.223095 | 0.215250 | 0.214784 | 0.214652 | 0.222966 | 0.244338 | 0.286626 | 0.341848 | 0.364100 | 0.429299 | 0.452614 | 0.447468 | 0.471872 | 0.424029 | 0.421212 | 0.401243 | 0.351256 | 0.336021 | 0.304953 | 0.307518 | 0.319125 | 0.263832 | 0.239564 | 0.232022 | 0.228780 | 0.252549 | 0.261074 | 0.245542 | 0.247023 | 0.277595 | 0.332785 | 0.355929 | 0.389998 | 0.408133 | 0.547699 | 0.530632 | 0.397583 | 0.307875 | 0.276900 | 0.227337 | 0.226120 | 0.222806 | 0.222634 | 0.216073 | 0.218164 | 0.218955 | 0.214750 | 0.214655 | 0.214626 | 0.215311 | 0.222679 | 0.246966 | 0.247124 | 0.232983 | 0.223311 | 0.214879 | 0.225324 | 0.243997 | 0.249193 | 0.260832 | 0.272041 | 0.285718 | 0.251003 | 0.229054 | 0.219275 | 0.220130 | 0.214928 | 0.214778 | 0.214682 | 0.215751 | 0.218251 | 0.234894 | 0.262251 | 0.340203 | 0.341187 | 0.327947 | 0.290047 | 0.271673 | 0.245293 | 0.229174 | 0.224683 | 0.232108 | 0.250117 | 0.284751 | 0.300682 | 0.316950 | 0.345508 | 0.394484 | 0.473995 | 0.564072 | 0.563509 | 0.615487 | 0.545674 | 0.507673 | 0.380848 | 0.295629 | 0.281303 | 0.316572 | 0.329717 | 0.385052 | 0.418247 | 0.551943 | 0.760667 |

Searchlight, spatiotemporal cluster permutation test

|  | start time | stop time | peak time | peak channel | cluster p | peak Cohen's d | direction |
| --- | --- | --- | --- | --- | --- | --- | --- |
| #1 | 120 | 545 | 155 | POz | 0.0077 | 1.801967 | positive |
| #2 | 550 | 1070 | 870 | O1 | 0.0249 | 0.745279 | positive |
